# A eutherian-specific metaviral gene, *RTL6*, coordinates microglial inflammatory responsiveness and state regulation

**DOI:** 10.64898/2026.07.15.738833

**Authors:** Fumitoshi Ishino, Masahito Irie, Hirosuke Shiura, Takashi Kohda, Tomoko Kaneko-Ishino

**Author notes:** Correspondence to: Fumitoshi Ishino and Tomko Kaneko-Ishino. Research Infrastructure Management Organization, Institute of Science Tokyo, Tokyo 113-8510, Japan (Fumitoshi Ishino). Nonprofit Organization Gene Information Bank, Fukuoka 810-0041, Japan (Masahito Irie).

## Abstract

*RTL6* (also known as *SIRH3*) is a eutherian-specific metavirus-derived gene highly conserved among placental mammals. We previously demonstrated that RTL6 is expressed in microglia and secreted into the brain extracellular space, where it mediates the trapping and clearance of bacterial lipopolysaccharide (LPS). However, its role in microglial TLR4 signaling remained unclear. To investigate the earliest LPS response, we performed RNA-seq analysis of wild-type and *Rtl6*-deficient primary microglia following 10 min of LPS stimulation. *Rtl6*-deficient microglia exhibited compensatory upregulation of residual transcripts derived from the untranslated region of *Rtl6*, suggesting feedback regulation of *Rtl6* expression. Loss of RTL6 attenuated the induction of immediate-early response genes, including *Fos, Jun, Nr4a1, Nr4a2, Egr2,* and *Egr3*, together with broad suppression of interferon-responsive and inflammatory transcriptional programs. Altered neuronal and oligodendrocyte/myelin interaction pathways indicated remodeling of microglial communication networks. Strikingly, genes involved in cell-cycle progression, DNA replication, chromatin assembly, DNA repair, and genome maintenance were coordinately upregulated, including multiple disease-associated microglia (DAM)-related genes. Collectively, these findings indicate that RTL6 couples extracellular LPS sensing to inflammatory transcriptional responses while regulating microglial functional state transitions. Our results identify *RTL6* as a previously unrecognized component of the regulatory network governing microglial functional states in placental mammals.

## 1. Introduction

In recent years, increasing attention has been directed toward virus-derived genes that have been co-opted during mammalian evolution (Kaneko-Ishino and Ishino, 2012, 2015; Wang and Han, 2021, 2022; Henriques *et al*., 2024; Shiura *et al*., 2026). Eutherian genomes contain numerous endogenous retroviral elements, including lineage-specific envelope (*ENV*)-derived genes such as *syncytins* (Mi *et al*., 2000; Lavialle *et al*., 2013). In addition to these, mammals also harbor a set of conserved genes derived from metavirus-like elements. These include the Sushi-ichi retrotransposon homologues (SIRH)/Retrotransposon Gag-like (RTL) gene family, which encodes proteins homologous to retroviral GAG (and in some cases POL) proteins, as well as Paraneoplastic Ma antigen (PNMA) genes, both derived from Ty3/gypsy-related retrotransposons (hereafter collectively referred to as metavirus-derived genes) (Kaneko-Ishino and Ishino, 2012, 2015; Henriques *et al*., 2024, Shiura *et al*., 2026). Notably, most of these genes are conserved across eutherians, and increasing evidence suggests that some have acquired important roles in mammalian development. Among them, Paternally expressed 10 (*PEG10*) is unique in being shared between eutherians and marsupials (Ono *et al*., 2001; Suzuki *et al*., 2007), whereas most other members are specific to eutherians, with a few found in marsupials (Charlier *et al*., 2001; Edwards *et al*., 2008; Kaneko-Ishino and Ishino, 2012, Wang and Han, 2021, Henriques *et al*., 2024, Shiura *et al*., 2026). Genetic studies using knockout (KO) mouse models have demonstrated that several of these genes, including *PEG10*, *RTL1* (*PEG11*), and *LDOC1* (*RTL7/SIRH7*), play essential roles in placental development (Ono *et al*., 2006; Sekita *et al*., 2008; Naruse *et al*., 2014; Kitazawa *et al*., 2017; Shiura *et al*., 2021, 2025). In contrast, other members, such as *RTL4* (*SIRH11/ZCCHC16*), *RTL5* (*SIRH8/RGAG4*), *RTL6* (*SIRH3/LDOC1L*), and *RTL9* (*SIRH10/RGAG1*), have been implicated in brain function, particularly in microglia, as revealed by a series of analyses of knock-in (KI) mice that produce a fusion protein with a fluorescent protein. These KI mouse studies have linked *Rtl4* to stress-related behavioral responses (Irie *et al*., 2015; Ishino *et al*., 2024), whereas *Rtl5*, *Rtl6,* and *Rtl9* have been implicated in host defense against viral, bacterial, and fungal components, respectively, possibly through the recognition and clearance of pathogen-associated molecular patterns (PAMPs) (Irie *et al*., 2022; Ishino *et al*., 2023).

Microglia are a conserved population of central nervous system (CNS)-resident macrophages that originate from yolk sac–derived progenitors and colonize the brain early in development (Ginhoux *et al.,* 2010; Ginhoux and Prinz, 2015). This ontogenetic program is broadly conserved across vertebrates, suggesting that microglia likely emerged with the establishment of the vertebrate CNS (Shimizu and Prinz, 2025). In mammals, certain lineage-specific biological innovations, such as soft skin and placentas, are associated with the co-option of virus-derived genes (Ono et al. 2006; Sekita et al., 2008; Matsui et al., 2011). Consistent with this concept, several eutherian-specific metavirus-derived genes, including *RTL4, RTL5*, *RTL6*, and *RTL9*, are selectively expressed in microglia, suggesting that they may contribute to the regulatory programs underlying microglial functions and state transitions. Among them, *RTL6* is an exceptionally well-conserved gene among eutherian mammals (Irie *et al*., 2022). Its dN/dS value (< 0.05) is substantially lower than the reported average value for typical housekeeping genes (0.093) (Zhang and Li, 2004), suggesting that *RTL6* has been maintained under strong purifying selection throughout eutherian evolution. Therefore, it is of particular interest to dissect the roles of *RTL6* in microglia.

In the present study, we investigated the role of *RTL6* in the earliest response to lipopolysaccharide (LPS) administration in microglia. Because we previously observed that although RTL6 reacts to LPS soon after its administration, *Tnfa* and *Il6* expression are largely comparable between WT and *Rtl6* KO microglia in 1 hr of LPS administration (Irie *et al*., 2022). Our RNA-seq results demonstrated that the loss of *Rtl6* is characterized by reduced TLR4-induced inflammatory responsiveness and increased expression of genes related to proliferation, as well as altered tissue remodeling in the presence of LPS. This implies that *RTL6* contributes to the coordination of microglial state transitions and inflammatory responsiveness in addition to the recognition and clearance of LPS.

## 2. Results

### Evidence for feedback regulation of *Rtl6* expression in KO microglia

To investigate the earliest transcriptional response to LPS, we compared RNA-seq profiles of WT and *Rtl6* KO microglia with or without 10 min of LPS treatment. *Rtl6* KO mice were generated by deleting the entire RTL6 open reading frame (ORF) while leaving the 5′ and 3′ untranslated regions (UTRs) intact (Irie et al., 2022). RNA-seq read coverage confirmed the complete absence of ORF-derived transcripts in KO microglia. In contrast, residual transcripts corresponding to the 5′ and 3′ UTRs were expressed at approximately threefold higher levels than in WT microglia regardless of LPS treatment (Figure 1A). These findings suggest feedback regulation of *Rtl6* transcription, implying that *Rtl6* expression is necessary for maintaining a normal state of microglia.

**Figure 1.**
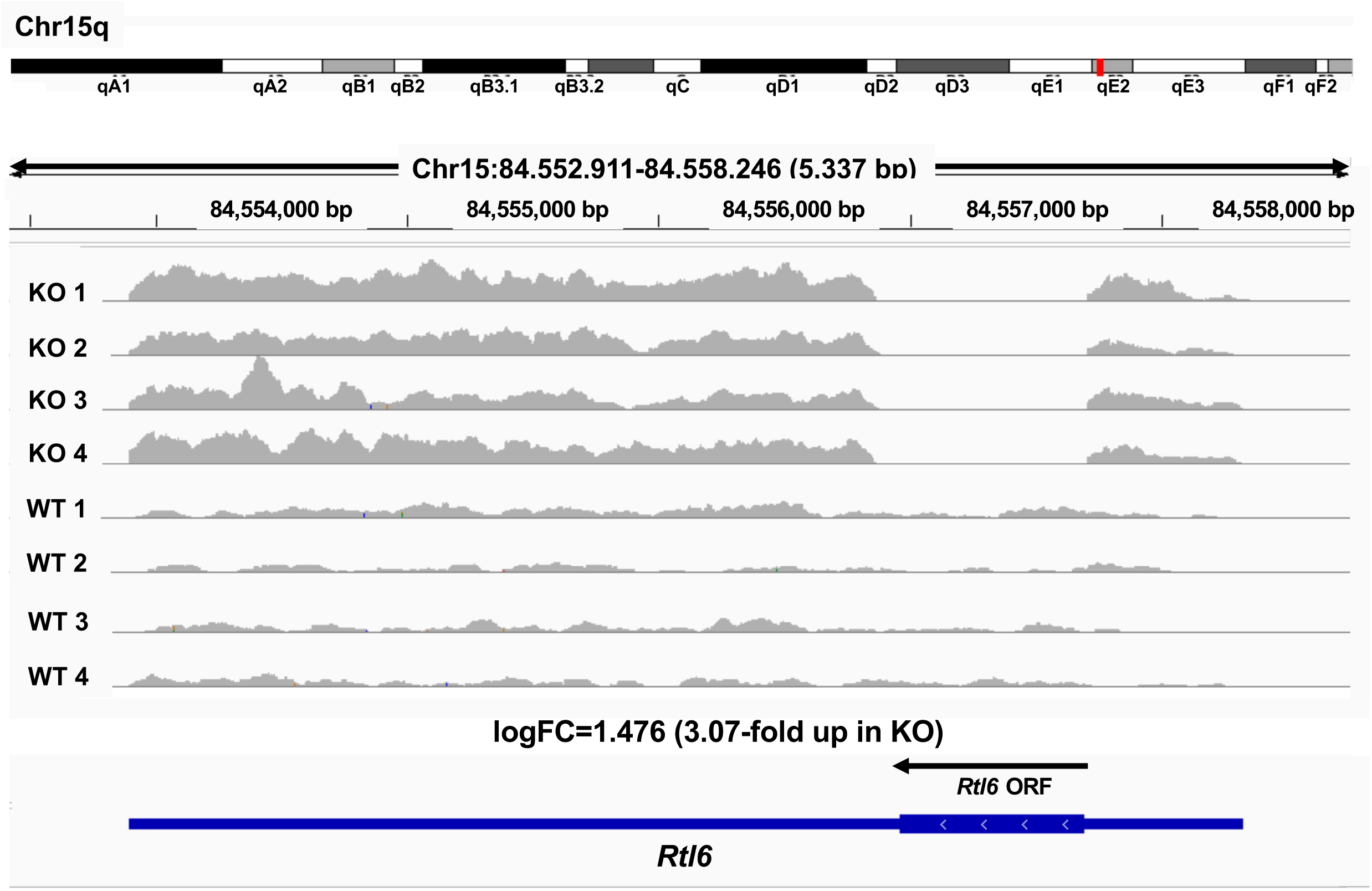
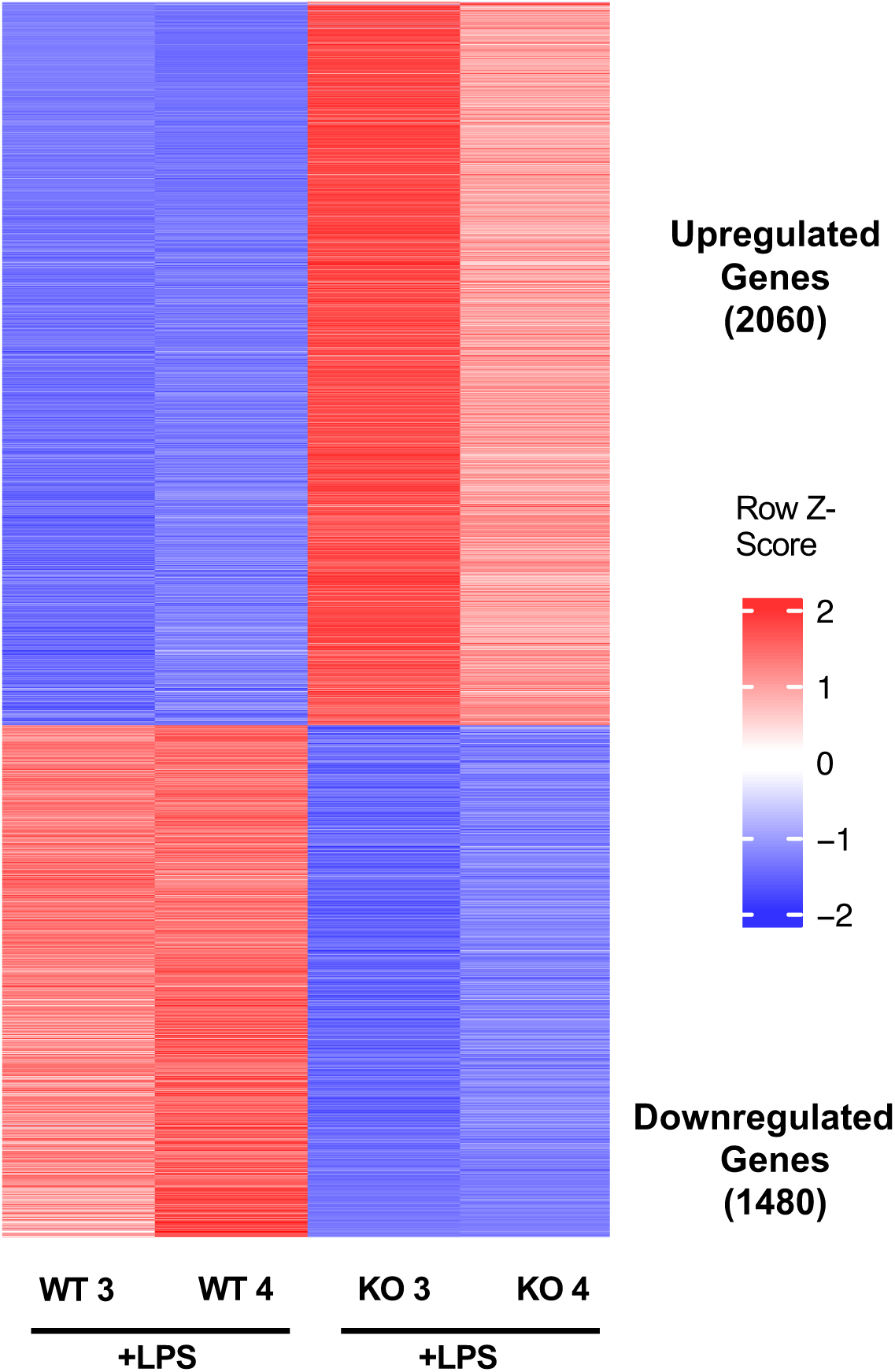
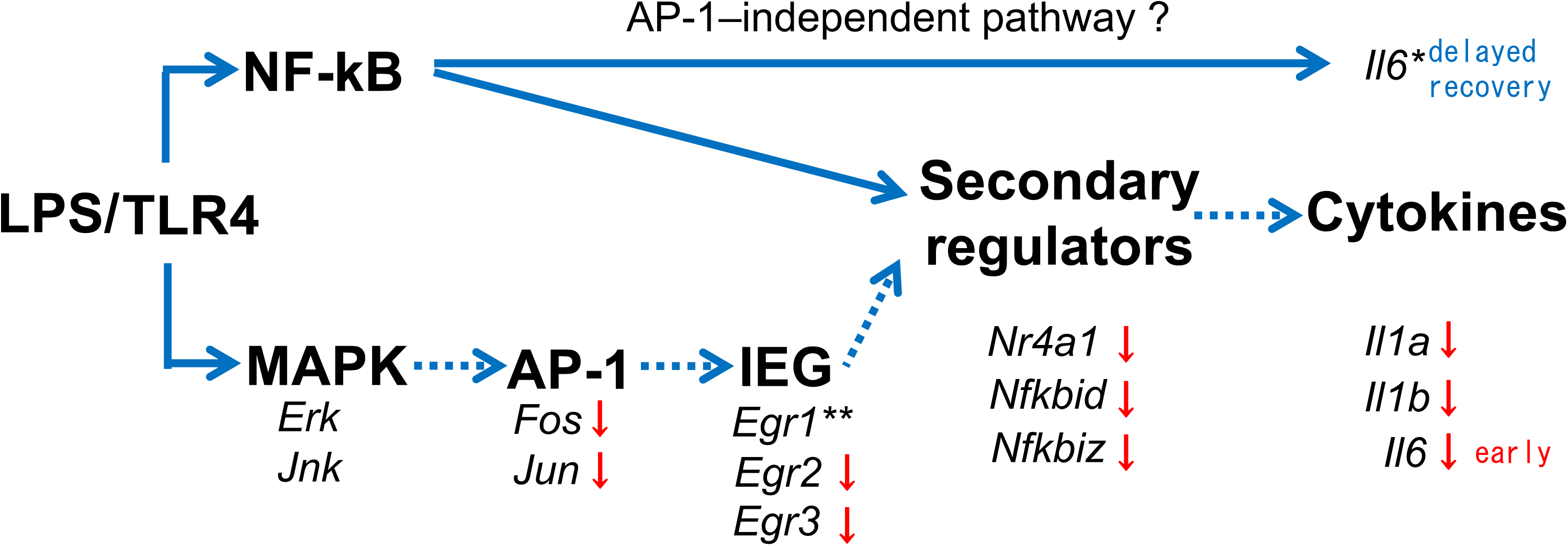
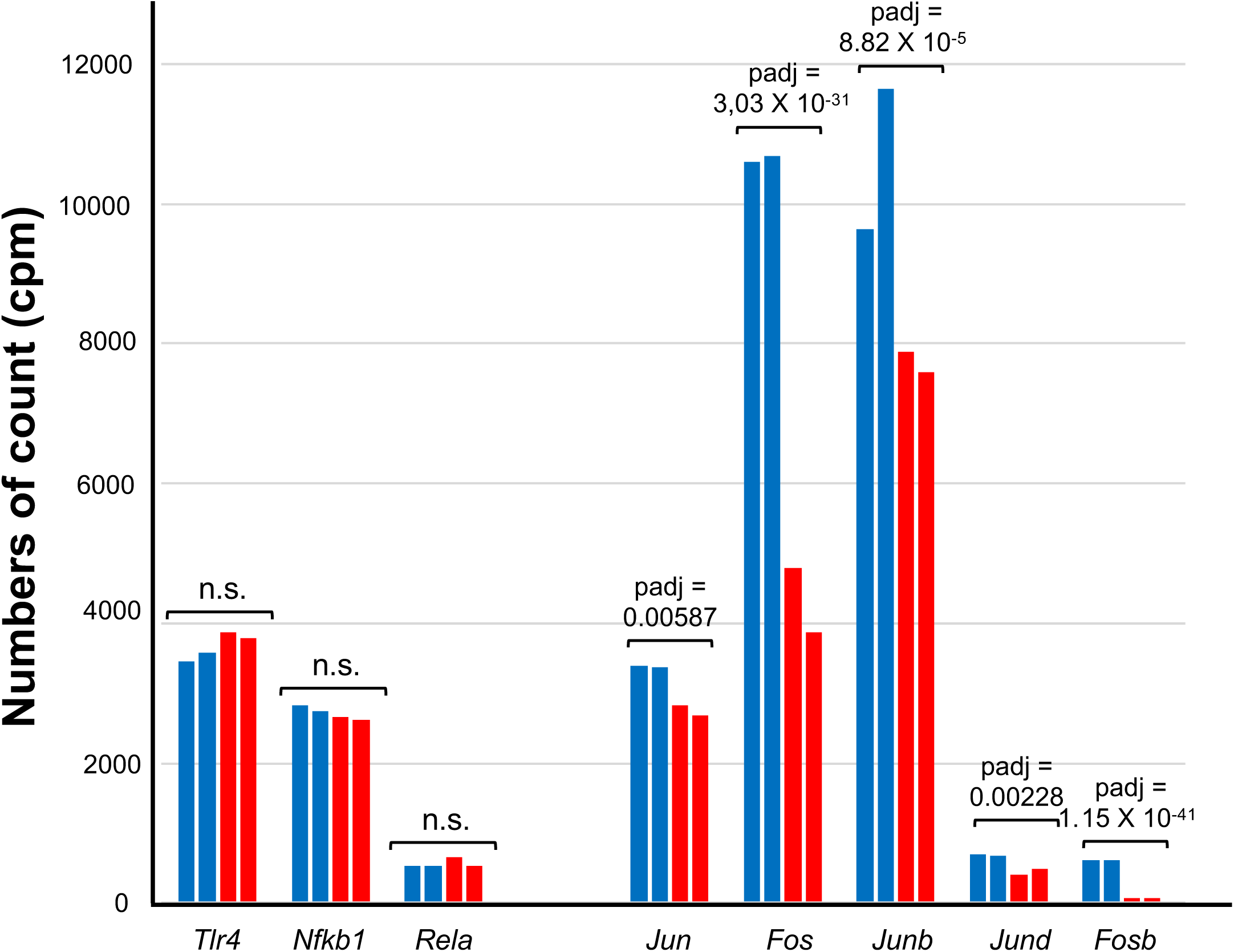
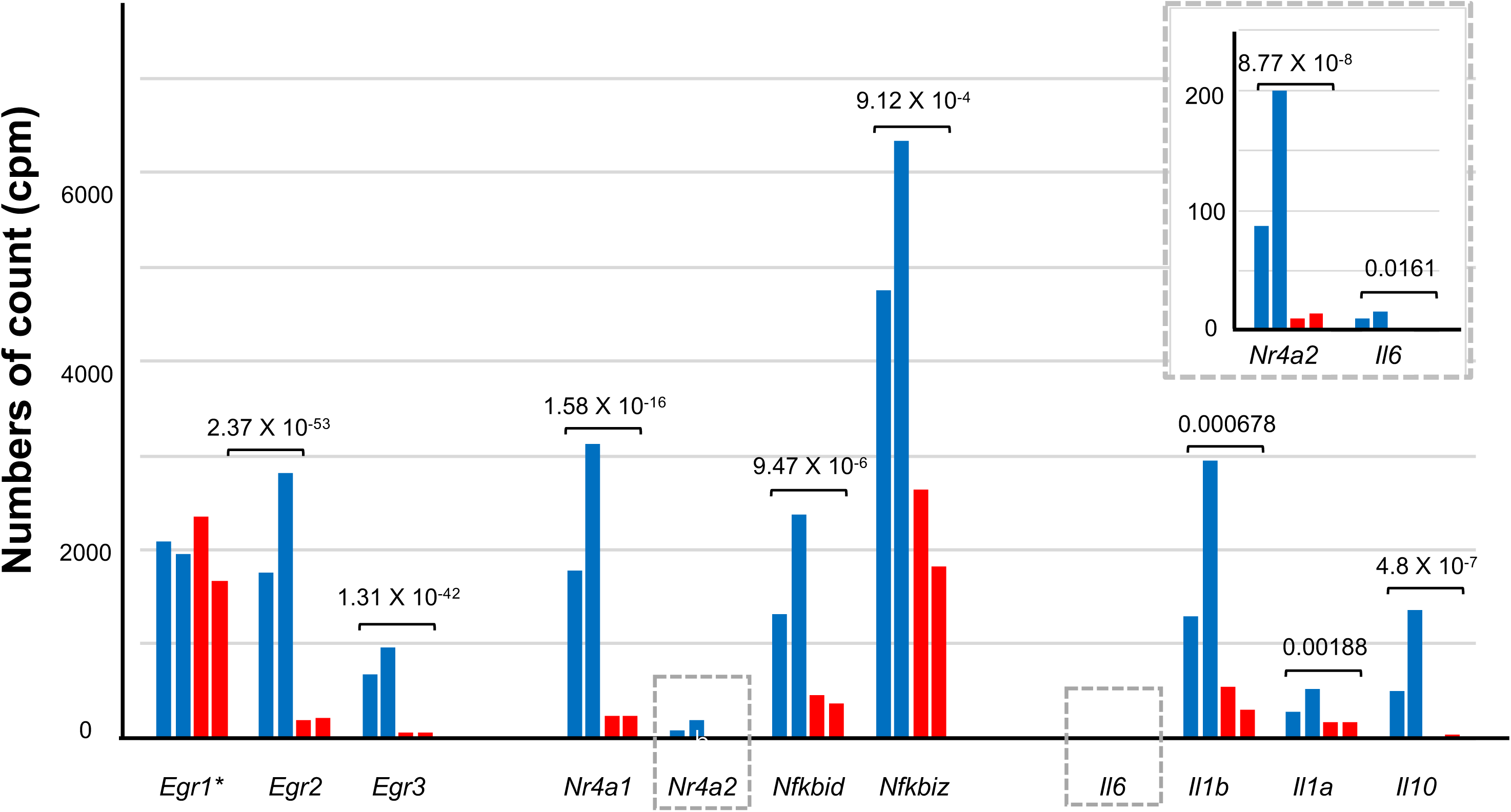
Transcriptional changes in *Rtl6*-deficient microglia. **(A)** Read coverage across the *Rtl6* locus. *Rtl6* is located on chromosome 15qE2 and is indicated by the red bar in the top track. In *Rtl6* KO mice, reads corresponding to the ORF were absent, whereas reads mapping to the remaining 5′ and 3′ untranslated regions (UTRs) were increased relative to WT controls (3.07-fold). WT1, WT2, KO1, and KO2: without LPS. WT3, WT4, KO3, and KO4: with LPS. **(B)** Heatmap of WT and *Rtl6* KO microglia following 10 min of LPS stimulation. Among the 3,540 significantly differentially expressed genes (Padj < 0.05), 2,060 were upregulated and 1,480 were downregulated. **(C)** Impaired MAPK–AP-1-mediated inflammatory responses in *Rtl6* KO microglia. \**Il6* expression was largely comparable between WT and KO microglia 1 h after LPS stimulation (Irie et al., 2022). **The preserved induction of *Egr1* in KO microglia may reflect its regulation by multiple upstream signaling pathways, including NF-κB, ERK–Elk1/SRF, calcium signaling, and general stress-response pathways, distinguishing it from the more AP-1-dependent regulation of *Egr2* and *Egr3* (Duclot and Kabbaj, 2017; Shan et al., 2019). **(D, E)** Changes in the LPS–TLR4 signaling pathway. **(D)** Upstream signaling components. **(E)** Downstream signaling components, including AP-1 family genes. Blue, WT; red, *Rtl6* KO. Adjusted *P* values (Padj) are shown for each gene. AP-1 component genes were significantly downregulated, whereas *Tlr4* (Padj = 0.376), *Nfkb1* (Padj = 0.662), and *Rela* (Padj = 0.735) showed no significant changes. Because *Nr4a2* and *Il6* were expressed at relatively low levels, they are additionally shown on an expanded scale (gray dashed box, upper right).

Unexpectedly, analysis of all samples revealed a marked difference in the extent of gene upregulation between the two untreated KO samples (KO1 and KO2), whereas widespread downregulation was consistently observed across all KO samples (Supplementary Fig. 1). These observations indicate substantial inter-sample variability among KO microglia under unstimulated conditions (see Discussion). We therefore focused subsequent analyses on LPS-induced transcriptional responses and compared the effects of LPS treatment between WT and KO microglia.

### Attenuation of immediate-early and interferon-responsive transcriptional programs

*Rtl6* KO microglia exhibited marked attenuation of immediate-early, inflammatory, and interferon-associated transcriptional programs. Comparison of gene expression profiles between WT and KO microglia following 10 min of LPS stimulation (WT3 and WT4 versus KO3 and KO4) identified 3,540 significantly differentially expressed genes (adjusted p-value < 0.05; Figure 1B; Table S1). To examine transcriptional changes under LPS-stimulated conditions in more detail, all differentially expressed genes, including 1,480 downregulated and 2,060 upregulated genes, were initially classified into 25 functional groups (Groups A–Y, Tables S2 and S3) and subsequently reorganized into five integrated functional modules (Modules 1–5, Table 1A and 1B) according to their biological relationships. These integrated modules capture the major transcriptional programs associated with RTL6 deficiency following LPS stimulation.

**Table 1.**
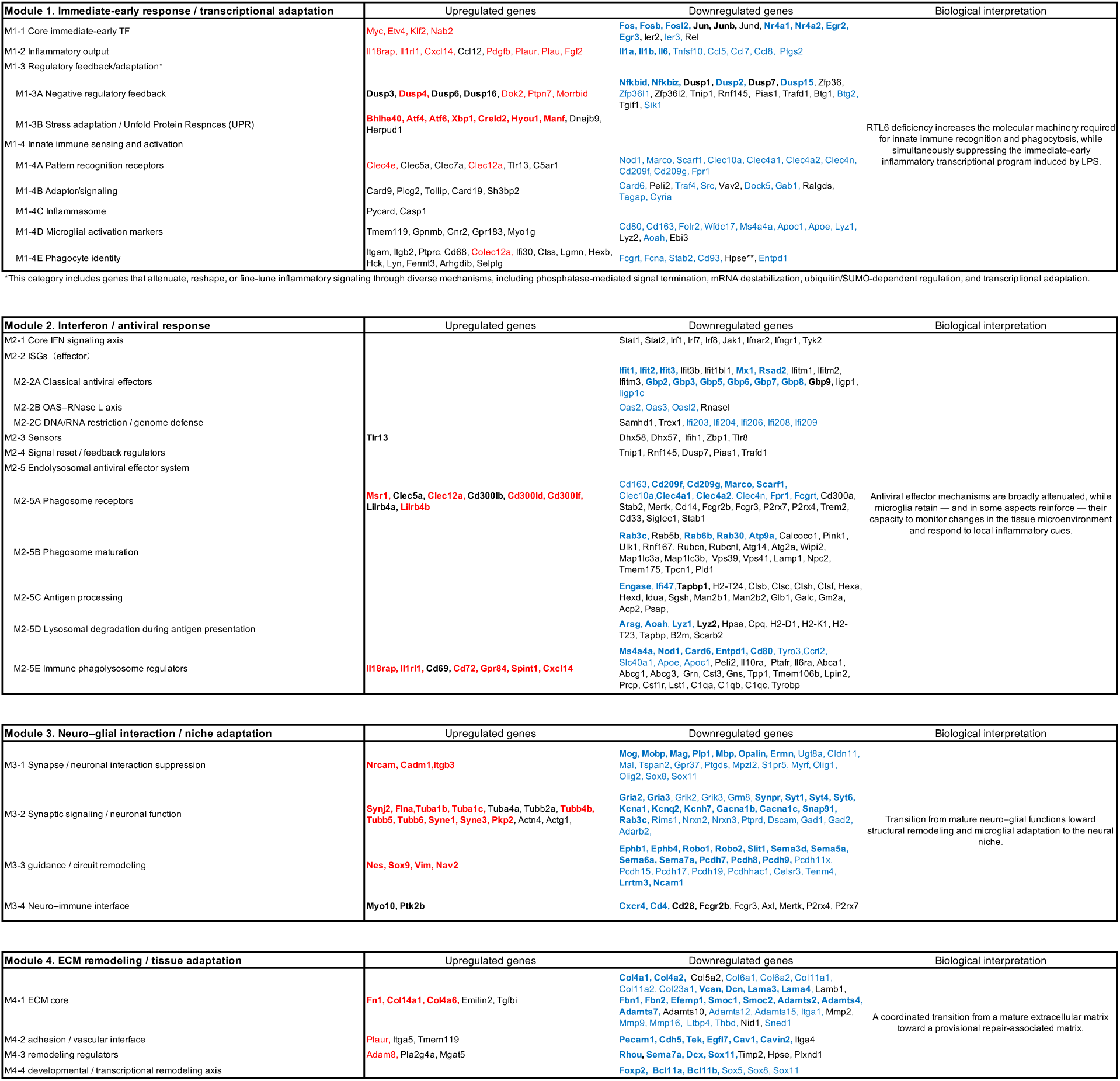
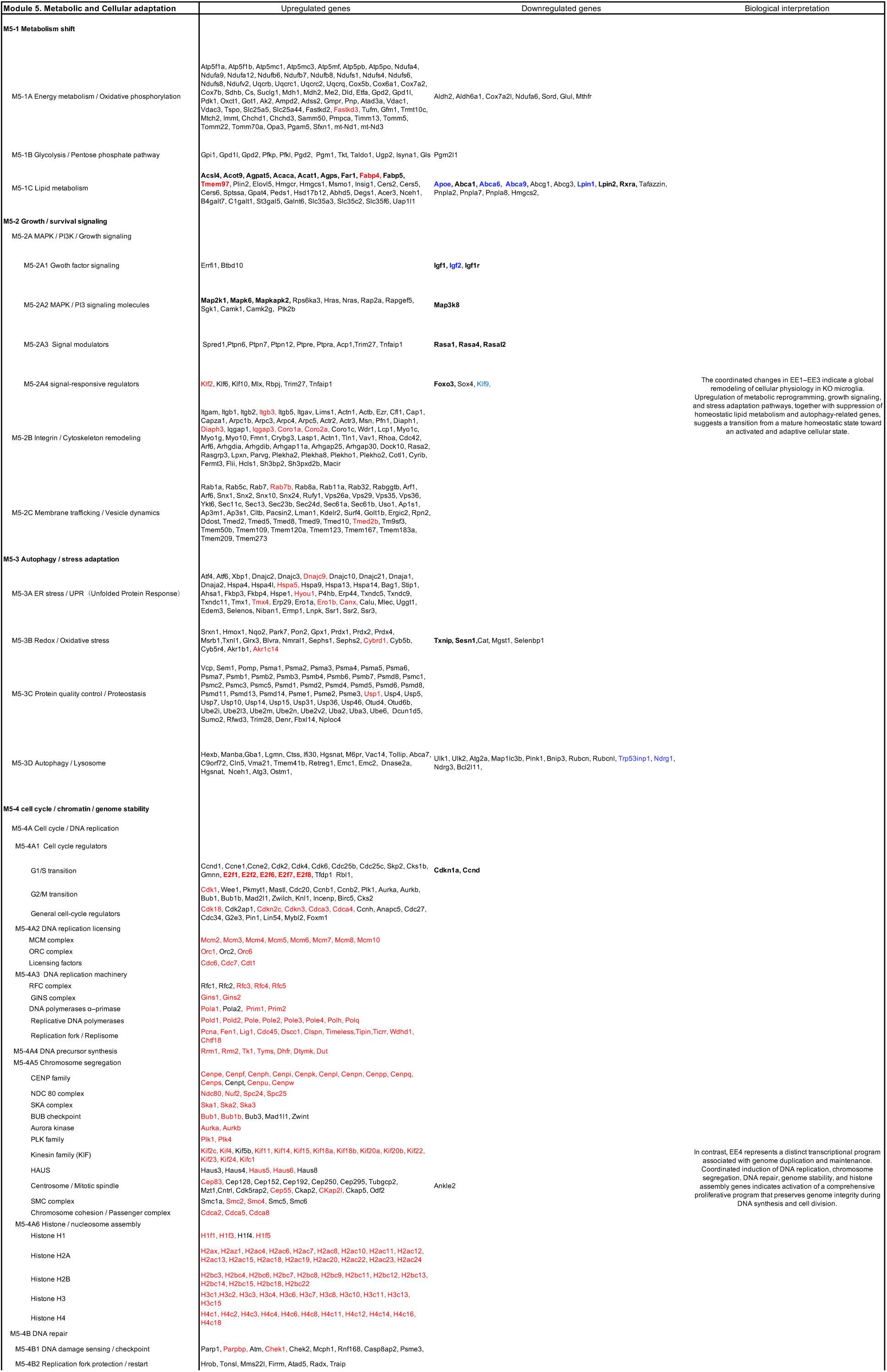
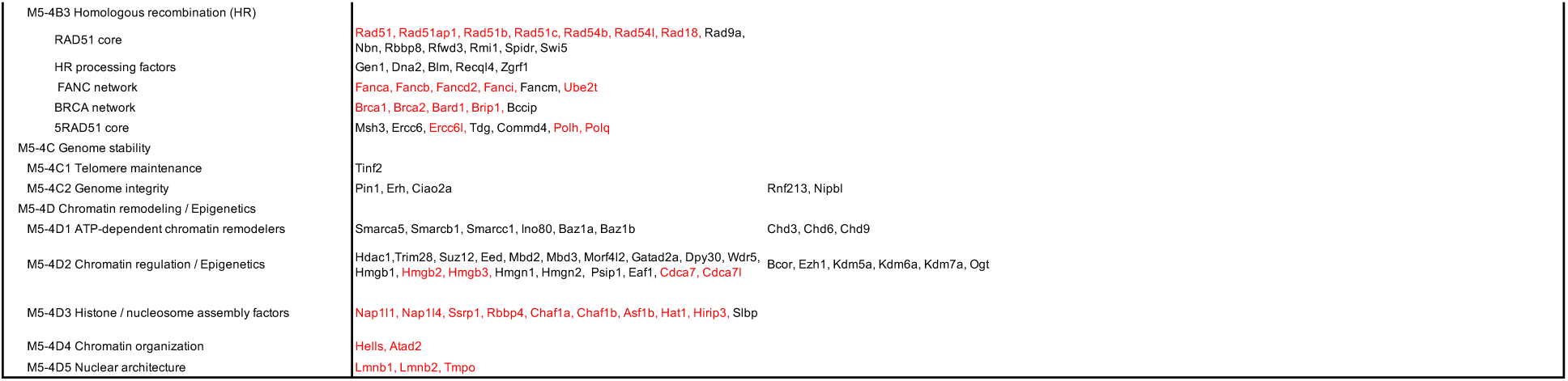
Five integrated functional modules characteristic of *Rtl6*-deficient microglia. (A) Modules 1–4 and (B) Module 5. Differentially expressed genes were classified into five integrated biological modules. The classification criteria differ slightly between Modules 1–4 and Module 5. Whereas Modules 1–4 are organized primarily according to biological processes, Module 5 is organized according to homeostatic programs associated with cellular adaptation. Some genes appear in more than one module because the classification is based on complementary biological perspectives (functional role versus regulatory program), rather than on mutually exclusive gene assignment. Genes shown in red and blue were upregulated or downregulated by more than twofold, respectively. Bold gene names indicate those discussed in the main text.

We first examined genes associated with the canonical LPS response (Modules 1 and 2). Module 1 was designed to represent the functional progression of the early inflammatory response. It begins with immediate-early transcription factor activation (M1-1), followed by inflammatory effector output (M1-2), then regulatory feedback and adaptive mechanisms that fine-tune signal intensity and duration (M1-3), and finally culminates in innate immune sensing and effector programs characteristic of activated microglia (M1-4). No significant change was detected in the expression of *Tlr4*, the major receptor for LPS (Kawai and Akira, 2010) (Figure 1C). However, the canonical immediate-early transcriptional network was markedly suppressed, with particularly large reductions in *Fos, Fosb, Jun, Junb, Nr4a1, Nr4a2, Egr2,* and *Egr3* (M1-1; Table 1A; Figures 1C–1E). In addition, several NF-κB-dependent feedback regulators, including *Nfkbid* and *Nfkbiz*, were also reduced (Fujioka *et al*., 2004; Tullai *et al*., 2011) (M1-3A; Figures 1C and 1E). Consistent with attenuation of these early transcriptional programs, expression of inflammatory effector genes, including *Il6, Il1b,* and *Il1a,* was significantly decreased (M1-2; Figures 1C and 1E). Within this framework, AP-1, composed of JUN and FOS family proteins, functions as a key convergence node integrating inputs from multiple MAPK branches, including ERK- and JNK-mediated pathways (Shaulian and Karin, 2002). ERK signaling primarily promotes the transcriptional induction of FOS, whereas JNK signaling enhances the activity of JUN through phosphorylation, and the combined activity of these components determines the magnitude of AP-1–dependent transcriptional output (Murphy *et al*., 2002; Weston and Davis, 2007).

A distinct set of stress-adaptive transcriptional regulators, including *Bhlhe40, Atf4, Atf6, Xbp1, Creld2, Hyou1*, and *Manf*, was upregulated (M1-3B). The numbers of up- and down-regulated genes are comparable in Module 1 (Table 1A). These findings indicate that RTL6 deficiency selectively attenuates the canonical immediate-early inflammatory transcriptional cascade while simultaneously promoting a stress-adaptive transcriptional program characterized by activation of the unfolded protein response (UPR) and extensive remodeling of innate immune signaling networks. Consistent with this model, differential regulation of DUSP family members further supports a selective alteration of downstream MAPK transcriptional output rather than upstream signaling itself. Immediate-early phosphatases such as *Dusp1* and *Dusp2*, along with additional MAPK-associated phosphatases including *Dusp7* and *Dusp15*, are reduced (M1-3A), consistent with impaired AP-1/IEG-dependent induction, whereas ERK-associated feedback regulators including *Dusp3*, *Dusp4*, *Dusp6*, and *Dusp16* are upregulated (M1-3A), consistent with preserved basal ERK signaling and compensatory feedback regulation (Caunt *et al*., 2008; Ham *et al*., 2015; Lake *et al*., 2016) (Supplementary Fig. 2). This divergence highlights a decoupling between stimulus-induced MAPK transcriptional responses and homeostatic MAPK feedback regulation. In the present study, although upstream MAPK signaling appears to be at least partially preserved—as suggested by relatively maintained ERK-associated signaling, including preserved *Egr1* expression (Duclot and Kabbaj, 2017)—the knockout condition impairs this convergence at the level of AP-1. This is evidenced by reduced *Jun* and *Fos* expression, accompanied by a marked attenuation of secondary immediate-early genes, including *Egr2* and *Egr3* (Figure 1C–1E).

Module 2 represents the sequential organization of the interferon-dependent antiviral response, encompassing interferon signaling (M2-1), antiviral effector induction (M2-2), intracellular nucleic acid sensing (M2-3), regulatory feedback (M2-4), and the endolysosomal antiviral effector system responsible for pathogen degradation and antigen processing (M2-5). Module 2 revealed coordinated suppression of the interferon-dependent antiviral response across all functional layers. Genes involved in interferon signaling, antiviral effector mechanisms, including *Ifit1–3, Mx1, Rsad2, and Gbp2–9* (M2-2A), pathogen-recognition and phagocytic receptors, including *Cd209f, Cd209g, Marco, Scarf1, Clec4a1, Clec4a2, Fpr1,* and *Fcgrt* (M2-5A), phagosome maturation, including *Rab3c, Rab6b, Rab30,* and *Atp9a* (M2-5B), lysosomal degradation and antigen processing, including *Engase, Ifi47,* and *Tapbp* (M2-5C), lysosomal degradation and antigen processing, including *Arsg, Aoah, Lyz1* and *Lyz2* (M2-5D), together with immune phagolysosome regulators, including *Ms4a4a, Nod1, Card6, Entpd1,* and *Cd80* (M2-5E), were broadly downregulated, indicating that RTL6 deficiency attenuates the antiviral defense network as an integrated functional program rather than affecting a single signaling pathway.

In contrast, pathogen and tissue sensing underwent extensive remodeling rather than being uniformly suppressed. As described above, several scavenger and phagocytic receptors, including *Marco, Scarf1,* Cd209 family members, and *Fcgrt*, were broadly downregulated (M2-5A), other environmental sensing receptors, including *Tlr13* (M2-3)*, Clec5a, Clec12a*, members of the Cd300 family, *Lilrb4a, Lilrb4b*, and *Msr1*, were maintained or induced (M2-5A). Thus, although the repertoire of pathogen-sensing receptors was substantially reorganized, the capacity for environmental surveillance was retained. Consistent with this pattern, immune regulatory molecules associated with tissue surveillance, including *Il18rap, Il1rl1 (ST2), Cd69, Cd72, Gpr84, Spint1,* and *Cxcl14*, were selectively upregulated (M2-5E). Totally, the number of downregulated genes is much larger than that of upregulated genes (Table 1A). Together, these findings indicate that RTL6 deficiency does not abolish innate immune sensing but instead functionally uncouples pathogen recognition from downstream antiviral effector functions, thereby shifting microglia toward an immune surveillance state characterized by selective environmental sensing and attenuated antimicrobial effector activity.

### Reduction of interaction between microglia and neuronal and other glial cells

Modules 3 and 4 revealed an additional layer of transcriptional rewiring involving microglial interactions with neighboring cells and the extracellular environment. Total number of downregulated genes is much larger than that of upregulated genes in Modules 3 and 4, like Module 2 (Table 1A). Module 3 represents functional interactions between microglia and neural tissue, including communication with neurons, oligodendrocytes, astrocytes, vascular cells, and other glial populations. Within Module 3, numerous genes associated with oligodendrocyte and myelin interactions, including *Mog, Mobp, Mag, Plp1, Mbp, Opalin,* and *Ermn,* were markedly downregulated (M3-1). Genes involved in synaptic signaling and neuronal function (M3-2), including *Gria2, Gria3, Synpr, Syt1, Syt4, Syt6, Kcna1, Kcnq2, Kcnh7, Cacna1b, Cacna1c, Snap91*, and *Rab3c*, together with genes associated with neuronal guidance and circuit remodeling (M3-3), including *Ephb1, Ephb4, Robo1, Robo2, Slit1, Sema3d, Sema5a, Pcdh7, Pcdh8, Pcdh9, Lrrtm3,* and *Ncam1*, were also broadly downregulated. Similarly, genes involved in the neuro–immune interface (M3-4), including *Cxcr4, Cd4, Cd28,* and *Fcgr2b*, showed reduced expression. These changes are consistent with the marked suppression of neuronal activity-dependent immediate-early genes, including *Fos, Fosb, Jun, Junb, Nr4a1, Nr4a2, Egr2,* and *Egr3* (Module 1 and Group D-7 in Table S2), indicating reduced responsiveness to neuronal activity (Schafer *et al*., 2012; Tyssowski *et al*., 2018; Safaivan *et al*., 2021).

In contrast, RTL6-deficient microglia upregulated a distinct set of genes associated with structural interactions and tissue adaptation. These included neuron–glia and interglial adhesion molecules (M3-1), such as *Cadm1, Nrcam,* and *Itgb3*; genes involved in synaptic organization and cytoskeletal remodeling (M3-2), including *Synj2, Flna, Tuba1b, Tuba1c, Tubb4b, Tubb5, Tubb6, Syne1, Syne3*, and *Pkp2*; neuronal guidance and structural remodeling genes (M3-3), including *Nes, Sox9, Vim,* and *Nav2*; and neuro–immune interface genes (M3-4), including *Myo10* and *Ptk2b*. Together, these findings indicate that RTL6 deficiency is associated with extensive remodeling of neuro–glial communication rather than a simple loss of neural interactions. While transcriptional programs linked to neuronal activity, synaptic communication, myelin interactions, and neuroimmune signaling were broadly attenuated, genes involved in structural adhesion, cytoskeletal organization, and trophic interactions were selectively induced, suggesting a transition from activity-responsive communication toward structural adaptation within the neural tissue niche.

### Remodeling of interactions between microglia and the extracellular matrix

Module 4 represents ECM remodeling, cell adhesion, and tissue adaptation pathways associated with changes in the local microenvironment. Numerous genes encoding core ECM components (M4-1), including *Col4a1, Col4a2, Vcan, Dcn, Lama3, Lama4, Fbn1, Fbn2, Efemp1, Smoc1, Smoc2, Adamts2, Adamts4, Adamts7, Itga1, Mmp9, Mmp16, Ltbp4, Thbd,* and *Sned1,* were markedly downregulated. Genes involved in cell adhesion and the vascular interface (M4-2), including *Pecam1, Cdh5, Tek, Egfl7, Cav1,* and *Cavin2,* were likewise reduced. In addition, remodeling regulators (M4-3), including *Rhou, Sema7a, Dcx,* and *Sox11*, together with transcriptional regulators involved in developmental remodeling (M4-4), including *Foxp2, Bcl11a* and *Bcl11b*, were also downregulated. In contrast, RTL6-deficient microglia selectively upregulated genes associated with ECM remodeling, cell–matrix adhesion, and tissue adaptation, including *Fn1, Col14a1,* and *Col4a6* (M4-1); *Plaur, Itga5,* and *Tmem119* (M4-2); and *Adam8, Pla2g4a,* and *Mgat5* (M4-3).

Together, these findings indicate that RTL6 deficiency is associated with coordinated remodeling of the extracellular matrix rather than generalized ECM activation. The simultaneous downregulation of mature basement membrane and vascular ECM components together with selective induction of fibronectin-associated matrix components and adhesion molecules suggests a transition toward a provisional, tissue-remodeling extracellular environment.

### Coordinated activation of metabolic adaptation and genome maintenance programs

Module 5 revealed a highly coordinated transcriptional program coupling metabolic remodeling with cell-cycle progression, DNA replication, and genome maintenance (Table 1B). In contrast to Modules 2–4, upregulated genes greatly outnumbered downregulated genes. Moreover, genes involved in metabolism, intracellular signaling, membrane trafficking, and proteostasis (Modules 5-1 to 5-3) were generally induced with modest fold changes, whereas genes involved in DNA replication, chromosome segregation, and chromatin organization (Module 5-4) exhibited substantially stronger induction. This pattern indicates coordinated remodeling of cellular physiology accompanied by activation of a proliferative transcriptional program. Because this module encompasses extensive metabolic and proliferative gene programs, detailed gene classifications and representative genes for each submodule are presented in the Supplementary Result 1.

Genes involved in oxidative phosphorylation (Complexes I–V), the tricarboxylic acid cycle, and mitochondrial bioenergetic and biogenesis pathways were broadly upregulated (M5-1A), indicating coordinated enhancement of mitochondrial bioenergetic capacity. These changes were accompanied by coordinated induction of glycolytic and pentose phosphate pathway enzymes (M5-1B), together with genes supporting glucose utilization and glutamine metabolism. Lipid metabolism was also extensively remodeled, with increased expression of genes involved in fatty acid activation, phospholipid synthesis, cholesterol biosynthesis, and membrane lipid production (M5-1C), including *Acsl4, Acot9, Agpat5, Acaca, Acat1, Agps, Far1, Fabp4, Fabp5*, and *Tmem97*, whereas genes associated with lipid storage and cholesterol efflux, such as *Apoe, Abca1, Abcg1, Lpin1, Lpin2,* and *Rxra,* were reduced, consistent with a shift from lipid homeostasis toward membrane biosynthesis.

Components of the MAPK-, PI3K-, small GTPase-, and calcium-dependent signaling pathways were coordinately induced together with multiple signaling modulators (M5-2A), such as *Map2k1, Mapk6,* and *Mapkapk2*. In contrast, *Map3k8* (*Tpl2*) (M5-2A2), the onlycanonical upstream MAPKKK of the TLR4–TPL2–ERK–AP-1 signaling axis, was significantly downregulated, accompanied by coordinated downregulation of multiple AP-1- and ERK-responsive immediate-early genes (e.g., *Jun/Junb/Jund, Fos/Fosb, Egr2/Egr3, Nr4a1/Nr4a2,* and *Dusp1*) as discussed in Module 1-1 (Dumitru et al., 2000; Ganke et al., 2012; Auther and Ley, 2013). The IGF– PI3K/AKT–FOXO signaling axis was also attenuated, as indicated by reduced expression of *Igf1, Igf1r, Igf2,* (M5-2A1), *Rasa1, Rasa4, Rasal2* (M5-2A3)*, Foxo3* (M5-2A4), and several stress-responsive genes. The IGF–PI3K/AKT–FOXO signaling axis, including *Igf1, Igf1r, Igf2* (M5-2A1), *Rasa1, Rasa4, Rasal2,* (M5-2A3) and *Foxo3* (M5-2A4), and several stress-responsive genes were also downregulated (Webb and Brunet, 2014; Manning and Toker, 2017). M5-2B represents a coordinated signaling module linking cell adhesion to actin cytoskeletal remodeling. The concerted induction of integrin-mediated actin cytoskeletal remodeling genes suggests enhanced capacity for cell adhesion, migration, morphological remodeling, and phagocytosis in macrophages and microglia. Membrane trafficking pathways (M5-2C) were also broadly activated, including Rab- and Arf-family GTPases, sorting nexins, COPII-mediated ER-to-Golgi transport, retromer components, and vesicle transport machinery, indicating enhanced intracellular membrane dynamics.

Proteostasis pathways were similarly activated (M5-3). ER protein-folding machinery, molecular chaperones, protein disulfide isomerases, ER quality-control systems, redox defense pathways, and ubiquitin–proteasome components were coordinately induced. Lysosomal pathways were likewise activated, whereas several canonical autophagy regulators were reduced, suggesting a shift toward lysosome-centered proteostatic adaptation with selective suppression of canonical autophagy. Collectively, Modules 5-1 to 5-3 indicate extensive remodeling of metabolism, intracellular signaling, membrane trafficking, and proteostasis, establishing a cellular state associated with activation of the proliferative genome-maintenance program described below.

Module 5-4 defined an integrated transcriptional program encompassing cell-cycle progression, DNA replication, genome maintenance, and chromatin remodeling. Genes regulating G1/S transition, G2/M transition, and general cell-cycle progression were broadly induced (M5-4A1). Notably, all major E2F transcription factors (*E2f1, E2f2, E2f6, E2f7,* and *E2f8*) were induced by more than twofold, indicating coordinated activation of the E2F transcriptional program controlling S-phase entry and DNA replication. Consistent with this, *Cdkn1a* was downregulated.

Genes required for DNA replication licensing, replication fork progression, DNA precursor synthesis, chromosome segregation, DNA repair, genome surveillance, and replication-coupled chromatin assembly were coordinately upregulated. Particularly prominent induction was observed for replication-coupled histone deposition factors, ATP-dependent chromatin remodelers, epigenetic regulators, chromatin organization factors, and nuclear architecture proteins, indicating activation of a comprehensive program coupling DNA replication with chromatin duplication and genome maintenance.

Together, these coordinated transcriptional changes demonstrate activation of an integrated genome-duplication program encompassing cell-cycle progression, DNA replication, chromosome segregation, DNA repair, genome surveillance, and replication-coupled chromatin assembly. Rather than reflecting independent pathway-specific responses, these coordinated transcriptional changes define a comprehensive molecular program supporting faithful genome duplication during cell proliferation (see more details in Supplementary Result 1).

### Disease-associated microglia (DAM)- and apoptosis-related genes

Interestingly, the transcriptional changes induced by RTL6 deficiency resemble the DAM program. For example, several genes associated with the *Spp1/Gpnmb/Fabp5* tissue-remodeling module were upregulated (Figure 2A), whereas components of the canonical *Trem2–Tyrobp–Apoe* signaling axis (Figure 2B), together with *Il10ra*, were downregulated (Group A1b, Tables S2 and S3) (Keren-Shaul *et al*., 2017; Krasemann *et al*., 2017; Prinz *et al*., 2019). The DAM-associated receptor *Axl* was also decreased, whereas the homeostatic microglial marker *Tmem119* was modestly increased. In addition, no significant changes were detected in two other characteristic DAM markers, *Cst7* and *Ctsz*. These findings indicate that RTL6-deficient microglia exhibit a non-canonical DAM-like transcriptional profile characterized by selective activation of tissue-remodeling pathways without coordinated induction of the core *Trem2*-dependent signaling network (Figure 2A and 2B).

**Figure 2.**
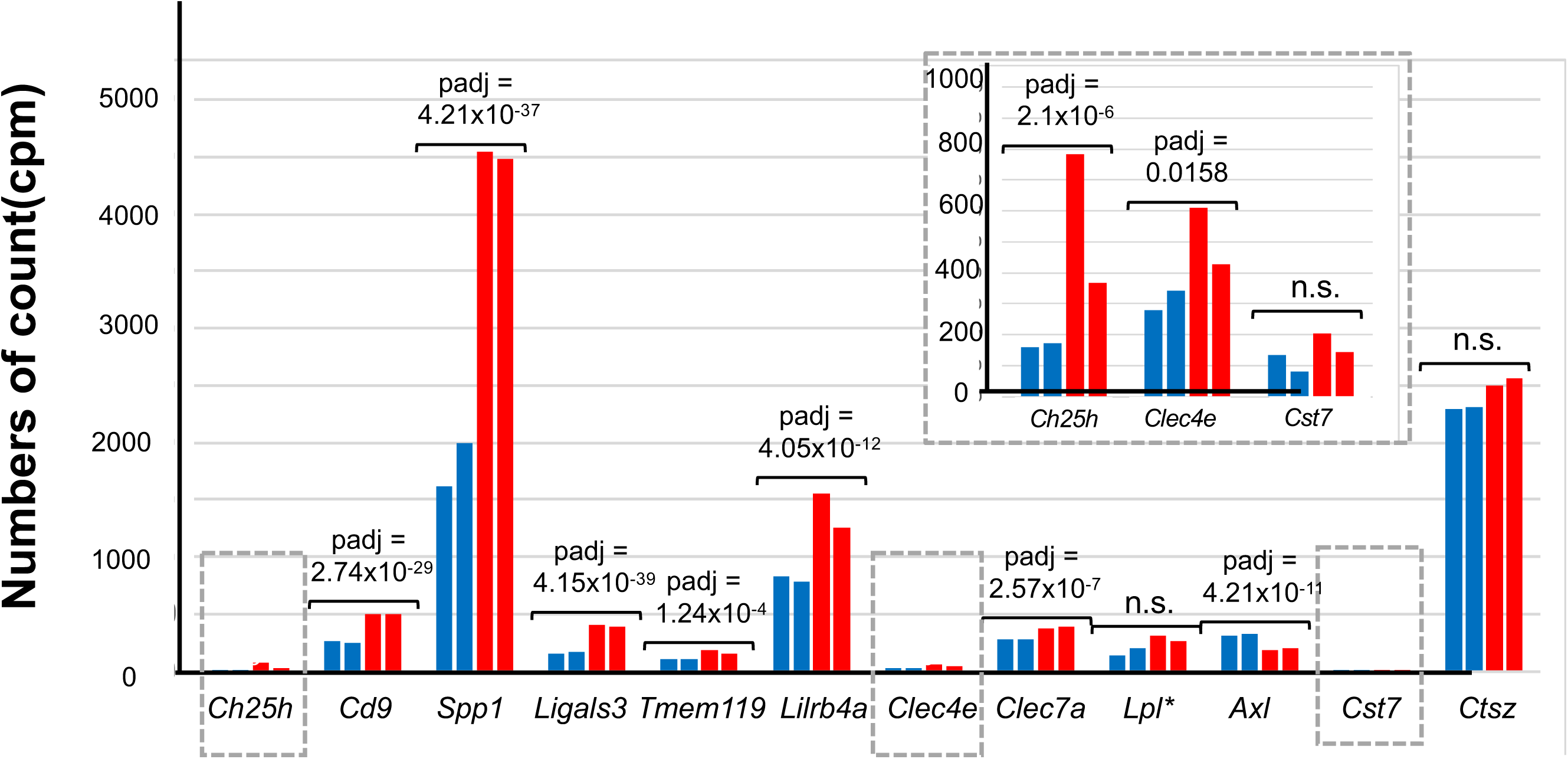
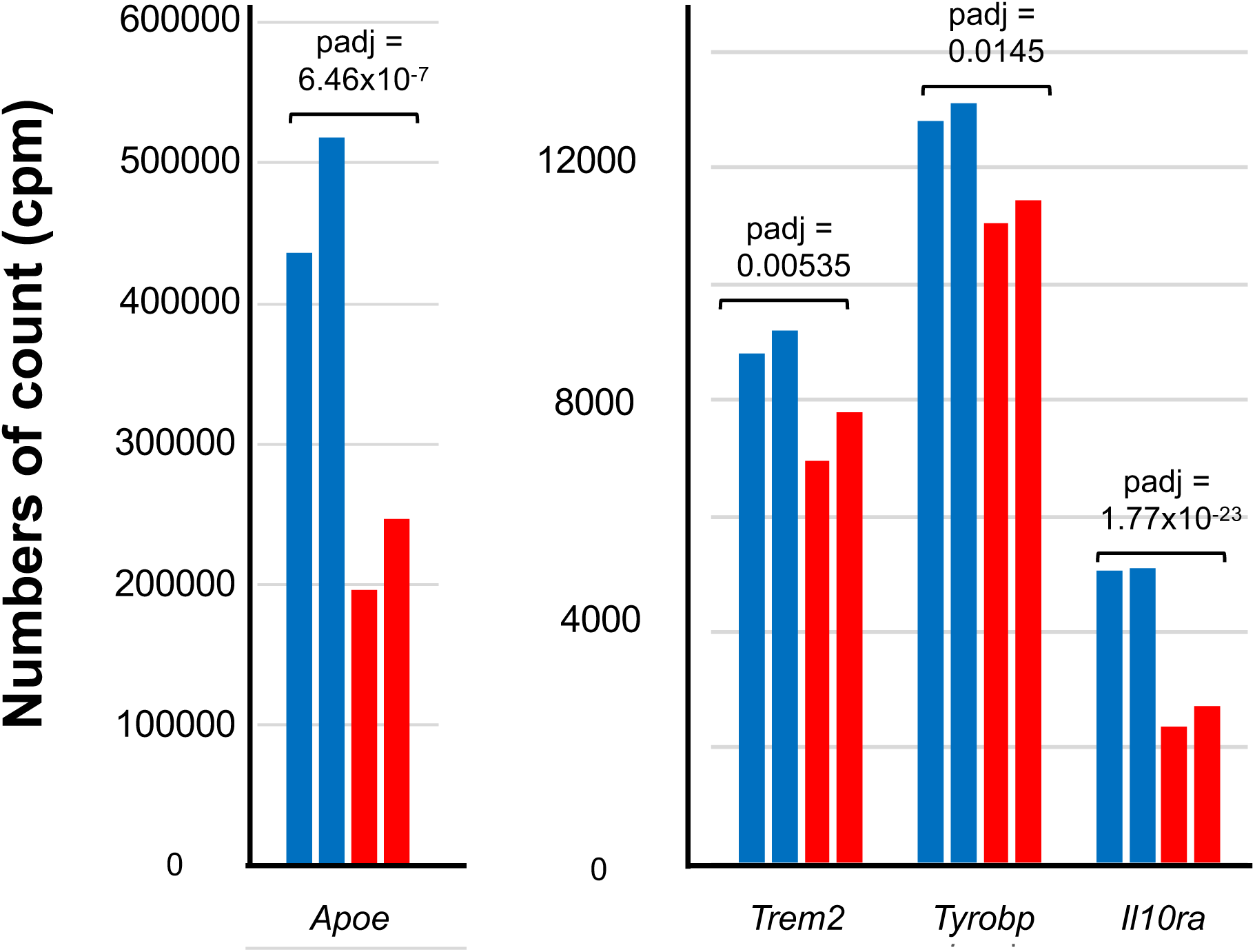
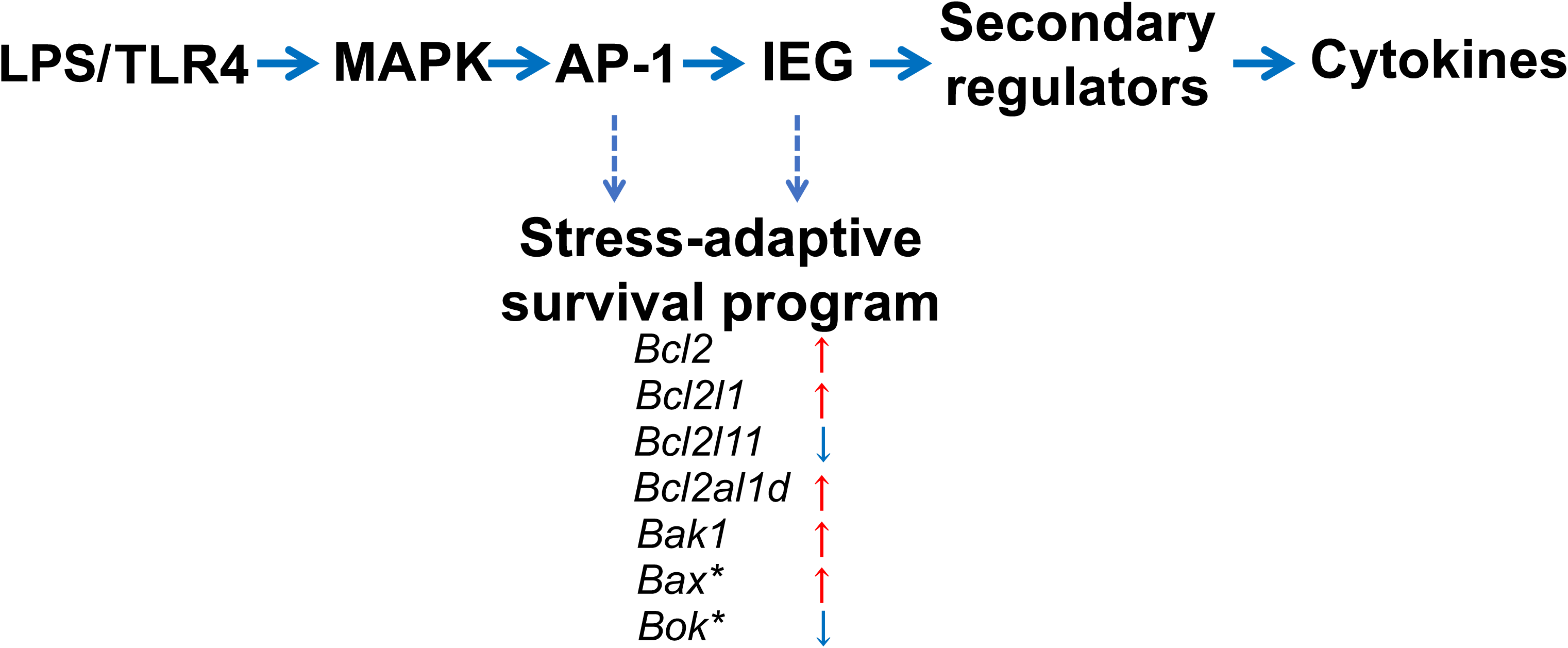
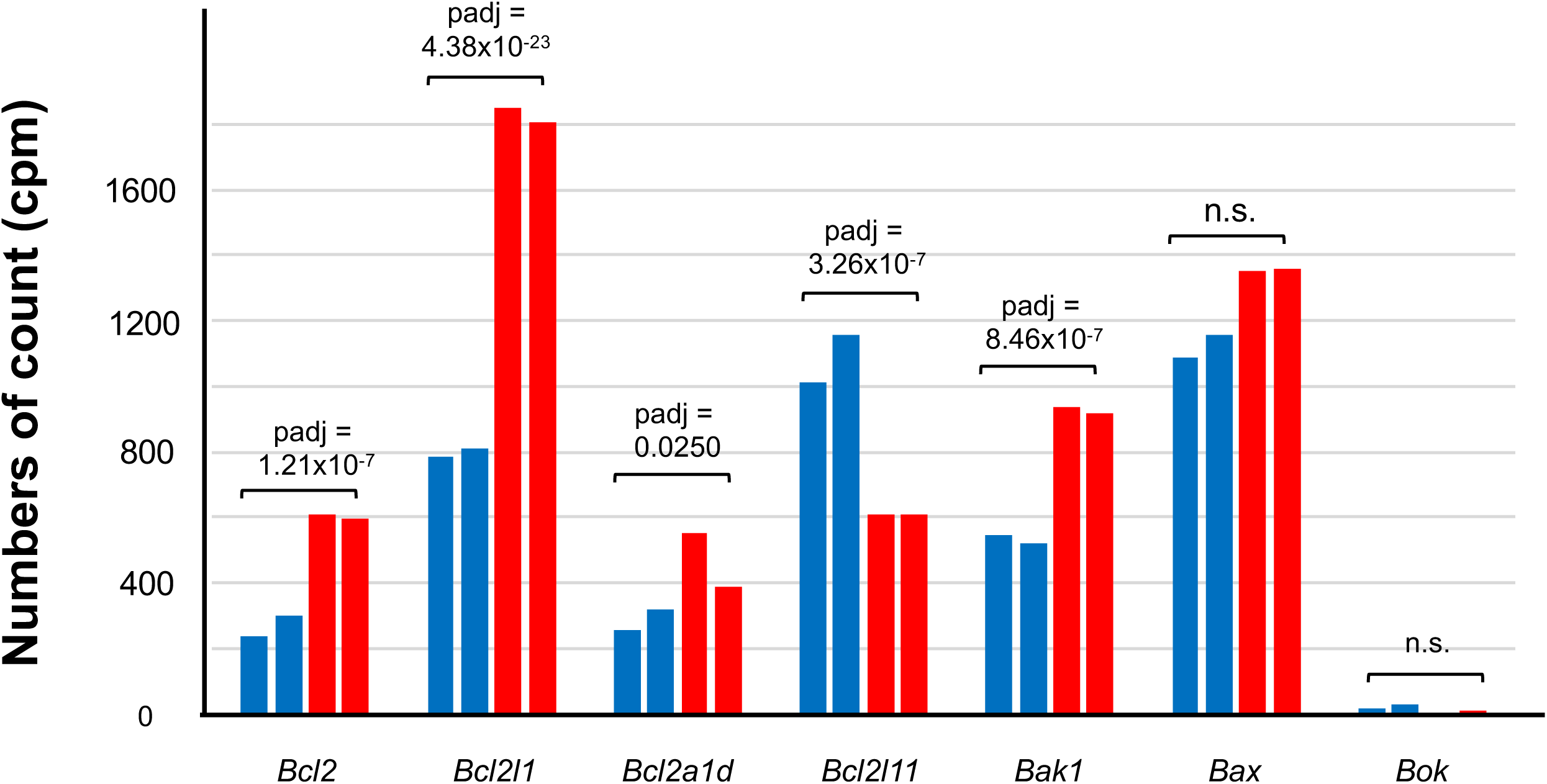
Transcriptional changes in DAM-related and apoptosis-related genes. **A** and **B**. Damage-associated microglia (DAM)-related genes. Several DAM-related genes were upregulated in Rtl6 KO, however, typical DAM genes, such as Cts7 and Ctsz exhibited no significant changes (A). Especially, *Trem2–Tyrobp–Apoe* signaling axis was downregulated (B), suggesting the observed change differs from the canonical DAM responses. Expression of Ch25h, Clec4e and Cst7, is small, therefore, they are also shown in the expanded scale (gray dashed box on the top right). **C** and **D**. *Bcl2-*related apoptosis genes. Their expression patterns suggest altered regulation of stress-responsive apoptotic pathways in *Rtl6*-deficient microglia.

We further examined genes associated with apoptosis and cell survival. Anti-apoptotic genes, including *Bcl2, Bcl2l1,* and *Bcl2a1d*, were upregulated together with the pro-apoptotic factors *Bak1*, whereas the BH3-only gene *Bcl2l11* was downregulated (Figure 2C and 2D, I1and I2, Tables S2 and S3). These coordinated changes indicate extensive remodeling of apoptosis- and survival-related pathways rather than activation of a canonical apoptotic program (Pihán *et al*., 2017; Youle and Strasser, 2008). Such reprogramming is consistent with a broader decoupling between stimulus-induced transcriptional responses and intrinsic cellular homeostasis (Wendeln *et al*., 2018).

## 3. Discussion

A major finding of the present study is that RTL6 deficiency attenuated the earliest transcriptional response to LPS and was accompanied by widespread reorganization of microglial transcriptional programs, rather than disruption of a single signaling pathway. Our previous studies demonstrated that the RTL6 protein is secreted into the brain extracellular space, where it directly binds bacterial LPS and contributes to LPS clearance *in vivo*. However, despite impaired LPS handling in *Rtl6* KO mice, cultured KO microglia exhibited largely normal induction of *Tnfa* and *Il6* one hour of LPS stimulation, leaving unresolved how RTL6 influences TLR4-mediated signaling (Irie et al., 2022).

To address this question, we focused on the earliest phase of the LPS response and analyzed transcriptomic changes only 10 min of LPS stimulation. Notably, the induction of immediate-early transcription factors, including *Egr2*, *Egr3*, *Nr4a1*, *Nr4a2*, *Fos* and *Fosb*, together with multiple inflammatory effector genes, were markedly attenuated in KO microglia (Figure 1C–1E). These findings indicate that RTL6 contributes to efficient initiation of the TLR4-dependent transcriptional program and suggest a broader role for RTL6 in regulating microglial activation than previously recognized. One possibility is that internalization of the RTL6–LPS complex facilitates optimal intracellular signaling following ligand recognition. Alternatively, RTL6 may possess additional functions independent of direct LPS binding, potentially through currently unidentified interacting proteins that influence transcriptional regulation. Although these mechanisms remain speculative, they are consistent with the approximately threefold increase in residual *Rtl6* untranslated-region transcripts observed in KO microglia, suggesting feedback regulation of *Rtl6* transcription itself (Figure 1A). Such feedback further implies that precise control of *RTL6* expression is important for maintaining microglial homeostasis and regulating transitions between functional states. From an evolutionary perspective, these findings may help explain the exceptional conservation of *RTL6* throughout eutherian mammals despite its origin as a metavirus-derived gene.

Unexpectedly, our transcriptomic analyses revealed large-scale but highly coordinated activation of proliferative, metabolic, and tissue-remodeling programs together with altered inflammatory responsiveness. However, it remains unclear whether these changes represent direct consequences of RTL6 deficiency or secondary adaptations that accumulated during long-term maintenance of the knockout line. The *Rtl6* KO mice used in this study have been maintained for many generations (Irie et al., 2022), raising the possibility that compensatory transcriptional reprogramming contributed to the observed phenotype. In contrast, attenuation of the immediate-early response immediately following LPS stimulation is more likely to reflect a primary defect in signaling responsiveness. Future studies employing inducible knockout, acute knockdown, and rescue experiments will therefore be important for distinguishing primary functions of *RTL6* from long-term adaptive changes.

Several experimental limitations should also be considered. First, the transcriptomic analyses were performed using primary cultured microglia rather than microglia directly isolated from the brain. Although these cultures preserve many intrinsic characteristics of microglia, they lack the complex cellular interactions normally provided by neurons, oligodendrocytes, vascular cells, and the intact extracellular environment. Given that microglial states are highly shaped by local cellular and extracellular cues (Butovsky *et al*., 2014; De Biase *et al*., 2017; Gosselin *et al*., 2017), the transcriptional changes observed here may be influenced by *in vitro* conditions and may not fully recapitulate *in vivo* conditions. Moreover, the KO microglia were maintained on *Rtl6*-deficient astrocyte feeder cells. Although RTL6 expression was not detected in astrocytes (Irie et al., 2022), the use of KO feeder cells raises the possibility that indirect alterations in the feeder-cell environment contributed to part of the observed transcriptional phenotype. It will therefore be important to determine whether similar phenotypes are observed when KO microglia are cultured on wild-type astrocyte feeders or examined directly *in vivo*. Future studies using acute brain isolation or single-cell transcriptomics will help clarify the contribution of the native brain microenvironment to the *RTL6*-dependent phenotype.

Another limitation is the restricted temporal scope of the present study. Transcriptomic analyses were performed only 10 min of LPS stimulation in order to capture the immediate-early response. However, our previous study showed that *Il6* induction in KO microglia was largely comparable to WT levels 1 h after stimulation (Irie et al., 2022), suggesting that that the early transcriptional differences observed here may reflect altered response kinetics rather than complete loss of signaling capacity. Whether the extensive transcriptional reprogramming identified in the present study follows a similar temporal trajectory remains to be determined. Future time-course transcriptomic analyses will therefore be important for defining the temporal progression of RTL6-dependent microglial state transitions.

An additional consideration is the unexpectedly large transcriptional difference observed between the two untreated KO samples. Owing to this variability, we were unable to define a precise basal transcriptional signature associated with RTL6 deficiency. Interestingly, one untreated KO sample exhibited widespread transcriptional activation resembling an LPS-responsive state while maintaining reduced expression of canonical immediate-early genes, arguing against inadvertent LPS contamination. Instead, this pattern may reflect spontaneous entry into a partially activated state driven by culture-associated stress or stochastic variation in microglial activation. These observations raise the possibility that RTL6 deficiency increases transcriptional instability under basal conditions, with LPS stimulation subsequently amplifying this latent variability. Additional biological replicates and detailed time-course analyses will be required to clarify the relationship between basal-state instability and stimulus-induced responses.

Despite these experimental limitations, loss of RTL6 was consistently associated with a substantial shift in microglial state following LPS stimulation compared to WT (Table 1A and 1B; Figures 1–2), indicating that RTL6 functions as a regulator of microglial state rather than merely facilitating extracellular LPS clearance. Microglia are a conserved population of CNS-resident macrophages across vertebrates, originating from yolk sac-derived progenitors (or functionally analogous hematopoietic tissues in fish) that colonize the CNS during early development (Ginhoux and Prinz, 2015; Shimizu and Prinz, 2025). Core identity genes, including *P2RY12, TMEM119, HEXB*, and *SALL1*, are broadly conserved from zebrafish to humans (Bennett et al., 2016; Masuda et al., 2020). Microglia exhibit intrinsic plasticity, transitioning between transcriptional states in response to environmental cues. In mammals, such states have been characterized as DAM (Keren-Shaul et al., 2017; Masuda et al., 2020); however, the capacity for state transitions itself represents a conserved vertebrate feature, whereas the underlying molecular programs and the organization of the microglial state likely diverge across species. Likewise, microglial contributions to neural circuit formation, including synaptic pruning, are broadly conserved across vertebrates, although the extent and underlying molecular mechanisms vary among species (Paolicelli *et al*., 2011; Schafer *et al*., 2012).

One of the most notable findings of the present study is that the transcriptional changes induced by RTL6 deficiency were not randomly distributed but instead formed a highly coordinated series of functional programs spanning inflammatory signaling, tissue adaptation, and cellular homeostasis. Module 1 revealed that RTL6 deficiency attenuates the canonical immediate-early inflammatory cascade while simultaneously inducing stress-adaptive transcriptional regulators associated with the unfolded protein response (UPR), indicating a redirection of the earliest inflammatory response rather than simple suppression. Module 2 further revealed broad suppression of interferon-dependent antiviral execution pathways while preserving, or even enhancing receptors involved in environmental sensing and immune surveillance. These observations suggest that *Rtl6*-deficient microglia retain the capacity to detect changes in the tissue environment but exhibit reduced commitment to full antimicrobial effector functions.

Modules 3 and 4 indicate that this altered immune state is accompanied by extensive remodeling of interactions with the neural microenvironment. Reduced expression of genes associated with neuronal activity, oligodendrocytes, and myelin was accompanied by increased expression of trophic factors, cell-adhesion molecules, and extracellular matrix remodeling genes, suggesting a transition from inflammatory activation toward structural adaptation within the neural niche. These findings suggest that RTL6 deficiency shifts microglia from an inflammatory effector state toward a tissue-remodeling phenotype specialized for communication with the neural microenvironment.

The most unexpected observation was the robust activation of Module 5. The coordinated M5-1–M5-3 responses suggest that RTL6 deficiency induces a systems-level metabolic and physiological remodeling rather than isolated changes in inflammatory signaling. Enhanced mitochondrial metabolism, glycolysis, lipid biosynthesis, intracellular membrane trafficking, and proteostasis collectively indicate that KO microglia adopt a metabolically active state capable of supporting increased biosynthetic and proliferative demands. At the same time, suppression of homeostatic lipid metabolism, IGF signaling, and canonical autophagy suggests departure from the resting microglial phenotype toward a stress-adapted cellular state.

A particularly striking observation was that these metabolic and adaptive changes closely parallel the extensive activation of DNA replication, chromatin assembly, and genome maintenance pathways observed in M5-4. *Rtl6*-deficient microglia coordinately induced genes involved in DNA replication licensing, replication fork progression, chromosome segregation, histone synthesis, chromatin assembly, DNA repair, and genome maintenance. Such coordinated activation is unlikely to reflect cell-cycle entry alone. Instead, it suggests that proliferation is coupled to active reinforcement of genome integrity during cellular state transitions. Similar coordination between proliferative programs and genome maintenance has been described in activated immune cells and regenerating tissues, where extensive transcriptional remodeling requires simultaneous protection of genome stability (Ramirez-Carrozzi et al., 2006; Keren-Shaul et al., 2017; Jackson and Bartek, 2009; Ciccia and Elledge, 2010; Prinz et al., 2019; Masuda et al., 2020). Collectively, these observations indicate that RTL6 deficiency induces a fundamental reorganization of microglial physiology that extends far beyond attenuation of inflammatory signaling, encompassing coordinated metabolic adaptation, tissue remodeling, and genome maintenance. Taken together, these coordinated transcriptional programs support the idea that RTL6 functions as a key regulator of microglial state transitions, coupling inflammatory responsiveness with maintenance of neuroimmune homeostasis.

Our findings suggest that RTL6 deficiency promotes a DAM-like transcriptional state in mice (Keren-Shaul et al., 2017; Masuda *et al*., 2020), although this state remains distinct from canonical DAM states described in mouse and human studies, particularly in its retention of certain homeostatic features and lack of full induction of key DAM markers (Figure 2A and 2B). These observations raise the possibility that *RTL6* contributes to eutherian-specific tuning of microglial state regulation, a hypothesis that will require validation in human microglial systems.

In mammals, microglia are established as a largely self-maintaining lineage, with minimal dependence on peripheral monocyte input, sustained by local proliferation within a CNS environment tightly constrained by the blood–brain barrier (Ginhoux et al., 2010; Shimizu and Prinz, 2025). Thus, mammalian microglia represent a system in which conserved microglial properties are more tightly integrated, reflecting adaptation to a relatively insulated CNS niche. In this context, RTL6 appears to contribute to the integration of inflammatory responsiveness, microglial state transitions, tissue adaptation, proliferative adaptation, and genome maintenance within mammalian microglia. These findings identify *RTL6* as a previously unrecognized component of the regulatory network that shapes microglial functional states in placental mammals.

Collectively, our results suggest that *RTL6* has evolved beyond its extracellular LPS-binding activity to function as an integrative regulator of microglial state transitions. By coordinating early inflammatory signaling with antiviral defense, tissue adaptation, proliferative adaptation, and genome maintenance, *RTL6* appears to couple innate immune activation to the establishment of distinct microglial functional states. As a eutherian-specific metavirus-derived gene, *RTL6* therefore represents a previously unrecognized evolutionary innovation that contributes to the diversification of microglial regulatory networks in placental mammals.

## Materials and Methods

### 4.1. Animals

All of the animal experiments were reviewed and approved by the Institutional Animal Care and Use Committee of Tokai University (Isehara-234011) and Institute of Science Tokyo (IST) (A2022-057A) and were performed in accordance with the Guideline for the Care and Use of Laboratory Animals of Tokai University. Mice were maintained under specific pathogen-free conditions on a 12:12 h light–dark cycle and were provided free access to a standard chow diet and water.

### 4.2 Microglia isolation and primary culture

Primary mixed glial cultures were prepared according to the protocol of Lian et al. with modifications (Lian *et al*., 2016). Briefly, P0–P2 brains were homogenized in phosphate-buffered saline (PBS) supplemented with trypsin and DNase (final concentration: 0.05% and 0.1 mg/mL, respectively) by repeated pipetting 10 times and incubated in a 37◦C water bath for 5 min, and this process was repeated 3 times. After the addition of Dulbecco’s modified Eagle medium (DMEM) (Thermo Fisher Science, GibcoTM Catalog No. 11995065, Waltham, MA, USA) along with 10% fetal bovine serum (FBS) (Thermo Fisher Science, Catalog No.10437028), the debris was removed through a cell strainer (EASYstrainer, mesh size: 100 µm, Greiner, Cat. No. 542000, Kremsmunster, Austria). After centrifugation (400×g, at 4◦C) for 5 min, the supernatants were aspirated and the pellet was resuspended with 5 mL of warmed culture medium. After determining the cell density, the mixed glial cells were plated into 35 mm glass-base dishes (IWAKI, Code No. 3910-035, Osaka, Japan) or 25 cm2 flasks (Canted Neck, Corning, Cat. No. 430639, New York, NY, USA) and incubated in a CO2 incubator with 5% CO2 and 100% humidity at 37◦C. Media were changed the next day and every 3–4 days thereafter. Ten minutes after administration of LPS (final concentration 10 ng/ml), microglia cells were collected by tapping the culture flasks vigorously for 2 min and the suspension were centrifugated (400×g, at 4◦C) for 5 min.

#### RNAseq

For RNA isolation and quality control, the integrity and quantity of the total RNA was measured using an Agilent Technologies 2200 TapeStation (Agilent Technologies Inc., Santa Clara, CA). For library preparation and sequencing, total RNA obtained from each sample was subjected to a sequencing library construction using the TruSeq Stranded mRNA Library Prep Kit (Illumina Inc., San Diego, CA) according to the manufacturer’s protocols. Sample libraries were sequenced using NovaSeq 6000 (Illumina Inc., San Diego, CA) in 100 base pair (bp) paired-end reads. For data analysis, raw sequencing reads were processed using the nf-core/rnaseq pipeline (v3.2.0; Patel *et al.,* 2022). Reads were aligned to the mouse reference genome (GRCm39, Ensembl release 113) using STAR, and gene-level counts were quantified with RSEM. Expected counts were imported into R (v4.5.1) and analyzed with DESeq2 (v1.48.2; Love *et al*., 2014). Differentially expressed genes (3,540 genes) were initially classified into 25 functional categories (Groups A–Y; Tables S2 and S3) and subsequently reorganized into five integrated functional modules (Modules 1-5; Table 1A and 1B) based on biological function. Functional annotation, iterative organization of gene categories, and refinement of the classification scheme were performed by the authors with assistance from ChatGPT (OpenAI, GPT-5.5).

## Supporting information

Supplementary Results, Supplementary Table S1-3 and Supplementary Figs 1 and 2

## Ethics

All of the animal experiments were reviewed and approved by the Institutional Animal Care and Use Committee of Tokai University (Isehara-234011) and Institute of Science Tokyo (IST) (A2022-057A) and were performed in accordance with the Guideline for the Care and Use of Laboratory Animals of Tokai University. This work did not require ethical approval from a human subject.

## Data accessibility

We have deposited the results of RNAseq data on WT and *Rtl6* KO microglia with and without LPS. Accession to cite for these SRA data: PRJNA1472295

Electronic supplementary material is available online.

## Declaration of AI use

We have used AI-assisted technologies in English editing this article.

## Author contributions

Conceptualization: F.I., T.K.-I.; Methodology: F.I., M.I., H.S., T.K.; Validation: F.I., M.I, T.K.-I.; Formal analysis: F.I., M.I, H.S., T.K., T.K.-I; Investigation: M.I, F.I., T.K.-I.; Data curation: M.I, F.I., T.K.-I.; Writing - original draft: F.I., T.K.-I.; Writing - review & editing: F.I., I.M., H. S., T.K.-I.; Visualization: F.I., M.I.; Supervision: F.I., T.K.-I. All authors gave final approval for publication and agreed to be held accountable for the work performed therein.

## Acknowledgements

The authors would like to thank Ms Naoko Takayasu (Tokai university) for technical assistance in preparing mRNA from brains and performing RT-PCR experiments, Mr Akinori Takase and Ms Keiko Yokoyama, Life Science Support center, Tokai University, for analysis of RNAseq data.

