## Supplementary Results, Supplementary Table S1-3 and Supplementary Figs 1 and 2 for "A eutherian-specific metaviral gene, *RTL6*, coordinates microglial inflammatory responsiveness and state regulation"

**Supplementary Result 1**

**Coordinated activation of cell-cycle and genome maintenance programs**

Module 5 revealed a highly coordinated transcriptional program integrating metabolic remodeling with cell-cycle progression, DNA replication, and genome maintenance (Table 1B).

**Module 5-1 Metabolism shift**

Rather than affecting a single metabolic pathway, these transcriptional changes involve multiple interconnected processes that support cellular activation and adaptation. Within M5-1, genes involved in oxidative phosphorylation (M5-1A) comprising Complexes I–V, such as *Ndufa4, Ndufa9, Ndufa12, Ndufb6, Ndufb7, Ndufb8, Ndufs1, Ndufs4, Ndufs6, Ndufs8, Ndufv2, mt-Nd1* and *mt-Nd3* (Complex I); *Sdhb* (Complex II); *Uqcrb, Uqcrc1, Uqcrc2* and *Uqcrq* (Complex III); *Cox5b, Cox6a1, Cox7a2,* and *Cox7b* (Complex IV); and *Atp5f1a, Atp5f1b, Atp5mc1, Atp5mc3, Atp5mf, Atp5pb,* and *Atp5po* (Complex V). Genes involved in the tricarboxylic acid cycle, represented by *Cs, Sdhb, Suclg1, Mdh2,* and *Dld,* ATP production and mitochondrial energy metabolism, comprising *Ak2, Ampd2, Adss2, Gmpr, Pnp, Oxct1, Pdk1,* and *Gpd2,* Mitochondrial intermediary metabolism, such as *Me2, Got1, Etfa, Gpd1l, Oxct1,* and *Pdk1,* were similarly induced. Mitochondrial bioenergetic and biogenesis pathways, including *Tomm5, Tomm22, Tomm70a, Timm13, and Pmpca* (protein import); *Tufm, Gfm1, Fastkd2, Fastkd3,* and *Trmt10c* (Mitochondrial translation); *Immt, Chchd1, Chchd3,* and *Samm50* (Cristae organization/mitochondrial architecture); *Atad3a, Mtch2, Opa3, Pgam5,* and *Sfxn1* (Mitochondrial membrane organization) and *Slc25a5, Slc25a44, Vdac1, Vdac3, Tspo* (Metabolite transport), were also coordinately upregulated, indicating enhanced mitochondrial activity and bioenergetic capacity. Although a small number of mitochondrial metabolic genes, including *Aldh2, Aldh6a1, Cox7a2l, Ndufa6, Glul,* and *Mthfr*, were modestly downregulated, they represented only a small minority relative to the coordinated induction of genes involved in oxidative phosphorylation, the TCA cycle, mitochondrial biogenesis, and ATP production, supporting an overall increase in mitochondrial bioenergetic capacity. Glycolytic and pentose phosphate pathway enzymes (M5-1B), including *Gpi, Pfkp, Pfkl,* and *Pgm1* (Glycolysis); *Pdg, Tkt*, and *Taldo1* (Pentose phosphate pathway); *Ugp2* and *Isyna1* (Glucose utilization/carbohydrate metabolism) and *Gls* (Glycolysis-linked anaplerotic metabolism), were coordinately induced, suggesting coordinated reinforcement of both energy production and anabolic metabolism. In parallel, lipid metabolic pathways were extensively remodeled. Genes involved in fatty acid activation, phospholipid synthesis, cholesterol biosynthesis, sphingolipid metabolism, and membrane glycosylation (M5-1C), including *Acsl4, Acot9, Agpat5, Acaca, Acat1, Agps, Far1, Fabp4, Fabp5*, and *Tmem97*, were predominantly increased, whereas genes associated with lipid storage, cholesterol efflux, and homeostatic lipid metabolism, including *Apoe, Abca1, Abcg1, Lpin1, Lpin2,* and *Rxra* (M5-1C), were reduced. These changes indicate a shift from lipid storage and homeostasis toward membrane biosynthesis and anabolic metabolism required for proliferative activation.

**Module 5-2 Growth/survival signalinga**

Module 5-2 revealed coordinated activation of intracellular signaling pathways that support cellular activation, cytoskeletal remodeling, and membrane dynamics. Genes involved in growth factor signaling (M5-2A1), including Errfi1 and Btbd10, were upregulated. In contrast, expression of *Igf1, Igf1r, Igf2* (M5-2A1), together with the RasGAP family members, *Rasa1, Rasa4, and Rasal2,* (M5-2A3) and *Foxo3* (M5-2A4) was reduced, accompanied by downregulation of the stress-responsive genes *Txnip, Sesn1* (M5-3B), *Ndrg1,* and *Ndrg3* (M5-3D), indicating coordinated suppression of the IGF–PI3K/AKT–FOXO signaling axis and its associated stress-adaptation program (Webb and Brunet, 2014; Manning and Toker, 2017). Components of MAPK-, PI3K-, small GTPase-, and calcium-dependent signaling pathways, including *Map2k1, Mapk6, Mapkapk2, Ptk2b, Camk1, Camk2g* (M5-2A2), *Spred1, Ptpn6, Ptpn7, Ptpn12* (M5-2A3), *Trim27*, and *Tnfaip1* (M5-2A3), were broadly induced together with multiple phosphatases and signaling modulators. In contrast, *Map3k8* (*Tpl2*), the only canonical upstream MAPKKK, was significantly downregulated among the MAPK signaling molecules. It preferentially attenuates the TLR4–TPL2 (MAP3K8)–ERK–AP-1 signaling axis, while other MAPK branches, including p38-associated pathways, remain relatively preserved or show compensatory regulation. This interpretation is supported by the concomitant downregulation of immediate-early response genes (*Jun, Junb, Jund, Fos, Fosb, Egr2, Egr3, Nr4a1, Nr4a2* and *Dusp1*) (M1-1) despite maintained or increased expression of several downstream or non-canonical MAPK components (e.g., *Map2k1, Mapk6,* and *Mapkapk2*) (Dumitru et al., 2000; Ganke et al., 2012; Auther and Ley, 2013). Genes in Integrin/Cytosckelton remodeling (M5-2B), including integrin receptors, such as *Itgam, Itgb1, Itgb2, Itgb3, Itgb5* and *Itgav,* integrin-associated adaptors, represented by *Lims1, Tln1, Fermt3, Lpxn, Parvg,* and *Lasp1*, actin polymerization machinery, involving *Actb, Actn1, Ezr, Cfl1, Cap1, Capza1, Arpc1b, Arpc3, Arpc4, Arpc5, Actr2, Actr3, Pfn1, Diaph1, Diaph3, Fmn1,*

*Wdr1, Lcp1, Msn*, and *Crybg3,* and Rho-family GTPase regulators, such as *Rhoa ,Cdc42, Arf6, Vav1, Dock10, Rasgrp3, Rasa2, Arhgdia, Arhgdib, Arhgap11a, Arhgap25,* and *Arhgap30,* were coordinately upregulated. Collectively, M5-2B represents a coordinated signaling module linking cell adhesion to actin cytoskeletal remodeling. The concerted induction of integrin-mediated actin cytoskeletal remodeling genes suggests an enhanced capacity for cell adhesion, migration, morphological remodeling, and phagocytosis in macrophages and microglia.

Membrane trafficking pathways were also extensively activated, including Rab- and Arf-family GTPases (*Rab1a, Rab5c, Rab7, Rab7b, Arf1* and *Arf6*), sorting nexins (*Snx1, Snx2, Snx10* and *Snx24*), COPII-mediated ER-to-Golgi transport components (*Sec11c, Sec13, Sec23b* and *Sec24d*), the retromer complex (*Vps26a, Vps29* and *Vps35*), the ESCRT-associated factor *Vps36*, the vesicle SNARE *Ykt6*, and additional vesicle tethering and endomembrane transport machinery (M5-2C), indicating coordinated activation of the membrane trafficking machinery required for vesicle transport, endosomal recycling, and membrane remodeling during microglial activation (Pfeffer, 1994).

**Module 5-3 Autophagy/stress adaptation**

The M5-3A module revealed coordinated activation of the ER proteostasis network, including UPR-associated transcription factors (*Atf4, Atf6* and *Xbp1*), molecular chaperones (*Hspa4, Hspa5, Hspa9, Hspa13, Hspa14,* Dnaja/Dnajc family members, *Bag1* and *Stip1*), protein disulfide isomerases and oxidative folding enzymes (*P4hb, Erp44, Txndc5, Txndc9, Txndc11, Ero1a* and *Ero1b*), and ER quality-control factors (*Canx, Uggt1, Edem3, Mlec* and *Selenos*), indicating coordinated reinforcement of ER proteostasis pathways (Ron and Walter, 2007; Hetz, 2012). Redox defense systems (M5-3B) and ubiquitin–proteasome pathways (M5-3C) were similarly activated, extending this proteostatic program beyond the ER and reinforcing protein quality control and oxidative stress resistance.

In parallel, genes involved in lysosomal biogenesis, lysosomal function, and autophagy (M5-3D)**,** lysosomal biogenesis and function (*Hexb, Manba, Gba1, Lgmn, Ctss, Ifi30, Hgsnat*, and *M6pr*), endolysosomal membrane trafficking (*Vac14, Tollip, Abca7, C9orf72, Cln5,* and *Vma21*), and ER-associated membrane remodeling (*Retreg1* and *Tmem41b*) were coordinately induced. In contrast, multiple regulators of canonical and stress-induced autophagy, including *Ulk1, Ulk2, Atg2a, Map1lc3b, Pink1, Bnip3, Rubcn, Rubcnl, Trp53inp1, Ndrg1,* and *Ndrg3*, were downregulated. Thus, lysosomal remodeling and proteostatic adaptation appear to be enhanced independently of canonical autophagy induction.

Collectively, the coordinated M5-1–M5-3 responses indicate that RTL6 deficiency induces extensive remodeling of cellular metabolism, intracellular signaling, membrane trafficking, and proteostatic adaptation, thereby establishing a cellular state that supports the coordinated activation of the DNA replication, chromatin duplication, and genome maintenance programs described in Module 5-4.

**Module 5-4 Cell cycle/chromatin/genome stability**

For subsequent analyses, Module 5-4 was subdivided into nine functional categories representing sequential stages of genome duplication and maintenance: (M5-4A1) cell cycle regulators, (M5-4A2) DNA replication licensing, (M5-4A3) DNA replication machinery, (M5-4A4) DNA precursor synthesis, (M5-4A5) chromosome segregation, (M5-4A6) histone/nucleosome assembly, (M5-4B) DNA repair, (M5-4C) genome stability, and (M5-4D) chromatin remodeling/epigenetic regulation.

M5-4 comprised an integrated transcriptional program coordinating cell-cycle progression, DNA replication, genome maintenance, and chromatin remodeling. Within this module, M5-4A1 showed broad induction of genes regulating the cell cycle, including those controlling the G1/S transition, G2/M transition, and general cell-cycle progression. Genes regulating the G1/S transition were coordinately induced, including cyclins (*Ccnd1, Ccne1,* and *Ccne2*), cyclin-dependent kinases (*Cdk2, Cdk4,* and *Cdk6*), the CDK-activating phosphatases *Cdc25b* and *Cdc25c,* the SCF-associated regulators *Skp2* and *Cks1b*, and multiple E2F transcription factors (*E2f1, E2f2, E2f6, E2f7,* and *E2f8*). Notably, the complete E2F transcriptional network was coordinately activated, with all major E2F family members induced by more than twofold, including the activator E2Fs (*E2f1* and *E2f2*) together with the atypical repressor E2Fs (*E2f6, E2f7,* and *E2f8*), indicating activation of the complete E2F transcriptional program coordinating S-phase entry and DNA replication. In contrast, the cyclin-dependent kinase inhibitor *Cdkn1a* (p21) was downregulated, consistent with release of G1 checkpoint restraint and promotion of S-phase entry. The concomitant reduction of *Ccnd2* suggests selective cyclin remodeling rather than generalized induction of all D-type cyclins, consistent with preferential utilization of the Ccnd1-driven proliferative program. Core regulators governing G2/M transition (*Cdk1*–Cyclin B, *Wee1, Pkmyt1, Mastl*), mitotic entry (*Plk1, Aurka*), spindle assembly checkpoint (*Bub1, Bub1b, Mad2l1, Zwilch, Knl1*), chromosome segregation (*Aurkb–Incenp–Birc5*), and mitotic exit (*Cdc20*) were coordinately upregulated, indicating activation of the complete mitotic progression program. Key regulators of G2/M transcription (*Foxm1, Mybl2, Lin54*), APC/C-mediated ubiquitin-dependent cell-cycle progression (*Anapc5, Cdc27*), and multiple checkpoint-associated regulators (*Cdca3, Cdkn3, G2e3, Pin1*) were coordinately induced, indicating coordinated activation of the regulatory circuitry governing mitotic progression.

Genes involved in DNA replication initiation (M5-4A2), including members of the minichromosome maintenance (MCM) and origin recognition (ORC) complexes together with additional licensing factors (*Cdc6, Cdc7* and *Cdt1*), were coordinately upregulated. Genes required for replication fork progression (M5-4A3), including components of the replication fork C (RFC) complex (*Rfc1-Rfc5*) and go-ichi-ni-san (GINS) complexes (*Gins1* and *Gins2*), DNA replication machinery, including primase (*Pola1, Pola2, Prim1, Prim2*) and replicative polymerase (*Pold1, Pold2, Pole, Pole2, Pole3* and *Pole4*), Replication fork (*Pcna, Fen1,* and *Lig1*), were likewise increased. Genes required for de novo and salvage synthesis of DNA precursors（M5-4A4）, including ribonucleotide reduction (*Rrm1, Rrm2*), thymidylate synthesis (*Tyms, Dhfr, Dtymk*), thymidine salvage (*Tk1*), and dUTP quality control (*Dut*), were coordinately upregulated, indicating enhanced nucleotide biosynthesis to support DNA replication during cell proliferation. Numerous genes involved in chromosome segregation (M5-4A5), including members of the centromere protein (CENP), kinesin-super family protein (KIF), HAUS complex components, and structural maintenance of chromosome (SMC) families together with centrosome- (*Cep83, Cep128, Cep152*) and spindle-associated proteins (*Ckap2, CKap2l, Ckap5*), chromosome cohesion and chromosome passenger complex components (*Cdca2, Cdca5, Cdca8*) were also induced. Thus, most centrosome and mitotic spindle components were coordinately upregulated, consistent with activation of the centrosome biogenesis program, whereas *Ankle2*, which primarily functions in nuclear envelope reassembly during mitotic exit rather than centrosome assembly, was the only gene modestly downregulated within this module. Extensive upregulation of canonical histone genes representing the H1, H2A, H2B, H3, and H4 families (M5-4A6) was observed. This was accompanied by increased expression of nucleosome assembly factors, including *Nap1l1, Nap1l4, Ssrp1*and *Rbb4* (M5-4D). Genes involved in DNA repair (M5-5B) were similarly upregulated. Genes involved in DNA damage sensing (M5-4B1), including *Parp1, Parpbp, Atm, Chek1* and *Chek2*, replication fork protection (M5-4B2), including *Hrob, Tonsl, Mms22l* and *Firrm,* were upregulated. Homologous recombination (HR) genes (M5-4B3), including Rad51 family members, were markedly upregulated. Fanconi amenia complex (FANC), BRCA-associated repair factors, and repair networks, including *Msh3, Ercc6, Ercc6l,*

*Tdg, Commd4, Polh* and *Polq*, were also markedly upregulated, consistent with replication-associated stress (Zeman and Cimprich, 2014, Hammond *et al.*, 2019).

Genes involved in genome maintenance (M5-4C) were also coordinately regulated. Telomere protection (*Tinf2*) together with genome integrity factors (*Pin1***,** *Erh***,** and *Ciao2a*) were upregulated, whereas *Nipbl***,** a cohesin-loading factor involved in higher-order chromosome organization, was modestly reduced. These findings suggest reinforcement of genome surveillance during DNA replication.

Extensive remodeling of chromatin regulatory pathways accompanied the DNA replication program (M5-4D). ATP-dependent chromatin remodelers (M5-4D1) involved in nucleosome remodeling and DNA replication, including SWI/SNF (*Smarcb1* and *Smarcc1*), ISWI (*Scarca5*), and INO80 complexes, together with *Baz1a,* and *Baz1b*, were coordinately upregulated, whereas CHD family genes (*Chd3, Chd6,* and *Chd9*), which are more closely associated with chromatin organization and transcriptional regulation, were modestly downregulated. These findings suggest preferential activation of chromatin-remodeling machinery that supports DNA replication and cell-cycle progression rather than a uniform induction of all ATP-dependent chromatin remodelers.

Chromatin regulators involved in chromatin organization, nucleosome dynamics, and maintenance of proliferative chromatin states (M5-4D2)—including *Hdac1*, together with the transcriptional corepressor *TRIM28* (*KAP1*), PRC2 components (*Suz12* and *Eed*), Chromatin corepressors, such as MBD/NuRD-associated proteins (*Mbd2, Mbd3, Gatad2A* and *Morf4l2*), COMPASS components (*Wdr5* and *Dpy30*), and HMGB/HMGN chromatin architectural proteins (*Hmgb1-3* and *Hmgn1-2*)—were coordinately upregulated. In contrast, several epigenetic modifiers associated primarily with developmental gene regulation or differentiation, including *Ezh1, Bcor, Kdm5a, Kdm6a, Kdm7a,* and *Ogt,* were downregulated, suggesting selective reinforcement of chromatin programs supporting proliferation rather than global activation of all epigenetic regulators. Replication-coupled histone deposition factors) (M5-4D3) **(***Chaf1a, Chaf1b, Asf1b, Hat1, Hirip3, Nap1l1, Nap1l4, Ssrp1, Rbbp4,* and *Slbp***)** were also strongly induced, consistent with extensive nucleosome assembly during DNA replication. In parallel, chromatin organization factors (M5-4D4) (*Hells* and *Atad2*) together with nuclear architecture components (M5-4D5) (*Lmnb1, Lmnb2,* and *Tmpo*) were upregulated, suggesting coordinated reconstruction of higher-order chromatin structure accompanying genome duplication.

Together, these coordinated transcriptional changes define an integrated genome duplication program encompassing cell-cycle progression, DNA replication, chromosome segregation, DNA repair, genome surveillance, and replication-coupled chromatin assembly. Rather than reflecting isolated pathway-specific changes, RTL6 deficiency induces coordinated activation of the molecular machinery required for faithful genome duplication during cell proliferation.

**Legends for Supplementary materials**

**Supplementary Table S1 RNA-seq data from WT and KO following 10 min of LPS stimulation.** Data of all the 3,540 significantly differentially expressed genes (padj < 0.05) identified between WT3-4, and KO 3-4 are listed together with their logFC values, adjusted *P* values (padj), and gene descriptions (see also Figure 1B).

**Supplementary Table S2 Gene classification of downregulated genes.**
Differentially expressed genes were classified nine biological process categories (A-I) and 25 Cellular machinery/homeostatic cellular system categories (J-Y). The numbers genes assigned to each category is indicated.

**Supplementary Table S3 Gene classification of upregulated genes.**
Differentially expressed genes were classified as described for Supplementary Table S2.

**Supplementary Figure S1 Heatmap of WT and *Rtl6* KO microglia with and without LPS stimulation.** Among the 4,734 significantly differentially expressed genes (padj < 0.05), 2,674 were upregulated and 2,055 were downregulated. KO1 (without LPS) exhibited a distinct transcriptome profile characterized by a weaker upregulation response while maintaining a downregulation pattern similar to that of other KO samples.

**Supplementary Figure S2 Differential expression of *Dusp* family genes in *Rtl6* KO.** *Dusp1, Dusp2, Dusp7,* and *Dusp15* were downregulated, consistent with impaired AP-1/immediate-early gene (IEG)-dependent induction, whereas *Dusp3, Dusp4, Dusp6,* and *Dusp18* were upregulated, consistent with preserved basal ERK signaling and compensatory negative-feedback regulation (Caunt et al., 2008; Ham et al., 2015; Lake et al., 2016).

#### Supplementary Table S1

| id | gene_name | logFC | baseMean | lfcSE | stat | PValue | padj.orig | padj | KO 3 | KO 4 | WT 3 | WT 4 | entrez_id | gene_description |
|---|---|---|---|---|---|---|---|---|---|---|---|---|---|---|
| ENSMUSG00000119892 | Gm23971 | -9.31442 | 59.66765 | 1.545717 | 6.025953 | 1.68E-09 | 3.65E-08 | 4.42E-08 | 0 | 0 | 156.3716 | 82.29899 | 115487918 | predicted gene, 23971 [Source:MGI Symbol;Acc:MGI:5453748] |
| ENSMUSG00000121339 | | -7.36727 | 15.48839 | 1.724115 | 4.273072 | 1.93E-05 | 0.000193 | 0.000234 | 0 | 0 | 31.09143 | 30.86212 | 100041085 | major facilitator superfamily domain containing 4B3, pseudogene [Source:NCBI gene (formerly Entrezgene);Acc:100041085] |
| ENSMUSG00000039349 | C130074G19Rik | -7.28029 | 14.60246 | 1.778046 | 4.094548 | 4.23E-05 | 0.000394 | 0.000478 | 0 | 0 | 18.28908 | 40.12076 | 226777 | RIKEN cDNA C130074G19 gene [Source:MGI Symbol;Acc:MGI:2444831] |
| ENSMUSG00000050121 | Opalin | -6.86755 | 10.97326 | 1.898235 | 3.61786 | 0.000297 | 0.002186 | 0.002649 | 0 | 0 | 10.97345 | 32.9196 | 226115 | oligodendrocytic myelin paranodal and inner loop protein [Source:MGI Symbol;Acc:MGI:2657025] |
| ENSMUSG00000113073 | Gm48261 | -6.64336 | 9.373059 | 1.895 | 3.505731 | 0.000455 | 0.00319 | 0.003864 | 0 | 0 | 21.03244 | 16.4598 | | predicted gene, 48261 [Source:MGI Symbol;Acc:MGI:6097679] |
| ENSMUSG00000004359 | Spic | -6.62014 | 9.230158 | 1.898285 | 3.48743 | 0.000488 | 0.003391 | 0.004107 | 0 | 0 | 17.37462 | 19.54601 | 20728 | Spi-C transcription factor (Spi-1/PU.1 related) [Source:MGI Symbol;Acc:MGI:1341168] |
| ENSMUSG00000095526 | Gm10243 | -6.56664 | 8.887261 | 1.917748 | 3.424139 | 0.000617 | 0.004151 | 0.005028 | 0 | 0 | 20.11798 | 15.43106 | | predicted gene 10243 [Source:MGI Symbol;Acc:MGI:3704266] |
| ENSMUSG00000025746 | Il6 | -6.19083 | 6.858311 | 2.044866 | 3.027497 | 0.002466 | | 0.016082 | 0 | 0 | 10.97345 | 16.4598 | 16193 | interleukin 6 [Source:MGI Symbol;Acc:MGI:96559] |
| ENSMUSG00000058626 | Capn11 | -6.18264 | 6.829694 | 2.192347 | 2.820103 | 0.004801 | | 0.028074 | 0 | 0 | 3.657815 | 23.66096 | 268958 | calpain 11 [Source:MGI Symbol;Acc:MGI:1352490] |
| ENSMUSG00000026418 | Tnni1 | -6.10898 | 6.486797 | 2.130083 | 2.867953 | 0.004131 | | 0.024807 | 0 | 0 | 6.401177 | 19.54601 | 21952 | troponin I, skeletal, slow 1 [Source:MGI Symbol;Acc:MGI:105073] |
| ENSMUSG00000053693 | Mast1 | -6.10046 | 6.429701 | 2.088654 | 2.920759 | 0.003492 | | 0.021528 | 0 | 0 | 16.46017 | 9.258636 | 56527 | microtubule associated serine/threonine kinase 1 [Source:MGI Symbol;Acc:MGI:1861901] |
| ENSMUSG00000103255 | Pcdhac1 | -5.98726 | 5.943903 | 2.136491 | 2.80238 | 0.005073 | | 0.029439 | 0 | 0 | 15.54571 | 8.229899 | 353236 | protocadherin alpha subfamily C, 1 [Source:MGI Symbol;Acc:MGI:1891442] |
| ENSMUSG00000024650 | Slc22a6 | -5.9834 | 5.943857 | 2.149161 | 2.784063 | 0.005368 | | 0.030782 | 0 | 0 | 7.315631 | 16.4598 | 18399 | solute carrier family 22 (organic anion transporter), member 6 [Source:MGI Symbol;Acc:MGI:892001] |
| ENSMUSG00000115471 | C230086J09Rik | -5.95611 | 16.15286 | 1.690949 | 3.522349 | 0.000428 | 0.003027 | 0.003666 | 1.057752 | 0 | 26.51916 | 37.03454 | | RIKEN cDNA C230086J09 gene [Source:MGI Symbol;Acc:MGI:2443423] |
| ENSMUSG00000086825 | Gm15675 | -5.82523 | 29.40106 | 1.213533 | 4.800225 | 1.58E-06 | 2E-05 | 2.43E-05 | 1.057752 | 0.983552 | 63.09731 | 52.4656 | | predicted gene 15675 [Source:MGI Symbol;Acc:MGI:3783117] |
| ENSMUSG00000076439 | Mog | -5.63827 | 77.80086 | 0.82665 | 6.820624 | 9.06E-12 | 2.71E-10 | 3.28E-10 | 3.173256 | 2.950655 | 77.72857 | 227.351 | 17441 | myelin oligodendrocyte glycoprotein [Source:MGI Symbol;Acc:MGI:97435] |
| ENSMUSG00000055333 | Fat2 | -5.56324 | 12.35229 | 1.784793 | 3.117023 | 0.001827 | 0.010367 | 0.012558 | 1.057752 | 0 | 32.92034 | 15.43106 | 245827 | FAT atypical cadherin 2 [Source:MGI Symbol;Acc:MGI:2685369] |
| ENSMUSG00000041670 | Rims1 | -5.35109 | 10.59059 | 1.862184 | 2.873553 | 0.004059 | 0.020225 | 0.024459 | 0 | 0.983552 | 31.09143 | 10.28737 | 116837 | regulating synaptic membrane exocytosis 1 [Source:MGI Symbol;Acc:MGI:2152971] |
| ENSMUSG00000037014 | Sstr4 | -5.31366 | 10.36188 | 1.830935 | 2.902156 | 0.003706 | 0.018707 | 0.022631 | 0 | 0.983552 | 13.71681 | 26.74717 | 20608 | somatostatin receptor 4 [Source:MGI Symbol;Acc:MGI:105372] |
| ENSMUSG00000142392 | Gm68273 | -5.3087 | 21.01833 | 1.598771 | 3.320487 | 0.000899 | 0.005737 | 0.00695 | 2.115504 | 0 | 76.81412 | 5.143687 | | predicted gene, 68273 [Source:MGI Symbol;Acc:MGI:7819375] |
| ENSMUSG00000000303 | Cdh1 | -5.2226 | 80.75788 | 1.989639 | 2.624898 | 0.008668 | 0.037918 | 0.045801 | 8.462015 | 0 | 199.3509 | 115.2186 | 12550 | cadherin 1 [Source:MGI Symbol;Acc:MGI:88354] |
| ENSMUSG00000020436 | Gabrg2 | -5.2185 | 9.704659 | 1.827696 | 2.855233 | 0.004301 | 0.021185 | 0.025613 | 0 | 0.983552 | 18.28908 | 19.54601 | 14406 | gamma-aminobutyric acid type A receptor, subunit gamma 2 [Source:MGI Symbol;Acc:MGI:95623] |
| ENSMUSG00000059742 | Kcnh7 | -5.12994 | 9.133149 | 1.852435 | 2.769295 | 0.005618 | 0.02643 | 0.031927 | 0 | 0.983552 | 20.11798 | 15.43106 | 170738 | potassium voltage-gated channel, subfamily H (eag-related), member 7 [Source:MGI Symbol;Acc:MGI:2159566] |
| ENSMUSG00000032517 | Mobp | -4.88081 | 370.1106 | 0.805815 | 6.05698 | 1.39E-09 | 3.06E-08 | 3.71E-08 | 19.03953 | 29.50655 | 371.2683 | 1060.628 | 17433 | myelin-associated oligodendrocytic basic protein [Source:MGI Symbol;Acc:MGI:108511] |
| ENSMUSG00000016529 | Il10 | -4.85223 | 481.3957 | 0.867361 | 5.594251 | 2.22E-08 | 4.11E-07 | 4.98E-07 | 20.09729 | 44.25983 | 500.2062 | 1361.02 | 16153 | interleukin 10 [Source:MGI Symbol;Acc:MGI:96537] |
| ENSMUSG00000051906 | Cd209f | -4.81321 | 103.9894 | 0.547905 | 8.784753 | 1.57E-18 | 9.5E-17 | 1.15E-16 | 7.404263 | 6.884862 | 160.9439 | 240.7245 | 69142 | CD209f antigen [Source:MGI Symbol;Acc:MGI:1916392] |
| ENSMUSG00000027875 | Hmgcs2 | -4.78251 | 14.54132 | 1.362974 | 3.508875 | 0.00045 | 0.00316 | 0.003828 | 1.057752 | 0.983552 | 28.34807 | 27.77591 | 15360 | 3-hydroxy-3-methylglutaryl-Coenzyme A synthase 2 [Source:MGI Symbol;Acc:MGI:101939] |
| ENSMUSG00000087684 | 1200007C13Rik | -4.69903 | 13.93117 | 1.42016 | 3.308803 | 0.000937 | 0.005932 | 0.007186 | 2.115504 | 0 | 23.7758 | 29.83338 | | RIKEN cDNA 1200007C13 gene [Source:MGI Symbol;Acc:MGI:1921369] |
| ENSMUSG00000079168 | Cd209g | -4.69597 | 20.85392 | 1.179249 | 3.982175 | 6.83E-05 | 0.000596 | 0.000722 | 3.173256 | 0 | 41.15042 | 39.09202 | 70192 | CD209g antigen [Source:MGI Symbol;Acc:MGI:1917442] |
| ENSMUSG00000045087 | S1pr5 | -4.51902 | 12.03655 | 1.586154 | 2.849043 | 0.004385 | 0.021504 | 0.025999 | 0 | 1.967103 | 9.144538 | 37.03454 | 94226 | sphingosine-1-phosphate receptor 5 [Source:MGI Symbol;Acc:MGI:2150641] |
| ENSMUSG00000124378 | Gm67753 | -4.48659 | 29.62421 | 0.97555 | 4.599039 | 4.24E-06 | 4.89E-05 | 5.93E-05 | 1.057752 | 3.934207 | 38.40706 | 75.09783 | | predicted gene, 67753 [Source:MGI Symbol;Acc:MGI:7818337] |
| ENSMUSG00000129763 | 6720473M11Rik | -4.44435 | 17.33053 | 1.184698 | 3.75146 | 0.000176 | 0.001372 | 0.001662 | 1.057752 | 1.967103 | 37.49261 | 28.80465 | 115488623 | RIKEN cDNA 6720473M11 gene [Source:MGI Symbol;Acc:MGI:1925150] |
| ENSMUSG00000098051 | Gm27032 | -4.38434 | 27.46252 | 1.013542 | 4.325761 | 1.52E-05 | 0.000157 | 0.00019 | 0 | 4.917758 | 49.38051 | 55.55182 | | predicted gene, 27032 [Source:MGI Symbol;Acc:MGI:5504147] |
| ENSMUSG00000032289 | Thsd4 | -4.35036 | 245.8646 | 0.33068 | 13.1558 | 1.58E-39 | 7.71E-37 | 9.34E-37 | 25.38605 | 20.65459 | 417.9054 | 519.5124 | 207596 | thrombospondin, type I, domain containing 4 [Source:MGI Symbol;Acc:MGI:2672033] |
| ENSMUSG00000041361 | Myzap | -4.2953 | 16.05304 | 1.29481 | 3.317319 | 0.000909 | 0.005785 | 0.007008 | 3.173256 | 0 | 21.94689 | 39.09202 | 102371 | myocardial zonula adherens protein [Source:MGI Symbol;Acc:MGI:2142908] |
| ENSMUSG00000066755 | Tnfsf18 | -4.28661 | 108.398 | 0.651706 | 6.577521 | 4.78E-11 | 1.3E-09 | 1.57E-09 | 4.231008 | 16.72038 | 114.3067 | 298.3338 | 240873 | tumor necrosis factor (ligand) superfamily, member 18 [Source:MGI Symbol;Acc:MGI:2673064] |
| ENSMUSG00000024261 | Syt4 | -4.2791 | 104.0156 | 0.579786 | 7.38048 | 1.58E-13 | 5.78E-12 | 7E-12 | 9.519767 | 10.81907 | 117.9645 | 277.7591 | 20983 | synaptotagmin IV [Source:MGI Symbol;Acc:MGI:101759] |
| ENSMUSG00000042662 | Dusp15 | -4.27533 | 10.23626 | 1.618598 | 2.64138 | 0.008257 | 0.036427 | 0.043999 | 0 | 1.967103 | 9.144538 | 29.83338 | 252864 | dual specificity phosphatase-like 15 [Source:MGI Symbol;Acc:MGI:1934928] |
| ENSMUSG00000031425 | Plp1 | -4.21017 | 1719.72 | 0.823112 | 5.114936 | 3.14E-07 | 4.59E-06 | 5.56E-06 | 132.219 | 220.3156 | 1580.176 | 4946.169 | 18823 | proteolipid protein (myelin) 1 [Source:MGI Symbol;Acc:MGI:97623] |
| ENSMUSG00000035864 | Syt1 | -4.16754 | 9.816222 | 1.572971 | 2.649474 | 0.008062 | 0.035659 | 0.043071 | 2.115504 | 0 | 23.7758 | 13.37359 | 20979 | synaptotagmin I [Source:MGI Symbol;Acc:MGI:99667] |
| ENSMUSG00000060716 | Plekhh1 | -4.16685 | 33.81081 | 0.892025 | 4.671222 | 2.99E-06 | 3.57E-05 | 4.32E-05 | 3.173256 | 3.934207 | 36.57815 | 91.55762 | 211945 | pleckstrin homology domain containing, family H (with MyTH4 domain) member 1 [Source:MGI Symbol;Acc:MGI:2144989] |
| ENSMUSG00000016346 | Kcnq2 | -4.15845 | 14.35855 | 1.234037 | 3.36979 | 0.000752 | 0.004917 | 0.005956 | 1.057752 | 1.967103 | 25.60471 | 28.80465 | 16536 | potassium voltage-gated channel, subfamily Q, member 2 [Source:MGI Symbol;Acc:MGI:1309503] |
| ENSMUSG00000029832 | Nfe2l3 | -4.14125 | 9.511871 | 1.49791 | 2.764687 | 0.005698 | 0.026725 | 0.032284 | 1.057752 | 0.983552 | 16.46017 | 19.54601 | 18025 | nuclear factor, erythroid derived 2, like 3 [Source:MGI Symbol;Acc:MGI:1339958] |
| [illegible] | [illegible] | [illegible] | [illegible] | [illegible] | [illegible] | [illegible] | [illegible] | [illegible] | [illegible] | [illegible] | [illegible] | [illegible] | [illegible] | [illegible] |

| | | | | | | | | | | | | | |
|---|---|---|---|---|---|---|---|---|---|---|---|---|---|
| ENSMUSG00000036585 | Fgf1 | -1.7184 | 51.5701 | 0.533613 | 3.220304 | 0.001281 | 0.007717 | 0.009349 | 26.4438 | 21.63814 | 67.66958 | 90.52889 | 14164 | fibroblast growth factor 1 [Source:MGI Symbol;Acc:MGI:95515] |
| ENSMUSG00000059146 | Ntrk3;Ntrk3 | -1.71764 | 71.79617 | 0.459538 | 3.737753 | 0.000186 | 0.001439 | 0.001743 | 28.5593 | 38.35852 | 122.5368 | 97.73005 | 18213;414121 | neurotrophic tyrosine kinase, receptor, type 3 [Source:MGI Symbol;Acc:MGI:97385];neurotrophic tyrosine kinase, receptor, type 3 [Source:MGI Symbol;Acc:MGI:[illegible] |
| ENSMUSG00000024935 | Slc1a1 | -1.71741 | 63.33838 | 0.62649 | 2.741314 | 0.006119 | 0.028358 | 0.034258 | 12.69302 | 46.22693 | 98.76101 | 95.67257 | 20510 | solute carrier family 1 (neuronal/epithelial high affinity glutamate transporter, system Xag), member 1 [Source:MGI Symbol;Acc:MGI:105083] |
| ENSMUSG00000018427 | Ypel2 | -1.71697 | 201.2784 | 0.306814 | 5.596133 | 2.19E-08 | 4.07E-07 | 4.93E-07 | 79.33139 | 108.1907 | 268.8494 | 348.742 | 77864 | yippee like 2 [Source:MGI Symbol;Acc:MGI:1925114] |
| ENSMUSG00000031618 | Nr3c2 | -1.7162 | 110.5629 | 0.403469 | 4.253624 | 2.1E-05 | 0.000209 | 0.000253 | 38.07907 | 64.91441 | 158.2005 | 181.0578 | 110784 | nuclear receptor subfamily 3, group C, member 2 [Source:MGI Symbol;Acc:MGI:99459] |
| ENSMUSG00000039304 | Tnfsf10 | -1.71195 | 41.01899 | 0.653028 | 2.62156 | 0.008753 | 0.038193 | 0.046133 | 13.75077 | 24.58879 | 80.47194 | 45.26444 | 22035 | tumor necrosis factor (ligand) superfamily, member 10 [Source:MGI Symbol;Acc:MGI:107414] |
| ENSMUSG00000003617 | Cp | -1.70485 | 1663.182 | 0.383503 | 4.445479 | 8.77E-06 | 9.59E-05 | 0.000116 | 582.8213 | 978.6339 | 2329.114 | 2762.16 | 12870 | ceruloplasmin [Source:MGI Symbol;Acc:MGI:88476] |
| ENSMUSG00000024598 | Fbn2 | -1.70414 | 195.9129 | 0.422864 | 4.030003 | 5.58E-05 | 0.0005 | 0.000605 | 132.219 | 52.12824 | 341.0913 | 258.2131 | 14119 | fibrillin 2 [Source:MGI Symbol;Acc:MGI:95490] |
| ENSMUSG00000030486 | Zfp108 | -1.70291 | 47.20925 | 0.614157 | 2.772759 | 0.005558 | 0.026215 | 0.031667 | 13.75077 | 30.4901 | 61.26841 | 83.32773 | 54678 | zinc finger protein 108 [Source:MGI Symbol;Acc:MGI:1891198] |
| ENSMUSG00000116097 | Gm36738 | -1.69821 | 56.50299 | 0.503415 | 3.373381 | 0.000743 | 0.004865 | 0.005894 | 30.67481 | 22.62169 | 83.2153 | 89.50015 | | predicted gene, 36738 [Source:MGI Symbol;Acc:MGI:5595897] |
| ENSMUSG00000056073 | Grik2 | -1.68684 | 41.50342 | 0.623975 | 2.70337 | 0.006864 | 0.031208 | 0.037703 | 12.69302 | 26.5559 | 64.01177 | 62.75298 | 14806 | glutamate receptor, ionotropic, kainate 2 (beta 2) [Source:MGI Symbol;Acc:MGI:95815] |
| ENSMUSG00000055022 | Cntn1 | -1.68008 | 533.4785 | 0.274074 | 6.130027 | 8.79E-10 | 1.99E-08 | 2.41E-08 | 176.6446 | 330.4734 | 853.1854 | 773.6105 | 12805 | contactin 1 [Source:MGI Symbol;Acc:MGI:105980] |
| ENSMUSG00000030643 | Rab30 | -1.6799 | 60.32035 | 0.55919 | 3.004171 | 0.002663 | 0.014119 | 0.017097 | 16.92403 | 40.32562 | 91.44538 | 92.58636 | 75985 | RAB30, member RAS oncogene family [Source:MGI Symbol;Acc:MGI:1923235] |
| ENSMUSG00000027581 | Stmn3 | -1.6778 | 43.51826 | 0.592709 | 2.830734 | 0.004644 | 0.022577 | 0.027297 | 14.80853 | 26.5559 | 65.84068 | 66.86793 | 20262 | stathmin-like 3 [Source:MGI Symbol;Acc:MGI:1277137] |
| ENSMUSG00000024501 | Dpysl3 | -1.67565 | 685.5787 | 0.480948 | 3.48406 | 0.000494 | 0.003427 | 0.004152 | 217.8969 | 435.7134 | 1091.858 | 996.8465 | 22240 | dihydropyrimidinase-like 3 [Source:MGI Symbol;Acc:MGI:1349762] |
| ENSMUSG00000026380 | Tfcp2l1 | -1.67408 | 52.11207 | 0.520918 | 3.213709 | 0.00131 | 0.007875 | 0.009539 | 21.15504 | 28.523 | 79.55748 | 79.21278 | 81879 | transcription factor CP2-like 1 [Source:MGI Symbol;Acc:MGI:2444691] |
| ENSMUSG00000028351 | Brinp1 | -1.67296 | 41.23626 | 0.626614 | 2.669849 | 0.007589 | 0.033978 | 0.041051 | 13.75077 | 25.57234 | 73.15631 | 52.4656 | 56710 | bone morphogenic protein/retinoic acid inducible neural specific 1 [Source:MGI Symbol;Acc:MGI:1928478] |
| ENSMUSG00000050953 | Gja1 | -1.66205 | 852.038 | 0.221737 | 7.495617 | 6.6E-14 | 2.51E-12 | 3.05E-12 | 342.7116 | 475.0555 | 1069.911 | 1520.474 | 14609 | gap junction protein, alpha 1 [Source:MGI Symbol;Acc:MGI:95713] |
| ENSMUSG00000042428 | Mgat3 | -1.66012 | 90.48606 | 0.447384 | 3.710738 | 0.000207 | 0.001584 | 0.001919 | 31.73256 | 55.07889 | 119.7935 | 155.3393 | 17309 | mannoside acetylglucosaminyltransferase 3 [Source:MGI Symbol;Acc:MGI:104532] |
| ENSMUSG00000026564 | Stysl2 | -1.6596 | 48.15207 | 0.578034 | 2.871106 | 0.00409 | 0.020349 | 0.024609 | 30.67481 | 15.73683 | 73.15631 | 73.04035 | 240892 | serine/threonine/tyrosine interacting like 2 [Source:MGI Symbol;Acc:MGI:2685055] |
| ENSMUSG00000030074 | Gxylt2 | -1.65707 | 68.46772 | 0.51235 | 3.234262 | 0.00122 | 0.007409 | 0.008975 | 44.42558 | 21.63814 | 98.76101 | 109.0462 | 232313 | glucoside xylosyltransferase 2 [Source:MGI Symbol;Acc:MGI:2682940] |
| ENSMUSG00000039960 | Rhou | -1.65245 | 61.2559 | 0.561466 | 2.943091 | 0.00325 | 0.016748 | 0.020274 | 28.5593 | 30.4901 | 59.4395 | 126.5347 | 69581 | ras homolog family member U [Source:MGI Symbol;Acc:MGI:1916831] |
| ENSMUSG00000057706 | Mex3b | -1.64841 | 354.5882 | 0.264516 | 6.231808 | 4.61E-10 | 1.09E-08 | 1.32E-08 | 131.1612 | 211.4636 | 521.2387 | 554.4894 | 108797 | mex3 RNA binding family member B [Source:MGI Symbol;Acc:MGI:1918252] |
| ENSMUSG00000024176 | Sox8 | -1.64665 | 75.61797 | 0.568028 | 2.898888 | 0.003745 | 0.018884 | 0.022845 | 17.98178 | 55.07889 | 123.4513 | 105.9599 | 20681 | SRY (sex determining region Y)-box 8 [Source:MGI Symbol;Acc:MGI:98370] |
| ENSMUSG00000109244 | Gm44751 | -1.64549 | 615.3219 | 0.262803 | 6.261286 | 3.82E-10 | 9.07E-09 | 1.1E-08 | 338.4806 | 257.6905 | 699.5572 | 1165.559 | | predicted gene 44751 [Source:MGI Symbol;Acc:MGI:5753327] |
| ENSMUSG00000034855 | Cxcl10 | -1.64141 | 275.2029 | 0.343143 | 4.783448 | 1.72E-06 | 2.15E-05 | 2.61E-05 | 165.0093 | 102.2894 | 310.9143 | 522.5986 | 15945 | C-X-C motif chemokine ligand 10 [Source:MGI Symbol;Acc:MGI:1352450] |
| ENSMUSG00000028716 | Pdzk1ip1 | -1.63763 | 50.07582 | 0.555647 | 2.947247 | 0.003206 | 0.016553 | 0.020038 | 20.09729 | 28.523 | 62.18286 | 89.50015 | 67182 | PDZK1 interacting protein 1 [Source:MGI Symbol;Acc:MGI:1914432] |
| ENSMUSG00000040564 | Apoc1 | -1.63605 | 698.0189 | 0.207622 | 7.879941 | 3.28E-15 | 1.44E-13 | 1.75E-13 | 301.4593 | 377.6839 | 897.9937 | 1214.939 | 11812 | apolipoprotein C-I [Source:MGI Symbol;Acc:MGI:88053] |
| ENSMUSG00000034936 | Arl4d | -1.6336 | 45.81608 | 0.552662 | 2.955878 | 0.003118 | 0.016151 | 0.019559 | 19.03953 | 25.57234 | 67.66958 | 70.98288 | 80981 | ADP-ribosylation factor-like 4D [Source:MGI Symbol;Acc:MGI:1933155] |
| ENSMUSG00000101389 | Ms4a4a | -1.63302 | 518.392 | 0.178294 | 9.159098 | 5.23E-20 | 3.93E-18 | 4.76E-18 | 234.8209 | 270.4767 | 778.2002 | 790.0703 | 666907 | membrane-spanning 4-domains, subfamily A, member 4A [Source:MGI Symbol;Acc:MGI:3643932] |
| ENSMUSG00000103197 | Gm37642 | -1.6291 | 55.86005 | 0.557794 | 2.920616 | 0.003493 | 0.01779 | 0.021528 | 37.02132 | 17.70393 | 90.53093 | 78.18404 | | predicted gene, 37642 [Source:MGI Symbol;Acc:MGI:5610870] |
| ENSMUSG00000046160 | Olig1 | -1.62868 | 160.9308 | 0.314089 | 5.1854 | 2.16E-07 | 3.28E-06 | 3.97E-06 | 66.63837 | 90.48676 | 238.6724 | 247.9257 | 50914 | oligodendrocyte transcription factor 1 [Source:MGI Symbol;Acc:MGI:1355334] |
| ENSMUSG00000121865 | | -1.6256 | 293.1154 | 0.288402 | 5.636558 | 1.73E-08 | 3.25E-07 | 3.94E-07 | 133.2767 | 153.4341 | 345.6635 | 540.0871 | 381650 | THAP domain containing 6 [Source:NCBI gene (formerly Entrezgene);Acc:381650] |
| ENSMUSG00000033453 | Adamts15 | -1.62103 | 65.843 | 0.478189 | 3.389947 | 0.000699 | 0.004617 | 0.005592 | 39.13682 | 25.57234 | 97.84656 | 100.8163 | 235130 | ADAM metallopeptidase with thrombospondin type 1 motif 15 [Source:MGI Symbol;Acc:MGI:2449569] |
| ENSMUSG00000102856 | Gm37084 | -1.61875 | 2491.961 | 0.341652 | 4.738015 | 2.16E-06 | 2.64E-05 | 3.2E-05 | 1453.351 | 995.3543 | 4312.564 | 3206.574 | | predicted gene, 37084 [Source:MGI Symbol;Acc:MGI:5610312] |
| ENSMUSG00000040836 | Gpr161 | -1.61867 | 52.35822 | 0.567544 | 2.852056 | 0.004344 | 0.021357 | 0.025821 | 16.92403 | 34.42431 | 85.04421 | 73.04035 | 240888 | G protein-coupled receptor 161 [Source:MGI Symbol;Acc:MGI:2685054] |
| ENSMUSG00000033960 | Jcad | -1.61445 | 167.1454 | 0.329238 | 4.903601 | 9.41E-07 | 1.25E-05 | 1.51E-05 | 71.92713 | 92.45386 | 213.0677 | 291.1327 | 240185 | junctional cadherin 5 associated [Source:MGI Symbol;Acc:MGI:2685174] |
| ENSMUSG00000025272 | Tro | -1.6097 | 98.45734 | 0.409935 | 3.926734 | 8.61E-05 | 0.000731 | 0.000885 | 38.07907 | 59.0131 | 160.9439 | 135.7933 | 56191 | trophinin [Source:MGI Symbol;Acc:MGI:1928994] |
| ENSMUSG00000063455 | D630045J12Rik | -1.60687 | 50.97929 | 0.580076 | 2.770112 | 0.005604 | 0.026377 | 0.031864 | 15.86628 | 34.42431 | 79.55748 | 74.06909 | 330286 | RIKEN cDNA D630045J12 gene [Source:MGI Symbol;Acc:MGI:2669829] |
| ENSMUSG00000022773 | Ypel1 | -1.59838 | 36.04892 | 0.615088 | 2.598615 | 0.00936 | 0.040332 | 0.048718 | 20.09729 | 15.73683 | 54.86723 | 53.49434 | 106369 | yippee like 1 [Source:MGI Symbol;Acc:MGI:1913303] |
| ENSMUSG00000020641 | Rsad2 | -1.59673 | 575.8557 | 0.190367 | 8.387665 | 4.96E-17 | 2.64E-15 | 3.2E-15 | 281.362 | 291.1313 | 966.5777 | 764.3519 | 58185 | radical S-adenosyl methionine domain containing 2 [Source:MGI Symbol;Acc:MGI:1929628] |
| ENSMUSG00000025104 | Hdgfl3 | -1.59527 | 140.1183 | 0.381657 | 4.179849 | 2.92E-05 | 0.000282 | 0.000341 | 48.65659 | 90.48676 | 209.4099 | 211.9199 | 29877 | HDGF like 3 [Source:MGI Symbol;Acc:MGI:1352760] |
| ENSMUSG00000009376 | Met | -1.59265 | 77.27401 | 0.453452 | 3.512278 | 0.000444 | 0.003126 | 0.003787 | 29.61705 | 47.21048 | 108.82 | 123.4485 | 17295 | met proto-oncogene [Source:MGI Symbol;Acc:MGI:96969] |
| ENSMUSG00000120977 | Gm32164 | -1.59086 | 44.86304 | 0.552044 | 2.881768 | 0.003955 | 0.019809 | 0.023957 | 20.09729 | 24.58879 | 65.84068 | 68.9254 | | predicted gene, 32164 [Source:MGI Symbol;Acc:MGI:5591323] |
| ENSMUSG00000001555 | Fkbp10 | -1.58795 | 74.96713 | 0.453599 | 3.500777 | 0.000464 | 0.003245 | 0.00393 | 44.42558 | 30.4901 | 101.5044 | 123.4485 | 14230 | FK506 binding protein 10 [Source:MGI Symbol;Acc:MGI:104769] |
| ENSMUSG00000109498 | Gm45222 | -1.58731 | 961.1939 | 0.181118 | 8.763945 | 1.89E-18 | 1.14E-16 | 1.38E-16 | 553.2043 | 407.1904 | 1335.103 | 1549.278 | | predicted gene 45222 [Source:MGI Symbol;Acc:MGI:5753798] |
| ENSMUSG00000033152 | Podxl2 | -1.58531 | 72.75853 | 0.484513 | 3.271956 | 0.001068 | 0.006635 | 0.008037 | 25.38605 | 47.21048 | 104.2477 | 114.1898 | 319655 | podocalyxin-like 2 [Source:MGI Symbol;Acc:MGI:2442488] |
| ENSMUSG00000067879 | Vxn | -1.57299 | 173.1119 | 0.484806 | 3.244577 | 0.001176 | 0.007195 | 0.008715 | 40.19457 | 133.763 | 279.8229 | 238.6671 | 76982 | vexin [Source:MGI Symbol;Acc:MGI:1924232] |
| ENSMUSG00000019647 | Sema6a | -1.5725 | 64.44739 | 0.513129 | 3.064524 | 0.00218 | 0.011969 | 0.014499 | 25.38605 | 39.34207 | 76.81412 | 116.2473 | 20358 | sema domain, transmembrane domain (TM), and cytoplasmic domain, (semaphorin) 6A [Source:MGI Symbol;Acc:MGI:1203727] |
| ENSMUSG00000106946 | Gm42856 | -1.57048 | 70.23157 | 0.465341 | 3.374902 | 0.000738 | 0.004847 | 0.005871 | 42.31008 | 28.523 | 96.9321 | 113.1611 | | predicted gene 42856 [Source:MGI Symbol;Acc:MGI:5662993] |
| ENSMUSG00000040624 | Plekhg1 | -1.57023 | 76.26508 | 0.580587 | 2.704561 | 0.006839 | 0.031125 | 0.037602 | 27.50155 | 49.17758 | 65.84068 | 162.5405 | 213783 | pleckstrin homology domain containing, family G (with RhoGef domain) member 1 [Source:MGI Symbol;Acc:MGI:2676551] |
| ENSMUSG00000042659 | Arrdc4 | -1.56636 | 697.584 | 0.18842 | 8.313171 | 9.32E-17 | 4.79E-15 | 5.8E-15 | 302.517 | 401.2891 | 977.5511 | 1108.979 | 66412 | arrestin domain containing 4 [Source:MGI Symbol;Acc:MGI:1913662] |
| ENSMUSG00000052727 | Map1b | -1.56271 | 527.6884 | 0.246834 | 6.331011 | 2.44E-10 | 5.95E-09 | 7.2E-09 | 199.9151 | 333.424 | 770.8846 | 806.5301 | 17755 | microtubule-associated protein 1B [Source:MGI Symbol;Acc:MGI:1306778] |
| ENSMUSG00000046463 | 5930403N24Rik | -1.56232 | 133.729 | 0.332179 | 4.703251 | 2.56E-06 | 3.09E-05 | 3.74E-05 | 62.40736 | 72.78283 | 184.7197 | 215.0061 | | RIKEN cDNA 5930403N24 gene [Source:MGI Symbol;Acc:MGI:2444171] |
| ENSMUSG00000040093 | Bmf | -1.56221 | 705.4695 | 0.173893 | 8.983737 | 2.62E-19 | 1.79E-17 | 2.17E-17 | 313.0946 | 400.3055 | 1048.879 | 1059.599 | 171543 | BCL2 modifying factor [Source:MGI Symbol;Acc:MGI:2176433] |
| ENSMUSG00000005125 | Ndrg1 | -1.56149 | 175.0445 | 0.454351 | 3.436754 | 0.000589 | 0.003989 | 0.004832 | 53.94535 | 122.944 | 183.8052 | 339.4833 | 17988 | N-myc downstream regulated gene 1 [Source:MGI Symbol;Acc:MGI:1341799] |
| ENSMUSG00000048388 | Fam171b | -1.55727 | 158.2996 | 0.373008 | 4.174891 | 2.98E-05 | 0.000287 | 0.000348 | 56.06085 | 104.2565 | 216.7256 | 256.1556 | 241520 | family with sequence similarity 171, member B [Source:MGI Symbol;Acc:MGI:2444579] |
| ENSMUSG00000064373 | Selenop | -1.55682 | 72349.36 | 0.229818 | 6.774152 | 1.25E-11 | 3.65E-10 | 4.42E-10 | 31241.76 | 42170.76 | 101853.7 | 114131.2 | 20363 | selenoprotein P [Source:MGI Symbol;Acc:MGI:894288] |
| ENSMUSG00000039713 | Plekhg5 | -1.55676 | 251.4293 | 0.247259 | 6.296081 | 3.05E-10 | 7.37E-09 | 8.93E-09 | 115.295 | 139.6643 | 372.1827 | 378.5753 | 269608 | pleckstrin homology domain containing, family G (with RhoGef domain) member 5 [Source:MGI Symbol;Acc:MGI:2652860] |
| ENSMUSG00000086782 | E130102H24Rik | -1.55314 | 95.30686 | 0.444289 | 3.495775 | 0.000473 | 0.003297 | 0.003993 | 32.79031 | 63.93086 | 143.5692 | 140.937 | 77866 | RIKEN cDNA E130102H24 gene [Source:MGI Symbol;Acc:MGI:1925116] |
| ENSMUSG00000042284 | Itga1 | -1.55254 | 87.49145 | 0.485083 | 3.200573 | 0.001372 | 0.008188 | 0.009918 | 31.73256 | 57.046 | 99.67547 | 161.5118 | 109700 | integrin alpha 1 [Source:MGI Symbol;Acc:MGI:96599] |
| ENSMUSG00000130847 | Gm67095 | -1.55224 | 265.9122 | 0.406518 | 3.818371 | 0.000134 | 0.001081 | 0.00131 | 187.2221 | 83.60189 | 504.7785 | 288.0465 | | predicted gene, 67095 [Source:MGI Symbol;Acc:MGI:7817032] |
| ENSMUSG00000039542 | Ncam1 | -1.54988 | 269.3634 | 0.383215 | 4.044427 | 5.25E-05 | 0.000473 | 0.000573 | 78.27364 | 195.7268 | 419.7343 | 383.719 | 17967 | neural cell adhesion molecule 1 [Source:MGI Symbol;Acc:MGI:97281] |
| ENSMUSG00000028226 | Mmp16 | -1.5479 | 148.9511 | 0.348528 | 4.441481 | 8.93E-06 | 9.74E-05 | 0.000118 | 58.17636 | 93.43741 | 215.8111 | 228.3797 | 17389 | matrix metallopeptidase 16 [Source:MGI Symbol;Acc:MGI:1276107] |
| ENSMUSG00000023274 | Cd4 | -1.54237 | 249.2344 | 0.243379 | 6.337323 | 2.34E-10 | 5.74E-09 | 6.95E-09 | 127.988 | 126.8782 | 392.3007 | 349.7707 | 12504 | CD4 antigen [Source:MGI Symbol;Acc:MGI:88335] |
| ENSMUSG00000031503 | Col4a2 | -1.53505 | 342.9692 | 0.254732 | 6.026148 | 1.68E-09 | 3.65E-08 | 4.42E-08 | 212.6081 | 139.6643 | 501.1207 | 518.4836 | 12827 | collagen, type IV, alpha 2 [Source:MGI Symbol;Acc:MGI:88455] |
| ENSMUSG00000008999 | Bmp7 | -1.53328 | 66.38232 | 0.45772 | 3.349817 | 0.000809 | 0.005238 | 0.006346 | 31.73256 | 36.39141 | 91.44538 | 105.9599 | 12162 | bone morphogenetic protein 7 [Source:MGI Symbol;Acc:MGI:103302] |
| ENSMUSG00000036158 | Prickle1 | -1.52987 | 73.63272 | 0.475582 | 3.216838 | 0.001296 | 0.007799 | 0.009447 | 42.31008 | 33.44076 | 85.04421 | 133.7359 | 106042 | prickle planar cell polarity protein 1 [Source:MGI Symbol;Acc:MGI:1916034] |
| ENSMUSG00000022816 | Fstl1 | -1.52624 | 872.0385 | 0.201529 | 7.573343 | 3.64E-14 | 1.42E-12 | 1.72E-12 | 505.6054 | 393.4207 | 1096.43 | 1492.698 | 14314 | follistatin-like 1 [Source:MGI Symbol;Acc:MGI:102793] |
| ENSMUSG00000043953 | Ccrl2 | -1.52576 | 1920.448 | 0.325922 | 4.681366 | 2.85E-06 | 3.41E-05 | 4.13E-05 | 1083.138 | 896.9991 | 2260.53 | 3441.126 | 54199 | C-C motif chemokine receptor-like 2 [Source:MGI Symbol;Acc:MGI:1920904] |
| ENSMUSG00000020297 | Nsg2 | -1.52469 | 176.5029 | 0.407706 | 3.739667 | 0.000184 | 0.001428 | 0.00173 | 53.94535 | 127.8617 | 250.5603 | 273.6441 | 18197 | neuron specific gene family member 2 [Source:MGI Symbol;Acc:MGI:1202070] |
| ENSMUSG00000006344 | Ggt5 | -1.52353 | 169.4161 | 0.354172 | 4.30166 | 1.7E-05 | 0.000173 | 0.000209 | 63.46511 | 111.1413 | 230.4424 | 272.6154 | 23887 | gamma-glutamyltransferase 5 [Source:MGI Symbol;Acc:MGI:1346063] |
| ENSMUSG00000029843 | Slc13a4 | -1.51781 | 116.7062 | 0.48164 | 3.151348 | 0.001625 | 0.009416 | 0.011407 | 83.5624 | 37.37496 | 129.8524 | 216.0348 | 243755 | solute carrier family 13 (sodium/sulfate symporters), member 4 [Source:MGI Symbol;Acc:MGI:2442367] |
| ENSMUSG00000060206 | Zfp462 | -1.51512 | 86.89357 | 0.57756 | 2.623316 | 0.008708 | 0.03804 | 0.045948 | 20.09729 | 69.83217 | 144.4837 | 113.1611 | 242466 | zinc finger protein 462 [Source:MGI Symbol;Acc:MGI:107690] |
| ENSMUSG00000037762 | Slc16a9 | -1.51395 | 61.94095 | 0.506574 | 2.9886 | 0.002803 | 0.01474 | 0.017849 | 31.73256 | 32.45721 | 70.41294 | 113.1611 | 66859 | solute carrier family 16 (monocarboxylic acid transporters), member 9 [Source:MGI Symbol;Acc:MGI:1914109] |
| ENSMUSG00000027030 | Stk39 | -1.50659 | 128.1913 | 0.428819 | 3.513342 | 0.000443 | 0.003115 | 0.003773 | 80.38915 | 53.11179 | 134.4247 | 244.8395 | 53416 | serine/threonine kinase 39 [Source:MGI Symbol;Acc:MGI:1858416] |
| ENSMUSG00000026938 | Fcna | -1.50103 | 668.8045 | 0.23698 | 6.334012 | 2.39E-10 | 5.85E-09 | 7.09E-09 | 261.2647 | 436.697 | 1028.761 | 948.4958 | 14133 | ficolin A [Source:MGI Symbol;Acc:MGI:1340905] |
| ENSMUSG00000063611 | Gm10134 | -1.50063 | 146.6037 | 0.33028 | 4.543509 | 5.53E-06 | 6.28E-05 | 7.61E-05 | 63.46511 | 89.5032 | 219.4689 | 213.9774 | | predicted gene 10134 [Source:MGI Symbol;Acc:MGI:3642322] |
| ENSMUSG00000002409 | Dyrk1b | -1.49513 | 76.48571 | 0.447402 | 3.3418 | 0.000832 | 0.005362 | 0.006495 | 33.84806 | 46.22693 | 127.1091 | 98.75879 | 13549 | dual-specificity tyrosine phosphorylation regulated kinase 1b [Source:MGI Symbol;Acc:MGI:1330302] |
| ENSMUSG00000106379 | Lhfpl3 | -1.4925 | 157.7338 | 0.479239 | 3.114304 | 0.001844 | 0.010436 | 0.012642 | 42.31008 | 122.944 | 274.3361 | 191.3451 | 269629 | lipoma HMGIC fusion partner-like 3 [Source:MGI Symbol;Acc:MGI:1925076] |
| ENSMUSG00000050334 | Mir9-2hg;Mir9- | -1.49231 | 175.9529 | 0.371112 | 4.021171 | 5.79E-05 | 0.000517 | 0.000627 | 61.34961 | 122.944 | 255.1326 | 264.3855 | 72677;320201 | Mir9-2 host gene [Source:MGI Symbol;Acc:MGI:2443574];Mir9-2 host gene [Source:MGI Symbol;Acc:MGI:2443574] |
| ENSMUSG00000037379 | Spon2 | -1.4913 | 62.7625 | 0.516281 | 2.888544 | 0.00387 | 0.019445 | 0.023516 | 43.36783 | 22.62169 | 91.44538 | 93.6151 | 100689 | spondin 2, extracellular matrix protein [Source:MGI Symbol;Acc:MGI:1923724] |
| ENSMUSG00000032725 | Folr2 | -1.48658 | 418.5003 | 0.228258 | 6.51271 | 7.38E-11 | 1.95E-09 | 2.37E-09 | 185.1066 | 254.7399 | 570.6192 | 663.5356 | 14276 | folate receptor beta [Source:MGI Symbol;Acc:MGI:95569] |
| ENSMUSG00000000948 | Gm38393 | -1.48643 | 48.94762 | 0.572361 | 2.59701 | 0.009404 | 0.040474 | 0.048891 | 17.98178 | 33.44076 | 79.55748 | 64.81045 | | predicted gene, 38393 [Source:MGI Symbol;Acc:MGI:5613898] |
| ENSMUSG00000026620 | Mark1 | -1.48172 | 47.41396 | 0.545442 | 2.71655 | 0.006597 | 0.030155 | 0.03643 | 25.38605 | 24.58879 | 59.4395 | 80.24151 | 226778 | MAP/microtubule affinity regulating kinase 1 [Source:MGI Symbol;Acc:MGI:2664902] |
| ENSMUSG00000042675 | Ypel3 | -1.48088 | 553.0089 | 0.177751 | 8.331222 | 8E-17 | 4.14E-15 | 5.02E-15 | 265.4957 | 317.6872 | 820.2651 | 808.5876 | 66090 | yippee like 3 [Source:MGI Symbol;Acc:MGI:1913340] |
| ENSMUSG00000003420 | Fcgrt | -1.48077 | 5053.048 | 0.141968 | 10.4303 | 1.8E-25 | 2.49E-23 | 3.01E-23 | 2226.568 | 3104.089 | 7236.988 | 7644.547 | 14132 | Fc fragment of IgG receptor and transporter [Source:MGI Symbol;Acc:MGI:103017] |
| ENSMUSG00000040435 | Ppp1r15a | -1.47576 | 2612.572 | 0.1688 | 8.742633 | 2.28E-18 | 1.35E-16 | 1.64E-16 | 1527.394 | 1236.324 | 3203.332 | 4483.237 | 17872 | protein phosphatase 1, regulatory subunit 15A [Source:MGI Symbol;Acc:MGI:1927072] |
| ENSMUSG00000021356 | Irf4 | -1.4729 | 79.84657 | 0.41186 | 3.576217 | 0.000349 | 0.002529 | 0.003064 | 42.31008 | 42.29272 | 121.6224 | 113.1611 | 16364 | interferon regulatory factor 4 [Source:MGI Symbol;Acc:MGI:1096873] |
| ENSMUSG00000036862 | Dchs1 | -1.47211 | 167.0861 | 0.343439 | 4.286396 | 1.82E-05 | 0.000183 | 0.000222 | 68.75387 | 108.1907 | 274.3361 | 217.0636 | 233651 | dachsous cadherin related 1 [Source:MGI Symbol;Acc:MGI:2685011] |
| ENSMUSG00000021895 | Arhgef3 | -1.45708 | 1188.632 | 0.157615 | 9.244573 | 2.36E-20 | 1.87E-18 | 2.27E-18 | 546.8577 | 721.9269 | 1742.035 | 1743.71 | 71704 | Rho guanine nucleotide exchange factor 3 [Source:MGI Symbol;Acc:MGI:1918954] |
| ENSMUSG00000039997 | Ifi203 | -1.45559 | 342.8673 | 0.231071 | 6.299341 | 2.99E-10 | 7.23E-09 | 8.76E-09 | 159.7205 | 206.5459 | 534.041 | 471.1617 | 15950 | interferon activated gene 203 [Source:MGI Symbol;Acc:MGI:96428] |
| ENSMUSG00000050288 | Fzd2 | -1.4554 | 62.58591 | 0.501176 | 2.903965 | 0.003685 | 0.018617 | 0.022523 | 26.4438 | 40.32562 | 78.64303 | 104.9312 | 57265 | frizzled class receptor 2 [Source:MGI Symbol;Acc:MGI:1888513] |
| ENSMUSG00000024548 | Setbp1 | -1.45369 | 55.76119 | 0.532481 | 2.730032 | 0.006333 | 0.029175 | 0.035246 | 22.21279 | 37.37496 | 91.44538 | 72.01161 | 240427 | SET binding protein 1 [Source:MGI Symbol;Acc:MGI:1933199] |
| ENSMUSG00000017144 | Rnd3 | -1.45024 | 166.3834 | 0.399469 | 3.630405 | 0.000283 | 0.002093 | 0.002535 | 56.06085 | 121.9604 | 223.1267 | 264.3855 | 74194 | Rho family GTPase 3 [Source:MGI Symbol;Acc:MGI:1921444] |
| ENSMUSG00000119979 | Gm57253 | -1.44957 | 58.8762 | 0.483324 | 2.999172 | 0.002707 | 0.014313 | 0.017332 | 30.67481 | 32.45721 | 94.18874 | 78.18404 | | predicted gene, 57253 [Source:MGI Symbol;Acc:MGI:7338419] |
| ENSMUSG00000029999 | Tgfa | -1.44785 | 240.1965 | 0.267732 | 5.407811 | 6.38E-08 | 1.1E-06 | 1.33E-06 | 108.9484 | 148.5163 | 340.1768 | 363.1443 | 21802 | transforming growth factor alpha [Source:MGI Symbol;Acc:MGI:98724] |
| ENSMUSG00000120122 | Gm57067 | -1.44758 | 49.71039 | 0.548774 | 2.637845 | 0.008343 | 0.036723 | 0.044356 | 32.79031 | 20.65459 | 79.55748 | 65.83919 | | predicted gene, 57067 [Source:MGI Symbol;Acc:MGI:7338047] |
| ENSMUSG00000031502 | Col4a1 | -1.44721 | 945.7484 | 0.237344 | 6.097543 | 1.08E-09 | 2.41E-08 | 2.92E-08 | 636.7667 | 378.6674 | 1216.224 | 1551.336 | 12826 | collagen, type IV, alpha 1 [Source:MGI Symbol;Acc:MGI:88454] |
| ENSMUSG00000029126 | Nsg1 | -1.44359 | 96.59314 | 0.377566 | 3.82342 | 0.000132 | 0.001063 | 0.001287 | 49.71434 | 54.09534 | 134.4247 | 148.1382 | 18196 | neuron specific gene family member 1 [Source:MGI Symbol;Acc:MGI:109149] |
| ENSMUSG00000002489 | Tiam1 | -1.44087 | 58.40191 | 0.494155 | 2.915826 | 0.003547 | 0.018021 | 0.021809 | 26.4438 | 36.39141 | 90.53093 | 80.24151 | 21844 | T cell lymphoma invasion and metastasis 1 [Source:MGI Symbol;Acc:MGI:103306] |
| ENSMUSG00000045932 | Ifit2 | -1.43751 | 1132.864 | 0.15843 | 9.073485 | 1.15E-19 | 8.24E-18 | 9.99E-18 | 564.8395 | 657.0125 | 1826.164 | 1483.439 | 15958 | interferon-induced protein with tetratricopeptide repeats 2 [Source:MGI Symbol;Acc:MGI:99449] |
| ENSMUSG00000027546 | Atp9a | -1.43494 | 223.8761 | 0.349408 | 4.106775 | 4.01E-05 | 0.000377 | 0.000457 | 90.96666 | 150.4834 | 256.9615 | 397.0926 | 11981 | ATPase, class II, type 9A [Source:MGI Symbol;Acc:MGI:1330826] |
| ENSMUSG00000087354 | 4930404I05Rik | -1.43324 | 55.48242 | 0.497472 | 2.881057 | 0.003963 | 0.019841 | 0.023995 | 27.50155 | 32.45721 | 86.87311 | 75.09783 | | RIKEN cDNA 4930404I05 gene [Source:MGI Symbol;Acc:MGI:1914644] |
| ENSMUSG00000032690 | Oas2 | -1.43156 | 221.412 | 0.253819 | 5.640075 | 1.7E-08 | 3.19E-07 | 3.86E-07 | 120.5837 | 119.0098 | 337.4335 | 308.6212 | 246728 | 2'-5' oligoadenylate synthetase 2 [Source:MGI Symbol;Acc:MGI:2180852] |
| ENSMUSG00000047793 | Sned1 | -1.431 | 107.95 | 0.477198 | 2.998745 | 0.002711 | 0.014328 | 0.01735 | 83.5624 | 33.44076 | 148.1415 | 166.6555 | 208777 | sushi, nidogen and EGF-like domains 1 [Source:MGI Symbol;Acc:MGI:3045960] |
| ENSMUSG00000042035 | Igsf3 | -1.43057 | 56.54549 | 0.48918 | 2.924417 | 0.003451 | 0.017601 | 0.021307 | 31.73256 | 29.50655 | 87.78757 | 77.1553 | 78908 | immunoglobulin superfamily, member 3 [Source:MGI Symbol;Acc:MGI:1926158] |
| ENSMUSG00000069515 | Lyz1 | -1.42897 | 60.06379 | 0.475527 | 3.005029 | 0.002656 | 0.014084 | 0.017055 | 29.61705 | 35.40786 | 87.78757 | 87.44267 | 17110 | lysozyme 1 [Source:MGI Symbol;Acc:MGI:96902] |
| ENSMUSG00000020593 | Lpin1 | -1.42533 | 578.7172 | 0.176681 | 8.067248 | 7.19E-16 | 3.39E-14 | 4.1E-14 | 283.4775 | 344.2431 | 839.4686 | 847.6796 | 14245 | lipin 1 [Source:MGI Symbol;Acc:MGI:1891340] |
| ENSMUSG00000030930 | Chst15 | -1.4207 | 396.9879 | 0.195067 | 7.283149 | 3.26E-13 | 1.16E-11 | 1.4E-11 | 211.5504 | 220.3156 | 602.6251 | 553.4607 | 77590 | carbohydrate sulfotransferase 15 [Source:MGI Symbol;Acc:MGI:1924840] |
| ENSMUSG00000029108 | Pcdh7 | -1.4161 | 184.2175 | 0.368983 | 3.837852 | 0.000124 | 0.001007 | 0.00122 | 68.75387 | 131.7959 | 235.9291 | 300.3913 | 54216 | protocadherin 7 [Source:MGI Symbol;Acc:MGI:1860487] |
| ENSMUSG00000053965 | Pde5a | -1.4159 | 89.51352 | 0.450986 | 3.139554 | 0.001692 | 0.009722 | 0.011777 | 60.29186 | 37.37496 | 106.0766 | 154.3106 | 242202 | phosphodiesterase 5A, cGMP-specific [Source:MGI Symbol;Acc:MGI:2651499] |
| ENSMUSG00000014164 | Klhl3 | -1.41443 | 57.80029 | 0.482662 | 2.930468 | 0.003385 | 0.017303 | 0.020946 | 29.61705 | 33.44076 | 86.87311 | 81.27025 | 100503085 | kelch-like 3 [Source:MGI Symbol;Acc:MGI:2445185] |
| ENSMUSG00000039830 | Olig2 | -1.41159 | 97.30161 | 0.529544 | 2.665674 | 0.007683 | 0.034347 | 0.041496 | 27.50155 | 78.68414 | 163.6872 | 119.3335 | 50913 | oligodendrocyte transcription factor 2 [Source:MGI Symbol;Acc:MGI:1355331] |
| ENSMUSG00000019944 | Rhobtb1 | -1.4114 | 304.6061 | 0.220648 | 6.396623 | 1.59E-10 | 4.02E-09 | 4.87E-09 | 160.7783 | 172.1215 | 462.7136 | 422.8111 | 69288 | Rho-related BTB domain containing 1 [Source:MGI Symbol;Acc:MGI:1916538] |
| ENSMUSG00000023473 | Celsr3 | -1.40967 | 57.26413 | 0.515195 | 2.73619 | 0.006216 | 0.028707 | 0.03468 | 39.13682 | 23.60524 | 85.04421 | 81.27025 | 107934 | cadherin, EGF LAG seven-pass G-type receptor 3 [Source:MGI Symbol;Acc:MGI:1858236] |
| ENSMUSG00000038668 | Lpar1 | -1.40715 | 123.6747 | 0.434723 | 3.23689 | 0.001208 | 0.007368 | 0.008925 | 48.65659 | 86.55255 | 136.2536 | 223.236 | 14745 | lysophosphatidic acid receptor 1 [Source:MGI Symbol;Acc:MGI:108429] |
| ENSMUSG00000039910 | Cited2 | -1.40232 | 662.6469 | 0.224729 | 6.240038 | 4.37E-10 | 1.03E-08 | 1.25E-08 | 417.812 | 309.8188 | 804.7194 | 1118.238 | 17684 | Cbp/p300-interacting transactivator, with Glu/Asp-rich carboxy-terminal domain, 2 [Source:MGI Symbol;Acc:MGI:1306784] |
| ENSMUSG00000022454 | Nell2 | -1.402 | 77.31651 | 0.462863 | 3.028969 | 0.002454 | 0.013218 | 0.016012 | 30.67481 | 54.09534 | 113.3923 | 111.1036 | 54003 | NEL-like 2 [Source:MGI Symbol;Acc:MGI:1858510] |
| ENSMUSG00000001741 | Il16 | -1.39763 | 1926.787 | 0.125778 | 11.11188 | 1.1E-28 | 1.89E-26 | 2.29E-26 | 958.3232 | 1161.575 | 2863.155 | 2724.097 | 16170 | interleukin 16 [Source:MGI Symbol;Acc:MGI:1270855] |
| ENSMUSG00000028525 | Pde4b | -1.39395 | 657.8776 | 0.186313 | 7.481779 | 7.33E-14 | 2.79E-12 | 3.38E-12 | 368.0977 | 357.0293 | 830.3241 | 1076.059 | 18578 | phosphodiesterase 4B, cAMP specific [Source:MGI Symbol;Acc:MGI:99557] |
| ENSMUSG00000010175 | Prox1 | -1.39346 | 267.386 | 0.248035 | 5.617996 | 1.93E-08 | 3.6E-07 | 4.36E-07 | 150.2008 | 144.5821 | 344.7491 | 430.0122 | 19130 | prospero homeobox 1 [Source:MGI Symbol;Acc:MGI:97772] |
| ENSMUSG00000096764 | Gm21985 | -1.38826 | 160.1012 | 0.448514 | 3.095238 | 0.001967 | 0.011011 | 0.013339 | 129.0457 | 48.19403 | 212.1533 | 251.0119 | | predicted gene 21985 [Source:MGI Symbol;Acc:MGI:5439454] |
| ENSMUSG00000021838 | Samd4 | -1.38733 | 517.9547 | 0.22112 | 6.274121 | 3.52E-10 | 8.4E-09 | 1.02E-08 | 239.0519 | 333.424 | 679.4392 | 819.9037 | 74480 | sterile alpha motif domain containing 4 [Source:MGI Symbol;Acc:MGI:1921730] |
| ENSMUSG00000032024 | Clmp | -1.38694 | 88.09485 | 0.420649 | 3.297133 | 0.000977 | 0.006153 | 0.007454 | 59.23411 | 38.35852 | 126.1946 | 128.5922 | 71566 | CXADR-like membrane protein [Source:MGI Symbol;Acc:MGI:1918816] |
| ENSMUSG00000063063 | Ctnna2 | -1.38414 | 133.1156 | 0.478838 | 2.890626 | 0.003845 | 0.019349 | 0.0234 | 38.07907 | 109.1742 | 185.6341 | 199.575 | 12386 | catenin alpha 2 [Source:MGI Symbol;Acc:MGI:88275] |
| ENSMUSG00000028019 | Pdgfc | -1.38216 | 141.8355 | 0.317183 | 4.357614 | 1.31E-05 | 0.000138 | 0.000167 | 83.5624 | 73.76638 | 192.9498 | 217.0636 | 54635 | platelet-derived growth factor, C polypeptide [Source:MGI Symbol;Acc:MGI:1859631] |
| ENSMUSG00000104214 | Gm36638 | -1.37968 | 58.06175 | 0.494694 | 2.788949 | 0.005288 | 0.025198 | 0.030448 | 37.02132 | 27.53945 | 90.53093 | 77.1553 | | predicted gene, 36638 [Source:MGI Symbol;Acc:MGI:5595797] |
| ENSMUSG00000017386 | Traf4 | -1.37787 | 60.98444 | 0.501405 | 2.748012 | 0.005996 | 0.027913 | 0.03372 | 25.38605 | 42.29272 | 87.78757 | 88.47141 | 22032 | TNF receptor associated factor 4 [Source:MGI Symbol;Acc:MGI:1202880] |
| ENSMUSG00000039109 | F13a1 | -1.37376 | 2398.791 | 0.12168 | 11.28997 | 1.47E-29 | 2.9E-27 | 3.51E-27 | 1199.491 | 1471.393 | 3433.774 | 3490.506 | 74145 | coagulation factor XIII, A1 subunit [Source:MGI Symbol;Acc:MGI:1921395] |
| ENSMUSG00000023826 | Prkn | -1.37128 | 74.72569 | 0.43628 | 3.143111 | 0.001672 | 0.009641 | 0.011679 | 38.07907 | 45.24338 | 118.879 | 96.70131 | 50873 | parkin RBR E3 ubiquitin protein ligase [Source:MGI Symbol;Acc:MGI:1355296] |
| ENSMUSG00000033450 | Tagap | -1.3691 | 1548.193 | 0.15194 | 9.010793 | 2.05E-19 | 1.42E-17 | 1.72E-17 | 944.5725 | 783.8907 | 2003.568 | 2460.74 | 72536 | T cell activation Rho GTPase activating protein [Source:MGI Symbol;Acc:MGI:3615484] |
| ENSMUSG00000032265 | Tent5a | -1.3658 | 1539.273 | 0.134843 | 10.12879 | 4.12E-24 | 4.77E-22 | 5.77E-22 | 833.5085 | 887.1636 | 2012.713 | 2423.705 | 212943 | terminal nucleotidyltransferase 5A [Source:MGI Symbol;Acc:MGI:2670964] |
| ENSMUSG00000037921 | Ddx60 | -1.36455 | 1070.997 | 0.142402 | 9.582391 | 9.48E-22 | 8.61E-20 | 1.04E-19 | 546.8577 | 651.1112 | 1582.005 | 1504.014 | 234311 | DExD/H box helicase 60 [Source:MGI Symbol;Acc:MGI:2384570] |
| ENSMUSG00000021701 | Plk2 | -1.36406 | 2159.397 | 0.318122 | 4.287857 | 1.8E-05 | 0.000182 | 0.00022 | 1276.707 | 1139.936 | 2443.421 | 3777.524 | 20620 | polo like kinase 2 [Source:MGI Symbol;Acc:MGI:1099790] |
| ENSMUSG00000109429 | Gm45223 | -1.35798 | 1256.336 | 0.147116 | 9.230658 | 2.69E-20 | 2.12E-18 | 2.57E-18 | 779.5632 | 631.4402 | 1898.406 | 1715.934 | | predicted gene 45223 [Source:MGI Symbol;Acc:MGI:5753799] |
| ENSMUSG00000032515 | Csrnp1 | -1.35172 | 374.8591 | 0.281962 | 4.794001 | 1.63E-06 | 2.06E-05 | 2.49E-05 | 209.4349 | 212.4472 | 399.6163 | 677.9379 | 215418 | cysteine-serine-rich nuclear protein 1 [Source:MGI Symbol;Acc:MGI:2387989] |
| ENSMUSG00000002289 | Angptl4 | -1.35028 | 111.7195 | 0.363461 | 3.715059 | 0.000203 | 0.001561 | 0.001891 | 71.92713 | 54.09534 | 165.5161 | 155.3393 | 57875 | angiopoietin-like 4 [Source:MGI Symbol;Acc:MGI:1888999] |
| ENSMUSG00000000386 | Mx1 | -1.34447 | 252.1016 | 0.25499 | 5.272645 | 1.34E-07 | 2.13E-06 | 2.58E-06 | 136.45 | 148.5163 | 401.4452 | 321.9948 | | MX dynamin-like GTPase 1 [Source:MGI Symbol;Acc:MGI:97243] |
| ENSMUSG00000029135 | Fosl2 | -1.3396 | 880.7753 | 0.227284 | 5.893923 | 3.77E-09 | 7.83E-08 | 9.49E-08 | 507.7209 | 489.8087 | 954.6898 | 1570.882 | 14284 | fos-like antigen 2 [Source:MGI Symbol;Acc:MGI:102858] |
| ENSMUSG00000058070 | Eml1 | -1.33661 | 125.0088 | 0.513824 | 2.601294 | 0.009287 | 0.040063 | 0.048394 | 41.25232 | 100.3223 | 119.7935 | 238.6671 | 68519 | echinoderm microtubule associated protein like 1 [Source:MGI Symbol;Acc:MGI:1915769] |
| ENSMUSG00000121280 | Gm57189 | -1.33454 | 701.4557 | 0.170409 | 7.831406 | 4.82E-15 | 2.1E-13 | 2.55E-13 | 437.9093 | 358.9964 | 956.5187 | 1052.398 | | predicted gene, 57189 [Source:MGI Symbol;Acc:MGI:7338291] |
| ENSMUSG00000003452 | Bicd1 | -1.33213 | 78.75751 | 0.511526 | 2.604219 | 0.009208 | 0.03978 | 0.048051 | 26.4438 | 62.94731 | 112.4778 | 113.1611 | 12121 | BICD cargo adaptor 1 [Source:MGI Symbol;Acc:MGI:1101760] |
| ENSMUSG00000113216 | Gm40841 | -1.32667 | 75.89042 | 0.42014 | 3.157697 | 0.00159 | 0.009263 | 0.011221 | 41.25232 | 45.24338 | 106.9911 | 110.0749 | | predicted gene, 40841 [Source:MGI Symbol;Acc:MGI:5623726] |
| ENSMUSG00000069792 | Wfdc17 | -1.32665 | 1695.012 | 0.328104 | 4.043372 | 5.27E-05 | 0.000475 | 0.000575 | 780.6209 | 1151.739 | 2121.533 | 2726.154 | 100034251 | WAP four-disulfide core domain 17 [Source:MGI Symbol;Acc:MGI:3649773] |
| ENSMUSG00000035356 | Nfkbiz | -1.30621 | 3889.508 | 0.333365 | 3.918267 | 8.92E-05 | 0.000753 | 0.000912 | 2647.553 | 1832.357 | 4744.186 | 6333.936 | 80859 | nuclear factor of kappa light polypeptide gene enhancer in B cells inhibitor, zeta [Source:MGI Symbol;Acc:MGI:1931595] |
| ENSMUSG00000034675 | Dbn1 | -1.30575 | 98.64341 | 0.392327 | 3.32822 | 0.000874 | 0.005597 | 0.00678 | 47.59884 | 65.89796 | 129.8524 | 151.2244 | 56320 | drebrin 1 [Source:MGI Symbol;Acc:MGI:1931838] |
| ENSMUSG00000036745 | Ttll7 | -1.29911 | 90.19512 | 0.478092 | 2.717276 | 0.006582 | 0.030098 | 0.036361 | 39.13682 | 64.91441 | 94.18874 | 162.5405 | 70892 | tubulin tyrosine ligase-like family, member 7 [Source:MGI Symbol;Acc:MGI:1918142] |
| ENSMUSG00000041132 | N4bp2l1 | -1.2987 | 500.0444 | 0.175929 | 7.381985 | 1.56E-13 | 5.73E-12 | 6.94E-12 | 273.9577 | 303.9175 | 703.215 | 719.0874 | 100637 | NEDD4 binding protein 2-like 1 [Source:MGI Symbol;Acc:MGI:2140872] |
| ENSMUSG00000041775 | Mapk1ip1 | -1.29693 | 84.6489 | 0.398444 | 3.254989 | 0.001134 | 0.00697 | 0.008443 | 50.77209 | 47.21048 | 124.3657 | 116.2473 | 69546 | mitogen-activated protein kinase 1 interacting protein 1 [Source:MGI Symbol;Acc:MGI:1916796] |
| ENSMUSG00000021250 | Fos | -1.29691 | 7484.681 | 0.106911 | 12.13077 | 7.26E-34 | 2.5E-31 | 3.03E-31 | 4782.096 | 3879.128 | 10593.03 | 10684.47 | 14281 | FBJ osteosarcoma oncogene [Source:MGI Symbol;Acc:MGI:95574] |
| ENSMUSG00000049037 | Clec4a1 | -1.29687 | 347.0098 | 0.202744 | 6.396613 | 1.59E-10 | 4.02E-09 | 4.87E-09 | 198.8574 | 202.6116 | 485.575 | 500.9951 | 269799 | C-type lectin domain family 4, member a1 [Source:MGI Symbol;Acc:MGI:3036291] |
| ENSMUSG00000052534 | Pbx1 | -1.29605 | 297.3271 | 0.269724 | 4.805088 | 1.55E-06 | 1.96E-05 | 2.38E-05 | 135.3922 | 208.513 | 438.0234 | 407.38 | 18514 | pre B cell leukemia homeobox 1 [Source:MGI Symbol;Acc:MGI:97495] |
| ENSMUSG00000027646 | Src | -1.29573 | 224.5791 | 0.356734 | 3.63221 | 0.000281 | 0.002082 | 0.002522 | 84.62015 | 175.0722 | 306.342 | 332.2822 | 20779 | Rous sarcoma oncogene [Source:MGI Symbol;Acc:MGI:98397] |
| ENSMUSG00000041797 | Abca9 | -1.29284 | 8452.958 | 0.09275 | 13.93895 | 3.67E-44 | 2.78E-41 | 3.37E-41 | 4556.795 | 5242.331 | 11615.39 | 12397.31 | 217262 | ATP-binding cassette, sub-family A member 9 [Source:MGI Symbol;Acc:MGI:2386796] |
| ENSMUSG00000037605 | Adgrl3 | -1.29137 | 139.4028 | 0.452507 | 2.853811 | 0.00432 | 0.021267 | 0.025712 | 45.48333 | 116.0591 | 213.9822 | 182.0865 | 319387 | adhesion G protein-coupled receptor L3 [Source:MGI Symbol;Acc:MGI:2441950] |
| ENSMUSG00000045382 | Cxcr4 | -1.29022 | 1567.283 | 0.182431 | 7.072378 | 1.52E-12 | 4.98E-11 | 6.03E-11 | 772.1589 | 1046.499 | 1924.925 | 2525.55 | 12767 | C-X-C motif chemokine receptor 4 [Source:MGI Symbol;Acc:MGI:109563] |
| ENSMUSG00000046743 | Fat4 | -1.28954 | 161.3639 | 0.347299 | 3.713058 | 0.000205 | 0.001572 | 0.001904 | 88.85116 | 98.35517 | 177.404 | 280.8453 | 329628 | FAT atypical cadherin 4 [Source:MGI Symbol;Acc:MGI:3045256] |
| ENSMUSG00000002265 | Peg3 | -1.28931 | 291.7311 | 0.227506 | 5.667141 | 1.45E-08 | 2.76E-07 | 3.34E-07 | 174.5291 | 164.2531 | 386.814 | 441.3283 | 18616 | paternally expressed 3 [Source:MGI Symbol;Acc:MGI:104748] |
| ENSMUSG00000048120 | Entpd1 | -1.28873 | 3375.343 | 0.134392 | 9.589341 | 8.86E-22 | 8.1E-20 | 9.81E-20 | 1694.519 | 2225.777 | 4606.104 | 4974.974 | 12495 | ectonucleoside triphosphate diphosphohydrolase 1 [Source:MGI Symbol;Acc:MGI:102805] |
| ENSMUSG00000143907 | Gm74406 | -1.28783 | 605.4744 | 0.222797 | 5.780279 | 7.46E-09 | 1.48E-07 | 1.79E-07 | 353.2891 | 350.1444 | 687.6693 | 1030.795 | | predicted gene, 74406 [Source:MGI Symbol;Acc:MGI:7831548] |
| ENSMUSG00000043832 | Clec4a3 | -1.28761 | 599.5529 | 0.164799 | 7.813247 | 5.57E-15 | 2.4E-13 | 2.91E-13 | 326.8453 | 369.8154 | 847.6987 | 853.852 | 73149 | C-type lectin domain family 4, member a3 [Source:MGI Symbol;Acc:MGI:1920399] |
| ENSMUSG00000061603 | Akap6 | -1.28025 | 120.554 | 0.470079 | 2.723479 | 0.00646 | 0.029655 | 0.035825 | 39.13682 | 101.3058 | 179.2329 | 162.5405 | 238161 | A kinase anchor protein 6 [Source:MGI Symbol;Acc:MGI:3050566] |
| ENSMUSG00000048249 | Crebrf | -1.28018 | 1786.543 | 0.133287 | 9.604675 | 7.64E-22 | 7.02E-20 | 8.5E-20 | 929.7639 | 1153.706 | 2458.966 | 2603.734 | 77128 | CREB3 regulatory factor [Source:MGI Symbol;Acc:MGI:1924378] |
| ENSMUSG00000008575 | Nfib | -1.27355 | 409.1235 | 0.30124 | 4.227696 | 2.36E-05 | 0.000231 | 0.00028 | 160.7783 | 317.6872 | 578.8493 | 579.1791 | 18028 | nuclear factor I/B [Source:MGI Symbol;Acc:MGI:103188] |
| ENSMUSG00000027298 | Tyro3 | -1.27204 | 65.99332 | 0.474966 | 2.678167 | 0.007403 | 0.033258 | 0.040181 | 34.90581 | 42.29272 | 77.72857 | 109.0462 | 22174 | TYRO3 protein tyrosine kinase 3 [Source:MGI Symbol;Acc:MGI:104294] |
| ENSMUSG00000030157 | Clec2d | -1.27095 | 685.5517 | 0.15363 | 8.272765 | 1.31E-16 | 6.55E-15 | 7.93E-15 | 420.9853 | 382.6016 | 959.2621 | 979.358 | 93694 | C-type lectin domain family 2, member d [Source:MGI Symbol;Acc:MGI:2135589] |
| ENSMUSG00000027435 | Cd93 | -1.26751 | 3422.736 | 0.095908 | 13.21585 | 7.11E-40 | 3.72E-37 | 4.5E-37 | 1897.607 | 2119.554 | 4745.101 | 4928.681 | 17064 | CD93 antigen [Source:MGI Symbol;Acc:MGI:106664] |
| ENSMUSG00000004044 | Cavin1 | -1.26646 | 252.978 | 0.257815 | 4.912276 | 9E-07 | 1.2E-05 | 1.45E-05 | 154.4318 | 142.615 | 313.6577 | 401.2076 | 19285 | caveolae associated 1 [Source:MGI Symbol;Acc:MGI:1277968] |
| ENSMUSG00000028456 | Unc13b | -1.26518 | 181.1696 | 0.304096 | 4.160445 | 3.18E-05 | 0.000304 | 0.000369 | 87.79341 | 124.9111 | 249.6459 | 262.328 | 22249 | unc-13 homolog B [Source:MGI Symbol;Acc:MGI:1342278] |
| ENSMUSG00000033565 | Rbfox2 | -1.26384 | 229.4227 | 0.248751 | 5.080735 | 3.76E-07 | 5.4E-06 | 6.54E-06 | 129.0457 | 140.6479 | 313.6577 | 334.3396 | 93686 | RNA binding protein, fox-1 homolog (C. elegans) 2 [Source:MGI Symbol;Acc:MGI:1933973] |
| ENSMUSG00000041849 | Card6 | -1.26368 | 61.78901 | 0.472544 | 2.674207 | 0.007491 | 0.033574 | 0.040562 | 41.25232 | 31.47365 | 85.95866 | 88.47141 | 239319 | caspase recruitment domain family, member 6 [Source:MGI Symbol;Acc:MGI:3032959] |
| ENSMUSG00000035835 | Plppr3 | -1.26088 | 133.1391 | 0.365981 | 3.445217 | 0.000571 | 0.003885 | 0.004706 | 59.23411 | 97.37162 | 185.6341 | 190.3164 | 216152 | phospholipid phosphatase related 3 [Source:MGI Symbol;Acc:MGI:2388640] |
| ENSMUSG00000039323 | Igfbp2 | -1.25973 | 817.5046 | 0.243435 | 5.174824 | 2.28E-07 | 3.44E-06 | 4.17E-06 | 448.4868 | 514.3975 | 850.4421 | 1456.692 | 16008 | insulin-like growth factor binding protein 2 [Source:MGI Symbol;Acc:MGI:96437] |
| ENSMUSG00000033857 | Engase | -1.25299 | 609.4363 | 0.217247 | 5.76758 | 8.04E-09 | 1.59E-07 | 1.92E-07 | 299.3438 | 420.9601 | 951.032 | 766.4093 | 217364 | endo-beta-N-acetylglucosaminidase [Source:MGI Symbol;Acc:MGI:2443788] |
| ENSMUSG00000025997 | Ikzf2 | -1.25263 | 176.2222 | 0.288093 | 4.347986 | 1.37E-05 | 0.000143 | 0.000174 | 94.13992 | 114.092 | 240.5014 | 256.1556 | 22779 | IKAROS family zinc finger 2 [Source:MGI Symbol;Acc:MGI:1342541] |
| ENSMUSG00000040537 | Adam22 | -1.24437 | 196.5513 | 0.39501 | 3.150222 | 0.001631 | 0.009449 | 0.011446 | 69.81163 | 163.2696 | 274.3361 | 278.7878 | 11496 | a disintegrin and metallopeptidase domain 22 [Source:MGI Symbol;Acc:MGI:1340046] |
| ENSMUSG00000038151 | Prdm1 | -1.24367 | 278.6067 | 0.262256 | 4.742205 | 2.11E-06 | 2.6E-05 | 3.14E-05 | 156.5473 | 174.0886 | 330.1178 | 453.6732 | 12142 | PR domain containing 1, with ZNF domain [Source:MGI Symbol;Acc:MGI:99655] |
| ENSMUSG00000054939 | Zfp174 | -1.24209 | 69.43022 | 0.438833 | 2.830448 | 0.004648 | 0.02259 | 0.027313 | 40.19457 | 42.29272 | 92.35984 | 102.8737 | 385674 | zinc finger protein 174 [Source:MGI Symbol;Acc:MGI:2686600] |
| ENSMUSG00000039670 | Oxld1 | -1.23978 | 67.85102 | 0.449425 | 2.758598 | 0.005805 | 0.027159 | 0.032808 | 43.36783 | 37.37496 | 87.78757 | 102.8737 | 66431 | oxidoreductase like domain containing 1 [Source:MGI Symbol;Acc:MGI:1913681] |
| ENSMUSG00000133244 | Gm70605 | -1.23587 | 106.9946 | 0.386403 | 3.198394 | 0.001382 | 0.00823 | 0.00997 | 64.52287 | 62.94731 | 122.5368 | 177.9716 | | predicted gene, 70605 [Source:MGI Symbol;Acc:MGI:7824005] |
| ENSMUSG00000016918 | Sulf1 | -1.23447 | 309.6241 | 0.357313 | 3.454856 | 0.000551 | 0.003771 | 0.004568 | 248.5717 | 120.9769 | 348.4069 | 520.5411 | 240725 | sulfatase 1 [Source:MGI Symbol;Acc:MGI:2138563] |
| ENSMUSG00000099041 | Gm28035 | -1.23428 | 189.1199 | 0.319256 | 3.866123 | 0.000111 | 0.000908 | 0.0011 | 86.73566 | 138.608 | 269.7639 | 261.2993 | | predicted gene, 28035 [Source:MGI Symbol;Acc:MGI:5547771] |
| ENSMUSG00000121474 | | -1.23272 | 396.2173 | 0.282016 | 4.371089 | 1.24E-05 | 0.000131 | 0.000158 | 242.2252 | 231.1346 | 704.1294 | 407.38 | 627984 | NLR family, pyrin domain containing 1C, pseudogene [Source:NCBI gene (formerly Entrezgene);Acc:627984] |
| ENSMUSG00000099843 | Gm7160 | -1.23226 | 94.64108 | 0.377513 | 3.264163 | 0.001098 | 0.006793 | 0.008229 | 55.0031 | 58.02955 | 138.997 | 126.5347 | | predicted gene 7160 [Source:MGI Symbol;Acc:MGI:3646078] |
| ENSMUSG00000028268 | Gbp3 | -1.23178 | 208.553 | 0.302067 | 4.07782 | 4.55E-05 | 0.000419 | 0.000508 | 98.37093 | 150.4834 | 279.8229 | 305.535 | 55932 | guanylate binding protein 3 [Source:MGI Symbol;Acc:MGI:1926263] |
| ENSMUSG00000027583 | Zbtb46 | -1.22845 | 140.2854 | 0.326267 | 3.765151 | 0.000166 | 0.001306 | 0.001582 | 77.21589 | 90.48676 | 177.404 | 216.0348 | 72147 | zinc finger and BTB domain containing 46 [Source:MGI Symbol;Acc:MGI:1919397] |
| ENSMUSG00000022665 | Ccdc80 | -1.22661 | 110.5735 | 0.444117 | 2.76191 | 0.005746 | 0.026918 | 0.032517 | 47.59884 | 84.58545 | 119.7935 | 190.3164 | 67896 | coiled-coil domain containing 80 [Source:MGI Symbol;Acc:MGI:1915146] |
| ENSMUSG00000027399 | Il1a | -1.22604 | 282.5706 | 0.329897 | 3.716419 | 0.000202 | 0.001554 | 0.001882 | 167.1248 | 171.138 | 272.5072 | 519.5124 | 16175 | interleukin 1 alpha [Source:MGI Symbol;Acc:MGI:96542] |
| ENSMUSG00000031714 | Gab1 | -1.22437 | 643.0036 | 0.2073 | 5.906299 | 3.5E-09 | 7.31E-08 | 8.85E-08 | 313.0946 | 457.3515 | 919.9405 | 881.6279 | 14388 | growth factor receptor bound protein 2-associated protein 1 [Source:MGI Symbol;Acc:MGI:108088] |
| ENSMUSG00000024164 | C3 | -1.22403 | 1825.509 | 0.419348 | 2.918896 | 0.003513 | 0.017874 | 0.02163 | 775.3322 | 1413.364 | 2588.819 | 2524.521 | 12266 | complement component 3 [Source:MGI Symbol;Acc:MGI:88227] |
| ENSMUSG00000022548 | Apod | -1.22303 | 374.8384 | 0.300607 | 4.068532 | 4.73E-05 | 0.000433 | 0.000525 | 208.3771 | 240.9702 | 373.0972 | 676.9092 | 11815 | apolipoprotein D [Source:MGI Symbol;Acc:MGI:88056] |
| ENSMUSG00000037347 | Chst7 | -1.22231 | 208.0951 | 0.260405 | 4.693902 | 2.68E-06 | 3.22E-05 | 3.9E-05 | 123.757 | 125.8946 | 276.1651 | 306.5637 | 60322 | carbohydrate sulfotransferase 7 [Source:MGI Symbol;Acc:MGI:1891767] |
| ENSMUSG00000025020 | Slit1 | -1.2198 | 182.5263 | 0.300182 | 4.063538 | 4.83E-05 | 0.000441 | 0.000534 | 97.31318 | 121.9604 | 283.4807 | 227.351 | 20562 | slit guidance ligand 1 [Source:MGI Symbol;Acc:MGI:1315203] |
| ENSMUSG00000001986 | Gria3 | -1.21824 | 77.25706 | 0.434202 | 2.805703 | 0.005021 | 0.024122 | 0.029156 | 49.71434 | 43.27627 | 123.4513 | 92.58636 | 53623 | glutamate receptor, ionotropic, AMPA3 (alpha 3) [Source:MGI Symbol;Acc:MGI:95810] |
| ENSMUSG00000030352 | Tspan9 | -1.21714 | 323.1448 | 0.240259 | 5.065969 | 4.06E-07 | 5.81E-06 | 7.03E-06 | 167.1248 | 221.2991 | 420.6488 | 483.5066 | 109246 | tetraspanin 9 [Source:MGI Symbol;Acc:MGI:1924558] |
| ENSMUSG00000103165 | Gm37560 | -1.21513 | 143.9902 | 0.350506 | 3.466789 | 0.000527 | 0.003627 | 0.004394 | 103.6597 | 69.83217 | 179.2329 | 223.236 | | predicted gene, 37560 [Source:MGI Symbol;Acc:MGI:5610788] |
| ENSMUSG00000020135 | Apc2 | -1.2095 | 90.44308 | 0.433756 | 2.788434 | 0.005296 | 0.025223 | 0.030477 | 40.19457 | 68.84862 | 117.9645 | 134.7646 | 23805 | APC regulator of WNT signaling pathway 2 [Source:MGI Symbol;Acc:MGI:1346052] |
| ENSMUSG00000060548 | Tnfrsf19 | -1.20861 | 125.9689 | 0.431523 | 2.800797 | 0.005098 | 0.024445 | 0.029537 | 48.65659 | 103.2729 | 155.4571 | 196.4888 | 29820 | tumor necrosis factor receptor superfamily, member 19 [Source:MGI Symbol;Acc:MGI:1352474] |
| ENSMUSG00000070436 | Serpinh1 | -1.20748 | 594.7089 | 0.255883 | 4.718869 | 2.37E-06 | 2.88E-05 | 3.48E-05 | 388.195 | 330.4734 | 619.0852 | 1041.082 | 12406 | serine (or cysteine) peptidase inhibitor, clade H, member 1 [Source:MGI Symbol;Acc:MGI:88283] |
| ENSMUSG00000049281 | Scn3b | -1.20537 | 100.1501 | 0.366863 | 3.28562 | 0.001018 | 0.006386 | 0.007736 | 58.17636 | 62.94731 | 134.4247 | 145.052 | 235281 | sodium channel, voltage-gated, type III, beta [Source:MGI Symbol;Acc:MGI:1918882] |
| ENSMUSG00000029561 | Oasl2 | -1.20176 | 400.7976 | 0.192449 | 6.244573 | 4.25E-10 | 1.01E-08 | 1.22E-08 | 234.8209 | 250.8057 | 540.4422 | 577.1217 | 23962 | 2'-5' oligoadenylate synthetase-like 2 [Source:MGI Symbol;Acc:MGI:1344390] |
| ENSMUSG00000087624 | 9230111E07Rik | -1.19951 | 89.79119 | 0.386003 | 3.10751 | 0.001887 | 0.010639 | 0.012888 | 52.8876 | 56.06245 | 129.8524 | 120.3623 | 100048985 | RIKEN cDNA 9230111E07 gene [Source:MGI Symbol;Acc:MGI:1924998] |
| ENSMUSG00000033446 | Lpar6 | -1.19943 | 2100.281 | 0.102708 | 11.67807 | 1.65E-31 | 4.24E-29 | 5.14E-29 | 1289.4 | 1258.946 | 2812.86 | 3039.919 | 67168 | lysophosphatidic acid receptor 6 [Source:MGI Symbol;Acc:MGI:1914418] |
| ENSMUSG00000042810 | Krba1 | -1.19866 | 155.6131 | 0.301324 | 3.977991 | 6.95E-05 | 0.000605 | 0.000732 | 100.4864 | 88.51965 | 227.699 | 205.7475 | 77827 | KRAB-A domain containing 1 [Source:MGI Symbol;Acc:MGI:1925077] |
| ENSMUSG00000041773 | Enc1 | -1.19803 | 2212.77 | 0.105084 | 11.40067 | 4.15E-30 | 8.86E-28 | 1.07E-27 | 1306.324 | 1379.923 | 2941.798 | 3223.034 | 13803 | ectodermal-neural cortex 1 [Source:MGI Symbol;Acc:MGI:109610] |
| ENSMUSG00000003541 | Ier3 | -1.19719 | 3921.386 | 0.129896 | 9.216573 | 3.07E-20 | 2.41E-18 | 2.92E-18 | 2478.313 | 2284.791 | 4771.62 | 6150.821 | 15937 | immediate early response 3 [Source:MGI Symbol;Acc:MGI:104814] |
| ENSMUSG00000042207 | Kdm5b | -1.19649 | 993.7498 | 0.138914 | 8.613218 | 7.1E-18 | 4.1E-16 | 4.96E-16 | 594.4566 | 612.7527 | 1296.696 | 1471.094 | 75605 | lysine demethylase 5B [Source:MGI Symbol;Acc:MGI:1922855] |
| ENSMUSG00000023349 | Clec4n | -1.19473 | 1387.662 | 0.174521 | 6.845775 | 7.61E-12 | 2.3E-10 | 2.78E-10 | 744.6573 | 942.2425 | 1672.536 | 2191.211 | 56620 | C-type lectin domain family 4, member n [Source:MGI Symbol;Acc:MGI:1861231] |
| ENSMUSG00000032549 | Rab6b | -1.193 | 147.4048 | 0.359065 | 3.322524 | 0.000892 | 0.005698 | 0.006902 | 87.79341 | 91.47031 | 157.2861 | 253.0694 | 270192 | RAB6B, member RAS oncogene family [Source:MGI Symbol;Acc:MGI:107283] |
| ENSMUSG00000027589 | Pcmtd2 | -1.19207 | 758.678 | 0.171847 | 6.936827 | 4.01E-12 | 1.27E-10 | 1.54E-10 | 404.0612 | 519.3153 | 1034.247 | 1077.088 | 245867 | protein-L-isoaspartate (D-aspartate) O-methyltransferase domain containing 2 [Source:MGI Symbol;Acc:MGI:1923927] |
| ENSMUSG00000038175 | Mylip | -1.18801 | 2711.671 | 0.125905 | 9.435793 | 3.88E-21 | 3.23E-19 | 3.92E-19 | 1542.202 | 1765.475 | 3429.202 | 4109.806 | 218203 | myosin regulatory light chain interacting protein [Source:MGI Symbol;Acc:MGI:2388271] |
| ENSMUSG00000104030 | 5330406M23Rik | -1.18593 | 109.3918 | 0.398998 | 2.972272 | 0.002956 | 0.015435 | 0.018691 | 69.81163 | 63.93086 | 186.5486 | 117.2761 | | RIKEN cDNA 5330406M23 gene [Source:MGI Symbol;Acc:MGI:1923921] |
| ENSMUSG00000020034 | Tcp11l2 | -1.18553 | 352.8268 | 0.327031 | 3.625144 | 0.000289 | 0.002134 | 0.002585 | 138.5655 | 292.1149 | 508.4363 | 472.1904 | 216198 | t-complex 11 (mouse) like 2 [Source:MGI Symbol;Acc:MGI:2444679] |
| ENSMUSG00000029761 | Cald1 | -1.18493 | 271.8151 | 0.27184 | 4.358928 | 1.31E-05 | 0.000137 | 0.000166 | 176.6446 | 155.4012 | 311.8288 | 443.3858 | 109624 | caldesmon 1 [Source:MGI Symbol;Acc:MGI:88250] |
| ENSMUSG00000038390 | Gpr162 | -1.18125 | 445.2132 | 0.207875 | 5.68249 | 1.33E-08 | 2.54E-07 | 3.08E-07 | 234.8209 | 309.8188 | 611.7696 | 624.4436 | 14788 | G protein-coupled receptor 162 [Source:MGI Symbol;Acc:MGI:1315214] |
| ENSMUSG00000057137 | Tmem140 | -1.18093 | 1474.095 | 0.155023 | 7.617775 | 2.58E-14 | 1.02E-12 | 1.24E-12 | 773.2166 | 1030.762 | 1983.45 | 2108.912 | 68487 | transmembrane protein 140 [Source:MGI Symbol;Acc:MGI:1915737] |
| ENSMUSG00000037972 | Snn | -1.17908 | 1061.983 | 0.141355 | 8.341249 | 7.35E-17 | 3.83E-15 | 4.64E-15 | 596.5721 | 704.223 | 1433.864 | 1513.273 | 20621 | stannin [Source:MGI Symbol;Acc:MGI:1276549] |
| ENSMUSG00000062901 | Klhl24 | -1.17755 | 3726.616 | 0.120425 | 9.778333 | 1.39E-22 | 1.37E-20 | 1.66E-20 | 2023.479 | 2545.432 | 5072.475 | 5265.078 | 75785 | kelch-like 24 [Source:MGI Symbol;Acc:MGI:1923035] |
| ENSMUSG00000053411 | Cbx7 | -1.17722 | 156.5469 | 0.310674 | 3.789256 | 0.000151 | 0.0012 | 0.001453 | 84.62015 | 107.2071 | 203.9232 | 230.4372 | 52609 | chromobox 7 [Source:MGI Symbol;Acc:MGI:1196439] |
| ENSMUSG00000030691 | Fchsd2 | -1.17514 | 1851.588 | 0.10442 | 11.25398 | 2.21E-29 | 4.25E-27 | 5.15E-27 | 1109.582 | 1163.542 | 2627.226 | 2506.004 | 207278 | FCH and double SH3 domains 2 [Source:MGI Symbol;Acc:MGI:2448475] |
| ENSMUSG00000053835 | H2-T24 | -1.17408 | 260.2687 | 0.253034 | 4.640027 | 3.48E-06 | 4.09E-05 | 4.96E-05 | 137.5077 | 181.9571 | 358.4659 | 363.1443 | 15042 | histocompatibility 2, T region locus 24 [Source:MGI Symbol;Acc:MGI:95958] |
| ENSMUSG00000024042 | Sik1 | -1.17272 | 151.5837 | 0.328363 | 3.571426 | 0.000355 | 0.002568 | 0.003111 | 87.79341 | 98.35517 | 177.404 | 242.782 | 17691 | salt inducible kinase 1 [Source:MGI Symbol;Acc:MGI:104754] |
| ENSMUSG00000033863 | Klf9 | -1.16899 | 100.642 | 0.387832 | 3.014169 | 0.002577 | 0.013758 | 0.01666 | 53.94535 | 69.83217 | 123.4513 | 155.3393 | 16601 | Kruppel-like transcription factor 9 [Source:MGI Symbol;Acc:MGI:1333856] |
| ENSMUSG00000076617 | Ighm | -1.16523 | 229.4381 | 0.25115 | 4.639566 | 3.49E-06 | 4.09E-05 | 4.96E-05 | 152.3163 | 130.8124 | 313.6577 | 320.9661 | | immunoglobulin heavy constant mu [Source:MGI Symbol;Acc:MGI:96448] |
| ENSMUSG00000001249 | Hpn | -1.1645 | 236.9594 | 0.24533 | 4.746673 | 2.07E-06 | 2.55E-05 | 3.08E-05 | 142.7965 | 149.4999 | 310.9143 | 344.627 | 15451 | hepsin [Source:MGI Symbol;Acc:MGI:1196620] |
| ENSMUSG00000032216 | Nedd4 | -1.16404 | 1636.98 | 0.332043 | 3.505684 | 0.000455 | 0.00319 | 0.003864 | 798.6027 | 1221.571 | 2039.232 | 2488.516 | 17999 | neural precursor cell expressed, developmentally down-regulated 4 [Source:MGI Symbol;Acc:MGI:97297] |
| ENSMUSG00000030313 | Dennd5b | -1.15839 | 72.96051 | 0.440439 | 2.630068 | 0.008537 | 0.037444 | 0.045227 | 38.07907 | 52.12824 | 98.76101 | 102.8737 | 320560 | DENN domain containing 5B [Source:MGI Symbol;Acc:MGI:2444273] |
| ENSMUSG00000021589 | Rhobtb3 | -1.15684 | 148.658 | 0.350585 | 3.299736 | 0.000968 | 0.006102 | 0.007391 | 69.81163 | 114.092 | 203.9232 | 206.7762 | 73296 | Rho-related BTB domain containing 3 [Source:MGI Symbol;Acc:MGI:1920546] |
| ENSMUSG00000021200 | Asb2 | -1.15601 | 190.7887 | 0.283241 | 4.081371 | 4.48E-05 | 0.000414 | 0.000502 | 133.2767 | 103.2729 | 264.2772 | 262.328 | 65256 | ankyrin repeat and SOCS box-containing 2 [Source:MGI Symbol;Acc:MGI:1929743] |
| ENSMUSG00000026360 | Rgs2 | -1.15434 | 2743.248 | 0.098452 | 11.72487 | 9.5E-32 | 2.48E-29 | 3.01E-29 | 1613.072 | 1788.097 | 3833.39 | 3738.432 | 19735 | regulator of G-protein signaling 2 [Source:MGI Symbol;Acc:MGI:1098271] |
| ENSMUSG00000120054 | Gm57214 | -1.14569 | 134.9971 | 0.371045 | 3.087737 | 0.002017 | 0.011194 | 0.013559 | 80.38915 | 87.5361 | 142.6548 | 229.4084 | | predicted gene, 57214 [Source:MGI Symbol;Acc:MGI:7338341] |
| ENSMUSG00000021281 | Tnfaip2 | -1.13853 | 2131.697 | 0.165979 | 6.859472 | 6.91E-12 | 2.1E-10 | 2.54E-10 | 1235.454 | 1427.134 | 2441.592 | 3422.609 | 21928 | tumor necrosis factor, alpha-induced protein 2 [Source:MGI Symbol;Acc:MGI:104960] |
| ENSMUSG00000028270 | Gbp2 | -1.13775 | 278.9042 | 0.264292 | 4.30488 | 1.67E-05 | 0.000171 | 0.000207 | 141.7388 | 206.5459 | 371.2683 | 396.0639 | 14469 | guanylate binding protein 2 [Source:MGI Symbol;Acc:MGI:102772] |
| ENSMUSG00000121122 | Gm56904 | -1.1366 | 131.7887 | 0.352519 | 3.224239 | 0.001263 | 0.007639 | 0.009254 | 92.02442 | 72.78283 | 154.5427 | 207.8049 | 102640901 | predicted gene, 56904 [Source:MGI Symbol;Acc:MGI:7337721] |
| ENSMUSG00000027966 | Col11a1 | -1.13571 | 292.8458 | 0.349623 | 3.248371 | 0.001161 | 0.007102 | 0.008603 | 115.295 | 250.8057 | 429.7933 | 375.4891 | 12814 | collagen, type XI, alpha 1 [Source:MGI Symbol;Acc:MGI:88446] |
| ENSMUSG00000039831 | Arhgap29 | -1.13567 | 354.0124 | 0.2315 | 4.905705 | 9.31E-07 | 1.23E-05 | 1.49E-05 | 226.3589 | 216.3814 | 418.8198 | 554.4894 | 214137 | Rho GTPase activating protein 29 [Source:MGI Symbol;Acc:MGI:2443818] |
| ENSMUSG00000024975 | Pdcd4 | -1.13451 | 230.4431 | 0.296322 | 3.828641 | 0.000129 | 0.001043 | 0.001263 | 112.1217 | 176.0558 | 305.4276 | 328.1672 | 18569 | programmed cell death 4 [Source:MGI Symbol;Acc:MGI:107490] |
| ENSMUSG00000038065 | Mturn | -1.13225 | 201.8173 | 0.366138 | 3.092423 | 0.001985 | 0.011084 | 0.013426 | 82.50465 | 170.1544 | 287.1385 | 267.4717 | 68235 | maturin, neural progenitor differentiation regulator homolog (Xenopus) [Source:MGI Symbol;Acc:MGI:1915485] |
| ENSMUSG00000032531 | Amotl2 | -1.1303 | 131.6454 | 0.358285 | 3.154756 | 0.001606 | 0.009332 | 0.011305 | 65.58062 | 99.33872 | 168.2595 | 193.4026 | 56332 | angiomotin-like 2 [Source:MGI Symbol;Acc:MGI:1929286] |
| ENSMUSG00000038119 | Cdon | -1.13007 | 103.2571 | 0.385536 | 2.931152 | 0.003377 | 0.01728 | 0.020918 | 64.52287 | 64.91441 | 117.9645 | 165.6267 | 57810 | cell adhesion molecule-related/down-regulated by oncogenes [Source:MGI Symbol;Acc:MGI:1926387] |
| ENSMUSG00000020423 | Btg2 | -1.12809 | 6196.725 | 0.099484 | 11.33945 | 8.37E-30 | 1.74E-27 | 2.11E-27 | 3944.357 | 3835.852 | 7806.692 | 9199.998 | 12227 | BTG anti-proliferation factor 2 [Source:MGI Symbol;Acc:MGI:108384] |
| ENSMUSG00000026764 | Kif5c | -1.12745 | 257.5612 | 0.236225 | 4.772764 | 1.82E-06 | 2.26E-05 | 2.74E-05 | 151.2585 | 172.1215 | 352.9792 | 353.8856 | 16574 | kinesin family member 5C [Source:MGI Symbol;Acc:MGI:1098269] |
| ENSMUSG00000055322 | Tns1 | -1.12406 | 194.7998 | 0.275596 | 4.078642 | 4.53E-05 | 0.000418 | 0.000506 | 114.2372 | 130.8124 | 286.224 | 247.9257 | 21961 | tensin 1 [Source:MGI Symbol;Acc:MGI:104552] |
| ENSMUSG00000027204 | Fbn1 | -1.12184 | 274.2439 | 0.37663 | 2.978638 | 0.002895 | 0.015139 | 0.018332 | 244.3407 | 101.3058 | 359.3804 | 391.9489 | 14118 | fibrillin 1 [Source:MGI Symbol;Acc:MGI:95489] |
| ENSMUSG00000038267 | Slc22a23 | -1.12092 | 335.8902 | 0.360043 | 3.113282 | 0.00185 | 0.010464 | 0.012676 | 124.8147 | 298.0162 | 436.1945 | 484.5353 | 73102 | solute carrier family 22, member 23 [Source:MGI Symbol;Acc:MGI:1920352] |
| ENSMUSG00000018339 | Gpx3 | -1.1208 | 398.1617 | 0.189022 | 5.929451 | 3.04E-09 | 6.37E-08 | 7.72E-08 | 253.8605 | 247.855 | 547.7578 | 543.1733 | 14778 | glutathione peroxidase 3 [Source:MGI Symbol;Acc:MGI:105102] |
| ENSMUSG00000028698 | Pik3r3 | -1.11209 | 138.0582 | 0.362553 | 3.06739 | 0.002159 | 0.011885 | 0.014397 | 90.96666 | 83.60189 | 147.2271 | 230.4372 | 18710 | phosphoinositide-3-kinase regulatory subunit 3 [Source:MGI Symbol;Acc:MGI:109277] |
| ENSMUSG00000002985 | Apoe | -1.10767 | 348931.5 | 0.19975 | 5.545284 | 2.93E-08 | 5.34E-07 | 6.46E-07 | 196167.5 | 246221.3 | 435606.5 | 517730.6 | 11816 | apolipoprotein E [Source:MGI Symbol;Acc:MGI:88057] |
| ENSMUSG00000074794 | Arrdc3 | -1.10569 | 1002.878 | 0.160977 | 6.868642 | 6.48E-12 | 1.99E-10 | 2.41E-10 | 553.2043 | 718.9763 | 1376.253 | 1363.077 | 105171 | arrestin domain containing 3 [Source:MGI Symbol;Acc:MGI:2145242] |
| ENSMUSG00000103706 | 6820402A03Rik | -1.09984 | 234.9251 | 0.283182 | 3.88387 | 0.000103 | 0.000853 | 0.001033 | 151.2585 | 147.5328 | 263.3627 | 377.5466 | | RIKEN cDNA 6820402A03 gene [Source:MGI Symbol;Acc:MGI:1922992] |
| ENSMUSG00000075122 | Cd80 | -1.09876 | 167.2839 | 0.289937 | 3.78964 | 0.000151 | 0.001199 | 0.001452 | 106.8329 | 106.2236 | 244.1592 | 211.9199 | 12519 | CD80 antigen [Source:MGI Symbol;Acc:MGI:101775] |
| ENSMUSG00000032232 | Cgnl1 | -1.09493 | 147.7813 | 0.333943 | 3.278785 | 0.001043 | 0.006508 | 0.007884 | 107.8907 | 80.65124 | 178.3185 | 224.2647 | 68178 | cingulin-like 1 [Source:MGI Symbol;Acc:MGI:1915428] |
| ENSMUSG00000024049 | Myom1 | -1.09351 | 220.5244 | 0.264854 | 4.128733 | 3.65E-05 | 0.000345 | 0.000418 | 127.988 | 153.4341 | 321.8877 | 278.7878 | 17929 | myomesin 1 [Source:MGI Symbol;Acc:MGI:1341430] |
| ENSMUSG00000048078 | Tenm4 | -1.09296 | 75.08987 | 0.41793 | 2.615179 | 0.008918 | 0.038769 | 0.046829 | 48.65659 | 47.21048 | 100.5899 | 103.9025 | 23966 | teneurin transmembrane protein 4 [Source:MGI Symbol;Acc:MGI:2447063] |
| ENSMUSG00000090272 | Mndal | -1.09207 | 408.5668 | 0.191916 | 5.690334 | 1.27E-08 | 2.44E-07 | 2.96E-07 | 276.0732 | 245.8879 | 549.5867 | 562.7193 | 100040462 | myeloid nuclear differentiation antigen like [Source:MGI Symbol;Acc:MGI:3780953] |
| ENSMUSG00000022994 | Adcy6 | -1.09147 | 115.8562 | 0.375454 | 2.90707 | 0.003648 | 0.018446 | 0.022315 | 60.29186 | 87.5361 | 141.7403 | 173.8566 | 11512 | adenylate cyclase 6 [Source:MGI Symbol;Acc:MGI:87917] |
| ENSMUSG00000053040 | Aph1c | -1.09051 | 865.2265 | 0.136365 | 7.997016 | 1.27E-15 | 5.84E-14 | 7.07E-14 | 536.2802 | 569.4764 | 1182.389 | 1172.761 | 68318 | aph1 homolog C, gamma secretase subunit [Source:MGI Symbol;Acc:MGI:1915568] |
| ENSMUSG00000020883 | Fbxl20 | -1.09019 | 1232.179 | 0.117081 | 9.311419 | 1.26E-20 | 1.02E-18 | 1.23E-18 | 777.4477 | 797.6604 | 1688.082 | 1665.526 | 72194 | F-box and leucine-rich repeat protein 20 [Source:MGI Symbol;Acc:MGI:1919444] |
| ENSMUSG00000037709 | Fam13a | -1.08626 | 128.8078 | 0.366939 | 2.96032 | 0.003073 | 0.015942 | 0.019305 | 63.46511 | 101.3058 | 167.345 | 183.1152 | 58909 | family with sequence similarity 13, member A [Source:MGI Symbol;Acc:MGI:1889842] |
| ENSMUSG00000067818 | Myl9 | -1.08442 | 98.90738 | 0.380766 | 2.847994 | 0.0044 | 0.021554 | 0.02606 | 68.75387 | 58.02955 | 120.7079 | 148.1382 | 98932 | myosin, light polypeptide 9, regulatory [Source:MGI Symbol;Acc:MGI:2138915] |
| ENSMUSG00000023961 | Enpp4 | -1.08135 | 379.6167 | 0.197452 | 5.476516 | 4.34E-08 | 7.62E-07 | 9.23E-07 | 243.2829 | 243.9208 | 490.1472 | 541.1158 | 224794 | ectonucleotide pyrophosphatase/phosphodiesterase 4 [Source:MGI Symbol;Acc:MGI:2682634] |
| ENSMUSG00000038058 | Nod1 | -1.08118 | 269.6315 | 0.228121 | 4.739517 | 2.14E-06 | 2.63E-05 | 3.18E-05 | 166.0671 | 179.99 | 362.1237 | 370.3454 | 107607 | nucleotide-binding oligomerization domain containing 1 [Source:MGI Symbol;Acc:MGI:1341839] |
| ENSMUSG00000028389 | Zfp37 | -1.07966 | 84.21858 | 0.400769 | 2.693957 | 0.007061 | 0.03196 | 0.038611 | 57.1186 | 51.14469 | 121.6224 | 106.9887 | 22696 | zinc finger protein 37 [Source:MGI Symbol;Acc:MGI:99181] |
| ENSMUSG00000069206 | Zfp874a | -1.07622 | 222.8194 | 0.26995 | 3.986714 | 6.7E-05 | 0.000586 | 0.00071 | 130.1035 | 156.3847 | 272.5072 | 332.2822 | 238692 | zinc finger protein 874a [Source:MGI Symbol;Acc:MGI:3040703] |
| ENSMUSG00000024421 | Lama3 | -1.07395 | 857.1251 | 0.195665 | 5.488713 | 4.05E-08 | 7.17E-07 | 8.69E-07 | 451.6601 | 652.0948 | 1228.111 | 1096.634 | 16774 | laminin, alpha 3 [Source:MGI Symbol;Acc:MGI:99909] |
| ENSMUSG00000039763 | Dnajc28 | -1.07315 | 129.3906 | 0.327507 | 3.276735 | 0.00105 | 0.006545 | 0.007928 | 76.15814 | 90.48676 | 171.9173 | 179.0003 | 246738 | DnaJ heat shock protein family (Hsp40) member C28 [Source:MGI Symbol;Acc:MGI:2181053] |
| ENSMUSG00000059839 | Zfp874b | -1.07264 | 235.2821 | 0.243876 | 4.398295 | 1.09E-05 | 0.000117 | 0.000141 | 143.8543 | 159.3354 | 320.0588 | 317.8798 | 408067 | zinc finger protein 874b [Source:MGI Symbol;Acc:MGI:3040702] |
| ENSMUSG00000043432 | Lrrc9 | -1.06796 | 81.86378 | 0.404608 | 2.6395 | 0.008303 | 0.036576 | 0.044179 | 48.65659 | 57.046 | 110.6489 | 111.1036 | 243813 | leukocyte receptor cluster (LRC) member 9 [Source:MGI Symbol;Acc:MGI:2444509] |
| ENSMUSG00000020604 | Arsg | -1.06299 | 146.7678 | 0.322316 | 3.297976 | 0.000974 | 0.006137 | 0.007435 | 84.62015 | 105.24 | 180.1474 | 217.0636 | 74008 | arylsulfatase G [Source:MGI Symbol;Acc:MGI:1921258] |
| ENSMUSG00000095041 | | -1.06182 | 3662.68 | 0.088897 | 11.94445 | 6.94E-33 | 2.06E-30 | 2.5E-30 | 2269.936 | 2474.616 | 4968.228 | 4937.939 | | |
| ENSMUSG00000025993 | Slc40a1 | -1.06 | 6085.846 | 0.0752 | 14.09567 | 4.04E-45 | 4.71E-42 | 5.71E-42 | 3884.065 | 4006.99 | 8381.884 | 8070.445 | 53945 | solute carrier family 40 (iron-regulated transporter), member 1 [Source:MGI Symbol;Acc:MGI:1315204] |
| ENSMUSG00000028211 | Trp53inp1 | -1.05982 | 1734.414 | 0.153835 | 6.8893 | 5.61E-12 | 1.74E-10 | 2.1E-10 | 965.7275 | 1282.551 | 2211.149 | 2478.228 | 60599 | transformation related protein 53 inducible nuclear protein 1 [Source:MGI Symbol;Acc:MGI:1926609] |
| ENSMUSG00000026604 | Ptpn14 | -1.05652 | 111.1078 | 0.354689 | 2.978727 | 0.002894 | 0.015139 | 0.018332 | 65.58062 | 78.68414 | 158.2005 | 141.9658 | 19250 | protein tyrosine phosphatase, non-receptor type 14 [Source:MGI Symbol;Acc:MGI:102467] |
| ENSMUSG00000051373 | Plpp7 | -1.05471 | 114.3558 | 0.344802 | 3.058885 | 0.002222 | 0.012161 | 0.014732 | 70.86938 | 77.70058 | 146.3126 | 162.5405 | 227721 | phospholipid phosphatase 7 (inactive) [Source:MGI Symbol;Acc:MGI:2445183] |
| ENSMUSG00000030577 | Cd22 | -1.04954 | 350.7104 | 0.22947 | 4.573762 | 4.79E-06 | 5.49E-05 | 6.65E-05 | 199.9151 | 256.707 | 438.0234 | 508.1963 | 12483 | CD22 antigen [Source:MGI Symbol;Acc:MGI:88322] |
| ENSMUSG00000019539 | Rcn3 | -1.04856 | 130.9084 | 0.399504 | 2.624663 | 0.008673 | 0.037934 | 0.04582 | 74.04263 | 96.38807 | 128.938 | 224.2647 | 52377 | reticulocalbin 3, EF-hand calcium binding domain [Source:MGI Symbol;Acc:MGI:1277122] |
| ENSMUSG00000106847 | Peg13 | -1.04783 | 171.8868 | 0.311379 | 3.365123 | 0.000765 | 0.004985 | 0.006039 | 124.8147 | 99.33872 | 202.0943 | 261.2993 | 353342 | paternally expressed 13 [Source:MGI Symbol;Acc:MGI:2663476] |
| ENSMUSG00000090086 | AI480526 | -1.04668 | 82.77266 | 0.398955 | 2.623547 | 0.008702 | 0.038025 | 0.04593 | 52.8876 | 55.07889 | 107.9056 | 115.2186 | 100675 | expressed sequence AI480526 [Source:MGI Symbol;Acc:MGI:2140910] |
| ENSMUSG00000078920 | Ifi47 | -1.03615 | 266.9798 | 0.231814 | 4.46976 | 7.83E-06 | 8.62E-05 | 0.000104 | 182.9911 | 167.2038 | 373.0972 | 344.627 | 15953 | interferon gamma inducible protein 47 [Source:MGI Symbol;Acc:MGI:99448] |
| ENSMUSG00000054404 | Slfn5 | -1.03544 | 2126.069 | 0.099645 | 10.39132 | 2.72E-25 | 3.61E-23 | 4.38E-23 | 1356.038 | 1432.051 | 2802.801 | 2913.384 | 327978 | schlafen 5 [Source:MGI Symbol;Acc:MGI:1329004] |
| ENSMUSG00000034652 | Cd300a | -1.03233 | 2744.101 | 0.103069 | 10.01585 | 1.3E-23 | 1.42E-21 | 1.72E-21 | 1694.519 | 1909.074 | 3590.146 | 3782.667 | 217303 | CD300A molecule [Source:MGI Symbol;Acc:MGI:2443411] |
| ENSMUSG00000023951 | Vegfa | -1.03127 | 97.78185 | 0.373183 | 2.763434 | 0.00572 | 0.026809 | 0.032386 | 64.52287 | 63.93086 | 120.7079 | 141.9658 | 22339 | vascular endothelial growth factor A [Source:MGI Symbol;Acc:MGI:103178] |
| ENSMUSG00000021322 | Aoah | -1.02984 | 2481.439 | 0.091029 | 11.31328 | 1.13E-29 | 2.25E-27 | 2.73E-27 | 1620.476 | 1642.531 | 3320.382 | 3342.368 | 27052 | acyloxyacyl hydrolase [Source:MGI Symbol;Acc:MGI:1350928] |
| ENSMUSG00000026674 | Ddr2 | -1.02959 | 187.2702 | 0.325224 | 3.1658 | 0.001547 | 0.009051 | 0.010964 | 147.0275 | 99.33872 | 216.7256 | 285.989 | 18214 | discoidin domain receptor family, member 2 [Source:MGI Symbol;Acc:MGI:1345277] |
| ENSMUSG00000022108 | Itm2b | -1.02798 | 40300.86 | 0.073615 | 13.9642 | 2.58E-44 | 2.06E-41 | 2.49E-41 | 25389.22 | 27651.57 | 52524.4 | 55638.23 | 16432 | integral membrane protein 2B [Source:MGI Symbol;Acc:MGI:1309517] |
| ENSMUSG00000045551 | Fpr1 | -1.02782 | 268.8097 | 0.236337 | 4.348949 | 1.37E-05 | 0.000143 | 0.000173 | 161.836 | 191.7926 | 350.2358 | 371.3742 | 14293 | formyl peptide receptor 1 [Source:MGI Symbol;Acc:MGI:107443] |
| ENSMUSG00000042978 | Sbk1 | -1.02473 | 172.047 | 0.32225 | 3.17992 | 0.001473 | 0.008682 | 0.010517 | 89.90891 | 136.7137 | 241.4158 | 220.1498 | 104175 | SH3-binding kinase 1 [Source:MGI Symbol;Acc:MGI:2135937] |
| ENSMUSG00000021127 | Zfp36l1 | -1.0232 | 2008.428 | 0.121799 | 8.400768 | 4.44E-17 | 2.38E-15 | 2.88E-15 | 1445.947 | 1203.867 | 2654.659 | 2729.24 | 12192 | zinc finger protein 36, C3H type-like 1 [Source:MGI Symbol;Acc:MGI:107946] |
| ENSMUSG00000103593 | Gm37352 | -1.01597 | 152.8998 | 0.306204 | 3.317946 | 0.000907 | 0.005775 | 0.006996 | 108.9484 | 93.43741 | 191.1208 | 218.0923 | | predicted gene, 37352 [Source:MGI Symbol;Acc:MGI:5610580] |
| ENSMUSG00000020589 | Cyria | -1.01408 | 639.1116 | 0.158109 | 6.413779 | 1.42E-10 | 3.64E-09 | 4.4E-09 | 400.888 | 445.5489 | 862.33 | 847.6796 | 76820 | CYFIP related Rac1 interactor A [Source:MGI Symbol;Acc:MGI:1261783] |
| ENSMUSG00000030148 | Clec4a2 | -1.01329 | 572.3146 | 0.164831 | 6.147475 | 7.87E-10 | 1.8E-08 | 2.17E-08 | 377.6174 | 380.6345 | 730.6486 | 800.3577 | 26888 | C-type lectin domain family 4, member a2 [Source:MGI Symbol;Acc:MGI:1349412] |
| ENSMUSG00000056749 | Nfil3 | -1.01132 | 200.1465 | 0.3557 | 2.843175 | 0.004467 | 0.021855 | 0.026423 | 136.45 | 128.8453 | 186.5486 | 348.742 | 18030 | nuclear factor, interleukin 3, regulated [Source:MGI Symbol;Acc:MGI:109495] |
| ENSMUSG00000044447 | Dock5 | -1.01073 | 203.4443 | 0.311054 | 3.24937 | 0.001157 | 0.00708 | 0.008577 | 161.836 | 108.1907 | 242.3303 | 301.42 | 68813 | dedicator of cytokinesis 5 [Source:MGI Symbol;Acc:MGI:2652871] |
| ENSMUSG00000058254 | Tspan7 | -1.00999 | 472.9405 | 0.295668 | 3.415954 | 0.000636 | 0.004259 | 0.005159 | 220.0124 | 407.1904 | 541.3567 | 723.2024 | 21912 | tetraspanin 7 [Source:MGI Symbol;Acc:MGI:1298407] |
| ENSMUSG00000027002 | Nckap1 | -1.00309 | 355.9233 | 0.322561 | 3.109756 | 0.001872 | 0.01057 | 0.012804 | 159.7205 | 313.753 | 414.2476 | 535.9722 | 50884 | NCK-associated protein 1 [Source:MGI Symbol;Acc:MGI:1355333] |
| ENSMUSG00000038352 | Arl5c | -0.99804 | 504.1311 | 0.180911 | 5.516747 | 3.45E-08 | 6.19E-07 | 7.5E-07 | 355.4046 | 317.6872 | 708.7017 | 634.7309 | 217151 | ADP-ribosylation factor-like 5C [Source:MGI Symbol;Acc:MGI:3028577] |
| ENSMUSG00000032089 | Il10ra | -0.99522 | 3814.429 | 0.094935 | 10.48309 | 1.03E-25 | 1.46E-23 | 1.77E-23 | 2392.635 | 2703.784 | 5050.528 | 5110.767 | 16154 | interleukin 10 receptor, alpha [Source:MGI Symbol;Acc:MGI:96538] |
| ENSMUSG00000009621 | Vav2 | -0.99448 | 1734.375 | 0.112069 | 8.87379 | 7.07E-19 | 4.58E-17 | 5.55E-17 | 1102.177 | 1215.67 | 2250.471 | 2369.182 | 22325 | vav 2 oncogene [Source:MGI Symbol;Acc:MGI:102718] |
| ENSMUSG00000026821 | Ralgds | -0.99407 | 1270.408 | 0.145769 | 6.819471 | 9.14E-12 | 2.72E-10 | 3.3E-10 | 760.5236 | 937.3248 | 1611.268 | 1772.514 | 19730 | ral guanine nucleotide dissociation stimulator [Source:MGI Symbol;Acc:MGI:107485] |
| ENSMUSG00000098557 | Kctd12 | -0.98624 | 18845.32 | 0.076051 | 12.96815 | 1.85E-38 | 8.79E-36 | 1.06E-35 | 12408.49 | 12877.64 | 23848.96 | 26246.18 | 239217 | potassium channel tetramerisation domain containing 12 [Source:MGI Symbol;Acc:MGI:2145823] |
| ENSMUSG00000070737 | Tmem35b | -0.98482 | 157.2908 | 0.308273 | 3.194649 | 0.0014 | 0.008318 | 0.010077 | 92.02442 | 119.0098 | 202.0943 | 216.0348 | 100039968 | transmembrane protein 35B [Source:MGI Symbol;Acc:MGI:3758095] |
| ENSMUSG00000015243 | Abca1 | -0.98224 | 51123.72 | 0.087533 | 11.22136 | 3.2E-29 | 5.92E-27 | 7.17E-27 | 31614.09 | 37110.39 | 67254.42 | 68515.97 | 11303 | ATP-binding cassette, sub-family A member 1 [Source:MGI Symbol;Acc:MGI:99607] |
| ENSMUSG00000048058 | Ldlrad3 | -0.98172 | 174.6798 | 0.32384 | 3.031501 | 0.002433 | 0.013125 | 0.0159 | 92.02442 | 142.615 | 221.2978 | 242.782 | 241576 | low density lipoprotein receptor class A domain containing 3 [Source:MGI Symbol;Acc:MGI:2138856] |
| ENSMUSG00000076431 | Sox4 | -0.98139 | 1418.046 | 0.142253 | 6.898915 | 5.24E-12 | 1.63E-10 | 1.98E-10 | 848.317 | 1058.302 | 1947.787 | 1817.779 | 20677 | SRY (sex determining region Y)-box 4 [Source:MGI Symbol;Acc:MGI:98366] |
| ENSMUSG00000020638 | Cmpk2 | -0.98093 | 303.1935 | 0.227476 | 4.312247 | 1.62E-05 | 0.000166 | 0.000201 | 191.4531 | 216.3814 | 432.5367 | 372.4029 | 22169 | cytidine/uridine monophosphate kinase 2 [Source:MGI Symbol;Acc:MGI:99830] |
| ENSMUSG00000059336 | Slc14a1 | -0.98038 | 127.1854 | 0.359273 | 2.728788 | 0.006357 | 0.029253 | 0.035339 | 67.69612 | 103.2729 | 170.0884 | 167.6842 | 108052 | solute carrier family 14 (urea transporter), member 1 [Source:MGI Symbol;Acc:MGI:1351654] |
| ENSMUSG00000021846 | Peli2;Peli2 | -0.97964 | 608.1981 | 0.179293 | 5.463925 | 4.66E-08 | 8.15E-07 | 9.88E-07 | 364.9244 | 453.4173 | 836.7252 | 777.7254 | 93834;624367 | pellino 2 [Source:MGI Symbol;Acc:MGI:1891445];pellino 2 [Source:MGI Symbol;Acc:MGI:1891445] |
| ENSMUSG00000067928 | Zfp760 | -0.9773 | 189.5909 | 0.272589 | 3.585244 | 0.000337 | 0.002449 | 0.002967 | 137.5077 | 118.0262 | 248.7314 | 254.0981 | 240034 | zinc finger protein 760 [Source:MGI Symbol;Acc:MGI:2679257] |
| ENSMUSG00000002900 | Lamb1 | -0.9765 | 162.2032 | 0.309526 | 3.154832 | 0.001606 | 0.009332 | 0.011305 | 115.295 | 103.2729 | 187.463 | 242.782 | 16777 | laminin B1 [Source:MGI Symbol;Acc:MGI:96743] |
| ENSMUSG00000035064 | Eef2k | -0.97594 | 1290.723 | 0.133043 | 7.335533 | 2.21E-13 | 7.99E-12 | 9.68E-12 | 797.5449 | 942.2425 | 1749.35 | 1673.756 | 13631 | eukaryotic elongation factor-2 kinase [Source:MGI Symbol;Acc:MGI:1195261] |
| ENSMUSG00000009828 | Cilk1 | -0.97516 | 190.1845 | 0.274434 | 3.553342 | 0.00038 | 0.002726 | 0.003302 | 123.757 | 132.7795 | 270.6783 | 233.5234 | 56542 | ciliogenesis associated kinase 1 [Source:MGI Symbol;Acc:MGI:1934157] |
| ENSMUSG00000026235 | Epha4 | -0.97246 | 134.1206 | 0.373051 | 2.606765 | 0.00914 | 0.039554 | 0.047778 | 68.75387 | 112.1249 | 159.115 | 196.4888 | 13838 | Eph receptor A4 [Source:MGI Symbol;Acc:MGI:98277] |
| ENSMUSG00000121069 | Gm56730 | -0.97235 | 115.3982 | 0.338321 | 2.874034 | 0.004053 | 0.020221 | 0.024454 | 76.15814 | 79.66769 | 154.5427 | 151.2244 | | predicted gene, 56730 [Source:MGI Symbol;Acc:MGI:7337373] |
| ENSMUSG00000047747 | Rnf150 | -0.97202 | 1916.291 | 0.101912 | 9.537882 | 1.46E-21 | 1.26E-19 | 1.52E-19 | 1268.245 | 1319.926 | 2591.562 | 2485.429 | 330812 | ring finger protein 150 [Source:MGI Symbol;Acc:MGI:2443860] |
| ENSMUSG00000023055 | Calcoco1 | -0.97069 | 316.0289 | 0.210907 | 4.602444 | 4.18E-06 | 4.82E-05 | 5.84E-05 | 214.7236 | 212.4472 | 431.6222 | 405.3225 | 67488 | calcium binding and coiled coil domain 1 [Source:MGI Symbol;Acc:MGI:1914738] |
| ENSMUSG00000031216 | Stard8 | -0.96516 | 4712.807 | 0.105385 | 9.15845 | 5.26E-20 | 3.93E-18 | 4.76E-18 | 2904.587 | 3479.806 | 6144.215 | 6322.62 | 236920 | StAR related lipid transfer domain containing 8 [Source:MGI Symbol;Acc:MGI:2448556] |
| ENSMUSG00000032717 | Mdfi | -0.9651 | 204.4569 | 0.279972 | 3.447127 | 0.000567 | 0.003861 | 0.004677 | 118.4682 | 158.3518 | 264.2772 | 276.7303 | 17240 | MyoD family inhibitor [Source:MGI Symbol;Acc:MGI:107687] |
| ENSMUSG00000035273 | Hpse | -0.96401 | 673.5194 | 0.155054 | 6.217209 | 5.06E-10 | 1.19E-08 | 1.44E-08 | 434.736 | 478.0061 | 867.8167 | 913.5188 | 15442 | heparanase [Source:MGI Symbol;Acc:MGI:1343124] |
| ENSMUSG00000024030 | Abcg1 | -0.96071 | 2993.206 | 0.148245 | 6.480566 | 9.14E-11 | 2.38E-09 | 2.88E-09 | 1734.713 | 2328.067 | 3662.388 | 4247.657 | 11307 | ATP binding cassette subfamily G member 1 [Source:MGI Symbol;Acc:MGI:107704] |
| ENSMUSG00000048621 | Gm6377 | -0.9603 | 1467.948 | 0.149627 | 6.417949 | 1.38E-10 | 3.55E-09 | 4.3E-09 | 924.4752 | 1068.137 | 1730.147 | 2149.032 | 622976 | predicted gene 6377 [Source:MGI Symbol;Acc:MGI:3647255] |
| ENSMUSG00000041633 | Kctd12b | -0.95851 | 358.2211 | 0.215075 | 4.456636 | 8.33E-06 | 9.13E-05 | 0.000111 | 223.1857 | 263.5919 | 504.7785 | 441.3283 | 207474 | potassium channel tetramerisation domain containing 12b [Source:MGI Symbol;Acc:MGI:2444667] |
| ENSMUSG00000047945 | Marcksl1 | -0.95818 | 4476.242 | 0.126368 | 7.582464 | 3.39E-14 | 1.33E-12 | 1.61E-12 | 3250.472 | 2833.612 | 5254.452 | 6566.431 | 17357 | MARCKS-like 1 [Source:MGI Symbol;Acc:MGI:97143] |
| ENSMUSG00000053560 | Ier2 | -0.9539 | 3095.234 | 0.107797 | 8.849042 | 8.83E-19 | 5.58E-17 | 6.76E-17 | 2121.85 | 2092.998 | 3762.063 | 4404.025 | 15936 | immediate early response 2 [Source:MGI Symbol;Acc:MGI:104815] |
| ENSMUSG00000029510 | Gpc2 | -0.95159 | 143.273 | 0.314445 | 3.02627 | 0.002476 | 0.013317 | 0.016127 | 107.8907 | 87.5361 | 188.3775 | 189.2877 | 71951 | glypican 2 cerebroglycan [Source:MGI Symbol;Acc:MGI:1919201] |
| ENSMUSG00000025558 | Dock9 | -0.94816 | 416.1105 | 0.1982 | 4.783846 | 1.72E-06 | 2.15E-05 | 2.61E-05 | 258.0915 | 309.8188 | 535.8699 | 560.6619 | 105445 | dedicator of cytokinesis 9 [Source:MGI Symbol;Acc:MGI:106321] |
| ENSMUSG00000029401 | Rilpl2 | -0.94641 | 804.7128 | 0.211116 | 4.48291 | 7.36E-06 | 8.14E-05 | 9.86E-05 | 454.8333 | 644.2264 | 925.4273 | 1194.364 | 80291 | Rab interacting lysosomal protein-like 2 [Source:MGI Symbol;Acc:MGI:1933112] |
| ENSMUSG00000003206 | Ebi3 | -0.94601 | 209.1313 | 0.259799 | 3.641325 | 0.000271 | 0.002019 | 0.002446 | 134.3345 | 151.467 | 285.3096 | 265.4142 | 50498 | Epstein-Barr virus induced gene 3 [Source:MGI Symbol;Acc:MGI:1354171] |
| ENSMUSG00000043664 | Tmem221 | -0.94587 | 159.5588 | 0.290661 | 3.254209 | 0.001137 | 0.00698 | 0.008456 | 107.8907 | 110.1578 | 202.0943 | 218.0923 | 434325 | transmembrane protein 221 [Source:MGI Symbol;Acc:MGI:3525074] |
| ENSMUSG00000032652 | Crebl2 | -0.94466 | 193.3431 | 0.274995 | 3.435204 | 0.000592 | 0.004006 | 0.004853 | 121.6415 | 142.615 | 239.5869 | 269.5292 | 232430 | cAMP responsive element binding protein-like 2 [Source:MGI Symbol;Acc:MGI:1889385] |
| ENSMUSG00000029512 | Ulk1 | -0.937 | 450.6473 | 0.219106 | 4.276476 | 1.9E-05 | 0.000191 | 0.000231 | 278.1888 | 340.3089 | 666.6368 | 517.4549 | 22241 | unc-51 like kinase 1 [Source:MGI Symbol;Acc:MGI:1270126] |
| ENSMUSG00000058318 | Phf21a | -0.93155 | 319.4943 | 0.22737 | 4.097092 | 4.18E-05 | 0.000391 | 0.000473 | 210.4926 | 229.1675 | 461.7992 | 376.5179 | 192285 | PHD finger protein 21A [Source:MGI Symbol;Acc:MGI:2384756] |
| ENSMUSG00000021767 | Kat6b | -0.93001 | 226.3796 | 0.270311 | 3.440508 | 0.000581 | 0.003941 | 0.004774 | 170.2981 | 141.6314 | 329.2034 | 264.3855 | 54169 | K(lysine) acetyltransferase 6B [Source:MGI Symbol;Acc:MGI:1858746] |
| ENSMUSG00000026012 | Cd28 | -0.92466 | 1308.031 | 0.149902 | 6.168416 | 6.9E-10 | 1.59E-08 | 1.92E-08 | 867.3566 | 937.3248 | 1517.079 | 1910.365 | 12487 | CD28 antigen [Source:MGI Symbol;Acc:MGI:88327] |
| ENSMUSG00000103103 | 4833445I07Rik | -0.92434 | 179.5028 | 0.273566 | 3.378854 | 0.000728 | 0.004788 | 0.0058 | 124.8147 | 122.944 | 229.5279 | 240.7245 | | RIKEN cDNA 4833445I07 gene [Source:MGI Symbol;Acc:MGI:2143268] |
| ENSMUSG00000038393 | Txnip | -0.91965 | 6523.102 | 0.30795 | 2.986362 | 0.002823 | 0.014838 | 0.017968 | 4337.841 | 4685.64 | 10329.67 | 6739.258 | 56338 | thioredoxin interacting protein [Source:MGI Symbol;Acc:MGI:1889549] |
| ENSMUSG00000066975 | Cryba4 | -0.90845 | 188.1846 | 0.28648 | 3.171078 | 0.001519 | 0.008912 | 0.010796 | 123.757 | 137.6972 | 217.64 | 273.6441 | 12959 | crystallin, beta A4 [Source:MGI Symbol;Acc:MGI:102716] |
| ENSMUSG00000028957 | Per3 | -0.90701 | 334.3473 | 0.222308 | 4.079948 | 4.5E-05 | 0.000416 | 0.000504 | 207.3194 | 257.6905 | 452.6546 | 419.7248 | 18628 | period circadian clock 3 [Source:MGI Symbol;Acc:MGI:1277134] |
| ENSMUSG00000027499 | Pkia | -0.90665 | 175.3009 | 0.341406 | 2.655626 | 0.007916 | 0.035169 | 0.042478 | 94.13992 | 149.4999 | 199.3509 | 258.2131 | 18767 | protein kinase inhibitor, alpha [Source:MGI Symbol;Acc:MGI:104747] |
| ENSMUSG00000040746 | Rnf167 | -0.90628 | 584.8105 | 0.178372 | 5.080844 | 3.76E-07 | 5.4E-06 | 6.54E-06 | 369.1554 | 444.5654 | 799.2326 | 726.2886 | 70510 | ring finger protein 167 [Source:MGI Symbol;Acc:MGI:1917760] |
| ENSMUSG00000038178 | Slc43a2 | -0.90452 | 4498.921 | 0.076733 | 11.7879 | 4.51E-32 | 1.27E-29 | 1.53E-29 | 3139.408 | 3126.711 | 5869.879 | 5859.688 | 215113 | solute carrier family 43, member 2 [Source:MGI Symbol;Acc:MGI:2442746] |
| ENSMUSG00000014158 | Trpv4 | -0.89937 | 172.6859 | 0.321234 | 2.799731 | 0.005115 | 0.024503 | 0.029607 | 96.25542 | 144.5821 | 211.2388 | 238.6671 | 63873 | transient receptor potential cation channel, subfamily V, member 4 [Source:MGI Symbol;Acc:MGI:1926945] |
| ENSMUSG00000030083 | Abtb1 | -0.8983 | 595.9001 | 0.172633 | 5.203531 | 1.96E-07 | 3E-06 | 3.64E-06 | 377.6174 | 454.4009 | 780.0291 | 771.553 | 80283 | ankyrin repeat and BTB domain containing 1 [Source:MGI Symbol;Acc:MGI:1933148] |
| ENSMUSG00000120996 | Gm56817 | -0.89663 | 241.1298 | 0.238326 | 3.762194 | 0.000168 | 0.001321 | 0.0016 | 162.8938 | 174.0886 | 312.7432 | 314.7936 | | predicted gene, 56817 [Source:MGI Symbol;Acc:MGI:7337547] |
| ENSMUSG00000097451 | Rian | -0.89626 | 241.2162 | 0.307571 | 2.913994 | 0.003568 | 0.018084 | 0.021884 | 133.2767 | 203.5952 | 267.935 | 360.0581 | | RNA imprinted and accumulated in nucleus [Source:MGI Symbol;Acc:MGI:1922995] |
| ENSMUSG00000022419 | Deptor | -0.89338 | 169.1744 | 0.289267 | 3.088418 | 0.002012 | 0.011178 | 0.013541 | 107.8907 | 128.8453 | 224.9556 | 215.0061 | 97998 | DEP domain containing MTOR-interacting protein [Source:MGI Symbol;Acc:MGI:2146322] |
| ENSMUSG00000134402 | | -0.88554 | 232.8799 | 0.242756 | 3.647846 | 0.000264 | 0.001977 | 0.002395 | 161.836 | 165.2367 | 291.7108 | 312.7362 | 631323 | predicted gene 12250 [Source:NCBI gene (formerly Entrezgene);Acc:631323] |
| ENSMUSG00000050212 | Eva1b | -0.88548 | 155.7298 | 0.295065 | 3.000972 | 0.002691 | 0.01424 | 0.017244 | 103.6597 | 115.0755 | 206.6666 | 197.5176 | 230752 | eva-1 homolog B [Source:MGI Symbol;Acc:MGI:1922063] |
| ENSMUSG00000031129 | Slc9a9 | -0.88542 | 5182.893 | 0.087232 | 10.15013 | 3.31E-24 | 3.86E-22 | 4.68E-22 | 3476.831 | 3803.394 | 6531.029 | 6920.316 | 331004 | solute carrier family 9 (sodium/hydrogen exchanger), member 9 [Source:MGI Symbol;Acc:MGI:2679732] |
| ENSMUSG00000020023 | Tmcc3 | -0.88514 | 1089.343 | 0.129595 | 6.829998 | 8.49E-12 | 2.54E-10 | 3.08E-10 | 739.3686 | 790.7756 | 1356.135 | 1471.094 | 319880 | transmembrane and coiled coil domains 3 [Source:MGI Symbol;Acc:MGI:2442900] |
| ENSMUSG00000079484 | Phyhd1 | -0.88504 | 981.348 | 0.140759 | 6.287614 | 3.22E-10 | 7.73E-09 | 9.37E-09 | 637.8244 | 740.6144 | 1252.802 | 1294.152 | 227696 | phytanoyl-CoA dioxygenase domain containing 1 [Source:MGI Symbol;Acc:MGI:3612860] |
| ENSMUSG00000007837 | Prrg2 | -0.88344 | 121.7496 | 0.328617 | 2.688373 | 0.00718 | 0.032402 | 0.039146 | 84.62015 | 86.55255 | 156.3716 | 159.4543 | 65116 | proline-rich Gla (G-carboxyglutamic acid) polypeptide 2 [Source:MGI Symbol;Acc:MGI:1929596] |
| ENSMUSG00000034265 | Zdhhc14 | -0.88147 | 843.4294 | 0.140087 | 6.29229 | 3.13E-10 | 7.52E-09 | 9.11E-09 | 615.6116 | 571.4435 | 1065.339 | 1121.324 | 224454 | zinc finger, DHHC domain containing 14 [Source:MGI Symbol;Acc:MGI:2653229] |
| ENSMUSG00000079481 | Nhsl2 | -0.88118 | 356.9152 | 0.216757 | 4.065288 | 4.8E-05 | 0.000438 | 0.000531 | 280.3043 | 222.2827 | 459.0558 | 466.018 | 100042480 | NHS like 2 [Source:MGI Symbol;Acc:MGI:3645090] |
| ENSMUSG00000109245 | Gm44860 | -0.87965 | 184.2999 | 0.31039 | 2.834009 | 0.004597 | 0.022383 | 0.027062 | 155.4895 | 104.2565 | 229.5279 | 247.9257 | | predicted gene 44860 [Source:MGI Symbol;Acc:MGI:5753436] |
| ENSMUSG00000047180 | Neurl3 | -0.87859 | 1145.986 | 0.122601 | 7.166286 | 7.71E-13 | 2.61E-11 | 3.16E-11 | 781.6787 | 833.0683 | 1504.277 | 1464.922 | 214854 | neuralized E3 ubiquitin protein ligase 3 [Source:MGI Symbol;Acc:MGI:2429944] |
| ENSMUSG00000060591 | Ifitm2 | -0.87529 | 1016.494 | 0.154598 | 5.661691 | 1.5E-08 | 2.84E-07 | 3.44E-07 | 686.481 | 747.4993 | 1181.474 | 1450.52 | 80876 | interferon induced transmembrane protein 2 [Source:MGI Symbol;Acc:MGI:1933382] |
| ENSMUSG00000056592 | Zfp175 | -0.87446 | 347.4119 | 0.228162 | 3.83264 | 0.000127 | 0.001028 | 0.001245 | 278.1888 | 212.4472 | 428.8788 | 470.133 | 210104 | zinc finger protein 175 [Source:MGI Symbol;Acc:MGI:2652821] |
| ENSMUSG00000000184 | Ccnd2 | -0.8737 | 392.5441 | 0.188933 | 4.624379 | 3.76E-06 | 4.39E-05 | 5.31E-05 | 275.0155 | 279.3287 | 506.6074 | 509.225 | 12444 | cyclin D2 [Source:MGI Symbol;Acc:MGI:88314] |
| ENSMUSG00000036833 | Pnpla7 | -0.87294 | 4876.42 | 0.118985 | 7.336595 | 2.19E-13 | 7.95E-12 | 9.63E-12 | 3046.326 | 3841.753 | 6114.953 | 6502.649 | 241274 | patatin-like phospholipase domain containing 7 [Source:MGI Symbol;Acc:MGI:2385325] |
| ENSMUSG00000027514 | Zbp1 | -0.87123 | 171.0411 | 0.295064 | 2.952693 | 0.00315 | 0.016297 | 0.019735 | 110.0062 | 131.7959 | 238.6724 | 203.69 | 58203 | Z-DNA binding protein 1 [Source:MGI Symbol;Acc:MGI:1927449] |
| ENSMUSG00000030539 | Sema4b | -0.86833 | 337.4039 | 0.236519 | 3.671281 | 0.000241 | 0.001822 | 0.002208 | 200.9729 | 276.378 | 428.8788 | 443.3858 | 20352 | sema domain, immunoglobulin domain (Ig), transmembrane domain (TM) and short cytoplasmic domain, (semaphorin) 4B [Source:MGI Symbol;Acc:MGI:107559] |
| ENSMUSG00000109005 | Gm45221 | -0.86726 | 203.0744 | 0.264802 | 3.275121 | 0.001056 | 0.006571 | 0.00796 | 148.0853 | 139.6643 | 280.7373 | 243.8108 | | predicted gene 45221 [Source:MGI Symbol;Acc:MGI:5753797] |
| ENSMUSG00000005397 | Nid1 | -0.86619 | 206.2861 | 0.293821 | 2.948013 | 0.003198 | 0.016517 | 0.019995 | 173.4713 | 119.0098 | 256.9615 | 275.7016 | 18073 | nidogen 1 [Source:MGI Symbol;Acc:MGI:97342] |
| ENSMUSG00000042834 | Nrep | -0.8654 | 365.6435 | 0.232009 | 3.730026 | 0.000191 | 0.001479 | 0.001792 | 218.9546 | 298.9997 | 450.8257 | 493.7939 | 27528 | neuronal regeneration related protein [Source:MGI Symbol;Acc:MGI:99444] |
| ENSMUSG00000048895 | Cdk5r1 | -0.86299 | 197.4613 | 0.264579 | 3.261743 | 0.001107 | 0.006831 | 0.008275 | 139.6233 | 140.6479 | 268.8494 | 240.7245 | 12569 | cyclin dependent kinase 5, regulatory subunit 1 [Source:MGI Symbol;Acc:MGI:101764] |
| ENSMUSG00000040433 | Zbtb38 | -0.86144 | 1367.984 | 0.112551 | 7.65378 | 1.95E-14 | 7.87E-13 | 9.53E-13 | 951.9767 | 990.4366 | 1762.153 | 1767.371 | 245007 | zinc finger and BTB domain containing 38 [Source:MGI Symbol;Acc:MGI:2442866] |
| ENSMUSG00000023495 | Pcbp4 | -0.86114 | 155.3372 | 0.301936 | 2.852051 | 0.004344 | 0.021357 | 0.025821 | 100.4864 | 119.9933 | 192.0353 | 208.8337 | 59092 | poly(rC) binding protein 4 [Source:MGI Symbol;Acc:MGI:1890471] |
| ENSMUSG00000039908 | Slc26a11 | -0.85982 | 669.064 | 0.172088 | 4.996364 | 5.84E-07 | 8.06E-06 | 9.76E-06 | 424.1585 | 526.2002 | 842.212 | 883.6854 | 268512 | solute carrier family 26, member 11 [Source:MGI Symbol;Acc:MGI:2444589] |
| ENSMUSG00000046402 | Rbp1 | -0.85975 | 127.8332 | 0.322411 | 2.666645 | 0.007661 | 0.034258 | 0.041389 | 94.13992 | 87.5361 | 160.9439 | 168.7129 | 19659 | retinol binding protein 1, cellular [Source:MGI Symbol;Acc:MGI:97876] |
| ENSMUSG00000023044 | Csad | -0.85972 | 589.2795 | 0.174223 | 4.93457 | 8.03E-07 | 1.08E-05 | 1.31E-05 | 379.7329 | 457.3515 | 744.3654 | 775.668 | 246277 | cysteine sulfinic acid decarboxylase [Source:MGI Symbol;Acc:MGI:2180098] |
| ENSMUSG00000032554 | Trf | -0.85881 | 24340.64 | 0.096057 | 8.940623 | 3.87E-19 | 2.6E-17 | 3.15E-17 | 16016.48 | 18587.16 | 29561.55 | 33197.35 | 22041 | transferrin [Source:MGI Symbol;Acc:MGI:98821] |
| ENSMUSG00000058402 | Zfp420 | -0.85855 | 128.4791 | 0.324258 | 2.647738 | 0.008103 | 0.035812 | 0.043255 | 94.13992 | 88.51965 | 156.3716 | 174.8853 | 233058 | zinc finger protein 420 [Source:MGI Symbol;Acc:MGI:2444666] |
| ENSMUSG00000057143 | Trim12c | -0.85671 | 750.3138 | 0.147558 | 5.805909 | 6.4E-09 | 1.28E-07 | 1.55E-07 | 508.7787 | 558.6574 | 948.2886 | 985.5304 | 319236 | tripartite motif-containing 12C [Source:MGI Symbol;Acc:MGI:4821183] |
| ENSMUSG00000040414 | Slc25a28 | -0.85637 | 517.1108 | 0.254033 | 3.371102 | 0.000749 | 0.0049 | 0.005935 | 393.4837 | 342.276 | 506.6074 | 826.0761 | 246696 | solute carrier family 25, member 28 [Source:MGI Symbol;Acc:MGI:2180509] |
| ENSMUSG00000030201 | Lrp6 | -0.85603 | 9685.147 | 0.089354 | 9.580208 | 9.68E-22 | 8.74E-20 | 1.06E-19 | 6396.226 | 7389.424 | 12562.77 | 12392.17 | 16974 | low density lipoprotein receptor-related protein 6 [Source:MGI Symbol;Acc:MGI:1298218] |
| ENSMUSG00000026042 | Col5a2 | -0.84813 | 442.1798 | 0.299443 | 2.832374 | 0.00462 | 0.022469 | 0.027166 | 396.657 | 235.0689 | 450.8257 | 686.1678 | 12832 | collagen, type V, alpha 2 [Source:MGI Symbol;Acc:MGI:88458] |
| ENSMUSG00000038068 | Rnf144b | -0.84637 | 637.2078 | 0.164286 | 5.151812 | 2.58E-07 | 3.85E-06 | 4.67E-06 | 419.9275 | 490.7923 | 828.4952 | 809.6163 | 218215 | ring finger protein 144B [Source:MGI Symbol;Acc:MGI:2384986] |
| ENSMUSG00000039007 | Cpq | -0.84563 | 608.1019 | 0.167617 | 5.045027 | 4.53E-07 | 6.44E-06 | 7.8E-06 | 404.0612 | 465.22 | 753.5099 | 809.6163 | 54381 | carboxypeptidase Q [Source:MGI Symbol;Acc:MGI:1889205] |
| ENSMUSG00000027602 | Map1lc3a | -0.84429 | 163.5401 | 0.297789 | 2.835194 | 0.00458 | 0.022308 | 0.026971 | 108.9484 | 124.9111 | 192.9498 | 227.351 | 66734 | microtubule-associated protein 1 light chain 3 alpha [Source:MGI Symbol;Acc:MGI:1915661] |
| ENSMUSG00000038578 | Susd1 | -0.84336 | 785.9975 | 0.139749 | 6.034817 | 1.59E-09 | 3.48E-08 | 4.21E-08 | 561.6663 | 563.5751 | 1026.017 | 992.7315 | 634731 | sushi domain containing 1 [Source:MGI Symbol;Acc:MGI:3651543] |
| ENSMUSG00000130119 | Gm74436 | -0.84179 | 185.0551 | 0.312394 | 2.694636 | 0.007047 | 0.031923 | 0.038567 | 147.0275 | 118.0262 | 198.4365 | 276.7303 | | predicted gene, 74436 [Source:MGI Symbol;Acc:MGI:7831608] |
| ENSMUSG00000028756 | Pink1 | -0.84161 | 1507.153 | 0.19461 | 4.324613 | 1.53E-05 | 0.000158 | 0.000191 | 836.6818 | 1321.893 | 1893.834 | 1976.204 | 68943 | PTEN induced putative kinase 1 [Source:MGI Symbol;Acc:MGI:1916193] |
| ENSMUSG00000078350 | Smim1 | -0.83974 | 293.6718 | 0.22858 | 3.673743 | 0.000239 | 0.001808 | 0.00219 | 194.6264 | 226.2169 | 355.7225 | 398.1214 | 68859 | small integral membrane protein 1 [Source:MGI Symbol;Acc:MGI:1916109] |
| ENSMUSG00000026648 | Dclre1c | -0.83962 | 1113.567 | 0.124736 | 6.731223 | 1.68E-11 | 4.84E-10 | 5.86E-10 | 777.4477 | 819.2986 | 1467.698 | 1389.824 | 227525 | DNA cross-link repair 1C [Source:MGI Symbol;Acc:MGI:2441769] |
| ENSMUSG00000033083 | Tbc1d4 | -0.83841 | 2314.528 | 0.111374 | 7.527868 | 5.16E-14 | 2E-12 | 2.42E-12 | 1534.798 | 1785.146 | 2936.311 | 3001.856 | 210789 | TBC1 domain family, member 4 [Source:MGI Symbol;Acc:MGI:2429660] |
| ENSMUSG00000069516 | Lyz2 | -0.83583 | 180304.2 | 0.072234 | 11.57109 | 5.77E-31 | 1.37E-28 | 1.66E-28 | 123057.8 | 135917 | 234621.4 | 227620.5 | 17105 | lysozyme 2 [Source:MGI Symbol;Acc:MGI:96897] |
| ENSMUSG00000058163 | Gm5431 | -0.83449 | 1063.718 | 0.126309 | 6.60674 | 3.93E-11 | 1.08E-09 | 1.3E-09 | 779.5632 | 749.4664 | 1401.858 | 1323.985 | 432555 | predicted gene 5431 [Source:MGI Symbol;Acc:MGI:3645205] |
| ENSMUSG00000037235 | Mxd4 | -0.83436 | 1545.296 | 0.154597 | 5.397024 | 6.78E-08 | 1.15E-06 | 1.4E-06 | 946.688 | 1273.699 | 1974.306 | 1986.492 | 17122 | Max dimerization protein 4 [Source:MGI Symbol;Acc:MGI:104991] |
| ENSMUSG00000035713 | Usp35 | -0.83407 | 440.8557 | 0.200574 | 4.158433 | 3.2E-05 | 0.000307 | 0.000372 | 280.3043 | 353.0951 | 555.9879 | 574.0354 | 244144 | ubiquitin specific peptidase 35 [Source:MGI Symbol;Acc:MGI:2685339] |
| ENSMUSG00000029299 | Abcg3 | -0.83304 | 542.0329 | 0.185266 | 4.49647 | 6.91E-06 | 7.7E-05 | 9.32E-05 | 352.2314 | 426.8614 | 656.5778 | 732.461 | 27405 | ATP binding cassette subfamily G member 3 [Source:MGI Symbol;Acc:MGI:1351624] |
| ENSMUSG00000027381 | Bcl2l11 | -0.83235 | 846.2067 | 0.146756 | 5.671647 | 1.41E-08 | 2.69E-07 | 3.26E-07 | 608.2073 | 608.8185 | 1010.471 | 1157.33 | 12125 | BCL2-like 11 [Source:MGI Symbol;Acc:MGI:1197519] |
| ENSMUSG00000021690 | Jmy | -0.83118 | 543.8959 | 0.171162 | 4.856088 | 1.2E-06 | 1.55E-05 | 1.88E-05 | 373.3864 | 409.1575 | 665.7224 | 727.3173 | 57748 | junction-mediating and regulatory protein [Source:MGI Symbol;Acc:MGI:1913096] |
| ENSMUSG00000019194 | Scn1b | -0.82987 | 1091.003 | 0.140848 | 5.891945 | 3.82E-09 | 7.92E-08 | 9.59E-08 | 715.0403 | 855.69 | 1389.055 | 1404.226 | 20266 | sodium channel, voltage-gated, type I, beta [Source:MGI Symbol;Acc:MGI:98247] |
| ENSMUSG00000047996 | Prrg1 | -0.82922 | 207.5912 | 0.25914 | 3.199895 | 0.001375 | 0.008201 | 0.009934 | 139.6233 | 159.3354 | 267.0205 | 264.3855 | 546336 | proline rich Gla (G-carboxyglutamic acid) 1 [Source:MGI Symbol;Acc:MGI:1917364] |
| ENSMUSG00000078429 | Ctdsp2 | -0.8284 | 1296.199 | 0.148455 | 5.580146 | 2.4E-08 | 4.41E-07 | 5.34E-07 | 823.9887 | 1043.548 | 1715.515 | 1601.744 | 52468 | CTD small phosphatase 2 [Source:MGI Symbol;Acc:MGI:1098748] |
| ENSMUSG00000017493 | Igfbp4 | -0.82839 | 5934.787 | 0.08044 | 10.29827 | 7.17E-25 | 8.99E-23 | 1.09E-22 | 4109.366 | 4442.703 | 7689.642 | 7497.438 | 16010 | insulin-like growth factor binding protein 4 [Source:MGI Symbol;Acc:MGI:96439] |
| ENSMUSG00000038332 | Sesn1 | -0.8281 | 4244.422 | 0.111498 | 7.427057 | 1.11E-13 | 4.17E-12 | 5.05E-12 | 2757.559 | 3358.829 | 5310.233 | 5551.067 | 140742 | sestrin 1 [Source:MGI Symbol;Acc:MGI:2155278] |
| ENSMUSG00000000982 | Ccl3 | -0.82782 | 4117.309 | 0.262141 | 3.157924 | 0.001589 | 0.00926 | 0.011217 | 3182.776 | 2751.978 | 4385.721 | 6148.763 | 20302 | C-C motif chemokine ligand 3 [Source:MGI Symbol;Acc:MGI:98260] |
| ENSMUSG00000072893 | 4933439C10Rik | -0.82689 | 149.4884 | 0.303092 | 2.728192 | 0.006368 | 0.029294 | 0.03539 | 99.42868 | 116.0591 | 191.1208 | 191.3451 | 74476 | RIKEN cDNA 4933439C10 gene [Source:MGI Symbol;Acc:MGI:1921726] |
| ENSMUSG00000023367 | Tmem176a | -0.82451 | 2293.796 | 0.108447 | 7.602839 | 2.9E-14 | 1.14E-12 | 1.38E-12 | 1542.202 | 1768.426 | 2916.193 | 2948.361 | 66058 | transmembrane protein 176A [Source:MGI Symbol;Acc:MGI:1913308] |
| ENSMUSG00000031903 | Pla2g15 | -0.82445 | 3441.964 | 0.08903 | 9.260304 | 2.04E-20 | 1.64E-18 | 1.98E-18 | 2401.097 | 2567.07 | 4306.163 | 4493.525 | 192654 | phospholipase A2, group XV [Source:MGI Symbol;Acc:MGI:2178076] |
| ENSMUSG00000006920 | Ezh1 | -0.82431 | 1009.322 | 0.138348 | 5.958195 | 2.55E-09 | 5.4E-08 | 6.54E-08 | 675.9035 | 780.94 | 1305.84 | 1274.606 | 14055 | enhancer of zeste 1 polycomb repressive complex 2 subunit [Source:MGI Symbol;Acc:MGI:1097695] |
| ENSMUSG00000040209 | Zfp704 | -0.82277 | 3340.274 | 0.082983 | 9.914895 | 3.59E-23 | 3.75E-21 | 4.54E-21 | 2416.963 | 2408.718 | 4288.788 | 4246.628 | 170753 | zinc finger protein 704 [Source:MGI Symbol;Acc:MGI:2180715] |
| ENSMUSG00000040848 | Sft2d2 | -0.82156 | 3738.533 | 0.091076 | 9.020584 | 1.87E-19 | 1.32E-17 | 1.6E-17 | 2572.453 | 2830.662 | 4698.464 | 4852.554 | 108735 | SFT2 domain containing 2 [Source:MGI Symbol;Acc:MGI:1917362] |
| ENSMUSG00000057596 | Trim30d | -0.8212 | 848.0721 | 0.143795 | 5.710901 | 1.12E-08 | 2.18E-07 | 2.64E-07 | 621.9581 | 603.9007 | 1021.445 | 1144.985 | 209387 | tripartite motif-containing 30D [Source:MGI Symbol;Acc:MGI:3035181] |
| ENSMUSG00000032594 | Ip6k1 | -0.82111 | 3743.349 | 0.106217 | 7.730502 | 1.07E-14 | 4.46E-13 | 5.41E-13 | 2487.832 | 2923.116 | 4639.939 | 4922.508 | 27399 | inositol hexaphosphate kinase 1 [Source:MGI Symbol;Acc:MGI:1351633] |

| | | | | | | | | | | | | | |
|---|---|---|---|---|---|---|---|---|---|---|---|---|---|
| ENSMUSG00000037007 | Zfp113 | -0.81764 | 187.603 | 0.270529 | 3.022379 | 0.002508 | 0.013433 | 0.016266 | 127.988 | 143.5985 | 243.2447 | 235.5809 | 56314 | zinc finger protein 113 [Source:MGI Symbol;Acc:MGI:1929116] |
| ENSMUSG00000098206 | Mir181a-1hg | -0.81739 | 279.9283 | 0.226962 | 3.60142 | 0.000316 | 0.002315 | 0.002804 | 214.7236 | 190.809 | 368.5249 | 345.6558 | | Mir181a-1 and Mir181b-1 host gene [Source:MGI Symbol;Acc:MGI:2443823] |
| ENSMUSG00000003134 | Tbc1d8 | -0.81693 | 1045.062 | 0.128442 | 6.360329 | 2.01E-10 | 4.99E-09 | 6.04E-09 | 789.0829 | 724.8776 | 1343.333 | 1322.956 | 54610 | TBC1 domain family, member 8 [Source:MGI Symbol;Acc:MGI:1927225] |
| ENSMUSG00000047153 | Khnyn | -0.81666 | 742.6293 | 0.14835 | 5.504937 | 3.69E-08 | 6.58E-07 | 7.97E-07 | 514.0674 | 561.608 | 972.0644 | 922.7774 | 219094 | KH and NYN domain containing [Source:MGI Symbol;Acc:MGI:2451333] |
| ENSMUSG00000040152 | Thbs1 | -0.8158 | 827.0947 | 0.190078 | 4.291941 | 1.77E-05 | 0.000179 | 0.000217 | 606.0918 | 592.0981 | 878.7901 | 1231.399 | 21825 | thrombospondin 1 [Source:MGI Symbol;Acc:MGI:98737] |
| ENSMUSG00000136278 | Gm73818 | -0.81406 | 295.5947 | 0.232856 | 3.495974 | 0.000472 | 0.003296 | 0.003993 | 239.0519 | 189.8255 | 374.9261 | 378.5753 | | predicted gene, 73818 [Source:MGI Symbol;Acc:MGI:7830382] |
| ENSMUSG00000020709 | Adap2 | -0.8127 | 1365.304 | 0.118357 | 6.866568 | 6.58E-12 | 2E-10 | 2.43E-10 | 1034.481 | 947.1603 | 1774.955 | 1704.618 | 216991 | ArfGAP with dual PH domains 2 [Source:MGI Symbol;Acc:MGI:2663075] |
| ENSMUSG00000056708 | Ier5 | -0.81165 | 2249.643 | 0.12625 | 6.428893 | 1.29E-10 | 3.31E-09 | 4.02E-09 | 1641.631 | 1623.844 | 2568.701 | 3164.396 | 15939 | immediate early response 5 [Source:MGI Symbol;Acc:MGI:1337072] |
| ENSMUSG00000092341 | Malat1 | -0.81026 | 51410.72 | 0.072146 | 11.23091 | 2.88E-29 | 5.38E-27 | 6.52E-27 | 37678.18 | 37006.13 | 68640.73 | 62317.82 | | metastasis associated lung adenocarcinoma transcript 1 (non-coding RNA) [Source:MGI Symbol;Acc:MGI:1919539] |
| ENSMUSG00000057440 | Mpp7 | -0.80904 | 347.909 | 0.229087 | 3.531594 | 0.000413 | 0.002932 | 0.003552 | 250.6872 | 254.7399 | 383.1562 | 503.0526 | 75739 | membrane protein, palmitoylated 7 (MAGUK p55 subfamily member 7) [Source:MGI Symbol;Acc:MGI:1922989] |
| ENSMUSG00000083282 | Ctsf | -0.80833 | 1309.439 | 0.137558 | 5.876262 | 4.2E-09 | 8.61E-08 | 1.04E-07 | 857.8368 | 1045.515 | 1677.108 | 1657.296 | 56464 | cathepsin F [Source:MGI Symbol;Acc:MGI:1861434] |
| ENSMUSG00000056185 | Snx32 | -0.80777 | 211.8887 | 0.257807 | 3.133242 | 0.001729 | 0.009904 | 0.011997 | 142.7965 | 165.2367 | 267.935 | 271.5867 | 225861 | sorting nexin 32 [Source:MGI Symbol;Acc:MGI:2444704] |
| ENSMUSG00000026536 | Ifi211 | -0.8062 | 516.6853 | 0.170803 | 4.72007 | 2.36E-06 | 2.86E-05 | 3.47E-05 | 385.0217 | 366.8648 | 632.802 | 682.0529 | 381308 | interferon activated gene 211 [Source:MGI Symbol;Acc:MGI:3041120] |
| ENSMUSG00000039193 | Nlrc4 | -0.80608 | 421.1939 | 0.193919 | 4.156766 | 3.23E-05 | 0.000309 | 0.000374 | 281.362 | 331.4569 | 543.1856 | 528.771 | 268973 | NLR family, CARD domain containing 4 [Source:MGI Symbol;Acc:MGI:3036243] |
| ENSMUSG00000036611 | Eepd1 | -0.80575 | 378.0524 | 0.245521 | 3.281808 | 0.001031 | 0.006465 | 0.007832 | 225.3012 | 324.5721 | 440.7667 | 521.5698 | 67484 | endonuclease/exonuclease/phosphatase family domain containing 1 [Source:MGI Symbol;Acc:MGI:1914734] |
| ENSMUSG00000048546 | Tob2 | -0.80523 | 593.2592 | 0.249041 | 3.233336 | 0.001224 | 0.00743 | 0.009 | 491.8546 | 371.7825 | 590.7372 | 918.6625 | 57259 | transducer of ERBB2, 2 [Source:MGI Symbol;Acc:MGI:1888525] |
| ENSMUSG00000069874 | Irgm2 | -0.80198 | 565.1698 | 0.173324 | 4.627047 | 3.71E-06 | 4.34E-05 | 5.25E-05 | 422.043 | 402.2726 | 771.799 | 664.5643 | 54396 | immunity-related GTPase family M member 2 [Source:MGI Symbol;Acc:MGI:1926262] |
| ENSMUSG00000067212 | H2-T23 | -0.80139 | 1391.626 | 0.117619 | 6.813402 | 9.53E-12 | 2.82E-10 | 3.42E-10 | 967.843 | 1061.252 | 1740.206 | 1797.204 | 15040 | histocompatibility 2, T region locus 23 [Source:MGI Symbol;Acc:MGI:95957] |
| ENSMUSG00000049502 | Dtx3l | -0.80071 | 1495.004 | 0.119203 | 6.717189 | 1.85E-11 | 5.3E-10 | 6.42E-10 | 1090.542 | 1090.759 | 2011.798 | 1786.917 | 209200 | deltex 3-like, E3 ubiquitin ligase [Source:MGI Symbol;Acc:MGI:2656973] |
| ENSMUSG00000031385 | Plxnb3 | -0.80057 | 201.5671 | 0.289131 | 2.768892 | 0.005625 | 0.026454 | 0.031956 | 122.6992 | 171.138 | 245.9881 | 266.443 | 140571 | plexin B3 [Source:MGI Symbol;Acc:MGI:2154240] |
| ENSMUSG00000027878 | Notch2 | -0.7986 | 4635.556 | 0.103786 | 7.69468 | 1.42E-14 | 5.83E-13 | 7.06E-13 | 3328.745 | 3440.464 | 6378.315 | 5394.699 | 18129 | notch 2 [Source:MGI Symbol;Acc:MGI:97364] |
| ENSMUSG00000040128 | Pnrc1 | -0.79729 | 826.9884 | 0.155503 | 5.127153 | 2.94E-07 | 4.33E-06 | 5.25E-06 | 589.1678 | 618.654 | 959.2621 | 1140.87 | 108767 | proline-rich nuclear receptor coactivator 1 [Source:MGI Symbol;Acc:MGI:1917838] |
| ENSMUSG00000034563 | Ccpg1 | -0.79587 | 1155.18 | 0.165157 | 4.818877 | 1.44E-06 | 1.84E-05 | 2.23E-05 | 724.5601 | 963.8807 | 1536.282 | 1395.997 | 72278 | cell cycle progression 1 [Source:MGI Symbol;Acc:MGI:1196419] |
| ENSMUSG00000025498 | Irf7 | -0.79162 | 386.7166 | 0.198431 | 3.989392 | 6.62E-05 | 0.000581 | 0.000704 | 262.3225 | 303.9175 | 491.9762 | 488.6502 | 54123 | interferon regulatory factor 7 [Source:MGI Symbol;Acc:MGI:1859212] |
| ENSMUSG00000026473 | Glul | -0.7907 | 13749.13 | 0.075215 | 10.51249 | 7.57E-26 | 1.08E-23 | 1.31E-23 | 9603.33 | 10541.71 | 17377.37 | 17474.13 | 14645 | glutamate-ammonia ligase [Source:MGI Symbol;Acc:MGI:95739] |
| ENSMUSG00000029650 | Slc46a3 | -0.78981 | 782.126 | 0.176515 | 4.474477 | 7.66E-06 | 8.45E-05 | 0.000102 | 492.9124 | 653.0783 | 978.4656 | 1004.048 | 71706 | solute carrier family 46, member 3 [Source:MGI Symbol;Acc:MGI:1918956] |
| ENSMUSG00000032333 | Stoml1 | -0.78859 | 295.0943 | 0.253206 | 3.114427 | 0.001843 | 0.010435 | 0.012641 | 179.8178 | 252.7728 | 387.7284 | 360.0581 | 69106 | stomatin-like 1 [Source:MGI Symbol;Acc:MGI:1916356] |
| ENSMUSG00000001156 | Mxd1 | -0.78714 | 189.3016 | 0.298233 | 2.639346 | 0.008307 | 0.036582 | 0.044186 | 115.295 | 162.286 | 228.6135 | 251.0119 | 17119 | MAX dimerization protein 1 [Source:MGI Symbol;Acc:MGI:96908] |
| ENSMUSG00000022848 | Slc49a4 | -0.78466 | 494.0141 | 0.182552 | 4.298252 | 1.72E-05 | 0.000175 | 0.000212 | 332.1341 | 393.4207 | 629.1442 | 621.3574 | 224132 | solute carrier family 49 member 4 [Source:MGI Symbol;Acc:MGI:2387188] |
| ENSMUSG00000000686 | Abhd15 | -0.78127 | 423.9274 | 0.203217 | 3.844516 | 0.000121 | 0.000983 | 0.001191 | 281.362 | 342.276 | 566.9614 | 505.11 | 67477 | abhydrolase domain containing 15 [Source:MGI Symbol;Acc:MGI:1914727] |
| ENSMUSG00000024174 | Pot1b | -0.7797 | 214.3522 | 0.255431 | 3.052465 | 0.00227 | 0.01238 | 0.014996 | 168.1826 | 147.5328 | 275.2506 | 266.443 | 72836 | protection of telomeres 1B [Source:MGI Symbol;Acc:MGI:1920086] |
| ENSMUSG00000031740 | Mmp2 | -0.77809 | 266.9639 | 0.27227 | 2.857805 | 0.004266 | 0.021042 | 0.02544 | 223.1857 | 170.1544 | 290.7963 | 383.719 | 17390 | matrix metallopeptidase 2 [Source:MGI Symbol;Acc:MGI:97009] |
| ENSMUSG00000073411 | H2-D1 | -0.7775 | 11920.15 | 0.068649 | 11.32577 | 9.78E-30 | 2E-27 | 2.43E-27 | 8628.082 | 8938.518 | 14752.88 | 15361.11 | 14964 | histocompatibility 2, D region locus 1 [Source:MGI Symbol;Acc:MGI:95896] |
| ENSMUSG00000022010 | Tsc22d1 | -0.7771 | 715.2699 | 0.175349 | 4.431752 | 9.35E-06 | 0.000102 | 0.000123 | 468.5841 | 585.2133 | 849.5276 | 957.7545 | 21807 | TSC22 domain family, member 1 [Source:MGI Symbol;Acc:MGI:109127] |
| ENSMUSG00000026712 | Mrc1 | -0.77532 | 45874.14 | 0.094082 | 8.240965 | 1.71E-16 | 8.44E-15 | 1.02E-14 | 30859.91 | 36810.41 | 56150.21 | 59676.03 | 17533 | mannose receptor, C type 1 [Source:MGI Symbol;Acc:MGI:97142] |
| ENSMUSG00000029298 | Gbp9 | -0.77388 | 779.1099 | 0.147004 | 5.264342 | 1.41E-07 | 2.22E-06 | 2.69E-06 | 545.8 | 603.9007 | 956.5187 | 1010.22 | 236573 | guanylate-binding protein 9 [Source:MGI Symbol;Acc:MGI:3605620] |
| ENSMUSG00000019947 | Arid5b | -0.77264 | 1052.806 | 0.153513 | 5.033057 | 4.83E-07 | 6.81E-06 | 8.25E-06 | 744.6573 | 810.4466 | 1465.869 | 1190.249 | 71371 | AT-rich interaction domain 5B [Source:MGI Symbol;Acc:MGI:2175912] |
| ENSMUSG00000041936 | Agrn | -0.77228 | 371.4806 | 0.272549 | 2.833542 | 0.004604 | 0.022402 | 0.027084 | 204.1461 | 344.2431 | 466.3714 | 471.1617 | 11603 | agrin [Source:MGI Symbol;Acc:MGI:87961] |
| ENSMUSG00000029765 | Plxna4 | -0.77049 | 965.5403 | 0.155703 | 4.948451 | 7.48E-07 | 1.01E-05 | 1.23E-05 | 754.1771 | 673.7329 | 1331.445 | 1102.806 | 243743 | plexin A4 [Source:MGI Symbol;Acc:MGI:2179061] |
| ENSMUSG00000032000 | Birc3 | -0.77014 | 728.6671 | 0.144987 | 5.311801 | 1.09E-07 | 1.76E-06 | 2.13E-06 | 553.2043 | 524.2331 | 914.4538 | 922.7774 | 11796 | baculoviral IAP repeat-containing 3 [Source:MGI Symbol;Acc:MGI:1197007] |
| ENSMUSG00000039377 | Hlx | -0.76982 | 397.5364 | 0.214933 | 3.581675 | 0.000341 | 0.002479 | 0.003003 | 317.3256 | 270.4767 | 449.9113 | 552.432 | 15284 | H2.0-like homeobox [Source:MGI Symbol;Acc:MGI:96109] |
| ENSMUSG00000022389 | Tef | -0.76818 | 1167.142 | 0.142508 | 5.390462 | 7.03E-08 | 1.19E-06 | 1.45E-06 | 777.4477 | 949.1274 | 1450.324 | 1491.669 | 21685 | thyrotroph embryonic factor [Source:MGI Symbol;Acc:MGI:98663] |
| ENSMUSG00000048118 | Arid4a | -0.76481 | 843.3701 | 0.142372 | 5.371936 | 7.79E-08 | 1.31E-06 | 1.59E-06 | 598.6876 | 651.1112 | 1100.088 | 1023.594 | 238247 | AT-rich interaction domain 4A [Source:MGI Symbol;Acc:MGI:2444354] |
| ENSMUSG00000022587 | Ly6e | -0.76481 | 2568.218 | 0.095489 | 8.009428 | 1.15E-15 | 5.34E-14 | 6.47E-14 | 1975.881 | 1830.39 | 3202.417 | 3264.184 | 17069 | lymphocyte antigen 6 family member E [Source:MGI Symbol;Acc:MGI:106651] |
| ENSMUSG00000036528 | Ppfibp2 | -0.76481 | 770.5298 | 0.148411 | 5.153297 | 2.56E-07 | 3.83E-06 | 4.64E-06 | 606.0918 | 536.0357 | 964.7488 | 975.243 | 19024 | PTPRF interacting protein, binding protein 2 (liprin beta 2) [Source:MGI Symbol;Acc:MGI:894649] |
| ENSMUSG00000074166 | AW146154 | -0.76457 | 230.3183 | 0.242933 | 3.147261 | 0.001648 | 0.009531 | 0.011545 | 168.1826 | 173.1051 | 281.6518 | 298.3338 | 101835 | expressed sequence AW146154 [Source:MGI Symbol;Acc:MGI:2142212] |
| ENSMUSG00000053559 | Smagp | -0.76295 | 529.9772 | 0.176554 | 4.321369 | 1.55E-05 | 0.00016 | 0.000193 | 362.8089 | 422.9272 | 651.0911 | 683.0816 | 207818 | small cell adhesion glycoprotein [Source:MGI Symbol;Acc:MGI:2448476] |
| ENSMUSG00000042439 | Zfp532 | -0.76225 | 176.8239 | 0.276765 | 2.754142 | 0.005885 | 0.027446 | 0.033156 | 137.5077 | 124.9111 | 226.7845 | 218.0923 | 328977 | zinc finger protein 532 [Source:MGI Symbol;Acc:MGI:3036282] |
| ENSMUSG00000027322 | Siglec1 | -0.76218 | 2771.151 | 0.142513 | 5.348165 | 8.89E-08 | 1.47E-06 | 1.78E-06 | 2116.562 | 1995.626 | 3987.019 | 2985.396 | 20612 | sialic acid binding Ig-like lectin 1, sialoadhesin [Source:MGI Symbol;Acc:MGI:99668] |
| ENSMUSG00000102555 | 6430511E19Rik | -0.76121 | 179.3754 | 0.28626 | 2.659167 | 0.007833 | 0.034866 | 0.042124 | 121.6415 | 144.5821 | 241.4158 | 209.8624 | | RIKEN cDNA 6430511E19 gene [Source:MGI Symbol;Acc:MGI:2443259] |
| ENSMUSG00000037410 | Tbc1d2b | -0.76065 | 2027.258 | 0.105006 | 7.243862 | 4.36E-13 | 1.52E-11 | 1.84E-11 | 1451.236 | 1557.946 | 2466.282 | 2633.568 | 67016 | TBC1 domain family, member 2B [Source:MGI Symbol;Acc:MGI:1914266] |
| ENSMUSG00000042190 | Cmklr1 | -0.75948 | 1136.987 | 0.137366 | 5.528871 | 3.22E-08 | 5.82E-07 | 7.06E-07 | 801.776 | 887.1636 | 1521.651 | 1337.359 | 14747 | chemerin chemokine-like receptor 1 [Source:MGI Symbol;Acc:MGI:109603] |
| ENSMUSG00000039307 | Hexd | -0.75826 | 169.7248 | 0.291662 | 2.599785 | 0.009328 | 0.040206 | 0.048566 | 116.3527 | 135.7301 | 198.4365 | 228.3797 | 238023 | hexosaminidase D [Source:MGI Symbol;Acc:MGI:3605542] |
| ENSMUSG00000029163 | Emilin1 | -0.75804 | 283.8145 | 0.244239 | 3.103665 | 0.001911 | 0.010766 | 0.013041 | 239.0519 | 182.9406 | 342.9202 | 370.3454 | 100952 | elastin microfibril interfacer 1 [Source:MGI Symbol;Acc:MGI:1926189] |
| ENSMUSG00000103094 | Gm37558 | -0.75523 | 485.2293 | 0.258317 | 2.923652 | 0.00346 | 0.017638 | 0.021352 | 357.5201 | 364.8977 | 761.74 | 456.7594 | | predicted gene, 37558 [Source:MGI Symbol;Acc:MGI:5610786] |
| ENSMUSG00000059824 | Dbp | -0.75297 | 215.2124 | 0.288442 | 2.61046 | 0.009042 | 0.039178 | 0.047324 | 135.3922 | 184.9077 | 243.2447 | 297.3051 | 13170 | D site albumin promoter binding protein [Source:MGI Symbol;Acc:MGI:94866] |
| ENSMUSG00000095253 | Zfp799 | -0.75201 | 277.7683 | 0.227721 | 3.302338 | 0.000959 | 0.006053 | 0.007332 | 192.5108 | 221.2991 | 347.4925 | 349.7707 | 240064 | zinc finger protein 799 [Source:MGI Symbol;Acc:MGI:2443934] |
| ENSMUSG00000009090 | Ap1b1 | -0.75138 | 5717.798 | 0.089509 | 8.394384 | 4.68E-17 | 2.5E-15 | 3.03E-15 | 4014.169 | 4508.601 | 7329.347 | 7019.075 | 11764 | adaptor protein complex AP-1, beta 1 subunit [Source:MGI Symbol;Acc:MGI:1096368] |
| ENSMUSG00000033355 | Rtp4 | -0.74906 | 830.6902 | 0.141599 | 5.289986 | 1.22E-07 | 1.96E-06 | 2.37E-06 | 601.8608 | 637.3415 | 1001.327 | 1082.232 | 67775 | receptor transporter protein 4 [Source:MGI Symbol;Acc:MGI:1915025] |
| ENSMUSG00000074733 | Zfp950 | -0.74803 | 462.1811 | 0.189078 | 3.956181 | 7.62E-05 | 0.000658 | 0.000797 | 375.5019 | 314.7365 | 591.6516 | 566.8343 | 414758 | zinc finger protein 950 [Source:MGI Symbol;Acc:MGI:2652824] |
| ENSMUSG00000024621 | Csf1r | -0.74689 | 49464.29 | 0.065355 | 11.42817 | 3.02E-30 | 6.55E-28 | 7.94E-28 | 35922.31 | 37955.26 | 63216.19 | 60763.4 | 12978 | colony stimulating factor 1 receptor [Source:MGI Symbol;Acc:MGI:1339758] |
| ENSMUSG00000020668 | Kif3c | -0.7465 | 297.4938 | 0.231525 | 3.224274 | 0.001263 | 0.007639 | 0.009254 | 204.1461 | 239.9866 | 345.6635 | 400.1788 | 16570 | kinesin family member 3C [Source:MGI Symbol;Acc:MGI:107979] |
| ENSMUSG00000066258 | Trim12a | -0.74583 | 467.8947 | 0.181299 | 4.113813 | 3.89E-05 | 0.000367 | 0.000445 | 334.2496 | 364.8977 | 611.7696 | 560.6619 | 76681 | tripartite motif-containing 12A [Source:MGI Symbol;Acc:MGI:1923931] |
| ENSMUSG00000059851 | Kmt5c | -0.74457 | 302.9315 | 0.223069 | 3.33784 | 0.000844 | 0.005422 | 0.006569 | 217.8969 | 235.0689 | 406.932 | 351.8282 | 232811 | lysine methyltransferase 5C [Source:MGI Symbol;Acc:MGI:2385262] |
| ENSMUSG00000044629 | Cnrip1 | -0.74168 | 1347.423 | 0.173722 | 4.269319 | 1.96E-05 | 0.000196 | 0.000238 | 829.2775 | 1187.147 | 1670.707 | 1702.56 | 380686 | cannabinoid receptor interacting protein 1 [Source:MGI Symbol;Acc:MGI:1917505] |
| ENSMUSG00000023947 | Nfkbie | -0.74054 | 251.5281 | 0.25303 | 2.926683 | 0.003426 | 0.017497 | 0.021181 | 212.6081 | 164.2531 | 307.2565 | 321.9948 | 18037 | nuclear factor of kappa light polypeptide gene enhancer in B cells inhibitor, epsilon [Source:MGI Symbol;Acc:MGI:1194908] |
| ENSMUSG00000037526 | Atg14 | -0.74005 | 311.571 | 0.238886 | 3.097928 | 0.001949 | 0.010932 | 0.013243 | 202.0306 | 264.5754 | 412.4187 | 367.2592 | 100504663 | autophagy related 14 [Source:MGI Symbol;Acc:MGI:1261775] |
| ENSMUSG00000003031 | Cdkn1b | -0.73949 | 942.5486 | 0.15442 | 4.788809 | 1.68E-06 | 2.11E-05 | 2.55E-05 | 635.7089 | 776.0223 | 1131.179 | 1227.284 | 12576 | cyclin dependent kinase inhibitor 1B [Source:MGI Symbol;Acc:MGI:104565] |
| ENSMUSG00000038545 | Cul7 | -0.73874 | 513.8372 | 0.184188 | 4.010789 | 6.05E-05 | 0.000537 | 0.000651 | 346.9426 | 422.9272 | 637.3743 | 648.1045 | 66515 | cullin 7 [Source:MGI Symbol;Acc:MGI:1913765] |
| ENSMUSG00000078995 | Zfp456 | -0.73824 | 216.3343 | 0.256402 | 2.879226 | 0.003987 | 0.01993 | 0.024103 | 149.143 | 175.0722 | 271.5928 | 269.5292 | 408065 | zinc finger protein 456 [Source:MGI Symbol;Acc:MGI:3040694] |
| ENSMUSG00000102336 | Gm37233 | -0.73791 | 355.9451 | 0.205072 | 3.598298 | 0.00032 | 0.002341 | 0.002835 | 278.1888 | 255.7234 | 469.1148 | 420.7536 | | predicted gene, 37233 [Source:MGI Symbol;Acc:MGI:5610461] |
| ENSMUSG00000018931 | Natd1 | -0.73611 | 397.13 | 0.192204 | 3.829833 | 0.000128 | 0.001039 | 0.001258 | 294.055 | 301.9504 | 520.3242 | 472.1904 | 24083 | N-acetyltransferase domain containing 1 [Source:MGI Symbol;Acc:MGI:1344388] |
| ENSMUSG00000032827 | Ppp1r9a | -0.73407 | 1881.24 | 0.099987 | 7.341671 | 2.11E-13 | 7.67E-12 | 9.29E-12 | 1397.29 | 1428.117 | 2376.665 | 2322.889 | 243725 | protein phosphatase 1, regulatory subunit 9A [Source:MGI Symbol;Acc:MGI:2442401] |
| ENSMUSG00000018395 | Kif3a | -0.73394 | 1032.373 | 0.169227 | 4.336998 | 1.44E-05 | 0.00015 | 0.000182 | 658.9794 | 891.0978 | 1281.15 | 1298.267 | 16568 | kinesin family member 3A [Source:MGI Symbol;Acc:MGI:107689] |
| ENSMUSG00000024235 | Map3k8 | -0.73381 | 750.4656 | 0.155228 | 4.727301 | 2.28E-06 | 2.78E-05 | 3.36E-05 | 557.4353 | 569.4764 | 867.8167 | 1007.134 | 26410 | mitogen-activated protein kinase kinase kinase 8 [Source:MGI Symbol;Acc:MGI:1346878] |
| ENSMUSG00000029470 | P2rx4 | -0.7327 | 2806.316 | 0.092021 | 7.962294 | 1.69E-15 | 7.6E-14 | 9.2E-14 | 2064.732 | 2152.011 | 3409.998 | 3598.523 | 18438 | purinergic receptor P2X, ligand-gated ion channel 4 [Source:MGI Symbol;Acc:MGI:1338859] |
| ENSMUSG00000029810 | Tmem176b | -0.73047 | 5235.715 | 0.081917 | 8.917184 | 4.78E-19 | 3.14E-17 | 3.8E-17 | 3781.463 | 4093.542 | 6479.82 | 6588.034 | 65963 | transmembrane protein 176B [Source:MGI Symbol;Acc:MGI:1916348] |
| ENSMUSG00000026365 | Cfh | -0.72727 | 11782.11 | 0.082384 | 8.827806 | 1.07E-18 | 6.66E-17 | 8.07E-17 | 8345.663 | 9400.787 | 14535.24 | 14846.74 | 12628 | complement component factor h [Source:MGI Symbol;Acc:MGI:88385] |
| ENSMUSG00000073412 | Lst1 | -0.72486 | 880.8565 | 0.135759 | 5.339342 | 9.33E-08 | 1.54E-06 | 1.86E-06 | 647.3442 | 680.6178 | 1068.997 | 1126.467 | 16988 | leukocyte specific transcript 1 [Source:MGI Symbol;Acc:MGI:1096324] |
| ENSMUSG00000024180 | Pgap6 | -0.72462 | 879.1556 | 0.174157 | 4.160726 | 3.17E-05 | 0.000304 | 0.000369 | 566.955 | 758.3184 | 1068.997 | 1122.352 | 60455 | post-glycosylphosphatidylinositol attachment to proteins 6 [Source:MGI Symbol;Acc:MGI:1926283] |
| ENSMUSG00000037966 | Ninj1 | -0.72455 | 2948.193 | 0.107836 | 6.719061 | 1.83E-11 | 5.24E-10 | 6.35E-10 | 2068.963 | 2376.261 | 3528.877 | 3818.673 | 18081 | ninjurin 1 [Source:MGI Symbol;Acc:MGI:1196617] |
| ENSMUSG00000026034 | Clk1 | -0.72327 | 2406.401 | 0.094751 | 7.633377 | 2.29E-14 | 9.1E-13 | 1.1E-12 | 1785.485 | 1845.143 | 2926.252 | 3068.724 | 12747 | CDC-like kinase 1 [Source:MGI Symbol;Acc:MGI:107403] |
| ENSMUSG00000028630 | Dyrk2 | -0.72276 | 1028.125 | 0.129735 | 5.571024 | 2.53E-08 | 4.62E-07 | 5.6E-07 | 761.5814 | 789.792 | 1229.94 | 1331.186 | 69181 | dual-specificity tyrosine phosphorylation regulated kinase 2 [Source:MGI Symbol;Acc:MGI:1330301] |
| ENSMUSG00000048878 | Hexim1 | -0.72253 | 1508.728 | 0.139363 | 5.184534 | 2.17E-07 | 3.29E-06 | 3.98E-06 | 1118.044 | 1158.624 | 1677.108 | 2081.136 | 192231 | hexamethylene bis-acetamide inducible 1 [Source:MGI Symbol;Acc:MGI:2385923] |
| ENSMUSG00000056234 | Ncoa4 | -0.71998 | 9403.39 | 0.104502 | 6.889565 | 5.6E-12 | 1.74E-10 | 2.1E-10 | 6376.129 | 7832.022 | 11700.44 | 11704.97 | 27057 | nuclear receptor coactivator 4 [Source:MGI Symbol;Acc:MGI:1350932] |
| ENSMUSG00000035824 | Tk2 | -0.71806 | 971.9035 | 0.138015 | 5.202733 | 1.96E-07 | 3.01E-06 | 3.65E-06 | 700.2318 | 769.1374 | 1154.955 | 1263.289 | 57813 | thymidine kinase 2, mitochondrial [Source:MGI Symbol;Acc:MGI:1913266] |
| ENSMUSG00000030067 | Foxp1 | -0.71587 | 590.7991 | 0.171288 | 4.179327 | 2.92E-05 | 0.000282 | 0.000342 | 425.2163 | 469.1542 | 783.6869 | 685.1391 | 108655 | forkhead box P1 [Source:MGI Symbol;Acc:MGI:1914004] |
| ENSMUSG00000104291 | A130071D04Rik | -0.71485 | 378.5853 | 0.209286 | 3.415687 | 0.000636 | 0.004261 | 0.005162 | 297.2283 | 276.378 | 514.8375 | 425.8973 | | RIKEN cDNA A130071D04 gene [Source:MGI Symbol;Acc:MGI:2444749] |
| ENSMUSG00000139722 | Gm66006 | -0.71429 | 418.9851 | 0.21669 | 3.296373 | 0.000979 | 0.006167 | 0.007471 | 355.4046 | 279.3287 | 476.4304 | 564.7768 | | predicted gene, 66006 [Source:MGI Symbol;Acc:MGI:7814887] |
| ENSMUSG00000017119 | Nbr1 | -0.71179 | 3794.903 | 0.103405 | 6.88349 | 5.84E-12 | 1.8E-10 | 2.18E-10 | 2649.669 | 3104.089 | 4646.34 | 4779.514 | 17966 | NBR1, autophagy cargo receptor [Source:MGI Symbol;Acc:MGI:108498] |
| ENSMUSG00000062822 | 4833420G17Rik | -0.71106 | 704.4725 | 0.155421 | 4.575065 | 4.76E-06 | 5.46E-05 | 6.62E-05 | 543.6845 | 525.2166 | 929.0851 | 819.9037 | 67392 | RIKEN cDNA 4833420G17 gene [Source:MGI Symbol;Acc:MGI:1914642] |
| ENSMUSG00000002602 | Axl | -0.7096 | 2542.006 | 0.099596 | 7.1248 | 1.04E-12 | 3.47E-11 | 4.21E-11 | 1839.431 | 2018.248 | 3086.282 | 3224.063 | 26362 | AXL receptor tyrosine kinase [Source:MGI Symbol;Acc:MGI:1347244] |
| ENSMUSG00000041777 | Cir1 | -0.70887 | 243.7444 | 0.243018 | 2.916955 | 0.003535 | 0.017973 | 0.02175 | 170.2981 | 199.661 | 303.5987 | 301.42 | 66935 | corepressor interacting with RBPJ, 1 [Source:MGI Symbol;Acc:MGI:1914185] |
| ENSMUSG00000040296 | Rigi | -0.70861 | 1322.675 | 0.122096 | 5.803694 | 6.49E-09 | 1.29E-07 | 1.57E-07 | 984.767 | 1023.877 | 1725.574 | 1556.48 | 230073 | RNA sensor RIG-I [Source:MGI Symbol;Acc:MGI:2442858] |
| ENSMUSG00000006611 | Hfe | -0.70787 | 1905.056 | 0.108251 | 6.53915 | 6.19E-11 | 1.65E-09 | 2E-09 | 1368.731 | 1524.505 | 2354.719 | 2372.268 | 15216 | homeostatic iron regulator [Source:MGI Symbol;Acc:MGI:109191] |
| ENSMUSG00000061232 | H2-K1 | -0.70708 | 6988.449 | 0.071531 | 9.884918 | 4.84E-23 | 4.99E-21 | 6.05E-21 | 5214.717 | 5403.633 | 8633.359 | 8702.089 | 14972 | histocompatibility 2, K1, K region [Source:MGI Symbol;Acc:MGI:95904] |
| ENSMUSG00000005534 | Insr | -0.70646 | 1046.461 | 0.134997 | 5.233145 | 1.67E-07 | 2.59E-06 | 3.14E-06 | 806.007 | 784.8743 | 1378.996 | 1215.968 | 16337 | insulin receptor [Source:MGI Symbol;Acc:MGI:96575] |
| ENSMUSG00000037341 | Slc9a7 | -0.70641 | 410.6994 | 0.213513 | 3.308506 | 0.000938 | 0.005936 | 0.00719 | 270.7845 | 353.0951 | 490.1472 | 528.771 | 236727 | solute carrier family 9 (sodium/hydrogen exchanger), member 7 [Source:MGI Symbol;Acc:MGI:2444530] |
| ENSMUSG00000020377 | Ltc4s | -0.70567 | 999.4955 | 0.14319 | 4.928175 | 8.3E-07 | 1.11E-05 | 1.35E-05 | 698.1163 | 821.2657 | 1272.92 | 1205.68 | 17001 | leukotriene C4 synthase [Source:MGI Symbol;Acc:MGI:107498] |
| ENSMUSG00000031613 | Hpgd | -0.70538 | 3918.787 | 0.088884 | 7.936008 | 2.09E-15 | 9.31E-14 | 1.13E-13 | 2941.608 | 3016.553 | 4646.34 | 5070.646 | 15446 | hydroxyprostaglandin dehydrogenase 15 (NAD) [Source:MGI Symbol;Acc:MGI:108085] |
| ENSMUSG00000034422 | Parp14 | -0.70438 | 4490.203 | 0.07835 | 8.990125 | 2.47E-19 | 1.69E-17 | 2.05E-17 | 3478.946 | 3351.944 | 5547.991 | 5581.929 | 547253 | poly (ADP-ribose) polymerase family, member 14 [Source:MGI Symbol;Acc:MGI:1919489] |
| ENSMUSG00000039382 | Wdr45 | -0.70402 | 268.9175 | 0.227633 | 3.092803 | 0.001983 | 0.011078 | 0.013419 | 194.6264 | 214.4143 | 329.2034 | 337.4259 | 54636 | WD repeat domain 45 [Source:MGI Symbol;Acc:MGI:1859606] |
| ENSMUSG00000002996 | Hbp1 | -0.70351 | 2208.106 | 0.105003 | 6.699846 | 2.09E-11 | 5.94E-10 | 7.19E-10 | 1584.512 | 1775.311 | 2735.131 | 2737.47 | 73389 | high mobility group box transcription factor 1 [Source:MGI Symbol;Acc:MGI:894659] |
| ENSMUSG00000041685 | Fcho2 | -0.69888 | 3076.385 | 0.084939 | 8.227949 | 1.9E-16 | 9.35E-15 | 1.13E-14 | 2363.018 | 2328.067 | 3813.272 | 3801.185 | 218503 | FCH domain only 2 [Source:MGI Symbol;Acc:MGI:3505790] |
| ENSMUSG00000029468 | P2rx7 | -0.69789 | 2680.327 | 0.098424 | 7.090674 | 1.33E-12 | 4.4E-11 | 5.33E-11 | 1980.112 | 2108.735 | 3447.491 | 3184.971 | 18439 | purinergic receptor P2X, ligand-gated ion channel, 7 [Source:MGI Symbol;Acc:MGI:1339957] |
| ENSMUSG00000033032 | Afap1l1 | -0.69729 | 506.027 | 0.175704 | 3.968543 | 7.23E-05 | 0.000627 | 0.000759 | 361.7512 | 410.1411 | 615.4274 | 636.7884 | 106877 | actin filament associated protein 1-like 1 [Source:MGI Symbol;Acc:MGI:2147199] |
| ENSMUSG00000021703 | Serinc5 | -0.69692 | 809.3926 | 0.168894 | 4.126382 | 3.69E-05 | 0.000349 | 0.000422 | 539.4535 | 695.371 | 1005.899 | 996.8465 | 218442 | serine incorporator 5 [Source:MGI Symbol;Acc:MGI:2444223] |
| ENSMUSG00000020893 | Per1 | -0.69679 | 230.0477 | 0.243633 | 2.859986 | 0.004237 | 0.020918 | 0.02529 | 182.9911 | 168.1873 | 287.1385 | 281.874 | 18626 | period circadian clock 1 [Source:MGI Symbol;Acc:MGI:1098283] |
| ENSMUSG00000024672 | Ms4a7 | -0.69467 | 8703.922 | 0.086895 | 7.994296 | 1.3E-15 | 5.93E-14 | 7.19E-14 | 6567.582 | 6727.494 | 10080.02 | 11440.59 | 109225 | membrane-spanning 4-domains, subfamily A, member 7 [Source:MGI Symbol;Acc:MGI:1918846] |
| ENSMUSG00000029385 | Ccng2 | -0.69458 | 963.8824 | 0.129918 | 5.346302 | 8.98E-08 | 1.48E-06 | 1.8E-06 | 715.0403 | 757.3348 | 1205.25 | 1177.904 | 12452 | cyclin G2 [Source:MGI Symbol;Acc:MGI:1095734] |
| ENSMUSG00000066800 | Rnasel | -0.69021 | 3648.428 | 0.086679 | 7.962793 | 1.68E-15 | 7.59E-14 | 9.19E-14 | 2689.863 | 2893.609 | 4498.198 | 4512.042 | 24014 | ribonuclease L (2', 5'-oligoisoadenylate synthetase-dependent) [Source:MGI Symbol;Acc:MGI:1098272] |
| ENSMUSG00000118087 | 4833438C02Rik | -0.68985 | 206.793 | 0.259466 | 2.658741 | 0.007843 | 0.03489 | 0.042154 | 152.3163 | 164.2531 | 268.8494 | 241.7533 | 100503392 | RIKEN cDNA 4833438C02 gene [Source:MGI Symbol;Acc:MGI:2147599] |
| ENSMUSG00000060441 | Trim5 | -0.6891 | 289.0884 | 0.244865 | 2.81421 | 0.00489 | 0.023562 | 0.028479 | 196.7419 | 245.8879 | 388.6429 | 325.081 | 667823 | tripartite motif-containing 5 [Source:MGI Symbol;Acc:MGI:3646853] |
| ENSMUSG00000042628 | Zfyve1 | -0.6891 | 1131.095 | 0.125701 | 5.482041 | 4.2E-08 | 7.43E-07 | 9E-07 | 840.9128 | 891.0978 | 1445.751 | 1346.617 | 217695 | zinc finger, FYVE domain containing 1 [Source:MGI Symbol;Acc:MGI:3026685] |
| ENSMUSG00000034908 | Sidt2 | -0.68901 | 1446.417 | 0.133518 | 5.160422 | 2.46E-07 | 3.7E-06 | 4.48E-06 | 1020.731 | 1194.032 | 1883.775 | 1687.129 | 214597 | SID1 transmembrane family, member 2 [Source:MGI Symbol;Acc:MGI:2446134] |
| ENSMUSG00000035236 | Scai | -0.68831 | 418.4084 | 0.201761 | 3.411516 | 0.000646 | 0.004315 | 0.005227 | 285.593 | 355.0622 | 517.5809 | 515.3974 | 320271 | suppressor of cancer cell invasion [Source:MGI Symbol;Acc:MGI:2443716] |
| ENSMUSG00000044471 | Lncpint | -0.68612 | 295.183 | 0.241987 | 2.835354 | 0.004577 | 0.022304 | 0.026966 | 258.0915 | 194.7432 | 374.0116 | 353.8856 | 232685 | long non-protein coding RNA, Trp53 induced transcript [Source:MGI Symbol;Acc:MGI:2673128] |
| ENSMUSG00000056153 | Socs6 | -0.68537 | 2661.96 | 0.10392 | 6.595192 | 4.25E-11 | 1.16E-09 | 1.4E-09 | 1909.242 | 2172.666 | 3238.995 | 3326.937 | 54607 | suppressor of cytokine signaling 6 [Source:MGI Symbol;Acc:MGI:1924885] |
| ENSMUSG00000068036 | Afdn | -0.68217 | 287.4068 | 0.244056 | 2.795123 | 0.005188 | 0.024769 | 0.029929 | 217.8969 | 223.2662 | 307.2565 | 401.2076 | 17356 | afadin, adherens junction formation factor [Source:MGI Symbol;Acc:MGI:1314653] |
| ENSMUSG00000052253 | Zfp622 | -0.68074 | 1160.884 | 0.152014 | 4.478136 | 7.53E-06 | 8.32E-05 | 0.000101 | 997.4601 | 786.8414 | 1388.141 | 1471.094 | 52521 | zinc finger protein 622 [Source:MGI Symbol;Acc:MGI:1289282] |
| ENSMUSG00000040855 | Reps2 | -0.67893 | 2964.695 | 0.098725 | 6.876998 | 6.11E-12 | 1.88E-10 | 2.28E-10 | 2157.814 | 2400.85 | 3562.712 | 3737.403 | 194590 | RALBP1 associated Eps domain containing protein 2 [Source:MGI Symbol;Acc:MGI:2663511] |
| ENSMUSG00000040033 | Stat2 | -0.67796 | 558.3464 | 0.180597 | 3.754011 | 0.000174 | 0.001359 | 0.001646 | 414.6387 | 444.5654 | 750.7666 | 623.4148 | 20847 | signal transducer and activator of transcription 2 [Source:MGI Symbol;Acc:MGI:103039] |
| ENSMUSG00000050103 | Agmo | -0.67662 | 292.7552 | 0.220727 | 3.065393 | 0.002174 | 0.011954 | 0.014481 | 211.5504 | 239.0031 | 359.3804 | 361.0868 | 319660 | alkylglycerol monooxygenase [Source:MGI Symbol;Acc:MGI:2442495] |
| ENSMUSG00000044952 | Kctd21 | -0.67656 | 218.6061 | 0.255976 | 2.643078 | 0.008216 | 0.036266 | 0.043804 | 154.4318 | 181.9571 | 263.3627 | 274.6729 | 622320 | potassium channel tetramerisation domain containing 21 [Source:MGI Symbol;Acc:MGI:3643121] |
| ENSMUSG00000006235 | Epor | -0.67607 | 232.957 | 0.258344 | 2.616921 | 0.008873 | 0.038649 | 0.046684 | 160.7783 | 197.6939 | 268.8494 | 304.5063 | 13857 | erythropoietin receptor [Source:MGI Symbol;Acc:MGI:95408] |
| ENSMUSG00000051439 | Cd14 | -0.67442 | 4842.767 | 0.140307 | 4.806699 | 1.53E-06 | 1.95E-05 | 2.36E-05 | 4060.71 | 3401.122 | 5156.605 | 6752.632 | 12475 | CD14 antigen [Source:MGI Symbol;Acc:MGI:88318] |
| ENSMUSG00000026571 | Dcaf6 | -0.67317 | 493.6751 | 0.188408 | 3.572956 | 0.000353 | 0.002556 | 0.003096 | 409.35 | 352.1115 | 647.4333 | 565.8055 | 74106 | DDB1 and CUL4 associated factor 6 [Source:MGI Symbol;Acc:MGI:1921356] |
| ENSMUSG00000027428 | Rbbp9 | -0.67315 | 217.8814 | 0.252812 | 2.662639 | 0.007753 | 0.034576 | 0.041774 | 159.7205 | 176.0558 | 256.9615 | 278.7878 | 26450 | retinoblastoma binding protein 9, serine hydrolase [Source:MGI Symbol;Acc:MGI:1347074] |
| ENSMUSG00000033444 | Specc1l | -0.66914 | 1646.324 | 0.105669 | 6.332431 | 2.41E-10 | 5.9E-09 | 7.15E-09 | 1247.09 | 1295.338 | 2050.205 | 1992.664 | 74392 | sperm antigen with calponin homology and coiled-coil domains 1-like [Source:MGI Symbol;Acc:MGI:1921642] |
| ENSMUSG00000046718 | Bst2 | -0.66908 | 2703.125 | 0.107807 | 6.206323 | 5.42E-10 | 1.27E-08 | 1.54E-08 | 2042.519 | 2131.357 | 3085.367 | 3553.259 | 69550 | bone marrow stromal cell antigen 2 [Source:MGI Symbol;Acc:MGI:1916800] |
| ENSMUSG00000040026 | Saa3 | -0.66888 | 769.3129 | 0.163214 | 4.098151 | 4.16E-05 | 0.000389 | 0.000471 | 579.648 | 608.8185 | 1036.991 | 851.7945 | 20210 | serum amyloid A 3 [Source:MGI Symbol;Acc:MGI:98223] |
| ENSMUSG00000008540 | Mgst1 | -0.66884 | 1029.959 | 0.173133 | 3.863168 | 0.000112 | 0.000918 | 0.001112 | 683.3077 | 906.8347 | 1176.902 | 1352.79 | 56615 | microsomal glutathione S-transferase 1 [Source:MGI Symbol;Acc:MGI:1913850] |
| ENSMUSG00000024248 | Cox7a2l | -0.66824 | 3752.615 | 0.097803 | 6.832543 | 8.34E-12 | 2.5E-10 | 3.03E-10 | 2714.191 | 3082.451 | 4507.343 | 4706.473 | 20463 | cytochrome c oxidase subunit 7A2 like [Source:MGI Symbol;Acc:MGI:106015] |
| ENSMUSG00000033423 | Eri3 | -0.66819 | 971.3786 | 0.147988 | 4.515187 | 6.33E-06 | 7.07E-05 | 8.57E-05 | 680.1345 | 820.2821 | 1214.395 | 1170.703 | 140546 | exoribonuclease 3 [Source:MGI Symbol;Acc:MGI:2153887] |
| ENSMUSG00000034445 | Cyb561a3 | -0.66642 | 616.4825 | 0.193554 | 3.443102 | 0.000575 | 0.00391 | 0.004737 | 414.6387 | 538.0028 | 699.5572 | 813.7312 | 225912 | cytochrome b561 family, member A3 [Source:MGI Symbol;Acc:MGI:2686925] |
| ENSMUSG00000042500 | Ago4 | -0.66641 | 617.0682 | 0.153692 | 4.335991 | 1.45E-05 | 0.00015 | 0.000182 | 477.0461 | 477.0226 | 758.0822 | 756.122 | 76850 | argonaute RISC catalytic subunit 4 [Source:MGI Symbol;Acc:MGI:1924100] |
| ENSMUSG00000034858 | Atosa | -0.66353 | 423.3597 | 0.187311 | 3.542374 | 0.000397 | 0.00283 | 0.003428 | 324.7298 | 330.4734 | 491.9762 | 546.2595 | 235493 | atos homolog A [Source:MGI Symbol;Acc:MGI:2387648] |
| ENSMUSG00000033066 | Gas7 | -0.66235 | 1679.128 | 0.122166 | 5.421697 | 5.9E-08 | 1.02E-06 | 1.24E-06 | 1252.378 | 1348.449 | 2200.176 | 1915.509 | 14457 | growth arrest specific 7 [Source:MGI Symbol;Acc:MGI:1202388] |
| ENSMUSG00000050994 | Adgb | -0.66222 | 655.8161 | 0.178436 | 3.711273 | 0.000206 | 0.001582 | 0.001916 | 490.7969 | 525.2166 | 894.3358 | 712.915 | 215772 | androglobin [Source:MGI Symbol;Acc:MGI:3605549] |
| ENSMUSG00000035051 | Dhx57 | -0.65973 | 734.1045 | 0.159388 | 4.139135 | 3.49E-05 | 0.000331 | 0.000401 | 518.2984 | 619.6376 | 903.4804 | 895.0015 | 106794 | DExH-box helicase 57 [Source:MGI Symbol;Acc:MGI:2147067] |
| ENSMUSG00000032596 | Uba7 | -0.6584 | 1761.765 | 0.137627 | 4.783957 | 1.72E-06 | 2.15E-05 | 2.61E-05 | 1213.241 | 1519.587 | 2241.326 | 2072.906 | 74153 | ubiquitin-like modifier activating enzyme 7 [Source:MGI Symbol;Acc:MGI:1349462] |
| ENSMUSG00000036053 | Fmnl2 | -0.65654 | 979.7485 | 0.134036 | 4.898266 | 9.67E-07 | 1.28E-05 | 1.55E-05 | 719.2713 | 801.5946 | 1192.448 | 1205.68 | 71409 | formin-like 2 [Source:MGI Symbol;Acc:MGI:1918659] |
| ENSMUSG00000021327 | Zkscan3 | -0.65623 | 797.2145 | 0.147116 | 4.460638 | 8.17E-06 | 8.98E-05 | 0.000109 | 592.3411 | 645.2099 | 931.8284 | 1019.479 | 72739 | zinc finger with KRAB and SCAN domains 3 [Source:MGI Symbol;Acc:MGI:1919989] |
| ENSMUSG00000045216 | Hs6st1 | -0.65591 | 953.6328 | 0.129465 | 5.066345 | 4.06E-07 | 5.8E-06 | 7.03E-06 | 725.6178 | 755.3677 | 1182.389 | 1151.157 | 50785 | heparan sulfate 6-O-sulfotransferase 1 [Source:MGI Symbol;Acc:MGI:1354958] |
| ENSMUSG00000029060 | Mib2 | -0.65512 | 809.2743 | 0.154009 | 4.253779 | 2.1E-05 | 0.000209 | 0.000253 | 573.3015 | 683.5684 | 980.2945 | 999.9327 | 76580 | mindbomb E3 ubiquitin protein ligase 2 [Source:MGI Symbol;Acc:MGI:2679684] |
| ENSMUSG00000028793 | Rnf19b | -0.65506 | 1262.043 | 0.124039 | 5.281075 | 1.28E-07 | 2.05E-06 | 2.48E-06 | 939.2837 | 1020.927 | 1484.159 | 1603.802 | 75234 | ring finger protein 19B [Source:MGI Symbol;Acc:MGI:1922484] |
| ENSMUSG00000022106 | Rcbtb2 | -0.65441 | 4095.882 | 0.107515 | 6.086678 | 1.15E-09 | 2.57E-08 | 3.11E-08 | 2897.182 | 3467.02 | 4918.847 | 5100.48 | 105670 | regulator of chromosome condensation (RCC1) and BTB (POZ) domain containing protein 2 [Source:MGI Symbol;Acc:MGI:1917200] |
| ENSMUSG00000079110 | Capn3 | -0.65425 | 356.9697 | 0.202287 | 3.234269 | 0.00122 | 0.007409 | 0.008975 | 264.438 | 290.1477 | 420.6488 | 452.6444 | 12335 | calpain 3 [Source:MGI Symbol;Acc:MGI:107437] |
| ENSMUSG00000030737 | Slco2b1 | -0.65293 | 4658.708 | 0.077181 | 8.459728 | 2.68E-17 | 1.47E-15 | 1.78E-15 | 3634.436 | 3609.635 | 5609.26 | 5781.504 | 101488 | solute carrier organic anion transporter family, member 2b1 [Source:MGI Symbol;Acc:MGI:1351872] |
| ENSMUSG00000034610 | Tut4 | -0.65111 | 655.0424 | 0.169903 | 3.832263 | 0.000127 | 0.001029 | 0.001246 | 558.493 | 461.2857 | 825.7518 | 774.6392 | 230594 | terminal uridylyl transferase 4 [Source:MGI Symbol;Acc:MGI:2445126] |
| ENSMUSG00000048280 | Zfp738 | -0.64928 | 581.8993 | 0.157969 | 4.11017 | 3.95E-05 | 0.000373 | 0.000452 | 453.7756 | 452.4338 | 702.3005 | 719.0874 | 408068 | zinc finger protein 738 [Source:MGI Symbol;Acc:MGI:3040706] |
| ENSMUSG00000030161 | Gabarapl1 | -0.64377 | 1439.743 | 0.12128 | 5.308106 | 1.11E-07 | 1.79E-06 | 2.17E-06 | 1058.81 | 1188.13 | 1704.542 | 1807.492 | 57436 | GABA type A receptor associated protein like 1 [Source:MGI Symbol;Acc:MGI:1914980] |
| ENSMUSG00000071669 | Snx29 | -0.6419 | 858.2739 | 0.157696 | 4.070454 | 4.69E-05 | 0.00043 | 0.000521 | 683.3077 | 657.9961 | 1149.468 | 942.3234 | 74478 | sorting nexin 29 [Source:MGI Symbol;Acc:MGI:1921728] |
| ENSMUSG00000028859 | Csf3r | -0.64113 | 800.602 | 0.139194 | 4.605987 | 4.11E-06 | 4.75E-05 | 5.75E-05 | 615.6116 | 635.3744 | 955.6042 | 995.8178 | 12986 | colony stimulating factor 3 receptor [Source:MGI Symbol;Acc:MGI:1339755] |
| ENSMUSG00000020432 | Tcn2 | -0.63911 | 4834.756 | 0.100514 | 6.358413 | 2.04E-10 | 5.03E-09 | 6.1E-09 | 3475.773 | 4085.674 | 5954.923 | 5822.653 | 21452 | transcobalamin 2 [Source:MGI Symbol;Acc:MGI:98534] |
| ENSMUSG00000071076 | Jund | -0.63846 | 564.5178 | 0.174367 | 3.661607 | 0.000251 | 0.001885 | 0.002284 | 403.0035 | 479.9732 | 694.0704 | 681.0241 | 16478 | jun D proto-oncogene [Source:MGI Symbol;Acc:MGI:96648] |
| ENSMUSG00000018340 | Anxa6 | -0.63787 | 2650.315 | 0.094976 | 6.716154 | 1.87E-11 | 5.33E-10 | 6.45E-10 | 1995.978 | 2151.028 | 3177.727 | 3276.528 | 11749 | annexin A6 [Source:MGI Symbol;Acc:MGI:88255] |
| ENSMUSG00000060301 | 2610008E11Rik | -0.63359 | 416.191 | 0.18321 | 3.458281 | 0.000544 | 0.003728 | 0.004516 | 327.9031 | 324.5721 | 502.0351 | 510.2537 | 72128 | RIKEN cDNA 2610008E11 gene [Source:MGI Symbol;Acc:MGI:1919378] |
| ENSMUSG00000036561 | Ppp6r2 | -0.63309 | 663.3104 | 0.161118 | 3.929324 | 8.52E-05 | 0.000724 | 0.000877 | 481.2771 | 558.6574 | 821.1795 | 792.1278 | 71474 | protein phosphatase 6, regulatory subunit 2 [Source:MGI Symbol;Acc:MGI:1918724] |
| ENSMUSG00000026594 | Ralgps2 | -0.63228 | 1342.456 | 0.116627 | 5.421428 | 5.91E-08 | 1.02E-06 | 1.24E-06 | 1063.041 | 1042.565 | 1571.946 | 1692.273 | 78255 | Ral GEF with PH domain and SH3 binding motif 2 [Source:MGI Symbol;Acc:MGI:1925505] |
| ENSMUSG00000025509 | Pnpla2 | -0.63118 | 249.2332 | 0.232752 | 2.711793 | 0.006692 | 0.030545 | 0.036902 | 191.4531 | 199.661 | 305.4276 | 300.3913 | 66853 | patatin-like phospholipase domain containing 2 [Source:MGI Symbol;Acc:MGI:1914103] |
| ENSMUSG00000044080 | S100a1 | -0.63071 | 613.864 | 0.193328 | 3.262416 | 0.001105 | 0.006819 | 0.008261 | 412.5232 | 550.7889 | 778.2002 | 713.9437 | 20193 | S100 calcium binding protein A1 [Source:MGI Symbol;Acc:MGI:1338917] |
| ENSMUSG00000049659 | Aftph | -0.62942 | 3164.222 | 0.093951 | 6.699462 | 2.09E-11 | 5.94E-10 | 7.2E-10 | 2394.75 | 2573.955 | 3717.255 | 3970.926 | 216549 | aftiphilin [Source:MGI Symbol;Acc:MGI:1923012] |
| ENSMUSG00000029009 | Mthfr | -0.62871 | 2004.217 | 0.106578 | 5.899014 | 3.66E-09 | 7.62E-08 | 9.23E-08 | 1495.661 | 1652.367 | 2388.553 | 2480.286 | 17769 | methylenetetrahydrofolate reductase [Source:MGI Symbol;Acc:MGI:106639] |
| ENSMUSG00000036908 | Unc93b1 | -0.62829 | 8329.79 | 0.07537 | 8.336072 | 7.68E-17 | 3.99E-15 | 4.83E-15 | 6313.721 | 6773.721 | 9980.349 | 10251.37 | 54445 | unc-93 homolog B1, TLR signaling regulator [Source:MGI Symbol;Acc:MGI:1859307] |
| ENSMUSG00000047414 | Flrt2 | -0.62715 | 334.9686 | 0.22661 | 2.767514 | 0.005649 | 0.026548 | 0.03207 | 239.0519 | 287.1971 | 371.2683 | 442.3571 | 399558 | fibronectin leucine rich transmembrane protein 2 [Source:MGI Symbol;Acc:MGI:3603594] |
| ENSMUSG00000110185 | Iglp | -0.62656 | 294.0498 | 0.230175 | 2.722104 | 0.006487 | 0.029725 | 0.03591 | 208.3771 | 253.7563 | 344.7491 | 369.3167 | 109169 | IgA inducing protein [Source:MGI Symbol;Acc:MGI:1924271] |
| ENSMUSG00000039656 | Rxrb | -0.62632 | 697.4308 | 0.171145 | 3.65958 | 0.000253 | 0.001899 | 0.002301 | 488.6814 | 607.8349 | 873.3034 | 819.9037 | 20182 | retinoid X receptor beta [Source:MGI Symbol;Acc:MGI:98215] |
| ENSMUSG00000055116 | Bmal1 | -0.62521 | 383.6353 | 0.197889 | 3.159371 | 0.001581 | 0.009232 | 0.011183 | 301.4593 | 301.9504 | 435.28 | 495.8514 | 11865 | basic helix-loop-helix ARNT like 1 [Source:MGI Symbol;Acc:MGI:1096381] |
| ENSMUSG00000021238 | Aldh6a1 | -0.62404 | 625.7574 | 0.170208 | 3.666318 | 0.000246 | 0.001854 | 0.002245 | 449.5446 | 535.0521 | 732.4775 | 785.9553 | 104776 | aldehyde dehydrogenase family 6, subfamily A1 [Source:MGI Symbol;Acc:MGI:1915077] |
| ENSMUSG00000005533 | Igf1r | -0.624 | 1065.595 | 0.143754 | 4.340727 | 1.42E-05 | 0.000148 | 0.000179 | 883.2228 | 794.7098 | 1389.055 | 1195.393 | 16001 | insulin-like growth factor I receptor [Source:MGI Symbol;Acc:MGI:96433] |
| ENSMUSG00000040430 | Pitpnc1 | -0.62399 | 1109.449 | 0.127034 | 4.911986 | 9.02E-07 | 1.2E-05 | 1.45E-05 | 833.5085 | 912.736 | 1371.681 | 1319.87 | 71795 | phosphatidylinositol transfer protein, cytoplasmic 1 [Source:MGI Symbol;Acc:MGI:1919045] |
| ENSMUSG00000020400 | Tnip1 | -0.62357 | 319.111 | 0.21234 | 2.936672 | 0.003318 | 0.017035 | 0.020622 | 247.5139 | 254.7399 | 365.7815 | 408.4087 | 57783 | TNFAIP3 interacting protein 1 [Source:MGI Symbol;Acc:MGI:1926194] |
| ENSMUSG00000027522 | Stx16 | -0.62334 | 810.8173 | 0.140224 | 4.44534 | 8.78E-06 | 9.59E-05 | 0.000116 | 662.1527 | 614.7198 | 992.1824 | 974.2143 | 228960 | syntaxin 16 [Source:MGI Symbol;Acc:MGI:1923396] |
| ENSMUSG00000025283 | Sat1 | -0.62256 | 8467.796 | 0.076087 | 8.182146 | 2.79E-16 | 1.36E-14 | 1.64E-14 | 6440.651 | 6895.681 | 10057.16 | 10477.69 | 20229 | spermidine/spermine N1-acetyl transferase 1 [Source:MGI Symbol;Acc:MGI:98233] |
| ENSMUSG00000030729 | Pgm2l1 | -0.62063 | 1406.442 | 0.112789 | 5.502566 | 3.74E-08 | 6.66E-07 | 8.07E-07 | 1079.965 | 1136.986 | 1731.976 | 1676.842 | 70974 | phosphoglucomutase 2-like 1 [Source:MGI Symbol;Acc:MGI:1918224] |
| ENSMUSG00000030126 | Tmcc1 | -0.62047 | 650.9304 | 0.163218 | 3.801478 | 0.000144 | 0.00115 | 0.001393 | 542.6267 | 483.9074 | 830.3241 | 746.8633 | 330401 | transmembrane and coiled coil domains 1 [Source:MGI Symbol;Acc:MGI:2442368] |
| ENSMUSG00000004952 | Rasa4 | -0.62041 | 1594.736 | 0.127341 | 4.872086 | 1.1E-06 | 1.45E-05 | 1.75E-05 | 1144.488 | 1369.104 | 1947.787 | 1917.566 | 54153 | RAS p21 protein activator 4 [Source:MGI Symbol;Acc:MGI:1858600] |
| ENSMUSG00000108621 | Gm33989 | -0.62036 | 890.1089 | 0.145721 | 4.257172 | 2.07E-05 | 0.000206 | 0.00025 | 686.481 | 717.0092 | 1154.955 | 1001.99 | 102637087 | predicted gene, 33989 [Source:MGI Symbol;Acc:MGI:5593148] |
| ENSMUSG00000013495 | Tmem175 | -0.62034 | 461.467 | 0.182821 | 3.393176 | 0.000691 | 0.004573 | 0.005539 | 345.8849 | 381.6181 | 583.4215 | 534.9434 | 72392 | transmembrane protein 175 [Source:MGI Symbol;Acc:MGI:1919642] |
| ENSMUSG00000036622 | Atp13a2 | -0.61992 | 6550.064 | 0.10445 | 5.935059 | 2.94E-09 | 6.19E-08 | 7.49E-08 | 4686.899 | 5640.669 | 8155.099 | 7717.588 | 74772 | ATPase type 13A2 [Source:MGI Symbol;Acc:MGI:1922022] |
| ENSMUSG00000042129 | Rassf4 | -0.61953 | 3910.822 | 0.090518 | 6.844257 | 7.69E-12 | 2.32E-10 | 2.81E-10 | 3176.429 | 2991.964 | 4928.906 | 4545.99 | 213391 | Ras association (RalGDS/AF-6) domain family member 4 [Source:MGI Symbol;Acc:MGI:2386853] |
| ENSMUSG00000019872 | Smpdl3a | -0.61948 | 1207.758 | 0.135615 | 4.567944 | 4.93E-06 | 5.64E-05 | 6.83E-05 | 896.9736 | 1007.157 | 1373.51 | 1553.393 | 57319 | sphingomyelin phosphodiesterase, acid-like 3A [Source:MGI Symbol;Acc:MGI:1931437] |
| ENSMUSG00000106874 | Gm20186 | -0.61941 | 352.1903 | 0.219784 | 2.818247 | 0.004829 | 0.02332 | 0.028187 | 280.3043 | 275.3945 | 475.516 | 377.5466 | | predicted gene, 20186 [Source:MGI Symbol;Acc:MGI:5012371] |
| ENSMUSG00000006127 | Inpp5k | -0.61902 | 667.8533 | 0.155278 | 3.986508 | 6.71E-05 | 0.000586 | 0.00071 | 522.5294 | 531.1179 | 851.3565 | 766.4093 | 19062 | inositol polyphosphate 5-phosphatase K [Source:MGI Symbol;Acc:MGI:1194899] |
| ENSMUSG00000024190 | Dusp1 | -0.61886 | 718.3417 | 0.176303 | 3.510202 | 0.000448 | 0.003148 | 0.003813 | 618.7849 | 514.3975 | 793.7459 | 946.4384 | 19252 | dual specificity phosphatase 1 [Source:MGI Symbol;Acc:MGI:105120] |
| ENSMUSG00000002763 | Pex6 | -0.61737 | 581.668 | 0.157964 | 3.908261 | 9.3E-05 | 0.000779 | 0.000943 | 454.8333 | 463.2528 | 697.7283 | 710.8575 | 224824 | peroxisomal biogenesis factor 6 [Source:MGI Symbol;Acc:MGI:2385054] |
| ENSMUSG00000000056 | Narf | -0.6141 | 930.702 | 0.142566 | 4.307472 | 1.65E-05 | 0.000169 | 0.000205 | 704.4628 | 766.1868 | 1059.852 | 1192.307 | 67608 | nuclear prelamin A recognition factor [Source:MGI Symbol;Acc:MGI:1914858] |
| ENSMUSG00000021549 | Rasa1 | -0.61371 | 3977.595 | 0.092915 | 6.605091 | 3.97E-11 | 1.09E-09 | 1.32E-09 | 2993.438 | 3293.915 | 4657.313 | 4965.715 | 218397 | RAS p21 protein activator 1 [Source:MGI Symbol;Acc:MGI:97860] |
| ENSMUSG00000018169 | Mfng | -0.61173 | 417.8241 | 0.187017 | 3.270966 | 0.001072 | 0.006655 | 0.008062 | 325.7876 | 335.3911 | 527.6399 | 482.4778 | 17305 | MFNG O-fucosylpeptide 3-beta-N-acetylglucosaminyltransferase [Source:MGI Symbol;Acc:MGI:1095404] |
| ENSMUSG00000037661 | Gpr160 | -0.60951 | 636.1951 | 0.175527 | 3.472479 | 0.000516 | 0.003562 | 0.004315 | 448.4868 | 558.6574 | 751.681 | 785.9553 | 71862 | G protein-coupled receptor 160 [Source:MGI Symbol;Acc:MGI:1919112] |
| ENSMUSG00000049744 | Arhgap15 | -0.60871 | 442.2659 | 0.18024 | 3.377229 | 0.000732 | 0.004812 | 0.00583 | 341.6539 | 358.9964 | 546.8434 | 521.5698 | 76117 | Rho GTPase activating protein 15 [Source:MGI Symbol;Acc:MGI:1923367] |
| ENSMUSG00000112964 | E430024I08Rik | -0.60761 | 260.8964 | 0.228854 | 2.655016 | 0.00793 | 0.035223 | 0.042543 | 200.9729 | 212.4472 | 308.1709 | 321.9948 | | RIKEN cDNA E430024I08 gene [Source:MGI Symbol;Acc:MGI:3649010] |
| ENSMUSG00000035208 | Slfn8 | -0.60611 | 1024.771 | 0.124346 | 4.874395 | 1.09E-06 | 1.43E-05 | 1.73E-05 | 819.7577 | 805.5288 | 1237.256 | 1236.542 | 276950 | schlafen 8 [Source:MGI Symbol;Acc:MGI:2672859] |
| ENSMUSG00000047514 | Tspyl1 | -0.60375 | 1134.931 | 0.145607 | 4.146416 | 3.38E-05 | 0.000322 | 0.00039 | 844.086 | 956.9958 | 1258.288 | 1480.353 | 22110 | testis-specific protein, Y-encoded-like 1 [Source:MGI Symbol;Acc:MGI:1298395] |
| ENSMUSG00000049421 | Zfp260 | -0.6021 | 894.9481 | 0.140149 | 4.296144 | 1.74E-05 | 0.000176 | 0.000214 | 753.1194 | 668.8152 | 1065.339 | 1092.519 | 26466 | zinc finger protein 260 [Source:MGI Symbol;Acc:MGI:1347071] |
| ENSMUSG00000039531 | Zup1 | -0.60123 | 761.1061 | 0.153352 | 3.92056 | 8.83E-05 | 0.000747 | 0.000905 | 647.3442 | 562.5916 | 944.6308 | 889.8578 | 72580 | zinc finger containing ubiquitin peptidase 1 [Source:MGI Symbol;Acc:MGI:1919830] |
| ENSMUSG00000073489 | Ifi204 | -0.60118 | 3080.97 | 0.089442 | 6.721436 | 1.8E-11 | 5.17E-10 | 6.26E-10 | 2386.288 | 2510.024 | 3794.069 | 3633.5 | 15951 | interferon activated gene 204 [Source:MGI Symbol;Acc:MGI:96429] |
| ENSMUSG00000025375 | Aatk | -0.6004 | 452.111 | 0.190056 | 3.159075 | 0.001583 | 0.009237 | 0.01119 | 342.7116 | 375.7167 | 505.693 | 584.3228 | 11302 | apoptosis-associated tyrosine kinase [Source:MGI Symbol;Acc:MGI:1197518] |
| ENSMUSG00000034959 | Rubcnl | -0.6 | 541.899 | 0.163795 | 3.6631 | 0.000249 | 0.001876 | 0.002273 | 419.9275 | 441.6147 | 653.8345 | 652.2195 | 271221 | RUN and cysteine rich domain containing beclin 1 interacting protein like [Source:MGI Symbol;Acc:MGI:2685590] |
| ENSMUSG00000075595 | Zfp652 | -0.59838 | 2130.184 | 0.110609 | 5.409862 | 6.31E-08 | 1.09E-06 | 1.32E-06 | 1689.23 | 1700.561 | 2736.046 | 2394.901 | 268469 | zinc finger protein 652 [Source:MGI Symbol;Acc:MGI:2442221] |
| ENSMUSG00000052698 | Tln2 | -0.598 | 2634.016 | 0.109611 | 5.45563 | 4.88E-08 | 8.51E-07 | 1.03E-06 | 1988.574 | 2202.172 | 2987.521 | 3357.799 | 70549 | talin 2 [Source:MGI Symbol;Acc:MGI:1917799] |
| ENSMUSG00000060376 | Bckdha | -0.59746 | 775.2428 | 0.183781 | 3.250946 | 0.00115 | 0.007044 | 0.008533 | 521.4717 | 712.0914 | 944.6308 | 922.7774 | 12039 | branched chain ketoacid dehydrogenase E1, alpha polypeptide [Source:MGI Symbol;Acc:MGI:107701] |
| ENSMUSG00000026641 | Usf1 | -0.59611 | 1086.83 | 0.128988 | 4.621442 | 3.81E-06 | 4.44E-05 | 5.38E-05 | 819.7577 | 910.7689 | 1294.867 | 1321.927 | 22278 | upstream transcription factor 1 [Source:MGI Symbol;Acc:MGI:99542] |
| ENSMUSG00000042726 | Trafd1 | -0.59527 | 1975.454 | 0.113112 | 5.262658 | 1.42E-07 | 2.24E-06 | 2.71E-06 | 1462.871 | 1683.84 | 2384.896 | 2370.211 | 231712 | TRAF type zinc finger domain containing 1 [Source:MGI Symbol;Acc:MGI:1923551] |
| ENSMUSG00000036478 | Btg1 | -0.5951 | 3156.37 | 0.090177 | 6.599209 | 4.13E-11 | 1.13E-09 | 1.37E-09 | 2531.2 | 2498.221 | 3935.809 | 3660.247 | 12226 | BTG anti-proliferation factor 1 [Source:MGI Symbol;Acc:MGI:88215] |
| ENSMUSG00000050619 | Zscan29 | -0.59467 | 785.4099 | 0.141705 | 4.196547 | 2.71E-05 | 0.000263 | 0.000318 | 610.3229 | 641.2757 | 969.321 | 920.7199 | 99334 | zinc finger SCAN domains 29 [Source:MGI Symbol;Acc:MGI:2139317] |
| ENSMUSG00000028599 | Tnfrsf1b | -0.59355 | 3385.92 | 0.084001 | 7.065986 | 1.59E-12 | 5.2E-11 | 6.3E-11 | 2708.903 | 2689.03 | 3998.907 | 4146.84 | 21938 | tumor necrosis factor receptor superfamily, member 1b [Source:MGI Symbol;Acc:MGI:1314883] |
| ENSMUSG00000046876 | Atxn1 | -0.59355 | 978.5149 | 0.133417 | 4.448815 | 8.63E-06 | 9.45E-05 | 0.000114 | 776.3899 | 783.8907 | 1234.513 | 1119.266 | 20238 | ataxin 1 [Source:MGI Symbol;Acc:MGI:104783] |
| ENSMUSG00000060224 | Pyroxd2 | -0.59294 | 421.706 | 0.185205 | 3.201547 | 0.001367 | 0.00817 | 0.009897 | 334.2496 | 338.3418 | 527.6399 | 486.5928 | 74580 | pyridine nucleotide-disulphide oxidoreductase domain 2 [Source:MGI Symbol;Acc:MGI:1921830] |
| ENSMUSG00000026199 | Ankzf1 | -0.59265 | 350.8218 | 0.197993 | 2.993289 | 0.00276 | 0.014551 | 0.01762 | 278.1888 | 281.2958 | 417.9054 | 425.8973 | 52231 | ankyrin repeat and zinc finger domain containing 1 [Source:MGI Symbol;Acc:MGI:1098746] |
| ENSMUSG00000002625 | Akap8l | -0.58801 | 703.9548 | 0.163194 | 3.603155 | 0.000314 | 0.002305 | 0.002792 | 509.8364 | 614.7198 | 839.4686 | 851.7945 | 54194 | A kinase anchor protein 8-like [Source:MGI Symbol;Acc:MGI:1860606] |
| ENSMUSG00000074749 | Kiz | -0.58704 | 309.257 | 0.209638 | 2.800276 | 0.005106 | 0.024469 | 0.029567 | 247.5139 | 246.8715 | 371.2683 | 371.3742 | 228730 | kizuna centrosomal protein [Source:MGI Symbol;Acc:MGI:2684960] |
| ENSMUSG00000070327 | Rnf213 | -0.58565 | 16289.47 | 0.074499 | 7.86119 | 3.81E-15 | 1.67E-13 | 2.03E-13 | 12915.15 | 13141.23 | 20445.36 | 18656.15 | 672511 | ring finger protein 213 [Source:MGI Symbol;Acc:MGI:1289196] |
| ENSMUSG00000044465 | Fhip1b | -0.58332 | 380.8162 | 0.20463 | 2.850585 | 0.004364 | 0.02142 | 0.025898 | 295.1128 | 314.7365 | 494.7195 | 418.6961 | 74349 | FHF complex subunit HOOK interacting protein 1B [Source:MGI Symbol;Acc:MGI:1921599] |
| ENSMUSG00000026104 | Stat1 | -0.5833 | 3804.711 | 0.081684 | 7.140863 | 9.27E-13 | 3.1E-11 | 3.75E-11 | 3070.654 | 3021.471 | 4644.511 | 4482.209 | 20846 | signal transducer and activator of transcription 1 [Source:MGI Symbol;Acc:MGI:103063] |
| ENSMUSG00000004207 | Psap | -0.58265 | 78268.67 | 0.087384 | 6.667655 | 2.6E-11 | 7.31E-10 | 8.86E-10 | 57695.08 | 67654.59 | 95404.05 | 92320.95 | 19156 | prosaposin [Source:MGI Symbol;Acc:MGI:97783] |
| ENSMUSG00000027639 | Samhd1 | -0.58062 | 1974.623 | 0.109011 | 5.326225 | 1E-07 | 1.64E-06 | 1.99E-06 | 1514.701 | 1650.4 | 2459.881 | 2273.51 | 56045 | SAM domain and HD domain, 1 [Source:MGI Symbol;Acc:MGI:1927468] |
| ENSMUSG00000041560 | Nop53 | -0.57976 | 1315.853 | 0.119778 | 4.840277 | 1.3E-06 | 1.67E-05 | 2.02E-05 | 1011.211 | 1098.627 | 1617.669 | 1535.905 | 68077 | NOP53 ribosome biogenesis factor [Source:MGI Symbol;Acc:MGI:2154441] |
| ENSMUSG00000059439 | Bcas3 | -0.57866 | 429.3656 | 0.204568 | 2.828713 | 0.004674 | 0.022677 | 0.027417 | 328.9608 | 359.9799 | 569.7047 | 458.8169 | 192197 | BCAS3 microtubule associated cell migration factor [Source:MGI Symbol;Acc:MGI:2385848] |
| ENSMUSG00000022641 | Bbx | -0.57721 | 2394.121 | 0.101552 | 5.683854 | 1.32E-08 | 2.53E-07 | 3.07E-07 | 1977.996 | 1864.814 | 2745.19 | 2988.482 | 70508 | bobby sox HMG box containing [Source:MGI Symbol;Acc:MGI:1917758] |
| ENSMUSG00000035671 | Zswim4 | -0.57602 | 328.4682 | 0.205283 | 2.805958 | 0.005017 | 0.024118 | 0.029151 | 271.8422 | 255.7234 | 392.3007 | 394.0064 | 212168 | zinc finger SWIM-type containing 4 [Source:MGI Symbol;Acc:MGI:2443726] |
| ENSMUSG00000021830 | Txndc16 | -0.57506 | 1654.702 | 0.127968 | 4.493809 | 7E-06 | 7.78E-05 | 9.43E-05 | 1209.01 | 1448.772 | 1931.326 | 2029.699 | 70561 | thioredoxin domain containing 16 [Source:MGI Symbol;Acc:MGI:1917811] |
| ENSMUSG00000048756 | Foxo3 | -0.57493 | 2329.004 | 0.103047 | 5.579309 | 2.41E-08 | 4.42E-07 | 5.35E-07 | 1768.561 | 1973.005 | 2809.202 | 2765.246 | 56484 | forkhead box O3 [Source:MGI Symbol;Acc:MGI:1890081] |
| ENSMUSG00000040111 | Gramd1b | -0.57445 | 2635.692 | 0.120673 | 4.760344 | 1.93E-06 | 2.4E-05 | 2.9E-05 | 1901.838 | 2332.985 | 3105.485 | 3202.459 | 235283 | GRAM domain containing 1B [Source:MGI Symbol;Acc:MGI:1925037] |
| ENSMUSG00000031304 | Il2rg | -0.57141 | 355.6746 | 0.218895 | 2.610416 | 0.009043 | 0.039178 | 0.047324 | 281.362 | 291.1313 | 473.6871 | 376.5179 | 16186 | interleukin 2 receptor, gamma chain [Source:MGI Symbol;Acc:MGI:96551] |
| ENSMUSG00000059498 | Fcgr3 | -0.5707 | 15429.35 | 0.085932 | 6.641343 | 3.11E-11 | 8.62E-10 | 1.04E-09 | 11537.96 | 13294.67 | 18427.16 | 18457.61 | 14131 | Fc receptor, IgG, low affinity III [Source:MGI Symbol;Acc:MGI:95500] |
| ENSMUSG00000004730 | Adgre1 | -0.57062 | 26164.85 | 0.062134 | 9.183696 | 4.17E-20 | 3.22E-18 | 3.9E-18 | 20938.2 | 21174.88 | 30678.1 | 31868.23 | 13733 | adhesion G protein-coupled receptor E1 [Source:MGI Symbol;Acc:MGI:106912] |
| ENSMUSG00000069893 | 9930111J21Rik1 | -0.57025 | 1186.268 | 0.16455 | 3.465509 | 0.000529 | 0.003639 | 0.004409 | 955.15 | 955.0287 | 1624.07 | 1210.824 | 667214 | RIKEN cDNA 9930111J21 gene 1 [Source:MGI Symbol;Acc:MGI:3041173] |
| ENSMUSG00000056144 | Trim34a | -0.56711 | 640.8581 | 0.155857 | 3.638651 | 0.000274 | 0.002039 | 0.00247 | 492.9124 | 539.9699 | 775.4568 | 755.0932 | 94094 | tripartite motif-containing 34A [Source:MGI Symbol;Acc:MGI:2137359] |
| ENSMUSG00000090386 | Mir99ahg | -0.56498 | 968.7964 | 0.14096 | 4.008066 | 6.12E-05 | 0.000543 | 0.000657 | 821.8732 | 741.598 | 1217.138 | 1094.577 | 77994 | Mir99a and Mirlet7c-1 host gene (non-protein coding) [Source:MGI Symbol;Acc:MGI:1919929] |
| ENSMUSG00000056529 | Ptafr | -0.56484 | 2056.843 | 0.110642 | 5.105108 | 3.31E-07 | 4.81E-06 | 5.82E-06 | 1551.722 | 1766.459 | 2493.716 | 2415.475 | 19204 | platelet-activating factor receptor [Source:MGI Symbol;Acc:MGI:106066] |
| ENSMUSG00000046668 | Cxxc5 | -0.56463 | 404.4646 | 0.187572 | 3.010191 | 0.002611 | 0.013905 | 0.016838 | 328.9608 | 323.5885 | 466.3714 | 498.9376 | 67393 | CXXC finger 5 [Source:MGI Symbol;Acc:MGI:1914643] |
| ENSMUSG00000032870 | Smap2 | -0.56429 | 1152.869 | 0.132729 | 4.251413 | 2.12E-05 | 0.000211 | 0.000255 | 996.4023 | 864.5419 | 1401.858 | 1348.675 | 69780 | small ArfGAP 2 [Source:MGI Symbol;Acc:MGI:1917030] |
| ENSMUSG00000027187 | Cat | -0.56377 | 3471.501 | 0.091102 | 6.188298 | 6.08E-10 | 1.41E-08 | 1.71E-08 | 2675.055 | 2928.033 | 4193.685 | 4089.231 | 12359 | catalase [Source:MGI Symbol;Acc:MGI:88271] |
| ENSMUSG00000018474 | Chd3 | -0.56224 | 1141.259 | 0.121184 | 4.639553 | 3.49E-06 | 4.09E-05 | 4.96E-05 | 944.5725 | 898.9662 | 1378.996 | 1342.502 | 216848 | chromodomain helicase DNA binding protein 3 [Source:MGI Symbol;Acc:MGI:1344395] |
| ENSMUSG00000040466 | Blvrb | -0.55952 | 2111.445 | 0.110588 | 5.059544 | 4.2E-07 | 5.98E-06 | 7.25E-06 | 1597.205 | 1816.62 | 2573.273 | 2458.682 | 233016 | biliverdin reductase B [Source:MGI Symbol;Acc:MGI:2385271] |
| ENSMUSG00000023067 | Cdkn1a | -0.55913 | 494.5242 | 0.208165 | 2.685973 | 0.007232 | 0.032607 | 0.039394 | 451.6601 | 348.1773 | 540.4422 | 637.8172 | 12575 | cyclin dependent kinase inhibitor 1A [Source:MGI Symbol;Acc:MGI:104556] |
| ENSMUSG00000042286 | Stab1 | -0.55866 | 148343.4 | 0.074874 | 7.461385 | 8.56E-14 | 3.25E-12 | 3.93E-12 | 118077.9 | 121872.9 | 186402.3 | 167020.7 | 192187 | stabilin 1 [Source:MGI Symbol;Acc:MGI:2178742] |
| ENSMUSG00000037415 | Ranbp10 | -0.55851 | 925.2647 | 0.149804 | 3.728293 | 0.000193 | 0.001487 | 0.001801 | 729.8488 | 767.1703 | 1197.935 | 1006.105 | 74334 | RAN binding protein 10 [Source:MGI Symbol;Acc:MGI:1921584] |
| ENSMUSG00000032741 | Tpcn1 | -0.55827 | 539.776 | 0.212772 | 2.623782 | 0.008696 | 0.03801 | 0.045912 | 437.9093 | 435.7134 | 752.5955 | 532.8859 | 252972 | two pore channel 1 [Source:MGI Symbol;Acc:MGI:2182472] |
| ENSMUSG00000026554 | Dcaf8 | -0.55747 | 2102.784 | 0.114764 | 4.857531 | 1.19E-06 | 1.55E-05 | 1.87E-05 | 1579.224 | 1823.505 | 2580.589 | 2427.82 | 98193 | DDB1 and CUL4 associated factor 8 [Source:MGI Symbol;Acc:MGI:91860] |
| ENSMUSG00000029030 | Tprg1l | -0.55726 | 1056.187 | 0.137267 | 4.059692 | 4.91E-05 | 0.000446 | 0.000541 | 819.7577 | 889.1307 | 1180.56 | 1335.301 | 67808 | transformation related protein 63 regulated 1 like [Source:MGI Symbol;Acc:MGI:1915058] |
| ENSMUSG00000047407 | Tgif1 | -0.55574 | 1461.615 | 0.178372 | 3.115628 | 0.001836 | 0.010412 | 0.012613 | 1280.938 | 1085.841 | 1444.837 | 2034.842 | 21815 | TGFB-induced factor homeobox 1 [Source:MGI Symbol;Acc:MGI:1194497] |
| ENSMUSG00000033880 | Lgals3bp | -0.55489 | 2538.047 | 0.091578 | 6.059167 | 1.37E-09 | 3.03E-08 | 3.67E-08 | 2100.695 | 2011.363 | 3034.158 | 3005.971 | 19039 | lectin, galactoside-binding, soluble, 3 binding protein [Source:MGI Symbol;Acc:MGI:99554] |
| ENSMUSG00000047789 | Slc38a9 | -0.55396 | 686.1573 | 0.150793 | 3.673632 | 0.000239 | 0.001808 | 0.00219 | 558.493 | 553.7396 | 849.5276 | 782.8691 | 268706 | solute carrier family 38, member 9 [Source:MGI Symbol;Acc:MGI:1918839] |
| ENSMUSG00000068457 | Uty | -0.55359 | 1348.48 | 0.123306 | 4.489607 | 7.14E-06 | 7.93E-05 | 9.6E-05 | 1023.904 | 1161.575 | 1623.156 | 1585.284 | 22290 | ubiquitously transcribed tetratricopeptide repeat containing, Y-linked [Source:MGI Symbol;Acc:MGI:894810] |
| ENSMUSG00000068874 | Selenbp1 | -0.55283 | 349.5172 | 0.202987 | 2.723462 | 0.00646 | 0.029655 | 0.035825 | 267.6112 | 298.9997 | 417.9054 | 413.5524 | 20341 | selenium binding protein 1 [Source:MGI Symbol;Acc:MGI:96825] |
| ENSMUSG00000040327 | Cul9 | -0.55242 | 447.1001 | 0.188225 | 2.93486 | 0.003337 | 0.017119 | 0.020723 | 338.4806 | 386.5358 | 554.159 | 509.225 | 78309 | cullin 9 [Source:MGI Symbol;Acc:MGI:1925559] |
| ENSMUSG00000041762 | Gpr155 | -0.55226 | 559.9587 | 0.160246 | 3.446302 | 0.000568 | 0.003871 | 0.004689 | 453.7756 | 454.4009 | 671.2091 | 660.4494 | 68526 | G protein-coupled receptor 155 [Source:MGI Symbol;Acc:MGI:1915776] |
| ENSMUSG00000049734 | Trex1 | -0.55202 | 835.8306 | 0.168532 | 3.275468 | 0.001055 | 0.006566 | 0.007954 | 638.8822 | 717.0092 | 1103.746 | 883.6854 | 22040 | three prime repair exonuclease 1 [Source:MGI Symbol;Acc:MGI:1328317] |
| ENSMUSG00000074829 | 2010315B03Rik | -0.55141 | 503.875 | 0.170473 | 3.234607 | 0.001218 | 0.007406 | 0.008971 | 424.1585 | 393.4207 | 597.1383 | 600.7826 | 630836 | RIKEN cDNA 2010315B03 gene [Source:MGI Symbol;Acc:MGI:1919321] |
| ENSMUSG00000039536 | Stau1 | -0.54976 | 814.6307 | 0.151499 | 3.628826 | 0.000285 | 0.002105 | 0.00255 | 688.5965 | 634.3908 | 1033.333 | 902.2027 | 20853 | staufen double-stranded RNA binding protein 1 [Source:MGI Symbol;Acc:MGI:1338864] |
| ENSMUSG00000121295 | | -0.54889 | 432.1047 | 0.189958 | 2.889529 | 0.003858 | 0.019401 | 0.023463 | 328.9608 | 372.7661 | 534.9555 | 491.7365 | 545551 | cingulin pseudogene [Source:NCBI gene (formerly Entrezgene);Acc:545551] |
| ENSMUSG00000044066 | Cep68 | -0.54652 | 1179.363 | 0.128875 | 4.240665 | 2.23E-05 | 0.00022 | 0.000267 | 1019.673 | 897.9827 | 1419.232 | 1380.566 | 216543 | centrosomal protein 68 [Source:MGI Symbol;Acc:MGI:2667663] |
| ENSMUSG00000020134 | Peli1 | -0.5463 | 1467.655 | 0.108621 | 5.029461 | 4.92E-07 | 6.92E-06 | 8.38E-06 | 1207.953 | 1178.295 | 1753.008 | 1731.365 | 67245 | pellino 1 [Source:MGI Symbol;Acc:MGI:1914495] |
| ENSMUSG00000054843 | Atrnl1 | -0.54583 | 675.8305 | 0.153376 | 3.558803 | 0.000373 | 0.002675 | 0.00324 | 521.4717 | 577.3448 | 809.2916 | 795.214 | 226255 | attractin like 1 [Source:MGI Symbol;Acc:MGI:2147749] |
| ENSMUSG00000052056 | Zfp217 | -0.54286 | 1602.437 | 0.109277 | 4.967773 | 6.77E-07 | 9.25E-06 | 1.12E-05 | 1323.248 | 1285.502 | 1837.138 | 1963.86 | 228913 | zinc finger protein 217 [Source:MGI Symbol;Acc:MGI:2685408] |
| ENSMUSG00000020102 | Slc16a7 | -0.54189 | 1457.909 | 0.11568 | 4.684348 | 2.81E-06 | 3.36E-05 | 4.07E-05 | 1191.029 | 1183.213 | 1647.846 | 1809.549 | 20503 | solute carrier family 16 (monocarboxylic acid transporters), member 7 [Source:MGI Symbol;Acc:MGI:1330284] |
| ENSMUSG00000039621 | Prex1 | -0.54121 | 4212.791 | 0.084516 | 6.403588 | 1.52E-10 | 3.86E-09 | 4.68E-09 | 3532.891 | 3331.29 | 5111.797 | 4875.186 | 277360 | phosphatidylinositol-3,4,5-trisphosphate-dependent Rac exchange factor 1 [Source:MGI Symbol;Acc:MGI:3040696] |
| ENSMUSG00000010307 | Tmem86a | -0.54116 | 2219.815 | 0.112219 | 4.822326 | 1.42E-06 | 1.82E-05 | 2.2E-05 | 1674.421 | 1941.531 | 2590.648 | 2672.66 | 67893 | transmembrane protein 86A [Source:MGI Symbol;Acc:MGI:1915143] |
| ENSMUSG00000060126 | Tpt1 | -0.54114 | 16137.47 | 0.063447 | 8.529084 | 1.48E-17 | 8.25E-16 | 1E-15 | 13307.58 | 12984.85 | 19321.49 | 18935.97 | 22070 | tumor protein, translationally-controlled 1 [Source:MGI Symbol;Acc:MGI:104890] |
| ENSMUSG00000060791 | Gmfg | -0.53962 | 986.5255 | 0.133855 | 4.031411 | 5.54E-05 | 0.000497 | 0.000602 | 774.2744 | 834.0518 | 1214.395 | 1123.381 | 63986 | glia maturation factor, gamma [Source:MGI Symbol;Acc:MGI:1927135] |
| ENSMUSG00000070923 | Klhl9 | -0.53854 | 4951.464 | 0.085467 | 6.301202 | 2.95E-10 | 7.15E-09 | 8.67E-09 | 3845.986 | 4229.272 | 5902.799 | 5827.797 | 242521 | kelch-like 9 [Source:MGI Symbol;Acc:MGI:2180122] |
| ENSMUSG00000035629 | Rubcn | -0.53833 | 1690.36 | 0.124226 | 4.333443 | 1.47E-05 | 0.000152 | 0.000184 | 1259.783 | 1496.966 | 2018.2 | 1986.492 | 100502698 | RUN domain and cysteine-rich domain containing, Beclin 1-interacting protein [Source:MGI Symbol;Acc:MGI:1915160] |
| ENSMUSG00000027488 | Snta1 | -0.53769 | 524.2486 | 0.175104 | 3.070704 | 0.002136 | 0.011771 | 0.014259 | 397.7147 | 457.3515 | 607.1973 | 634.7309 | 20648 | syntrophin, acidic 1 [Source:MGI Symbol;Acc:MGI:101772] |
| ENSMUSG00000030604 | Zfp626 | -0.53736 | 416.7282 | 0.195411 | 2.74991 | 0.005961 | 0.027778 | 0.033556 | 312.0368 | 367.8483 | 506.6074 | 480.4203 | 71163 | zinc finger protein 626 [Source:MGI Symbol;Acc:MGI:1918413] |
| ENSMUSG00000017466 | Timp2 | -0.53606 | 3707.465 | 0.093255 | 5.74833 | 9.01E-09 | 1.77E-07 | 2.14E-07 | 2866.508 | 3185.724 | 4309.821 | 4467.806 | 21858 | tissue inhibitor of metalloproteinase 2 [Source:MGI Symbol;Acc:MGI:98753] |
| ENSMUSG00000044786 | Zfp36 | -0.53345 | 3613.673 | 0.11565 | 4.612647 | 3.98E-06 | 4.62E-05 | 5.59E-05 | 3250.472 | 2656.573 | 4388.464 | 4159.185 | 22695 | zinc finger protein 36 [Source:MGI Symbol;Acc:MGI:99180] |
| ENSMUSG00000013707 | Tnfaip8l2 | -0.53331 | 2077.679 | 0.106473 | 5.008869 | 5.48E-07 | 7.62E-06 | 9.23E-06 | 1603.552 | 1792.031 | 2479.084 | 2436.05 | 69769 | tumor necrosis factor, alpha-induced protein 8-like 2 [Source:MGI Symbol;Acc:MGI:1917019] |
| ENSMUSG00000029455 | Aldh2 | -0.53316 | 4753.661 | 0.096957 | 5.498996 | 3.82E-08 | 6.79E-07 | 8.22E-07 | 3614.338 | 4155.506 | 5748.257 | 5496.544 | 11669 | aldehyde dehydrogenase 2, mitochondrial [Source:MGI Symbol;Acc:MGI:99600] |
| ENSMUSG00000018750 | Zbtb4 | -0.53286 | 963.546 | 0.133389 | 3.994779 | 6.48E-05 | 0.00057 | 0.000691 | 765.8124 | 809.463 | 1183.303 | 1095.605 | 75580 | zinc finger and BTB domain containing 4 [Source:MGI Symbol;Acc:MGI:1922830] |
| ENSMUSG00000028016 | Ints12 | -0.53251 | 721.909 | 0.181446 | 2.934829 | 0.003337 | 0.017119 | 0.020723 | 554.262 | 625.5389 | 748.0232 | 959.8119 | 71793 | integrator complex subunit 12 [Source:MGI Symbol;Acc:MGI:1919043] |
| ENSMUSG00000060510 | Zfp266 | -0.53141 | 1077.713 | 0.124759 | 4.259489 | 2.05E-05 | 0.000204 | 0.000247 | 902.2624 | 860.6077 | 1244.572 | 1303.41 | 77519 | zinc finger protein 266 [Source:MGI Symbol;Acc:MGI:1924769] |
| ENSMUSG00000024187 | Fam234a | -0.53098 | 1643.063 | 0.130863 | 4.057566 | 4.96E-05 | 0.00045 | 0.000545 | 1216.415 | 1471.393 | 2000.825 | 1883.618 | 106581 | family with sequence similarity 234, member A [Source:MGI Symbol;Acc:MGI:2146854] |
| ENSMUSG00000038886 | Man2a2 | -0.53083 | 2861.796 | 0.097278 | 5.456881 | 4.85E-08 | 8.46E-07 | 1.03E-06 | 2218.106 | 2463.797 | 3405.426 | 3359.856 | 140481 | mannosidase 2, alpha 2 [Source:MGI Symbol;Acc:MGI:2150656] |
| ENSMUSG00000027227 | Sord | -0.52982 | 2085.138 | 0.104251 | 5.08212 | 3.73E-07 | 5.37E-06 | 6.5E-06 | 1623.649 | 1789.081 | 2492.801 | 2435.021 | 20322 | sorbitol dehydrogenase [Source:MGI Symbol;Acc:MGI:98266] |
| ENSMUSG00000020091 | Eif4ebp2 | -0.52848 | 1663.686 | 0.159245 | 3.318672 | 0.000904 | 0.005762 | 0.00698 | 1487.199 | 1238.292 | 2218.465 | 1710.79 | 13688 | eukaryotic translation initiation factor 4E binding protein 2 [Source:MGI Symbol;Acc:MGI:109198] |
| ENSMUSG00000040363 | Bcor | -0.52841 | 613.2692 | 0.165473 | 3.193343 | 0.001406 | 0.008343 | 0.010106 | 511.9519 | 492.7594 | 774.5424 | 673.823 | 71458 | BCL6 interacting corepressor [Source:MGI Symbol;Acc:MGI:1918708] |
| ENSMUSG00000051343 | Rab11fip5 | -0.52499 | 3946.851 | 0.119022 | 4.41088 | 1.03E-05 | 0.000111 | 0.000134 | 2902.471 | 3570.293 | 4877.697 | 4436.944 | 52055 | RAB11 family interacting protein 5 (class I) [Source:MGI Symbol;Acc:MGI:1098586] |
| ENSMUSG00000066152 | Slc31a2 | -0.52486 | 830.016 | 0.140696 | 3.730459 | 0.000191 | 0.001477 | 0.001789 | 702.3473 | 658.9796 | 946.4597 | 1012.278 | 20530 | solute carrier family 31, member 2 [Source:MGI Symbol;Acc:MGI:1333844] |
| ENSMUSG00000042408 | Zmym6 | -0.52457 | 551.4814 | 0.161756 | 3.24299 | 0.001183 | 0.007232 | 0.008761 | 444.2558 | 460.3022 | 650.1767 | 651.1907 | 100177 | zinc finger, MYM-type 6 [Source:MGI Symbol;Acc:MGI:106505] |
| ENSMUSG00000058446 | Znrf2 | -0.52329 | 4389.322 | 0.081966 | 6.384195 | 1.72E-10 | 4.31E-09 | 5.22E-09 | 3615.396 | 3587.997 | 5015.779 | 5338.118 | 387524 | zinc and ring finger 2 [Source:MGI Symbol;Acc:MGI:1196246] |
| ENSMUSG00000029501 | Ankle2 | -0.5228 | 2317.272 | 0.094297 | 5.544219 | 2.95E-08 | 5.36E-07 | 6.5E-07 | 1872.221 | 1931.696 | 2778.111 | 2687.062 | 71782 | ankyrin repeat and LEM domain containing 2 [Source:MGI Symbol;Acc:MGI:1261856] |
| ENSMUSG00000059495 | Arhgef12 | -0.5232 | 1168.695 | 0.127559 | 4.094708 | 4.23E-05 | 0.000394 | 0.000478 | 904.3779 | 1014.042 | 1355.221 | 1401.14 | 69632 | Rho guanine nucleotide exchange factor 12 [Source:MGI Symbol;Acc:MGI:1916882] |
| ENSMUSG00000046185 | Zfp84 | -0.52123 | 582.5685 | 0.178919 | 2.913202 | 0.003577 | 0.018124 | 0.021933 | 431.5628 | 525.2166 | 715.1029 | 658.3919 | 74352 | zinc finger protein 84 [Source:MGI Symbol;Acc:MGI:107780] |
| ENSMUSG00000035623 | Rsf1 | -0.52081 | 1181.648 | 0.122181 | 4.262592 | 2.02E-05 | 0.000202 | 0.000244 | 1008.038 | 933.3906 | 1371.681 | 1413.485 | 233532 | remodeling and spacing factor 1 [Source:MGI Symbol;Acc:MGI:2682305] |
| ENSMUSG00000018927 | Ccl6 | -0.52067 | 947.8723 | 0.129591 | 4.017771 | 5.88E-05 | 0.000524 | 0.000634 | 761.5814 | 795.6933 | 1129.35 | 1104.864 | 20305 | C-C motif chemokine ligand 6 [Source:MGI Symbol;Acc:MGI:98263] |
| ENSMUSG00000058761 | Rnf169 | -0.52029 | 1091.787 | 0.128578 | 4.046516 | 5.2E-05 | 0.000469 | 0.000569 | 944.5725 | 849.7887 | 1293.038 | 1279.749 | 108937 | ring finger protein 169 [Source:MGI Symbol;Acc:MGI:1920257] |
| ENSMUSG00000059810 | Rgs3 | -0.51925 | 438.1635 | 0.185296 | 2.802266 | 0.005075 | 0.024365 | 0.02944 | 340.5961 | 379.651 | 530.3832 | 502.0238 | 50780 | regulator of G-protein signaling 3 [Source:MGI Symbol;Acc:MGI:1354734] |
| ENSMUSG00000046897 | Zfp740 | -0.51849 | 1572.106 | 0.114232 | 4.538924 | 5.65E-06 | 6.41E-05 | 7.76E-05 | 1260.84 | 1323.861 | 1767.639 | 1936.084 | 68744 | zinc finger protein 740 [Source:MGI Symbol;Acc:MGI:1915994] |
| ENSMUSG00000019818 | Cd164 | -0.51783 | 4397.172 | 0.101372 | 5.108231 | 3.25E-07 | 4.74E-06 | 5.74E-06 | 3515.967 | 3715.858 | 4794.481 | 5562.383 | 53599 | CD164 antigen [Source:MGI Symbol;Acc:MGI:1859568] |
| ENSMUSG00000020101 | Vsir | -0.5173 | 7584.134 | 0.073544 | 7.033806 | 2.01E-12 | 6.47E-11 | 7.84E-11 | 6066.207 | 6410.79 | 8835.453 | 9024.084 | 74048 | V-set immunoregulatory receptor [Source:MGI Symbol;Acc:MGI:1921298] |
| ENSMUSG00000074024 | 4632427E13Rik | -0.51682 | 744.3628 | 0.164465 | 3.142457 | 0.001675 | 0.009655 | 0.011696 | 643.1132 | 582.2626 | 953.7753 | 798.3002 | | RIKEN cDNA 4632427E13 gene [Source:MGI Symbol;Acc:MGI:1915436] |
| ENSMUSG00000037058 | Paip2 | -0.5168 | 1438.957 | 0.130612 | 3.956731 | 7.6E-05 | 0.000657 | 0.000796 | 1078.907 | 1288.453 | 1672.536 | 1715.934 | 67869 | polyadenylate-binding protein-interacting protein 2 [Source:MGI Symbol;Acc:MGI:1915119] |
| ENSMUSG00000025232 | Hexa | -0.51557 | 24719.79 | 0.071511 | 7.20972 | 5.61E-13 | 1.94E-11 | 2.35E-11 | 19543.02 | 21154.23 | 28525.47 | 29656.44 | 15211 | hexosaminidase A [Source:MGI Symbol;Acc:MGI:96073] |
| ENSMUSG00000032449 | Slc25a36 | -0.51459 | 3129.332 | 0.093382 | 5.510583 | 3.58E-08 | 6.4E-07 | 7.75E-07 | 2685.632 | 2468.715 | 3602.948 | 3760.035 | 192287 | solute carrier family 25, member 36 [Source:MGI Symbol;Acc:MGI:1924909] |
| ENSMUSG00000030560 | Ctsc | -0.51402 | 20929.9 | 0.109231 | 4.705833 | 2.53E-06 | 3.05E-05 | 3.7E-05 | 17459.25 | 17020.36 | 21710.96 | 27529.01 | 13032 | cathepsin C [Source:MGI Symbol;Acc:MGI:109553] |
| ENSMUSG00000000594 | Gm2a | -0.51246 | 1873.907 | 0.105334 | 4.865089 | 1.14E-06 | 1.5E-05 | 1.81E-05 | 1478.737 | 1610.074 | 2216.636 | 2190.182 | 14667 | GM2 ganglioside activator protein [Source:MGI Symbol;Acc:MGI:95762] |
| ENSMUSG00000040415 | Dtx3 | -0.51243 | 517.1851 | 0.193691 | 2.645629 | 0.008154 | 0.036025 | 0.043513 | 372.3287 | 479.9732 | 613.5985 | 602.8401 | 80904 | deltex 3, E3 ubiquitin ligase [Source:MGI Symbol;Acc:MGI:2135752] |
| ENSMUSG00000033540 | Idua | -0.51031 | 776.5861 | 0.145537 | 3.506402 | 0.000454 | 0.003186 | 0.003859 | 629.3624 | 652.0948 | 955.6042 | 869.2831 | 15932 | iduronidase, alpha-L [Source:MGI Symbol;Acc:MGI:96418] |
| ENSMUSG00000029426 | Scarb2 | -0.50975 | 9158.431 | 0.079503 | 6.411668 | 1.44E-10 | 3.67E-09 | 4.45E-09 | 7302.719 | 7810.384 | 10368.08 | 11152.54 | 12492 | scavenger receptor class B, member 2 [Source:MGI Symbol;Acc:MGI:1196458] |
| ENSMUSG00000041058 | Wwp1 | -0.50918 | 10584.19 | 0.074701 | 6.816215 | 9.35E-12 | 2.77E-10 | 3.36E-10 | 8381.626 | 9089.001 | 12532.59 | 12333.53 | 107568 | WW domain containing E3 ubiquitin protein ligase 1 [Source:MGI Symbol;Acc:MGI:1861728] |
| ENSMUSG00000021457 | Syk | -0.50878 | 4577.988 | 0.083506 | 6.09271 | 1.11E-09 | 2.48E-08 | 3E-08 | 3630.205 | 3927.322 | 5375.16 | 5379.268 | 20963 | spleen tyrosine kinase [Source:MGI Symbol;Acc:MGI:99515] |
| ENSMUSG00000001627 | Ifrd1 | -0.50852 | 1483.087 | 0.176519 | 2.880841 | 0.003966 | 0.019848 | 0.024003 | 1398.348 | 1050.433 | 1528.967 | 1954.601 | 15982 | interferon-related developmental regulator 1 [Source:MGI Symbol;Acc:MGI:1316717] |
| ENSMUSG00000014361 | Mertk | -0.50591 | 3309.16 | 0.09845 | 5.138694 | 2.77E-07 | 4.09E-06 | 4.96E-06 | 2625.34 | 2843.448 | 3697.137 | 4070.714 | 17289 | MER proto-oncogene tyrosine kinase [Source:MGI Symbol;Acc:MGI:96965] |
| ENSMUSG00000028842 | Ago3 | -0.50478 | 2847.488 | 0.095803 | 5.268978 | 1.37E-07 | 2.17E-06 | 2.62E-06 | 2464.562 | 2244.465 | 3298.435 | 3382.488 | 214150 | argonaute RISC catalytic subunit 3 [Source:MGI Symbol;Acc:MGI:2446634] |
| ENSMUSG00000020009 | Ifngr1 | -0.50391 | 4657.892 | 0.087398 | 5.765771 | 8.13E-09 | 1.6E-07 | 1.94E-07 | 3731.749 | 3972.565 | 5251.708 | 5675.544 | 15979 | interferon gamma receptor 1 [Source:MGI Symbol;Acc:MGI:107655] |
| ENSMUSG00000052102 | Gnpda1 | -0.50365 | 2296.974 | 0.105666 | 4.766433 | 1.88E-06 | 2.33E-05 | 2.82E-05 | 1783.37 | 2016.281 | 2702.211 | 2686.033 | 26384 | glucosamine-6-phosphate deaminase 1 [Source:MGI Symbol;Acc:MGI:1347054] |
| ENSMUSG00000032238 | Rora | -0.50359 | 449.3982 | 0.186827 | 2.695488 | 0.007029 | 0.031861 | 0.038492 | 345.8849 | 397.3549 | 511.1797 | 543.1733 | 19883 | RAR-related orphan receptor alpha [Source:MGI Symbol;Acc:MGI:104661] |
| ENSMUSG00000025492 | Ifitm3 | -0.50262 | 679.3921 | 0.171322 | 2.933775 | 0.003349 | 0.017166 | 0.02078 | 502.4322 | 621.6047 | 773.6279 | 819.9037 | 66141 | interferon induced transmembrane protein 3 [Source:MGI Symbol;Acc:MGI:1913391] |
| ENSMUSG00000004798 | Ulk2 | -0.50196 | 3757.208 | 0.089872 | 5.585262 | 2.33E-08 | 4.3E-07 | 5.21E-07 | 3034.69 | 3184.74 | 4228.434 | 4580.967 | 29869 | unc-51 like kinase 2 [Source:MGI Symbol;Acc:MGI:1352758] |
| ENSMUSG00000032067 | Pts | -0.50188 | 525.668 | 0.175949 | 2.852444 | 0.004338 | 0.021345 | 0.025806 | 401.9457 | 468.1706 | 624.572 | 607.9838 | 19286 | 6-pyruvoyl-tetrahydropterin synthase [Source:MGI Symbol;Acc:MGI:1338783] |
| ENSMUSG00000020100 | Slc29a3 | -0.50033 | 806.2041 | 0.153867 | 3.251726 | 0.001147 | 0.00703 | 0.008516 | 720.3291 | 615.7034 | 979.38 | 909.4038 | 71279 | solute carrier family 29 (nucleoside transporters), member 3 [Source:MGI Symbol;Acc:MGI:1918529] |
| ENSMUSG00000024773 | Atg2a | -0.49999 | 531.0832 | 0.167614 | 2.982984 | 0.002855 | 0.014961 | 0.018118 | 446.3713 | 433.7463 | 646.5189 | 597.6964 | 329015 | autophagy related 2A [Source:MGI Symbol;Acc:MGI:1916291] |
| ENSMUSG00000053774 | Ubxn7 | -0.49902 | 1384.559 | 0.113463 | 4.398141 | 1.09E-05 | 0.000117 | 0.000141 | 1150.834 | 1143.871 | 1572.861 | 1670.669 | 224111 | UBX domain protein 7 [Source:MGI Symbol;Acc:MGI:2146388] |
| ENSMUSG00000061666 | Gdpd1 | -0.4973 | 651.1765 | 0.158281 | 3.141903 | 0.001679 | 0.009666 | 0.01171 | 516.1829 | 563.5751 | 729.7341 | 795.214 | 66569 | glycerophosphodiester phosphodiesterase domain containing 1 [Source:MGI Symbol;Acc:MGI:1913819] |
| ENSMUSG00000027806 | Tsc22d2 | -0.49722 | 1205.887 | 0.125761 | 3.953716 | 7.69E-05 | 0.000663 | 0.000803 | 1015.442 | 984.5352 | 1336.017 | 1487.554 | 72033 | TSC22 domain family, member 2 [Source:MGI Symbol;Acc:MGI:1919283] |
| ENSMUSG00000118866 | Rn7s1 | -0.49698 | 549000.3 | 0.067449 | 7.368191 | 1.73E-13 | 6.32E-12 | 7.66E-12 | 456984.8 | 453744.9 | 615464.9 | 669806.8 | 103948 | 7S RNA 1 [Source:MGI Symbol;Acc:MGI:97950] |
| ENSMUSG00000020696 | Rffl | -0.49666 | 544.5373 | 0.18897 | 2.628234 | 0.008583 | 0.037625 | 0.045446 | 396.657 | 506.5291 | 647.4333 | 627.5298 | 67338 | ring finger and FYVE like domain containing protein [Source:MGI Symbol;Acc:MGI:1914588] |
| ENSMUSG00000031431 | Tsc22d3 | -0.4963 | 900.2021 | 0.14436 | 3.437944 | 0.000586 | 0.003975 | 0.004815 | 694.943 | 798.644 | 1083.628 | 1023.594 | 14605 | TSC22 domain family, member 3 [Source:MGI Symbol;Acc:MGI:1196284] |
| ENSMUSG00000053477 | Tcf4 | -0.49537 | 3684.323 | 0.094144 | 5.26181 | 1.43E-07 | 2.24E-06 | 2.72E-06 | 2934.204 | 3180.806 | 4136.989 | 4485.295 | 21413 | transcription factor 4 [Source:MGI Symbol;Acc:MGI:98506] |
| ENSMUSG00000002992 | Apoc2 | -0.49521 | 908.9473 | 0.174978 | 2.830147 | 0.004653 | 0.022604 | 0.02733 | 642.0554 | 866.509 | 1088.2 | 1039.025 | 11813 | apolipoprotein C2 [Source:MGI Symbol;Acc:MGI:88054] |
| ENSMUSG00000021676 | Iqgap2 | -0.49506 | 2925.548 | 0.101454 | 4.87966 | 1.06E-06 | 1.4E-05 | 1.69E-05 | 2277.34 | 2578.873 | 3360.618 | 3485.362 | 544963 | IQ motif containing GTPase activating protein 2 [Source:MGI Symbol;Acc:MGI:2449975] |
| ENSMUSG00000039879 | Heca | -0.49484 | 1141.845 | 0.128267 | 3.857902 | 0.000114 | 0.000935 | 0.001132 | 988.998 | 906.8347 | 1284.808 | 1386.738 | 380629 | hdc homolog, cell cycle regulator [Source:MGI Symbol;Acc:MGI:2685715] |
| ENSMUSG00000002103 | Acp2 | -0.49279 | 3375.725 | 0.083683 | 5.88874 | 3.89E-09 | 8.05E-08 | 9.75E-08 | 2852.757 | 2756.895 | 3936.724 | 3956.524 | 11432 | acid phosphatase 2, lysosomal [Source:MGI Symbol;Acc:MGI:87882] |
| ENSMUSG00000038520 | Tbc1d17 | -0.49256 | 963.3042 | 0.146174 | 3.369663 | 0.000753 | 0.004917 | 0.005956 | 753.1194 | 847.8216 | 1198.849 | 1053.427 | 233204 | TBC1 domain family, member 17 [Source:MGI Symbol;Acc:MGI:2449973] |
| ENSMUSG00000112449 | Srp54b | -0.49192 | 494.1066 | 0.185356 | 2.653903 | 0.007957 | 0.035287 | 0.042621 | 418.8698 | 402.2726 | 526.7254 | 628.5585 | 665155 | signal recognition particle 54B [Source:MGI Symbol;Acc:MGI:3714357] |
| ENSMUSG00000031214 | Ophn1 | -0.49188 | 4044.129 | 0.086028 | 5.717628 | 1.08E-08 | 2.1E-07 | 2.54E-07 | 3374.229 | 3348.994 | 4912.446 | 4540.847 | 94190 | oligophrenin 1 [Source:MGI Symbol;Acc:MGI:2151070] |
| ENSMUSG00000029119 | Man2b2 | -0.49183 | 2800.583 | 0.10327 | 4.762554 | 1.91E-06 | 2.37E-05 | 2.87E-05 | 2253.012 | 2402.817 | 3460.293 | 3086.212 | 17160 | mannosidase 2, alpha B2 [Source:MGI Symbol;Acc:MGI:1195262] |
| ENSMUSG00000039852 | Rere | -0.49097 | 892.709 | 0.161223 | 3.045305 | 0.002324 | 0.012615 | 0.015281 | 812.3535 | 672.7494 | 1115.634 | 970.0993 | 68703 | arginine glutamic acid dipeptide (RE) repeats [Source:MGI Symbol;Acc:MGI:2683486] |
| ENSMUSG00000017830 | Dhx58 | -0.48987 | 680.3588 | 0.147844 | 3.313457 | 0.000922 | 0.005849 | 0.007085 | 558.493 | 573.4106 | 805.6338 | 783.8979 | 80861 | DExH-box helicase 58 [Source:MGI Symbol;Acc:MGI:1931560] |
| ENSMUSG00000066442 | Mthfs | -0.48857 | 486.6871 | 0.174768 | 2.795515 | 0.005182 | 0.024754 | 0.029911 | 418.8698 | 391.4536 | 587.0794 | 549.3457 | 107885 | 5, 10-methenyltetrahydrofolate synthetase [Source:MGI Symbol;Acc:MGI:1340032] |
| ENSMUSG00000039501 | Znfx1 | -0.48847 | 3746.804 | 0.081808 | 5.970901 | 2.36E-09 | 5.04E-08 | 6.1E-08 | 3090.751 | 3146.382 | 4447.903 | 4302.18 | 98999 | zinc finger, NFX1-type containing 1 [Source:MGI Symbol;Acc:MGI:2138982] |
| ENSMUSG00000020275 | Rel | -0.48782 | 802.0095 | 0.164696 | 2.961966 | 0.003057 | 0.015873 | 0.019222 | 678.019 | 657.0125 | 833.9819 | 1039.025 | 19696 | reticuloendotheliosis oncogene [Source:MGI Symbol;Acc:MGI:97897] |
| ENSMUSG00000000378 | Ccm2 | -0.48738 | 871.7346 | 0.142058 | 3.430855 | 0.000602 | 0.004064 | 0.004923 | 682.25 | 769.1374 | 998.5836 | 1036.967 | 216527 | cerebral cavernous malformation 2 [Source:MGI Symbol;Acc:MGI:2384924] |
| ENSMUSG00000060802 | B2m | -0.48516 | 24391.35 | 0.06867 | 7.065106 | 1.6E-12 | 5.22E-11 | 6.33E-11 | 20298.26 | 20357.55 | 27396.12 | 29513.45 | 12010 | beta-2 microglobulin [Source:MGI Symbol;Acc:MGI:88127] |
| ENSMUSG00000057367 | Birc2 | -0.48505 | 690.6384 | 0.152099 | 3.189043 | 0.001427 | 0.008455 | 0.010242 | 563.7818 | 587.1804 | 769.0557 | 842.5359 | 11797 | baculoviral IAP repeat-containing 2 [Source:MGI Symbol;Acc:MGI:1197009] |
| ENSMUSG00000089809 | Rasgef1b | -0.48496 | 1219.84 | 0.18215 | 2.662427 | 0.007758 | 0.034588 | 0.041788 | 1066.214 | 966.8313 | 1170.501 | 1675.813 | 320292 | RasGEF domain family, member 1B [Source:MGI Symbol;Acc:MGI:2443755] |
| ENSMUSG00000024308 | Tapbp | -0.48489 | 2209.581 | 0.124143 | 3.905912 | 9.39E-05 | 0.000786 | 0.000952 | 1802.409 | 1881.534 | 2828.406 | 2325.975 | 21356 | TAP binding protein [Source:MGI Symbol;Acc:MGI:1201689] |
| ENSMUSG00000046688 | Tifa | -0.48428 | 2321.126 | 0.092603 | 5.229643 | 1.7E-07 | 2.63E-06 | 3.19E-06 | 1949.437 | 1920.876 | 2683.922 | 2730.269 | 211550 | TRAF-interacting protein with forkhead-associated domain [Source:MGI Symbol;Acc:MGI:2182965] |
| ENSMUSG00000032359 | Ctsh | -0.48408 | 4432.698 | 0.087887 | 5.507949 | 3.63E-08 | 6.48E-07 | 7.85E-07 | 3528.66 | 3862.407 | 5063.331 | 5276.394 | 13036 | cathepsin H [Source:MGI Symbol;Acc:MGI:107285] |
| ENSMUSG00000000275 | Trim25 | -0.4838 | 1136.018 | 0.120511 | 4.014589 | 5.95E-05 | 0.00053 | 0.000642 | 960.4387 | 934.3741 | 1347.905 | 1301.353 | 217069 | tripartite motif-containing 25 [Source:MGI Symbol;Acc:MGI:102749] |
| ENSMUSG00000028995 | Hycc1 | -0.48282 | 1022.885 | 0.132777 | 3.636311 | 0.000277 | 0.002056 | 0.00249 | 837.7395 | 868.4761 | 1130.265 | 1255.06 | 84652 | hyccin PI4KA lipid kinase complex subunit 1 [Source:MGI Symbol;Acc:MGI:2149839] |
| ENSMUSG00000028559 | Osbpl9 | -0.48181 | 3602.758 | 0.096733 | 4.980844 | 6.33E-07 | 8.68E-06 | 1.05E-05 | 2833.717 | 3178.839 | 4092.181 | 4306.295 | 100273 | oxysterol binding protein-like 9 [Source:MGI Symbol;Acc:MGI:1923784] |
| ENSMUSG00000030990 | Pgap2 | -0.48131 | 566.8875 | 0.159934 | 3.00941 | 0.002618 | 0.013926 | 0.016864 | 464.3531 | 481.9403 | 655.6634 | 665.5931 | 233575 | post-GPI attachment to proteins 2 [Source:MGI Symbol;Acc:MGI:2385286] |
| ENSMUSG00000006373 | Pgrmc1 | -0.48015 | 1211.11 | 0.152137 | 3.156021 | 0.001599 | 0.00931 | 0.011277 | 1026.019 | 996.3379 | 1240.914 | 1581.169 | 53328 | progesterone receptor membrane component 1 [Source:MGI Symbol;Acc:MGI:1858305] |
| ENSMUSG00000086841 | 2410006H16Rik | -0.48014 | 514.0277 | 0.167083 | 2.873663 | 0.004057 | 0.020224 | 0.024459 | 434.736 | 423.9108 | 609.0262 | 588.4378 | | RIKEN cDNA 2410006H16 gene [Source:MGI Symbol;Acc:MGI:1916471] |
| ENSMUSG00000010110 | Stx5a | -0.48002 | 1019.863 | 0.132139 | 3.632657 | 0.000281 | 0.00208 | 0.00252 | 859.9523 | 843.8874 | 1249.144 | 1126.467 | 56389 | syntaxin 5A [Source:MGI Symbol;Acc:MGI:1928483] |
| ENSMUSG00000029922 | Mkrn1 | -0.47979 | 1253.87 | 0.119556 | 4.013116 | 5.99E-05 | 0.000533 | 0.000645 | 1012.269 | 1081.907 | 1426.548 | 1494.755 | 54484 | makorin, ring finger protein, 1 [Source:MGI Symbol;Acc:MGI:1859353] |
| ENSMUSG00000069892 | 9930111J21Rik2 | -0.47942 | 2234.604 | 0.104275 | 4.597649 | 4.27E-06 | 4.92E-05 | 5.96E-05 | 1899.722 | 1833.34 | 2469.94 | 2735.413 | 245240 | RIKEN cDNA 9930111J21 gene 2 [Source:MGI Symbol;Acc:MGI:3711310] |
| ENSMUSG00000132705 | Gm60833 | -0.4781 | 874.9962 | 0.166314 | 2.874687 | 0.004044 | 0.020185 | 0.024412 | 722.4446 | 740.6144 | 1143.982 | 892.944 | | predicted gene, 60833 [Source:MGI Symbol;Acc:MGI:7804711] |
| ENSMUSG00000019518 | Ap4m1 | -0.47801 | 493.5868 | 0.176295 | 2.711428 | 0.006699 | 0.03057 | 0.036931 | 388.195 | 436.697 | 565.1325 | 584.3228 | 11781 | adaptor-related protein complex AP-4, mu 1 [Source:MGI Symbol;Acc:MGI:1337063] |
| ENSMUSG00000026896 | Ifih1 | -0.477 | 997.3674 | 0.127409 | 3.743861 | 0.000181 | 0.001408 | 0.001705 | 841.9705 | 826.1834 | 1189.704 | 1131.611 | 71586 | interferon induced with helicase C domain 1 [Source:MGI Symbol;Acc:MGI:1918836] |
| ENSMUSG00000030223 | Ptpro | -0.47586 | 1054.73 | 0.122869 | 3.872901 | 0.000108 | 0.000887 | 0.001074 | 887.4539 | 877.3281 | 1238.17 | 1215.968 | 19277 | protein tyrosine phosphatase receptor type O [Source:MGI Symbol;Acc:MGI:1097152] |
| ENSMUSG00000059142 | Zfp945 | -0.47562 | 478.9821 | 0.172309 | 2.760285 | 0.005775 | 0.027035 | 0.032659 | 406.1767 | 395.3878 | 566.0469 | 548.317 | 240041 | zinc finger protein 945 [Source:MGI Symbol;Acc:MGI:2445132] |
| ENSMUSG00000032580 | Rbm5 | -0.475 | 2667.157 | 0.088338 | 5.377032 | 7.57E-08 | 1.28E-06 | 1.55E-06 | 2240.319 | 2223.81 | 3118.288 | 3086.212 | 83486 | RNA binding motif protein 5 [Source:MGI Symbol;Acc:MGI:1933204] |
| ENSMUSG00000071337 | Tia1 | -0.47479 | 850.1792 | 0.140129 | 3.388251 | 0.000703 | 0.004639 | 0.00562 | 717.1558 | 706.1901 | 1036.076 | 941.2947 | 21841 | cytotoxic granule-associated RNA binding protein 1 [Source:MGI Symbol;Acc:MGI:107914] |
| ENSMUSG00000096210 | H1f0 | -0.47446 | 1384.815 | 0.122236 | 3.881522 | 0.000104 | 0.00086 | 0.001041 | 1087.369 | 1230.423 | 1584.748 | 1636.721 | 14958 | H1.0 linker histone [Source:MGI Symbol;Acc:MGI:95893] |
| ENSMUSG00000036469 | Marchf1 | -0.47312 | 1334.759 | 0.117621 | 4.02238 | 5.76E-05 | 0.000515 | 0.000624 | 1103.235 | 1132.068 | 1489.645 | 1614.089 | 72925 | membrane associated ring-CH-type finger 1 [Source:MGI Symbol;Acc:MGI:1920175] |
| ENSMUSG00000026482 | Rgl1 | -0.4731 | 1856.076 | 0.146776 | 3.223274 | 0.001267 | 0.007656 | 0.009274 | 1347.576 | 1760.558 | 2061.179 | 2254.992 | 19731 | ral guanine nucleotide dissociation stimulator,-like 1 [Source:MGI Symbol;Acc:MGI:107484] |
| ENSMUSG00000020064 | Herc4 | -0.47306 | 1493.13 | 0.10785 | 4.386287 | 1.15E-05 | 0.000123 | 0.000149 | 1258.725 | 1242.226 | 1715.515 | 1756.055 | 67345 | hect domain and RLD 4 [Source:MGI Symbol;Acc:MGI:1914595] |
| ENSMUSG00000031012 | Cask | -0.47209 | 674.8761 | 0.156241 | 3.021539 | 0.002515 | 0.013465 | 0.016306 | 553.2043 | 577.3448 | 739.7931 | 829.1623 | 12361 | calcium/calmodulin dependent serine protein kinase [Source:MGI Symbol;Acc:MGI:1309489] |
| ENSMUSG00000021514 | Zfp369 | -0.47088 | 479.4709 | 0.173742 | 2.710206 | 0.006724 | 0.030664 | 0.037045 | 412.5232 | 391.4536 | 569.7047 | 544.2021 | 170936 | zinc finger protein 369 [Source:MGI Symbol;Acc:MGI:2176229] |
| ENSMUSG00000044712 | Slc38a6 | -0.47017 | 746.1975 | 0.149516 | 3.144626 | 0.001663 | 0.009602 | 0.011632 | 653.6907 | 597.9994 | 899.8226 | 833.2773 | 625098 | solute carrier family 38, member 6 [Source:MGI Symbol;Acc:MGI:3648156] |
| ENSMUSG00000037306 | Man1c1 | -0.47016 | 3612.009 | 0.081942 | 5.73775 | 9.59E-09 | 1.88E-07 | 2.27E-07 | 3009.304 | 3048.027 | 4253.125 | 4137.582 | 230815 | mannosidase, alpha, class 1C, member 1 [Source:MGI Symbol;Acc:MGI:2446214] |
| ENSMUSG00000040997 | Abhd4 | -0.46869 | 1525.968 | 0.140009 | 3.347589 | 0.000815 | 0.005269 | 0.006382 | 1133.91 | 1426.15 | 1795.987 | 1747.825 | 105501 | abhydrolase domain containing 4 [Source:MGI Symbol;Acc:MGI:1915938] |
| ENSMUSG00000021482 | Prxl2c | -0.46818 | 889.0449 | 0.136781 | 3.422854 | 0.00062 | 0.004167 | 0.005048 | 711.867 | 779.9565 | 1023.274 | 1041.082 | 66129 | peroxiredoxin like 2C [Source:MGI Symbol;Acc:MGI:1913379] |
| ENSMUSG00000002957 | Ap2a2 | -0.4676 | 5351.007 | 0.090305 | 5.177938 | 2.24E-07 | 3.39E-06 | 4.1E-06 | 4243.701 | 4738.752 | 6382.888 | 6038.688 | 11772 | adaptor-related protein complex 2, alpha 2 subunit [Source:MGI Symbol;Acc:MGI:101920] |
| ENSMUSG00000055435 | Maf | -0.46733 | 9899.641 | 0.067694 | 6.903553 | 5.07E-12 | 1.59E-10 | 1.93E-10 | 8380.568 | 8240.196 | 11673 | 11304.79 | 17132 | MAF bZIP transcription factor [Source:MGI Symbol;Acc:MGI:96909] |
| ENSMUSG00000043467 | Zbtb37 | -0.46712 | 613.7304 | 0.155369 | 3.006522 | 0.002643 | 0.014025 | 0.016983 | 529.9337 | 500.6278 | 711.4451 | 712.915 | 240869 | zinc finger and BTB domain containing 37 [Source:MGI Symbol;Acc:MGI:2444467] |
| ENSMUSG00000033685 | Ucp2 | -0.46694 | 1720.663 | 0.156021 | 2.992807 | 0.002764 | 0.014569 | 0.017642 | 1658.555 | 1231.407 | 2089.527 | 1903.164 | 22228 | uncoupling protein 2 (mitochondrial, proton carrier) [Source:MGI Symbol;Acc:MGI:109354] |
| ENSMUSG00000029798 | Herc6 | -0.46612 | 616.7893 | 0.166975 | 2.791531 | 0.005246 | 0.025022 | 0.030234 | 552.1465 | 483.9074 | 682.1826 | 748.9208 | 67138 | hect domain and RLD 6 [Source:MGI Symbol;Acc:MGI:1914388] |
| ENSMUSG00000020865 | Abcc3 | -0.46348 | 3710.915 | 0.08701 | 5.32675 | 1E-07 | 1.64E-06 | 1.99E-06 | 3014.593 | 3225.066 | 4382.063 | 4221.938 | 76408 | ATP-binding cassette, sub-family C member 3 [Source:MGI Symbol;Acc:MGI:1923658] |
| ENSMUSG00000040522 | Tlr8 | -0.46335 | 2112.669 | 0.10774 | 4.300666 | 1.7E-05 | 0.000173 | 0.00021 | 1714.616 | 1837.275 | 2330.028 | 2568.757 | 170744 | toll-like receptor 8 [Source:MGI Symbol;Acc:MGI:2176887] |
| ENSMUSG00000057191 | AB124611 | -0.4629 | 948.1587 | 0.134048 | 3.453215 | 0.000554 | 0.003787 | 0.004587 | 758.4081 | 836.0189 | 1096.43 | 1101.778 | 382062 | cDNA sequence AB124611 [Source:MGI Symbol;Acc:MGI:3043001] |
| ENSMUSG00000026349 | Ccnt2 | -0.46282 | 1227.094 | 0.117463 | 3.940697 | 8.12E-05 | 0.000694 | 0.000841 | 1058.81 | 1005.019 | 1424.719 | 1419.658 | 72949 | cyclin T2 [Source:MGI Symbol;Acc:MGI:1920199] |
| ENSMUSG00000020876 | Snx11 | -0.46194 | 597.9548 | 0.165793 | 2.786257 | 0.005332 | 0.025369 | 0.030654 | 470.6996 | 535.0521 | 671.2091 | 713.9437 | 74479 | sorting nexin 11 [Source:MGI Symbol;Acc:MGI:1921729] |
| ENSMUSG00000031938 | 4931406C07Rik | -0.46095 | 864.0974 | 0.143825 | 3.204957 | 0.001351 | 0.008086 | 0.009795 | 715.0403 | 739.6309 | 1063.51 | 938.2085 | 70984 | RIKEN cDNA 4931406C07 gene [Source:MGI Symbol;Acc:MGI:1918234] |
| ENSMUSG00000031652 | N4bp1 | -0.46042 | 1151.135 | 0.129047 | 3.567851 | 0.00036 | 0.002598 | 0.003147 | 1008.038 | 930.4399 | 1394.542 | 1271.519 | 80750 | NEDD4 binding protein 1 [Source:MGI Symbol;Acc:MGI:2136825] |
| ENSMUSG00000031770 | Herpud1 | -0.46013 | 1191.796 | 0.121683 | 3.781406 | 0.000156 | 0.001232 | 0.001493 | 1042.943 | 963.8807 | 1364.365 | 1395.997 | 64209 | homocysteine-inducible, endoplasmic reticulum stress-inducible, ubiquitin-like domain member 1 [Source:MGI Symbol;Acc:MGI:1927406] |
| ENSMUSG00000052837 | Junb | -0.46007 | 9187.845 | 0.102045 | 4.508506 | 6.53E-06 | 7.28E-05 | 8.82E-05 | 7883.425 | 7586.134 | 9636.514 | 11645.31 | 16477 | jun B proto-oncogene [Source:MGI Symbol;Acc:MGI:96647] |
| ENSMUSG00000026014 | Raph1 | -0.45971 | 1879.074 | 0.105093 | 4.374314 | 1.22E-05 | 0.000129 | 0.000156 | 1649.035 | 1515.653 | 2164.512 | 2187.096 | 77300 | Ras association (RalGDS/AF-6) and pleckstrin homology domains 1 [Source:MGI Symbol;Acc:MGI:1924550] |
| ENSMUSG00000037111 | Setd7 | -0.45953 | 822.2047 | 0.142044 | 3.235127 | 0.001216 | 0.007405 | 0.00897 | 666.3837 | 717.9927 | 919.9405 | 984.5016 | 73251 | SET domain containing (lysine methyltransferase) 7 [Source:MGI Symbol;Acc:MGI:1920501] |
| ENSMUSG00000040651 | Tasor | -0.45881 | 1822.767 | 0.100795 | 4.551887 | 5.32E-06 | 6.06E-05 | 7.34E-05 | 1541.145 | 1529.423 | 2077.639 | 2142.86 | 218850 | transcription activation suppressor [Source:MGI Symbol;Acc:MGI:1921694] |
| ENSMUSG00000034616 | Ssh3 | -0.45862 | 1476.751 | 0.118542 | 3.868828 | 0.000109 | 0.000899 | 0.001089 | 1178.336 | 1309.107 | 1662.477 | 1757.083 | 245857 | slingshot protein phosphatase 3 [Source:MGI Symbol;Acc:MGI:2683546] |
| ENSMUSG00000017707 | Serinc3 | -0.45852 | 33615.62 | 0.062832 | 7.297533 | 2.93E-13 | 1.05E-11 | 1.27E-11 | 27688.77 | 28946.91 | 38476.56 | 39350.23 | 26943 | serine incorporator 3 [Source:MGI Symbol;Acc:MGI:1349457] |
| ENSMUSG00000005043 | Sgsh | -0.45738 | 1185.476 | 0.118505 | 3.859559 | 0.000114 | 0.00093 | 0.001126 | 992.1713 | 1006.173 | 1400.029 | 1343.531 | 27029 | N-sulfoglucosamine sulfohydrolase (sulfamidase) [Source:MGI Symbol;Acc:MGI:1350341] |
| ENSMUSG00000029998 | Pcyox1 | -0.45657 | 4130.943 | 0.089653 | 5.0926 | 3.53E-07 | 5.11E-06 | 6.18E-06 | 3350.958 | 3613.569 | 4616.163 | 4943.083 | 66881 | prenylcysteine oxidase 1 [Source:MGI Symbol;Acc:MGI:1914131] |
| ENSMUSG00000026923 | Notch1 | -0.45581 | 1002.592 | 0.132324 | 3.44461 | 0.000572 | 0.00389 | 0.004713 | 818.7 | 872.4104 | 1206.165 | 1113.094 | 18128 | notch 1 [Source:MGI Symbol;Acc:MGI:97363] |
| ENSMUSG00000042599 | Kdm7a | -0.4554 | 2096.014 | 0.104823 | 4.344481 | 1.4E-05 | 0.000145 | 0.000176 | 1759.041 | 1776.294 | 2302.595 | 2546.125 | 338523 | lysine (K)-specific demethylase 7A [Source:MGI Symbol;Acc:MGI:2443388] |
| ENSMUSG00000022051 | Bnip3l | -0.45446 | 1743.291 | 0.103915 | 4.373436 | 1.22E-05 | 0.000129 | 0.000157 | 1455.467 | 1486.147 | 1967.905 | 2063.647 | 12177 | BCL2/adenovirus E1B interacting protein 3-like [Source:MGI Symbol;Acc:MGI:1332659] |
| ENSMUSG00000037369 | Kdm6a | -0.45288 | 872.0378 | 0.134576 | 3.365273 | 0.000765 | 0.004985 | 0.006039 | 715.0403 | 757.3348 | 1000.412 | 1015.364 | 22289 | lysine (K)-specific demethylase 6A [Source:MGI Symbol;Acc:MGI:1095419] |
| ENSMUSG00000046879 | Irgm1 | -0.45264 | 1236.681 | 0.115801 | 3.908791 | 9.28E-05 | 0.000778 | 0.000942 | 1028.135 | 1060.269 | 1428.377 | 1429.945 | 15944 | immunity-related GTPase family M member 1 [Source:MGI Symbol;Acc:MGI:107567] |
| ENSMUSG00000005580 | Adcy9 | -0.45223 | 592.3095 | 0.162102 | 2.789796 | 0.005274 | 0.025148 | 0.030387 | 489.7391 | 510.4633 | 652.0056 | 717.0299 | 11515 | adenylate cyclase 9 [Source:MGI Symbol;Acc:MGI:108450] |
| ENSMUSG00000085438 | Oip5os1 | -0.45127 | 3604.035 | 0.090969 | 4.960645 | 7.03E-07 | 9.57E-06 | 1.16E-05 | 3088.636 | 3000.816 | 3975.131 | 4351.559 | | Opa interacting protein 5, opposite strand 1 [Source:MGI Symbol;Acc:MGI:1913852] |
| ENSMUSG00000028522 | Mier1 | -0.45088 | 1391.918 | 0.112307 | 4.01473 | 5.95E-05 | 0.00053 | 0.000642 | 1191.029 | 1161.575 | 1644.188 | 1570.882 | 71148 | MEIR1 treanscription regulator [Source:MGI Symbol;Acc:MGI:1918398] |
| ENSMUSG00000026932 | Nacc2 | -0.45073 | 1325.647 | 0.145879 | 3.089788 | 0.002003 | 0.011141 | 0.013496 | 987.9403 | 1252.061 | 1539.026 | 1523.56 | 67991 | nucleus accumbens associated 2, BEN and BTB (POZ) domain containing [Source:MGI Symbol;Acc:MGI:1915241] |
| ENSMUSG00000140119 | Gm73145 | -0.45059 | 938.9452 | 0.139803 | 3.223051 | 0.001268 | 0.007659 | 0.009278 | 749.9461 | 837.0025 | 1125.693 | 1043.14 | | predicted gene, 73145 [Source:MGI Symbol;Acc:MGI:7829048] |
| ENSMUSG00000057133 | Chd6 | -0.45059 | 1306.924 | 0.11387 | 3.957086 | 7.59E-05 | 0.000656 | 0.000795 | 1108.524 | 1100.594 | 1537.197 | 1481.382 | 71389 | chromodomain helicase DNA binding protein 6 [Source:MGI Symbol;Acc:MGI:1918639] |
| ENSMUSG00000025154 | Arhgap19 | -0.44965 | 6940.853 | 0.077243 | 5.821201 | 5.84E-09 | 1.17E-07 | 1.42E-07 | 5822.924 | 5912.129 | 7724.391 | 8303.968 | 71085 | Rho GTPase activating protein 19 [Source:MGI Symbol;Acc:MGI:1918335] |
| ENSMUSG00000021242 | Npc2 | -0.4496 | 12573.32 | 0.078566 | 5.722578 | 1.05E-08 | 2.04E-07 | 2.47E-07 | 10320.49 | 10938.08 | 13881.41 | 15153.3 | 67963 | NPC intracellular cholesterol transporter 2 [Source:MGI Symbol;Acc:MGI:1915213] |
| ENSMUSG00000030330 | Ing4 | -0.44899 | 1276.246 | 0.113918 | 3.941341 | 8.1E-05 | 0.000693 | 0.000839 | 1088.427 | 1070.104 | 1472.271 | 1474.181 | 28019 | inhibitor of growth family, member 4 [Source:MGI Symbol;Acc:MGI:107307] |
| ENSMUSG00000049866 | Arl4c | -0.44869 | 5234.715 | 0.108017 | 4.153926 | 3.27E-05 | 0.000312 | 0.000378 | 4126.29 | 4726.949 | 5613.832 | 6471.787 | 320982 | ADP-ribosylation factor-like 4C [Source:MGI Symbol;Acc:MGI:2445172] |
| ENSMUSG00000005886 | Ncoa2 | -0.44849 | 1267.194 | 0.117738 | 3.809204 | 0.000139 | 0.001117 | 0.001353 | 1090.542 | 1053.384 | 1505.191 | 1419.658 | 17978 | nuclear receptor coactivator 2 [Source:MGI Symbol;Acc:MGI:1276533] |
| ENSMUSG00000025464 | Paox | -0.44849 | 1298.645 | 0.127375 | 3.52099 | 0.00043 | 0.003041 | 0.003683 | 1094.773 | 1101.578 | 1395.457 | 1602.773 | 212503 | polyamine oxidase (exo-N4-amino) [Source:MGI Symbol;Acc:MGI:1916983] |
| ENSMUSG00000041935 | Rmc1 | -0.44812 | 1076.713 | 0.123361 | 3.632608 | 0.000281 | 0.00208 | 0.00252 | 926.5907 | 895.032 | 1219.881 | 1265.347 | 106064 | RAB7A interacting MON1-CCZ1 complex subunit 1 [Source:MGI Symbol;Acc:MGI:2146232] |

| | | | | | | | | | | | | | | |
|---|---|---|---|---|---|---|---|---|---|---|---|---|---|---|
| ENSMUSG00000021775 | Nr1d2 | -0.44742 | 1060.757 | 0.143444 | 3.119118 | 0.001814 | 0.010309 | 0.012488 | 821.8732 | 972.7326 | 1176.902 | 1271.519 | 353187 | nuclear receptor subfamily 1, group D, member 2 [Source:MGI Symbol;Acc:MGI:2449205] |
| ENSMUSG00000036667 | Tcaf1 | -0.44702 | 551.1418 | 0.16371 | 2.730557 | 0.006323 | 0.02914 | 0.035203 | 452.7178 | 479.9732 | 622.7431 | 649.1333 | 77574 | TRPM8 channel-associated factor 1 [Source:MGI Symbol;Acc:MGI:1914665] |
| ENSMUSG00000059474 | Mbtd1 | -0.4453 | 722.8576 | 0.148067 | 3.007456 | 0.002634 | 0.013997 | 0.016949 | 583.8791 | 640.2922 | 833.9819 | 833.2773 | 103537 | mbt domain containing 1 [Source:MGI Symbol;Acc:MGI:2143977] |
| ENSMUSG00000025575 | Cant1 | -0.44505 | 1029.797 | 0.130063 | 3.421766 | 0.000622 | 0.004182 | 0.005066 | 844.086 | 899.9498 | 1145.811 | 1229.341 | 76025 | calcium activated nucleotidase 1 [Source:MGI Symbol;Acc:MGI:1923275] |
| ENSMUSG00000031666 | Rbl2 | -0.44432 | 1453.817 | 0.112572 | 3.947009 | 7.91E-05 | 0.000679 | 0.000822 | 1238.627 | 1224.522 | 1617.669 | 1734.451 | 19651 | RB transcriptional corepressor like 2 [Source:MGI Symbol;Acc:MGI:105085] |
| ENSMUSG00000038437 | Mllt6 | -0.44377 | 1008.652 | 0.128468 | 3.454341 | 0.000552 | 0.003773 | 0.00457 | 827.162 | 882.2459 | 1183.303 | 1141.898 | 246198 | myeloid/lymphoid or mixed-lineage leukemia; translocated to, 6 [Source:MGI Symbol;Acc:MGI:1935145] |
| ENSMUSG00000090733 | Rps27 | -0.44024 | 4185.635 | 0.082061 | 5.364754 | 8.11E-08 | 1.36E-06 | 1.64E-06 | 3464.138 | 3639.141 | 4741.443 | 4897.819 | 57294 | ribosomal protein S27 [Source:MGI Symbol;Acc:MGI:1888676] |
| ENSMUSG00000038170 | Pde4dip | -0.44019 | 928.2478 | 0.129388 | 3.402122 | 0.000669 | 0.004441 | 0.00538 | 782.7364 | 792.7427 | 1079.97 | 1057.542 | 83679 | phosphodiesterase 4D interacting protein (myomegalin) [Source:MGI Symbol;Acc:MGI:1891434] |
| ENSMUSG00000030921 | Trim30a | -0.43628 | 3284.498 | 0.086863 | 5.022619 | 5.1E-07 | 7.16E-06 | 8.67E-06 | 2721.596 | 2861.152 | 3719.084 | 3836.162 | 20128 | tripartite motif-containing 30A [Source:MGI Symbol;Acc:MGI:98178] |
| ENSMUSG00000024013 | Fgd2 | -0.43535 | 2026.64 | 0.109677 | 3.969349 | 7.21E-05 | 0.000625 | 0.000757 | 1622.591 | 1823.505 | 2384.896 | 2275.567 | 26382 | FYVE, RhoGEF and PH domain containing 2 [Source:MGI Symbol;Acc:MGI:1347084] |
| ENSMUSG00000031447 | Lamp1 | -0.4342 | 40823.76 | 0.073192 | 5.932339 | 2.99E-09 | 6.28E-08 | 7.61E-08 | 33191.2 | 36260.6 | 45744.64 | 48098.61 | 16783 | lysosomal-associated membrane protein 1 [Source:MGI Symbol;Acc:MGI:96745] |
| ENSMUSG00000027634 | Ndrg3 | -0.4333 | 581.8285 | 0.164339 | 2.63663 | 0.008373 | 0.036823 | 0.044477 | 469.6418 | 520.2988 | 650.1767 | 687.1966 | 29812 | N-myc downstream regulated gene 3 [Source:MGI Symbol;Acc:MGI:1352499] |
| ENSMUSG00000047875 | Gpr157 | -0.43208 | 991.2046 | 0.134266 | 3.218093 | 0.00129 | 0.007771 | 0.009413 | 795.4294 | 892.0814 | 1146.725 | 1130.582 | 269604 | G protein-coupled receptor 157 [Source:MGI Symbol;Acc:MGI:2442046] |
| ENSMUSG00000028497 | Hacd4 | -0.43189 | 5513.998 | 0.080646 | 5.355455 | 8.53E-08 | 1.42E-06 | 1.72E-06 | 4513.427 | 4875.466 | 6295.1 | 6371.999 | 66775 | 3-hydroxyacyl-CoA dehydratase 4 [Source:MGI Symbol;Acc:MGI:1914025] |
| ENSMUSG00000063268 | Parp10 | -0.43185 | 678.7341 | 0.162973 | 2.649861 | 0.008052 | 0.035639 | 0.043047 | 533.107 | 622.5882 | 809.2916 | 749.9495 | 671535 | poly (ADP-ribose) polymerase family, member 10 [Source:MGI Symbol;Acc:MGI:3712326] |
| ENSMUSG00000027215 | Cd82 | -0.43177 | 617.8021 | 0.156134 | 2.765369 | 0.005686 | 0.026684 | 0.032234 | 542.6267 | 509.4798 | 695.8994 | 723.2024 | 12521 | CD82 antigen [Source:MGI Symbol;Acc:MGI:104651] |
| ENSMUSG00000026457 | Adipor1 | -0.43092 | 7221.389 | 0.079269 | 5.436203 | 5.44E-08 | 9.42E-07 | 1.14E-06 | 5979.472 | 6321.287 | 8013.359 | 8571.44 | 72674 | adiponectin receptor 1 [Source:MGI Symbol;Acc:MGI:1919924] |
| ENSMUSG00000027695 | Pld1 | -0.4303 | 935.0814 | 0.136436 | 3.15385 | 0.001611 | 0.00935 | 0.011327 | 788.0252 | 805.5288 | 1129.35 | 1017.421 | 18805 | phospholipase D1 [Source:MGI Symbol;Acc:MGI:109585] |
| ENSMUSG00000088185 | Scarna2 | -0.42908 | 765.7883 | 0.147295 | 2.913074 | 0.003579 | 0.018125 | 0.021934 | 630.4201 | 674.7165 | 840.3831 | 917.6337 | | small Cajal body-specific RNA 2 [Source:MGI Symbol;Acc:MGI:3819484] |
| ENSMUSG00000056579 | Tug1 | -0.4289 | 2918.189 | 0.097197 | 4.412648 | 1.02E-05 | 0.00011 | 0.000133 | 2553.413 | 2421.504 | 3196.016 | 3501.822 | 544752 | taurine upregulated gene 1 [Source:MGI Symbol;Acc:MGI:2144114] |
| ENSMUSG00000037652 | Phc3 | -0.42856 | 1804.214 | 0.106154 | 4.037135 | 5.41E-05 | 0.000486 | 0.000589 | 1530.567 | 1546.143 | 2152.624 | 1987.521 | 241915 | polyhomeotic 3 [Source:MGI Symbol;Acc:MGI:2181434] |
| ENSMUSG00000060261 | Gtf2i | -0.42742 | 2113.497 | 0.097954 | 4.363451 | 1.28E-05 | 0.000135 | 0.000163 | 1851.066 | 1754.656 | 2446.164 | 2402.102 | 14886 | general transcription factor II I [Source:MGI Symbol;Acc:MGI:1202722] |
| ENSMUSG00000018001 | Cyth3 | -0.42605 | 1106.546 | 0.124283 | 3.42809 | 0.000608 | 0.004098 | 0.004964 | 910.7244 | 977.6504 | 1251.887 | 1285.922 | 19159 | cytohesin 3 [Source:MGI Symbol;Acc:MGI:1335107] |
| ENSMUSG00000056216 | Cebpg | -0.42515 | 1655.283 | 0.105948 | 4.012827 | 6E-05 | 0.000533 | 0.000646 | 1377.193 | 1448.772 | 1875.545 | 1919.624 | 12611 | CCAAT/enhancer binding protein gamma [Source:MGI Symbol;Acc:MGI:104982] |
| ENSMUSG00000003847 | Nfat5 | -0.42284 | 1103.873 | 0.135429 | 3.122253 | 0.001795 | 0.010211 | 0.01237 | 990.0558 | 896.9991 | 1335.103 | 1193.335 | 54446 | nuclear factor of activated T cells 5 [Source:MGI Symbol;Acc:MGI:1859333] |
| ENSMUSG00000021411 | Pxdc1 | -0.42282 | 893.9992 | 0.143139 | 2.95388 | 0.003138 | 0.016245 | 0.019672 | 742.5418 | 784.8743 | 960.1765 | 1088.404 | 66895 | PX domain containing 1 [Source:MGI Symbol;Acc:MGI:1914145] |
| ENSMUSG00000066026 | Dhrs3 | -0.42193 | 4692.776 | 0.098107 | 4.300691 | 1.7E-05 | 0.000173 | 0.00021 | 3723.287 | 4299.104 | 5494.953 | 5253.762 | 20148 | dehydrogenase/reductase 3 [Source:MGI Symbol;Acc:MGI:1315215] |
| ENSMUSG00000045817 | Zfp36l2 | -0.42171 | 2396.575 | 0.129367 | 3.259777 | 0.001115 | 0.006873 | 0.008325 | 2229.741 | 1868.748 | 2950.942 | 2536.866 | 12193 | zinc finger protein 36, C3H type-like 2 [Source:MGI Symbol;Acc:MGI:107945] |
| ENSMUSG00000059555 | Tor4a | -0.42103 | 1825.2 | 0.099891 | 4.214938 | 2.5E-05 | 0.000244 | 0.000295 | 1565.473 | 1555.979 | 2077.639 | 2101.71 | 227612 | torsin family 4, member A [Source:MGI Symbol;Acc:MGI:2442720] |
| ENSMUSG00000068479 | Mfap1a | -0.4195 | 688.9999 | 0.147179 | 2.850262 | 0.004368 | 0.021429 | 0.025908 | 583.8791 | 595.0488 | 773.6279 | 803.4439 | 67532 | microfibrillar-associated protein 1A [Source:MGI Symbol;Acc:MGI:1914782] |
| ENSMUSG00000013622 | Atraid | -0.41895 | 870.7719 | 0.140526 | 2.98132 | 0.00287 | 0.015033 | 0.018204 | 702.3473 | 787.8249 | 994.0113 | 998.904 | 381629 | all-trans retinoic acid induced differentiation factor [Source:MGI Symbol;Acc:MGI:1918918] |
| ENSMUSG00000029461 | Fam168a | -0.41888 | 795.4196 | 0.138535 | 3.023626 | 0.002498 | 0.013391 | 0.016216 | 674.8457 | 686.5191 | 893.4214 | 926.8924 | 319604 | family with sequence similarity 168, member A [Source:MGI Symbol;Acc:MGI:2442372] |
| ENSMUSG00000022884 | Eif4a2 | -0.41884 | 5706.984 | 0.073447 | 5.702678 | 1.18E-08 | 2.28E-07 | 2.76E-07 | 4846.619 | 4921.693 | 6455.13 | 6604.494 | 13682 | eukaryotic translation initiation factor 4A2 [Source:MGI Symbol;Acc:MGI:106906] |
| ENSMUSG00000024299 | Adamts10 | -0.41703 | 780.9565 | 0.141807 | 2.940829 | 0.003273 | 0.016836 | 0.020381 | 644.1709 | 693.4039 | 894.3358 | 891.9153 | 224697 | ADAM metallopeptidase with thrombospondin type 1 motif 10 [Source:MGI Symbol;Acc:MGI:2449112] |
| ENSMUSG00000028552 | Eps15 | -0.41659 | 5002.28 | 0.084468 | 4.931928 | 8.14E-07 | 1.1E-05 | 1.33E-05 | 4405.537 | 4164.358 | 5517.814 | 5921.412 | 13858 | epidermal growth factor receptor pathway substrate 15 [Source:MGI Symbol;Acc:MGI:104583] |
| ENSMUSG00000022228 | Zscan26 | -0.41604 | 1230.842 | 0.12015 | 3.462662 | 0.000535 | 0.003675 | 0.004451 | 1031.308 | 1077.973 | 1453.067 | 1361.02 | 432731 | zinc finger and SCAN domain containing 26 [Source:MGI Symbol;Acc:MGI:3531417] |
| ENSMUSG00000060147 | Serpinb6a | -0.41512 | 9139.571 | 0.087278 | 4.756337 | 1.97E-06 | 2.44E-05 | 2.96E-05 | 7314.354 | 8352.321 | 10544.57 | 10347.04 | 20719 | serine (or cysteine) peptidase inhibitor, clade B, member 6a [Source:MGI Symbol;Acc:MGI:103123] |
| ENSMUSG00000038371 | Sbf2 | -0.4148 | 3818.844 | 0.081331 | 5.100101 | 3.39E-07 | 4.92E-06 | 5.96E-06 | 3266.338 | 3281.128 | 4444.246 | 4283.662 | 319934 | SET binding factor 2 [Source:MGI Symbol;Acc:MGI:1921831] |
| ENSMUSG00000022564 | Grina | -0.41414 | 2113 | 0.108317 | 3.823448 | 0.000132 | 0.001063 | 0.001287 | 1710.385 | 1913.008 | 2479.999 | 2348.607 | 66168 | glutamate receptor, ionotropic, N-methyl D-aspartate-associated protein 1 (glutamate binding) [Source:MGI Symbol;Acc:MGI:1913418] |
| ENSMUSG00000037257 | Aagab | -0.41399 | 2163.01 | 0.101576 | 4.075656 | 4.59E-05 | 0.000422 | 0.000512 | 1781.254 | 1927.761 | 2420.559 | 2522.464 | 66939 | alpha- and gamma-adaptin binding protein [Source:MGI Symbol;Acc:MGI:1914189] |
| ENSMUSG00000030123 | Plxnd1 | -0.41387 | 931.8941 | 0.154543 | 2.678026 | 0.007406 | 0.033262 | 0.040186 | 806.007 | 792.7427 | 1174.159 | 954.6683 | 67784 | plexin D1 [Source:MGI Symbol;Acc:MGI:2154244] |
| ENSMUSG00000071637 | Cebpd | -0.41308 | 607.678 | 0.155462 | 2.657127 | 0.007881 | 0.035033 | 0.042327 | 533.107 | 509.4798 | 688.5837 | 699.5414 | 12609 | CCAAT/enhancer binding protein delta [Source:MGI Symbol;Acc:MGI:103573] |
| ENSMUSG00000015291 | Gdi1 | -0.41294 | 959.5045 | 0.140753 | 2.933808 | 0.003348 | 0.017166 | 0.02078 | 762.6391 | 883.2294 | 1087.286 | 1104.864 | 14567 | GDP dissociation inhibitor 1 [Source:MGI Symbol;Acc:MGI:99846] |
| ENSMUSG00000022553 | Maf1 | -0.41208 | 1301.227 | 0.120389 | 3.422863 | 0.00062 | 0.004167 | 0.005048 | 1068.329 | 1164.525 | 1448.495 | 1523.56 | 68877 | MAF1 homolog, negative regulator of RNA polymerase III [Source:MGI Symbol;Acc:MGI:1916127] |
| ENSMUSG00000045092 | S1pr1 | -0.41057 | 3060.096 | 0.102811 | 3.993422 | 6.51E-05 | 0.000573 | 0.000694 | 2457.158 | 2797.221 | 3400.854 | 3585.15 | 13609 | sphingosine-1-phosphate receptor 1 [Source:MGI Symbol;Acc:MGI:1096355] |
| ENSMUSG00000022901 | Cd86 | -0.41051 | 718.769 | 0.150037 | 2.736089 | 0.006217 | 0.028707 | 0.03468 | 646.2864 | 588.1639 | 800.1471 | 840.4784 | 12524 | CD86 antigen [Source:MGI Symbol;Acc:MGI:101773] |
| ENSMUSG00000032405 | Pias1 | -0.41034 | 1217.722 | 0.122279 | 3.355788 | 0.000791 | 0.005144 | 0.006231 | 996.4023 | 1094.693 | 1381.74 | 1398.054 | 56469 | protein inhibitor of activated STAT 1 [Source:MGI Symbol;Acc:MGI:1913125] |
| ENSMUSG00000045868 | Gvin1 | -0.40886 | 2284.791 | 0.0948 | 4.31293 | 1.61E-05 | 0.000166 | 0.000201 | 1919.82 | 2006.445 | 2631.798 | 2581.102 | 74558 | GTPase, very large interferon inducible 1 [Source:MGI Symbol;Acc:MGI:1921808] |
| ENSMUSG00000026656 | Fcgr2b | -0.40848 | 6074.204 | 0.096572 | 4.229765 | 2.34E-05 | 0.00023 | 0.000278 | 5211.544 | 5227.577 | 6391.118 | 7466.576 | 14130 | Fc receptor, IgG, low affinity IIb [Source:MGI Symbol;Acc:MGI:95499] |
| ENSMUSG00000031393 | Mecp2 | -0.40741 | 715.1287 | 0.144107 | 2.827104 | 0.004697 | 0.022784 | 0.027547 | 620.9004 | 608.8185 | 821.1795 | 809.6163 | 17257 | methyl CpG binding protein 2 [Source:MGI Symbol;Acc:MGI:99918] |
| ENSMUSG00000035248 | Tut7 | -0.40713 | 2735.752 | 0.090787 | 4.48446 | 7.31E-06 | 8.1E-05 | 9.81E-05 | 2302.726 | 2401.833 | 3175.898 | 3062.551 | 214290 | terminal uridylyl transferase 7 [Source:MGI Symbol;Acc:MGI:2387179] |
| ENSMUSG00000020721 | Helz | -0.40675 | 1199.855 | 0.129991 | 3.129048 | 0.001754 | 0.010016 | 0.012133 | 1075.734 | 988.4695 | 1442.094 | 1293.123 | 78455 | helicase with zinc finger domain [Source:MGI Symbol;Acc:MGI:1925705] |
| ENSMUSG00000101970 | Chaserr | -0.40655 | 693.5544 | 0.148545 | 2.736857 | 0.006203 | 0.028658 | 0.03462 | 616.6694 | 576.3613 | 781.858 | 799.3289 | 69170 | CHD2 adjacent suppressive regulatory RNA [Source:MGI Symbol;Acc:MGI:1916420] |
| ENSMUSG00000029007 | Agtrap | -0.40635 | 699.7851 | 0.146695 | 2.770042 | 0.005605 | 0.026377 | 0.031864 | 585.9946 | 617.6705 | 799.2326 | 796.2427 | 11610 | angiotensin II, type I receptor-associated protein [Source:MGI Symbol;Acc:MGI:1339977] |
| ENSMUSG00000002395 | Use1 | -0.40621 | 1188.963 | 0.134711 | 3.015398 | 0.002566 | 0.013717 | 0.01661 | 940.3415 | 1104.529 | 1331.445 | 1379.537 | 67023 | unconventional SNARE in the ER 1 homolog (S. cerevisiae) [Source:MGI Symbol;Acc:MGI:1914273] |
| ENSMUSG00000060657 | Marf1 | -0.40568 | 1764.244 | 0.104135 | 3.895731 | 9.79E-05 | 0.000815 | 0.000987 | 1524.221 | 1511.719 | 2068.495 | 1952.543 | 223989 | meiosis regulator and mRNA stability 1 [Source:MGI Symbol;Acc:MGI:2444505] |
| ENSMUSG00000032633 | Flcn | -0.40481 | 2822.963 | 0.08682 | 4.662677 | 3.12E-06 | 3.69E-05 | 4.47E-05 | 2444.465 | 2414.619 | 3226.193 | 3206.574 | 216805 | folliculin [Source:MGI Symbol;Acc:MGI:2442184] |
| ENSMUSG00000092837 | Rpph1 | -0.40455 | 162381.3 | 0.0671 | 6.029051 | 1.65E-09 | 3.59E-08 | 4.35E-08 | 135084.4 | 144439.5 | 180672.3 | 189328.8 | 85029 | ribonuclease P RNA component H1 [Source:MGI Symbol;Acc:MGI:1934664] |
| ENSMUSG00000027447 | Cst3 | -0.40417 | 15518.25 | 0.073573 | 5.493524 | 3.94E-08 | 6.99E-07 | 8.46E-07 | 12801.97 | 13914.31 | 17409.37 | 17947.35 | 13010 | cystatin C [Source:MGI Symbol;Acc:MGI:102519] |
| ENSMUSG00000034708 | Grn | -0.40375 | 15331.2 | 0.072358 | 5.579943 | 2.41E-08 | 4.41E-07 | 5.34E-07 | 13513.84 | 12885.51 | 16876.25 | 18049.2 | 14824 | granulin [Source:MGI Symbol;Acc:MGI:95832] |
| ENSMUSG00000021709 | Erbin | -0.40375 | 8776.541 | 0.068661 | 5.880331 | 4.09E-09 | 8.42E-08 | 1.02E-07 | 7487.826 | 7624.493 | 9850.497 | 10143.35 | 59079 | Erbb2 interacting protein [Source:MGI Symbol;Acc:MGI:1890169] |
| ENSMUSG00000024052 | Lpin2 | -0.40345 | 3429.941 | 0.08629 | 4.675492 | 2.93E-06 | 3.5E-05 | 4.24E-05 | 3044.21 | 2863.119 | 3924.836 | 3887.598 | 64898 | lipin 2 [Source:MGI Symbol;Acc:MGI:1891341] |
| ENSMUSG00000000149 | Gna12 | -0.40157 | 10086.45 | 0.072026 | 5.57533 | 2.47E-08 | 4.51E-07 | 5.47E-07 | 8569.906 | 8812.623 | 11138.96 | 11824.31 | 14673 | guanine nucleotide binding protein, alpha 12 [Source:MGI Symbol;Acc:MGI:95767] |
| ENSMUSG00000041515 | Irf8 | -0.40135 | 4145.797 | 0.082527 | 4.863312 | 1.15E-06 | 1.51E-05 | 1.83E-05 | 3661.937 | 3483.74 | 4635.366 | 4802.146 | 15900 | interferon regulatory factor 8 [Source:MGI Symbol;Acc:MGI:96395] |
| ENSMUSG00000025742 | Prps2 | -0.40065 | 6023.263 | 0.093939 | 4.265002 | 2E-05 | 0.0002 | 0.000242 | 4977.781 | 5405.6 | 6447.814 | 7261.857 | 110639 | phosphoribosyl pyrophosphate synthetase 2 [Source:MGI Symbol;Acc:MGI:97776] |
| ENSMUSG00000009995 | Tafazzin | -0.39991 | 810.2373 | 0.141917 | 2.817907 | 0.004834 | 0.023335 | 0.028204 | 665.326 | 731.7625 | 919.0261 | 924.8349 | 66826 | tafazzin, phospholipid-lysophospholipid transacylase [Source:MGI Symbol;Acc:MGI:109626] |
| ENSMUSG00000078771 | Evi2a | -0.39967 | 2747.517 | 0.108957 | 3.668145 | 0.000244 | 0.001842 | 0.002232 | 2315.419 | 2422.488 | 2893.332 | 3358.827 | 14017 | ecotropic viral integration site 2a [Source:MGI Symbol;Acc:MGI:95458] |
| ENSMUSG00000060012 | Kif13b | -0.39847 | 4773.278 | 0.079691 | 5.000127 | 5.73E-07 | 7.93E-06 | 9.6E-06 | 4058.594 | 4178.128 | 5563.537 | 5292.854 | 16554 | kinesin family member 13B [Source:MGI Symbol;Acc:MGI:1098265] |
| ENSMUSG00000018899 | Irf1 | -0.39797 | 1311.694 | 0.141381 | 2.814884 | 0.004879 | 0.023525 | 0.028435 | 1214.299 | 1049.45 | 1377.167 | 1605.859 | 16362 | interferon regulatory factor 1 [Source:MGI Symbol;Acc:MGI:96590] |
| ENSMUSG00000037266 | Rsrp1 | -0.39791 | 4700.83 | 0.079439 | 5.009003 | 5.47E-07 | 7.62E-06 | 9.23E-06 | 4087.153 | 4026.661 | 5483.065 | 5206.44 | 27981 | arginine/serine rich protein 1 [Source:MGI Symbol;Acc:MGI:106498] |
| ENSMUSG00000020032 | Nuak1 | -0.39718 | 1273.697 | 0.117086 | 3.392213 | 0.000693 | 0.004585 | 0.005554 | 1085.253 | 1113.381 | 1405.516 | 1490.64 | 77976 | NUAK family, SNF1-like kinase, 1 [Source:MGI Symbol;Acc:MGI:1925226] |
| ENSMUSG00000045098 | Kmt5b | -0.39608 | 1081.209 | 0.121387 | 3.262925 | 0.001103 | 0.006813 | 0.008254 | 933.9949 | 933.3906 | 1219.881 | 1237.571 | 225888 | lysine methyltransferase 5B [Source:MGI Symbol;Acc:MGI:2444557] |
| ENSMUSG00000032412 | Atp1b3 | -0.39477 | 5503.686 | 0.104767 | 3.76804 | 0.000165 | 0.001294 | 0.001568 | 4580.066 | 4929.561 | 5722.652 | 6782.465 | 11933 | ATPase, Na+/K+ transporting, beta 3 polypeptide [Source:MGI Symbol;Acc:MGI:107788] |
| ENSMUSG00000034160 | Ogt | -0.39346 | 3053.794 | 0.09186 | 4.283202 | 1.84E-05 | 0.000185 | 0.000225 | 2615.82 | 2664.442 | 3603.863 | 3331.052 | 108155 | O-linked N-acetylglucosamine (GlcNAc) transferase (UDP-N-acetylglucosamine:polypeptide-N-acetylglucosaminyl transferase) [Source:MGI Symbol;Acc:MGI:1339639] |
| ENSMUSG00000004071 | Cdip1 | -0.39333 | 745.2576 | 0.146171 | 2.690885 | 0.007126 | 0.032214 | 0.038919 | 617.7271 | 670.7823 | 862.33 | 830.191 | 66626 | cell death inducing Trp53 target 1 [Source:MGI Symbol;Acc:MGI:1913876] |
| ENSMUSG00000027339 | Rassf2 | -0.39234 | 3180.225 | 0.089793 | 4.369355 | 1.25E-05 | 0.000131 | 0.000159 | 2784.003 | 2716.57 | 3484.984 | 3735.345 | 215653 | Ras association (RalGDS/AF-6) domain family member 2 [Source:MGI Symbol;Acc:MGI:2442060] |
| ENSMUSG00000019873 | Reep3 | -0.39177 | 4129.284 | 0.095336 | 4.109389 | 3.97E-05 | 0.000374 | 0.000453 | 3631.262 | 3512.263 | 4397.608 | 4976.003 | 28193 | receptor accessory protein 3 [Source:MGI Symbol;Acc:MGI:88930] |
| ENSMUSG00000038664 | Herc1 | -0.38985 | 6685.27 | 0.080125 | 4.865536 | 1.14E-06 | 1.49E-05 | 1.81E-05 | 5908.602 | 5667.225 | 7872.533 | 7292.719 | 235439 | HECT and RLD domain containing E3 ubiquitin protein ligase family member 1 [Source:MGI Symbol;Acc:MGI:2384589] |
| ENSMUSG00000042349 | Ikbke | -0.38953 | 1571.382 | 0.114385 | 3.405393 | 0.000661 | 0.0044 | 0.00533 | 1403.637 | 1317.959 | 1857.256 | 1706.675 | 56489 | inhibitor of kappaB kinase epsilon [Source:MGI Symbol;Acc:MGI:1929612] |
| ENSMUSG00000045658 | Pid1 | -0.38946 | 1848.958 | 0.105718 | 3.683915 | 0.00023 | 0.001747 | 0.002116 | 1638.458 | 1563.847 | 2169.999 | 2023.526 | 98496 | phosphotyrosine interaction domain containing 1 [Source:MGI Symbol;Acc:MGI:2138391] |
| ENSMUSG00000028618 | Tmem59 | -0.3875 | 3303.347 | 0.089566 | 4.326433 | 1.52E-05 | 0.000157 | 0.00019 | 2815.736 | 2908.362 | 3613.007 | 3876.282 | 56374 | transmembrane protein 59 [Source:MGI Symbol;Acc:MGI:1929278] |
| ENSMUSG00000054263 | Lifr | -0.38732 | 1394.988 | 0.112314 | 3.448518 | 0.000564 | 0.003845 | 0.004657 | 1176.22 | 1241.242 | 1571.032 | 1591.457 | 16880 | LIF receptor alpha [Source:MGI Symbol;Acc:MGI:96788] |
| ENSMUSG00000022450 | Ndufa6 | -0.38564 | 857.4279 | 0.133719 | 2.883972 | 0.003927 | 0.019691 | 0.023813 | 742.5418 | 744.5486 | 984.8668 | 957.7545 | 67130 | NADH:ubiquinone oxidoreductase subunit A6 [Source:MGI Symbol;Acc:MGI:1914380] |
| ENSMUSG00000029311 | Hsd17b11 | -0.38518 | 1300.778 | 0.117205 | 3.286419 | 0.001015 | 0.006371 | 0.007718 | 1089.484 | 1166.492 | 1450.324 | 1496.813 | 114664 | hydroxysteroid (17-beta) dehydrogenase 11 [Source:MGI Symbol;Acc:MGI:2149821] |
| ENSMUSG00000003363 | Pld3 | -0.38406 | 3554.932 | 0.101589 | 3.780563 | 0.000156 | 0.001236 | 0.001497 | 2872.854 | 3295.882 | 4131.502 | 3919.489 | 18807 | phospholipase D family member 3 [Source:MGI Symbol;Acc:MGI:1333782] |
| ENSMUSG00000048720 | Tbc1d12 | -0.38197 | 864.6002 | 0.136234 | 2.80376 | 0.005051 | 0.02426 | 0.029323 | 723.5023 | 777.9894 | 985.7812 | 971.1281 | 209478 | TBC1D12: TBC1 domain family, member 12 [Source:MGI Symbol;Acc:MGI:2384803] |
| ENSMUSG00000051285 | Pcmtd1 | -0.38151 | 2446.227 | 0.097928 | 3.895803 | 9.79E-05 | 0.000815 | 0.000987 | 2031.941 | 2216.926 | 2754.335 | 2781.706 | 319263 | protein-L-isoaspartate (D-aspartate) O-methyltransferase domain containing 1 [Source:MGI Symbol;Acc:MGI:2441773] |
| ENSMUSG00000026773 | Pfkfb3 | -0.38126 | 1159.09 | 0.120252 | 3.1705 | 0.001522 | 0.008926 | 0.010813 | 996.4023 | 1016.992 | 1278.406 | 1344.56 | 170768 | 6-phosphofructo-2-kinase/fructose-2,6-biphosphatase 3 [Source:MGI Symbol;Acc:MGI:2181202] |
| ENSMUSG00000073725 | Lmbrd1 | -0.38124 | 2219.518 | 0.104661 | 3.642572 | 0.00027 | 0.002012 | 0.002438 | 1889.145 | 1966.12 | 2383.067 | 2639.74 | 68421 | LMBR1 domain containing 1 [Source:MGI Symbol;Acc:MGI:1915671] |
| ENSMUSG00000070565 | Rasal2 | -0.3809 | 1070.077 | 0.122904 | 3.099155 | 0.001941 | 0.010891 | 0.013193 | 913.8977 | 945.1932 | 1197.02 | 1224.197 | 226525 | RAS protein activator like 2 [Source:MGI Symbol;Acc:MGI:2443881] |
| ENSMUSG00000045636 | Mtus1 | -0.38022 | 2045.237 | 0.100251 | 3.792675 | 0.000149 | 0.001186 | 0.001437 | 1815.102 | 1739.903 | 2373.008 | 2252.935 | 102103 | mitochondrial tumor suppressor 1 [Source:MGI Symbol;Acc:MGI:2142572] |
| ENSMUSG00000035437 | Rabgap1 | -0.37932 | 1755.801 | 0.104336 | 3.635577 | 0.000277 | 0.002061 | 0.002496 | 1504.123 | 1548.11 | 1934.07 | 2036.9 | 227800 | RAB GTPase activating protein 1 [Source:MGI Symbol;Acc:MGI:2385139] |
| ENSMUSG00000058655 | Eif4b | -0.3788 | 6238.712 | 0.083599 | 4.531204 | 5.86E-06 | 6.63E-05 | 8.03E-05 | 5171.349 | 5677.06 | 7191.265 | 6915.172 | 75705 | eukaryotic translation initiation factor 4B [Source:MGI Symbol;Acc:MGI:95304] |
| ENSMUSG00000030213 | Atf7ip | -0.37815 | 934.4191 | 0.140624 | 2.689111 | 0.007164 | 0.03234 | 0.039072 | 854.6635 | 771.1045 | 1103.746 | 1008.163 | 54343 | activating transcription factor 7 interacting protein [Source:MGI Symbol;Acc:MGI:1858965] |
| ENSMUSG00000000711 | Rab5b | -0.37618 | 2305.446 | 0.109825 | 3.425285 | 0.000614 | 0.004137 | 0.005011 | 1872.221 | 2140.208 | 2536.695 | 2672.66 | 19344 | RAB5B, member RAS oncogene family [Source:MGI Symbol;Acc:MGI:105938] |
| ENSMUSG00000054640 | Slc8a1 | -0.37543 | 5327.481 | 0.076086 | 4.934335 | 8.04E-07 | 1.08E-05 | 1.31E-05 | 4711.227 | 4565.647 | 6108.552 | 5924.498 | 20541 | solute carrier family 8 (sodium/calcium exchanger), member 1 [Source:MGI Symbol;Acc:MGI:107956] |
| ENSMUSG00000000958 | Slc7a7 | -0.37535 | 1201.724 | 0.142672 | 2.630844 | 0.008517 | 0.037391 | 0.045163 | 951.9767 | 1139.936 | 1291.209 | 1423.772 | 20540 | solute carrier family 7 (cationic amino acid transporter, y+ system), member 7 [Source:MGI Symbol;Acc:MGI:1337120] |
| ENSMUSG00000005871 | Apc | -0.37516 | 2183.791 | 0.100621 | 3.728388 | 0.000193 | 0.001487 | 0.001801 | 1906.069 | 1897.271 | 2567.786 | 2364.038 | 11789 | APC, WNT signaling pathway regulator [Source:MGI Symbol;Acc:MGI:88039] |
| ENSMUSG00000058594 | Fbh1 | -0.3736 | 2097.21 | 0.095516 | 3.911342 | 9.18E-05 | 0.000772 | 0.000935 | 1812.987 | 1841.209 | 2351.061 | 2383.584 | 50755 | F-box DNA helicase 1 [Source:MGI Symbol;Acc:MGI:1354699] |
| ENSMUSG00000026031 | Cflar | -0.37345 | 3711.42 | 0.084895 | 4.398945 | 1.09E-05 | 0.000117 | 0.000141 | 3336.15 | 3131.629 | 4174.482 | 4203.421 | 12633 | CASP8 and FADD-like apoptosis regulator [Source:MGI Symbol;Acc:MGI:1336166] |
| ENSMUSG00000032175 | Tyk2 | -0.37331 | 1461.103 | 0.122167 | 3.055713 | 0.002245 | 0.012273 | 0.014867 | 1250.263 | 1296.321 | 1755.751 | 1542.077 | 54721 | tyrosine kinase 2 [Source:MGI Symbol;Acc:MGI:1929470] |
| ENSMUSG00000019189 | Rnf145 | -0.37254 | 3143.648 | 0.093988 | 3.963678 | 7.38E-05 | 0.000639 | 0.000774 | 2621.109 | 2858.201 | 3464.866 | 3630.414 | 74315 | ring finger protein 145 [Source:MGI Symbol;Acc:MGI:1921565] |
| ENSMUSG00000029729 | Zkscan1 | -0.37231 | 1105.025 | 0.141148 | 2.637752 | 0.008346 | 0.036723 | 0.044356 | 898.0314 | 1027.812 | 1172.33 | 1321.927 | 74570 | zinc finger with KRAB and SCAN domains 1 [Source:MGI Symbol;Acc:MGI:1921820] |
| ENSMUSG00000031154 | Otud5 | -0.37118 | 1254.25 | 0.115615 | 3.210511 | 0.001325 | 0.00795 | 0.009631 | 1078.907 | 1108.463 | 1394.542 | 1435.089 | 54644 | OTU domain containing 5 [Source:MGI Symbol;Acc:MGI:1859615] |
| ENSMUSG00000025907 | Rb1cc1 | -0.37063 | 1242.147 | 0.124227 | 2.983472 | 0.00285 | 0.014953 | 0.018107 | 1116.986 | 1050.433 | 1465.869 | 1335.301 | 12421 | RB1-inducible coiled-coil 1 [Source:MGI Symbol;Acc:MGI:1341850] |
| ENSMUSG00000030180 | Kdm5a | -0.37023 | 2268.791 | 0.094853 | 3.903191 | 9.49E-05 | 0.000793 | 0.00096 | 1991.747 | 1967.103 | 2614.423 | 2501.889 | 214899 | lysine demethylase 5A [Source:MGI Symbol;Acc:MGI:2136980] |
| ENSMUSG00000015149 | Sirt2 | -0.3682 | 1327.722 | 0.118218 | 3.114557 | 0.001842 | 0.010435 | 0.012641 | 1106.409 | 1211.736 | 1496.961 | 1495.784 | 64383 | sirtuin 2 [Source:MGI Symbol;Acc:MGI:1927664] |
| ENSMUSG00000053716 | Dusp7 | -0.36814 | 1012.486 | 0.125634 | 2.930258 | 0.003387 | 0.017308 | 0.020953 | 893.8004 | 874.3775 | 1160.442 | 1121.324 | 235584 | dual specificity phosphatase 7 [Source:MGI Symbol;Acc:MGI:2387100] |
| ENSMUSG00000026842 | Abl1 | -0.36756 | 2095.925 | 0.102015 | 3.603055 | 0.000314 | 0.002305 | 0.002792 | 1885.972 | 1775.311 | 2432.447 | 2289.969 | 11350 | c-abl oncogene 1, non-receptor tyrosine kinase [Source:MGI Symbol;Acc:MGI:87859] |
| ENSMUSG00000033767 | Tmem131l | -0.36641 | 870.0071 | 0.14104 | 2.597909 | 0.009379 | 0.040403 | 0.048804 | 799.6604 | 720.9434 | 1006.814 | 952.6108 | 229473 | transmembrane 131 like [Source:MGI Symbol;Acc:MGI:2443399] |
| ENSMUSG00000037110 | Ralgapa2 | -0.36605 | 797.4007 | 0.137574 | 2.660779 | 0.007796 | 0.034717 | 0.041944 | 700.2318 | 693.4039 | 907.1382 | 888.8291 | 241694 | Ral GTPase activating protein, alpha subunit 2 (catalytic) [Source:MGI Symbol;Acc:MGI:3036245] |
| ENSMUSG00000031812 | Map1lc3b | -0.36587 | 2843.566 | 0.092526 | 3.954195 | 7.68E-05 | 0.000662 | 0.000801 | 2393.693 | 2575.922 | 3237.167 | 3167.482 | 67443 | microtubule-associated protein 1 light chain 3 beta [Source:MGI Symbol;Acc:MGI:1914693] |
| ENSMUSG00000031996 | Aplp2 | -0.36565 | 5897.51 | 0.086969 | 4.204351 | 2.62E-05 | 0.000254 | 0.000308 | 4902.68 | 5404.617 | 6447.814 | 6834.931 | 11804 | amyloid beta precursor-like protein 2 [Source:MGI Symbol;Acc:MGI:88047] |
| ENSMUSG00000091264 | Smim13 | -0.36559 | 997.4192 | 0.13436 | 2.720965 | 0.006509 | 0.0298 | 0.036001 | 904.3779 | 838.9696 | 1073.569 | 1172.761 | 108934 | small integral membrane protein 13 [Source:MGI Symbol;Acc:MGI:2652854] |
| ENSMUSG00000025314 | Ptprj | -0.36516 | 8877.522 | 0.068014 | 5.368909 | 7.92E-08 | 1.33E-06 | 1.61E-06 | 7864.385 | 7655.966 | 10056.25 | 9933.488 | 19271 | protein tyrosine phosphatase receptor type J [Source:MGI Symbol;Acc:MGI:104574] |
| ENSMUSG00000034707 | Gns | -0.36493 | 10682.67 | 0.070169 | 5.200759 | 1.98E-07 | 3.04E-06 | 3.68E-06 | 9078.685 | 9598.481 | 12136.63 | 11916.89 | 75612 | glucosamine (N-acetyl)-6-sulfatase [Source:MGI Symbol;Acc:MGI:1922862] |
| ENSMUSG00000027947 | Il6ra | -0.36468 | 1561.589 | 0.109724 | 3.323652 | 0.000888 | 0.005677 | 0.006877 | 1409.983 | 1320.91 | 1784.099 | 1731.365 | 16194 | interleukin 6 receptor, alpha [Source:MGI Symbol;Acc:MGI:105304] |
| ENSMUSG00000034850 | Tmem127 | -0.36405 | 1319.24 | 0.12693 | 2.868128 | 0.004129 | 0.020514 | 0.024807 | 1202.664 | 1104.529 | 1400.943 | 1568.824 | 69470 | transmembrane protein 127 [Source:MGI Symbol;Acc:MGI:1916720] |
| ENSMUSG00000041354 | Rgl2 | -0.36381 | 2397.465 | 0.100015 | 3.637565 | 0.000275 | 0.002047 | 0.002479 | 1994.92 | 2198.238 | 2708.612 | 2688.091 | 19732 | ral guanine nucleotide dissociation stimulator-like 2 [Source:MGI Symbol;Acc:MGI:107483] |
| ENSMUSG00000022014 | Epsti1 | -0.36292 | 864.8539 | 0.13624 | 2.663802 | 0.007726 | 0.034508 | 0.041691 | 754.1771 | 759.3019 | 1007.728 | 938.2085 | 108670 | epithelial stromal interaction 1 [Source:MGI Symbol;Acc:MGI:1915168] |
| ENSMUSG00000066151 | Fkbp15 | -0.36209 | 3564.252 | 0.085403 | 4.239716 | 2.24E-05 | 0.000221 | 0.000267 | 3072.769 | 3166.053 | 4114.128 | 3904.058 | 338355 | FK506 binding protein 15 [Source:MGI Symbol;Acc:MGI:2444782] |
| ENSMUSG00000022353 | Mtss1 | -0.36174 | 1780.236 | 0.122089 | 2.962957 | 0.003047 | 0.015835 | 0.019175 | 1684.999 | 1432.051 | 2049.291 | 1954.601 | 211401 | MTSS I-BAR domain containing 1 [Source:MGI Symbol;Acc:MGI:2384818] |
| ENSMUSG00000073079 | Srp54a | -0.3614 | 1384.935 | 0.124263 | 2.90838 | 0.003633 | 0.018381 | 0.022237 | 1202.664 | 1221.571 | 1453.067 | 1662.44 | 24067 | signal recognition particle 54A [Source:MGI Symbol;Acc:MGI:1346087] |
| ENSMUSG00000045917 | Tmem268 | -0.35917 | 1264.246 | 0.114324 | 3.141701 | 0.00168 | 0.009669 | 0.011713 | 1114.871 | 1100.594 | 1431.12 | 1410.399 | 230279 | transmembrane protein 268 [Source:MGI Symbol;Acc:MGI:1913920] |
| ENSMUSG00000001768 | Rin2;Rin2 | -0.35898 | 5178.97 | 0.08933 | 4.018542 | 5.86E-05 | 0.000523 | 0.000633 | 4355.822 | 4719.081 | 5574.51 | 6066.464 | 74030;408057 | Ras and Rab interactor 2 [Source:MGI Symbol;Acc:MGI:1921280];Ras and Rab interactor 2 [Source:MGI Symbol;Acc:MGI:1921280] |
| ENSMUSG00000029578 | Wipi2 | -0.35777 | 1069.479 | 0.132883 | 2.692358 | 0.007095 | 0.032092 | 0.038771 | 883.2228 | 991.4201 | 1167.758 | 1235.514 | 74781 | WD repeat domain, phosphoinositide interacting 2 [Source:MGI Symbol;Acc:MGI:1923831] |
| ENSMUSG00000053436 | Mapk14 | -0.35587 | 2225.4 | 0.09372 | 3.797165 | 0.000146 | 0.001166 | 0.001412 | 1949.437 | 1955.301 | 2524.807 | 2472.056 | 26416 | mitogen-activated protein kinase 14 [Source:MGI Symbol;Acc:MGI:1346865] |
| ENSMUSG00000089417 | Gm22009 | -0.35535 | 2494.884 | 0.111624 | 3.183484 | 0.001455 | 0.008586 | 0.010401 | 2017.133 | 2360.524 | 2754.335 | 2847.545 | 115489854 | predicted gene, 22009 [Source:MGI Symbol;Acc:MGI:5451786] |
| ENSMUSG00000040188 | Scamp2 | -0.35359 | 3151.524 | 0.084793 | 4.170071 | 3.05E-05 | 0.000293 | 0.000355 | 2750.155 | 2784.435 | 3581.001 | 3490.506 | 24044 | secretory carrier membrane protein 2 [Source:MGI Symbol;Acc:MGI:1346518] |
| ENSMUSG00000071713 | Csf2rb | -0.35288 | 1557.953 | 0.114118 | 3.092268 | 0.001986 | 0.011085 | 0.013428 | 1372.962 | 1364.186 | 1835.309 | 1659.353 | 12983 | colony stimulating factor 2 receptor, beta, low-affinity (granulocyte-macrophage) [Source:MGI Symbol;Acc:MGI:1339759] |
| ENSMUSG00000017132 | Cyth1 | -0.35251 | 1276.896 | 0.135428 | 2.602968 | 0.009242 | 0.039914 | 0.048213 | 1027.077 | 1215.67 | 1384.483 | 1480.353 | 19157 | cytohesin 1 [Source:MGI Symbol;Acc:MGI:1334257] |
| ENSMUSG00000028822 | Tmem50a | -0.3519 | 3035.088 | 0.095582 | 3.681624 | 0.000232 | 0.00176 | 0.002132 | 2607.358 | 2725.422 | 3248.14 | 3559.431 | 71817 | transmembrane protein 50A [Source:MGI Symbol;Acc:MGI:1919067] |
| ENSMUSG00000001211 | Agpat3 | -0.35182 | 2467.012 | 0.104959 | 3.351969 | 0.000802 | 0.005206 | 0.006307 | 2048.865 | 2285.774 | 2676.606 | 2856.804 | 28169 | 1-acylglycerol-3-phosphate O-acyltransferase 3 [Source:MGI Symbol;Acc:MGI:1336186] |
| ENSMUSG00000030315 | Vgll4 | -0.34969 | 1014.858 | 0.128364 | 2.724175 | 0.006446 | 0.029628 | 0.035793 | 862.0678 | 922.5715 | 1117.463 | 1157.33 | 232334 | vestigial like family member 4 [Source:MGI Symbol;Acc:MGI:2652840] |
| ENSMUSG00000022307 | Oxr1 | -0.34887 | 2051.539 | 0.104277 | 3.345606 | 0.000821 | 0.005302 | 0.006423 | 1843.662 | 1765.475 | 2201.09 | 2395.929 | 170719 | oxidation resistance 1 [Source:MGI Symbol;Acc:MGI:2179326] |
| ENSMUSG00000061119 | Prcp | -0.34762 | 1690.855 | 0.106791 | 3.255127 | 0.001133 | 0.006969 | 0.008442 | 1491.43 | 1485.163 | 1958.76 | 1828.066 | 72461 | prolylcarboxypeptidase (angiotensinase C) [Source:MGI Symbol;Acc:MGI:1919711] |
| ENSMUSG00000033790 | Tubgcp5 | -0.34756 | 1252.704 | 0.129009 | 2.694071 | 0.007059 | 0.031958 | 0.03861 | 1052.463 | 1152.723 | 1479.586 | 1326.042 | 233276 | tubulin, gamma complex component 5 [Source:MGI Symbol;Acc:MGI:2178836] |
| ENSMUSG00000028134 | Ptbp2 | -0.34755 | 1134.871 | 0.119523 | 2.907786 | 0.00364 | 0.01841 | 0.022271 | 1012.269 | 985.5188 | 1278.406 | 1263.289 | 56195 | polypyrimidine tract binding protein 2 [Source:MGI Symbol;Acc:MGI:1860489] |
| ENSMUSG00000034593 | Myo5a | -0.34713 | 15727.44 | 0.062877 | 5.520743 | 3.38E-08 | 6.06E-07 | 7.35E-07 | 13740.2 | 13948.73 | 17780.64 | 17440.18 | 17918 | myosin VA [Source:MGI Symbol;Acc:MGI:105976] |
| ENSMUSG00000028603 | Scp2 | -0.34657 | 2798.598 | 0.091268 | 3.797317 | 0.000146 | 0.001165 | 0.001412 | 2385.231 | 2542.481 | 3114.63 | 3152.051 | 20280 | sterol carrier protein 2, liver [Source:MGI Symbol;Acc:MGI:98254] |
| ENSMUSG00000029038 | Ssu72 | -0.34634 | 1553.991 | 0.115966 | 2.986563 | 0.002821 | 0.014833 | 0.017963 | 1290.457 | 1445.821 | 1724.66 | 1755.026 | 68991 | Ssu72 RNA polymerase II CTD phosphatase homolog (yeast) [Source:MGI Symbol;Acc:MGI:1916241] |
| ENSMUSG00000017802 | Retreg3 | -0.34487 | 1391.357 | 0.115072 | 2.996992 | 0.002727 | 0.0144 | 0.017438 | 1249.205 | 1202.884 | 1610.353 | 1502.985 | 67998 | reticulophagy regulator family member 3 [Source:MGI Symbol;Acc:MGI:1915248] |
| ENSMUSG00000076161 | Rn7sk | -0.34455 | 306150.4 | 0.078607 | 4.38314 | 1.17E-05 | 0.000124 | 0.00015 | 252655.7 | 286874.5 | 337408.8 | 347662.8 | 19817 | RNA, 7SK, nuclear [Source:MGI Symbol;Acc:MGI:103186] |
| ENSMUSG00000005142 | Man2b1 | -0.34397 | 16985.35 | 0.063971 | 5.37701 | 7.57E-08 | 1.28E-06 | 1.55E-06 | 14717.56 | 15222.43 | 19180.67 | 18820.75 | 17159 | mannosidase 2, alpha B1 [Source:MGI Symbol;Acc:MGI:107286] |
| ENSMUSG00000001248 | Gramd1a | -0.34149 | 2026.44 | 0.1117 | 3.057163 | 0.002234 | 0.012218 | 0.014801 | 1665.959 | 1909.074 | 2278.819 | 2251.906 | 52857 | GRAM domain containing 1A [Source:MGI Symbol;Acc:MGI:105490] |
| ENSMUSG00000021003 | Galc | -0.34094 | 2349.265 | 0.093882 | 3.631534 | 0.000282 | 0.002086 | 0.002527 | 2105.984 | 2039.886 | 2580.589 | 2670.602 | 14420 | galactosylceramidase [Source:MGI Symbol;Acc:MGI:95636] |
| ENSMUSG00000026880 | Stom | -0.33849 | 3115.963 | 0.096289 | 3.515398 | 0.000439 | 0.003097 | 0.003751 | 2614.763 | 2888.691 | 3399.939 | 3560.46 | 13830 | stomatin [Source:MGI Symbol;Acc:MGI:95403] |
| ENSMUSG00000045594 | Glb1 | -0.33703 | 2868.671 | 0.09389 | 3.589598 | 0.000331 | 0.002414 | 0.002925 | 2485.717 | 2584.774 | 3323.125 | 3081.068 | 12091 | galactosidase, beta 1 [Source:MGI Symbol;Acc:MGI:88151] |
| ENSMUSG00000056608 | Chd9 | -0.33506 | 2718.856 | 0.093932 | 3.567087 | 0.000361 | 0.002604 | 0.003154 | 2411.674 | 2397.899 | 3149.379 | 2916.47 | 109151 | chromodomain helicase DNA binding protein 9 [Source:MGI Symbol;Acc:MGI:1924001] |
| ENSMUSG00000021939 | Ctsb | -0.33495 | 149031.9 | 0.061964 | 5.405553 | 6.46E-08 | 1.11E-06 | 1.34E-06 | 129036.2 | 134580.4 | 163023.3 | 169487.6 | 13030 | cathepsin B [Source:MGI Symbol;Acc:MGI:88561] |
| ENSMUSG00000032434 | Cmtm6 | -0.33438 | 4960.196 | 0.0808 | 4.13834 | 3.5E-05 | 0.000332 | 0.000402 | 4247.932 | 4527.288 | 5453.803 | 5611.762 | 67213 | CKLF-like MARVEL transmembrane domain containing 6 [Source:MGI Symbol;Acc:MGI:2447165] |
| ENSMUSG00000027598 | Itch | -0.33428 | 3348.757 | 0.082511 | 4.051345 | 5.09E-05 | 0.000461 | 0.000558 | 2974.398 | 2950.655 | 3726.399 | 3743.575 | 16396 | itchy, E3 ubiquitin protein ligase [Source:MGI Symbol;Acc:MGI:1202301] |
| ENSMUSG00000072946 | Ptgr2 | -0.33383 | 1279.08 | 0.127666 | 2.614876 | 0.008926 | 0.038781 | 0.046844 | 1051.405 | 1211.736 | 1428.377 | 1424.801 | 77219 | prostaglandin reductase 2 [Source:MGI Symbol;Acc:MGI:1916372] |
| ENSMUSG00000074748 | Atxn7l3b | -0.33219 | 1714.807 | 0.126792 | 2.61995 | 0.008794 | 0.038362 | 0.046338 | 1494.603 | 1541.226 | 1740.206 | 2083.193 | 382423 | ataxin 7-like 3B [Source:MGI Symbol;Acc:MGI:1914971] |
| ENSMUSG00000023942 | Slc29a1 | -0.33131 | 1435.402 | 0.113439 | 2.920572 | 0.003494 | 0.01779 | 0.021528 | 1273.533 | 1268.782 | 1539.94 | 1659.353 | 63959 | solute carrier family 29 (nucleoside transporters), member 1 [Source:MGI Symbol;Acc:MGI:1927073] |
| ENSMUSG00000055447 | Cd47 | -0.32953 | 1591.882 | 0.115359 | 2.856546 | 0.004283 | 0.021112 | 0.025524 | 1454.409 | 1367.137 | 1688.082 | 1857.9 | 16423 | CD47 antigen (Rh-related antigen, integrin-associated signal transducer) [Source:MGI Symbol;Acc:MGI:96617] |
| ENSMUSG00000045210 | Vcpip1 | -0.32851 | 2401.159 | 0.091057 | 3.607708 | 0.000309 | 0.002267 | 0.002746 | 2132.428 | 2125.455 | 2663.804 | 2682.947 | 70675 | valosin containing protein (p97)/p47 complex interacting protein 1 [Source:MGI Symbol;Acc:MGI:1917925] |
| ENSMUSG00000039220 | Ppp1r10 | -0.32706 | 1159.328 | 0.120089 | 2.7235 | 0.006459 | 0.029655 | 0.035825 | 1046.117 | 1011.091 | 1316.814 | 1263.289 | 52040 | protein phosphatase 1, regulatory subunit 10 [Source:MGI Symbol;Acc:MGI:1289273] |
| ENSMUSG00000027009 | Itga4 | -0.3242 | 2957.523 | 0.093834 | 3.455064 | 0.00055 | 0.00377 | 0.004566 | 2742.751 | 2511.007 | 3320.382 | 3255.954 | 16401 | integrin alpha 4 [Source:MGI Symbol;Acc:MGI:96603] |
| ENSMUSG00000042350 | Arel1 | -0.32279 | 1557.72 | 0.105759 | 3.052121 | 0.002272 | 0.012389 | 0.015008 | 1384.597 | 1383.857 | 1747.521 | 1714.905 | 68497 | apoptosis resistant E3 ubiquitin protein ligase 1 [Source:MGI Symbol;Acc:MGI:1915747] |
| ENSMUSG00000040699 | Limd2 | -0.32277 | 2401.942 | 0.096001 | 3.362108 | 0.000773 | 0.005034 | 0.006098 | 2205.413 | 2063.491 | 2636.37 | 2702.493 | 67803 | LIM domain containing 2 [Source:MGI Symbol;Acc:MGI:1915053] |
| ENSMUSG00000023030 | Slc11a2 | -0.32269 | 1752.901 | 0.115841 | 2.785599 | 0.005343 | 0.025404 | 0.030697 | 1600.379 | 1514.67 | 1829.822 | 2066.733 | 18174 | solute carrier family 11 (proton-coupled divalent metal ion transporters), member 2 [Source:MGI Symbol;Acc:MGI:1345279] |
| ENSMUSG00000022971 | Ifnar2 | -0.3206 | 2091.696 | 0.100179 | 3.200247 | 0.001373 | 0.008194 | 0.009926 | 1798.178 | 1921.86 | 2288.878 | 2357.866 | 15976 | interferon (alpha and beta) receptor 2 [Source:MGI Symbol;Acc:MGI:1098243] |
| ENSMUSG00000023150 | Ivns1abp | -0.32033 | 5704.773 | 0.072372 | 4.426205 | 9.59E-06 | 0.000104 | 0.000126 | 5072.978 | 5075.127 | 6354.54 | 6316.447 | 117198 | influenza virus NS1A binding protein [Source:MGI Symbol;Acc:MGI:2152389] |
| ENSMUSG00000036887 | C1qa | -0.31946 | 55982.7 | 0.06639 | 4.811813 | 1.5E-06 | 1.91E-05 | 2.31E-05 | 49684.72 | 49933.94 | 59872.04 | 64440.11 | 12259 | complement component 1, q subcomponent, alpha polypeptide [Source:MGI Symbol;Acc:MGI:88223] |
| ENSMUSG00000020190 | Mknk2 | -0.31842 | 1992.235 | 0.114182 | 2.788727 | 0.005292 | 0.025208 | 0.030459 | 1647.977 | 1898.255 | 2254.129 | 2168.578 | 17347 | MAP kinase-interacting serine/threonine kinase 2 [Source:MGI Symbol;Acc:MGI:894279] |
| ENSMUSG00000033392 | Clasp2 | -0.31825 | 2596.689 | 0.090511 | 3.516086 | 0.000438 | 0.003092 | 0.003745 | 2290.033 | 2332.985 | 2932.653 | 2831.085 | 76499 | CLIP associating protein 2 [Source:MGI Symbol;Acc:MGI:1923749] |
| ENSMUSG00000027620 | Rbm39 | -0.3164 | 5653.771 | 0.072977 | 4.335628 | 1.45E-05 | 0.00015 | 0.000182 | 5060.285 | 5012.179 | 6221.029 | 6321.591 | 170791 | RNA binding motif protein 39 [Source:MGI Symbol;Acc:MGI:2157953] |
| ENSMUSG00000004508 | Gab2 | -0.31583 | 1688.991 | 0.119181 | 2.650023 | 0.008049 | 0.035633 | 0.043038 | 1448.062 | 1561.88 | 1988.937 | 1757.083 | 14389 | growth factor receptor bound protein 2-associated protein 2 [Source:MGI Symbol;Acc:MGI:1333854] |
| ENSMUSG00000028581 | Laptm5 | -0.3157 | 11301.73 | 0.067349 | 4.687576 | 2.76E-06 | 3.32E-05 | 4.02E-05 | 9862.479 | 10277.13 | 12453.95 | 12613.35 | 16792 | lysosomal-associated protein transmembrane 5 [Source:MGI Symbol;Acc:MGI:108046] |
| ENSMUSG00000034724 | Cnot6l | -0.31522 | 1805.062 | 0.104569 | 3.014471 | 0.002574 | 0.013749 | 0.016649 | 1647.977 | 1569.749 | 2052.034 | 1950.486 | 231464 | CCR4-NOT transcription complex, subunit 6-like [Source:MGI Symbol;Acc:MGI:2443154] |
| ENSMUSG00000040785 | Ttc3 | -0.31455 | 2344.907 | 0.093076 | 3.379479 | 0.000726 | 0.004779 | 0.00579 | 2052.039 | 2128.406 | 2609.851 | 2589.332 | 22129 | tetratricopeptide repeat domain 3 [Source:MGI Symbol;Acc:MGI:1276539] |
| ENSMUSG00000043300 | B3galnt1 | -0.31447 | 1527.079 | 0.114603 | 2.744008 | 0.006069 | 0.028213 | 0.034082 | 1370.846 | 1351.4 | 1609.439 | 1776.629 | 26879 | UDP-GalNAc:betaGlcNAc beta 1,3-galactosaminyltransferase, polypeptide 1 [Source:MGI Symbol;Acc:MGI:1349405] |
| ENSMUSG00000024835 | Coro1b | -0.3126 | 3322.189 | 0.090994 | 3.435333 | 0.000592 | 0.004006 | 0.004853 | 2832.66 | 3094.254 | 3686.163 | 3675.679 | 23789 | coronin, actin binding protein 1B [Source:MGI Symbol;Acc:MGI:1345963] |
| ENSMUSG00000024425 | Ndfip1 | -0.31239 | 2303.504 | 0.103229 | 3.026148 | 0.002477 | 0.013317 | 0.016127 | 1993.862 | 2115.62 | 2437.019 | 2667.516 | 65113 | Nedd4 family interacting protein 1 [Source:MGI Symbol;Acc:MGI:1929601] |
| ENSMUSG00000039497 | Dse | -0.31189 | 5731.6 | 0.075092 | 4.153461 | 3.27E-05 | 0.000313 | 0.000379 | 5216.832 | 5012.179 | 6283.212 | 6414.177 | 212898 | dermatan sulfate epimerase [Source:MGI Symbol;Acc:MGI:2443455] |
| ENSMUSG00000047126 | Cltc | -0.31016 | 44165.59 | 0.074216 | 4.17919 | 2.93E-05 | 0.000282 | 0.000342 | 41245.98 | 37626.75 | 47587.26 | 50202.38 | 67300 | clathrin heavy chain [Source:MGI Symbol;Acc:MGI:2388633] |
| ENSMUSG00000030275 | Etnk1 | -0.30958 | 3017.29 | 0.094508 | 3.275675 | 0.001054 | 0.006564 | 0.007952 | 2817.851 | 2571.988 | 3286.547 | 3392.776 | 75320 | ethanolamine kinase 1 [Source:MGI Symbol;Acc:MGI:1922570] |
| ENSMUSG00000022807 | Osbpl11 | -0.30853 | 3097.872 | 0.096252 | 3.205424 | 0.001349 | 0.008079 | 0.009786 | 2622.167 | 2913.28 | 3477.668 | 3378.373 | 106326 | oxysterol binding protein-like 11 [Source:MGI Symbol;Acc:MGI:2146553] |
| ENSMUSG00000038116 | Phf20 | -0.30632 | 1766.484 | 0.107024 | 2.862172 | 0.004207 | 0.020808 | 0.025157 | 1645.862 | 1513.686 | 1940.471 | 1965.917 | 228829 | PHD finger protein 20 [Source:MGI Symbol;Acc:MGI:2444148] |
| ENSMUSG00000023026 | Dip2b | -0.30629 | 2893.74 | 0.089593 | 3.418699 | 0.000629 | 0.004223 | 0.005116 | 2603.127 | 2571.988 | 3110.972 | 3288.873 | 239667 | disco interacting protein 2 homolog B [Source:MGI Symbol;Acc:MGI:2145977] |
| ENSMUSG00000048811 | Cd300c2 | -0.30607 | 2834.744 | 0.089722 | 3.411327 | 0.000646 | 0.004316 | 0.005229 | 2477.255 | 2592.642 | 3087.196 | 3181.885 | 140497 | CD300C molecule 2 [Source:MGI Symbol;Acc:MGI:2153249] |
| ENSMUSG00000036093 | Arl5a | -0.30459 | 2209.855 | 0.093659 | 3.252144 | 0.001145 | 0.007026 | 0.008511 | 1962.13 | 1992.676 | 2443.421 | 2441.194 | 75423 | ADP-ribosylation factor-like 5A [Source:MGI Symbol;Acc:MGI:1922673] |
| ENSMUSG00000037416 | Dmxl1 | -0.30436 | 7327.674 | 0.071772 | 4.2406 | 2.23E-05 | 0.00022 | 0.000267 | 6636.336 | 6478.655 | 7949.347 | 8246.359 | 240283 | Dmx-like 1 [Source:MGI Symbol;Acc:MGI:2443926] |
| ENSMUSG00000053470 | Kdm3a | -0.30435 | 1365.626 | 0.113712 | 2.676545 | 0.007439 | 0.033394 | 0.040345 | 1188.913 | 1255.012 | 1487.816 | 1530.761 | 104263 | lysine (K)-specific demethylase 3A [Source:MGI Symbol;Acc:MGI:98847] |
| ENSMUSG00000035697 | Arhgap45 | -0.30401 | 5875.168 | 0.077852 | 3.904919 | 9.43E-05 | 0.000788 | 0.000954 | 5204.139 | 5313.146 | 6694.716 | 6288.671 | 70719 | Rho GTPase activating protein 45 [Source:MGI Symbol;Acc:MGI:1917969] |
| ENSMUSG00000036402 | Gng12 | -0.30399 | 3673.749 | 0.085377 | 3.560558 | 0.00037 | 0.002659 | 0.003222 | 3233.548 | 3342.109 | 3946.783 | 4172.559 | 14701 | guanine nucleotide binding protein (G protein), gamma 12 [Source:MGI Symbol;Acc:MGI:1336171] |
| ENSMUSG00000058099 | Nfam1 | -0.30327 | 2105.992 | 0.098939 | 3.0652 | 0.002175 | 0.011954 | 0.014481 | 1867.99 | 1903.173 | 2396.783 | 2256.021 | 74039 | Nfat activating molecule with ITAM motif 1 [Source:MGI Symbol;Acc:MGI:1921289] |
| ENSMUSG00000036257 | Pnpla8 | -0.30239 | 2268.871 | 0.097433 | 3.103572 | 0.001912 | 0.010766 | 0.013041 | 1982.227 | 2081.195 | 2444.335 | 2567.728 | 67452 | patatin-like phospholipase domain containing 8 [Source:MGI Symbol;Acc:MGI:1914702] |
| ENSMUSG00000000222 | Rmnd5a | -0.30228 | 2572.545 | 0.089759 | 3.367667 | 0.000758 | 0.004946 | 0.005992 | 2329.17 | 2278.889 | 2820.176 | 2861.947 | 68477 | required for meiotic nuclear division 5 homolog A [Source:MGI Symbol;Acc:MGI:1915727] |
| ENSMUSG00000021945 | Zmym2 | -0.30204 | 1809.791 | 0.100179 | 3.014982 | 0.00257 | 0.013731 | 0.016627 | 1617.303 | 1624.827 | 2013.627 | 1983.406 | 76007 | zinc finger, MYM-type 2 [Source:MGI Symbol;Acc:MGI:1923257] |
| ENSMUSG00000066150 | Slc31a1 | -0.30086 | 2083.899 | 0.102092 | 2.946933 | 0.003209 | 0.016561 | 0.020047 | 1818.276 | 1915.959 | 2223.952 | 2377.412 | 20529 | solute carrier family 31, member 1 [Source:MGI Symbol;Acc:MGI:1333843] |
| ENSMUSG00000027180 | Fbxo3 | -0.30084 | 1613.223 | 0.104195 | 2.88726 | 0.003886 | 0.019512 | 0.023597 | 1438.543 | 1452.706 | 1785.014 | 1776.629 | 57443 | F-box protein 3 [Source:MGI Symbol;Acc:MGI:1929084] |
| ENSMUSG00000036362 | P2ry13 | -0.3005 | 2621.286 | 0.088991 | 3.376786 | 0.000733 | 0.004818 | 0.005837 | 2355.614 | 2342.82 | 2864.069 | 2922.643 | 74191 | purinergic receptor P2Y, G-protein coupled 13 [Source:MGI Symbol;Acc:MGI:1921441] |
| ENSMUSG00000097589 | Dleu2 | -0.30034 | 1687.92 | 0.110813 | 2.710358 | 0.006721 | 0.030659 | 0.037039 | 1540.087 | 1486.147 | 1947.787 | 1777.658 | | deleted in lymphocytic leukemia, 2 [Source:MGI Symbol;Acc:MGI:1934030] |
| ENSMUSG00000066406 | Akap13 | -0.29946 | 4487.457 | 0.087729 | 3.413499 | 0.000641 | 0.004293 | 0.005201 | 3921.086 | 4125.999 | 5160.263 | 4742.479 | 75547 | A kinase anchor protein 13 [Source:MGI Symbol;Acc:MGI:2676556] |
| ENSMUSG00000036896 | C1qc | -0.29929 | 41686.99 | 0.068042 | 4.398621 | 1.09E-05 | 0.000117 | 0.000141 | 36522.06 | 38233.6 | 44520.18 | 47472.11 | 12262 | complement component 1, q subcomponent, C chain [Source:MGI Symbol;Acc:MGI:88225] |
| ENSMUSG00000030516 | Tjp1 | -0.29921 | 2041.429 | 0.106984 | 2.796798 | 0.005161 | 0.02468 | 0.029821 | 1833.084 | 1828.423 | 2375.751 | 2128.458 | 21872 | tight junction protein 1 [Source:MGI Symbol;Acc:MGI:98759] |
| ENSMUSG00000037685 | Atp8a1 | -0.29894 | 3972.881 | 0.079564 | 3.757207 | 0.000172 | 0.001344 | 0.001628 | 3524.429 | 3600.783 | 4350.971 | 4415.341 | 11980 | ATPase phospholipid transporting 8A1 [Source:MGI Symbol;Acc:MGI:1330848] |
| ENSMUSG00000026782 | Abi2 | -0.29862 | 2173.727 | 0.096691 | 3.088365 | 0.002013 | 0.011178 | 0.013541 | 1997.036 | 1902.189 | 2373.008 | 2422.676 | 329165 | abl interactor 2 [Source:MGI Symbol;Acc:MGI:106913] |
| ENSMUSG00000002428 | Hltf | -0.29736 | 1915.197 | 0.100456 | 2.960087 | 0.003076 | 0.015949 | 0.019313 | 1762.215 | 1674.989 | 2102.329 | 2121.256 | 20585 | helicase-like transcription factor [Source:MGI Symbol;Acc:MGI:1196437] |
| ENSMUSG00000043004 | Gng2 | -0.29673 | 2077.689 | 0.099205 | 2.991041 | 0.00278 | 0.014641 | 0.017729 | 1924.051 | 1805.801 | 2297.108 | 2283.797 | 14702 | guanine nucleotide binding protein (G protein), gamma 2 [Source:MGI Symbol;Acc:MGI:102705] |
| ENSMUSG00000028436 | Dcaf12 | -0.29644 | 1485.73 | 0.111657 | 2.654882 | 0.007934 | 0.035226 | 0.042547 | 1377.193 | 1290.42 | 1664.306 | 1611.003 | 68970 | DDB1 and CUL4 associated factor 12 [Source:MGI Symbol;Acc:MGI:1916220] |
| ENSMUSG00000029571 | Tmem106b | -0.296 | 3688.435 | 0.090911 | 3.25597 | 0.00113 | 0.006954 | 0.008424 | 3192.295 | 3429.645 | 3929.408 | 4202.392 | 71900 | transmembrane protein 106B [Source:MGI Symbol;Acc:MGI:1919150] |
| ENSMUSG00000026469 | Xpr1 | -0.29415 | 1747.67 | 0.101829 | 2.888663 | 0.003869 | 0.019444 | 0.023515 | 1583.455 | 1556.962 | 1945.043 | 1905.222 | 19775 | xenotropic and polytropic retrovirus receptor 1 [Source:MGI Symbol;Acc:MGI:97932] |
| ENSMUSG00000030894 | Tpp1 | -0.29371 | 3682.133 | 0.087975 | 3.338602 | 0.000842 | 0.005412 | 0.006556 | 3344.612 | 3273.26 | 4213.803 | 3896.857 | 12751 | tripeptidyl peptidase I [Source:MGI Symbol;Acc:MGI:1336194] |
| ENSMUSG00000052684 | Jun | -0.29342 | 3068.918 | 0.086944 | 3.374785 | 0.000739 | 0.004847 | 0.005871 | 2823.14 | 2692.965 | 3385.308 | 3374.259 | 16476 | jun proto-oncogene [Source:MGI Symbol;Acc:MGI:96646] |
| ENSMUSG00000034354 | Mtmr3 | -0.29337 | 2474.1 | 0.092184 | 3.182469 | 0.00146 | 0.008609 | 0.010428 | 2206.47 | 2240.531 | 2780.854 | 2668.545 | 74302 | myotubularin related protein 3 [Source:MGI Symbol;Acc:MGI:1921552] |
| ENSMUSG00000027291 | Vps39 | -0.292 | 1752.057 | 0.103888 | 2.810671 | 0.004944 | 0.023798 | 0.028764 | 1571.819 | 1578.6 | 1876.459 | 1981.348 | 269338 | VPS39 HOPS complex subunit [Source:MGI Symbol;Acc:MGI:2443189] |
| ENSMUSG00000023992 | Trem2 | -0.29197 | 8179.607 | 0.085783 | 3.403593 | 0.000665 | 0.004419 | 0.005353 | 6932.506 | 7775.96 | 8800.704 | 9209.257 | 83433 | triggering receptor expressed on myeloid cells 2 [Source:MGI Symbol;Acc:MGI:1913150] |
| ENSMUSG00000029098 | Acox3 | -0.29052 | 2093.701 | 0.099134 | 2.930518 | 0.003384 | 0.017303 | 0.020946 | 1840.488 | 1926.778 | 2355.633 | 2251.906 | 80911 | acyl-Coenzyme A oxidase 3, pristanoyl [Source:MGI Symbol;Acc:MGI:1933156] |
| ENSMUSG00000028668 | Eloa | -0.28907 | 2437.669 | 0.091262 | 3.167499 | 0.001538 | 0.009002 | 0.010904 | 2198.008 | 2190.37 | 2646.429 | 2715.867 | 27224 | elongin A [Source:MGI Symbol;Acc:MGI:1351315] |
| ENSMUSG00000046609 | Cd33 | -0.28765 | 4253.551 | 0.092496 | 3.109846 | 0.001872 | 0.01057 | 0.012804 | 3966.57 | 3696.187 | 4895.986 | 4455.461 | 12489 | CD33 molecule [Source:MGI Symbol;Acc:MGI:99440] |
| ENSMUSG00000029178 | Klf3 | -0.28606 | 2128.686 | 0.101503 | 2.818269 | 0.004828 | 0.02332 | 0.028187 | 1925.108 | 1912.024 | 2437.019 | 2240.59 | 16599 | Kruppel-like transcription factor 3 (basic) [Source:MGI Symbol;Acc:MGI:1342773] |
| ENSMUSG00000018398 | Septin8 | -0.28442 | 2235.364 | 0.103722 | 2.742131 | 0.006104 | 0.028331 | 0.034225 | 1951.552 | 2079.228 | 2350.146 | 2560.527 | 20362 | septin 8 [Source:MGI Symbol;Acc:MGI:894310] |
| ENSMUSG00000040710 | St8sia4 | -0.28296 | 2487.571 | 0.091544 | 3.09096 | 0.001995 | 0.011108 | 0.013456 | 2284.744 | 2204.139 | 2709.527 | 2751.872 | 20452 | ST8 alpha-N-acetyl-neuraminide alpha-2,8-sialyltransferase 4 [Source:MGI Symbol;Acc:MGI:106018] |
| ENSMUSG00000029647 | Pan3 | -0.2827 | 2250.763 | 0.094747 | 2.983772 | 0.002847 | 0.014944 | 0.018096 | 2075.309 | 1986.774 | 2469.94 | 2471.027 | 72587 | PAN3 poly(A) specific ribonuclease subunit [Source:MGI Symbol;Acc:MGI:1919837] |
| ENSMUSG00000037940 | Inpp4b | -0.28213 | 3826.369 | 0.090463 | 3.11875 | 0.001816 | 0.010314 | 0.012494 | 3563.566 | 3343.092 | 4040.057 | 4358.76 | 234515 | inositol polyphosphate-4-phosphatase, type II [Source:MGI Symbol;Acc:MGI:2158925] |
| ENSMUSG00000069662 | Marcks | -0.28049 | 14779.44 | 0.069854 | 4.015354 | 5.94E-05 | 0.000529 | 0.000641 | 13343.54 | 13351.71 | 16769.25 | 15653.27 | 17118 | myristoylated alanine rich protein kinase C substrate [Source:MGI Symbol;Acc:MGI:96907] |
| ENSMUSG00000003429 | Rps11 | -0.28016 | 5396.307 | 0.083386 | 3.35979 | 0.00078 | 0.005074 | 0.006146 | 4883.641 | 4863.663 | 5653.154 | 6184.769 | 27207 | ribosomal protein S11 [Source:MGI Symbol;Acc:MGI:1351329] |
| ENSMUSG00000014353 | Tmem87b | -0.27797 | 2957.089 | 0.092432 | 3.007323 | 0.002636 | 0.013998 | 0.01695 | 2772.368 | 2573.955 | 3185.957 | 3296.074 | 72477 | transmembrane protein 87B [Source:MGI Symbol;Acc:MGI:1919727] |
| ENSMUSG00000034892 | Rps29 | -0.27757 | 7257.581 | 0.074914 | 3.705209 | 0.000211 | 0.001613 | 0.001954 | 6418.439 | 6703.888 | 7765.542 | 8142.456 | 20090 | ribosomal protein S29 [Source:MGI Symbol;Acc:MGI:107681] |
| ENSMUSG00000041225 | Arhgap12 | -0.27703 | 2870.629 | 0.087857 | 3.153235 | 0.001615 | 0.009366 | 0.011346 | 2554.471 | 2636.902 | 3108.229 | 3182.913 | 75415 | Rho GTPase activating protein 12 [Source:MGI Symbol;Acc:MGI:1922665] |
| ENSMUSG00000060090 | Rp2 | -0.27569 | 3326.867 | 0.08327 | 3.310761 | 0.00093 | 0.005901 | 0.007148 | 2977.572 | 3042.125 | 3620.323 | 3667.449 | 19889 | retinitis pigmentosa 2 homolog [Source:MGI Symbol;Acc:MGI:1277953] |
| ENSMUSG00000032122 | Slc37a2 | -0.27433 | 5285.069 | 0.082295 | 3.333529 | 0.000858 | 0.005503 | 0.006666 | 4745.075 | 4822.354 | 5551.649 | 6021.2 | 56857 | solute carrier family 37 (glycerol-3-phosphate transporter), member 2 [Source:MGI Symbol;Acc:MGI:1929693] |
| ENSMUSG00000020590 | Snx13 | -0.2701 | 2638.602 | 0.101684 | 2.656236 | 0.007902 | 0.035116 | 0.042414 | 2354.556 | 2429.373 | 2722.329 | 3048.149 | 217463 | sorting nexin 13 [Source:MGI Symbol;Acc:MGI:2661416] |
| ENSMUSG00000040462 | Os9 | -0.26881 | 3101.545 | 0.097098 | 2.768456 | 0.005632 | 0.026481 | 0.031989 | 2654.957 | 2971.31 | 3362.447 | 3417.465 | 216440 | amplified in osteosarcoma [Source:MGI Symbol;Acc:MGI:1924301] |
| ENSMUSG00000028530 | Jak1 | -0.26723 | 4705.72 | 0.081915 | 3.262326 | 0.001105 | 0.006819 | 0.008261 | 4424.576 | 4118.131 | 5136.487 | 5143.687 | 16451 | Janus kinase 1 [Source:MGI Symbol;Acc:MGI:96628] |
| ENSMUSG00000102869 | Norad | -0.26232 | 6245.009 | 0.078564 | 3.33893 | 0.00084 | 0.00541 | 0.006554 | 5839.848 | 5517.725 | 6643.507 | 6978.954 | 347740 | non-coding RNA activated by DNA damage [Source:MGI Symbol;Acc:MGI:1914767] |
| ENSMUSG00000079419 | Ms4a6c | -0.26034 | 4961.811 | 0.074989 | 3.471723 | 0.000517 | 0.003571 | 0.004325 | 4553.622 | 4477.127 | 5397.106 | 5419.388 | 73656 | membrane-spanning 4-domains, subfamily A, member 6C [Source:MGI Symbol;Acc:MGI:2385644] |
| ENSMUSG00000040713 | Creg1 | -0.2603 | 5368.87 | 0.079333 | 3.281071 | 0.001034 | 0.006477 | 0.007846 | 4800.078 | 4970.87 | 5683.33 | 6021.2 | 433375 | cellular repressor of E1A-stimulated genes 1 [Source:MGI Symbol;Acc:MGI:1344382] |
| ENSMUSG00000074212 | Dnajb14 | -0.25805 | 2070.518 | 0.096948 | 2.661709 | 0.007775 | 0.034641 | 0.041853 | 1920.877 | 1851.044 | 2270.589 | 2239.561 | 70604 | DnaJ heat shock protein family (Hsp40) member B14 [Source:MGI Symbol;Acc:MGI:1917854] |
| ENSMUSG00000078606 | Gvin2 | -0.25761 | 2496.422 | 0.098654 | 2.611267 | 0.009021 | 0.039114 | 0.047246 | 2381 | 2167.748 | 2740.618 | 2696.321 | 100042856 | GTPase, very large interferon inducible, family member 2 [Source:MGI Symbol;Acc:MGI:3782245] |
| ENSMUSG00000015846 | Rxra | -0.25713 | 3403.628 | 0.092195 | 2.789001 | 0.005287 | 0.025198 | 0.030448 | 2950.07 | 3251.622 | 3689.821 | 3723 | 20181 | retinoid X receptor alpha [Source:MGI Symbol;Acc:MGI:98214] |
| ENSMUSG00000041236 | Vps41 | -0.25457 | 2645.181 | 0.091403 | 2.78513 | 0.005351 | 0.025425 | 0.030722 | 2398.981 | 2425.438 | 2802.801 | 2953.505 | 218035 | VPS41 HOPS complex subunit [Source:MGI Symbol;Acc:MGI:1929215] |
| ENSMUSG00000030432 | Rpl28 | -0.25426 | 2825.344 | 0.092139 | 2.759556 | 0.005788 | 0.027087 | 0.032722 | 2504.757 | 2648.705 | 3003.066 | 3144.85 | 19943 | ribosomal protein L28 [Source:MGI Symbol;Acc:MGI:101839] |
| ENSMUSG00000026867 | Gapvd1 | -0.25404 | 3989.51 | 0.08039 | 3.160052 | 0.001577 | 0.009214 | 0.011161 | 3641.84 | 3636.191 | 4259.526 | 4420.484 | 66691 | GTPase activating protein and VPS9 domains 1 [Source:MGI Symbol;Acc:MGI:1913941] |
| ENSMUSG00000051495 | Irf2bp2 | -0.25336 | 4288.107 | 0.085876 | 2.950269 | 0.003175 | 0.016414 | 0.019877 | 3787.81 | 4036.496 | 4531.119 | 4797.002 | 270110 | interferon regulatory factor 2 binding protein 2 [Source:MGI Symbol;Acc:MGI:2443921] |
| ENSMUSG00000035901 | Dennd5a | -0.25135 | 5271.254 | 0.088637 | 2.835713 | 0.004572 | 0.022293 | 0.026953 | 4564.2 | 5061.357 | 5586.398 | 5873.062 | 19347 | DENN domain containing 5A [Source:MGI Symbol;Acc:MGI:1201681] |
| ENSMUSG00000026339 | Ccdc93 | -0.24992 | 2393.495 | 0.092221 | 2.709993 | 0.006728 | 0.030675 | 0.037058 | 2151.467 | 2221.843 | 2614.423 | 2586.246 | 70829 | coiled-coil domain containing 93 [Source:MGI Symbol;Acc:MGI:1918079] |
| ENSMUSG00000022141 | Nipbl | -0.24798 | 2820.508 | 0.093489 | 2.652487 | 0.00799 | 0.035403 | 0.042761 | 2667.65 | 2490.353 | 3128.347 | 2995.683 | 71175 | NIPBL cohesin loading factor [Source:MGI Symbol;Acc:MGI:1913976] |
| ENSMUSG00000045128 | Rpl18a | -0.24757 | 4583.345 | 0.082498 | 3.000934 | 0.002692 | 0.01424 | 0.017244 | 4246.874 | 4134.851 | 4810.027 | 5141.629 | 76808 | ribosomal protein L18A [Source:MGI Symbol;Acc:MGI:1924058] |
| ENSMUSG00000029063 | Nadk | -0.24722 | 3398.598 | 0.086411 | 2.860983 | 0.004223 | 0.020873 | 0.025235 | 3203.931 | 3012.619 | 3681.591 | 3696.253 | 192185 | NAD kinase [Source:MGI Symbol;Acc:MGI:2183149] |
| ENSMUSG00000036905 | C1qb | -0.24622 | 24965.3 | 0.071671 | 3.435364 | 0.000592 | 0.004006 | 0.004853 | 22208.56 | 23470.49 | 26146.98 | 28035.15 | 12260 | complement component 1, q subcomponent, beta polypeptide [Source:MGI Symbol;Acc:MGI:88224] |
| ENSMUSG00000025612 | Bach1 | -0.24004 | 3076.792 | 0.085865 | 2.795539 | 0.005181 | 0.024754 | 0.029911 | 2778.714 | 2864.103 | 3365.19 | 3299.161 | 12013 | BTB and CNC homology 1, basic leucine zipper transcription factor 1 [Source:MGI Symbol;Acc:MGI:894680] |
| ENSMUSG00000023010 | Tmbim6 | -0.23756 | 9000.648 | 0.073264 | 3.242518 | 0.001185 | 0.007239 | 0.008769 | 7981.796 | 8540.179 | 9709.671 | 9770.947 | 110213 | transmembrane BAX inhibitor motif containing 6 [Source:MGI Symbol;Acc:MGI:99682] |
| ENSMUSG00000024431 | Nr3c1 | -0.23407 | 3726.37 | 0.08458 | 2.767433 | 0.00565 | 0.026548 | 0.03207 | 3371.055 | 3477.839 | 3922.092 | 4134.495 | 14815 | nuclear receptor subfamily 3, group C, member 1 [Source:MGI Symbol;Acc:MGI:95824] |
| ENSMUSG00000074785 | Plxnc1 | -0.23292 | 3938.355 | 0.082647 | 2.818211 | 0.004829 | 0.02332 | 0.028187 | 3672.515 | 3570.293 | 4355.544 | 4155.07 | 54712 | plexin C1 [Source:MGI Symbol;Acc:MGI:1890127] |
| ENSMUSG00000071415 | Rpl23 | -0.23155 | 9551.637 | 0.070035 | 3.306152 | 0.000946 | 0.005981 | 0.007245 | 8861.846 | 8711.317 | 10067.22 | 10566.16 | 65019 | ribosomal protein L23 [Source:MGI Symbol;Acc:MGI:1929455] |
| ENSMUSG00000030844 | Rgs10 | -0.22988 | 3249.787 | 0.084467 | 2.721583 | 0.006497 | 0.029763 | 0.035956 | 2952.186 | 3030.323 | 3468.523 | 3548.115 | 67865 | regulator of G-protein signalling 10 [Source:MGI Symbol;Acc:MGI:1915115] |
| ENSMUSG00000067288 | Rps28 | -0.22904 | 3813.414 | 0.087536 | 2.616569 | 0.008882 | 0.038666 | 0.046705 | 3378.46 | 3644.059 | 4188.198 | 4042.938 | 54127 | ribosomal protein S28 [Source:MGI Symbol;Acc:MGI:1859516] |
| ENSMUSG00000027423 | Snx5 | -0.22721 | 12662.3 | 0.067483 | 3.366907 | 0.00076 | 0.004958 | 0.006005 | 11375.06 | 11959.01 | 13663.77 | 13651.34 | 69178 | sorting nexin 5 [Source:MGI Symbol;Acc:MGI:1916428] |
| ENSMUSG00000030064 | Frmd4b | -0.22721 | 3594.388 | 0.086488 | 2.627068 | 0.008612 | 0.03771 | 0.045549 | 3219.797 | 3404.072 | 3958.671 | 3795.012 | 232288 | FERM domain containing 4B [Source:MGI Symbol;Acc:MGI:2141794] |
| ENSMUSG00000101249 | Gm29216 | -0.22667 | 10509.84 | 0.080934 | 2.800678 | 0.0051 | 0.024447 | 0.029539 | 9290.235 | 10080.42 | 10911.26 | 11757.44 | | predicted gene 29216 [Source:MGI Symbol;Acc:MGI:5579922] |
| ENSMUSG00000027293 | Ehd4 | -0.22663 | 7715.004 | 0.080213 | 2.825323 | 0.004723 | 0.022882 | 0.027665 | 6780.19 | 7439.585 | 8220.025 | 8420.215 | 98878 | EH-domain containing 4 [Source:MGI Symbol;Acc:MGI:1919619] |
| ENSMUSG00000005656 | Snx6 | -0.22572 | 3822.994 | 0.083516 | 2.70276 | 0.006877 | 0.031247 | 0.03775 | 3468.369 | 3580.128 | 4023.597 | 4219.881 | 72183 | sorting nexin 6 [Source:MGI Symbol;Acc:MGI:1919433] |
| ENSMUSG00000015305 | Sash1 | -0.22412 | 7024.667 | 0.076133 | 2.943791 | 0.003242 | 0.016716 | 0.020235 | 6325.356 | 6635.04 | 7759.141 | 7379.133 | 70097 | SAM and SH3 domain containing 1 [Source:MGI Symbol;Acc:MGI:1917347] |
| ENSMUSG00000074129 | Rpl13a | -0.21741 | 11972.22 | 0.064777 | 3.356326 | 0.00079 | 0.005136 | 0.006221 | 11042.93 | 11100.36 | 12722.8 | 13022.79 | 22121 | ribosomal protein L13A [Source:MGI Symbol;Acc:MGI:1351455] |
| ENSMUSG00000020053 | Igf1 | -0.21478 | 11279.53 | 0.064878 | 3.310498 | 0.000931 | 0.005901 | 0.007148 | 10459.05 | 10423.68 | 12006.78 | 12228.6 | 16000 | insulin-like growth factor 1 [Source:MGI Symbol;Acc:MGI:96432] |
| ENSMUSG00000006333 | Rps9 | -0.21439 | 8244.261 | 0.071225 | 3.010074 | 0.002612 | 0.013906 | 0.016839 | 7676.106 | 7589.085 | 8642.503 | 9069.348 | 76846 | ribosomal protein S9 [Source:MGI Symbol;Acc:MGI:1924096] |
| ENSMUSG00000030579 | Tyrobp | -0.20547 | 12103.68 | 0.067073 | 3.063358 | 0.002189 | 0.012011 | 0.01455 | 11032.35 | 11453.46 | 12803.27 | 13125.66 | 22177 | TYRO protein tyrosine kinase binding protein [Source:MGI Symbol;Acc:MGI:1277211] |
| ENSMUSG00000026872 | Zeb2 | -0.18331 | 8810.713 | 0.066959 | 2.73768 | 0.006187 | 0.028619 | 0.034573 | 8291.717 | 8211.673 | 9326.515 | 9412.947 | 24136 | zinc finger E-box binding homeobox 2 [Source:MGI Symbol;Acc:MGI:1344407] |
| ENSMUSG00000038642 | Ctss | 0.159505 | 32734.5 | 0.059433 | -2.68377 | 0.00728 | 0.032783 | 0.039607 | 34621.28 | 34463.65 | 31294.44 | 30558.64 | 13040 | cathepsin S [Source:MGI Symbol;Acc:MGI:107341] |
| ENSMUSG00000036103 | Colec12 | 0.179189 | 11793.31 | 0.063958 | -2.80166 | 0.005084 | 0.024403 | 0.029487 | 12539.65 | 12509.79 | 11023.74 | 11100.08 | 140792 | collectin sub-family member 12 [Source:MGI Symbol;Acc:MGI:2152907] |
| ENSMUSG00000021190 | Lgmn | 0.18213 | 49325.05 | 0.061911 | -2.9418 | 0.003263 | 0.016801 | 0.020338 | 51745.22 | 53122.61 | 45238.03 | 47194.35 | 19141 | legumain [Source:MGI Symbol;Acc:MGI:1330838] |
| ENSMUSG00000024121 | Atp6v0c | 0.184604 | 6674.497 | 0.070415 | -2.62166 | 0.00875 | 0.038192 | 0.046132 | 7157.807 | 7044.197 | 6248.463 | 6247.522 | 11984 | ATPase, H+ transporting, lysosomal V0 subunit C [Source:MGI Symbol;Acc:MGI:88116] |
| ENSMUSG00000015120 | Ube2i | 0.201918 | 4707.728 | 0.07665 | -2.63429 | 0.008431 | 0.037056 | 0.044759 | 5068.747 | 5004.311 | 4319.88 | 4437.973 | 22196 | ubiquitin-conjugating enzyme E2I [Source:MGI Symbol;Acc:MGI:107365] |
| ENSMUSG00000024740 | Ddb1 | 0.202655 | 7494.465 | 0.070858 | -2.86001 | 0.004236 | 0.020918 | 0.02529 | 8044.203 | 7996.275 | 7103.477 | 6833.902 | 13194 | damage specific DNA binding protein 1 [Source:MGI Symbol;Acc:MGI:1202384] |
| ENSMUSG00000026395 | Ptprc | 0.203434 | 9653.503 | 0.070832 | -2.87206 | 0.004078 | 0.020301 | 0.024551 | 10627.23 | 10039.11 | 8953.417 | 8994.251 | 19264 | protein tyrosine phosphatase receptor type C [Source:MGI Symbol;Acc:MGI:97810] |
| ENSMUSG00000027111 | Itga6 | 0.205771 | 11217.14 | 0.067458 | -3.05037 | 0.002286 | 0.012449 | 0.01508 | 12100.68 | 11931.47 | 10632.35 | 10204.05 | 16403 | integrin alpha 6 [Source:MGI Symbol;Acc:MGI:96605] |
| ENSMUSG00000031246 | Sh3bgrl | 0.206169 | 6078.691 | 0.073262 | -2.81413 | 0.004891 | 0.023562 | 0.028479 | 6548.542 | 6475.704 | 5547.077 | 5743.441 | 56726 | SH3-binding domain glutamic acid-rich protein like [Source:MGI Symbol;Acc:MGI:1930849] |
| ENSMUSG00000052593 | Adam17 | 0.20652 | 5344.209 | 0.076462 | -2.70095 | 0.006914 | 0.031408 | 0.037945 | 5861.003 | 5591.491 | 4979.201 | 4945.14 | 11491 | a disintegrin and metallopeptidase domain 17 [Source:MGI Symbol;Acc:MGI:1096335] |
| ENSMUSG00000018707 | Dync1h1 | 0.208465 | 15343.37 | 0.075743 | -2.75228 | 0.005918 | 0.027586 | 0.033325 | 17205.39 | 15695.52 | 14477.63 | 13994.94 | 13424 | dynein cytoplasmic 1 heavy chain 1 [Source:MGI Symbol;Acc:MGI:103147] |
| ENSMUSG00000019055 | Plod1 | 0.209638 | 4422.803 | 0.077367 | -2.70967 | 0.006735 | 0.030695 | 0.037083 | 4710.169 | 4777.111 | 4142.476 | 4061.455 | 18822 | procollagen-lysine, 2-oxoglutarate 5-dioxygenase 1 [Source:MGI Symbol;Acc:MGI:99907] |
| ENSMUSG00000043131 | Mob1a | 0.2166 | 8067.611 | 0.078021 | -2.77617 | 0.0055 | 0.026015 | 0.031426 | 8813.189 | 8530.344 | 7162.002 | 7764.91 | 232157 | MOB kinase activator 1A [Source:MGI Symbol;Acc:MGI:2442631] |
| ENSMUSG00000079477 | Rab7 | 0.222151 | 3837.02 | 0.080999 | -2.74264 | 0.006095 | 0.028304 | 0.034193 | 4087.153 | 4176.16 | 3491.385 | 3593.38 | 19349 | RAB7, member RAS oncogene family [Source:MGI Symbol;Acc:MGI:105068] |
| ENSMUSG00000031555 | Adam9 | 0.2229 | 5967.379 | 0.0723 | -3.08297 | 0.002049 | 0.011338 | 0.013734 | 6490.366 | 6364.563 | 5483.065 | 5531.521 | 11502 | ADAM metallopeptidase domain 9 [Source:MGI Symbol;Acc:MGI:105376] |
| ENSMUSG00000003812 | Dnase2a;Dnase | 0.223754 | 3098.269 | 0.08516 | -2.62745 | 0.008603 | 0.037679 | 0.045511 | 3289.608 | 3386.368 | 2861.326 | 2855.775 | 13423;100503 | deoxyribonuclease II alpha [Source:MGI Symbol;Acc:MGI:1329019];deoxyribonuclease II alpha [Source:MGI Symbol;Acc:MGI:1329019] |
| ENSMUSG00000019173 | Rab5c | 0.229332 | 4859.728 | 0.076234 | -3.00828 | 0.002627 | 0.013968 | 0.016915 | 5309.915 | 5180.367 | 4431.443 | 4517.186 | 19345 | RAB5C, member RAS oncogene family [Source:MGI Symbol;Acc:MGI:105306] |
| ENSMUSG00000027303 | Ptpra | 0.229588 | 4579.383 | 0.087273 | -2.63069 | 0.008521 | 0.037397 | 0.04517 | 4913.258 | 4971.854 | 4009.88 | 4422.542 | 19262 | protein tyrosine phosphatase receptor type A [Source:MGI Symbol;Acc:MGI:97808] |
| ENSMUSG00000018846 | Pank3 | 0.230637 | 4144.916 | 0.084255 | -2.73736 | 0.006194 | 0.028632 | 0.034589 | 4562.084 | 4388.608 | 3697.137 | 3931.834 | 211347 | pantothenate kinase 3 [Source:MGI Symbol;Acc:MGI:2387464] |
| ENSMUSG00000021987 | Mtmr6 | 0.231717 | 3400.904 | 0.083785 | -2.76562 | 0.005681 | 0.026679 | 0.032229 | 3724.344 | 3622.421 | 3086.282 | 3170.569 | 219135 | myotubularin related protein 6 [Source:MGI Symbol;Acc:MGI:2145637] |
| ENSMUSG00000048865 | Arhgap30 | 0.233682 | 3634.374 | 0.085 | -2.7492 | 0.005974 | 0.027829 | 0.033618 | 3820.6 | 4035.513 | 3385.308 | 3296.074 | 226652 | Rho GTPase activating protein 30 [Source:MGI Symbol;Acc:MGI:2684948] |
| ENSMUSG00000024725 | Ostf1 | 0.234219 | 4072.376 | 0.089713 | -2.61076 | 0.009034 | 0.039161 | 0.047303 | 4563.142 | 4241.075 | 3612.093 | 3873.196 | 20409 | osteoclast stimulating factor 1 [Source:MGI Symbol;Acc:MGI:700012] |
| ENSMUSG00000018774 | Cd68 | 0.235679 | 9379.374 | 0.073795 | -3.19371 | 0.001405 | 0.008342 | 0.010105 | 10089.9 | 10198.45 | 8924.155 | 8304.997 | 12514 | CD68 antigen [Source:MGI Symbol;Acc:MGI:88342] |
| ENSMUSG00000048076 | Arf1 | 0.235943 | 5433.787 | 0.076453 | -3.08612 | 0.002028 | 0.011243 | 0.013619 | 5937.161 | 5816.725 | 4877.697 | 5103.566 | 11840 | ARF GTPase 1 [Source:MGI Symbol;Acc:MGI:99431] |
| ENSMUSG00000017631 | Abr | 0.235958 | 3524.942 | 0.088749 | -2.65871 | 0.007844 | 0.03489 | 0.042154 | 3847.044 | 3778.806 | 3366.105 | 3107.816 | 109934 | active BCR-related gene [Source:MGI Symbol;Acc:MGI:107771] |
| ENSMUSG00000029465 | Arpc3 | 0.235999 | 6024.313 | 0.072066 | -3.27476 | 0.001058 | 0.006577 | 0.007967 | 6528.445 | 6503.244 | 5478.493 | 5587.073 | 56378 | actin related protein 2/3 complex, subunit 3 [Source:MGI Symbol;Acc:MGI:1928375] |
| ENSMUSG00000034088 | Hdlbp | 0.244195 | 4569.578 | 0.085156 | -2.86763 | 0.004136 | 0.020533 | 0.024825 | 5149.136 | 4762.357 | 4263.184 | 4103.633 | 110611 | high density lipoprotein (HDL) binding protein [Source:MGI Symbol;Acc:MGI:99256] |
| ENSMUSG00000037805 | Rpl10a | 0.247373 | 5195.921 | 0.082286 | -3.00624 | 0.002645 | 0.014033 | 0.016993 | 5778.499 | 5502.972 | 4907.874 | 4594.341 | 19896 | ribosomal protein L10A [Source:MGI Symbol;Acc:MGI:1343877] |
| ENSMUSG00000004771 | Rab11a | 0.247511 | 2302.184 | 0.094103 | -2.63021 | 0.008533 | 0.037439 | 0.045221 | 2553.413 | 2445.11 | 2102.329 | 2107.883 | 53869 | RAB11A, member RAS oncogene family [Source:MGI Symbol;Acc:MGI:1858202] |
| ENSMUSG00000036046 | 5031439G07Rik | 0.24776 | 3669.184 | 0.094838 | -2.61245 | 0.008989 | 0.039012 | 0.047123 | 4216.199 | 3751.266 | 3367.933 | 3341.339 | 223739 | RIKEN cDNA 5031439G07 gene [Source:MGI Symbol;Acc:MGI:2444899] |
| ENSMUSG00000021660 | Btf3 | 0.248165 | 2925.39 | 0.088847 | -2.79317 | 0.005219 | 0.024903 | 0.030091 | 3245.183 | 3108.023 | 2716.842 | 2631.51 | 218490 | basic transcription factor 3 [Source:MGI Symbol;Acc:MGI:1202875] |
| ENSMUSG00000002812 | Flii | 0.248844 | 2504.171 | 0.094816 | -2.6245 | 0.008678 | 0.037941 | 0.045828 | 2758.617 | 2681.162 | 2362.034 | 2214.872 | 14248 | flightless I actin binding protein [Source:MGI Symbol;Acc:MGI:1342286] |
| ENSMUSG00000042228 | Lyn | 0.248954 | 6647.532 | 0.074235 | -3.35358 | 0.000798 | 0.00518 | 0.006275 | 7402.148 | 7037.312 | 6023.507 | 6127.16 | 17096 | LYN proto-oncogene, Src family tyrosine kinase [Source:MGI Symbol;Acc:MGI:96892] |
| ENSMUSG00000037260 | Hgsnat | 0.251057 | 6388.768 | 0.079765 | -3.14745 | 0.001647 | 0.009528 | 0.011542 | 6833.077 | 7054.033 | 6058.257 | 5609.705 | 52120 | heparan-alpha-glucosaminide N-acetyltransferase [Source:MGI Symbol;Acc:MGI:1196297] |
| ENSMUSG00000024644 | Cndp2 | 0.251073 | 4886.458 | 0.076198 | -3.29501 | 0.000984 | 0.006192 | 0.007501 | 5257.027 | 5364.291 | 4517.402 | 4407.111 | 66054 | CNDP dipeptidase 2 [Source:MGI Symbol;Acc:MGI:1913304] |
| ENSMUSG00000014769 | Psmb1 | 0.251397 | 2334.039 | 0.095288 | -2.63829 | 0.008333 | 0.036686 | 0.044312 | 2540.72 | 2532.646 | 2068.495 | 2194.297 | 19170 | proteasome (prosome, macropain) subunit, beta type 1 [Source:MGI Symbol;Acc:MGI:104884] |
| ENSMUSG00000017781 | Pitpna | 0.25153 | 3275.865 | 0.086254 | -2.91616 | 0.003544 | 0.018013 | 0.021799 | 3655.591 | 3466.036 | 2982.034 | 2999.798 | 18738 | phosphatidylinositol transfer protein, alpha [Source:MGI Symbol;Acc:MGI:99887] |
| ENSMUSG00000061306 | Slc38a10 | 0.251535 | 3888.954 | 0.088005 | -2.85819 | 0.004261 | 0.021023 | 0.025417 | 4405.537 | 4049.282 | 3591.975 | 3509.023 | 72055 | solute carrier family 38, member 10 [Source:MGI Symbol;Acc:MGI:1919305] |
| ENSMUSG00000032562 | Gnai2 | 0.252573 | 7084.226 | 0.07641 | -3.3055 | 0.000948 | 0.005992 | 0.007259 | 7461.382 | 7944.147 | 6575.837 | 6355.539 | 14678 | G protein subunit alpha i2 [Source:MGI Symbol;Acc:MGI:95772] |
| ENSMUSG00000007458 | M6pr | 0.253156 | 7054.085 | 0.092221 | -2.74512 | 0.006049 | 0.028126 | 0.033978 | 7873.905 | 7468.108 | 6003.389 | 6870.937 | 17113 | mannose-6-phosphate receptor, cation dependent [Source:MGI Symbol;Acc:MGI:96904] |
| ENSMUSG00000079426 | Arpc4 | 0.25369 | 6305.512 | 0.071404 | -3.55291 | 0.000381 | 0.002729 | 0.003305 | 6872.214 | 6844.536 | 5697.047 | 5808.251 | 68089 | actin related protein 2/3 complex, subunit 4 [Source:MGI Symbol;Acc:MGI:1915339] |
| ENSMUSG00000032518 | Rpsa | 0.254623 | 12525.93 | 0.069918 | -3.64175 | 0.000271 | 0.002017 | 0.002443 | 14053.29 | 13204.18 | 11499.26 | 11346.97 | 16785 | ribosomal protein SA [Source:MGI Symbol;Acc:MGI:105381] |
| ENSMUSG00000024965 | Fermt3 | 0.257018 | 5482.116 | 0.084494 | -3.04186 | 0.002351 | 0.012742 | 0.015435 | 5869.465 | 6069.497 | 5223.36 | 4766.14 | 108101 | fermitin family member 3 [Source:MGI Symbol;Acc:MGI:2147790] |
| ENSMUSG00000027698 | Nceh1 | 0.259074 | 4430.3 | 0.082845 | -3.12721 | 0.001765 | 0.010071 | 0.0122 | 4869.89 | 4783.012 | 3894.659 | 4172.559 | 320024 | neutral cholesterol ester hydrolase 1 [Source:MGI Symbol;Acc:MGI:2443191] |
| ENSMUSG00000004266 | Ptpn6 | 0.260381 | 4362.028 | 0.089306 | -2.91559 | 0.00355 | 0.018022 | 0.021809 | 4658.339 | 4849.893 | 3777.609 | 4162.271 | 15170 | protein tyrosine phosphatase, non-receptor type 6 [Source:MGI Symbol;Acc:MGI:96055] |
| ENSMUSG00000038900 | Rpl12 | 0.260449 | 7211.13 | 0.081782 | -3.1847 | 0.001449 | 0.008563 | 0.010373 | 8246.234 | 7474.993 | 6612.416 | 6510.879 | 269261 | ribosomal protein L12 [Source:MGI Symbol;Acc:MGI:98002] |
| ENSMUSG00000033047 | Eif3l | 0.26095 | 2056.003 | 0.097301 | -2.68189 | 0.007321 | 0.032944 | 0.039801 | 2283.686 | 2199.222 | 1855.427 | 1885.676 | 223691 | eukaryotic translation initiation factor 3, subunit L [Source:MGI Symbol;Acc:MGI:2386251] |
| ENSMUSG00000052459 | Atp6v1a | 0.262419 | 11785.45 | 0.065731 | -3.9923 | 6.54E-05 | 0.000575 | 0.000696 | 12774.47 | 12933.7 | 10553.71 | 10879.93 | 11964 | ATPase, H+ transporting, lysosomal V1 subunit A [Source:MGI Symbol;Acc:MGI:1201780] |
| ENSMUSG00000056537 | Rlim | 0.262904 | 4623.282 | 0.081622 | -3.221 | 0.001277 | 0.007705 | 0.009333 | 5117.404 | 4968.903 | 4074.806 | 4332.013 | 19820 | ring finger protein, LIM domain interacting [Source:MGI Symbol;Acc:MGI:1342291] |
| ENSMUSG00000028691 | Prdx1 | 0.263825 | 14379.56 | 0.065489 | -4.02854 | 5.61E-05 | 0.000503 | 0.000609 | 15626.17 | 15754.53 | 12796.87 | 13340.67 | 18477 | peroxiredoxin 1 [Source:MGI Symbol;Acc:MGI:99523] |
| ENSMUSG00000029064 | Gnb1 | 0.26496 | 3918.397 | 0.08281 | -3.19961 | 0.001376 | 0.008206 | 0.00994 | 4190.813 | 4363.035 | 3491.385 | 3628.357 | 14688 | guanine nucleotide binding protein (G protein), beta 1 [Source:MGI Symbol;Acc:MGI:95781] |
| ENSMUSG00000036707 | Cab39 | 0.265314 | 5677.436 | 0.081256 | -3.26517 | 0.001094 | 0.006776 | 0.008208 | 6233.332 | 6161.951 | 4944.452 | 5370.009 | 12283 | calcium binding protein 39 [Source:MGI Symbol;Acc:MGI:107438] |
| ENSMUSG00000025534 | Gusb | 0.26614 | 10063.71 | 0.068534 | -3.88335 | 0.000103 | 0.000854 | 0.001035 | 10887.44 | 11090.53 | 8953.417 | 9323.447 | 110006 | glucuronidase, beta [Source:MGI Symbol;Acc:MGI:95872] |
| ENSMUSG00000014956 | Ppp1cb | 0.266277 | 2723.429 | 0.092476 | -2.87943 | 0.003984 | 0.019924 | 0.024095 | 3048.441 | 2899.51 | 2415.073 | 2530.694 | 19046 | protein phosphatase 1 catalytic subunit beta [Source:MGI Symbol;Acc:MGI:104871] |
| ENSMUSG00000032328 | Tmem30a | 0.266531 | 4184.916 | 0.094545 | -2.81909 | 0.004816 | 0.023293 | 0.028154 | 4802.194 | 4338.447 | 3656.901 | 3942.122 | 69981 | transmembrane protein 30A [Source:MGI Symbol;Acc:MGI:106402] |
| ENSMUSG00000029175 | Slc35f6 | 0.266984 | 2228.089 | 0.09617 | -2.77616 | 0.005501 | 0.026015 | 0.031426 | 2374.653 | 2492.32 | 1997.167 | 2048.216 | 74919 | solute carrier family 35, member F6 [Source:MGI Symbol;Acc:MGI:1922169] |
| ENSMUSG00000001642 | Akr1b1 | 0.267338 | 2999.202 | 0.085373 | -3.13143 | 0.00174 | 0.009957 | 0.012062 | 3281.146 | 3271.293 | 2697.639 | 2746.729 | 11677 | aldo-keto reductase family 1 member B [Source:MGI Symbol;Acc:MGI:1353494] |
| ENSMUSG00000037072 | Selenof | 0.268588 | 4983.879 | 0.074936 | -3.58423 | 0.000338 | 0.002457 | 0.002977 | 5397.708 | 5495.103 | 4518.316 | 4524.387 | 93684 | selenoprotein F [Source:MGI Symbol;Acc:MGI:1927947] |
| ENSMUSG00000022378 | Cyrib | 0.27005 | 3779.778 | 0.091014 | -2.96714 | 0.003006 | 0.015663 | 0.018967 | 4305.05 | 3960.763 | 3524.305 | 3328.994 | 223601 | CYFIP related Rac1 interactor B [Source:MGI Symbol;Acc:MGI:1923520] |
| ENSMUSG00000038633 | Degs1 | 0.270253 | 2509.339 | 0.094809 | -2.85049 | 0.004365 | 0.02142 | 0.025898 | 2742.751 | 2744.109 | 2194.689 | 2355.809 | 13244 | delta 4-desaturase, sphingolipid 1 [Source:MGI Symbol;Acc:MGI:1097711] |
| ENSMUSG00000019998 | Stx7 | 0.271824 | 4701.495 | 0.081587 | -3.33169 | 0.000863 | 0.005534 | 0.006704 | 5287.702 | 4998.41 | 4174.482 | 4345.387 | 53331 | syntaxin 7 [Source:MGI Symbol;Acc:MGI:1858210] |
| ENSMUSG00000020368 | Canx | 0.272083 | 15328.1 | 0.064574 | -4.2135 | 2.51E-05 | 0.000245 | 0.000297 | 16994.9 | 16543.34 | 13694.86 | 14079.3 | 12330 | calnexin [Source:MGI Symbol;Acc:MGI:88261] |
| ENSMUSG00000003283 | Hck | 0.273201 | 2114.888 | 0.10233 | -2.6698 | 0.00759 | 0.033978 | 0.041051 | 2404.27 | 2224.794 | 1872.801 | 1957.687 | 15162 | hemopoietic cell kinase [Source:MGI Symbol;Acc:MGI:96052] |
| ENSMUSG00000076432 | Ywhaq | 0.275239 | 4003.448 | 0.089383 | -3.07931 | 0.002075 | 0.011469 | 0.013894 | 4509.196 | 4258.779 | 3478.582 | 3767.236 | 22630 | tyrosine 3-monooxygenase/tryptophan 5-monooxygenase activation protein theta [Source:MGI Symbol;Acc:MGI:891963] |
| ENSMUSG00000022789 | Dnm1l | 0.275498 | 1691.802 | 0.103781 | -2.65461 | 0.00794 | 0.03524 | 0.042564 | 1889.145 | 1816.62 | 1523.48 | 1537.962 | 74006 | dynamin 1-like [Source:MGI Symbol;Acc:MGI:1921256] |
| ENSMUSG00000038965 | Ube2l3 | 0.277497 | 2907.607 | 0.095588 | -2.90305 | 0.003695 | 0.018666 | 0.022581 | 3328.745 | 3044.092 | 2578.76 | 2678.832 | 22195 | ubiquitin-conjugating enzyme E2L 3 [Source:MGI Symbol;Acc:MGI:109240] |
| ENSMUSG00000025503 | Taldo1 | 0.277502 | 3565.493 | 0.093937 | -2.95413 | 0.003136 | 0.016238 | 0.019663 | 3958.108 | 3857.49 | 3397.196 | 3049.178 | 21351 | transaldolase 1 [Source:MGI Symbol;Acc:MGI:1274789] |
| ENSMUSG00000021248 | Tmed10 | 0.277567 | 2504.168 | 0.098572 | -2.81588 | 0.004864 | 0.023467 | 0.028365 | 2868.623 | 2620.182 | 2235.84 | 2292.027 | 68581 | transmembrane p24 trafficking protein 10 [Source:MGI Symbol;Acc:MGI:1915831] |
| ENSMUSG00000028048 | Gba1 | 0.27903 | 3181.934 | 0.087418 | -3.1919 | 0.001413 | 0.008381 | 0.010153 | 3392.21 | 3585.046 | 2898.819 | 2851.66 | 14466 | glucosylceramidase beta 1 [Source:MGI Symbol;Acc:MGI:95665] |
| ENSMUSG00000003970 | Rpl8 | 0.27951 | 8104.903 | 0.070851 | -3.94505 | 7.98E-05 | 0.000683 | 0.000827 | 9071.28 | 8704.432 | 7393.359 | 7250.541 | 26961 | ribosomal protein L8 [Source:MGI Symbol;Acc:MGI:1350927] |
| ENSMUSG00000044533 | Rps2 | 0.279751 | 11227.13 | 0.070264 | -3.98144 | 6.85E-05 | 0.000597 | 0.000723 | 12683.5 | 11941.3 | 10091 | 10192.73 | 16898 | ribosomal protein S2 [Source:MGI Symbol;Acc:MGI:105110] |
| ENSMUSG00000022817 | Itgb5 | 0.279823 | 9937.569 | 0.068208 | -4.10251 | 4.09E-05 | 0.000383 | 0.000464 | 11045.05 | 10752.19 | 9116.19 | 8836.854 | 16419 | integrin beta 5 [Source:MGI Symbol;Acc:MGI:96614] |
| ENSMUSG00000029622 | Arpc1b | 0.280987 | 10087.92 | 0.071028 | -3.95602 | 7.62E-05 | 0.000658 | 0.000797 | 11397.28 | 10737.43 | 9062.237 | 9154.734 | 11867 | actin related protein 2/3 complex, subunit 1B [Source:MGI Symbol;Acc:MGI:1343142] |
| ENSMUSG00000030447 | Cyfip1 | 0.281495 | 11355.29 | 0.065445 | -4.30125 | 1.7E-05 | 0.000173 | 0.00021 | 12588.31 | 12330.79 | 10163.24 | 10338.81 | 20430 | cytoplasmic FMR1 interacting protein 1 [Source:MGI Symbol;Acc:MGI:1338801] |
| ENSMUSG00000036427 | Gpi1 | 0.28171 | 4123.197 | 0.0782 | -3.60244 | 0.000315 | 0.002308 | 0.002796 | 4520.832 | 4528.272 | 3747.432 | 3696.253 | 14751 | glucose-6-phosphate isomerase 1 [Source:MGI Symbol;Acc:MGI:95797] |
| ENSMUSG00000094483 | Purb | 0.282282 | 4510.183 | 0.079569 | -3.54761 | 0.000389 | 0.002776 | 0.003363 | 5024.322 | 4875.466 | 3987.933 | 4153.013 | 19291 | purine rich element binding protein B [Source:MGI Symbol;Acc:MGI:1338779] |
| ENSMUSG00000000078 | Klf6 | 0.283154 | 6064.309 | 0.104271 | -2.71556 | 0.006616 | 0.030236 | 0.036528 | 7172.616 | 6142.28 | 5180.381 | 5761.958 | 23849 | Kruppel-like transcription factor 6 [Source:MGI Symbol;Acc:MGI:1346318] |
| ENSMUSG00000004789 | Dlst | 0.283542 | 1736.787 | 0.101828 | -2.78452 | 0.005361 | 0.025465 | 0.03077 | 1887.029 | 1926.778 | 1573.775 | 1559.566 | 78920 | dihydrolipoamide S-succinyltransferase [Source:MGI Symbol;Acc:MGI:1926170] |
| ENSMUSG00000005161 | Prdx2 | 0.283784 | 2301.734 | 0.101309 | -2.80116 | 0.005092 | 0.024426 | 0.029515 | 2431.772 | 2623.132 | 2139.822 | 2012.21 | 21672 | peroxiredoxin 2 [Source:MGI Symbol;Acc:MGI:109486] |
| ENSMUSG00000021114 | Atp6v1d | 0.284709 | 2226.879 | 0.093587 | -3.0422 | 0.002349 | 0.012732 | 0.015423 | 2464.562 | 2427.406 | 2021.857 | 1993.693 | 73834 | ATPase, H+ transporting, lysosomal V1 subunit D [Source:MGI Symbol;Acc:MGI:1921084] |
| ENSMUSG00000020078 | Vps26a | 0.284875 | 3743.526 | 0.088997 | -3.20094 | 0.00137 | 0.00818 | 0.009909 | 4266.971 | 3956.828 | 3294.777 | 3455.529 | 30930 | VPS26 retromer complex component A [Source:MGI Symbol;Acc:MGI:1353654] |
| ENSMUSG00000008475 | Arpc5 | 0.284962 | 7362.511 | 0.079868 | -3.56789 | 0.00036 | 0.002598 | 0.003147 | 8323.45 | 7850.71 | 6403.92 | 6871.966 | 67771 | actin related protein 2/3 complex, subunit 5 [Source:MGI Symbol;Acc:MGI:1915021] |
| ENSMUSG00000017677 | Wsb1 | 0.285958 | 3044.143 | 0.091901 | -3.1116 | 0.001861 | 0.010512 | 0.012734 | 3472.6 | 3217.198 | 2704.04 | 2782.735 | 78889 | WD repeat and SOCS box-containing 1 [Source:MGI Symbol;Acc:MGI:1926139] |
| ENSMUSG00000024085 | Man2a1 | 0.287102 | 4003.709 | 0.093662 | -3.06531 | 0.002174 | 0.011954 | 0.014481 | 4655.166 | 4146.654 | 3559.969 | 3653.046 | 17158 | mannosidase 2, alpha 1 [Source:MGI Symbol;Acc:MGI:104669] |
| ENSMUSG00000034714 | Ttyh2 | 0.288099 | 2186.165 | 0.105816 | -2.72264 | 0.006476 | 0.029695 | 0.035874 | 2360.902 | 2447.077 | 2076.725 | 1859.957 | 117160 | tweety family member 2 [Source:MGI Symbol;Acc:MGI:2157091] |
| ENSMUSG00000030082 | Sec61a1 | 0.288512 | 3210.233 | 0.085587 | -3.37098 | 0.000749 | 0.0049 | 0.005935 | 3594.241 | 3466.036 | 2854.925 | 2925.729 | 53421 | SEC61 translocon subunit alpha 1 [Source:MGI Symbol;Acc:MGI:1858417] |
| ENSMUSG00000027184 | Caprin1 | 0.289274 | 8805.281 | 0.079987 | -3.61652 | 0.000299 | 0.002197 | 0.002661 | 10141.73 | 9228.666 | 7795.719 | 8055.013 | 53872 | cell cycle associated protein 1 [Source:MGI Symbol;Acc:MGI:1858234] |
| ENSMUSG00000016921 | Srsf6 | 0.289379 | 2728.113 | 0.090619 | -3.19337 | 0.001406 | 0.008343 | 0.010106 | 3053.73 | 2947.704 | 2405.014 | 2506.004 | 67996 | serine and arginine-rich splicing factor 6 [Source:MGI Symbol;Acc:MGI:1915246] |
| ENSMUSG00000034667 | Xpot | 0.289534 | 2255.329 | 0.104731 | -2.76455 | 0.0057 | 0.026726 | 0.032285 | 2604.185 | 2357.573 | 1974.306 | 2085.251 | 73192 | exportin, tRNA (nuclear export receptor for tRNAs) [Source:MGI Symbol;Acc:MGI:1920442] |
| ENSMUSG00000028465 | Tln1 | 0.289635 | 14039.91 | 0.071059 | -4.07598 | 4.58E-05 | 0.000422 | 0.000511 | 15755.21 | 15134.89 | 13025.48 | 12244.03 | 21894 | talin 1 [Source:MGI Symbol;Acc:MGI:1099832] |
| ENSMUSG00000006699 | Cdc42 | 0.290264 | 14969.41 | 0.070632 | -4.10954 | 3.96E-05 | 0.000374 | 0.000453 | 17058.37 | 15882.39 | 13356.51 | 13580.36 | 12540 | cell division cycle 42 [Source:MGI Symbol;Acc:MGI:106211] |
| ENSMUSG00000071042 | Rasgrp3 | 0.290719 | 2669.964 | 0.098337 | -2.95635 | 0.003113 | 0.016132 | 0.019535 | 3061.134 | 2814.925 | 2332.772 | 2471.027 | 240168 | RAS, guanyl releasing protein 3 [Source:MGI Symbol;Acc:MGI:3028579] |
| ENSMUSG00000026618 | Iars2 | 0.291116 | 1432.39 | 0.109059 | -2.66934 | 0.0076 | 0.034014 | 0.041095 | 1592.974 | 1559.913 | 1286.637 | 1290.037 | 381314 | isoleucine-tRNA synthetase 2, mitochondrial [Source:MGI Symbol;Acc:MGI:1919586] |
| ENSMUSG00000039286 | Fndc3b | 0.291258 | 2387.067 | 0.110833 | -2.62791 | 0.008591 | 0.03765 | 0.045477 | 2809.389 | 2445.11 | 2084.04 | 2209.728 | 72007 | fibronectin type III domain containing 3B [Source:MGI Symbol;Acc:MGI:1919257] |
| ENSMUSG00000015745 | Plekho1 | 0.291448 | 3167.639 | 0.08377 | -3.47916 | 0.000503 | 0.003484 | 0.004221 | 3487.408 | 3485.707 | 2862.24 | 2835.2 | 67220 | pleckstrin homology domain containing, family O member 1 [Source:MGI Symbol;Acc:MGI:1914470] |
| ENSMUSG00000038024 | Dennd4c | 0.29218 | 2246.192 | 0.111567 | -2.61889 | 0.00882 | 0.03846 | 0.046456 | 2611.589 | 2333.968 | 1923.096 | 2116.113 | 329877 | DENN domain containing 4C [Source:MGI Symbol;Acc:MGI:1914769] |
| ENSMUSG00000024054 | Smchd1 | 0.292415 | 3078.599 | 0.109966 | -2.65914 | 0.007834 | 0.034866 | 0.042124 | 3605.876 | 3172.938 | 2609.851 | 2925.729 | 74355 | SMC hinge domain containing 1 [Source:MGI Symbol;Acc:MGI:1921605] |
| ENSMUSG00000021423 | Ly86 | 0.293158 | 6376.03 | 0.070963 | -4.13116 | 3.61E-05 | 0.000342 | 0.000414 | 6984.336 | 7058.951 | 5756.487 | 5704.349 | 17084 | lymphocyte antigen 86 [Source:MGI Symbol;Acc:MGI:1321404] |
| ENSMUSG00000019810 | Fuca2 | 0.293387 | 2026.24 | 0.10576 | -2.77408 | 0.005536 | 0.026149 | 0.031587 | 2127.139 | 2335.935 | 1859.085 | 1782.802 | 66848 | fucosidase, alpha-L- 2, plasma [Source:MGI Symbol;Acc:MGI:1914098] |
| ENSMUSG00000039630 | Hnrnpu | 0.293544 | 8660.487 | 0.088049 | -3.33389 | 0.000856 | 0.005498 | 0.00666 | 10176.63 | 8901.143 | 7833.211 | 7730.961 | 51810 | heterogeneous nuclear ribonucleoprotein U [Source:MGI Symbol;Acc:MGI:1858195] |
| ENSMUSG00000057113 | Npm1 | 0.293645 | 5423.914 | 0.081974 | -3.58216 | 0.000341 | 0.002476 | 0.002999 | 6204.773 | 5743.942 | 4949.939 | 4797.002 | 18148 | nucleophosmin 1 [Source:MGI Symbol;Acc:MGI:106184] |
| ENSMUSG00000034663 | Bmp2k | 0.29389 | 7899.386 | 0.078232 | -3.75663 | 0.000172 | 0.001346 | 0.001631 | 9020.508 | 8382.811 | 7262.592 | 6931.632 | 140780 | BMP2 inducible kinase [Source:MGI Symbol;Acc:MGI:2155456] |
| ENSMUSG00000016541 | Atxn10 | 0.294126 | 1369.761 | 0.113592 | -2.58932 | 0.009617 | 0.041319 | 0.049911 | 1523.163 | 1494.999 | 1266.519 | 1194.364 | 54138 | ataxin 10 [Source:MGI Symbol;Acc:MGI:1859293] |
| ENSMUSG00000017421 | Zfp207 | 0.294462 | 2286.501 | 0.095925 | -3.06973 | 0.002143 | 0.011805 | 0.0143 | 2549.182 | 2489.369 | 2109.645 | 1997.808 | 22680 | zinc finger protein 207 [Source:MGI Symbol;Acc:MGI:1340045] |
| ENSMUSG00000019960 | Dusp6 | 0.294487 | 7454.295 | 0.10047 | -2.93109 | 0.003378 | 0.01728 | 0.020918 | 8326.623 | 8099.548 | 7293.684 | 6097.326 | 67603 | dual specificity phosphatase 6 [Source:MGI Symbol;Acc:MGI:1914853] |
| ENSMUSG00000034854 | Mfsd12 | 0.295102 | 2419.295 | 0.102409 | -2.8816 | 0.003957 | 0.019813 | 0.023962 | 2536.489 | 2795.254 | 2230.353 | 2115.084 | 73822 | major facilitator superfamily domain containing 12 [Source:MGI Symbol;Acc:MGI:3604804] |
| ENSMUSG00000004455 | Ppp1cc;Ppp1cc | 0.295428 | 6431.489 | 0.079859 | -3.69935 | 0.000216 | 0.001649 | 0.001998 | 7308.008 | 6867.158 | 5612.918 | 5937.872 | 19047;434233 | protein phosphatase 1 catalytic subunit gamma [Source:MGI Symbol;Acc:MGI:104872];protein phosphatase 1 catalytic subunit gamma [Source:MGI Symbol;Acc:MGI:104872] |
| ENSMUSG00000024695 | Zfp91 | 0.2969 | 3097.146 | 0.092132 | -3.22256 | 0.001271 | 0.007667 | 0.009287 | 3480.004 | 3348.994 | 2672.949 | 2886.637 | 109910 | zinc finger protein 91 [Source:MGI Symbol;Acc:MGI:104854] |
| ENSMUSG00000041133 | Smc1a | 0.297444 | 1610.856 | 0.106321 | -2.7976 | 0.005148 | 0.024634 | 0.029766 | 1800.294 | 1752.689 | 1475.928 | 1414.514 | 24061 | structural maintenance of chromosomes 1A [Source:MGI Symbol;Acc:MGI:1344345] |
| ENSMUSG00000025651 | Uqcrc1 | 0.297634 | 2329.164 | 0.097178 | -3.06276 | 0.002193 | 0.012027 | 0.014569 | 2481.486 | 2655.59 | 2116.961 | 2062.618 | 22273 | ubiquinol-cytochrome c reductase core protein 1 [Source:MGI Symbol;Acc:MGI:107876] |
| ENSMUSG00000066232 | Ipo7 | 0.297824 | 3302.777 | 0.113839 | -2.61619 | 0.008892 | 0.038687 | 0.04673 | 3950.703 | 3334.24 | 2826.577 | 3099.586 | 233726 | importin 7 [Source:MGI Symbol;Acc:MGI:2152414] |
| ENSMUSG00000024286 | Ccny | 0.299975 | 3636.144 | 0.08669 | -3.4603 | 0.00054 | 0.003705 | 0.004489 | 4149.561 | 3876.177 | 3220.706 | 3298.132 | 67974 | cyclin Y [Source:MGI Symbol;Acc:MGI:1915224] |
| ENSMUSG00000023277 | Twf2 | 0.30031 | 1712.748 | 0.104267 | -2.88019 | 0.003974 | 0.019882 | 0.024045 | 1911.358 | 1869.732 | 1571.032 | 1498.87 | 23999 | twinfilin actin binding protein 2 [Source:MGI Symbol;Acc:MGI:1346078] |
| ENSMUSG00000029713 | Gnb2 | 0.301002 | 3838.175 | 0.084783 | -3.55028 | 0.000385 | 0.002752 | 0.003333 | 4299.762 | 4175.177 | 3542.594 | 3335.166 | 14693 | guanine nucleotide binding protein (G protein), beta 2 [Source:MGI Symbol;Acc:MGI:95784] |
| ENSMUSG00000022710 | Usp7 | 0.301909 | 2542.931 | 0.09765 | -3.09173 | 0.00199 | 0.011097 | 0.013443 | 2935.262 | 2681.162 | 2279.733 | 2275.567 | 252870 | ubiquitin specific peptidase 7 [Source:MGI Symbol;Acc:MGI:2182061] |
| ENSMUSG00000029103 | Lrpap1 | 0.302376 | 1919.739 | 0.097951 | -3.087 | 0.002022 | 0.011213 | 0.013584 | 2121.85 | 2118.57 | 1724.66 | 1713.876 | 16976 | low density lipoprotein receptor-related protein associated protein 1 [Source:MGI Symbol;Acc:MGI:96829] |
| ENSMUSG00000039230 | Tbcd | 0.302897 | 1441.786 | 0.108986 | -2.77922 | 0.005449 | 0.025819 | 0.031189 | 1606.725 | 1578.6 | 1303.097 | 1278.721 | 108903 | tubulin-specific chaperone d [Source:MGI Symbol;Acc:MGI:1919686] |
| ENSMUSG00000050721 | Plekho2 | 0.303322 | 1952.306 | 0.106218 | -2.85566 | 0.004295 | 0.021164 | 0.025588 | 2076.367 | 2236.597 | 1690.825 | 1805.434 | 102595 | pleckstrin homology domain containing, family O member 2 [Source:MGI Symbol;Acc:MGI:2143132] |
| ENSMUSG00000033417 | Cacul1 | 0.303836 | 3909.731 | 0.081998 | -3.70542 | 0.000211 | 0.001613 | 0.001954 | 4345.245 | 4294.187 | 3416.399 | 3583.092 | 78832 | CDK2 associated, cullin domain 1 [Source:MGI Symbol;Acc:MGI:1926082] |
| ENSMUSG00000025016 | Tm9sf3 | 0.304119 | 9834.752 | 0.085312 | -3.5648 | 0.000364 | 0.002623 | 0.003177 | 11387.76 | 10346.96 | 8433.093 | 9171.193 | 107358 | transmembrane 9 superfamily member 3 [Source:MGI Symbol;Acc:MGI:1914262] |
| ENSMUSG00000032872 | Cyb5r4;Cyb5r4 | 0.304677 | 1687.732 | 0.106199 | -2.86892 | 0.004119 | 0.020483 | 0.024772 | 1903.953 | 1826.455 | 1472.271 | 1548.25 | 266690;12395 | cytochrome b5 reductase 4 [Source:MGI Symbol;Acc:MGI:2386848];cytochrome b5 reductase 4 [Source:MGI Symbol;Acc:MGI:2386848] |
| ENSMUSG00000024327 | Slc39a7 | 0.304725 | 1509.57 | 0.10783 | -2.82597 | 0.004714 | 0.022854 | 0.027632 | 1665.959 | 1671.054 | 1376.253 | 1325.014 | 14977 | solute carrier family 39 (zinc transporter), member 7 [Source:MGI Symbol;Acc:MGI:95909] |
| ENSMUSG00000028410 | Dnaja1 | 0.305306 | 2572.923 | 0.093609 | -3.26149 | 0.001108 | 0.006834 | 0.008278 | 2874.97 | 2812.958 | 2229.438 | 2374.326 | 15502 | DnaJ heat shock protein family (Hsp40) member A1 [Source:MGI Symbol;Acc:MGI:1270129] |
| ENSMUSG00000071533 | Pcnp | 0.305535 | 1606.104 | 0.108882 | -2.80611 | 0.005014 | 0.024115 | 0.029147 | 1725.193 | 1825.472 | 1403.687 | 1470.066 | 76302 | PEST proteolytic signal containing nuclear protein [Source:MGI Symbol;Acc:MGI:1923552] |
| ENSMUSG00000021374 | Nup153 | 0.305967 | 2463.681 | 0.106665 | -2.86848 | 0.004124 | 0.020498 | 0.02479 | 2872.854 | 2574.938 | 2116.961 | 2289.969 | 218210 | nucleoporin 153 [Source:MGI Symbol;Acc:MGI:2385621] |
| ENSMUSG00000030339 | Ltbr | 0.306342 | 1232.688 | 0.115989 | -2.64113 | 0.008263 | 0.036443 | 0.044018 | 1342.287 | 1383.857 | 1111.061 | 1093.548 | 17000 | lymphotoxin B receptor [Source:MGI Symbol;Acc:MGI:104875] |
| ENSMUSG00000035898 | Uba6 | 0.306398 | 1240.474 | 0.117109 | -2.61635 | 0.008888 | 0.03868 | 0.046722 | 1406.81 | 1336.647 | 1099.173 | 1119.266 | 231380 | ubiquitin-like modifier activating enzyme 6 [Source:MGI Symbol;Acc:MGI:1913894] |
| ENSMUSG00000042524 | Sun2 | 0.306459 | 1908.358 | 0.100896 | -3.03738 | 0.002386 | 0.012914 | 0.015644 | 2165.218 | 2055.623 | 1718.259 | 1694.33 | 223697 | Sad1 and UNC84 domain containing 2 [Source:MGI Symbol;Acc:MGI:2443011] |
| ENSMUSG00000028409 | Smu1 | 0.307057 | 1614.396 | 0.110619 | -2.77582 | 0.005506 | 0.026026 | 0.031439 | 1800.294 | 1770.393 | 1380.825 | 1506.071 | 74255 | smu-1 suppressor of mec-8 and unc-52 homolog (C. elegans) [Source:MGI Symbol;Acc:MGI:1915546] |
| ENSMUSG00000032612 | Usp4 | 0.307637 | 1636.725 | 0.104885 | -2.93307 | 0.003356 | 0.017199 | 0.02082 | 1807.698 | 1813.669 | 1491.474 | 1434.06 | 22258 | ubiquitin specific peptidase 4 (proto-oncogene) [Source:MGI Symbol;Acc:MGI:98905] |
| ENSMUSG00000007815 | Rhoa | 0.307838 | 5742.674 | 0.095493 | -3.22367 | 0.001266 | 0.00765 | 0.009267 | 6766.439 | 5940.652 | 5318.463 | 4945.14 | 11848 | ras homolog family member A [Source:MGI Symbol;Acc:MGI:1096342] |
| ENSMUSG00000024091 | Vapa | 0.308419 | 2445.992 | 0.095344 | -3.23482 | 0.001217 | 0.007405 | 0.00897 | 2780.83 | 2631.984 | 2139.822 | 2231.331 | 30960 | vesicle-associated membrane protein, associated protein A [Source:MGI Symbol;Acc:MGI:1353561] |
| ENSMUSG00000014496 | Ankrd28 | 0.309863 | 1840.187 | 0.108151 | -2.8651 | 0.004169 | 0.020653 | 0.02497 | 2042.519 | 2031.034 | 1560.973 | 1726.221 | 105522 | ankyrin repeat domain 28 [Source:MGI Symbol;Acc:MGI:2145661] |
| ENSMUSG00000007739 | Cct4 | 0.31004 | 2978.64 | 0.095744 | -3.23823 | 0.001203 | 0.007339 | 0.00889 | 3459.907 | 3135.563 | 2687.58 | 2631.51 | 12464 | chaperonin containing TCP1 subunit 4 [Source:MGI Symbol;Acc:MGI:104689] |
| ENSMUSG00000001833 | Septin7 | 0.310353 | 3900.132 | 0.083819 | -3.70266 | 0.000213 | 0.001629 | 0.001973 | 4191.871 | 4443.687 | 3445.662 | 3519.31 | 235072 | septin 7 [Source:MGI Symbol;Acc:MGI:1335094] |
| ENSMUSG00000007564 | Ppp2r1a | 0.310448 | 2396.765 | 0.118839 | -2.61234 | 0.008993 | 0.039014 | 0.047126 | 2885.547 | 2422.488 | 2210.235 | 2068.791 | 51792 | protein phosphatase 2, regulatory subunit A, alpha [Source:MGI Symbol;Acc:MGI:1926334] |
| ENSMUSG00000074781 | Ube2n | 0.310475 | 1564.935 | 0.117455 | -2.64336 | 0.008209 | 0.036257 | 0.043793 | 1650.093 | 1814.653 | 1342.418 | 1452.577 | 93765 | ubiquitin-conjugating enzyme E2N [Source:MGI Symbol;Acc:MGI:1934835] |
| ENSMUSG00000030272 | Camk1 | 0.310808 | 5045.407 | 0.077625 | -4.00398 | 6.23E-05 | 0.00055 | 0.000666 | 5550.024 | 5622.965 | 4395.78 | 4612.858 | 52163 | calcium/calmodulin-dependent protein kinase I [Source:MGI Symbol;Acc:MGI:1098535] |
| ENSMUSG00000023915 | Tnfrsf21 | 0.310881 | 1426.933 | 0.110033 | -2.82536 | 0.004723 | 0.022882 | 0.027665 | 1553.838 | 1606.14 | 1262.861 | 1284.893 | 94185 | tumor necrosis factor receptor superfamily, member 21 [Source:MGI Symbol;Acc:MGI:2151075] |
| ENSMUSG00000025809 | Itgb1 | 0.311282 | 9552.138 | 0.087173 | -3.57083 | 0.000356 | 0.002571 | 0.003114 | 11160.34 | 9996.819 | 8198.079 | 8853.314 | 16412 | integrin beta 1 (fibronectin receptor beta) [Source:MGI Symbol;Acc:MGI:96610] |
| ENSMUSG00000061136 | Prpf40a | 0.31189 | 2464.489 | 0.101552 | -3.07123 | 0.002132 | 0.011754 | 0.014239 | 2868.623 | 2591.659 | 2240.412 | 2157.262 | 56194 | pre-mRNA processing factor 40A [Source:MGI Symbol;Acc:MGI:1860512] |
| ENSMUSG00000028180 | Zranb2 | 0.312327 | 1502.05 | 0.109043 | -2.86426 | 0.00418 | 0.020685 | 0.025008 | 1668.075 | 1660.235 | 1374.424 | 1305.468 | 53861 | zinc finger, RAN-binding domain containing 2 [Source:MGI Symbol;Acc:MGI:1858211] |
| ENSMUSG00000043535 | Setx | 0.31274 | 3085.6 | 0.090842 | -3.44267 | 0.000576 | 0.003915 | 0.004742 | 3547.7 | 3289.98 | 2733.302 | 2771.418 | 269254 | senataxin [Source:MGI Symbol;Acc:MGI:2443480] |
| ENSMUSG00000041697 | Cox6a1 | 0.313373 | 1416.91 | 0.112001 | -2.79795 | 0.005143 | 0.024615 | 0.029743 | 1579.224 | 1560.897 | 1227.197 | 1300.324 | 12861 | cytochrome c oxidase subunit 6A1 [Source:MGI Symbol;Acc:MGI:103099] |
| ENSMUSG00000026159 | Agfg1 | 0.313521 | 2186.945 | 0.113769 | -2.75578 | 0.005855 | 0.027343 | 0.033031 | 2568.222 | 2278.889 | 1854.512 | 2046.159 | 15463 | ArfGAP with FG repeats 1 [Source:MGI Symbol;Acc:MGI:1333754] |

| | | | |
|---|---|---|---|
| [illegible] | [illegible] | [illegible] | myosin, light polypeptide 6, alkali, smooth muscle and non-muscle [Source:MGI Symbol;Acc:MGI:109318] |
| [illegible] | [illegible] | [illegible] | DnaJ heat shock protein family (Hsp40) member A2 [Source:MGI Symbol;Acc:MGI:1931882] |
| [illegible] | [illegible] | [illegible] | vav 1 oncogene [Source:MGI Symbol;Acc:MGI:98923] |
| [illegible] | [illegible] | [illegible] | annexin A7 [Source:MGI Symbol;Acc:MGI:88031] |
| [illegible] | [illegible] | [illegible] | STE20-like kinase [Source:MGI Symbol;Acc:MGI:103241] |
| [illegible] | [illegible] | [illegible] | hnRNP-associated with lethal yellow [Source:MGI Symbol;Acc:MGI:97850] |
| [illegible] | [illegible] | [illegible] | NLR family, apoptosis inhibitory protein 2 [Source:MGI Symbol;Acc:MGI:1298226] |
| [illegible] | [illegible] | [illegible] | v-maf musculoaponeurotic fibrosarcoma oncogene family, protein G (avian) [Source:MGI Symbol;Acc:MGI:96911] |
| [illegible] | [illegible] | [illegible] | deltex 4, E3 ubiquitin ligase [Source:MGI Symbol;Acc:MGI:2672905] |
| [illegible] | [illegible] | [illegible] | cytochrome c oxidase subunit 5B [Source:MGI Symbol;Acc:MGI:88475] |
| [illegible] | [illegible] | [illegible] | vesicle amine transport 1 [Source:MGI Symbol;Acc:MGI:1349450] |
| [illegible] | [illegible] | [illegible] | eukaryotic translation initiation factor 3, subunit C [Source:MGI Symbol;Acc:MGI:1926966] |
| [illegible] | [illegible] | [illegible] | heat shock protein 70 family, member 13 [Source:MGI Symbol;Acc:MGI:1309463] |
| [illegible] | [illegible] | [illegible] | proteasome (prosome, macropain) 26S subunit, non-ATPase, 4 [Source:MGI Symbol;Acc:MGI:1201670] |
| [illegible] | [illegible] | [illegible] | RAB8A, member RAS oncogene family [Source:MGI Symbol;Acc:MGI:96960] |
| [illegible] | [illegible] | [illegible] | disabled 2, mitogen-responsive phosphoprotein [Source:MGI Symbol;Acc:MGI:109175] |
| [illegible] | [illegible] | [illegible] | syntrophin, basic 2 [Source:MGI Symbol;Acc:MGI:101771] |
| [illegible] | [illegible] | [illegible] | signal peptidase complex subunit 2 homolog (S. cerevisiae) [Source:MGI Symbol;Acc:MGI:1913874] |
| [illegible] | [illegible] | [illegible] | short coiled-coil protein [Source:MGI Symbol;Acc:MGI:1927654] |
| [illegible] | [illegible] | [illegible] | annexin A4 [Source:MGI Symbol;Acc:MGI:88030] |
| [illegible] | [illegible] | [illegible] | [illegible] |

| | | | | | | | | | | | | | | |
|---|---|---|---|---|---|---|---|---|---|---|---|---|---|---|
| ENSMUSG00000025474 | Tubgcp2 | 0.473078 | 700.6953 | 0.148397 | -3.18791 | 0.001433 | 0.008475 | 0.010266 | 830.3353 | 798.644 | 567.8758 | 605.9263 | 74237 | tubulin, gamma complex component 2 [Source:MGI Symbol;Acc:MGI:1921487] |
| ENSMUSG00000046722 | Cdc42se1 | 0.473577 | 3544.322 | 0.082596 | -5.73369 | 9.83E-09 | 1.92E-07 | 2.33E-07 | 4170.716 | 4070.92 | 2931.739 | 3003.913 | 57912 | CDC42 small effector 1 [Source:MGI Symbol;Acc:MGI:1889510] |
| ENSMUSG00000079111 | Kdelr2 | 0.473699 | 1057.105 | 0.123527 | -3.83477 | 0.000126 | 0.00102 | 0.001235 | 1253.436 | 1204.851 | 883.3624 | 886.7716 | 66913 | KDEL (Lys-Asp-Glu-Leu) endoplasmic reticulum protein retention receptor 2 [Source:MGI Symbol;Acc:MGI:1914163] |
| ENSMUSG00000037361 | Sf3b6 | 0.474271 | 748.8038 | 0.145877 | -3.25118 | 0.001149 | 0.007041 | 0.008529 | 890.6271 | 850.7722 | 602.6251 | 651.1907 | 66055 | splicing factor 3B, subunit 6 [Source:MGI Symbol;Acc:MGI:1913305] |
| ENSMUSG00000022561 | Gpaa1 | 0.474712 | 665.6709 | 0.153561 | -3.09135 | 0.001992 | 0.011099 | 0.013445 | 807.0647 | 741.598 | 568.7903 | 545.2308 | 14731 | GPI anchor attachment protein 1 [Source:MGI Symbol;Acc:MGI:1202392] |
| ENSMUSG00000018102 | H2bc4 | 0.475454 | 1265.733 | 0.133718 | -3.55565 | 0.000377 | 0.002706 | 0.003278 | 1523.163 | 1421.232 | 976.6367 | 1141.898 | 68024 | H2B clustered histone 4 [Source:MGI Symbol;Acc:MGI:1915274] |
| ENSMUSG00000025858 | Get4 | 0.476557 | 451.5044 | 0.183669 | -2.59465 | 0.009469 | 0.040742 | 0.049213 | 555.3198 | 495.7101 | 387.7284 | 367.2592 | 67604 | golgi to ER traffic protein 4 [Source:MGI Symbol;Acc:MGI:1914854] |
| ENSMUSG00000038607 | Gng10 | 0.476828 | 730.9567 | 0.149551 | -3.1884 | 0.001431 | 0.008468 | 0.010257 | 895.9159 | 805.5288 | 607.1973 | 615.1849 | 14700 | guanine nucleotide binding protein (G protein), gamma 10 [Source:MGI Symbol;Acc:MGI:1336169] |
| ENSMUSG00000038975 | Rabggtb | 0.477374 | 550.0724 | 0.161733 | -2.95161 | 0.003161 | 0.016348 | 0.019797 | 629.3624 | 651.1112 | 459.9703 | 459.8456 | 19352 | Rab geranylgeranyl transferase, b subunit [Source:MGI Symbol;Acc:MGI:99537] |
| ENSMUSG00000028932 | Psmc2 | 0.477434 | 1029.775 | 0.12994 | -3.67425 | 0.000239 | 0.001805 | 0.002186 | 1220.646 | 1176.328 | 823.0084 | 899.1164 | 19181 | proteasome (prosome, macropain) 26S subunit, ATPase 2 [Source:MGI Symbol;Acc:MGI:109555] |
| ENSMUSG00000026926 | Pmpca | 0.477573 | 1281.024 | 0.117614 | -4.06052 | 4.9E-05 | 0.000446 | 0.00054 | 1512.585 | 1469.426 | 1035.162 | 1106.921 | 66865 | peptidase (mitochondrial processing) alpha [Source:MGI Symbol;Acc:MGI:1918568] |
| ENSMUSG00000041390 | Mdfic | 0.478194 | 2977.443 | 0.088839 | -5.38268 | 7.34E-08 | 1.24E-06 | 1.51E-06 | 3562.508 | 3370.632 | 2497.373 | 2479.257 | 16543 | MyoD family inhibitor domain containing [Source:MGI Symbol;Acc:MGI:104611] |
| ENSMUSG00000000902 | Smarcb1 | 0.479293 | 472.8027 | 0.175575 | -2.72985 | 0.006336 | 0.029176 | 0.035247 | 570.1283 | 531.1179 | 387.7284 | 402.2363 | 20587 | SWI/SNF related BAF chromatin remodeling complex subunit B1 [Source:MGI Symbol;Acc:MGI:1328366] |
| ENSMUSG00000032301 | Psma4 | 0.479614 | 1798.52 | 0.102486 | -4.67981 | 2.87E-06 | 3.43E-05 | 4.16E-05 | 2107.042 | 2082.179 | 1465.869 | 1538.991 | 26441 | proteasome subunit alpha 4 [Source:MGI Symbol;Acc:MGI:1347060] |
| ENSMUSG00000036966 | Spryd3 | 0.48033 | 426.2859 | 0.183626 | -2.61581 | 0.008902 | 0.038709 | 0.046756 | 485.5081 | 507.5127 | 343.8346 | 368.288 | 223918 | SPRY domain containing 3 [Source:MGI Symbol;Acc:MGI:2446175] |
| ENSMUSG00000028757 | Ddost | 0.480446 | 2796.79 | 0.08873 | -5.41469 | 6.14E-08 | 1.06E-06 | 1.28E-06 | 3319.226 | 3197.527 | 2354.719 | 2315.688 | 13200 | dolichyl-di-phosphooligosaccharide-protein glycotransferase [Source:MGI Symbol;Acc:MGI:1194508] |
| ENSMUSG00000035726 | Supt16 | 0.480505 | 2441.808 | 0.092961 | -5.16888 | 2.36E-07 | 3.54E-06 | 4.29E-06 | 2834.775 | 2854.267 | 1989.852 | 2088.337 | 114741 | SPT16, facilitates chromatin remodeling subunit [Source:MGI Symbol;Acc:MGI:1890948] |
| ENSMUSG00000031545 | Gpat4 | 0.480838 | 1473.974 | 0.131775 | -3.64892 | 0.000263 | 0.001971 | 0.002388 | 1789.716 | 1644.498 | 1128.436 | 1333.244 | 102247 | glycerol-3-phosphate acyltransferase 4 [Source:MGI Symbol;Acc:MGI:2142716] |
| ENSMUSG00000017801 | Mlx | 0.4814 | 641.7904 | 0.158677 | -3.03383 | 0.002415 | 0.013048 | 0.015807 | 780.6209 | 715.0421 | 513.923 | 557.5756 | 21428 | MAX-like protein X [Source:MGI Symbol;Acc:MGI:108398] |
| ENSMUSG00000000561 | Wdr77 | 0.481458 | 575.1771 | 0.159568 | -3.01725 | 0.002551 | 0.013643 | 0.016521 | 686.481 | 654.0619 | 474.6015 | 485.564 | 70465 | WD repeat domain 77 [Source:MGI Symbol;Acc:MGI:1917715] |
| ENSMUSG00000041057 | Wdr43 | 0.481788 | 1014.848 | 0.127959 | -3.76518 | 0.000166 | 0.001306 | 0.001582 | 1220.646 | 1144.854 | 834.8963 | 858.9957 | 72515 | WD repeat domain 43 [Source:MGI Symbol;Acc:MGI:1919765] |
| ENSMUSG00000078440 | Dohh | 0.482045 | 513.975 | 0.173603 | -2.7767 | 0.005491 | 0.025988 | 0.031393 | 633.5934 | 564.5587 | 421.5632 | 436.1846 | 102115 | deoxyhypusine hydroxylase/monooxygenase [Source:MGI Symbol;Acc:MGI:1915964] |
| ENSMUSG00000087260 | Lamtor5 | 0.482103 | 709.0161 | 0.155876 | -3.09286 | 0.001982 | 0.011078 | 0.013419 | 857.8368 | 794.7098 | 555.9879 | 627.5298 | 68576 | late endosomal/lysosomal adaptor, MAPK and MTOR activator 5 [Source:MGI Symbol;Acc:MGI:1915826] |
| ENSMUSG00000037331 | Larp1 | 0.482328 | 1510.675 | 0.12561 | -3.83989 | 0.000123 | 0.001 | 0.001212 | 1882.798 | 1639.581 | 1309.498 | 1210.824 | 73158 | La ribonucleoprotein 1, translational regulator [Source:MGI Symbol;Acc:MGI:1890165] |
| ENSMUSG00000025027 | Xpnpep1 | 0.482458 | 1209.736 | 0.118076 | -4.08599 | 4.39E-05 | 0.000407 | 0.000493 | 1419.503 | 1400.578 | 983.9523 | 1034.91 | 170750 | X-prolyl aminopeptidase (aminopeptidase P) 1, soluble [Source:MGI Symbol;Acc:MGI:2180003] |
| ENSMUSG00000026078 | Pdcl3 | 0.482553 | 652.3012 | 0.152236 | -3.16977 | 0.001526 | 0.008938 | 0.010828 | 747.8306 | 773.0716 | 560.5602 | 527.7423 | 68833 | phosducin-like 3 [Source:MGI Symbol;Acc:MGI:1916083] |
| ENSMUSG00000026229 | Psmd1 | 0.483315 | 2381.61 | 0.09982 | -4.84186 | 1.29E-06 | 1.66E-05 | 2.01E-05 | 2897.182 | 2656.573 | 1953.273 | 2019.411 | 70247 | proteasome (prosome, macropain) 26S subunit, non-ATPase, 1 [Source:MGI Symbol;Acc:MGI:1917497] |
| ENSMUSG00000109511 | Nup62 | 0.48392 | 607.2685 | 0.156663 | -3.08892 | 0.002009 | 0.011166 | 0.013526 | 730.9066 | 685.5355 | 507.5219 | 505.11 | 18226 | nucleoporin 62 [Source:MGI Symbol;Acc:MGI:1351500] |
| ENSMUSG00000021737 | Psmd6 | 0.484052 | 1426.234 | 0.118848 | -4.07287 | 4.64E-05 | 0.000426 | 0.000517 | 1740.002 | 1586.469 | 1152.212 | 1226.255 | 66413 | proteasome (prosome, macropain) 26S subunit, non-ATPase, 6 [Source:MGI Symbol;Acc:MGI:1913663] |
| ENSMUSG00000031485 | Plpbp | 0.484305 | 1025.371 | 0.124106 | -3.90236 | 9.53E-05 | 0.000794 | 0.000962 | 1201.606 | 1190.098 | 847.6987 | 862.0819 | 114863 | pyridoxal phosphate binding protein [Source:MGI Symbol;Acc:MGI:1891207] |
| ENSMUSG00000032299 | Commd4 | 0.484389 | 621.8462 | 0.156371 | -3.0977 | 0.00195 | 0.010936 | 0.013248 | 708.6938 | 741.598 | 501.1207 | 535.9722 | 66199 | COMM domain containing 4 [Source:MGI Symbol;Acc:MGI:1913449] |
| ENSMUSG00000021024 | Psma6 | 0.484796 | 1331.604 | 0.129343 | -3.74815 | 0.000178 | 0.001389 | 0.001682 | 1660.671 | 1445.821 | 1070.825 | 1149.1 | 26443 | proteasome subunit alpha 6 [Source:MGI Symbol;Acc:MGI:1347006] |
| ENSMUSG00000006998 | Psmd2 | 0.485349 | 4373.62 | 0.08136 | -5.96542 | 2.44E-09 | 5.19E-08 | 6.29E-08 | 5160.772 | 5043.653 | 3544.423 | 3745.633 | 21762 | proteasome (prosome, macropain) 26S subunit, non-ATPase, 2 [Source:MGI Symbol;Acc:MGI:1096584] |
| ENSMUSG00000023021 | Cers5 | 0.48535 | 912.2951 | 0.13591 | -3.57112 | 0.000355 | 0.00257 | 0.003114 | 1111.697 | 1016.992 | 748.9377 | 771.553 | 71949 | ceramide synthase 5 [Source:MGI Symbol;Acc:MGI:1919199] |
| ENSMUSG00000030986 | Dhx32 | 0.486054 | 583.3308 | 0.159726 | -3.04306 | 0.002342 | 0.0127 | 0.015385 | 661.0949 | 700.2888 | 495.634 | 476.3054 | 101437 | DEAH-box helicase 32 (putative) [Source:MGI Symbol;Acc:MGI:2141813] |
| ENSMUSG00000063884 | Ptcd3 | 0.486314 | 736.9779 | 0.145396 | -3.34476 | 0.000824 | 0.005314 | 0.006437 | 878.9918 | 840.9367 | 595.3094 | 632.6735 | 69956 | pentatricopeptide repeat domain 3 [Source:MGI Symbol;Acc:MGI:1917206] |
| ENSMUSG00000029538 | Srsf9 | 0.486625 | 712.1281 | 0.160915 | -3.0241 | 0.002494 | 0.013375 | 0.016197 | 898.0314 | 764.2197 | 575.1915 | 611.07 | 108014 | serine and arginine-rich splicing factor 9 [Source:MGI Symbol;Acc:MGI:104896] |
| ENSMUSG00000000058 | Cav2 | 0.48679 | 546.0128 | 0.166959 | -2.91563 | 0.00355 | 0.018022 | 0.021809 | 611.3806 | 662.9138 | 441.6812 | 468.0755 | 12390 | caveolin 2 [Source:MGI Symbol;Acc:MGI:107571] |
| ENSMUSG00000028455 | Stoml2 | 0.487513 | 662.6783 | 0.161512 | -3.01843 | 0.002541 | 0.0136 | 0.016468 | 765.8124 | 780.94 | 509.3508 | 594.6102 | 66592 | stomatin (Epb7.2)-like 2 [Source:MGI Symbol;Acc:MGI:1913842] |
| ENSMUSG00000006740 | Kif5b | 0.488128 | 5661.659 | 0.091821 | -5.31606 | 1.06E-07 | 1.72E-06 | 2.09E-06 | 6759.035 | 6460.951 | 4417.726 | 5008.922 | 16573 | kinesin family member 5B [Source:MGI Symbol;Acc:MGI:1098268] |
| ENSMUSG00000053581 | Zfand2a | 0.48837 | 729.3661 | 0.159494 | -3.062 | 0.002199 | 0.012053 | 0.014601 | 859.9523 | 842.9038 | 554.159 | 660.4494 | 100494 | zinc finger, AN1-type domain 2A [Source:MGI Symbol;Acc:MGI:2140729] |
| ENSMUSG00000079293 | Clec7a | 0.488655 | 3332.659 | 0.085603 | -5.70838 | 1.14E-08 | 2.21E-07 | 2.67E-07 | 3802.618 | 3980.434 | 2744.276 | 2803.309 | 56644 | C-type lectin domain family 7, member a [Source:MGI Symbol;Acc:MGI:1861431] |
| ENSMUSG00000002881 | Nab1 | 0.488743 | 1196.394 | 0.129334 | -3.77893 | 0.000158 | 0.001243 | 0.001505 | 1489.315 | 1305.173 | 992.1824 | 998.904 | 17936 | Ngfi-A binding protein 1 [Source:MGI Symbol;Acc:MGI:107564] |
| ENSMUSG00000000253 | Gmpr | 0.488794 | 476.3018 | 0.176747 | -2.7655 | 0.005684 | 0.026681 | 0.032231 | 530.9915 | 581.2791 | 388.6429 | 404.2938 | 66355 | guanosine monophosphate reductase [Source:MGI Symbol;Acc:MGI:1913605] |
| ENSMUSG00000030057 | Cnbp | 0.488892 | 3093.417 | 0.105453 | -4.6361 | 3.55E-06 | 4.16E-05 | 5.04E-05 | 3794.156 | 3430.628 | 2427.875 | 2721.01 | 12785 | cellular nucleic acid binding protein [Source:MGI Symbol;Acc:MGI:88431] |
| ENSMUSG00000025630 | Hprt1 | 0.489087 | 3006.225 | 0.110852 | -4.41207 | 1.02E-05 | 0.00011 | 0.000134 | 3714.825 | 3306.701 | 2341.002 | 2662.372 | 15452 | hypoxanthine phosphoribosyltransferase 1 [Source:MGI Symbol;Acc:MGI:96217] |
| ENSMUSG00000030588 | Yif1b | 0.48916 | 584.4625 | 0.157389 | -3.10797 | 0.001884 | 0.010626 | 0.012872 | 673.788 | 691.4368 | 490.1472 | 482.4778 | 77254 | Yip1 interacting factor homolog B (S. cerevisiae) [Source:MGI Symbol;Acc:MGI:1924504] |
| ENSMUSG00000061613 | U2af1 | 0.489517 | 436.1443 | 0.179434 | -2.72812 | 0.00637 | 0.029294 | 0.03539 | 517.2407 | 501.6114 | 358.4659 | 367.2592 | 108121 | U2 small nuclear ribonucleoprotein auxiliary factor (U2AF) 1 [Source:MGI Symbol;Acc:MGI:98884] |
| ENSMUSG00000040463 | Mybbp1a | 0.489692 | 887.5636 | 0.165208 | -2.96409 | 0.003036 | 0.015791 | 0.019123 | 1159.296 | 914.7031 | 765.3978 | 710.8575 | 18432 | MYB binding protein (P160) 1a [Source:MGI Symbol;Acc:MGI:106181] |
| ENSMUSG00000025239 | Limd1 | 0.490092 | 1389.478 | 0.114462 | -4.28171 | 1.85E-05 | 0.000187 | 0.000226 | 1681.826 | 1564.831 | 1154.955 | 1156.301 | 29806 | LIM domains containing 1 [Source:MGI Symbol;Acc:MGI:1352502] |
| ENSMUSG00000027067 | Ssrp1 | 0.49017 | 1696.668 | 0.112173 | -4.36975 | 1.24E-05 | 0.000131 | 0.000159 | 2085.887 | 1878.584 | 1396.371 | 1425.83 | 20833 | structure specific recognition protein 1 [Source:MGI Symbol;Acc:MGI:107912] |
| ENSMUSG00000014074 | Rnf168 | 0.490527 | 636.2631 | 0.174561 | -2.81006 | 0.004953 | 0.023833 | 0.028807 | 807.0647 | 679.6342 | 495.634 | 562.7193 | 70238 | ring finger protein 168 [Source:MGI Symbol;Acc:MGI:1917488] |
| ENSMUSG00000029649 | Pomp | 0.490608 | 1656.726 | 0.106885 | -4.59007 | 4.43E-06 | 5.1E-05 | 6.18E-05 | 1997.036 | 1874.65 | 1380.825 | 1374.393 | 66537 | proteasome maturation protein [Source:MGI Symbol;Acc:MGI:1913787] |
| ENSMUSG00000028729 | Ebna1bp2 | 0.490887 | 486.9733 | 0.179689 | -2.73187 | 0.006298 | 0.029042 | 0.035084 | 607.1496 | 531.1179 | 411.5042 | 398.1214 | 69072 | EBNA1 binding protein 2 [Source:MGI Symbol;Acc:MGI:1916322] |
| ENSMUSG00000041355 | Ssr2 | 0.491038 | 1234.228 | 0.123017 | -3.99163 | 6.56E-05 | 0.000576 | 0.000698 | 1514.701 | 1370.088 | 1030.589 | 1021.536 | 66256 | signal sequence receptor, beta [Source:MGI Symbol;Acc:MGI:1913506] |
| ENSMUSG00000031634 | Ufsp2 | 0.491398 | 902.7371 | 0.130426 | -3.76763 | 0.000165 | 0.001295 | 0.001569 | 1055.636 | 1054.367 | 748.9377 | 752.007 | 192169 | UFM1-specific peptidase 2 [Source:MGI Symbol;Acc:MGI:1913679] |
| ENSMUSG00000095115 | Itpripl2 | 0.491794 | 2587.32 | 0.098603 | -4.98762 | 6.11E-07 | 8.4E-06 | 1.02E-05 | 3159.505 | 2888.691 | 2107.816 | 2193.268 | 319622 | inositol 1,4,5-triphosphate receptor interacting protein-like 2 [Source:MGI Symbol;Acc:MGI:2442416] |
| ENSMUSG00000042029 | Ncapg2 | 0.491957 | 1983.284 | 0.119921 | -4.10234 | 4.09E-05 | 0.000383 | 0.000464 | 2503.699 | 2133.324 | 1695.397 | 1600.715 | 76044 | non-SMC condensin II complex, subunit G2 [Source:MGI Symbol;Acc:MGI:1923294] |
| ENSMUSG00000054091 | Arln | 0.492056 | 525.463 | 0.168187 | -2.92564 | 0.003437 | 0.017549 | 0.021245 | 613.4961 | 614.7198 | 417.9054 | 455.7306 | 67704 | allnegulin [Source:MGI Symbol;Acc:MGI:1914954] |
| ENSMUSG00000028034 | Fubp1 | 0.492082 | 1463.108 | 0.11584 | -4.24793 | 2.16E-05 | 0.000214 | 0.000259 | 1742.117 | 1678.923 | 1272.005 | 1159.387 | 51886 | far upstream element (FUSE) binding protein 1 [Source:MGI Symbol;Acc:MGI:1196294] |
| ENSMUSG00000028964 | Park7 | 0.492148 | 1201.533 | 0.118799 | -4.1427 | 3.43E-05 | 0.000326 | 0.000395 | 1427.965 | 1380.907 | 975.7222 | 1021.536 | 57320 | Parkinson disease (autosomal recessive, early onset) 7 [Source:MGI Symbol;Acc:MGI:2135637] |
| ENSMUSG00000029570 | Lfng | 0.492248 | 1387.836 | 0.123481 | -3.98641 | 6.71E-05 | 0.000586 | 0.00071 | 1728.367 | 1516.637 | 1161.356 | 1144.985 | 16848 | LFNG O-fucosylpeptide 3-beta-N-acetylglucosaminyltransferase [Source:MGI Symbol;Acc:MGI:1095413] |
| ENSMUSG00000026917 | Wdr5 | 0.492392 | 774.2218 | 0.141496 | -3.4799 | 0.000502 | 0.003478 | 0.004213 | 932.9372 | 877.3281 | 644.6899 | 641.9321 | 140858 | WD repeat domain 5 [Source:MGI Symbol;Acc:MGI:2155884] |
| ENSMUSG00000030603 | Psmc4 | 0.492511 | 1530 | 0.108991 | -4.51882 | 6.22E-06 | 6.97E-05 | 8.44E-05 | 1822.507 | 1754.656 | 1244.572 | 1298.267 | 23996 | proteasome (prosome, macropain) 26S subunit, ATPase, 4 [Source:MGI Symbol;Acc:MGI:1346093] |
| ENSMUSG00000060073 | Psma3 | 0.492861 | 2040.246 | 0.106152 | -4.64296 | 3.43E-06 | 4.04E-05 | 4.89E-05 | 2441.291 | 2329.05 | 1614.011 | 1776.629 | 19167 | proteasome subunit alpha 3 [Source:MGI Symbol;Acc:MGI:104883] |
| ENSMUSG00000019715 | Gle1 | 0.493378 | 402.7127 | 0.185278 | -2.6629 | 0.007747 | 0.03457 | 0.041766 | 470.6996 | 471.1213 | 334.6901 | 334.3396 | 74412 | GLE1 RNA export mediator [Source:MGI Symbol;Acc:MGI:1921662] |
| ENSMUSG00000046111 | Cep295 | 0.493666 | 630.3241 | 0.182997 | -2.69767 | 0.006983 | 0.031676 | 0.038268 | 829.2775 | 645.2099 | 522.1531 | 524.656 | 319675 | centrosomal protein 295 [Source:MGI Symbol;Acc:MGI:2442521] |
| ENSMUSG00000039234 | Sec24d | 0.493718 | 842.1306 | 0.135026 | -3.65646 | 0.000256 | 0.00192 | 0.002325 | 977.3628 | 992.4037 | 710.5306 | 688.2253 | 69608 | SEC24 homolog D, COPII coat complex component [Source:MGI Symbol;Acc:MGI:1916858] |
| ENSMUSG00000000581 | C1d | 0.493721 | 720.5977 | 0.150569 | -3.27904 | 0.001042 | 0.006508 | 0.007884 | 882.1651 | 803.5617 | 615.4274 | 581.2366 | 57316 | C1D nuclear receptor co-repressor [Source:MGI Symbol;Acc:MGI:1927354] |
| ENSMUSG00000049091 | Sephs2 | 0.494886 | 1978.786 | 0.121789 | -4.06347 | 4.83E-05 | 0.000441 | 0.000534 | 2436.003 | 2193.32 | 1522.566 | 1763.256 | 20768 | selenophosphate synthetase 2 [Source:MGI Symbol;Acc:MGI:108388] |
| ENSMUSG00000019795 | Pcmt1 | 0.494912 | 965.1369 | 0.132705 | -3.72943 | 0.000192 | 0.001482 | 0.001795 | 1114.871 | 1142.887 | 765.3978 | 837.3922 | 18537 | protein-L-isoaspartate (D-aspartate) O-methyltransferase 1 [Source:MGI Symbol;Acc:MGI:97502] |
| ENSMUSG00000038023 | Atp6v0a2 | 0.495283 | 853.2862 | 0.157706 | -3.14055 | 0.001686 | 0.009698 | 0.011748 | 918.1287 | 1077.973 | 662.979 | 754.0645 | 21871 | ATPase, H+ transporting, lysosomal V0 subunit A2 [Source:MGI Symbol;Acc:MGI:104855] |
| ENSMUSG00000031479 | Vps36 | 0.495382 | 685.6034 | 0.147889 | -3.34967 | 0.000809 | 0.005238 | 0.006346 | 803.8915 | 800.6111 | 583.4215 | 554.4894 | 70160 | vacuolar protein sorting 36 [Source:MGI Symbol;Acc:MGI:1917410] |
| ENSMUSG00000038544 | Inip | 0.495581 | 531.2217 | 0.172832 | -2.86742 | 0.004138 | 0.02054 | 0.024833 | 657.9217 | 585.2133 | 427.0499 | 454.7019 | 66209 | INTS3 and NABP interacting protein [Source:MGI Symbol;Acc:MGI:1913459] |
| ENSMUSG00000032330 | Cox7a2 | 0.495607 | 1302.838 | 0.11778 | -4.20792 | 2.58E-05 | 0.000251 | 0.000304 | 1573.935 | 1475.328 | 1105.575 | 1056.513 | 12866 | cytochrome c oxidase subunit 7A2 [Source:MGI Symbol;Acc:MGI:1316715] |
| ENSMUSG00000029816 | Gpnmb | 0.495856 | 20601.07 | 0.087655 | -5.65691 | 1.54E-08 | 2.9E-07 | 3.52E-07 | 22593.58 | 25620.54 | 17837.34 | 16352.81 | 93695 | glycoprotein (transmembrane) nmb [Source:MGI Symbol;Acc:MGI:1934765] |
| ENSMUSG00000052397 | Ezr | 0.496512 | 570.45 | 0.162866 | -3.04859 | 0.002299 | 0.0125 | 0.015142 | 638.8822 | 696.3546 | 476.4304 | 470.133 | 22350 | ezrin [Source:MGI Symbol;Acc:MGI:98931] |
| ENSMUSG00000024346 | Pfdn1 | 0.49664 | 522.1541 | 0.166779 | -2.97784 | 0.002903 | 0.015167 | 0.018367 | 627.2469 | 595.0488 | 427.0499 | 439.2708 | 67199 | prefoldin 1 [Source:MGI Symbol;Acc:MGI:1914449] |
| ENSMUSG00000027828 | Ssr3 | 0.496927 | 4555.914 | 0.080746 | -6.1542 | 7.55E-10 | 1.72E-08 | 2.09E-08 | 5464.346 | 5202.005 | 3842.535 | 3714.771 | 67437 | signal sequence receptor, gamma [Source:MGI Symbol;Acc:MGI:1914687] |
| ENSMUSG00000026643 | Nmt2 | 0.49712 | 707.6031 | 0.154037 | -3.22728 | 0.00125 | 0.007568 | 0.009167 | 873.7031 | 782.9071 | 567.8758 | 605.9263 | 18108 | N-myristoyltransferase 2 [Source:MGI Symbol;Acc:MGI:1202298] |
| ENSMUSG00000026234 | Ncl | 0.497838 | 1255.142 | 0.15603 | -3.19064 | 0.00142 | 0.008415 | 0.010193 | 1653.266 | 1286.486 | 1089.115 | 991.7028 | 17975 | nucleolin [Source:MGI Symbol;Acc:MGI:97286] |
| ENSMUSG00000025371 | Chmp6 | 0.49807 | 1140.528 | 0.121879 | -4.08659 | 4.38E-05 | 0.000406 | 0.000492 | 1359.211 | 1312.058 | 971.15 | 919.6912 | 208092 | charged multivesicular body protein 6 [Source:MGI Symbol;Acc:MGI:3583942] |
| ENSMUSG00000049130 | C5ar1 | 0.498137 | 2522.434 | 0.115152 | -4.32592 | 1.52E-05 | 0.000157 | 0.00019 | 3121.426 | 2785.418 | 1950.53 | 2232.36 | 12273 | complement component 5a receptor 1 [Source:MGI Symbol;Acc:MGI:88232] |
| ENSMUSG00000068921 | Dap3 | 0.498596 | 449.4133 | 0.179089 | -2.78407 | 0.005368 | 0.025482 | 0.030782 | 534.1647 | 518.3317 | 359.3804 | 385.7765 | 65111 | death associated protein 3 [Source:MGI Symbol;Acc:MGI:1929538] |
| ENSMUSG00000021149 | Gtpbp4 | 0.498627 | 831.1743 | 0.172545 | -2.88983 | 0.003855 | 0.019391 | 0.023451 | 1101.12 | 845.8545 | 673.038 | 704.6851 | 69237 | GTP binding protein 4 [Source:MGI Symbol;Acc:MGI:1916487] |
| ENSMUSG00000026655 | Fam107b | 0.499068 | 1160.142 | 0.133616 | -3.73508 | 0.000188 | 0.001452 | 0.001759 | 1384.597 | 1333.696 | 1032.418 | 889.8578 | 66540 | family with sequence similarity 107, member B [Source:MGI Symbol;Acc:MGI:1913790] |
| ENSMUSG00000021952 | Xpo4 | 0.499327 | 697.3165 | 0.164934 | -3.02743 | 0.002466 | 0.013281 | 0.016082 | 881.1073 | 752.417 | 547.7578 | 607.9838 | 57258 | exportin 4 [Source:MGI Symbol;Acc:MGI:1888526] |
| ENSMUSG00000025132 | Arhgdia | 0.499524 | 3255.632 | 0.093846 | -5.32284 | 1.02E-07 | 1.67E-06 | 2.02E-06 | 3936.953 | 3691.27 | 2801.887 | 2592.418 | 192662 | Rho GDP dissociation inhibitor alpha [Source:MGI Symbol;Acc:MGI:2178103] |
| ENSMUSG00000109865 | Hspa14 | 0.499532 | 675.2689 | 0.155158 | -3.21951 | 0.001284 | 0.007736 | 0.009371 | 835.624 | 746.5157 | 554.159 | 564.7768 | 50497 | heat shock protein 14 [Source:MGI Symbol;Acc:MGI:1354164] |
| ENSMUSG00000052337 | Immt | 0.499865 | 2160.734 | 0.099578 | -5.01985 | 5.17E-07 | 7.25E-06 | 8.78E-06 | 2618.994 | 2444.126 | 1812.447 | 1767.371 | 76614 | inner membrane protein, mitochondrial [Source:MGI Symbol;Acc:MGI:1923864] |
| ENSMUSG00000042688 | Mapk6 | 0.499878 | 1289.898 | 0.141479 | -3.53323 | 0.000411 | 0.002918 | 0.003535 | 1589.801 | 1432.051 | 971.15 | 1166.588 | 50772 | mitogen-activated protein kinase 6 [Source:MGI Symbol;Acc:MGI:1354946] |
| ENSMUSG00000030970 | Ctbp2 | 0.499896 | 894.6368 | 0.131057 | -3.81434 | 0.000137 | 0.001095 | 0.001327 | 1055.636 | 1040.598 | 741.622 | 740.6909 | 13017 | C-terminal binding protein 2 [Source:MGI Symbol;Acc:MGI:1201686] |
| ENSMUSG00000021504 | B4galt7 | 0.500237 | 613.8395 | 0.161404 | -3.09928 | 0.00194 | 0.01089 | 0.013192 | 731.9643 | 706.1901 | 479.1738 | 538.0296 | 218271 | beta-1,4-galactosyltransferase 7 [Source:MGI Symbol;Acc:MGI:2384987] |
| ENSMUSG00000029521 | Chek2 | 0.500792 | 693.4027 | 0.160039 | -3.12918 | 0.001753 | 0.010016 | 0.012133 | 865.2411 | 760.2855 | 600.7962 | 547.2883 | 50883 | checkpoint kinase 2 [Source:MGI Symbol;Acc:MGI:1355321] |
| ENSMUSG00000028416 | Bag1 | 0.501002 | 1929.761 | 0.104558 | -4.79161 | 1.65E-06 | 2.08E-05 | 2.52E-05 | 2351.383 | 2171.682 | 1574.689 | 1621.29 | 12017 | BCL2-associated athanogene 1 [Source:MGI Symbol;Acc:MGI:108047] |
| ENSMUSG00000044573 | Acp1 | 0.501347 | 956.6791 | 0.132023 | -3.79741 | 0.000146 | 0.001165 | 0.001412 | 1164.585 | 1077.973 | 782.7725 | 801.3864 | 11431 | acid phosphatase 1, soluble [Source:MGI Symbol;Acc:MGI:87881] |
| ENSMUSG00000025499 | Hras | 0.501731 | 606.5478 | 0.161719 | -3.10248 | 0.001919 | 0.010793 | 0.013075 | 721.3868 | 700.2888 | 473.6871 | 530.8285 | 15461 | Harvey rat sarcoma virus oncogene [Source:MGI Symbol;Acc:MGI:96224] |
| ENSMUSG00000056999 | Ide | 0.501996 | 1403.691 | 0.127315 | -3.94296 | 8.05E-05 | 0.000689 | 0.000834 | 1750.579 | 1540.242 | 1111.061 | 1212.881 | 15925 | insulin degrading enzyme [Source:MGI Symbol;Acc:MGI:96412] |
| ENSMUSG00000013822 | Elof1 | 0.502966 | 751.7548 | 0.15707 | -3.20218 | 0.001364 | 0.008155 | 0.009879 | 868.4143 | 894.0485 | 569.7047 | 674.8517 | 66126 | ELF1 homolog, elongation factor 1 [Source:MGI Symbol;Acc:MGI:1913376] |
| ENSMUSG00000027957 | Slc35a3 | 0.50302 | 1595.484 | 0.130274 | -3.86124 | 0.000113 | 0.000924 | 0.001119 | 2021.364 | 1720.232 | 1254.631 | 1385.709 | 229782 | solute carrier family 35 (UDP-N-acetylglucosamine (UDP-GlcNAc) transporter), member 3 [Source:MGI Symbol;Acc:MGI:1917648] |
| ENSMUSG00000030786 | Itgam | 0.503527 | 9828.971 | 0.069296 | -7.26629 | 3.69E-13 | 1.3E-11 | 1.57E-11 | 11670.18 | 11384.61 | 8298.668 | 7962.427 | 16409 | integrin alpha M [Source:MGI Symbol;Acc:MGI:96607] |
| ENSMUSG00000022285 | Ywhaz | 0.503759 | 4687.234 | 0.096922 | -5.19755 | 2.02E-07 | 3.08E-06 | 3.74E-06 | 5809.174 | 5185.285 | 3705.367 | 4049.11 | 22631 | tyrosine 3-monooxygenase/tryptophan 5-monooxygenase activation protein, zeta polypeptide [Source:MGI Symbol;Acc:MGI:109484] |
| ENSMUSG00000022682 | Rrn3 | 0.503981 | 994.8008 | 0.149498 | -3.37116 | 0.000749 | 0.0049 | 0.005935 | 1238.627 | 1094.693 | 758.0822 | 887.8003 | 106298 | RRN3 RNA polymerase I transcription factor homolog (yeast) [Source:MGI Symbol;Acc:MGI:1925255] |
| ENSMUSG00000022674 | Ube2v2 | 0.504116 | 526.4361 | 0.187437 | -2.68953 | 0.007155 | 0.032329 | 0.039058 | 683.3077 | 551.7725 | 425.221 | 445.4433 | 70620 | ubiquitin-conjugating enzyme E2 variant 2 [Source:MGI Symbol;Acc:MGI:1917870] |
| ENSMUSG00000028894 | Inpp5b | 0.504122 | 548.044 | 0.166664 | -3.02478 | 0.002488 | 0.013358 | 0.016175 | 629.3624 | 656.029 | 432.5367 | 474.2479 | 16330 | inositol polyphosphate-5-phosphatase B [Source:MGI Symbol;Acc:MGI:103257] |
| ENSMUSG00000075701 | Selenos | 0.504135 | 601.0462 | 0.16599 | -3.03714 | 0.002388 | 0.01292 | 0.015651 | 689.6542 | 719.9598 | 462.7136 | 531.8572 | 109815 | selenoprotein S [Source:MGI Symbol;Acc:MGI:95994] |
| ENSMUSG00000007041 | Clic1 | 0.504553 | 3954.064 | 0.081881 | -6.16201 | 7.18E-10 | 1.65E-08 | 2E-08 | 4622.376 | 4655.15 | 3349.644 | 3189.086 | 114584 | chloride intracellular channel 1 [Source:MGI Symbol;Acc:MGI:2148924] |
| ENSMUSG00000019179 | Mdh2 | 0.505198 | 795.2586 | 0.163002 | -3.09933 | 0.00194 | 0.01089 | 0.013192 | 1013.326 | 852.7393 | 615.4274 | 699.5414 | 17448 | malate dehydrogenase 2, NAD (mitochondrial) [Source:MGI Symbol;Acc:MGI:97050] |
| ENSMUSG00000017548 | Suz12 | 0.505243 | 1412.264 | 0.160769 | -3.14266 | 0.001674 | 0.009652 | 0.011693 | 1903.953 | 1410.413 | 1140.324 | 1194.364 | 52615 | SUZ12 polycomb repressive complex 2 subunit [Source:MGI Symbol;Acc:MGI:1261758] |
| ENSMUSG00000024098 | Twsg1 | 0.505759 | 582.1468 | 0.168585 | -3.00002 | 0.0027 | 0.014278 | 0.01729 | 699.174 | 666.848 | 447.1679 | 515.3974 | 65960 | twisted gastrulation BMP signaling modulator 1 [Source:MGI Symbol;Acc:MGI:2137520] |
| ENSMUSG00000027282 | Mtch2 | 0.506138 | 1793.997 | 0.105296 | -4.80681 | 1.53E-06 | 1.95E-05 | 2.36E-05 | 2181.084 | 2030.051 | 1473.185 | 1491.669 | 56428 | mitochondrial carrier 2 [Source:MGI Symbol;Acc:MGI:1929260] |
| ENSMUSG00000021537 | Cetn3 | 0.506388 | 579.4476 | 0.158344 | -3.19803 | 0.001384 | 0.008231 | 0.009971 | 685.4232 | 674.7165 | 470.0293 | 487.6215 | 12626 | centrin 3 [Source:MGI Symbol;Acc:MGI:1097706] |
| ENSMUSG00000008393 | Carhsp1 | 0.507022 | 949.8536 | 0.132824 | -3.81726 | 0.000135 | 0.001085 | 0.001315 | 1108.524 | 1121.249 | 750.7666 | 818.8749 | 52502 | calcium regulated heat stable protein 1 [Source:MGI Symbol;Acc:MGI:1196368] |
| ENSMUSG00000022533 | Atp13a3 | 0.508318 | 3184.54 | 0.106395 | -4.77764 | 1.77E-06 | 2.21E-05 | 2.68E-05 | 3975.032 | 3504.395 | 2503.775 | 2754.959 | 224088 | ATPase type 13A3 [Source:MGI Symbol;Acc:MGI:2685387] |
| ENSMUSG00000026377 | Nifk | 0.508355 | 576.0895 | 0.167273 | -3.03907 | 0.002373 | 0.012847 | 0.015562 | 709.7515 | 643.2428 | 454.4835 | 496.8801 | 67949 | nucleolar protein interacting with the FHA domain of MKI67 [Source:MGI Symbol;Acc:MGI:1915199] |
| ENSMUSG00000063273 | Naa15 | 0.508359 | 1798.904 | 0.14383 | -3.53445 | 0.000409 | 0.002908 | 0.003522 | 2376.769 | 1849.077 | 1540.855 | 1428.916 | 74838 | N(alpha)-acetyltransferase 15, NatA auxiliary subunit [Source:MGI Symbol;Acc:MGI:1922088] |
| ENSMUSG00000036167 | Pphln1 | 0.509011 | 587.4211 | 0.171291 | -2.97162 | 0.002962 | 0.015462 | 0.018724 | 744.6573 | 635.3744 | 472.7726 | 496.8801 | 223828 | periphilin 1 [Source:MGI Symbol;Acc:MGI:1917029] |
| ENSMUSG00000015671 | Psma2 | 0.50969 | 2002.944 | 0.107132 | -4.75761 | 1.96E-06 | 2.43E-05 | 2.94E-05 | 2419.079 | 2286.758 | 1575.604 | 1730.336 | 19166 | proteasome subunit alpha 2 [Source:MGI Symbol;Acc:MGI:104885] |
| ENSMUSG00000026121 | Sema4c | 0.510143 | 909.0148 | 0.141161 | -3.6139 | 0.000302 | 0.002216 | 0.002684 | 1137.083 | 999.2885 | 750.7666 | 748.9208 | 20353 | sema domain, immunoglobulin domain (Ig), transmembrane domain (TM) and short cytoplasmic domain, (semaphorin) 4C [Source:MGI Symbol;Acc:MGI:109252] |
| ENSMUSG00000027087 | Itgav | 0.510555 | 6017.528 | 0.078697 | -6.48764 | 8.72E-11 | 2.29E-09 | 2.77E-09 | 7324.932 | 6817.98 | 4939.88 | 4987.319 | 16410 | integrin alpha V [Source:MGI Symbol;Acc:MGI:96608] |
| ENSMUSG00000029169 | Dhx15 | 0.511838 | 3361.173 | 0.119235 | -4.2927 | 1.77E-05 | 0.000179 | 0.000216 | 4375.92 | 3527.016 | 2766.223 | 2775.533 | 13204 | DEAH-box helicase 15 [Source:MGI Symbol;Acc:MGI:1099786] |
| ENSMUSG00000041747 | Utp15 | 0.512363 | 773.2269 | 0.143641 | -3.56697 | 0.000361 | 0.002604 | 0.003154 | 931.8794 | 886.1801 | 615.4274 | 659.4206 | 105372 | UTP15 small subunit processome component [Source:MGI Symbol;Acc:MGI:2145443] |
| ENSMUSG00000021978 | Extl3 | 0.513389 | 2016.046 | 0.11504 | -4.46271 | 8.09E-06 | 8.91E-05 | 0.000108 | 2547.067 | 2195.287 | 1654.247 | 1667.583 | 54616 | exostosin-like glycosyltransferase 3 [Source:MGI Symbol;Acc:MGI:1860765] |
| ENSMUSG00000000290 | Itgb2 | 0.513425 | 3918.81 | 0.101907 | -5.03817 | 4.7E-07 | 6.64E-06 | 8.05E-06 | 4956.625 | 4261.729 | 3271.916 | 3184.971 | 16414 | integrin beta 2 [Source:MGI Symbol;Acc:MGI:96611] |
| ENSMUSG00000044147 | Arf6 | 0.513799 | 1897.66 | 0.109532 | -4.69086 | 2.72E-06 | 3.27E-05 | 3.96E-05 | 2354.556 | 2109.718 | 1547.256 | 1579.112 | 11845 | ARF GTPase 6 [Source:MGI Symbol;Acc:MGI:99435] |
| ENSMUSG00000028656 | Cap1 | 0.514364 | 10903.82 | 0.069399 | -7.41169 | 1.25E-13 | 4.61E-12 | 5.59E-12 | 13166.9 | 12488.16 | 9033.889 | 8926.354 | 12331 | cyclase associated actin cytoskeleton regulatory protein 1 [Source:MGI Symbol;Acc:MGI:88262] |
| ENSMUSG00000006304 | Arpc2 | 0.514569 | 5751.418 | 0.096172 | -5.3505 | 8.77E-08 | 1.45E-06 | 1.76E-06 | 7273.102 | 6260.307 | 4785.337 | 4686.927 | 76709 | actin related protein 2/3 complex, subunit 2 [Source:MGI Symbol;Acc:MGI:1923959] |
| ENSMUSG00000026321 | Tnfrsf11a | 0.514768 | 2139.352 | 0.131478 | -3.91525 | 9.03E-05 | 0.000761 | 0.000922 | 2789.292 | 2245.449 | 1825.25 | 1697.417 | 21934 | tumor necrosis factor receptor superfamily, member 11a, NFKB activator [Source:MGI Symbol;Acc:MGI:1314891] |
| ENSMUSG00000018965 | Ywhah | 0.515627 | 2089.349 | 0.113657 | -4.53668 | 5.71E-06 | 6.47E-05 | 7.84E-05 | 2636.976 | 2280.856 | 1699.97 | 1739.595 | 22629 | tyrosine 3-monooxygenase/tryptophan 5-monooxygenase activation protein, eta polypeptide [Source:MGI Symbol;Acc:MGI:109194] |
| ENSMUSG00000054814 | Usp46 | 0.515699 | 462.1303 | 0.191938 | -2.6868 | 0.007214 | 0.032536 | 0.039307 | 570.1283 | 517.3482 | 347.4925 | 413.5524 | 69727 | ubiquitin specific peptidase 46 [Source:MGI Symbol;Acc:MGI:1916977] |
| ENSMUSG00000019297 | Nop9 | 0.516207 | 503.6066 | 0.176933 | -2.91752 | 0.003528 | 0.017947 | 0.021718 | 603.9763 | 581.2791 | 386.814 | 442.3571 | 67842 | NOP9 nucleolar protein [Source:MGI Symbol;Acc:MGI:1915092] |
| ENSMUSG00000016487 | Ppfibp1 | 0.51632 | 1002.365 | 0.132035 | -3.91049 | 9.21E-05 | 0.000773 | 0.000937 | 1237.57 | 1122.232 | 818.4362 | 831.2198 | 67533 | PTPRF interacting protein, binding protein 1 (liprin beta 1) [Source:MGI Symbol;Acc:MGI:1914783] |
| ENSMUSG00000025487 | Psmd13 | 0.516453 | 1129.244 | 0.127423 | -4.05306 | 5.06E-05 | 0.000458 | 0.000555 | 1385.655 | 1272.716 | 899.8226 | 958.7832 | 23997 | proteasome (prosome, macropain) 26S subunit, non-ATPase, 13 [Source:MGI Symbol;Acc:MGI:1345192] |
| ENSMUSG00000024943 | Smc5 | 0.516685 | 940.3705 | 0.140794 | -3.66979 | 0.000243 | 0.001832 | 0.002219 | 1178.336 | 1035.68 | 755.3389 | 792.1278 | 226026 | structural maintenance of chromosomes 5 [Source:MGI Symbol;Acc:MGI:2385088] |
| ENSMUSG00000023110 | Prmt5 | 0.516927 | 582.1496 | 0.167207 | -3.09154 | 0.001991 | 0.011098 | 0.013444 | 722.4446 | 648.1606 | 459.0558 | 498.9376 | 27374 | protein arginine N-methyltransferase 5 [Source:MGI Symbol;Acc:MGI:1351645] |
| ENSMUSG00000026688 | Mgst3 | 0.517748 | 385.0623 | 0.196107 | -2.64014 | 0.008287 | 0.036529 | 0.044122 | 460.1221 | 446.5325 | 297.1975 | 336.3971 | 66447 | microsomal glutathione S-transferase 3 [Source:MGI Symbol;Acc:MGI:1913697] |
| ENSMUSG00000041025 | Iffo2 | 0.517802 | 504.3612 | 0.170097 | -3.04415 | 0.002333 | 0.012659 | 0.015334 | 612.4384 | 575.3777 | 407.8464 | 421.7823 | 212632 | intermediate filament family orphan 2 [Source:MGI Symbol;Acc:MGI:2140675] |
| ENSMUSG00000045414 | Dipk2a | 0.517812 | 1161.297 | 0.124428 | -4.16152 | 3.16E-05 | 0.000303 | 0.000368 | 1426.907 | 1308.124 | 940.0585 | 970.0993 | 68861 | divergent protein kinase domain 2A [Source:MGI Symbol;Acc:MGI:1916111] |
| ENSMUSG00000018293 | Pfn1 | 0.517964 | 4387.255 | 0.106973 | -4.84202 | 1.29E-06 | 1.66E-05 | 2.01E-05 | 5522.523 | 4811.535 | 3824.246 | 3390.718 | 18643 | profilin 1 [Source:MGI Symbol;Acc:MGI:97549] |
| ENSMUSG00000026627 | Pacc1 | 0.518163 | 1128.592 | 0.127949 | -4.04977 | 5.13E-05 | 0.000464 | 0.000562 | 1289.4 | 1369.104 | 971.15 | 884.7141 | 66950 | proton activated chloride channel 1 [Source:MGI Symbol;Acc:MGI:1914200] |
| ENSMUSG00000021719 | Rgs7bp | 0.518257 | 424.9053 | 0.188993 | -2.7422 | 0.006103 | 0.028331 | 0.034225 | 525.7027 | 475.0555 | 334.6901 | 364.173 | 52882 | regulator of G-protein signalling 7 binding protein [Source:MGI Symbol;Acc:MGI:106334] |
| ENSMUSG00000041974 | Spidr | 0.518789 | 669.6229 | 0.148234 | -3.49981 | 0.000466 | 0.003255 | 0.003943 | 797.5449 | 779.9565 | 549.5867 | 551.4032 | 224008 | scaffolding protein involved in DNA repair [Source:MGI Symbol;Acc:MGI:1924834] |
| ENSMUSG00000022439 | Parvg | 0.518921 | 769.8334 | 0.161029 | -3.22254 | 0.001271 | 0.007667 | 0.009287 | 932.9372 | 881.2623 | 693.156 | 571.978 | 64099 | parvin, gamma [Source:MGI Symbol;Acc:MGI:2158329] |
| ENSMUSG00000030880 | Polr3e | 0.519249 | 475.1658 | 0.187391 | -2.77093 | 0.00559 | 0.026346 | 0.031826 | 609.2651 | 510.4633 | 394.1296 | 386.8052 | 26939 | polymerase (RNA) III (DNA directed) polypeptide E [Source:MGI Symbol;Acc:MGI:1349452] |
| ENSMUSG00000042111 | Ccdc115 | 0.519723 | 804.7512 | 0.143645 | -3.61812 | 0.000297 | 0.002185 | 0.002647 | 979.4783 | 916.6702 | 634.631 | 688.2253 | 69668 | coiled-coil domain containing 115 [Source:MGI Symbol;Acc:MGI:1916918] |
| ENSMUSG00000028394 | Pole3 | 0.520381 | 403.9545 | 0.186919 | -2.784 | 0.005369 | 0.025482 | 0.030782 | 462.2376 | 489.8087 | 335.6046 | 328.1672 | 59001 | polymerase (DNA directed), epsilon 3 (p17 subunit) [Source:MGI Symbol;Acc:MGI:1933378] |
| ENSMUSG00000037070 | Rbmxl1 | 0.520391 | 751.6876 | 0.157186 | -3.31066 | 0.000931 | 0.005901 | 0.007148 | 959.381 | 812.4137 | 613.5985 | 621.3574 | 19656 | RNA binding motif protein, X-linked like-1 [Source:MGI Symbol;Acc:MGI:1343045] |
| ENSMUSG00000030835 | Nomo1 | 0.52151 | 740.6465 | 0.167027 | -3.12231 | 0.001794 | 0.010211 | 0.01237 | 958.3232 | 787.8249 | 580.6782 | 635.7597 | 211548 | nodal modulator 1 [Source:MGI Symbol;Acc:MGI:2385850] |
| ENSMUSG00000036371 | Serbp1 | 0.52211 | 4024.555 | 0.099015 | -5.27302 | 1.34E-07 | 2.13E-06 | 2.58E-06 | 5072.978 | 4417.131 | 3247.226 | 3360.885 | 66870 | serpine1 mRNA binding protein 1 [Source:MGI Symbol;Acc:MGI:1914120] |
| ENSMUSG00000002658 | Gtf2f1 | 0.522164 | 615.0413 | 0.158135 | -3.30202 | 0.00096 | 0.006057 | 0.007337 | 692.8275 | 757.3348 | 503.8641 | 506.1388 | 98053 | general transcription factor IIF, polypeptide 1 [Source:MGI Symbol;Acc:MGI:1923848] |
| ENSMUSG00000034111 | Tmed8 | 0.522481 | 1030.052 | 0.156229 | -3.34433 | 0.000825 | 0.00532 | 0.006444 | 1349.691 | 1079.94 | 877.8757 | 812.7025 | 382620 | transmembrane p24 trafficking protein 8 [Source:MGI Symbol;Acc:MGI:1923480] |
| ENSMUSG00000052214 | Opa3 | 0.522899 | 462.791 | 0.189596 | -2.75796 | 0.005816 | 0.027195 | 0.032852 | 568.0128 | 523.2495 | 348.4069 | 411.4949 | 403187 | optic atrophy 3 [Source:MGI Symbol;Acc:MGI:2686271] |
| ENSMUSG00000017715 | Pgs1 | 0.523289 | 1026.295 | 0.145317 | -3.60103 | 0.000317 | 0.002317 | 0.002807 | 1244.974 | 1175.344 | 767.2268 | 917.6337 | 74451 | phosphatidylglycerophosphate synthase 1 [Source:MGI Symbol;Acc:MGI:1921701] |
| ENSMUSG00000022272 | Myo10 | 0.523423 | 525.2407 | 0.16935 | -3.09078 | 0.001996 | 0.011108 | 0.013456 | 595.5143 | 643.2428 | 418.8198 | 443.3858 | 17909 | myosin X [Source:MGI Symbol;Acc:MGI:107716] |
| ENSMUSG00000024696 | Lpxn | 0.523755 | 2125.621 | 0.100511 | -5.21091 | 1.88E-07 | 2.89E-06 | 3.5E-06 | 2603.127 | 2411.669 | 1742.949 | 1744.739 | 107321 | leupaxin [Source:MGI Symbol;Acc:MGI:2147677] |
| ENSMUSG00000030793 | Pycard | 0.52445 | 1608.328 | 0.115891 | -4.52537 | 6.03E-06 | 6.8E-05 | 8.23E-05 | 1977.996 | 1817.604 | 1374.424 | 1263.289 | 66824 | PYD and CARD domain containing [Source:MGI Symbol;Acc:MGI:1931465] |
| ENSMUSG00000028902 | Sf3a3 | 0.525369 | 737.3466 | 0.149734 | -3.50867 | 0.00045 | 0.003161 | 0.00383 | 918.1287 | 822.2492 | 598.9673 | 610.0412 | 75062 | splicing factor 3a, subunit 3 [Source:MGI Symbol;Acc:MGI:1922312] |
| ENSMUSG00000030423 | Pop4 | 0.525636 | 374.5737 | 0.199438 | -2.63559 | 0.008399 | 0.036926 | 0.044602 | 444.2558 | 439.6476 | 286.224 | 328.1672 | 66161 | processing of precursor 4, ribonuclease P/MRP family, (S. cerevisiae) [Source:MGI Symbol;Acc:MGI:1913411] |
| ENSMUSG00000078713 | Tomm5 | 0.525651 | 479.0735 | 0.172276 | -3.05121 | 0.002279 | 0.012422 | 0.015048 | 572.2438 | 558.6574 | 399.6163 | 385.7765 | 68512 | translocase of outer mitochondrial membrane 5 [Source:MGI Symbol;Acc:MGI:1915762] |
| ENSMUSG00000021282 | Eif5 | 0.5257 | 3496.749 | 0.099474 | -5.28479 | 1.26E-07 | 2.01E-06 | 2.43E-06 | 4309.281 | 3944.042 | 2720.5 | 3013.172 | 217869 | eukaryotic translation initiation factor 5 [Source:MGI Symbol;Acc:MGI:95309] |
| ENSMUSG00000038991 | Txndc5 | 0.526151 | 1994.226 | 0.101482 | -5.18467 | 2.16E-07 | 3.29E-06 | 3.98E-06 | 2416.963 | 2290.692 | 1597.551 | 1671.698 | 105245 | thioredoxin domain containing 5 [Source:MGI Symbol;Acc:MGI:2145316] |
| ENSMUSG00000120144 | Gm56517 | 0.526277 | 1017.636 | 0.132318 | -3.97735 | 6.97E-05 | 0.000606 | 0.000734 | 1255.552 | 1146.821 | 810.2061 | 857.967 | | predicted gene, 56517 [Source:MGI Symbol;Acc:MGI:7336948] |
| ENSMUSG00000073838 | Tufm | 0.526449 | 707.4109 | 0.15975 | -3.29545 | 0.000983 | 0.006185 | 0.007492 | 834.5663 | 836.0189 | 628.2298 | 530.8285 | 233870 | Tu translation elongation factor, mitochondrial [Source:MGI Symbol;Acc:MGI:1923686] |
| ENSMUSG00000041736 | Tspo | 0.527541 | 1386.566 | 0.112692 | -4.68128 | 2.85E-06 | 3.41E-05 | 4.13E-05 | 1663.844 | 1611.058 | 1161.356 | 1110.008 | 12257 | translocator protein [Source:MGI Symbol;Acc:MGI:88222] |
| ENSMUSG00000069565 | Dazap1 | 0.527594 | 610.7714 | 0.163089 | -3.23501 | 0.001216 | 0.007405 | 0.00897 | 766.8701 | 675.7 | 497.4629 | 503.0526 | 70248 | DAZ associated protein 1 [Source:MGI Symbol;Acc:MGI:1917498] |
| ENSMUSG00000029554 | Mad1l1 | 0.527749 | 467.2891 | 0.180691 | -2.92073 | 0.003492 | 0.01779 | 0.021528 | 579.648 | 524.2331 | 395.9585 | 369.3167 | 17120 | MAD1 mitotic arrest deficient 1-like 1 [Source:MGI Symbol;Acc:MGI:1341857] |
| ENSMUSG00000032171 | Pin1 | 0.527838 | 512.5503 | 0.181068 | -2.91514 | 0.003555 | 0.018036 | 0.021826 | 639.9399 | 570.46 | 392.3007 | 447.5007 | 23988 | peptidyl-prolyl cis/trans isomerase, NIMA-interacting 1 [Source:MGI Symbol;Acc:MGI:1346036] |
| ENSMUSG00000059456 | Ptk2b | 0.529205 | 1172.362 | 0.153894 | -3.43877 | 0.000584 | 0.003964 | 0.004802 | 1531.625 | 1238.292 | 897.9937 | 1021.536 | 19229 | PTK2 protein tyrosine kinase 2 beta [Source:MGI Symbol;Acc:MGI:104908] |
| ENSMUSG00000027714 | Exosc9 | 0.52956 | 563.4997 | 0.178669 | -2.96391 | 0.003038 | 0.015795 | 0.019127 | 729.8488 | 601.9336 | 465.457 | 456.7594 | 50911 | exosome component 9 [Source:MGI Symbol;Acc:MGI:1355319] |
| ENSMUSG00000022956 | Atp5po | 0.529738 | 1651.04 | 0.104139 | -5.08685 | 3.64E-07 | 5.25E-06 | 6.36E-06 | 1976.938 | 1924.811 | 1358.878 | 1343.531 | 28080 | ATP synthase peripheral stalk subunit OSCP [Source:MGI Symbol;Acc:MGI:106341] |
| ENSMUSG00000054051 | Ercc6 | 0.530821 | 442.4355 | 0.20161 | -2.63291 | 0.008466 | 0.037185 | 0.044915 | 582.8213 | 463.2528 | 366.696 | 356.9719 | 319955 | excision repair cross-complementing rodent repair deficiency, complementation group 6 [Source:MGI Symbol;Acc:MGI:1100494] |
| ENSMUSG00000028330 | Ncbp1 | 0.531344 | 746.9502 | 0.146739 | -3.62101 | 0.000293 | 0.002164 | 0.002622 | 917.0709 | 848.8051 | 594.395 | 627.5298 | 433702 | nuclear cap binding protein subunit 1 [Source:MGI Symbol;Acc:MGI:1891840] |
| ENSMUSG00000010517 | Faf1 | 0.53135 | 961.936 | 0.135725 | -3.9149 | 9.04E-05 | 0.000761 | 0.000922 | 1195.26 | 1078.956 | 769.0557 | 804.4726 | 14084 | Fas-associated factor 1 [Source:MGI Symbol;Acc:MGI:109419] |
| ENSMUSG00000030759 | Far1 | 0.531424 | 2724.839 | 0.094569 | -5.61942 | 1.92E-08 | 3.57E-07 | 4.33E-07 | 3335.092 | 3107.04 | 2183.716 | 2273.51 | 67420 | fatty acyl CoA reductase 1 [Source:MGI Symbol;Acc:MGI:1914670] |
| ENSMUSG00000027236 | Eif3j1 | 0.531721 | 708.6943 | 0.167731 | -3.17008 | 0.001524 | 0.008933 | 0.010822 | 923.4174 | 752.417 | 571.5336 | 587.409 | 78655 | eukaryotic translation initiation factor 3, subunit J1 [Source:MGI Symbol;Acc:MGI:1925905] |
| ENSMUSG00000025190 | Got1 | 0.532263 | 928.7657 | 0.149054 | -3.57093 | 0.000356 | 0.002571 | 0.003114 | 1195.26 | 1001.256 | 748.0232 | 770.5243 | 14718 | glutamic-oxaloacetic transaminase 1, soluble [Source:MGI Symbol;Acc:MGI:95791] |
| ENSMUSG00000021474 | Sfxn1 | 0.532521 | 678.4292 | 0.150654 | -3.53473 | 0.000408 | 0.002906 | 0.00352 | 807.0647 | 797.6604 | 576.1059 | 532.8859 | 14057 | sideroflexin 1 [Source:MGI Symbol;Acc:MGI:2137677] |
| ENSMUSG00000037149 | Ddx1 | 0.533067 | 1572.523 | 0.112278 | -4.74774 | 2.06E-06 | 2.54E-05 | 3.07E-05 | 1930.397 | 1789.081 | 1252.802 | 1317.813 | 104721 | DEAD box helicase 1 [Source:MGI Symbol;Acc:MGI:2144727] |
| ENSMUSG00000029034 | Ints11 | 0.533509 | 634.2908 | 0.162652 | -3.28008 | 0.001038 | 0.006496 | 0.007869 | 792.2562 | 708.1572 | 495.634 | 541.1158 | 71957 | integrator complex subunit 11 [Source:MGI Symbol;Acc:MGI:1919207] |
| ENSMUSG00000029364 | Wsb2 | 0.533985 | 1871.614 | 0.102113 | -5.22937 | 1.7E-07 | 2.63E-06 | 3.19E-06 | 2170.507 | 2257.251 | 1496.046 | 1562.652 | 59043 | WD repeat and SOCS box-containing 2 [Source:MGI Symbol;Acc:MGI:2144041] |
| ENSMUSG00000033186 | Mzt1 | 0.534388 | 603.0511 | 0.169174 | -3.1588 | 0.001584 | 0.009239 | 0.011192 | 752.0616 | 674.7165 | 461.7992 | 523.6273 | 76789 | mitotic spindle organizing protein 1 [Source:MGI Symbol;Acc:MGI:1924039] |
| ENSMUSG00000024456 | Diaph1 | 0.534697 | 929.6651 | 0.146786 | -3.64269 | 0.00027 | 0.002012 | 0.002438 | 1192.086 | 1008.14 | 757.1678 | 761.2656 | 13367 | diaphanous related formin 1 [Source:MGI Symbol;Acc:MGI:1194490] |
| ENSMUSG00000053317 | Sec61b | 0.534799 | 919.4808 | 0.133937 | -3.99291 | 6.53E-05 | 0.000574 | 0.000695 | 1129.679 | 1046.499 | 758.9967 | 742.7484 | 66212 | SEC61 translocon subunit beta [Source:MGI Symbol;Acc:MGI:1913462] |
| ENSMUSG00000039298 | Cdk5rap2 | 0.535092 | 746.0588 | 0.151288 | -3.53691 | 0.000405 | 0.002883 | 0.003493 | 877.9341 | 888.1472 | 649.2622 | 568.8918 | 214444 | CDK5 regulatory subunit associated protein 2 [Source:MGI Symbol;Acc:MGI:2384875] |
| ENSMUSG00000030036 | Mogs | 0.535423 | 802.7139 | 0.157146 | -3.40716 | 0.000656 | 0.004375 | 0.0053 | 1037.655 | 862.5748 | 658.4068 | 652.2195 | 57377 | mannosyl-oligosaccharide glucosidase [Source:MGI Symbol;Acc:MGI:1929872] |
| ENSMUSG00000019373 | Cops3 | 0.535634 | 846.4951 | 0.13843 | -3.86935 | 0.000109 | 0.000897 | 0.001087 | 1041.886 | 961.9136 | 686.7548 | 695.4264 | 26572 | COP9 signalosome subunit 3 [Source:MGI Symbol;Acc:MGI:1349409] |
| ENSMUSG00000053907 | Mat2a | 0.53613 | 2276.188 | 0.114118 | -4.69803 | 2.63E-06 | 3.16E-05 | 3.83E-05 | 2807.274 | 2580.84 | 1730.147 | 1986.492 | 232087 | methionine adenosyltransferase 2A [Source:MGI Symbol;Acc:MGI:2443731] |
| ENSMUSG00000037960 | Card19 | 0.536418 | 644.3162 | 0.152671 | -3.51355 | 0.000442 | 0.003115 | 0.003773 | 785.9097 | 739.6309 | 523.982 | 527.7423 | 68480 | caspase recruitment domain family, member 19 [Source:MGI Symbol;Acc:MGI:1915730] |
| ENSMUSG00000043998 | Mgat2 | 0.536593 | 1947.223 | 0.100562 | -5.33593 | 9.51E-08 | 1.56E-06 | 1.89E-06 | 2331.285 | 2278.889 | 1548.17 | 1630.549 | 217664 | mannoside acetylglucosaminyltransferase 2 [Source:MGI Symbol;Acc:MGI:2384966] |
| ENSMUSG00000022186 | Oxct1 | 0.536692 | 5941.181 | 0.072384 | -7.41455 | 1.22E-13 | 4.55E-12 | 5.51E-12 | 7058.379 | 7008.789 | 4801.797 | 4895.761 | 67041 | 3-oxoacid CoA transferase 1 [Source:MGI Symbol;Acc:MGI:1914291] |
| ENSMUSG00000022009 | Nufip1 | 0.536769 | 356.8158 | 0.197775 | -2.71404 | 0.006647 | 0.030367 | 0.036686 | 434.736 | 410.1411 | 287.1385 | 295.2476 | 27275 | nuclear FMR1 interacting protein 1 [Source:MGI Symbol;Acc:MGI:1351474] |
| ENSMUSG00000034674 | Tdg | 0.536948 | 484.4126 | 0.181339 | -2.96102 | 0.003066 | 0.015912 | 0.019268 | 610.3229 | 537.0192 | 409.6753 | 380.6328 | 21665 | thymine DNA glycosylase [Source:MGI Symbol;Acc:MGI:108247] |
| ENSMUSG00000037851 | Iars1 | 0.536959 | 943.9403 | 0.176538 | -3.04161 | 0.002353 | 0.012748 | 0.015442 | 1265.071 | 970.7655 | 832.153 | 707.7713 | 105148 | isoleucyl-tRNA synthetase 1 [Source:MGI Symbol;Acc:MGI:2145219] |
| ENSMUSG00000058799 | Nap1l1 | 0.537153 | 1854.082 | 0.144535 | -3.71642 | 0.000202 | 0.001554 | 0.001882 | 2487.832 | 1903.173 | 1507.934 | 1517.388 | 53605 | nucleosome assembly protein 1-like 1 [Source:MGI Symbol;Acc:MGI:1855693] |
| ENSMUSG00000042426 | Dhx29 | 0.537662 | 503.8953 | 0.179872 | -2.98914 | 0.002798 | 0.014719 | 0.017824 | 635.7089 | 557.6738 | 393.2151 | 428.9835 | 218629 | DExH-box helicase 29 [Source:MGI Symbol;Acc:MGI:2145374] |
| ENSMUSG00000030471 | Zdhhc13 | 0.537908 | 488.09 | 0.173401 | -3.10211 | 0.001921 | 0.010799 | 0.013081 | 563.7818 | 592.0981 | 384.9851 | 411.4949 | 243983 | zinc finger, DHHC domain containing 13 [Source:MGI Symbol;Acc:MGI:1919227] |
| ENSMUSG00000029594 | Rbm19 | 0.538156 | 435.0947 | 0.189176 | -2.84474 | 0.004445 | 0.021755 | 0.026302 | 543.6845 | 486.8581 | 337.4335 | 372.4029 | 74111 | RNA binding motif protein 19 [Source:MGI Symbol;Acc:MGI:1921361] |
| ENSMUSG00000020708 | Psmc5 | 0.539283 | 1058.063 | 0.123168 | -4.37845 | 1.2E-05 | 0.000127 | 0.000154 | 1233.339 | 1273.699 | 864.1589 | 861.0532 | 19184 | protease (prosome, macropain) 26S subunit, ATPase 5 [Source:MGI Symbol;Acc:MGI:105047] |
| ENSMUSG00000002741 | Ykt6 | 0.539316 | 2086.489 | 0.105499 | -5.11203 | 3.19E-07 | 4.65E-06 | 5.63E-06 | 2589.377 | 2354.623 | 1663.391 | 1738.566 | 56418 | YKT6 v-SNARE homolog (S. cerevisiae) [Source:MGI Symbol;Acc:MGI:1927550] |
| ENSMUSG00000025156 | Gps1 | 0.539344 | 1310.542 | 0.128586 | -4.19443 | 2.74E-05 | 0.000265 | 0.000321 | 1642.689 | 1462.541 | 1018.702 | 1118.238 | 209318 | G protein pathway suppressor 1 [Source:MGI Symbol;Acc:MGI:2384801] |
| ENSMUSG00000019494 | Cops6 | 0.539918 | 632.9056 | 0.162763 | -3.3172 | 0.000909 | 0.005786 | 0.007008 | 777.4477 | 722.9105 | 547.7578 | 483.5066 | 26893 | COP9 signalosome subunit 6 [Source:MGI Symbol;Acc:MGI:1349439] |
| ENSMUSG00000021871 | Gm49342 | 0.540305 | 1298.387 | 0.124771 | -4.33037 | 1.49E-05 | 0.000154 | 0.000186 | 1624.707 | 1452.706 | 1028.761 | 1087.375 | | predicted gene, 49342 [Source:MGI Symbol;Acc:MGI:6121535] |
| ENSMUSG00000116564 | Riok2 | 0.540375 | 449.8667 | 0.187835 | -2.87686 | 0.004017 | 0.02006 | 0.024261 | 529.9337 | 536.0357 | 337.4335 | 396.0639 | 67045 | RIO kinase 2 [Source:MGI Symbol;Acc:MGI:1914295] |
| ENSMUSG00000022663 | Atg3 | 0.540712 | 663.3307 | 0.151339 | -3.57284 | 0.000353 | 0.002556 | 0.003096 | 811.2957 | 761.269 | 546.8434 | 533.9147 | 67841 | autophagy related 3 [Source:MGI Symbol;Acc:MGI:1915091] |
| ENSMUSG00000072620 | Slfn2 | 0.54081 | 2785.73 | 0.096851 | -5.58396 | 2.35E-08 | 4.33E-07 | 5.25E-07 | 3418.654 | 3184.74 | 2191.946 | 2347.579 | 20556 | schlafen 2 [Source:MGI Symbol;Acc:MGI:1313258] |
| ENSMUSG00000064341 | mt-Nd1 | 0.541149 | 2842.067 | 0.114537 | -4.72466 | 2.3E-06 | 2.81E-05 | 3.4E-05 | 3670.399 | 3067.698 | 2273.332 | 2356.837 | 17716 | mitochondrially encoded NADH dehydrogenase 1 [Source:MGI Symbol;Acc:MGI:101787] |
| ENSMUSG00000029407 | Uso1 | 0.541445 | 816.987 | 0.196365 | -2.75734 | 0.005827 | 0.027238 | 0.032904 | 1134.968 | 802.5782 | 705.9583 | 624.4436 | 56041 | USO1 vesicle docking factor [Source:MGI Symbol;Acc:MGI:1929095] |
| ENSMUSG00000030591 | Psmd8 | 0.542028 | 3123.574 | 0.091139 | -5.94726 | 2.73E-09 | 5.76E-08 | 6.98E-08 | 3781.463 | 3626.355 | 2630.884 | 2455.596 | 57296 | proteasome (prosome, macropain) 26S subunit, non-ATPase, 8 [Source:MGI Symbol;Acc:MGI:1888669] |
| ENSMUSG00000056201 | Cfl1 | 0.542055 | 12204.78 | 0.07925 | -6.83978 | 7.93E-12 | 2.39E-10 | 2.89E-10 | 15020.08 | 13923.16 | 10316.87 | 9559.027 | 12631 | cofilin 1, non-muscle [Source:MGI Symbol;Acc:MGI:101757] |
| ENSMUSG00000004897 | Hdgf | 0.542376 | 1294.529 | 0.133267 | -4.06984 | 4.7E-05 | 0.000431 | 0.000522 | 1663.844 | 1406.479 | 1046.135 | 1061.657 | 15191 | heparin binding growth factor [Source:MGI Symbol;Acc:MGI:1194494] |
| ENSMUSG00000001785 | Pwp1 | 0.542708 | 879.8179 | 0.141261 | -3.8419 | 0.000122 | 0.000993 | 0.001202 | 1070.445 | 1016.009 | 676.6958 | 756.122 | 103136 | PWP1 homolog, endonuclein [Source:MGI Symbol;Acc:MGI:1914735] |
| ENSMUSG00000022723 | Crybg3 | 0.543097 | 1980.929 | 0.122722 | -4.42544 | 9.62E-06 | 0.000104 | 0.000126 | 2540.72 | 2158.896 | 1678.937 | 1545.164 | 224273 | beta-gamma crystallin domain containing 3 [Source:MGI Symbol;Acc:MGI:2676311] |
| ENSMUSG00000044042 | Fmn1 | 0.543161 | 1035.076 | 0.155208 | -3.49956 | 0.000466 | 0.003256 | 0.003945 | 1368.731 | 1086.825 | 833.9819 | 850.7658 | 14260 | formin 1 [Source:MGI Symbol;Acc:MGI:101815] |
| ENSMUSG00000027722 | Afg2a | 0.543487 | 547.8356 | 0.202685 | -2.68143 | 0.007331 | 0.032964 | 0.039826 | 717.1558 | 582.2626 | 395.0441 | 496.8801 | 57815 | AFG2 AAA ATPase homolog A [Source:MGI Symbol;Acc:MGI:1927170] |
| ENSMUSG00000048612 | Myof | 0.543878 | 1426.878 | 0.127496 | -4.26586 | 1.99E-05 | 0.000199 | 0.000241 | 1692.403 | 1693.676 | 1254.631 | 1066.801 | 226101 | myoferlin [Source:MGI Symbol;Acc:MGI:1919192] |
| ENSMUSG00000031851 | Ntpcr | 0.544198 | 2011.196 | 0.112185 | -4.85089 | 1.23E-06 | 1.59E-05 | 1.93E-05 | 2532.258 | 2240.531 | 1682.595 | 1589.399 | 66566 | nucleoside-triphosphatase, cancer-related [Source:MGI Symbol;Acc:MGI:1913816] |
| ENSMUSG00000052533 | Nup188 | 0.544257 | 676.6577 | 0.199813 | -2.72384 | 0.006453 | 0.02965 | 0.035819 | 937.1682 | 668.8152 | 568.7903 | 531.8572 | 227699 | nucleoporin 188 [Source:MGI Symbol;Acc:MGI:2446190] |
| ENSMUSG00000014850 | Msh3 | 0.54431 | 370.1388 | 0.199082 | -2.7341 | 0.006255 | 0.028872 | 0.034879 | 462.2376 | 416.0424 | 292.6252 | 309.6499 | 17686 | mutS homolog 3 [Source:MGI Symbol;Acc:MGI:109519] |
| ENSMUSG00000030019 | Fbxl14 | 0.544398 | 594.295 | 0.174108 | -3.12679 | 0.001767 | 0.010082 | 0.012213 | 736.1953 | 673.7329 | 442.5956 | 524.656 | 101358 | F-box and leucine-rich repeat protein 14 [Source:MGI Symbol;Acc:MGI:2141676] |
| ENSMUSG00000025198 | Erlin1 | 0.545103 | 588.7274 | 0.209288 | -2.60456 | 0.009199 | 0.039752 | 0.048017 | 814.469 | 583.2462 | 506.6074 | 450.587 | 226144 | ER lipid raft associated 1 [Source:MGI Symbol;Acc:MGI:2387613] |
| ENSMUSG00000005899 | Smpd4 | 0.545131 | 516.4481 | 0.180256 | -3.02421 | 0.002493 | 0.013375 | 0.016197 | 664.2682 | 561.608 | 416.0765 | 423.8398 | 77626 | sphingomyelin phosphodiesterase 4 [Source:MGI Symbol;Acc:MGI:1924876] |
| ENSMUSG00000024583 | Txnl1 | 0.545196 | 1392.145 | 0.120044 | -4.54164 | 5.58E-06 | 6.33E-05 | 7.67E-05 | 1713.558 | 1590.403 | 1083.628 | 1180.99 | 53382 | thioredoxin-like 1 [Source:MGI Symbol;Acc:MGI:1860078] |
| ENSMUSG00000001380 | Hars1 | 0.545389 | 867.2685 | 0.146992 | -3.71034 | 0.000207 | 0.001586 | 0.001921 | 1091.6 | 966.8313 | 673.038 | 737.6047 | 15115 | histidyl-tRNA synthetase 1 [Source:MGI Symbol;Acc:MGI:108087] |
| ENSMUSG00000035478 | Mbd3 | 0.545937 | 425.6411 | 0.184126 | -2.96502 | 0.003027 | 0.015753 | 0.019076 | 523.5872 | 486.8581 | 339.2624 | 352.8569 | 17192 | methyl-CpG binding domain protein 3 [Source:MGI Symbol;Acc:MGI:1333812] |
| ENSMUSG00000025076 | Casp7 | 0.547037 | 581.258 | 0.170232 | -3.21348 | 0.001311 | 0.007878 | 0.009543 | 717.1558 | 662.9138 | 439.8523 | 505.11 | 12369 | caspase 7 [Source:MGI Symbol;Acc:MGI:109383] |
| ENSMUSG00000027613 | Eif6 | 0.54712 | 975.5934 | 0.149193 | -3.6672 | 0.000245 | 0.001848 | 0.002239 | 1254.494 | 1062.236 | 754.4244 | 831.2198 | 16418 | eukaryotic translation initiation factor 6 [Source:MGI Symbol;Acc:MGI:1196288] |
| ENSMUSG00000032803 | Cdv3 | 0.547136 | 2757.441 | 0.092669 | -5.90419 | 3.54E-09 | 7.39E-08 | 8.95E-08 | 3343.554 | 3204.411 | 2176.4 | 2305.4 | 321022 | carnitine deficiency-associated gene expressed in ventricle 3 [Source:MGI Symbol;Acc:MGI:2448759] |
| ENSMUSG00000020949 | Fkbp3 | 0.547772 | 637.0378 | 0.162242 | -3.37626 | 0.000735 | 0.004825 | 0.005845 | 772.1589 | 740.6144 | 481.9172 | 553.4607 | 30795 | FK506 binding protein 3 [Source:MGI Symbol;Acc:MGI:1353460] |
| ENSMUSG00000040681 | Hmgn1 | 0.54854 | 2447.882 | 0.10591 | -5.1793 | 2.23E-07 | 3.37E-06 | 4.08E-06 | 3064.307 | 2750.994 | 1916.695 | 2059.532 | 15312 | high mobility group nucleosomal binding domain 1 [Source:MGI Symbol;Acc:MGI:96120] |
| ENSMUSG00000057497 | Fam136a | 0.548688 | 339.8841 | 0.205937 | -2.66435 | 0.007714 | 0.034462 | 0.041635 | 423.1008 | 384.5687 | 284.3951 | 267.4717 | 66488 | family with sequence similarity 136, member A [Source:MGI Symbol;Acc:MGI:1913738] |
| ENSMUSG00000028953 | Abcf2 | 0.548701 | 1327.332 | 0.125422 | -4.37483 | 1.22E-05 | 0.000129 | 0.000156 | 1623.649 | 1529.423 | 1012.3 | 1143.956 | 27407 | ATP-binding cassette, sub-family F member 2 [Source:MGI Symbol;Acc:MGI:1351657] |
| ENSMUSG00000040354 | Mars1 | 0.548726 | 771.9085 | 0.161329 | -3.40127 | 0.000671 | 0.004451 | 0.005392 | 996.4023 | 837.986 | 656.5778 | 596.6677 | 216443 | methionine-tRNA synthetase 1 [Source:MGI Symbol;Acc:MGI:1345633] |
| ENSMUSG00000052997 | Uba2 | 0.54962 | 1927.507 | 0.114579 | -4.79684 | 1.61E-06 | 2.04E-05 | 2.47E-05 | 2426.483 | 2153.978 | 1505.191 | 1624.376 | 50995 | ubiquitin-like modifier activating enzyme 2 [Source:MGI Symbol;Acc:MGI:1858313] |
| ENSMUSG00000021629 | Slc30a5 | 0.550171 | 2162.589 | 0.104349 | -5.27243 | 1.35E-07 | 2.13E-06 | 2.58E-06 | 2691.979 | 2448.06 | 1718.259 | 1792.06 | 69048 | solute carrier family 30 (zinc transporter), member 5 [Source:MGI Symbol;Acc:MGI:1916298] |
| ENSMUSG00000024726 | Carnmt1 | 0.550756 | 549.6931 | 0.182602 | -3.01616 | 0.00256 | 0.013687 | 0.016574 | 707.636 | 598.983 | 417.9054 | 474.2479 | 67383 | carnosine N-methyltransferase 1 [Source:MGI Symbol;Acc:MGI:1914633] |
| ENSMUSG00000075705 | Msrb1 | 0.551137 | 757.0844 | 0.147179 | -3.74466 | 0.000181 | 0.001405 | 0.001702 | 853.6058 | 946.1767 | 615.4274 | 613.1275 | 27361 | methionine sulfoxide reductase B1 [Source:MGI Symbol;Acc:MGI:1351642] |
| ENSMUSG00000043067 | Dpy19l1 | 0.551176 | 1680.885 | 0.107258 | -5.1388 | 2.77E-07 | 4.09E-06 | 4.96E-06 | 2054.154 | 1942.515 | 1393.628 | 1333.244 | 244745 | dpy-19 like C-mannosyltransferase 1 [Source:MGI Symbol;Acc:MGI:1915685] |
| ENSMUSG00000018583 | G3bp1 | 0.551269 | 3305.573 | 0.107443 | -5.13081 | 2.88E-07 | 4.26E-06 | 5.16E-06 | 4244.758 | 3614.552 | 2632.713 | 2730.269 | 27041 | G3BP stress granule assembly factor 1 [Source:MGI Symbol;Acc:MGI:1351465] |
| ENSMUSG00000022370 | Mrpl13 | 0.551945 | 563.9027 | 0.182218 | -3.02904 | 0.002453 | 0.013218 | 0.016012 | 726.6756 | 614.7198 | 488.3183 | 425.8973 | 68537 | mitochondrial ribosomal protein L13 [Source:MGI Symbol;Acc:MGI:2137218] |
| ENSMUSG00000001089 | Luzp1 | 0.552021 | 1666.508 | 0.126155 | -4.37573 | 1.21E-05 | 0.000128 | 0.000155 | 2154.641 | 1808.752 | 1357.049 | 1345.588 | 269593 | leucine zipper protein 1 [Source:MGI Symbol;Acc:MGI:107629] |
| ENSMUSG00000072774 | Zfp951 | 0.552601 | 350.8503 | 0.203762 | -2.712 | 0.006688 | 0.030535 | 0.03689 | 441.0825 | 393.4207 | 279.8229 | 289.0752 | 626391 | zinc finger protein 951 [Source:MGI Symbol;Acc:MGI:2441896] |
| ENSMUSG00000028577 | Plaa | 0.552831 | 1557.215 | 0.130388 | -4.23989 | 2.24E-05 | 0.000221 | 0.000267 | 1986.458 | 1717.281 | 1188.79 | 1336.33 | 18786 | phospholipase A2, activating protein [Source:MGI Symbol;Acc:MGI:104810] |
| ENSMUSG00000039221 | Rpl22l1 | 0.552919 | 809.6804 | 0.15273 | -3.62024 | 0.000294 | 0.00217 | 0.002628 | 947.7457 | 978.6339 | 710.5306 | 601.8114 | 68028 | ribosomal protein L22 like 1 [Source:MGI Symbol;Acc:MGI:1915278] |
| ENSMUSG00000029500 | Pgam5 | 0.552976 | 532.1758 | 0.175317 | -3.15414 | 0.00161 | 0.009345 | 0.01132 | 672.7302 | 593.0817 | 413.3331 | 449.5582 | 72542 | phosphoglycerate mutase family member 5 [Source:MGI Symbol;Acc:MGI:1919792] |
| ENSMUSG00000031917 | Nip7 | 0.55316 | 457.5008 | 0.175166 | -3.15793 | 0.001589 | 0.00926 | 0.011217 | 539.4535 | 548.8218 | 370.3538 | 371.3742 | 66164 | NIP7, nucleolar pre-rRNA processing protein [Source:MGI Symbol;Acc:MGI:1913414] |
| ENSMUSG00000034330 | Plcg2 | 0.553207 | 3296.31 | 0.105515 | -5.24294 | 1.58E-07 | 2.47E-06 | 2.99E-06 | 4129.463 | 3712.908 | 2826.577 | 2516.292 | 234779 | phospholipase C, gamma 2 [Source:MGI Symbol;Acc:MGI:97616] |
| ENSMUSG00000030659 | Nucb2 | 0.553957 | 367.7772 | 0.203428 | -2.72311 | 0.006467 | 0.02967 | 0.035844 | 448.4868 | 426.8614 | 320.0588 | 275.7016 | 53322 | nucleobindin 2 [Source:MGI Symbol;Acc:MGI:1858179] |
| ENSMUSG00000021548 | Ccnh | 0.55435 | 378.6618 | 0.202114 | -2.74276 | 0.006092 | 0.028303 | 0.034191 | 461.1798 | 439.6476 | 282.5662 | 331.2534 | 66671 | cyclin H [Source:MGI Symbol;Acc:MGI:1913921] |
| ENSMUSG00000021550 | Qng1 | 0.55453 | 563.711 | 0.176167 | -3.14775 | 0.001645 | 0.009522 | 0.011535 | 728.7911 | 612.7527 | 446.2535 | 467.0468 | 70153 | Q-nucleotide N-glycosylase 1 [Source:MGI Symbol;Acc:MGI:1917403] |
| ENSMUSG00000047547 | Cltb | 0.55484 | 456.4742 | 0.190583 | -2.91128 | 0.003599 | 0.018223 | 0.022053 | 515.1252 | 571.4435 | 397.7874 | 341.5408 | 74325 | clathrin light chain B [Source:MGI Symbol;Acc:MGI:1921575] |
| ENSMUSG00000037364 | Srrt | 0.554886 | 1478.432 | 0.109023 | -5.0896 | 3.59E-07 | 5.18E-06 | 6.28E-06 | 1789.716 | 1729.084 | 1209.822 | 1185.105 | 83701 | serrate RNA effector molecule homolog (Arabidopsis) [Source:MGI Symbol;Acc:MGI:1933527] |
| ENSMUSG00000029415 | Sdad1 | 0.554889 | 400.0658 | 0.192815 | -2.87783 | 0.004004 | 0.020005 | 0.024194 | 495.0279 | 457.3515 | 339.2624 | 308.6212 | 231452 | SDA1 domain containing 1 [Source:MGI Symbol;Acc:MGI:2140779] |
| ENSMUSG00000028224 | Nbn | 0.555893 | 522.7973 | 0.182097 | -3.05273 | 0.002268 | 0.012373 | 0.014988 | 662.1527 | 582.2626 | 394.1296 | 452.6444 | 27354 | nibrin [Source:MGI Symbol;Acc:MGI:1351625] |
| ENSMUSG00000044408 | Sptssa | 0.55612 | 2255.043 | 0.093615 | -5.94052 | 2.84E-09 | 5.99E-08 | 7.26E-08 | 2658.131 | 2710.668 | 1840.796 | 1810.578 | 104725 | serine palmitoyltransferase, small subunit A [Source:MGI Symbol;Acc:MGI:1913399] |
| ENSMUSG00000030007 | Cct7 | 0.556195 | 2333.01 | 0.107535 | -5.17221 | 2.31E-07 | 3.48E-06 | 4.22E-06 | 2951.128 | 2603.461 | 1852.683 | 1924.768 | 12468 | chaperonin containing TCP1 subunit 7 [Source:MGI Symbol;Acc:MGI:107184] |
| ENSMUSG00000024208 | Uqcc2 | 0.556413 | 591.9611 | 0.158938 | -3.50082 | 0.000464 | 0.003245 | 0.00393 | 712.9248 | 696.3546 | 462.7136 | 495.8514 | 67267 | ubiquinol-cytochrome c reductase complex assembly factor 2 [Source:MGI Symbol;Acc:MGI:1914517] |
| ENSMUSG00000058006 | Mdn1 | 0.55654 | 1879.092 | 0.136943 | -4.06401 | 4.82E-05 | 0.00044 | 0.000533 | 2501.583 | 1973.005 | 1516.164 | 1525.617 | 100019 | midasin AAA ATPase 1 [Source:MGI Symbol;Acc:MGI:1926159] |
| ENSMUSG00000028882 | Ppp1r8 | 0.556857 | 558.5726 | 0.166004 | -3.35448 | 0.000795 | 0.005166 | 0.006258 | 690.712 | 639.3086 | 436.1945 | 468.0755 | 100336 | protein phosphatase 1, regulatory subunit 8 [Source:MGI Symbol;Acc:MGI:2140494] |
| ENSMUSG00000034484 | Snx2 | 0.556876 | 9493.922 | 0.079501 | -7.00465 | 2.48E-12 | 7.92E-11 | 9.59E-11 | 11711.43 | 10895.79 | 7415.306 | 7953.168 | 67804 | sorting nexin 2 [Source:MGI Symbol;Acc:MGI:1915054] |
| ENSMUSG00000024097 | Srsf7 | 0.557011 | 1493.833 | 0.126453 | -4.4049 | 1.06E-05 | 0.000114 | 0.000138 | 1841.546 | 1715.314 | 1120.206 | 1298.267 | 225027 | serine and arginine-rich splicing factor 7 [Source:MGI Symbol;Acc:MGI:1926232] |
| ENSMUSG00000069744 | Psmb3 | 0.557726 | 1315.959 | 0.11997 | -4.6489 | 3.34E-06 | 3.93E-05 | 4.76E-05 | 1638.458 | 1495.982 | 1046.135 | 1083.26 | 26446 | proteasome (prosome, macropain) subunit, beta type 3 [Source:MGI Symbol;Acc:MGI:1347014] |
| ENSMUSG00000061479 | Snrpa | 0.558267 | 447.6501 | 0.182539 | -3.05834 | 0.002226 | 0.012179 | 0.014753 | 547.9155 | 518.3317 | 344.7491 | 379.6041 | 53607 | small nuclear ribonucleoprotein polypeptide A [Source:MGI Symbol;Acc:MGI:1855690] |
| ENSMUSG00000019920 | Lims1 | 0.558529 | 3024.564 | 0.086984 | -6.42108 | 1.35E-10 | 3.48E-09 | 4.22E-09 | 3659.822 | 3545.704 | 2411.415 | 2481.314 | 110829 | LIM and senescent cell antigen-like domains 1 [Source:MGI Symbol;Acc:MGI:1195263] |
| ENSMUSG00000033596 | Rfwd3 | 0.558941 | 1247.219 | 0.139494 | -4.00692 | 6.15E-05 | 0.000544 | 0.000659 | 1618.36 | 1353.367 | 972.9789 | 1044.168 | 234736 | ring finger and WD repeat domain 3 [Source:MGI Symbol;Acc:MGI:2384584] |
| ENSMUSG00000050244 | Heatr1 | 0.559646 | 1999.496 | 0.105273 | -5.31616 | 1.06E-07 | 1.72E-06 | 2.09E-06 | 2494.179 | 2271.021 | 1601.209 | 1631.577 | 217995 | HEAT repeat containing 1 [Source:MGI Symbol;Acc:MGI:2442524] |
| ENSMUSG00000026315 | Serpinb8 | 0.559779 | 531.0624 | 0.1644 | -3.40498 | 0.000662 | 0.004401 | 0.005331 | 643.1132 | 622.5882 | 431.6222 | 426.926 | 20725 | serine (or cysteine) peptidase inhibitor, clade B, member 8 [Source:MGI Symbol;Acc:MGI:894657] |
| ENSMUSG00000060675 | Plaat3 | 0.560402 | 520.8663 | 0.178219 | -3.14446 | 0.001664 | 0.009604 | 0.011634 | 640.9977 | 600.9501 | 452.6546 | 388.8627 | 225845 | phospholipase A and acyltransferase 3 [Source:MGI Symbol;Acc:MGI:2179715] |
| ENSMUSG00000003808 | Farsa | 0.560527 | 730.0839 | 0.142772 | -3.92602 | 8.64E-05 | 0.000732 | 0.000887 | 875.8186 | 864.5419 | 592.5661 | 587.409 | 66590 | phenylalanyl-tRNA synthetase, alpha subunit [Source:MGI Symbol;Acc:MGI:1913840] |
| ENSMUSG00000001891 | Ugp2 | 0.560801 | 1702.833 | 0.104676 | -5.35747 | 8.44E-08 | 1.41E-06 | 1.7E-06 | 2079.54 | 1979.89 | 1366.194 | 1385.709 | 216558 | UDP-glucose pyrophosphorylase 2 [Source:MGI Symbol;Acc:MGI:2183447] |
| ENSMUSG00000028282 | Casp8ap2 | 0.561034 | 991.5189 | 0.146171 | -3.8382 | 0.000124 | 0.001007 | 0.001219 | 1286.226 | 1077.973 | 813.8639 | 788.0128 | 26885 | caspase 8 associated protein 2 [Source:MGI Symbol;Acc:MGI:1349399] |
| ENSMUSG00000034345 | Gtf2h5 | 0.561654 | 540.5473 | 0.163648 | -3.43208 | 0.000599 | 0.004047 | 0.004903 | 638.8822 | 650.1277 | 446.2535 | 426.926 | 66467 | general transcription factor IIH, polypeptide 5 [Source:MGI Symbol;Acc:MGI:107227] |
| ENSMUSG00000027429 | Sec23b | 0.56185 | 1146.355 | 0.124202 | -4.52369 | 6.08E-06 | 6.84E-05 | 8.29E-05 | 1372.962 | 1360.252 | 885.1913 | 967.0131 | 27054 | SEC23 homolog B, COPII coat complex component [Source:MGI Symbol;Acc:MGI:1350925] |
| ENSMUSG00000015143 | Actn1 | 0.562031 | 818.3577 | 0.170789 | -3.2908 | 0.000999 | 0.006275 | 0.007602 | 1089.484 | 862.5748 | 695.8994 | 625.4723 | 109711 | actinin, alpha 1 [Source:MGI Symbol;Acc:MGI:2137706] |
| ENSMUSG00000022131 | Gpr180 | 0.562362 | 322.3765 | 0.210111 | -2.6765 | 0.007439 | 0.033394 | 0.040345 | 404.0612 | 364.8977 | 263.3627 | 257.1843 | 58245 | G protein-coupled receptor 180 [Source:MGI Symbol;Acc:MGI:1930949] |
| ENSMUSG00000048677 | Tpcn2 | 0.562529 | 617.0718 | 0.163158 | -3.44775 | 0.000565 | 0.003854 | 0.004669 | 739.3686 | 732.746 | 532.2121 | 463.9605 | 233979 | two pore segment channel 2 [Source:MGI Symbol;Acc:MGI:2385297] |
| ENSMUSG00000024359 | Hspa9 | 0.562586 | 3710.352 | 0.114919 | -4.89549 | 9.81E-07 | 1.29E-05 | 1.57E-05 | 4739.786 | 4109.279 | 2774.453 | 3217.89 | 15526 | heat shock protein 9 [Source:MGI Symbol;Acc:MGI:96245] |
| ENSMUSG00000025358 | Cdk2 | 0.562776 | 736.3865 | 0.15386 | -3.65773 | 0.000254 | 0.001911 | 0.002315 | 938.2259 | 818.315 | 586.1649 | 602.8401 | 12566 | cyclin dependent kinase 2 [Source:MGI Symbol;Acc:MGI:104772] |
| ENSMUSG00000020766 | Galk1 | 0.563562 | 440.1046 | 0.178218 | -3.1622 | 0.001566 | 0.009156 | 0.011092 | 526.7605 | 523.2495 | 357.5514 | 352.8569 | 14635 | galactokinase 1 [Source:MGI Symbol;Acc:MGI:95730] |
| ENSMUSG00000032020 | Ubash3b | 0.563639 | 1580.911 | 0.11634 | -4.84476 | 1.27E-06 | 1.64E-05 | 1.98E-05 | 1971.65 | 1799.9 | 1228.111 | 1323.985 | 72828 | ubiquitin associated and SH3 domain containing, B [Source:MGI Symbol;Acc:MGI:1920078] |
| ENSMUSG00000029777 | Gars1 | 0.563742 | 1008.762 | 0.146134 | -3.85771 | 0.000114 | 0.000935 | 0.001133 | 1304.208 | 1102.561 | 783.6869 | 844.5934 | 353172 | glycyl-tRNA synthetase 1 [Source:MGI Symbol;Acc:MGI:2449057] |
| ENSMUSG00000026509 | Capn2 | 0.563935 | 790.5399 | 0.139548 | -4.04115 | 5.32E-05 | 0.000479 | 0.00058 | 947.7457 | 938.3083 | 621.8286 | 654.277 | 12334 | calpain 2 [Source:MGI Symbol;Acc:MGI:88264] |
| ENSMUSG00000044894 | Uqcrq | 0.564574 | 403.9114 | 0.203571 | -2.77335 | 0.005548 | 0.026182 | 0.031627 | 523.5872 | 440.6312 | 343.8346 | 307.5925 | 22272 | ubiquinol-cytochrome c reductase, complex III subunit VII [Source:MGI Symbol;Acc:MGI:107807] |
| ENSMUSG00000058267 | Mrps14 | 0.564705 | 547.5865 | 0.166367 | -3.39433 | 0.000688 | 0.004555 | 0.005518 | 683.3077 | 623.5718 | 438.0234 | 445.4433 | 64659 | mitochondrial ribosomal protein S14 [Source:MGI Symbol;Acc:MGI:1928141] |
| ENSMUSG00000062082 | Cd200r4 | 0.56491 | 793.66 | 0.143451 | -3.938 | 8.22E-05 | 0.000701 | 0.000849 | 946.688 | 947.1603 | 609.0262 | 671.7655 | 239849 | CD200 receptor 4 [Source:MGI Symbol;Acc:MGI:3036289] |
| ENSMUSG00000029147 | Ppm1g | 0.565225 | 511.3615 | 0.199498 | -2.83324 | 0.004608 | 0.022415 | 0.027101 | 690.712 | 530.1344 | 423.3921 | 401.2076 | 14208 | protein phosphatase 1G (formerly 2C), magnesium-dependent, gamma isoform [Source:MGI Symbol;Acc:MGI:106065] |
| ENSMUSG00000018433 | Nol11 | 0.565755 | 655.6205 | 0.159242 | -3.5528 | 0.000381 | 0.002729 | 0.003305 | 833.5085 | 731.7625 | 529.4688 | 527.7423 | 68979 | nucleolar protein 11 [Source:MGI Symbol;Acc:MGI:1916229] |
| ENSMUSG00000062352 | Itgb1bp1;Itgb1b | 0.566229 | 673.2496 | 0.151113 | -3.74705 | 0.000179 | 0.001394 | 0.001689 | 814.469 | 792.7427 | 523.0676 | 562.7193 | 16413;105245 | integrin beta 1 binding protein 1 [Source:MGI Symbol;Acc:MGI:1306802];integrin beta 1 binding protein 1 [Source:MGI Symbol;Acc:MGI:1306802] |
| ENSMUSG00000059208 | Hnrnpm | 0.566588 | 1865.72 | 0.105617 | -5.36454 | 8.12E-08 | 1.36E-06 | 1.64E-06 | 2213.875 | 2241.514 | 1568.288 | 1439.204 | 76936 | heterogeneous nuclear ribonucleoprotein M [Source:MGI Symbol;Acc:MGI:1926465] |
| ENSMUSG00000026617 | Bpnt1 | 0.566664 | 524.6757 | 0.179261 | -3.16111 | 0.001572 | 0.009184 | 0.011125 | 668.4992 | 584.2297 | 400.5308 | 445.4433 | 23827 | 3'(2'), 5'-bisphosphate nucleotidase 1 [Source:MGI Symbol;Acc:MGI:1338800] |
| ENSMUSG00000024287 | Thoc1 | 0.566934 | 920.2689 | 0.141661 | -4.00206 | 6.28E-05 | 0.000554 | 0.000671 | 1136.026 | 1061.252 | 696.8138 | 786.9841 | 225160 | THO complex 1 [Source:MGI Symbol;Acc:MGI:1919668] |
| ENSMUSG00000029782 | Tmem209 | 0.567023 | 452.8786 | 0.178093 | -3.18386 | 0.001453 | 0.008581 | 0.010395 | 554.262 | 527.1837 | 356.637 | 373.4317 | 72649 | transmembrane protein 209 [Source:MGI Symbol;Acc:MGI:1919899] |
| ENSMUSG00000028837 | Psmb2 | 0.567112 | 1238.03 | 0.130319 | -4.35171 | 1.35E-05 | 0.000141 | 0.000171 | 1577.108 | 1379.923 | 1026.017 | 969.0706 | 26445 | proteasome (prosome, macropain) subunit, beta type 2 [Source:MGI Symbol;Acc:MGI:1347045] |
| ENSMUSG00000059588 | Calcrl | 0.567632 | 524.7375 | 0.196027 | -2.89568 | 0.003783 | 0.019072 | 0.023065 | 706.5783 | 546.8547 | 412.4187 | 433.0984 | 54598 | calcitonin receptor-like [Source:MGI Symbol;Acc:MGI:1926944] |
| ENSMUSG00000034932 | Mrpl54 | 0.567765 | 346.4292 | 0.200714 | -2.82873 | 0.004673 | 0.022677 | 0.027417 | 426.274 | 401.2891 | 283.4807 | 274.6729 | 66047 | mitochondrial ribosomal protein L54 [Source:MGI Symbol;Acc:MGI:1913297] |
| ENSMUSG00000078652 | Psme3 | 0.56821 | 725.4985 | 0.148663 | -3.82214 | 0.000132 | 0.001068 | 0.001293 | 896.9736 | 836.0189 | 565.1325 | 603.8688 | 19192 | proteaseome (prosome, macropain) activator subunit 3 (PA28 gamma, Ki) [Source:MGI Symbol;Acc:MGI:1096366] |
| ENSMUSG00000006095 | Tbcb | 0.569621 | 803.7796 | 0.145148 | -3.92441 | 8.69E-05 | 0.000736 | 0.000892 | 907.5511 | 1013.058 | 639.2032 | 655.3057 | 66411 | tubulin folding cofactor B [Source:MGI Symbol;Acc:MGI:1913661] |
| ENSMUSG00000037370 | Enpp1 | 0.569852 | 1111.546 | 0.126321 | -4.51113 | 6.45E-06 | 7.21E-05 | 8.73E-05 | 1365.558 | 1291.403 | 928.1706 | 861.0532 | 18605 | ectonucleotide pyrophosphatase/phosphodiesterase 1 [Source:MGI Symbol;Acc:MGI:97370] |
| ENSMUSG00000000399 | Ndufa9 | 0.569985 | 1285.038 | 0.122699 | -4.64541 | 3.39E-06 | 3.99E-05 | 4.84E-05 | 1613.072 | 1458.607 | 1056.194 | 1012.278 | 66108 | NADH:ubiquinone oxidoreductase subunit A9 [Source:MGI Symbol;Acc:MGI:1913358] |
| ENSMUSG00000028792 | Ak2 | 0.570042 | 910.6761 | 0.135869 | -4.19554 | 2.72E-05 | 0.000264 | 0.00032 | 1136.026 | 1040.598 | 723.333 | 742.7484 | 11637 | adenylate kinase 2 [Source:MGI Symbol;Acc:MGI:87978] |
| ENSMUSG00000003868 | Ruvbl2 | 0.570078 | 547.7001 | 0.16362 | -3.48416 | 0.000494 | 0.003427 | 0.004152 | 673.788 | 635.3744 | 444.4246 | 437.2134 | 20174 | RuvB-like AAA ATPase 2 [Source:MGI Symbol;Acc:MGI:1342299] |
| ENSMUSG00000024516 | Sec11c | 0.570309 | 1446.676 | 0.119141 | -4.78682 | 1.69E-06 | 2.13E-05 | 2.57E-05 | 1824.622 | 1633.679 | 1182.389 | 1146.013 | 66286 | SEC11 homolog C, signal peptidase complex subunit [Source:MGI Symbol;Acc:MGI:1913536] |
| ENSMUSG00000020929 | Eftud2 | 0.570698 | 1090.056 | 0.128893 | -4.42768 | 9.53E-06 | 0.000103 | 0.000125 | 1357.096 | 1249.111 | 905.3093 | 848.7083 | 20624 | elongation factor Tu GTP binding domain containing 2 [Source:MGI Symbol;Acc:MGI:1336880] |
| ENSMUSG00000022241 | Tars1 | 0.570706 | 1020.451 | 0.143376 | -3.98049 | 6.88E-05 | 0.000599 | 0.000725 | 1323.248 | 1116.331 | 812.035 | 830.191 | 110960 | threonyl-tRNA synthetase 1 [Source:MGI Symbol;Acc:MGI:106314] |
| ENSMUSG00000005846 | Rsl1d1 | 0.571716 | 1135.96 | 0.131074 | -4.36179 | 1.29E-05 | 0.000136 | 0.000164 | 1440.658 | 1275.667 | 893.4214 | 934.0935 | 66409 | ribosomal L1 domain containing 1 [Source:MGI Symbol;Acc:MGI:1913659] |
| ENSMUSG00000044763 | Trmt10c | 0.57248 | 343.141 | 0.19961 | -2.868 | 0.004131 | 0.020516 | 0.024807 | 412.5232 | 408.174 | 276.1651 | 275.7016 | 52575 | tRNA methyltransferase 10C, mitochondrial RNase P subunit [Source:MGI Symbol;Acc:MGI:1196261] |
| ENSMUSG00000041879 | Ipo9 | 0.572584 | 1906.728 | 0.124816 | -4.58744 | 4.49E-06 | 5.15E-05 | 6.24E-05 | 2485.717 | 2075.294 | 1570.117 | 1495.784 | 226432 | importin 9 [Source:MGI Symbol;Acc:MGI:1918944] |
| ENSMUSG00000024556 | Me2 | 0.572828 | 608.9628 | 0.16254 | -3.52422 | 0.000425 | 0.003007 | 0.003642 | 772.1589 | 684.552 | 487.4039 | 491.7365 | 107029 | malic enzyme 2, NAD(+)-dependent, mitochondrial [Source:MGI Symbol;Acc:MGI:2147351] |
| ENSMUSG00000024078 | Ttc27 | 0.573745 | 395.5146 | 0.207426 | -2.76602 | 0.005675 | 0.026655 | 0.032199 | 520.4139 | 425.8779 | 304.5131 | 331.2534 | 74196 | tetratricopeptide repeat domain 27 [Source:MGI Symbol;Acc:MGI:1921446] |
| ENSMUSG00000022035 | Ccdc25 | 0.574036 | 393.5402 | 0.19338 | -2.96844 | 0.002993 | 0.015607 | 0.018899 | 497.1434 | 444.5654 | 314.5721 | 317.8798 | 67179 | coiled-coil domain containing 25 [Source:MGI Symbol;Acc:MGI:1914429] |
| ENSMUSG00000046949 | Nqo2 | 0.574377 | 786.7139 | 0.152807 | -3.75884 | 0.000171 | 0.001337 | 0.00162 | 1011.211 | 871.4268 | 618.1708 | 646.0471 | 18105 | N-ribosyldihydronicotinamide quinone reductase 2 [Source:MGI Symbol;Acc:MGI:104513] |
| ENSMUSG00000025580 | Eif4a3 | 0.574669 | 1413.589 | 0.126844 | -4.53052 | 5.88E-06 | 6.65E-05 | 8.05E-05 | 1818.276 | 1564.831 | 1121.12 | 1150.128 | 192170 | eukaryotic translation initiation factor 4A3 [Source:MGI Symbol;Acc:MGI:1923731] |
| ENSMUSG00000021196 | Pfkp | 0.574935 | 519.0812 | 0.222059 | -2.58911 | 0.009622 | 0.041333 | 0.049928 | 514.0674 | 727.8283 | 385.8995 | 448.5295 | 56421 | phosphofructokinase, platelet [Source:MGI Symbol;Acc:MGI:1891833] |
| ENSMUSG00000030983 | Bccip | 0.575121 | 425.6854 | 0.181424 | -3.17005 | 0.001524 | 0.008933 | 0.010822 | 517.2407 | 501.6114 | 339.2624 | 344.627 | 66165 | BRCA2 and CDKN1A interacting protein [Source:MGI Symbol;Acc:MGI:1913415] |
| ENSMUSG00000053641 | Dennd4a | 0.575547 | 1358.675 | 0.12379 | -4.64937 | 3.33E-06 | 3.93E-05 | 4.76E-05 | 1726.251 | 1526.472 | 1111.061 | 1070.916 | 102442 | DENN domain containing 4A [Source:MGI Symbol;Acc:MGI:2142979] |
| ENSMUSG00000063406 | Tmed5 | 0.575947 | 1951.454 | 0.124742 | -4.61712 | 3.89E-06 | 4.53E-05 | 5.49E-05 | 2474.082 | 2197.254 | 1449.409 | 1685.072 | 73130 | transmembrane p24 trafficking protein 5 [Source:MGI Symbol;Acc:MGI:1921586] |
| ENSMUSG00000064127 | Med14 | 0.576204 | 624.5371 | 0.210478 | -2.7376 | 0.006189 | 0.028619 | 0.034573 | 872.6453 | 622.5882 | 461.7992 | 541.1158 | 26896 | mediator complex subunit 14 [Source:MGI Symbol;Acc:MGI:1349442] |
| ENSMUSG00000068394 | Cep152 | 0.576342 | 295.5975 | 0.218958 | -2.6322 | 0.008483 | 0.037252 | 0.044996 | 373.3864 | 334.4076 | 235.9291 | 238.6671 | 99100 | centrosomal protein 152 [Source:MGI Symbol;Acc:MGI:2139083] |
| ENSMUSG00000047554 | Tmem41b | 0.576518 | 790.7165 | 0.138732 | -4.15563 | 3.24E-05 | 0.00031 | 0.000375 | 950.919 | 942.2425 | 623.6575 | 646.0471 | 233724 | transmembrane protein 41B [Source:MGI Symbol;Acc:MGI:1289225] |
| ENSMUSG00000031231 | Cox7b | 0.578658 | 1638.211 | 0.109652 | -5.27722 | 1.31E-07 | 2.08E-06 | 2.52E-06 | 2041.461 | 1883.501 | 1304.926 | 1322.956 | 66142 | cytochrome c oxidase subunit 7B [Source:MGI Symbol;Acc:MGI:1913392] |
| ENSMUSG00000023106 | Denr | 0.578744 | 419.4097 | 0.204113 | -2.83542 | 0.004577 | 0.022304 | 0.026966 | 557.4353 | 447.516 | 330.1178 | 342.5695 | 68184 | density-regulated protein [Source:MGI Symbol;Acc:MGI:1915434] |
| ENSMUSG00000078453 | Abracl | 0.579125 | 838.3172 | 0.15147 | -3.82337 | 0.000132 | 0.001063 | 0.001287 | 1085.253 | 923.555 | 659.3212 | 685.1391 | 73112 | ABRA C-terminal like [Source:MGI Symbol;Acc:MGI:1920362] |
| ENSMUSG00000029394 | Cdk2ap1 | 0.579365 | 603.4404 | 0.164751 | -3.5166 | 0.000437 | 0.003088 | 0.003741 | 737.2531 | 709.1408 | 515.752 | 451.6157 | 13445 | cyclin dependent kinase 2 associated protein 1 [Source:MGI Symbol;Acc:MGI:1202069] |
| ENSMUSG00000005413 | Hmox1 | 0.580099 | 6582.54 | 0.077246 | -7.50977 | 5.92E-14 | 2.26E-12 | 2.74E-12 | 8109.784 | 7666.785 | 5163.006 | 5390.584 | 15368 | heme oxygenase 1 [Source:MGI Symbol;Acc:MGI:96163] |
| ENSMUSG00000024896 | Minpp1 | 0.581543 | 547.0471 | 0.178061 | -3.26598 | 0.001091 | 0.00676 | 0.008188 | 676.9612 | 634.3908 | 400.5308 | 476.3054 | 17330 | multiple inositol polyphosphate histidine phosphatase 1 [Source:MGI Symbol;Acc:MGI:1336159] |
| ENSMUSG00000022125 | Cln5 | 0.581609 | 2097.3 | 0.098766 | -5.88875 | 3.89E-09 | 8.05E-08 | 9.75E-08 | 2561.875 | 2466.748 | 1640.53 | 1720.049 | 211286 | ceroid-lipofuscinosis, neuronal 5 [Source:MGI Symbol;Acc:MGI:2442253] |
| ENSMUSG00000034681 | Rnps1 | 0.58215 | 972.0569 | 0.13373 | -4.35316 | 1.34E-05 | 0.000141 | 0.00017 | 1188.913 | 1141.904 | 741.622 | 815.7887 | 19826 | RNA binding protein with serine rich domain 1 [Source:MGI Symbol;Acc:MGI:97960] |
| ENSMUSG00000027642 | Rpn2 | 0.582223 | 4630.417 | 0.090885 | -6.40613 | 1.49E-10 | 3.8E-09 | 4.61E-09 | 5834.56 | 5270.854 | 3810.529 | 3605.724 | 20014 | ribophorin II [Source:MGI Symbol;Acc:MGI:98085] |
| ENSMUSG00000024498 | Tcerg1 | 0.582951 | 650.53 | 0.193871 | -3.00689 | 0.002639 | 0.014013 | 0.016969 | 895.9159 | 664.8809 | 541.3567 | 499.9664 | 56070 | transcription elongation regulator 1 (CA150) [Source:MGI Symbol;Acc:MGI:1926421] |
| ENSMUSG00000035293 | G2e3 | 0.583938 | 799.0977 | 0.149475 | -3.90658 | 9.36E-05 | 0.000784 | 0.000949 | 1019.673 | 897.9827 | 658.4068 | 620.3286 | 217558 | G2/M-phase specific E3 ubiquitin ligase [Source:MGI Symbol;Acc:MGI:2444298] |
| ENSMUSG00000002210 | Smg9 | 0.58405 | 326.848 | 0.207546 | -2.81408 | 0.004892 | 0.023562 | 0.028479 | 395.5992 | 388.5029 | 249.6459 | 273.6441 | 71997 | SMG9 nonsense mediated mRNA decay factor [Source:MGI Symbol;Acc:MGI:1919247] |
| ENSMUSG00000056962 | Jmjd6 | 0.5843 | 519.7346 | 0.174441 | -3.34955 | 0.000809 | 0.005238 | 0.006346 | 662.1527 | 585.2133 | 425.221 | 406.3513 | 107817 | jumonji domain containing 6 [Source:MGI Symbol;Acc:MGI:1858910] |
| ENSMUSG00000040549 | Ckap5 | 0.584389 | 3830.675 | 0.083724 | -6.97992 | 2.95E-12 | 9.41E-11 | 1.14E-10 | 4724.978 | 4467.292 | 3044.217 | 3086.212 | 75786 | cytoskeleton associated protein 5 [Source:MGI Symbol;Acc:MGI:1923036] |
| ENSMUSG00000039183 | Nubp2 | 0.584563 | 502.2595 | 0.168334 | -3.47264 | 0.000515 | 0.003562 | 0.004315 | 594.4566 | 610.7856 | 400.5308 | 403.265 | 26426 | nucleotide binding protein 2 [Source:MGI Symbol;Acc:MGI:1347072] |
| ENSMUSG00000022498 | Txndc11 | 0.584852 | 593.5319 | 0.156237 | -3.74337 | 0.000182 | 0.001409 | 0.001707 | 719.2713 | 705.2066 | 476.4304 | 473.2192 | 106200 | thioredoxin domain containing 11 [Source:MGI Symbol;Acc:MGI:1923620] |
| ENSMUSG00000005103 | Wdr1 | 0.585149 | 10860.43 | 0.071981 | -8.12921 | 4.32E-16 | 2.07E-14 | 2.51E-14 | 13433.45 | 12633.72 | 8822.65 | 8551.894 | 22388 | WD repeat domain 1 [Source:MGI Symbol;Acc:MGI:1337100] |
| ENSMUSG00000090213 | Peds1 | 0.585291 | 1321.976 | 0.114189 | -5.12563 | 2.97E-07 | 4.37E-06 | 5.29E-06 | 1612.014 | 1560.897 | 1037.905 | 1077.088 | 407243 | plasmanylethanolamine desaturase 1 [Source:MGI Symbol;Acc:MGI:2142624] |
| ENSMUSG00000037280 | Galnt6 | 0.585523 | 792.832 | 0.140488 | -4.16777 | 3.08E-05 | 0.000296 | 0.000358 | 924.4752 | 978.6339 | 643.7755 | 624.4436 | 207839 | polypeptide N-acetylgalactosaminyltransferase 6 [Source:MGI Symbol;Acc:MGI:1891640] |
| ENSMUSG00000028403 | Zdhhc21 | 0.585615 | 913.145 | 0.173076 | -3.38357 | 0.000715 | 0.004713 | 0.005709 | 1242.858 | 949.1274 | 693.156 | 767.4381 | 68268 | zinc finger, DHHC domain containing 21 [Source:MGI Symbol;Acc:MGI:1915518] |
| ENSMUSG00000006494 | Pdk1 | 0.586064 | 1394.321 | 0.142747 | -4.10561 | 4.03E-05 | 0.000379 | 0.000459 | 1859.528 | 1488.114 | 1100.088 | 1129.554 | 228026 | pyruvate dehydrogenase kinase, isoenzyme 1 [Source:MGI Symbol;Acc:MGI:1926119] |
| ENSMUSG00000033096 | Apmap | 0.586124 | 555.6166 | 0.168466 | -3.47918 | 0.000503 | 0.003484 | 0.004221 | 706.5783 | 627.506 | 448.0824 | 440.2996 | 71881 | adipocyte plasma membrane associated protein [Source:MGI Symbol;Acc:MGI:1919131] |
| ENSMUSG00000035443 | Thyn1 | 0.58627 | 350.0457 | 0.205638 | -2.85098 | 0.004358 | 0.021408 | 0.025883 | 424.1585 | 416.0424 | 260.6193 | 299.3626 | 77862 | thymocyte nuclear protein 1 [Source:MGI Symbol;Acc:MGI:1925112] |
| ENSMUSG00000031467 | Agpat5 | 0.586533 | 886.8057 | 0.144528 | -4.05827 | 4.94E-05 | 0.000449 | 0.000544 | 1133.91 | 995.3543 | 688.5837 | 729.3748 | 52123 | 1-acylglycerol-3-phosphate O-acyltransferase 5 [Source:MGI Symbol;Acc:MGI:1196345] |
| ENSMUSG00000041836 | Ptpre | 0.587161 | 852.9887 | 0.138111 | -4.25137 | 2.12E-05 | 0.000211 | 0.000255 | 1064.098 | 984.5352 | 689.4982 | 673.823 | 19267 | protein tyrosine phosphatase receptor type E [Source:MGI Symbol;Acc:MGI:97813] |
| ENSMUSG00000026790 | Odf2 | 0.588007 | 848.571 | 0.138461 | -4.24674 | 2.17E-05 | 0.000215 | 0.00026 | 1056.694 | 981.5846 | 665.7224 | 690.2828 | 18286 | outer dense fiber of sperm tails 2 [Source:MGI Symbol;Acc:MGI:1098824] |
| ENSMUSG00000075232 | Amd1 | 0.588074 | 1534.487 | 0.143872 | -4.08749 | 4.36E-05 | 0.000405 | 0.00049 | 2024.537 | 1661.219 | 1138.495 | 1313.698 | 11702 | S-adenosylmethionine decarboxylase 1 [Source:MGI Symbol;Acc:MGI:88004] |
| ENSMUSG00000034659 | Tmem109 | 0.588277 | 469.5443 | 0.174158 | -3.37784 | 0.000731 | 0.004804 | 0.005819 | 571.186 | 556.6903 | 367.6104 | 382.6903 | 68539 | transmembrane protein 109 [Source:MGI Symbol;Acc:MGI:1915789] |
| ENSMUSG00000027889 | Ampd2 | 0.589057 | 437.3923 | 0.180864 | -3.25691 | 0.001126 | 0.006936 | 0.008403 | 541.569 | 509.4798 | 353.8936 | 344.627 | 109674 | adenosine monophosphate deaminase 2 [Source:MGI Symbol;Acc:MGI:88016] |
| ENSMUSG00000042699 | Dhx9 | 0.589067 | 3069.868 | 0.133432 | -4.41472 | 1.01E-05 | 0.000109 | 0.000132 | 4140.041 | 3236.869 | 2579.674 | 2322.889 | 13211 | DExH-box helicase 9 [Source:MGI Symbol;Acc:MGI:108177] |
| ENSMUSG00000062933 | Gm10123 | 0.591253 | 700.1645 | 0.157079 | -3.76405 | 0.000167 | 0.001311 | 0.001589 | 868.4143 | 815.3644 | 595.3094 | 521.5698 | | predicted pseudogene 10123 [Source:MGI Symbol;Acc:MGI:3704342] |
| ENSMUSG00000031826 | Usp10 | 0.591254 | 359.4271 | 0.199353 | -2.96586 | 0.003018 | 0.015717 | 0.019032 | 441.0825 | 422.9272 | 274.3361 | 299.3626 | 22224 | ubiquitin specific peptidase 10 [Source:MGI Symbol;Acc:MGI:894652] |
| ENSMUSG00000079470 | Utp14b | 0.591865 | 334.2002 | 0.202745 | -2.91925 | 0.003509 | 0.017859 | 0.021613 | 408.2922 | 395.3878 | 269.7639 | 263.3568 | 195434 | UTP14B small subunit processome component [Source:MGI Symbol;Acc:MGI:2445092] |
| ENSMUSG00000032120 | C2cd2l | 0.591901 | 540.1947 | 0.164707 | -3.59366 | 0.000326 | 0.002379 | 0.002882 | 669.557 | 629.4731 | 430.7077 | 431.041 | 71764 | C2 calcium-dependent domain containing 2-like [Source:MGI Symbol;Acc:MGI:1919014] |
| ENSMUSG00000021606 | Ndufs6 | 0.592043 | 390.4987 | 0.207828 | -2.84872 | 0.00439 | 0.021519 | 0.026017 | 518.2984 | 420.9601 | 318.2299 | 304.5063 | 407785 | NADH:ubiquinone oxidoreductase core subunit S6 [Source:MGI Symbol;Acc:MGI:107932] |
| ENSMUSG00000035439 | Haus8 | 0.592808 | 700.5282 | 0.152943 | -3.876 | 0.000106 | 0.000877 | 0.001063 | 884.2806 | 800.6111 | 543.1856 | 574.0354 | 76478 | 4HAUS augmin-like complex, subunit 8 [Source:MGI Symbol;Acc:MGI:1923728] |
| ENSMUSG00000043702 | Pde12 | 0.593558 | 482.269 | 0.184502 | -3.21709 | 0.001295 | 0.007795 | 0.009443 | 626.1891 | 534.0686 | 375.8405 | 392.9777 | 211948 | phosphodiesterase 12 [Source:MGI Symbol;Acc:MGI:2443226] |
| ENSMUSG00000027774 | Gfm1 | 0.596278 | 691.851 | 0.148762 | -4.00827 | 6.12E-05 | 0.000542 | 0.000657 | 846.2015 | 819.2986 | 534.041 | 567.863 | 28030 | G elongation factor, mitochondrial 1 [Source:MGI Symbol;Acc:MGI:107339] |
| ENSMUSG00000035367 | Rmi1 | 0.596316 | 616.484 | 0.169606 | -3.5159 | 0.000438 | 0.003092 | 0.003746 | 803.8915 | 680.6178 | 502.0351 | 479.3916 | 74386 | RecQ mediated genome instability 1 [Source:MGI Symbol;Acc:MGI:1921636] |
| ENSMUSG00000001270 | Ckb | 0.596464 | 5302.944 | 0.084778 | -7.03561 | 1.98E-12 | 6.4E-11 | 7.75E-11 | 6083.131 | 6684.217 | 4251.296 | 4193.133 | 12709 | creatine kinase, brain [Source:MGI Symbol;Acc:MGI:88407] |
| ENSMUSG00000032381 | Ciao2a | 0.596768 | 1252.764 | 0.117658 | -5.07206 | 3.94E-07 | 5.63E-06 | 6.83E-06 | 1545.376 | 1471.393 | 1015.958 | 978.3292 | 68250 | cytosolic iron-sulfur assembly component 2A [Source:MGI Symbol;Acc:MGI:1915500] |
| ENSMUSG00000030079 | Ruvbl1 | 0.596893 | 413.9425 | 0.20036 | -2.9791 | 0.002891 | 0.015126 | 0.018317 | 542.6267 | 454.4009 | 342.9202 | 315.8224 | 56505 | RuvB-like AAA ATPase 1 [Source:MGI Symbol;Acc:MGI:1928760] |
| ENSMUSG00000038187 | Btbd10 | 0.597686 | 474.8739 | 0.175908 | -3.39771 | 0.00068 | 0.004501 | 0.005453 | 584.9368 | 558.6574 | 363.9526 | 391.9489 | 68815 | BTB domain containing 10 [Source:MGI Symbol;Acc:MGI:1916065] |
| ENSMUSG00000058833 | Rex1bd | 0.598445 | 365.9074 | 0.195041 | -3.0683 | 0.002153 | 0.011857 | 0.014364 | 451.6601 | 429.8121 | 288.9674 | 293.1901 | 66462 | required for excision 1-B domain containing [Source:MGI Symbol;Acc:MGI:1913712] |
| ENSMUSG00000027175 | Tcp11l1 | 0.598876 | 458.344 | 0.201329 | -2.97461 | 0.002934 | 0.015323 | 0.018555 | 605.0341 | 499.6443 | 392.3007 | 336.3971 | 320554 | t-complex 11 like 1 [Source:MGI Symbol;Acc:MGI:2444263] |
| ENSMUSG00000031256 | Cstf2 | 0.598915 | 819.9128 | 0.158324 | -3.78283 | 0.000155 | 0.001226 | 0.001486 | 1070.445 | 904.8676 | 618.1708 | 686.1678 | 108062 | cleavage stimulation factor, 3' pre-RNA subunit 2 [Source:MGI Symbol;Acc:MGI:1343054] |
| ENSMUSG00000031976 | Urb2 | 0.599266 | 417.8064 | 0.188244 | -3.18346 | 0.001455 | 0.008586 | 0.010401 | 528.876 | 478.0061 | 339.2624 | 325.081 | 382038 | URB2 ribosome biogenesis 2 homolog (S. cerevisiae) [Source:MGI Symbol;Acc:MGI:2681124] |
| ENSMUSG00000030224 | Strap | 0.599271 | 1633.103 | 0.127583 | -4.6971 | 2.64E-06 | 3.17E-05 | 3.84E-05 | 2080.598 | 1853.995 | 1207.993 | 1389.824 | 20901 | serine/threonine kinase receptor associated protein [Source:MGI Symbol;Acc:MGI:1329037] |
| ENSMUSG00000028967 | Errfi1 | 0.599296 | 372.5377 | 0.216325 | -2.77035 | 0.0056 | 0.026377 | 0.031864 | 502.4322 | 395.3878 | 298.1119 | 294.2189 | 74155 | ERBB receptor feedback inhibitor 1 [Source:MGI Symbol;Acc:MGI:1921405] |
| ENSMUSG00000022136 | Dnajc3 | 0.599925 | 3554.821 | 0.091768 | -6.53744 | 6.26E-11 | 1.67E-09 | 2.02E-09 | 4398.132 | 4168.292 | 2708.612 | 2944.246 | 100037258 | DnaJ heat shock protein family (Hsp40) member C3 [Source:MGI Symbol;Acc:MGI:107373] |
| ENSMUSG00000028961 | Pgd | 0.599971 | 2436.363 | 0.105582 | -5.68253 | 1.33E-08 | 2.54E-07 | 3.08E-07 | 3051.614 | 2820.826 | 2034.66 | 1838.354 | 110208 | phosphogluconate dehydrogenase [Source:MGI Symbol;Acc:MGI:97553] |
| ENSMUSG00000056076 | Eif3b | 0.6008 | 1386.643 | 0.125682 | -4.78033 | 1.75E-06 | 2.18E-05 | 2.64E-05 | 1724.136 | 1617.943 | 1029.675 | 1174.818 | 27979 | eukaryotic translation initiation factor 3, subunit B [Source:MGI Symbol;Acc:MGI:106478] |
| ENSMUSG00000002728 | Naa20 | 0.601098 | 1302.846 | 0.126752 | -4.74231 | 2.11E-06 | 2.6E-05 | 3.14E-05 | 1675.479 | 1465.492 | 1024.188 | 1046.226 | 67877 | N(alpha)-acetyltransferase 20, NatB catalytic subunit [Source:MGI Symbol;Acc:MGI:1915127] |
| ENSMUSG00000030047 | Arhgap25 | 0.601517 | 1973.97 | 0.116877 | -5.1466 | 2.65E-07 | 3.95E-06 | 4.79E-06 | 2436.003 | 2324.133 | 1686.253 | 1449.491 | 232201 | Rho GTPase activating protein 25 [Source:MGI Symbol;Acc:MGI:2443687] |
| ENSMUSG00000029484 | Anxa3 | 0.601801 | 7876.38 | 0.075197 | -8.00296 | 1.21E-15 | 5.6E-14 | 6.78E-14 | 9716.509 | 9275.876 | 6424.038 | 6089.096 | 11745 | annexin A3 [Source:MGI Symbol;Acc:MGI:1201378] |
| ENSMUSG00000025393 | Atp5f1b | 0.602236 | 10467.59 | 0.081123 | -7.4237 | 1.14E-13 | 4.26E-12 | 5.17E-12 | 13197.57 | 12044.57 | 8052.68 | 8575.555 | 11947 | ATP synthase F1 subunit beta [Source:MGI Symbol;Acc:MGI:107801] |
| ENSMUSG00000031960 | Aars1 | 0.602272 | 2143.068 | 0.116188 | -5.1836 | 2.18E-07 | 3.3E-06 | 4E-06 | 2751.213 | 2416.587 | 1618.583 | 1785.888 | 234734 | alanyl-tRNA synthetase 1 [Source:MGI Symbol;Acc:MGI:2384560] |
| ENSMUSG00000026245 | Farsb | 0.602367 | 1304.767 | 0.118067 | -5.10193 | 3.36E-07 | 4.88E-06 | 5.92E-06 | 1627.88 | 1518.604 | 1015.044 | 1057.542 | 23874 | phenylalanyl-tRNA synthetase, beta subunit [Source:MGI Symbol;Acc:MGI:1346035] |
| ENSMUSG00000026483 | Niban1 | 0.602556 | 748.394 | 0.155603 | -3.87239 | 0.000108 | 0.000888 | 0.001075 | 972.074 | 833.0683 | 598.9673 | 589.4665 | 63913 | niban apoptosis regulator 1 [Source:MGI Symbol;Acc:MGI:2137237] |
| ENSMUSG00000073792 | Alg6 | 0.602874 | 259.4281 | 0.228297 | -2.64074 | 0.008272 | 0.036474 | 0.044056 | 309.9213 | 315.7201 | 201.1798 | 210.8912 | 320438 | ALG6 alpha-1,3-glucosyltransferase [Source:MGI Symbol;Acc:MGI:2444031] |
| ENSMUSG00000029036 | Atad3a | 0.602922 | 337.5927 | 0.21211 | -2.8425 | 0.004476 | 0.02188 | 0.026454 | 431.5628 | 382.6016 | 253.3037 | 282.9028 | 108888 | ATPase family, AAA domain containing 3A [Source:MGI Symbol;Acc:MGI:1919214] |
| ENSMUSG00000115338 | Pnp | 0.603113 | 3405.261 | 0.095846 | -6.29251 | 3.12E-10 | 7.52E-09 | 9.11E-09 | 4339.956 | 3874.21 | 2717.757 | 2689.119 | 18950 | purine-nucleoside phosphorylase [Source:MGI Symbol;Acc:MGI:97365] |
| ENSMUSG00000106918 | Mrpl33 | 0.603728 | 458.4967 | 0.177244 | -3.4062 | 0.000659 | 0.004389 | 0.005316 | 569.0705 | 537.0192 | 357.5514 | 370.3454 | 66845 | mitochondrial ribosomal protein L33 [Source:MGI Symbol;Acc:MGI:2137225] |
| ENSMUSG00000041707 | Tmem273 | 0.603763 | 1421.047 | 0.110242 | -5.47672 | 4.33E-08 | 7.62E-07 | 9.23E-07 | 1722.02 | 1706.462 | 1146.725 | 1108.979 | 69069 | transmembrane protein 273 [Source:MGI Symbol;Acc:MGI:1916319] |
| ENSMUSG00000031578 | Mak16 | 0.604139 | 590.7768 | 0.173963 | -3.4728 | 0.000515 | 0.003562 | 0.004315 | 775.3322 | 650.1277 | 457.2269 | 480.4203 | 67920 | MAK16 homolog [Source:MGI Symbol;Acc:MGI:1915170] |
| ENSMUSG00000029580 | Actb | 0.604308 | 140166.1 | 0.083121 | -7.27023 | 3.59E-13 | 1.26E-11 | 1.53E-11 | 181489.1 | 156712.2 | 111877.9 | 110585.2 | 11461 | actin, beta [Source:MGI Symbol;Acc:MGI:87904] |
| ENSMUSG00000025130 | P4hb | 0.604879 | 9544.432 | 0.072901 | -8.29726 | 1.07E-16 | 5.38E-15 | 6.52E-15 | 11899.71 | 11133.81 | 7621.058 | 7523.156 | 18453 | prolyl 4-hydroxylase, beta polypeptide [Source:MGI Symbol;Acc:MGI:97464] |
| ENSMUSG00000017999 | Ddx27 | 0.605233 | 554.6908 | 0.176622 | -3.42671 | 0.000611 | 0.004117 | 0.004987 | 715.0403 | 623.5718 | 415.162 | 464.9893 | 228889 | DEAD box helicase 27 [Source:MGI Symbol;Acc:MGI:2385884] |
| ENSMUSG00000004446 | Bid | 0.605497 | 754.8757 | 0.148856 | -4.06766 | 4.75E-05 | 0.000435 | 0.000527 | 927.6484 | 894.0485 | 565.1325 | 632.6735 | 12122 | BH3 interacting domain death agonist [Source:MGI Symbol;Acc:MGI:108093] |
| ENSMUSG00000051615 | Rap2a | 0.605592 | 5659.714 | 0.09184 | -6.59396 | 4.28E-11 | 1.17E-09 | 1.41E-09 | 7164.154 | 6496.359 | 4292.446 | 4685.899 | 76108 | RAS related protein 2a [Source:MGI Symbol;Acc:MGI:97855] |
| ENSMUSG00000063412 | Erhrt-ps | 0.606519 | 287.4214 | 0.226251 | -2.68074 | 0.007346 | 0.033023 | 0.039897 | 372.3287 | 321.6214 | 222.2123 | 233.5234 | | ERH mRNA splicing and mitosis factor retrotransposed, pseudogene [Source:MGI Symbol;Acc:MGI:3704308] |
| ENSMUSG00000042406 | Atf4 | 0.606741 | 2250.323 | 0.100257 | -6.05184 | 1.43E-09 | 3.16E-08 | 3.82E-08 | 2804.1 | 2629.034 | 1732.89 | 1835.267 | 11911 | activating transcription factor 4 [Source:MGI Symbol;Acc:MGI:88096] |
| ENSMUSG00000029253 | Cenpc1 | 0.606895 | 504.3945 | 0.172175 | -3.52488 | 0.000424 | 0.003002 | 0.003636 | 634.6511 | 583.2462 | 392.3007 | 407.38 | 12617 | centromere protein C1 [Source:MGI Symbol;Acc:MGI:99700] |
| ENSMUSG00000009207 | Lnpk | 0.607556 | 550.6827 | 0.197625 | -3.07429 | 0.00211 | 0.011643 | 0.014104 | 727.7333 | 601.9336 | 389.5573 | 483.5066 | 69605 | lunapark, ER junction formation factor [Source:MGI Symbol;Acc:MGI:1918115] |
| ENSMUSG00000025757 | Hspa4l | 0.6077 | 683.5508 | 0.215288 | -2.82273 | 0.004762 | 0.023053 | 0.027872 | 660.0372 | 990.4366 | 523.0676 | 560.6619 | 18415 | heat shock protein 4 like [Source:MGI Symbol;Acc:MGI:107422] |
| ENSMUSG00000108414 | Snhg1 | 0.607776 | 434.0484 | 0.213804 | -2.84267 | 0.004474 | 0.021877 | 0.02645 | 584.9368 | 463.2528 | 314.5721 | 373.4317 | | small nucleolar RNA host gene 1 [Source:MGI Symbol;Acc:MGI:3763743] |
| ENSMUSG00000021930 | Spryd7 | 0.608542 | 404.1404 | 0.190739 | -3.19045 | 0.001421 | 0.008417 | 0.010196 | 514.0674 | 462.2693 | 318.2299 | 321.9948 | 66674 | SPRY domain containing 7 [Source:MGI Symbol;Acc:MGI:1913924] |
| ENSMUSG00000029174 | Tbc1d1 | 0.608765 | 1314.132 | 0.14183 | -4.29221 | 1.77E-05 | 0.000179 | 0.000217 | 1752.695 | 1422.216 | 1025.103 | 1056.513 | 57915 | TBC1 domain family, member 1 [Source:MGI Symbol;Acc:MGI:1889508] |
| ENSMUSG00000098188 | Sowahc | 0.609302 | 1283.638 | 0.143087 | -4.25827 | 2.06E-05 | 0.000205 | 0.000248 | 1701.923 | 1399.594 | 977.5511 | 1055.485 | 268301 | sosondowah ankyrin repeat domain family member C [Source:MGI Symbol;Acc:MGI:3606051] |
| ENSMUSG00000042644 | Itpr3 | 0.609821 | 508.4454 | 0.193786 | -3.14689 | 0.00165 | 0.009539 | 0.011556 | 597.6298 | 631.4402 | 450.8257 | 353.8856 | 16440 | inositol 1,4,5-triphosphate receptor 3 [Source:MGI Symbol;Acc:MGI:96624] |
| ENSMUSG00000031924 | Cyb5b | 0.609988 | 1933.684 | 0.104246 | -5.85143 | 4.87E-09 | 9.91E-08 | 1.2E-07 | 2428.598 | 2244.465 | 1513.421 | 1548.25 | 66427 | cytochrome b5 type B [Source:MGI Symbol;Acc:MGI:1913677] |
| ENSMUSG00000021670 | Hmgcr | 0.610291 | 1046.496 | 0.135424 | -4.50653 | 6.59E-06 | 7.35E-05 | 8.9E-05 | 1344.403 | 1185.18 | 846.7842 | 809.6163 | 15357 | 3-hydroxy-3-methylglutaryl-Coenzyme A reductase [Source:MGI Symbol;Acc:MGI:96159] |
| ENSMUSG00000018379 | Srsf1 | 0.610352 | 4201.889 | 0.093158 | -6.55181 | 5.68E-11 | 1.52E-09 | 1.84E-09 | 5378.668 | 4777.111 | 3317.638 | 3334.138 | 110809 | serine and arginine-rich splicing factor 1 [Source:MGI Symbol;Acc:MGI:98283] |
| ENSMUSG00000026827 | Gpd2 | 0.611873 | 344.2157 | 0.203883 | -3.0011 | 0.00269 | 0.01424 | 0.017244 | 411.4655 | 420.9601 | 286.224 | 258.2131 | 14571 | glycerol phosphate dehydrogenase 2, mitochondrial [Source:MGI Symbol;Acc:MGI:99778] |
| ENSMUSG00000028609 | Magoh | 0.611908 | 364.2045 | 0.228663 | -2.67603 | 0.00745 | 0.033412 | 0.040366 | 478.1039 | 402.2726 | 252.3893 | 324.0523 | 17149 | mago homolog, exon junction complex core component [Source:MGI Symbol;Acc:MGI:1330312] |
| ENSMUSG00000016319 | Slc25a5 | 0.612541 | 5259.898 | 0.075734 | -8.08802 | 6.06E-16 | 2.87E-14 | 3.48E-14 | 6414.208 | 6305.55 | 4087.609 | 4232.225 | 11740 | solute carrier family 25 (mitochondrial carrier, adenine nucleotide translocator), member 5 [Source:MGI Symbol;Acc:MGI:1353496] |
| ENSMUSG00000035021 | Baz1a | 0.612831 | 1215.474 | 0.142218 | -4.30908 | 1.64E-05 | 0.000168 | 0.000204 | 1610.956 | 1328.778 | 934.5718 | 987.5879 | 217578 | bromodomain adjacent to zinc finger domain 1A [Source:MGI Symbol;Acc:MGI:1309478] |
| ENSMUSG00000041319 | Thoc6 | 0.613176 | 321.9059 | 0.218902 | -2.80115 | 0.005092 | 0.024426 | 0.029515 | 418.8698 | 359.9799 | 267.0205 | 241.7533 | 386612 | THO complex 6 [Source:MGI Symbol;Acc:MGI:2677480] |
| ENSMUSG00000063445 | Nmral1 | 0.613819 | 280.5784 | 0.224071 | -2.73939 | 0.006155 | 0.028499 | 0.034428 | 324.7298 | 354.0786 | 229.5279 | 213.9774 | 67824 | NmrA-like family domain containing 1 [Source:MGI Symbol;Acc:MGI:1915074] |

| | | | | | | | | | | | | | | |
|---|---|---|---|---|---|---|---|---|---|---|---|---|---|---|
| ENSMUSG00000021595 | Nsun2 | 0.613997 | 1303.561 | 0.128706 | -4.77053 | 1.84E-06 | 2.28E-05 | 2.77E-05 | 1691.345 | 1462.541 | 1022.359 | 1037.996 | 28114 | NOL1/NOP2/Sun domain family member 2 [Source:MGI Symbol;Acc:MGI:107252] |
| ENSMUSG00000008892 | Vdac3 | 0.614113 | 1787.143 | 0.118259 | -5.19298 | 2.07E-07 | 3.15E-06 | 3.82E-06 | 2298.495 | 2025.133 | 1357.964 | 1466.979 | 22335 | voltage-dependent anion channel 3 [Source:MGI Symbol;Acc:MGI:106922] |
| ENSMUSG00000022295 | Atp6v1c1 | 0.61445 | 3521.869 | 0.092174 | -6.6662 | 2.63E-11 | 7.33E-10 | 8.88E-10 | 4396.017 | 4125.016 | 2675.692 | 2890.752 | 66335 | ATPase, H+ transporting, lysosomal V1 subunit C1 [Source:MGI Symbol;Acc:MGI:1913585] |
| ENSMUSG00000001082 | Mfsd10 | 0.615238 | 296.8226 | 0.214651 | -2.86623 | 0.004154 | 0.020597 | 0.024902 | 356.4624 | 361.947 | 240.5014 | 228.3797 | 68294 | major facilitator superfamily domain containing 10 [Source:MGI Symbol;Acc:MGI:1915544] |
| ENSMUSG00000020022 | Ndufa12 | 0.615255 | 576.5776 | 0.171543 | -3.58659 | 0.000335 | 0.002439 | 0.002954 | 676.9612 | 717.9927 | 420.6488 | 490.7077 | 66414 | NADH:ubiquinone oxidoreductase subunit A12 [Source:MGI Symbol;Acc:MGI:1913664] |
| ENSMUSG00000016477 | E2f3 | 0.615442 | 451.9484 | 0.177959 | -3.45834 | 0.000544 | 0.003728 | 0.004516 | 562.724 | 531.1179 | 353.8936 | 360.0581 | 13557 | E2F transcription factor 3 [Source:MGI Symbol;Acc:MGI:1096340] |
| ENSMUSG00000090553 | Snrpe | 0.61553 | 641.7623 | 0.166533 | -3.69615 | 0.000219 | 0.001668 | 0.00202 | 835.624 | 717.9927 | 525.8109 | 487.6215 | 20643 | small nuclear ribonucleoprotein E [Source:MGI Symbol;Acc:MGI:98346] |
| ENSMUSG00000039985 | Sinhcaf | 0.615664 | 383.4627 | 0.196915 | -3.12654 | 0.001769 | 0.010082 | 0.012214 | 484.4504 | 443.5818 | 288.9674 | 316.8511 | 56306 | SIN3-HDAC complex associated factor [Source:MGI Symbol;Acc:MGI:1929091] |
| ENSMUSG00000017774 | Myo1c | 0.615784 | 2475.499 | 0.118275 | -5.2064 | 1.93E-07 | 2.96E-06 | 3.58E-06 | 3263.165 | 2729.356 | 2006.312 | 1903.164 | 17913 | myosin IC [Source:MGI Symbol;Acc:MGI:106612] |
| ENSMUSG00000020321 | Mdh1 | 0.616981 | 2932.242 | 0.086167 | -7.16027 | 8.05E-13 | 2.7E-11 | 3.27E-11 | 3581.548 | 3518.164 | 2305.338 | 2323.918 | 17449 | malate dehydrogenase 1, NAD (soluble) [Source:MGI Symbol;Acc:MGI:97051] |
| ENSMUSG00000017428 | Psmd11 | 0.617018 | 792.9752 | 0.150271 | -4.10604 | 4.02E-05 | 0.000378 | 0.000458 | 1027.077 | 893.0649 | 619.0852 | 632.6735 | 69077 | proteasome (prosome, macropain) 26S subunit, non-ATPase, 11 [Source:MGI Symbol;Acc:MGI:1916327] |
| ENSMUSG00000046324 | Ermp1 | 0.617523 | 835.2035 | 0.173415 | -3.56095 | 0.00037 | 0.002657 | 0.003218 | 1093.715 | 928.4728 | 586.1649 | 732.461 | 226090 | endoplasmic reticulum metallopeptidase 1 [Source:MGI Symbol;Acc:MGI:106250] |
| ENSMUSG00000031657 | Heatr3 | 0.617547 | 566.0459 | 0.16284 | -3.79235 | 0.000149 | 0.001187 | 0.001438 | 710.8093 | 659.9632 | 440.7667 | 452.6444 | 234549 | HEAT repeat containing 3 [Source:MGI Symbol;Acc:MGI:2444491] |
| ENSMUSG00000068039 | Tcp1 | 0.618656 | 3983.765 | 0.110863 | -5.58039 | 2.4E-08 | 4.41E-07 | 5.34E-07 | 5286.644 | 4364.019 | 3153.951 | 3130.448 | 21454 | t-complex protein 1 [Source:MGI Symbol;Acc:MGI:98535] |
| ENSMUSG00000066735 | Vkorc1l1 | 0.61875 | 423.669 | 0.197894 | -3.12667 | 0.001768 | 0.010082 | 0.012213 | 552.1465 | 474.0719 | 314.5721 | 353.8856 | 69568 | vitamin K epoxide reductase complex, subunit 1-like 1 [Source:MGI Symbol;Acc:MGI:1916818] |
| ENSMUSG00000024683 | Mrpl16 | 0.619003 | 330.4668 | 0.208568 | -2.96787 | 0.002999 | 0.01563 | 0.018928 | 394.5415 | 406.2068 | 275.2506 | 245.8682 | 94063 | mitochondrial ribosomal protein L16 [Source:MGI Symbol;Acc:MGI:2137219] |
| ENSMUSG00000038388 | Pals2 | 0.619676 | 665.9319 | 0.157004 | -3.94687 | 7.92E-05 | 0.000679 | 0.000822 | 816.5845 | 796.6769 | 492.8906 | 557.5756 | 56524 | protein associated with LIN7 2, MAGUK family member [Source:MGI Symbol;Acc:MGI:1927340] |
| ENSMUSG00000007589 | Tinf2 | 0.619945 | 338.6873 | 0.223495 | -2.77387 | 0.005539 | 0.026158 | 0.031599 | 454.8333 | 365.8812 | 254.2182 | 279.8166 | 28113 | Terf1 (TRF1)-interacting nuclear factor 2 [Source:MGI Symbol;Acc:MGI:107246] |
| ENSMUSG00000029472 | Anapc5 | 0.620702 | 3751.503 | 0.091557 | -6.77942 | 1.21E-11 | 3.53E-10 | 4.28E-10 | 4763.057 | 4329.595 | 2904.305 | 3009.057 | 59008 | anaphase-promoting complex subunit 5 [Source:MGI Symbol;Acc:MGI:1929722] |
| ENSMUSG00000039886 | Tmem120a | 0.621036 | 380.3695 | 0.190979 | -3.25185 | 0.001147 | 0.00703 | 0.008516 | 453.7756 | 468.1706 | 298.1119 | 301.42 | 215210 | transmembrane protein 120A [Source:MGI Symbol;Acc:MGI:2686991] |
| ENSMUSG00000030760 | Acer3 | 0.621743 | 875.608 | 0.133184 | -4.6683 | 3.04E-06 | 3.61E-05 | 4.38E-05 | 1072.56 | 1050.433 | 700.4716 | 678.9667 | 66190 | alkaline ceramidase 3 [Source:MGI Symbol;Acc:MGI:1913440] |
| ENSMUSG00000001440 | Kpnb1 | 0.622033 | 4927.927 | 0.102981 | -6.04029 | 1.54E-09 | 3.38E-08 | 4.09E-08 | 6467.095 | 5481.334 | 3791.326 | 3971.955 | 16211 | karyopherin subunit beta 1 [Source:MGI Symbol;Acc:MGI:107532] |
| ENSMUSG00000031422 | Morf4l2 | 0.622263 | 3177.843 | 0.098749 | -6.30146 | 2.95E-10 | 7.15E-09 | 8.67E-09 | 4039.555 | 3665.697 | 2409.586 | 2596.533 | 56397 | mortality factor 4 like 2 [Source:MGI Symbol;Acc:MGI:1927167] |
| ENSMUSG00000026750 | Psmb7 | 0.622796 | 1862.68 | 0.108845 | -5.72187 | 1.05E-08 | 2.05E-07 | 2.48E-07 | 2359.845 | 2157.912 | 1506.105 | 1426.859 | 19177 | proteasome (prosome, macropain) subunit, beta type 7 [Source:MGI Symbol;Acc:MGI:107637] |
| ENSMUSG00000022247 | Brix1 | 0.622826 | 274.7465 | 0.224359 | -2.77602 | 0.005503 | 0.026018 | 0.031429 | 341.6539 | 324.5721 | 208.4955 | 224.2647 | 67832 | BRX1, biogenesis of ribosomes [Source:MGI Symbol;Acc:MGI:1915082] |
| ENSMUSG00000027566 | Psma7 | 0.623703 | 1622.915 | 0.111478 | -5.59485 | 2.21E-08 | 4.1E-07 | 4.96E-07 | 2056.27 | 1880.551 | 1263.775 | 1291.065 | 26444 | proteasome subunit alpha 7 [Source:MGI Symbol;Acc:MGI:1347070] |
| ENSMUSG00000021831 | Ero1a | 0.624808 | 926.6247 | 0.161344 | -3.87253 | 0.000108 | 0.000888 | 0.001075 | 1248.147 | 1000.272 | 696.8138 | 761.2656 | 50527 | endoplasmic reticulum oxidoreductase 1 alpha [Source:MGI Symbol;Acc:MGI:1354385] |
| ENSMUSG00000006057 | Atp5mc1 | 0.625726 | 1289.08 | 0.119832 | -5.22171 | 1.77E-07 | 2.74E-06 | 3.32E-06 | 1608.841 | 1519.587 | 977.5511 | 1050.341 | 11951 | ATP synthase membrane subunit c locus 1 [Source:MGI Symbol;Acc:MGI:107653] |
| ENSMUSG00000024535 | Snx24 | 0.626251 | 1017.71 | 0.127487 | -4.91226 | 9E-07 | 1.2E-05 | 1.45E-05 | 1239.685 | 1230.423 | 773.6279 | 827.1048 | 69226 | sorting nexing 24 [Source:MGI Symbol;Acc:MGI:1916476] |
| ENSMUSG00000031007 | Atp6ap2 | 0.626639 | 5203.424 | 0.08737 | -7.17225 | 7.38E-13 | 2.51E-11 | 3.04E-11 | 6561.235 | 6070.481 | 3939.467 | 4242.513 | 70495 | ATPase, H+ transporting, lysosomal accessory protein 2 [Source:MGI Symbol;Acc:MGI:1917745] |
| ENSMUSG00000033732 | Sf3b3 | 0.6281 | 1970.941 | 0.120014 | -5.23357 | 1.66E-07 | 2.59E-06 | 3.13E-06 | 2594.665 | 2192.337 | 1545.427 | 1551.336 | 101943 | splicing factor 3b, subunit 3 [Source:MGI Symbol;Acc:MGI:1289341] |
| ENSMUSG00000042208 | Sanbr | 0.628133 | 814.3901 | 0.139936 | -4.48872 | 7.17E-06 | 7.95E-05 | 9.63E-05 | 968.9008 | 1009.124 | 660.2357 | 619.2999 | 71675 | SANT and BTB domain regulator of CSR [Source:MGI Symbol;Acc:MGI:1918925] |
| ENSMUSG00000062981 | Mrpl42 | 0.628533 | 584.4368 | 0.172968 | -3.63381 | 0.000279 | 0.002073 | 0.002511 | 765.8124 | 654.0619 | 475.516 | 442.3571 | 67270 | mitochondrial ribosomal protein L42 [Source:MGI Symbol;Acc:MGI:1333774] |
| ENSMUSG00000030245 | Golt1b | 0.628564 | 872.7894 | 0.164296 | -3.8258 | 0.00013 | 0.001054 | 0.001277 | 1167.758 | 952.078 | 641.9466 | 729.3748 | 66964 | golgi transport 1B [Source:MGI Symbol;Acc:MGI:1914214] |
| ENSMUSG00000042901 | Aida | 0.628921 | 781.8319 | 0.157229 | -4.00003 | 6.33E-05 | 0.000559 | 0.000677 | 1021.788 | 877.3281 | 585.2504 | 642.9608 | 108909 | axin interactor, dorsalization associated [Source:MGI Symbol;Acc:MGI:1919737] |
| ENSMUSG00000026421 | Csrp1 | 0.629413 | 966.9873 | 0.129433 | -4.86286 | 1.16E-06 | 1.51E-05 | 1.83E-05 | 1182.567 | 1166.492 | 737.0498 | 781.8404 | 13007 | cysteine and glycine-rich protein 1 [Source:MGI Symbol;Acc:MGI:88549] |
| ENSMUSG00000027968 | Larp7 | 0.629487 | 413.6367 | 0.18964 | -3.31937 | 0.000902 | 0.005753 | 0.006969 | 530.9915 | 474.0719 | 326.46 | 323.0235 | 28036 | La ribonucleoprotein 7, transcriptional regulator [Source:MGI Symbol;Acc:MGI:107634] |
| ENSMUSG00000067995 | Gtf2f2 | 0.63043 | 341.2733 | 0.216211 | -2.91581 | 0.003548 | 0.018021 | 0.021809 | 451.6601 | 377.6839 | 256.9615 | 278.7878 | 68705 | general transcription factor IIF, polypeptide 2 [Source:MGI Symbol;Acc:MGI:1915955] |
| ENSMUSG00000040274 | Cdk6 | 0.630837 | 1063.452 | 0.165582 | -3.80982 | 0.000139 | 0.001115 | 0.00135 | 1463.929 | 1121.249 | 872.3889 | 796.2427 | 12571 | cyclin dependent kinase 6 [Source:MGI Symbol;Acc:MGI:1277162] |
| ENSMUSG00000045983 | Eif4g1 | 0.63092 | 5481.719 | 0.095424 | -6.61172 | 3.8E-11 | 1.04E-09 | 1.26E-09 | 7141.941 | 6181.622 | 4264.098 | 4339.214 | 208643 | eukaryotic translation initiation factor 4, gamma 1 [Source:MGI Symbol;Acc:MGI:2384784] |
| ENSMUSG00000031604 | Msmo1 | 0.631186 | 793.5084 | 0.144203 | -4.37705 | 1.2E-05 | 0.000127 | 0.000154 | 1010.153 | 918.6373 | 613.5985 | 631.6447 | 66234 | methylsterol monoxygenase 1 [Source:MGI Symbol;Acc:MGI:1913484] |
| ENSMUSG00000038489 | Polr2l | 0.631461 | 335.2592 | 0.202753 | -3.11443 | 0.001843 | 0.010435 | 0.012641 | 399.8302 | 415.0588 | 259.7049 | 266.443 | 66491 | polymerase (RNA) II (DNA directed) polypeptide L [Source:MGI Symbol;Acc:MGI:1913741] |
| ENSMUSG00000030357 | Fkbp4 | 0.632842 | 1172.601 | 0.139992 | -4.52055 | 6.17E-06 | 6.92E-05 | 8.39E-05 | 1544.318 | 1307.14 | 884.2768 | 954.6683 | 14228 | FK506 binding protein 4 [Source:MGI Symbol;Acc:MGI:95543] |
| ENSMUSG00000029326 | Enoph1 | 0.633101 | 406.4745 | 0.201771 | -3.13772 | 0.001703 | 0.009776 | 0.011842 | 532.0492 | 456.368 | 299.0264 | 338.4546 | 67870 | enolase-phosphatase 1 [Source:MGI Symbol;Acc:MGI:1915120] |
| ENSMUSG00000022177 | Haus4 | 0.633164 | 269.3253 | 0.23344 | -2.71232 | 0.006681 | 0.030515 | 0.036865 | 349.0581 | 305.8846 | 201.1798 | 221.1785 | 219072 | HAUS augmin-like complex, subunit 4 [Source:MGI Symbol;Acc:MGI:1261794] |
| ENSMUSG00000021377 | Dek | 0.633555 | 2141.502 | 0.136946 | -4.62632 | 3.72E-06 | 4.35E-05 | 5.27E-05 | 2851.699 | 2356.59 | 1539.94 | 1817.779 | 110052 | DEK proto-oncogene [Source:MGI Symbol;Acc:MGI:1926209] |
| ENSMUSG00000004393 | Ddx56 | 0.633704 | 581.7645 | 0.171715 | -3.69045 | 0.000224 | 0.001705 | 0.002065 | 751.0039 | 663.8974 | 430.7077 | 481.4491 | 52513 | DEAD box helicase 56 [Source:MGI Symbol;Acc:MGI:1277172] |
| ENSMUSG00000005610 | Eif4g2 | 0.635374 | 12238.33 | 0.105183 | -6.04067 | 1.53E-09 | 3.37E-08 | 4.09E-08 | 16469.2 | 13312.37 | 9530.438 | 9641.326 | 13690 | eukaryotic translation initiation factor 4, gamma 2 [Source:MGI Symbol;Acc:MGI:109207] |
| ENSMUSG00000028063 | Lmna | 0.636504 | 537.9412 | 0.181565 | -3.50565 | 0.000455 | 0.00319 | 0.003864 | 684.3655 | 625.5389 | 458.1414 | 383.719 | 16905 | lamin A [Source:MGI Symbol;Acc:MGI:96794] |
| ENSMUSG00000079555 | Haus3 | 0.636901 | 465.98 | 0.193004 | -3.29993 | 0.000967 | 0.0061 | 0.007389 | 624.0736 | 510.4633 | 362.1237 | 367.2592 | 231123 | HAUS augmin-like complex, subunit 3 [Source:MGI Symbol;Acc:MGI:2387633] |
| ENSMUSG00000042747 | Krtcap2 | 0.637728 | 387.2154 | 0.198901 | -3.20626 | 0.001345 | 0.008063 | 0.009767 | 489.7391 | 453.4173 | 322.8022 | 282.9028 | 66059 | keratinocyte associated protein 2 [Source:MGI Symbol;Acc:MGI:1913309] |
| ENSMUSG00000041360 | Pum3 | 0.637874 | 673.9713 | 0.155419 | -4.10423 | 4.06E-05 | 0.00038 | 0.000461 | 864.1833 | 777.0058 | 516.6664 | 538.0296 | 52874 | pumilio RNA-binding family member 3 [Source:MGI Symbol;Acc:MGI:106253] |
| ENSMUSG00000041238 | Rbbp8 | 0.637965 | 652.0371 | 0.157143 | -4.05978 | 4.91E-05 | 0.000446 | 0.000541 | 831.393 | 756.3513 | 494.7195 | 525.6848 | 225182 | retinoblastoma binding protein 8, endonuclease [Source:MGI Symbol;Acc:MGI:2442995] |
| ENSMUSG00000071172 | Srsf3 | 0.639202 | 4132.072 | 0.10005 | -6.38883 | 1.67E-10 | 4.19E-09 | 5.07E-09 | 5392.419 | 4673.838 | 3296.606 | 3165.425 | 20383 | serine and arginine-rich splicing factor 3 [Source:MGI Symbol;Acc:MGI:98285] |
| ENSMUSG00000021131 | Erh | 0.639218 | 610.7545 | 0.161986 | -3.94613 | 7.94E-05 | 0.00068 | 0.000824 | 719.2713 | 768.1539 | 453.5691 | 502.0238 | 13877 | ERH mRNA splicing and mitosis factor [Source:MGI Symbol;Acc:MGI:108089] |
| ENSMUSG00000005374 | Tbl2 | 0.639222 | 309.498 | 0.21733 | -2.94125 | 0.003269 | 0.016825 | 0.020368 | 399.8302 | 354.0786 | 234.1002 | 249.9832 | 27368 | transducin (beta)-like 2 [Source:MGI Symbol;Acc:MGI:1351652] |
| ENSMUSG00000024067 | Dpy30 | 0.639709 | 328.2928 | 0.213564 | -2.99539 | 0.002741 | 0.014456 | 0.017505 | 406.1767 | 393.4207 | 236.8435 | 276.7303 | 66310 | dpy-30, histone methyltransferase complex regulatory subunit [Source:MGI Symbol;Acc:MGI:1913560] |
| ENSMUSG00000041219 | Arhgap11a | 0.640194 | 753.6274 | 0.163253 | -3.92147 | 8.8E-05 | 0.000744 | 0.000902 | 988.998 | 847.8216 | 627.3153 | 550.3745 | 228482 | Rho GTPase activating protein 11A [Source:MGI Symbol;Acc:MGI:2444300] |
| ENSMUSG00000020644 | Id2 | 0.640249 | 2969.753 | 0.106153 | -6.03139 | 1.63E-09 | 3.55E-08 | 4.3E-08 | 3751.846 | 3483.74 | 2167.256 | 2476.171 | 15902 | inhibitor of DNA binding 2 [Source:MGI Symbol;Acc:MGI:96397] |
| ENSMUSG00000021427 | Ssr1 | 0.640724 | 3338.724 | 0.118809 | -5.3929 | 6.93E-08 | 1.18E-06 | 1.43E-06 | 4418.23 | 3717.825 | 2444.335 | 2774.505 | 107513 | signal sequence receptor, alpha [Source:MGI Symbol;Acc:MGI:105082] |
| ENSMUSG00000022698 | Naa50 | 0.642274 | 2517.904 | 0.118864 | -5.40342 | 6.54E-08 | 1.12E-06 | 1.35E-06 | 3359.42 | 2779.517 | 1952.359 | 1980.319 | 72117 | N(alpha)-acetyltransferase 50, NatE catalytic subunit [Source:MGI Symbol;Acc:MGI:1919367] |
| ENSMUSG00000026496 | Parp1 | 0.642394 | 2872.874 | 0.097897 | -6.56197 | 5.31E-11 | 1.43E-09 | 1.73E-09 | 3679.919 | 3324.405 | 2207.492 | 2279.682 | 11545 | poly (ADP-ribose) polymerase family, member 1 [Source:MGI Symbol;Acc:MGI:1340806] |
| ENSMUSG00000056091 | St3gal5 | 0.643254 | 279.0038 | 0.242422 | -2.65344 | 0.007968 | 0.035315 | 0.042654 | 378.6752 | 301.9504 | 228.6135 | 206.7762 | 20454 | ST3 beta-galactoside alpha-2,3-sialyltransferase 5 [Source:MGI Symbol;Acc:MGI:1339963] |
| ENSMUSG00000018848 | Rars1 | 0.643589 | 1563.455 | 0.11732 | -5.48577 | 4.12E-08 | 7.28E-07 | 8.82E-07 | 2018.191 | 1794.982 | 1206.165 | 1234.485 | 104458 | arginyl-tRNA synthetase 1 [Source:MGI Symbol;Acc:MGI:1914297] |
| ENSMUSG00000028771 | Ptpn12 | 0.643689 | 571.5502 | 0.159179 | -4.04382 | 5.26E-05 | 0.000474 | 0.000574 | 704.4628 | 689.4697 | 441.6812 | 450.587 | 19248 | protein tyrosine phosphatase, non-receptor type 12 [Source:MGI Symbol;Acc:MGI:104673] |
| ENSMUSG00000030505 | Prmt3 | 0.643733 | 486.1488 | 0.178996 | -3.59635 | 0.000323 | 0.002356 | 0.002854 | 623.0159 | 562.5916 | 363.9526 | 395.0351 | 71974 | protein arginine N-methyltransferase 3 [Source:MGI Symbol;Acc:MGI:1919224] |
| ENSMUSG00000020018 | Snrpf | 0.64403 | 265.1208 | 0.240741 | -2.67519 | 0.007469 | 0.033485 | 0.040455 | 346.9426 | 299.9833 | 222.2123 | 191.3451 | 69878 | small nuclear ribonucleoprotein polypeptide F [Source:MGI Symbol;Acc:MGI:1917128] |
| ENSMUSG00000026806 | Ddx31 | 0.645396 | 489.8442 | 0.175326 | -3.68111 | 0.000232 | 0.001762 | 0.002135 | 602.9186 | 592.0981 | 362.1237 | 402.2363 | 227674 | DEAD/H box helicase 31 [Source:MGI Symbol;Acc:MGI:2682639] |
| ENSMUSG00000031715 | Smarca5 | 0.645519 | 2828.857 | 0.117764 | -5.48145 | 4.22E-08 | 7.45E-07 | 9.02E-07 | 3713.767 | 3188.675 | 2068.495 | 2344.492 | 93762 | SNF2 related chromatin remodeling ATPase 5 [Source:MGI Symbol;Acc:MGI:1935129] |
| ENSMUSG00000037465 | Klf10 | 0.646089 | 732.4398 | 0.164242 | -3.93376 | 8.36E-05 | 0.000712 | 0.000862 | 867.3566 | 920.6044 | 626.4009 | 515.3974 | 21847 | Kruppel-like transcription factor 10 [Source:MGI Symbol;Acc:MGI:1101353] |
| ENSMUSG00000056952 | Tatdn2 | 0.646991 | 1234.644 | 0.143469 | -4.50963 | 6.49E-06 | 7.25E-05 | 8.78E-05 | 1596.148 | 1417.298 | 877.8757 | 1047.255 | 381801 | TatD DNase domain containing 2 [Source:MGI Symbol;Acc:MGI:3576210] |
| ENSMUSG00000021957 | Tkt | 0.64754 | 4629.849 | 0.082981 | -7.80344 | 6.02E-15 | 2.57E-13 | 3.12E-13 | 5861.003 | 5442.975 | 3623.066 | 3592.351 | 21881 | transketolase [Source:MGI Symbol;Acc:MGI:105992] |
| ENSMUSG00000020437 | Myo1g | 0.64828 | 711.8055 | 0.171761 | -3.77431 | 0.00016 | 0.001264 | 0.001532 | 909.6666 | 829.1341 | 613.5985 | 494.8227 | 246177 | myosin IG [Source:MGI Symbol;Acc:MGI:1927091] |
| ENSMUSG00000045294 | Insig1 | 0.648555 | 1413.63 | 0.110745 | -5.85627 | 4.73E-09 | 9.63E-08 | 1.17E-07 | 1751.637 | 1700.561 | 1088.2 | 1114.123 | 231070 | insulin induced gene 1 [Source:MGI Symbol;Acc:MGI:1916289] |
| ENSMUSG00000024829 | Mrpl21 | 0.648566 | 302.6663 | 0.224343 | -2.89096 | 0.003841 | 0.019335 | 0.023383 | 399.8302 | 339.3253 | 227.699 | 243.8108 | 353242 | mitochondrial ribosomal protein L21 [Source:MGI Symbol;Acc:MGI:2660674] |
| ENSMUSG00000030751 | Psma1 | 0.649058 | 2515.263 | 0.10082 | -6.43778 | 1.21E-10 | 3.14E-09 | 3.8E-09 | 3232.49 | 2911.313 | 1968.819 | 1948.429 | 26440 | proteasome subunit alpha 1 [Source:MGI Symbol;Acc:MGI:1347005] |
| ENSMUSG00000044906 | 4930503L19Rik | 0.650039 | 350.5926 | 0.213176 | -3.04931 | 0.002294 | 0.012484 | 0.015122 | 464.3531 | 392.4371 | 285.3096 | 260.2706 | 269033 | RIKEN cDNA 4930503L19 gene [Source:MGI Symbol;Acc:MGI:1922045] |
| ENSMUSG00000025289 | Prdx4 | 0.650214 | 1205.921 | 0.125922 | -5.16364 | 2.42E-07 | 3.64E-06 | 4.4E-06 | 1543.26 | 1403.528 | 967.4921 | 909.4038 | 53381 | peroxiredoxin 4 [Source:MGI Symbol;Acc:MGI:1859815] |
| ENSMUSG00000028683 | Eif2b3 | 0.650545 | 243.7826 | 0.249716 | -2.60514 | 0.009184 | 0.039718 | 0.047976 | 315.2101 | 280.3122 | 172.8318 | 206.7762 | 108067 | eukaryotic translation initiation factor 2B, subunit 3 [Source:MGI Symbol;Acc:MGI:1313286] |
| ENSMUSG00000071014 | Ndufb6 | 0.650625 | 466.8801 | 0.20623 | -3.15485 | 0.001606 | 0.009332 | 0.011305 | 552.1465 | 588.1639 | 313.6577 | 413.5524 | 230075 | NADH:ubiquinone oxidoreductase subunit B6 [Source:MGI Symbol;Acc:MGI:2684983] |
| ENSMUSG00000020720 | Psmd12 | 0.650974 | 1098.894 | 0.121389 | -5.36273 | 8.2E-08 | 1.37E-06 | 1.66E-06 | 1333.825 | 1351.4 | 843.1264 | 867.2256 | 66997 | proteasome (prosome, macropain) 26S subunit, non-ATPase, 12 [Source:MGI Symbol;Acc:MGI:1914247] |
| ENSMUSG00000064360 | mt-Nd3 | 0.652391 | 1302.875 | 0.191223 | -3.41169 | 0.000646 | 0.004315 | 0.005227 | 1500.95 | 1684.824 | 1217.138 | 808.5876 | 17718 | mitochondrially encoded NADH dehydrogenase 3 [Source:MGI Symbol;Acc:MGI:102499] |
| ENSMUSG00000062867 | Impdh2 | 0.652454 | 1107.988 | 0.143227 | -4.55537 | 5.23E-06 | 5.96E-05 | 7.22E-05 | 1466.044 | 1243.209 | 900.737 | 821.9611 | 23918 | inosine monophosphate dehydrogenase 2 [Source:MGI Symbol;Acc:MGI:109367] |
| ENSMUSG00000025287 | Acot9 | 0.652462 | 617.5486 | 0.167772 | -3.88897 | 0.000101 | 0.000836 | 0.001013 | 815.5267 | 694.3875 | 481.9172 | 478.3629 | 56360 | acyl-CoA thioesterase 9 [Source:MGI Symbol;Acc:MGI:1928939] |
| ENSMUSG00000019838 | Slc16a10 | 0.652582 | 1571.309 | 0.112237 | -5.81434 | 6.09E-09 | 1.22E-07 | 1.48E-07 | 1993.862 | 1848.094 | 1250.973 | 1192.307 | 72472 | solute carrier family 16 (monocarboxylic acid transporters), member 10 [Source:MGI Symbol;Acc:MGI:1919722] |
| ENSMUSG00000040688 | Tbl3 | 0.652762 | 619.5804 | 0.167359 | -3.90036 | 9.6E-05 | 0.0008 | 0.000969 | 782.7364 | 731.7625 | 445.339 | 518.4836 | 213773 | transducin (beta)-like 3 [Source:MGI Symbol;Acc:MGI:2384863] |
| ENSMUSG00000039842 | Mcph1 | 0.654149 | 336.4424 | 0.214228 | -3.05352 | 0.002262 | 0.012347 | 0.014957 | 433.6783 | 389.4865 | 279.8229 | 242.782 | 244329 | microcephaly, primary autosomal recessive 1 [Source:MGI Symbol;Acc:MGI:2443308] |
| ENSMUSG00000029390 | Tmed2 | 0.655004 | 5808.223 | 0.09073 | -7.21927 | 5.23E-13 | 1.81E-11 | 2.19E-11 | 7479.364 | 6729.461 | 4346.399 | 4677.669 | 56334 | transmembrane p24 trafficking protein 2 [Source:MGI Symbol;Acc:MGI:1929269] |
| ENSMUSG00000025613 | Cct8 | 0.656141 | 2968.722 | 0.091446 | -7.17517 | 7.22E-13 | 2.47E-11 | 2.99E-11 | 3746.557 | 3518.164 | 2260.53 | 2349.636 | 12469 | chaperonin containing TCP1 subunit 8 [Source:MGI Symbol;Acc:MGI:107183] |
| ENSMUSG00000024309 | Pfdn6 | 0.656815 | 258.3979 | 0.246674 | -2.66269 | 0.007752 | 0.034576 | 0.041774 | 334.2496 | 298.0162 | 180.1474 | 221.1785 | 14976 | prefoldin subunit 6 [Source:MGI Symbol;Acc:MGI:95908] |
| ENSMUSG00000032802 | Srxn1 | 0.656928 | 896.5064 | 0.143139 | -4.58944 | 4.44E-06 | 5.11E-05 | 6.19E-05 | 1115.928 | 1077.973 | 648.3478 | 743.7771 | 76650 | sulfiredoxin 1 homolog (S. cerevisiae) [Source:MGI Symbol;Acc:MGI:104971] |

[illegible]

| | | | | | | | | | | | | | |
|---|---|---|---|---|---|---|---|---|---|---|---|---|---|
| ENSMUSG00000020392 | Cdkn2aipnl | 0.868457 | 922.8068 | 0.16337 | -5.31589 | 1.06E-07 | 1.72E-06 | 2.09E-06 | 1329.594 | 1055.351 | 627.3153 | 678.9667 | 52626 | CDKN2A interacting protein N-terminal like [Source:MGI Symbol;Acc:MGI:1261797] |
| ENSMUSG00000002984 | Tomm40 | 0.868532 | 428.2832 | 0.214266 | -4.05353 | 5.05E-05 | 0.000458 | 0.000554 | 614.5539 | 492.7594 | 330.1178 | 275.7016 | 53333 | translocase of outer mitochondrial membrane 40 [Source:MGI Symbol;Acc:MGI:1858259] |
| ENSMUSG00000022978 | Mis18a | 0.868903 | 304.3319 | 0.228701 | -3.7993 | 0.000145 | 0.001159 | 0.001404 | 431.5628 | 355.0622 | 208.4955 | 222.2073 | 66578 | MIS18 kinetochore protein A [Source:MGI Symbol;Acc:MGI:1913828] |
| ENSMUSG00000024844 | Banf1 | 0.870794 | 300.2305 | 0.238517 | -3.65087 | 0.000261 | 0.00196 | 0.002374 | 416.7543 | 359.9799 | 235.9291 | 188.2589 | 23825 | BAF nuclear assembly factor 1 [Source:MGI Symbol;Acc:MGI:1346330] |
| ENSMUSG00000024140 | Epas1 | 0.871749 | 128.325 | 0.323609 | -2.69384 | 0.007063 | 0.031962 | 0.038614 | 174.5291 | 157.3683 | 87.78757 | 93.6151 | 13819 | endothelial PAS domain protein 1 [Source:MGI Symbol;Acc:MGI:109169] |
| ENSMUSG00000027276 | Jag1 | 0.873603 | 278.3478 | 0.257017 | -3.399 | 0.000676 | 0.004482 | 0.005429 | 414.6387 | 305.8846 | 206.6666 | 186.2015 | 16449 | jagged 1 [Source:MGI Symbol;Acc:MGI:1095416] |
| ENSMUSG00000025007 | Aldh18a1 | 0.873633 | 401.0578 | 0.204367 | -4.27482 | 1.91E-05 | 0.000192 | 0.000233 | 566.955 | 471.1213 | 293.5397 | 272.6154 | 56454 | aldehyde dehydrogenase 18 family, member A1 [Source:MGI Symbol;Acc:MGI:1888908] |
| ENSMUSG00000020328 | Nudcd2 | 0.874013 | 475.6683 | 0.176688 | -4.94665 | 7.55E-07 | 1.02E-05 | 1.24E-05 | 640.9977 | 590.131 | 339.2624 | 332.2822 | 52653 | NudC domain containing 2 [Source:MGI Symbol;Acc:MGI:1277103] |
| ENSMUSG00000008384 | Sertad1 | 0.876697 | 258.9089 | 0.286206 | -3.06317 | 0.00219 | 0.012015 | 0.014554 | 284.5353 | 385.5523 | 153.6282 | 211.9199 | 55942 | SERTA domain containing 1 [Source:MGI Symbol;Acc:MGI:1913438] |
| ENSMUSG00000035351 | Nup37 | 0.878275 | 196.5442 | 0.274622 | -3.19812 | 0.001383 | 0.008231 | 0.009971 | 275.0155 | 234.0853 | 128.938 | 148.1382 | 69736 | nucleoporin 37 [Source:MGI Symbol;Acc:MGI:1919964] |
| ENSMUSG00000001467 | Cyp51 | 0.87892 | 596.4719 | 0.160854 | -5.46408 | 4.65E-08 | 8.15E-07 | 9.88E-07 | 786.9674 | 758.3184 | 402.3597 | 438.2421 | 13121 | cytochrome P450, family 51 [Source:MGI Symbol;Acc:MGI:106040] |
| ENSMUSG00000051627 | H1f4 | 0.879268 | 2567.381 | 0.164123 | -5.35739 | 8.44E-08 | 1.41E-06 | 1.7E-06 | 3592.125 | 3061.796 | 2118.79 | 1496.813 | 50709 | H1.4 linker histone, cluster member [Source:MGI Symbol;Acc:MGI:1931527] |
| ENSMUSG00000031843 | Mphosph6 | 0.879455 | 226.6159 | 0.249281 | -3.52797 | 0.000419 | 0.00297 | 0.003598 | 308.8636 | 278.3451 | 153.6282 | 165.6267 | 68533 | M phase phosphoprotein 6 [Source:MGI Symbol;Acc:MGI:1915783] |
| ENSMUSG00000066595 | Flvcr1 | 0.882923 | 860.492 | 0.144028 | -6.13022 | 8.78E-10 | 1.99E-08 | 2.41E-08 | 1053.521 | 1178.295 | 622.7431 | 587.409 | 226844 | feline leukemia virus subgroup C cellular receptor 1 [Source:MGI Symbol;Acc:MGI:2444881] |
| ENSMUSG00000019823 | Mical1 | 0.883401 | 440.1544 | 0.199583 | -4.42624 | 9.59E-06 | 0.000104 | 0.000126 | 563.7818 | 578.3284 | 343.8346 | 274.6729 | 171580 | microtubule associated monooxygenase, calponin and LIM domain containing 1 [Source:MGI Symbol;Acc:MGI:2385847] |
| ENSMUSG00000020899 | Pfas | 0.88433 | 663.7856 | 0.184391 | -4.79594 | 1.62E-06 | 2.04E-05 | 2.47E-05 | 972.074 | 750.4499 | 481.0027 | 451.6157 | 237823 | phosphoribosylformylglycinamidine synthase (FGAR amidotransferase) [Source:MGI Symbol;Acc:MGI:2684864] |
| ENSMUSG00000026999 | Nup35 | 0.886446 | 278.2233 | 0.232286 | -3.81619 | 0.000136 | 0.001088 | 0.001319 | 380.7907 | 341.2924 | 181.9763 | 208.8337 | 69482 | nucleoporin 35 [Source:MGI Symbol;Acc:MGI:1916732] |
| ENSMUSG00000049327 | Kmt5a | 0.887084 | 640.0207 | 0.160775 | -5.51754 | 3.44E-08 | 6.17E-07 | 7.47E-07 | 873.7031 | 787.8249 | 432.5367 | 466.018 | 67956 | lysine methyltransferase 5A [Source:MGI Symbol;Acc:MGI:1915206] |
| ENSMUSG00000029752 | Asns | 0.889409 | 199.0334 | 0.30413 | -2.92443 | 0.003451 | 0.017601 | 0.021307 | 306.7481 | 210.4801 | 147.2271 | 131.6784 | 27053 | asparagine synthetase [Source:MGI Symbol;Acc:MGI:1350929] |
| ENSMUSG00000061024 | Rrs1 | 0.890641 | 180.7249 | 0.288048 | -3.09199 | 0.001988 | 0.011092 | 0.013436 | 227.4167 | 241.9537 | 111.5634 | 141.9658 | 59014 | ribosome biogenesis regulator 1 [Source:MGI Symbol;Acc:MGI:1929721] |
| ENSMUSG00000020869 | Lrrc59 | 0.892673 | 2289.063 | 0.109527 | -8.15023 | 3.63E-16 | 1.75E-14 | 2.12E-14 | 3160.563 | 2790.336 | 1557.315 | 1648.037 | 98238 | leucine rich repeat containing 59 [Source:MGI Symbol;Acc:MGI:2138133] |
| ENSMUSG00000025364 | Pa2g4 | 0.89351 | 807.2844 | 0.15356 | -5.81863 | 5.93E-09 | 1.19E-07 | 1.44E-07 | 1107.466 | 991.4201 | 529.4688 | 600.7826 | 18813 | proliferation-associated 2G4 [Source:MGI Symbol;Acc:MGI:894684] |
| ENSMUSG00000057406 | Nsd2 | 0.893897 | 2020.968 | 0.141982 | -6.29584 | 3.06E-10 | 7.37E-09 | 8.93E-09 | 2959.59 | 2296.593 | 1467.698 | 1359.991 | 107823 | nuclear receptor binding SET domain protein 2 [Source:MGI Symbol;Acc:MGI:1276574] |
| ENSMUSG00000038126 | Mphosph9 | 0.894557 | 246.5604 | 0.272813 | -3.27901 | 0.001042 | 0.006508 | 0.007884 | 362.8089 | 278.3451 | 152.7138 | 192.3739 | 269702 | M-phase phosphoprotein 9 [Source:MGI Symbol;Acc:MGI:2443138] |
| ENSMUSG00000033721 | Vav3 | 0.894712 | 1143.29 | 0.149944 | -5.96698 | 2.42E-09 | 5.15E-08 | 6.23E-08 | 1651.151 | 1322.877 | 802.8905 | 796.2427 | 57257 | vav 3 oncogene [Source:MGI Symbol;Acc:MGI:1888518] |
| ENSMUSG00000049553 | Polr1a | 0.898787 | 605.0344 | 0.183535 | -4.89709 | 9.73E-07 | 1.28E-05 | 1.56E-05 | 880.0496 | 695.371 | 418.8198 | 425.8973 | 20019 | polymerase (RNA) I polypeptide A [Source:MGI Symbol;Acc:MGI:1096397] |
| ENSMUSG00000036208 | Nepro | 0.899184 | 206.3023 | 0.259456 | -3.46565 | 0.000529 | 0.003639 | 0.004408 | 282.4198 | 254.7399 | 139.9114 | 148.1382 | 212547 | nucleolus and neural progenitor protein [Source:MGI Symbol;Acc:MGI:2384836] |
| ENSMUSG00000056220 | Pla2g4a | 0.900617 | 1492.974 | 0.138616 | -6.49722 | 8.18E-11 | 2.16E-09 | 2.61E-09 | 2127.139 | 1761.541 | 987.6101 | 1095.605 | 18783 | phospholipase A2, group IVA (cytosolic, calcium-dependent) [Source:MGI Symbol;Acc:MGI:1195256] |
| ENSMUSG00000037722 | Gnpnat1 | 0.900884 | 274.7036 | 0.225845 | -3.98894 | 6.64E-05 | 0.000582 | 0.000705 | 373.3864 | 342.276 | 193.8642 | 189.2877 | 54342 | glucosamine-phosphate N-acetyltransferase 1 [Source:MGI Symbol;Acc:MGI:1858963] |
| ENSMUSG00000035311 | Gnptab | 0.901927 | 1265.327 | 0.125298 | -7.19823 | 6.1E-13 | 2.1E-11 | 2.54E-11 | 1738.944 | 1557.946 | 863.2444 | 901.1739 | 432486 | N-acetylglucosamine-1-phosphate transferase, alpha and beta subunits [Source:MGI Symbol;Acc:MGI:3643902] |
| ENSMUSG00000028367 | Txn1 | 0.902351 | 2093.268 | 0.101698 | -8.87288 | 7.13E-19 | 4.59E-17 | 5.56E-17 | 2797.754 | 2657.557 | 1504.277 | 1413.485 | 22166 | thioredoxin 1 [Source:MGI Symbol;Acc:MGI:98874] |
| ENSMUSG00000003810 | Mast2 | 0.902454 | 618.0391 | 0.16473 | -5.47837 | 4.29E-08 | 7.56E-07 | 9.16E-07 | 826.1042 | 784.8743 | 459.9703 | 401.2076 | 17776 | microtubule associated serine/threonine kinase 2 [Source:MGI Symbol;Acc:MGI:894676] |
| ENSMUSG00000042462 | Dctpp1 | 0.9038 | 304.1192 | 0.214268 | -4.21809 | 2.46E-05 | 0.000241 | 0.000292 | 408.2922 | 384.5687 | 215.8111 | 207.8049 | 66422 | dCTP pyrophosphatase 1 [Source:MGI Symbol;Acc:MGI:1913672] |
| ENSMUSG00000032344 | Cgas | 0.905581 | 192.1301 | 0.315666 | -2.8688 | 0.00412 | 0.020484 | 0.024773 | 300.4015 | 200.6445 | 123.4513 | 144.0232 | 214763 | cyclic GMP-AMP synthase [Source:MGI Symbol;Acc:MGI:2442261] |
| ENSMUSG00000048279 | Sacs | 0.90664 | 906.5563 | 0.1763 | -5.1426 | 2.71E-07 | 4.02E-06 | 4.87E-06 | 1351.807 | 1013.058 | 608.1118 | 653.2482 | 50720 | sacsin [Source:MGI Symbol;Acc:MGI:1354724] |
| ENSMUSG00000025403 | Shmt2 | 0.907315 | 320.0716 | 0.264577 | -3.4293 | 0.000605 | 0.004084 | 0.004947 | 469.6418 | 365.8812 | 260.6193 | 184.144 | 108037 | serine hydroxymethyltransferase 2 (mitochondrial) [Source:MGI Symbol;Acc:MGI:1277989] |
| ENSMUSG00000036678 | Aaas | 0.907733 | 287.3376 | 0.248114 | -3.65853 | 0.000254 | 0.001906 | 0.002309 | 407.2345 | 342.276 | 175.5751 | 224.2647 | 223921 | achalasia, adrenocortical insufficiency, alacrimia [Source:MGI Symbol;Acc:MGI:2443767] |
| ENSMUSG00000035024 | Ncapd3 | 0.907813 | 667.3699 | 0.180114 | -5.04022 | 4.65E-07 | 6.59E-06 | 7.98E-06 | 975.2473 | 766.1868 | 451.7402 | 476.3054 | 78658 | non-SMC condensin II complex, subunit D3 [Source:MGI Symbol;Acc:MGI:2142989] |
| ENSMUSG00000054808 | Actn4 | 0.908342 | 1948.935 | 0.15672 | -5.79596 | 6.79E-09 | 1.35E-07 | 1.64E-07 | 2913.049 | 2173.649 | 1445.751 | 1263.289 | 60595 | actinin alpha 4 [Source:MGI Symbol;Acc:MGI:1890773] |
| ENSMUSG00000030042 | Pole4 | 0.908601 | 577.0686 | 0.17308 | -5.2496 | 1.52E-07 | 2.39E-06 | 2.89E-06 | 799.6604 | 706.1901 | 378.5839 | 423.8398 | 66979 | polymerase (DNA-directed), epsilon 4 (p12 subunit) [Source:MGI Symbol;Acc:MGI:1914229] |
| ENSMUSG00000035704 | Alg8 | 0.908713 | 194.5806 | 0.282621 | -3.21531 | 0.001303 | 0.007834 | 0.00949 | 285.593 | 222.2827 | 132.5958 | 137.8508 | 381903 | ALG8 alpha-1,3-glucosyltransferase [Source:MGI Symbol;Acc:MGI:2141959] |
| ENSMUSG00000004356 | Utp20 | 0.910175 | 1214.36 | 0.146888 | -6.19641 | 5.78E-10 | 1.35E-08 | 1.63E-08 | 1727.309 | 1442.87 | 789.1736 | 898.0877 | 70683 | UTP20 small subunit processome component [Source:MGI Symbol;Acc:MGI:1917933] |
| ENSMUSG00000035352 | Ccl12 | 0.912733 | 795.9158 | 0.223572 | -4.08251 | 4.46E-05 | 0.000413 | 0.0005 | 1277.764 | 801.5946 | 531.2977 | 573.0067 | 20293 | C-C motif chemokine ligand 12 [Source:MGI Symbol;Acc:MGI:108224] |
| ENSMUSG00000063694 | Cycs | 0.914527 | 615.3613 | 0.188263 | -4.8577 | 1.19E-06 | 1.55E-05 | 1.87E-05 | 898.0314 | 710.1243 | 399.6163 | 453.6732 | 13063 | cytochrome c, somatic [Source:MGI Symbol;Acc:MGI:88578] |
| ENSMUSG00000036825 | Ssx2ip | 0.916207 | 539.938 | 0.171552 | -5.3407 | 9.26E-08 | 1.53E-06 | 1.85E-06 | 715.0403 | 696.3546 | 350.2358 | 398.1214 | 99167 | SSX family member 2 interacting protein [Source:MGI Symbol;Acc:MGI:2139150] |
| ENSMUSG00000020290 | Xpo1 | 0.917299 | 3702.487 | 0.114365 | -8.02081 | 1.05E-15 | 4.89E-14 | 5.92E-14 | 5313.088 | 4369.92 | 2504.689 | 2622.252 | 103573 | exportin 1 [Source:MGI Symbol;Acc:MGI:2144013] |
| ENSMUSG00000022391 | Rangap1 | 0.917315 | 1387.601 | 0.140412 | -6.53305 | 6.44E-11 | 1.71E-09 | 2.07E-09 | 2001.267 | 1627.778 | 940.973 | 980.3867 | 19387 | RAN GTPase activating protein 1 [Source:MGI Symbol;Acc:MGI:103071] |
| ENSMUSG00000025823 | Pdia4 | 0.917442 | 3400.751 | 0.083419 | -10.998 | 3.91E-28 | 6.51E-26 | 7.88E-26 | 4496.503 | 4397.46 | 2333.686 | 2375.355 | 12304 | protein disulfide isomerase associated 4 [Source:MGI Symbol;Acc:MGI:104864] |
| ENSMUSG00000055491 | Pprc1 | 0.918911 | 255.0628 | 0.285328 | -3.22054 | 0.001279 | 0.007714 | 0.009344 | 401.9457 | 265.559 | 181.9763 | 170.7704 | 226169 | peroxisome proliferative activated receptor, gamma, coactivator-related 1 [Source:MGI Symbol;Acc:MGI:2385096] |
| ENSMUSG00000019849 | Prep | 0.920758 | 718.0732 | 0.176378 | -5.22036 | 1.79E-07 | 2.76E-06 | 3.34E-06 | 1054.579 | 825.1999 | 503.8641 | 488.6502 | 19072 | prolyl endopeptidase [Source:MGI Symbol;Acc:MGI:1270863] |
| ENSMUSG00000026202 | Tuba4a | 0.923645 | 613.6033 | 0.157393 | -5.86842 | 4.4E-09 | 9.02E-08 | 1.09E-07 | 782.7364 | 824.2163 | 413.3331 | 434.1272 | 22145 | tubulin, alpha 4A [Source:MGI Symbol;Acc:MGI:1095410] |
| ENSMUSG00000020358 | Hnrnpab | 0.923988 | 3132.019 | 0.135772 | -6.80546 | 1.01E-11 | 2.97E-10 | 3.59E-10 | 4540.929 | 3662.747 | 1970.648 | 2353.751 | 15384 | heterogeneous nuclear ribonucleoprotein A/B [Source:MGI Symbol;Acc:MGI:1330294] |
| ENSMUSG00000060860 | Ube2s | 0.92452 | 1457.393 | 0.134921 | -6.85228 | 7.27E-12 | 2.2E-10 | 2.66E-10 | 2092.233 | 1726.133 | 1012.3 | 998.904 | 77891 | ubiquitin-conjugating enzyme E2S [Source:MGI Symbol;Acc:MGI:1925141] |
| ENSMUSG00000054115 | Skp2 | 0.924767 | 366.5999 | 0.246906 | -3.74543 | 0.00018 | 0.001401 | 0.001697 | 570.1283 | 390.47 | 249.6459 | 256.1556 | 27401 | S-phase kinase-associated protein 2 [Source:MGI Symbol;Acc:MGI:1351663] |
| ENSMUSG00000040212 | Emp3 | 0.925943 | 845.2438 | 0.14907 | -6.21146 | 5.25E-10 | 1.23E-08 | 1.49E-08 | 1159.296 | 1056.335 | 619.0852 | 546.2595 | 13732 | epithelial membrane protein 3 [Source:MGI Symbol;Acc:MGI:1098729] |
| ENSMUSG00000053801 | Grwd1 | 0.928753 | 234.037 | 0.265476 | -3.49845 | 0.000468 | 0.003269 | 0.003959 | 334.2496 | 279.3287 | 143.5692 | 179.0003 | 101612 | glutamate-rich WD repeat containing 1 [Source:MGI Symbol;Acc:MGI:2141989] |
| ENSMUSG00000030156 | Cd69 | 0.929343 | 284.4438 | 0.346532 | -2.68184 | 0.007322 | 0.032944 | 0.039801 | 501.3744 | 244.9044 | 209.4099 | 182.0865 | 12515 | CD69 antigen [Source:MGI Symbol;Acc:MGI:88343] |
| ENSMUSG00000118665 | Lin54 | 0.930193 | 562.2598 | 0.172586 | -5.38973 | 7.06E-08 | 1.2E-06 | 1.45E-06 | 784.8519 | 690.4533 | 402.3597 | 371.3742 | 231506 | lin-54 DREAM MuvB core complex component [Source:MGI Symbol;Acc:MGI:2140902] |
| ENSMUSG00000034317 | Trim59 | 0.930625 | 298.6425 | 0.264797 | -3.51449 | 0.000441 | 0.003106 | 0.003762 | 455.8911 | 327.5227 | 183.8052 | 227.351 | 66949 | tripartite motif-containing 59 [Source:MGI Symbol;Acc:MGI:1914199] |
| ENSMUSG00000063524 | Eno1 | 0.931724 | 7144.361 | 0.082016 | -11.3603 | 6.59E-30 | 1.39E-27 | 1.68E-27 | 9653.044 | 9096.87 | 5111.797 | 4715.732 | 13806 | enolase 1, alpha non-neuron [Source:MGI Symbol;Acc:MGI:95393] |
| ENSMUSG00000010048 | Ifrd2 | 0.935141 | 146.5001 | 0.302392 | -3.09248 | 0.001985 | 0.011084 | 0.013426 | 199.9151 | 184.9077 | 102.4188 | 98.75879 | 15983 | interferon-related developmental regulator 2 [Source:MGI Symbol;Acc:MGI:1316708] |
| ENSMUSG00000004099 | Dnmt1 | 0.935576 | 2640.879 | 0.115465 | -8.1027 | 5.38E-16 | 2.56E-14 | 3.1E-14 | 3763.481 | 3173.921 | 1861.828 | 1764.285 | 13433 | DNA methyltransferase 1 [Source:MGI Symbol;Acc:MGI:94912] |
| ENSMUSG00000030615 | Tmem126a | 0.935817 | 527.4917 | 0.19837 | -4.71754 | 2.39E-06 | 2.89E-05 | 3.5E-05 | 783.7942 | 601.9336 | 353.8936 | 370.3454 | 66271 | transmembrane protein 126A [Source:MGI Symbol;Acc:MGI:1913521] |
| ENSMUSG00000060288 | Ppih | 0.937462 | 268.3718 | 0.248862 | -3.767 | 0.000165 | 0.001298 | 0.001572 | 392.426 | 312.7694 | 172.8318 | 195.4601 | 66101 | peptidyl prolyl isomerase H [Source:MGI Symbol;Acc:MGI:106499] |
| ENSMUSG00000061533 | Cep128 | 0.939309 | 323.7427 | 0.223373 | -4.20511 | 2.61E-05 | 0.000254 | 0.000307 | 430.505 | 420.9601 | 245.9881 | 197.5176 | 75216 | centrosomal protein 128 [Source:MGI Symbol;Acc:MGI:1922466] |
| ENSMUSG00000060152 | Pop5 | 0.939499 | 227.743 | 0.252946 | -3.71423 | 0.000204 | 0.001566 | 0.001897 | 321.5566 | 277.3616 | 161.8583 | 150.1957 | 117109 | processing of precursor 5, ribonuclease P/MRP family (S. cerevisiae) [Source:MGI Symbol;Acc:MGI:2151221] |
| ENSMUSG00000004100 | Ppan | 0.94375 | 283.6952 | 0.241041 | -3.9153 | 9.03E-05 | 0.000761 | 0.000922 | 406.1767 | 340.3089 | 177.404 | 210.8912 | 235036 | peter pan homolog [Source:MGI Symbol;Acc:MGI:2178445] |
| ENSMUSG00000006442 | Srm | 0.948415 | 273.4055 | 0.25809 | -3.67475 | 0.000238 | 0.001802 | 0.002183 | 411.4655 | 308.8352 | 173.7462 | 199.575 | 20810 | spermidine synthase [Source:MGI Symbol;Acc:MGI:102690] |
| ENSMUSG00000030342 | Cd9 | 0.948852 | 3777.747 | 0.080852 | -11.7357 | 8.36E-32 | 2.26E-29 | 2.74E-29 | 4948.163 | 5006.278 | 2605.279 | 2551.269 | 12527 | CD9 antigen [Source:MGI Symbol;Acc:MGI:88348] |
| ENSMUSG00000033909 | Usp36 | 0.949158 | 225.5066 | 0.248343 | -3.82197 | 0.000132 | 0.001068 | 0.001293 | 312.0368 | 282.2793 | 155.4571 | 152.2531 | 72344 | ubiquitin specific peptidase 36 [Source:MGI Symbol;Acc:MGI:1919594] |
| ENSMUSG00000022893 | Adamts1 | 0.94965 | 660.5964 | 0.235158 | -4.03834 | 5.38E-05 | 0.000484 | 0.000587 | 1035.539 | 705.2066 | 383.1562 | 518.4836 | 11504 | ADAM metallopeptidase with thrombospondin type 1 motif 1 [Source:MGI Symbol;Acc:MGI:109249] |
| ENSMUSG00000031432 | Prps1 | 0.949986 | 206.4789 | 0.258461 | -3.67555 | 0.000237 | 0.001798 | 0.002178 | 285.593 | 258.6741 | 141.7403 | 139.9083 | 19139 | phosphoribosyl pyrophosphate synthetase 1 [Source:MGI Symbol;Acc:MGI:97775] |
| ENSMUSG00000004880 | Lbr | 0.950194 | 1060.672 | 0.137722 | -6.89935 | 5.22E-12 | 1.63E-10 | 1.98E-10 | 1492.488 | 1303.206 | 703.215 | 743.7771 | 98386 | lamin B receptor [Source:MGI Symbol;Acc:MGI:2138281] |
| ENSMUSG00000031904 | Slc7a6 | 0.951005 | 309.7761 | 0.233077 | -4.08022 | 4.5E-05 | 0.000416 | 0.000504 | 452.7178 | 363.9141 | 200.2654 | 222.2073 | 330836 | solute carrier family 7 (cationic amino acid transporter, y+ system), member 6 [Source:MGI Symbol;Acc:MGI:2142598] |
| ENSMUSG00000003038 | Hmgn2 | 0.951252 | 1974.788 | 0.124191 | -7.65962 | 1.86E-14 | 7.54E-13 | 9.13E-13 | 2808.331 | 2397.899 | 1278.406 | 1414.514 | 15331 | high mobility group nucleosomal binding domain 2 [Source:MGI Symbol;Acc:MGI:96136] |
| ENSMUSG00000053553 | Chlsn | 0.951476 | 127.8483 | 0.334125 | -2.84767 | 0.004404 | 0.02157 | 0.026078 | 176.6446 | 160.3189 | 77.72857 | 96.70131 | 73212 | cholesin [Source:MGI Symbol;Acc:MGI:1920462] |
| ENSMUSG00000020250 | Txnrd1 | 0.952142 | 3151.139 | 0.099437 | -9.57535 | 1.02E-21 | 9.08E-20 | 1.1E-19 | 4397.075 | 3912.569 | 2107.816 | 2187.096 | 50493 | thioredoxin reductase 1 [Source:MGI Symbol;Acc:MGI:1354175] |
| ENSMUSG00000038335 | Tsr1 | 0.95407 | 582.6063 | 0.194475 | -4.90588 | 9.3E-07 | 1.23E-05 | 1.49E-05 | 848.317 | 688.4862 | 358.4659 | 435.1559 | 104662 | TSR1 20S rRNA accumulation [Source:MGI Symbol;Acc:MGI:2144566] |
| ENSMUSG00000030208 | Emp1 | 0.954474 | 253.1047 | 0.234786 | -4.06529 | 4.8E-05 | 0.000438 | 0.000531 | 334.2496 | 333.424 | 163.6872 | 181.0578 | 13730 | epithelial membrane protein 1 [Source:MGI Symbol;Acc:MGI:107941] |
| ENSMUSG00000022503 | Nubp1 | 0.95574 | 424.2141 | 0.197112 | -4.84872 | 1.24E-06 | 1.61E-05 | 1.95E-05 | 609.2651 | 510.4633 | 288.053 | 289.0752 | 26425 | nucleotide binding protein 1 [Source:MGI Symbol;Acc:MGI:1347073] |
| ENSMUSG00000000751 | Rpa1 | 0.955953 | 1364.743 | 0.137948 | -6.92981 | 4.21E-12 | 1.33E-10 | 1.61E-10 | 1967.419 | 1634.663 | 897.0792 | 959.8119 | 68275 | replication protein A1 [Source:MGI Symbol;Acc:MGI:1915525] |
| ENSMUSG00000038482 | Tfdp1 | 0.958259 | 1283.555 | 0.129752 | -7.38529 | 1.52E-13 | 5.6E-12 | 6.78E-12 | 1817.218 | 1572.699 | 876.0468 | 868.2543 | 21781 | transcription factor Dp 1 [Source:MGI Symbol;Acc:MGI:101934] |
| ENSMUSG00000031353 | Rbbp7 | 0.959337 | 2155.769 | 0.122614 | -7.82405 | 5.12E-15 | 2.21E-13 | 2.68E-13 | 3077 | 2617.231 | 1390.884 | 1537.962 | 245688 | retinoblastoma binding protein 7, chromatin remodeling factor [Source:MGI Symbol;Acc:MGI:1194910] |
| ENSMUSG00000032902 | Slc16a1 | 0.959438 | 746.4712 | 0.191699 | -5.00492 | 5.59E-07 | 7.77E-06 | 9.41E-06 | 845.1438 | 1126.167 | 467.2859 | 547.2883 | 20501 | solute carrier family 16 (monocarboxylic acid transporters), member 1 [Source:MGI Symbol;Acc:MGI:106013] |
| ENSMUSG00000023909 | Paqr4 | 0.961197 | 248.6367 | 0.235548 | -4.08069 | 4.49E-05 | 0.000415 | 0.000503 | 336.3651 | 320.6379 | 163.6872 | 173.8566 | 76498 | progestin and adipoQ receptor family member IV [Source:MGI Symbol;Acc:MGI:1923748] |
| ENSMUSG00000115230 | AU022793;AU0 | 0.962964 | 127.859 | 0.357295 | -2.69515 | 0.007036 | 0.031883 | 0.038519 | 167.1248 | 171.138 | 104.2477 | 68.9254 | 102636079;10 | expressed sequence AU022793 [Source:MGI Symbol;Acc:MGI:2146144];expressed sequence AU022793 [Source:MGI Symbol;Acc:MGI:2146144] |
| ENSMUSG00000029207 | Apbb2 | 0.964354 | 757.6628 | 0.196206 | -4.915 | 8.88E-07 | 1.19E-05 | 1.44E-05 | 1171.989 | 832.0847 | 527.6399 | 498.9376 | 11787 | amyloid beta precursor protein binding family B member 2 [Source:MGI Symbol;Acc:MGI:108405] |
| ENSMUSG00000035104 | Eva1a | 0.965107 | 160.551 | 0.304893 | -3.1654 | 0.001549 | 0.00906 | 0.010975 | 236.9364 | 187.8584 | 112.4778 | 104.9312 | 232146 | eva-1 homolog A, regulator of programmed cell death [Source:MGI Symbol;Acc:MGI:2385247] |
| ENSMUSG00000070476 | Fam217b | 0.965784 | 463.9947 | 0.194783 | -4.95825 | 7.11E-07 | 9.66E-06 | 1.17E-05 | 673.788 | 553.7396 | 305.4276 | 323.0235 | 71532 | family with sequence similarity 217, member B [Source:MGI Symbol;Acc:MGI:1918782] |
| ENSMUSG00000035376 | Hacd2 | 0.967847 | 426.821 | 0.216512 | -4.47017 | 7.82E-06 | 8.61E-05 | 0.000104 | 621.9581 | 507.5127 | 257.876 | 319.9373 | 70757 | 3-hydroxyacyl-CoA dehydratase 2 [Source:MGI Symbol;Acc:MGI:1918007] |
| ENSMUSG00000058013 | Septin11 | 0.967882 | 681.7298 | 0.158862 | -6.09258 | 1.11E-09 | 2.48E-08 | 3E-08 | 962.5542 | 841.9202 | 455.398 | 467.0468 | 52398 | septin 11 [Source:MGI Symbol;Acc:MGI:1277214] |
| ENSMUSG00000023104 | Rfc2 | 0.967937 | 699.2304 | 0.151359 | -6.39499 | 1.61E-10 | 4.06E-09 | 4.92E-09 | 962.5542 | 888.1472 | 462.7136 | 483.5066 | 19718 | replication factor C (activator 1) 2 [Source:MGI Symbol;Acc:MGI:1341868] |
| ENSMUSG00000026305 | Lrrfip1 | 0.970078 | 1523.678 | 0.116837 | -8.30281 | 1.02E-16 | 5.17E-15 | 6.27E-15 | 2126.081 | 1909.074 | 1028.761 | 1030.795 | 16978 | leucine rich repeat (in FLII) interacting protein 1 [Source:MGI Symbol;Acc:MGI:1342770] |
| ENSMUSG00000028741 | Mrto4 | 0.970221 | 300.8973 | 0.222553 | -4.35951 | 1.3E-05 | 0.000137 | 0.000166 | 422.043 | 374.7332 | 193.8642 | 212.9486 | 69902 | mRNA turnover 4, ribosome maturation factor [Source:MGI Symbol;Acc:MGI:1917152] |
| ENSMUSG00000015176 | Nolc1 | 0.97025 | 571.7772 | 0.190043 | -5.10541 | 3.3E-07 | 4.8E-06 | 5.82E-06 | 851.4903 | 662.9138 | 385.8995 | 386.8052 | 70769 | nucleolar and coiled-body phosphoprotein 1 [Source:MGI Symbol;Acc:MGI:1918019] |
| ENSMUSG00000074505 | Fat3 | 0.973063 | 3843.168 | 0.118147 | -8.23607 | 1.78E-16 | 8.76E-15 | 1.06E-14 | 5616.663 | 4568.598 | 2687.58 | 2499.832 | 270120 | FAT atypical cadherin 3 [Source:MGI Symbol;Acc:MGI:2444314] |
| ENSMUSG00000038047 | Haus6 | 0.974389 | 374.8597 | 0.221973 | -4.38967 | 1.14E-05 | 0.000121 | 0.000146 | 533.107 | 460.3022 | 221.1267 | 282.9028 | 230376 | HAUS augmin-like complex, subunit 6 [Source:MGI Symbol;Acc:MGI:1923389] |
| ENSMUSG00000020399 | Havcr2 | 0.97458 | 918.3282 | 0.141916 | -6.86719 | 6.55E-12 | 2E-10 | 2.42E-10 | 1295.746 | 1138.953 | 625.4864 | 613.1275 | 171285 | hepatitis A virus cellular receptor 2 [Source:MGI Symbol;Acc:MGI:2159682] |
| ENSMUSG00000035198 | Tubg1 | 0.976247 | 244.2587 | 0.242607 | -4.02399 | 5.72E-05 | 0.000512 | 0.00062 | 345.8849 | 301.9504 | 164.6017 | 164.598 | 103733 | tubulin, gamma 1 [Source:MGI Symbol;Acc:MGI:101834] |
| ENSMUSG00000037458 | Azin1 | 0.978425 | 2196.492 | 0.122341 | -7.99749 | 1.27E-15 | 5.83E-14 | 7.07E-14 | 3038.921 | 2788.369 | 1349.734 | 1608.945 | 54375 | antizyme inhibitor 1 [Source:MGI Symbol;Acc:MGI:1859169] |
| ENSMUSG00000018417 | Myo1b | 0.979346 | 224.1395 | 0.286644 | -3.4166 | 0.000634 | 0.004251 | 0.005149 | 350.1159 | 244.9044 | 155.4571 | 146.0807 | 17912 | myosin IB [Source:MGI Symbol;Acc:MGI:107752] |
| ENSMUSG00000044934 | Zfp367 | 0.981595 | 480.9264 | 0.22317 | -4.39855 | 1.09E-05 | 0.000117 | 0.000141 | 731.9643 | 544.8876 | 289.8819 | 356.9719 | 238673 | zinc finger protein 367 [Source:MGI Symbol;Acc:MGI:2442266] |
| ENSMUSG00000057561 | Eif1a | 0.981672 | 2048.662 | 0.09745 | -10.0736 | 7.23E-24 | 8.3E-22 | 1.01E-21 | 2761.79 | 2678.211 | 1377.167 | 1377.479 | 13664 | eukaryotic translation initiation factor 1A [Source:MGI Symbol;Acc:MGI:95298] |
| ENSMUSG00000038028 | Tigar | 0.982216 | 231.4696 | 0.285474 | -3.44065 | 0.00058 | 0.003941 | 0.004773 | 341.6539 | 273.4274 | 180.1474 | 130.6496 | 319801 | Trp53 induced glycolysis regulatory phosphatase [Source:MGI Symbol;Acc:MGI:2442752] |
| ENSMUSG00000006589 | Aprt | 0.982743 | 959.1918 | 0.137224 | -7.16158 | 7.98E-13 | 2.68E-11 | 3.25E-11 | 1340.172 | 1207.801 | 660.2357 | 628.5585 | 11821 | adenine phosphoribosyl transferase [Source:MGI Symbol;Acc:MGI:88061] |
| ENSMUSG00000027580 | Helz2 | 0.982916 | 249.4169 | 0.275912 | -3.56242 | 0.000367 | 0.002644 | 0.003203 | 363.8667 | 298.9997 | 162.8465 | 140.937 | 229003 | helicase with zinc finger 2, transcriptional coactivator [Source:MGI Symbol;Acc:MGI:2385169] |
| ENSMUSG00000009575 | Cbx5 | 0.983821 | 3351.496 | 0.111598 | -8.81578 | 1.19E-18 | 7.36E-17 | 8.91E-17 | 4850.85 | 4053.217 | 2221.208 | 2280.711 | 12419 | chromobox 5 [Source:MGI Symbol;Acc:MGI:109372] |
| ENSMUSG00000004642 | Slbp | 0.985122 | 895.1194 | 0.154142 | -6.391 | 1.65E-10 | 4.16E-09 | 5.04E-09 | 1297.862 | 1080.923 | 583.4215 | 618.2711 | 20492 | stem-loop binding protein [Source:MGI Symbol;Acc:MGI:108402] |
| ENSMUSG00000037601 | Nme1 | 0.988481 | 890.8462 | 0.135652 | -7.2869 | 3.17E-13 | 1.13E-11 | 1.37E-11 | 1200.548 | 1168.459 | 576.1059 | 618.2711 | 18102 | NME/NM23 nucleoside diphosphate kinase 1 [Source:MGI Symbol;Acc:MGI:97355] |
| ENSMUSG00000029610 | Aimp2 | 0.989358 | 188.8962 | 0.283023 | -3.49568 | 0.000473 | 0.003297 | 0.003993 | 279.2465 | 223.2662 | 123.4513 | 129.6209 | 231872 | aminoacyl tRNA synthetase complex-interacting multifunctional protein 2 [Source:MGI Symbol;Acc:MGI:2385237] |
| ENSMUSG00000093938 | Evi2b;Evi2b | 0.991808 | 297.7283 | 0.233982 | -4.23883 | 2.25E-05 | 0.000221 | 0.000268 | 407.2345 | 385.5523 | 222.2123 | 175.9141 | 216984;10148[illegible] | [illegible] ecotropic viral integration site 2b [Source:MGI Symbol;Acc:MGI:1890682];ecotropic viral integration site 2b [Source:MGI Symbol;Acc:MGI:1890682] |
| ENSMUSG00000032279 | Idh3a | 0.99279 | 551.8343 | 0.181378 | -5.47361 | 4.41E-08 | 7.74E-07 | 9.37E-07 | 776.3899 | 692.4204 | 338.3479 | 400.1788 | 67834 | isocitrate dehydrogenase 3 (NAD+) alpha [Source:MGI Symbol;Acc:MGI:1915084] |
| ENSMUSG00000024640 | Psat1 | 0.992979 | 1963.236 | 0.138652 | -7.16165 | 7.97E-13 | 2.68E-11 | 3.25E-11 | 2887.663 | 2338.886 | 1226.283 | 1400.112 | 107272 | phosphoserine aminotransferase 1 [Source:MGI Symbol;Acc:MGI:2183441] |
| ENSMUSG00000021556 | Golm1 | 0.993193 | 2999.095 | 0.10953 | -9.06775 | 1.21E-19 | 8.65E-18 | 1.05E-17 | 4294.473 | 3691.27 | 2069.409 | 1941.227 | 105348 | golgi membrane protein 1 [Source:MGI Symbol;Acc:MGI:1917329] |
| ENSMUSG00000026281 | Dtymk | 0.994616 | 262.4734 | 0.236765 | -4.20086 | 2.66E-05 | 0.000258 | 0.000313 | 375.5019 | 323.5885 | 172.8318 | 177.9716 | 21915 | deoxythymidylate kinase [Source:MGI Symbol;Acc:MGI:108396] |
| ENSMUSG00000037725 | Ckap2 | 0.995692 | 405.5722 | 0.232656 | -4.27968 | 1.87E-05 | 0.000188 | 0.000228 | 623.0159 | 457.3515 | 248.7314 | 293.1901 | 80986 | cytoskeleton associated protein 2 [Source:MGI Symbol;Acc:MGI:1931797] |
| ENSMUSG00000030104 | Edem1 | 0.996407 | 6283.091 | 0.11048 | -9.01893 | 1.9E-19 | 1.33E-17 | 1.62E-17 | 9210.964 | 7530.072 | 4050.116 | 4341.272 | 192193 | ER degradation enhancer, mannosidase alpha-like 1 [Source:MGI Symbol;Acc:MGI:2180139] |
| ENSMUSG00000026020 | Nop58 | 0.997533 | 577.7716 | 0.185423 | -5.37978 | 7.46E-08 | 1.26E-06 | 1.53E-06 | 848.317 | 691.4368 | 363.9526 | 407.38 | 55989 | NOP58 ribonucleoprotein [Source:MGI Symbol;Acc:MGI:1933184] |
| ENSMUSG00000063229 | Ldha | 0.998028 | 5729.975 | 0.099939 | -9.98634 | 1.75E-23 | 1.85E-21 | 2.25E-21 | 8251.523 | 7021.576 | 3752.004 | 3894.8 | 16828 | lactate dehydrogenase A [Source:MGI Symbol;Acc:MGI:96759] |
| ENSMUSG00000026249 | Serpine2 | 0.998623 | 543.0093 | 0.166478 | -5.99854 | 1.99E-09 | 4.29E-08 | 5.19E-08 | 708.6938 | 738.6473 | 350.2358 | 374.4604 | 20720 | serine (or cysteine) peptidase inhibitor, clade E, member 2 [Source:MGI Symbol;Acc:MGI:101780] |
| ENSMUSG00000057469 | E2f6 | 1.003074 | 430.091 | 0.201504 | -4.97794 | 6.43E-07 | 8.79E-06 | 1.07E-05 | 629.3624 | 518.3317 | 274.3361 | 298.3338 | 50496 | E2F transcription factor 6 [Source:MGI Symbol;Acc:MGI:1354159] |
| ENSMUSG00000060981 | H4c8 | 1.003768 | 2565.703 | 0.131293 | -7.64524 | 2.09E-14 | 8.37E-13 | 1.01E-12 | 3734.922 | 3113.925 | 1847.197 | 1566.767 | 69386 | H4 clustered histone 8 [Source:MGI Symbol;Acc:MGI:2448427] |
| ENSMUSG00000034168 | Irf2bpl | 1.00383 | 517.6671 | 0.171101 | -5.86688 | 4.44E-09 | 9.08E-08 | 1.1E-07 | 680.1345 | 701.2724 | 329.2034 | 360.0581 | 238330 | interferon regulatory factor 2 binding protein-like [Source:MGI Symbol;Acc:MGI:2442463] |
| ENSMUSG00000002477 | Snrpd1 | 1.007019 | 549.4459 | 0.180952 | -5.5651 | 2.62E-08 | 4.77E-07 | 5.78E-07 | 802.8337 | 664.8809 | 364.8671 | 365.2018 | 20641 | small nuclear ribonucleoprotein D1 [Source:MGI Symbol;Acc:MGI:98344] |
| ENSMUSG00000052833 | Sae1 | 1.008731 | 1249.74 | 0.134019 | -7.52678 | 5.2E-14 | 2.01E-12 | 2.43E-12 | 1802.409 | 1537.291 | 840.3831 | 818.8749 | 56459 | SUMO1 activating enzyme subunit 1 [Source:MGI Symbol;Acc:MGI:1929264] |
| ENSMUSG00000052798 | Nup107 | 1.01144 | 403.4025 | 0.20879 | -4.84429 | 1.27E-06 | 1.64E-05 | 1.99E-05 | 598.6876 | 479.9732 | 263.3627 | 271.5867 | 103468 | nucleoporin 107 [Source:MGI Symbol;Acc:MGI:2143854] |
| ENSMUSG00000054079 | Utp18 | 1.0126 | 447.2485 | 0.193182 | -5.24168 | 1.59E-07 | 2.48E-06 | 3.01E-06 | 646.2864 | 549.8054 | 285.3096 | 307.5925 | 217109 | UTP18 small subunit processome component [Source:MGI Symbol;Acc:MGI:1923402] |
| ENSMUSG00000003814 | Calr | 1.012622 | 15620.31 | 0.063111 | -16.0452 | 6.18E-58 | 4.68E-54 | 5.67E-54 | 21053.49 | 20722.45 | 10454.95 | 10250.34 | 12317 | calreticulin [Source:MGI Symbol;Acc:MGI:88252] |
| ENSMUSG00000034641 | Cd300ld | 1.016066 | 176.5118 | 0.309649 | -3.28134 | 0.001033 | 0.006473 | 0.007842 | 273.9577 | 198.6774 | 124.3657 | 109.0462 | 217305 | CD300 molecule like family member d [Source:MGI Symbol;Acc:MGI:2442358] |
| ENSMUSG00000037278 | Tmem97 | 1.017176 | 411.277 | 0.202657 | -5.0192 | 5.19E-07 | 7.26E-06 | 8.8E-06 | 599.7453 | 501.6114 | 283.4807 | 260.2706 | 69071 | transmembrane protein 97 [Source:MGI Symbol;Acc:MGI:1916321] |
| ENSMUSG00000001056 | Nhp2 | 1.017662 | 400.6313 | 0.211924 | -4.80202 | 1.57E-06 | 1.99E-05 | 2.41E-05 | 599.7453 | 473.0884 | 264.2772 | 265.4142 | 52530 | NHP2 ribonucleoprotein [Source:MGI Symbol;Acc:MGI:1098547] |
| ENSMUSG00000039774 | Galnt12 | 1.017763 | 341.6516 | 0.223181 | -4.56026 | 5.11E-06 | 5.83E-05 | 7.07E-05 | 508.7787 | 406.2068 | 230.4424 | 221.1785 | 230145 | polypeptide N-acetylgalactosaminyltransferase 12 [Source:MGI Symbol;Acc:MGI:2444664] |
| ENSMUSG00000023953 | Polh | 1.018352 | 498.3503 | 0.187007 | -5.44554 | 5.16E-08 | 8.97E-07 | 1.09E-06 | 728.7911 | 605.8678 | 326.46 | 332.2822 | 80905 | polymerase (DNA directed), eta (RAD 30 related) [Source:MGI Symbol;Acc:MGI:1891457] |
| ENSMUSG00000062937 | Mtap | 1.018363 | 441.9815 | 0.20026 | -5.08519 | 3.67E-07 | 5.29E-06 | 6.41E-06 | 653.6907 | 530.1344 | 298.1119 | 285.989 | 66902 | methylthioadenosine phosphorylase [Source:MGI Symbol;Acc:MGI:1914152] |
| ENSMUSG00000044783 | Hjurp;Hjurp | 1.02417 | 1222.934 | 0.129074 | -7.93478 | 2.11E-15 | 9.38E-14 | 1.14E-13 | 1733.655 | 1546.143 | 832.153 | 779.7829 | 212427;38128 | Holliday junction recognition protein [Source:MGI Symbol;Acc:MGI:2685821];Holliday junction recognition protein [Source:MGI Symbol;Acc:MGI:2685821] |
| ENSMUSG00000021811 | Dnajc9 | 1.027722 | 482.4424 | 0.203911 | -5.04004 | 4.65E-07 | 6.59E-06 | 7.98E-06 | 718.2135 | 576.3613 | 292.6252 | 342.5695 | 108671 | DnaJ heat shock protein family (Hsp40) member C9 [Source:MGI Symbol;Acc:MGI:1915326] |
| ENSMUSG00000027490 | E2f1 | 1.028233 | 201.7538 | 0.258642 | -3.97551 | 7.02E-05 | 0.00061 | 0.000738 | 273.9577 | 267.5261 | 130.7669 | 134.7646 | 13555 | E2F transcription factor 1 [Source:MGI Symbol;Acc:MGI:101941] |
| ENSMUSG00000027597 | Ahcy | 1.030647 | 751.4248 | 0.161683 | -6.37451 | 1.84E-10 | 4.57E-09 | 5.54E-09 | 1094.773 | 923.555 | 512.0941 | 475.2767 | 269378 | S-adenosylhomocysteine hydrolase [Source:MGI Symbol;Acc:MGI:87968] |
| ENSMUSG00000023456 | Tpi1 | 1.030782 | 3926.479 | 0.118697 | -8.68417 | 3.82E-18 | 2.23E-16 | 2.7E-16 | 5853.599 | 4691.542 | 2563.214 | 2597.562 | 21991 | triosephosphate isomerase 1 [Source:MGI Symbol;Acc:MGI:98797] |
| ENSMUSG00000029552 | Tes | 1.031768 | 1084.405 | 0.150223 | -6.86825 | 6.5E-12 | 1.99E-10 | 2.41E-10 | 1608.841 | 1304.19 | 701.3861 | 723.2024 | 21753 | testin LIM domain protein [Source:MGI Symbol;Acc:MGI:105081] |
| ENSMUSG00000059743 | Fdps | 1.032868 | 496.0415 | 0.18603 | -5.55216 | 2.82E-08 | 5.14E-07 | 6.22E-07 | 720.3291 | 612.7527 | 338.3479 | 312.7362 | 110196 | farnesyl diphosphate synthetase [Source:MGI Symbol;Acc:MGI:104888] |
| ENSMUSG00000029915 | Clec5a | 1.035505 | 720.8726 | 0.166934 | -6.20309 | 5.54E-10 | 1.29E-08 | 1.56E-08 | 1056.694 | 881.2623 | 451.7402 | 493.7939 | 23845 | C-type lectin domain family 5, member a [Source:MGI Symbol;Acc:MGI:1345151] |
| ENSMUSG00000030707 | Coro1a | 1.0357 | 5756.909 | 0.100218 | -10.3345 | 4.92E-25 | 6.27E-23 | 7.6E-23 | 8297.006 | 7181.894 | 3935.809 | 3612.926 | 12721 | coronin, actin binding protein 1A [Source:MGI Symbol;Acc:MGI:1345961] |
| ENSMUSG00000038843 | Gcnt1 | 1.035903 | 2852.897 | 0.13219 | -7.83648 | 4.63E-15 | 2.02E-13 | 2.45E-13 | 4287.069 | 3383.418 | 1781.356 | 1959.745 | 14537 | glucosaminyl (N-acetyl) transferase 1, core 2 [Source:MGI Symbol;Acc:MGI:95676] |
| ENSMUSG00000024660 | Incenp | 1.036528 | 931.3073 | 0.163039 | -6.35756 | 2.05E-10 | 5.05E-09 | 6.12E-09 | 1397.29 | 1107.479 | 630.9731 | 589.4665 | 16319 | inner centromere protein [Source:MGI Symbol;Acc:MGI:1313288] |
| ENSMUSG00000021532 | Fastkd3 | 1.037148 | 380.9782 | 0.225333 | -4.60274 | 4.17E-06 | 4.82E-05 | 5.84E-05 | 548.9732 | 421.9437 | 232.2713 | 240.7245 | 69577 | FAST kinase domains 3 [Source:MGI Symbol;Acc:MGI:1916827] |
| ENSMUSG00000074813 | Morrbid | 1.037637 | 108.745 | 0.383693 | -2.70434 | 0.006844 | 0.031136 | 0.037616 | 174.5291 | 118.0262 | 70.41294 | 72.01161 | | myeloid RNA regulator of BCL2L11 induced cell death [Source:MGI Symbol;Acc:MGI:3652191] |
| ENSMUSG00000011752 | Pgam1 | 1.038843 | 5204.352 | 0.103983 | -9.99048 | 1.68E-23 | 1.8E-21 | 2.19E-21 | 7607.352 | 6395.053 | 3345.072 | 3469.931 | 18648 | phosphoglycerate mutase 1 [Source:MGI Symbol;Acc:MGI:97552] |
| ENSMUSG00000026864 | Hspa5 | 1.039285 | 16662.62 | 0.083078 | -12.5097 | 6.61E-36 | 2.71E-33 | 3.28E-33 | 23747.59 | 21087.35 | 10617.72 | 11197.81 | 14828 | heat shock protein 5 [Source:MGI Symbol;Acc:MGI:95835] |
| ENSMUSG00000006732 | Mettl1 | 1.041161 | 83.41626 | 0.399714 | -2.60477 | 0.009194 | 0.039747 | 0.048012 | 117.4105 | 107.2071 | 57.61059 | 51.43687 | 17299 | methyltransferase 1, tRNA methylguanosine [Source:MGI Symbol;Acc:MGI:1339986] |
| ENSMUSG00000020116 | Pno1 | 1.042495 | 529.0083 | 0.181957 | -5.72935 | 1.01E-08 | 1.97E-07 | 2.38E-07 | 742.5418 | 681.6013 | 316.401 | 375.4891 | 66249 | partner of NOB1 homolog [Source:MGI Symbol;Acc:MGI:1913499] |
| ENSMUSG00000070284 | Gmppb | 1.042943 | 299.9804 | 0.239833 | -4.34861 | 1.37E-05 | 0.000143 | 0.000173 | 453.7756 | 354.0786 | 188.3775 | 203.69 | 331026 | GDP-mannose pyrophosphorylase B [Source:MGI Symbol;Acc:MGI:2660880] |
| ENSMUSG00000096054 | Syne1 | 1.044255 | 917.6773 | 0.144398 | -7.23178 | 4.77E-13 | 1.64E-11 | 2E-11 | 1218.53 | 1254.028 | 644.6899 | 553.4607 | 64009 | spectrin repeat containing, nuclear envelope 1 [Source:MGI Symbol;Acc:MGI:1927152] |
| ENSMUSG00000030512 | Snrpa1 | 1.044355 | 463.4601 | 0.191881 | -5.44272 | 5.25E-08 | 9.1E-07 | 1.1E-06 | 678.019 | 570.46 | 292.6252 | 312.7362 | 68981 | small nuclear ribonucleoprotein polypeptide A' [Source:MGI Symbol;Acc:MGI:1916231] |
| ENSMUSG00000034424 | Gcsh | 1.044565 | 466.3645 | 0.177225 | -5.894 | 3.77E-09 | 7.83E-08 | 9.49E-08 | 638.8822 | 617.6705 | 313.6577 | 295.2476 | 68133 | glycine cleavage system protein H (aminomethyl carrier) [Source:MGI Symbol;Acc:MGI:1915383] |
| ENSMUSG00000055148 | Klf2 | 1.04509 | 275.7852 | 0.302853 | -3.45081 | 0.000559 | 0.003816 | 0.004622 | 406.1767 | 337.3582 | 225.8701 | 133.7359 | 16598 | Kruppel-like transcription factor 2 (lung) [Source:MGI Symbol;Acc:MGI:1342772] |
| ENSMUSG00000028896 | Rcc1 | [illegible] | [illegible] | [illegible] | [illegible] | [illegible] | [illegible] | [illegible] | [illegible] | [illegible] | [illegible] | [illegible] | [illegible] | regulator of chromosome condensation 1 [illegible] |
| ENSMUSG00000141779 | Gm70496 | 1.047397 | 214.1458 | 0.286936 | -3.65027 | 0.000262 | 0.001963 | 0.002378 | 336.3651 | 241.9702 | 136.2536 | 142.9445 | | predicted gene, 70496 [Source:MGI Symbol;Acc:MGI:7823791] |
| ENSMUSG00000053063 | Clec12a | 1.047502 | 1172.993 | 0.159648 | -6.56132 | 5.33E-11 | 1.43E-09 | 1.74E-09 | 1762.215 | 1400.578 | 819.3506 | 708.8288 | 232413 | C-type lectin domain family 12, member a [Source:MGI Symbol;Acc:MGI:3040968] |
| ENSMUSG00000002297 | Dbf4 | 1.047624 | 817.5729 | 0.145831 | -7.1838 | 6.78E-13 | 2.32E-11 | 2.81E-11 | 1162.469 | 1041.581 | 523.0676 | 543.1733 | 27214 | DBF4 zinc finger [Source:MGI Symbol;Acc:MGI:1351328] |
| ENSMUSG00000040269 | Mrps28 | 1.048497 | 216.2213 | 0.285076 | -3.67796 | 0.000235 | 0.001782 | 0.002159 | 336.3651 | 246.8715 | 149.9704 | 131.6784 | 66230 | mitochondrial ribosomal protein S28 [Source:MGI Symbol;Acc:MGI:1913480] |
| ENSMUSG00000055760 | Gemin6 | 1.050631 | 113.5405 | 0.364344 | -2.88362 | 0.003931 | 0.019706 | 0.023842 | 173.4713 | 132.7795 | 67.66958 | 80.24151 | 67242 | gem nuclear organelle associated protein 6 [Source:MGI Symbol;Acc:MGI:1914492] |
| ENSMUSG00000018189 | Uchl5 | 1.051218 | 509.0528 | 0.185004 | -5.68212 | 1.33E-08 | 2.55E-07 | 3.09E-07 | 731.9643 | 641.2757 | 309.0854 | 353.8856 | 56207 | ubiquitin carboxyl-terminal esterase L5 [Source:MGI Symbol;Acc:MGI:1914848] |
| ENSMUSG00000022369 | Mtbp | 1.052065 | 150.5381 | 0.321418 | -3.2732 | 0.001063 | 0.006611 | 0.008008 | 199.9151 | 206.5459 | 113.3923 | 82.29899 | 105837 | Mdm2, transformed 3T3 cell double minute p53 binding protein [Source:MGI Symbol;Acc:MGI:2146005] |
| ENSMUSG00000024436 | Mrps18b | 1.053018 | 331.298 | 0.215935 | -4.87655 | 1.08E-06 | 1.42E-05 | 1.72E-05 | 478.1039 | 416.0424 | 205.7521 | 225.2935 | 66973 | mitochondrial ribosomal protein S18B [Source:MGI Symbol;Acc:MGI:1914223] |
| ENSMUSG00000029246 | Ppat | 1.054365 | 403.0491 | 0.217476 | -4.84818 | 1.25E-06 | 1.61E-05 | 1.95E-05 | 607.1496 | 480.9568 | 243.2447 | 280.8453 | 231327 | phosphoribosyl pyrophosphate amidotransferase [Source:MGI Symbol;Acc:MGI:2387203] |
| ENSMUSG00000055541 | Lair1 | 1.056842 | 1813.114 | 0.118317 | -8.93227 | 4.17E-19 | 2.78E-17 | 3.36E-17 | 2622.167 | 2275.939 | 1155.87 | 1198.479 | 52855 | leukocyte-associated Ig-like receptor 1 [Source:MGI Symbol;Acc:MGI:105492] |
| ENSMUSG00000031438 | Rnf128 | 1.058592 | 1230.15 | 0.125239 | -8.45255 | 2.85E-17 | 1.54E-15 | 1.87E-15 | 1706.154 | 1617.943 | 758.0822 | 838.4209 | 66889 | ring finger protein 128 [Source:MGI Symbol;Acc:MGI:1914139] |
| ENSMUSG00000038685 | Rtel1 | 1.059699 | 290.4886 | 0.25053 | -4.22983 | 2.34E-05 | 0.00023 | 0.000278 | 417.812 | 367.8483 | 215.8111 | 160.483 | 269400 | regulator of telomere elongation helicase 1 [Source:MGI Symbol;Acc:MGI:2139369] |
| ENSMUSG00000112478 | Gm47761 | 1.062931 | 112.1403 | 0.34658 | -3.06691 | 0.002163 | 0.0119 | 0.014415 | 160.7783 | 142.615 | 73.15631 | 72.01161 | | predicted gene, 47761 [Source:MGI Symbol;Acc:MGI:6096912] |
| ENSMUSG00000040034 | Nup43 | 1.067662 | 150.3758 | 0.312949 | -3.41162 | 0.000646 | 0.004315 | 0.005227 | 224.2434 | 182.9406 | 91.44538 | 102.8737 | 69912 | nucleoporin 43 [Source:MGI Symbol;Acc:MGI:1917162] |
| ENSMUSG00000015217 | Hmgb3 | 1.06991 | 172.5823 | 0.293314 | -3.64766 | 0.000265 | 0.001978 | 0.002396 | 259.1492 | 208.513 | 111.5634 | 111.1036 | 15354 | high mobility group box 3 [Source:MGI Symbol;Acc:MGI:1098219] |
| ENSMUSG00000035842 | Ddx11 | 1.073333 | 300.3116 | 0.249645 | -4.29944 | 1.71E-05 | 0.000174 | 0.000211 | 452.7178 | 361.947 | 215.8111 | 170.7704 | 320209 | DEAD/H box helicase 11 [Source:MGI Symbol;Acc:MGI:2443590] |
| ENSMUSG00000042354 | Gnl3 | 1.073744 | 910.9149 | 0.161073 | -6.66618 | 2.63E-11 | 7.33E-10 | 8.88E-10 | 1365.558 | 1104.529 | 561.4746 | 612.0987 | 30877 | guanine nucleotide binding protein nucleolar 3 [Source:MGI Symbol;Acc:MGI:1353651] |
| ENSMUSG00000061458 | Nol10 | 1.074397 | 137.6312 | 0.332645 | -3.22986 | 0.001239 | 0.007512 | 0.0091 | 206.2616 | 167.2038 | 97.84656 | 79.21278 | 217431 | nucleolar protein 10 [Source:MGI Symbol;Acc:MGI:2684913] |
| ENSMUSG00000020024 | Cep83 | 1.075043 | 548.8759 | 0.174059 | -6.17632 | 6.56E-10 | 1.52E-08 | 1.84E-08 | 797.5449 | 691.4368 | 355.7225 | 350.7994 | 77048 | centrosomal protein 83 [Source:MGI Symbol;Acc:MGI:1924298] |
| ENSMUSG00000057666 | Gapdh | 1.078227 | 9696.398 | 0.092987 | -11.5955 | 4.34E-31 | 1.06E-28 | 1.29E-28 | 14095.6 | 12225.55 | 6443.242 | 6021.2 | 14433 | glyceraldehyde-3-phosphate dehydrogenase [Source:MGI Symbol;Acc:MGI:95640] |
| ENSMUSG00000000489 | Pdgfb | 1.080494 | 246.4716 | 0.250673 | -4.31037 | 1.63E-05 | 0.000167 | 0.000203 | 361.7512 | 307.8517 | 169.174 | 147.1094 | 18591 | platelet derived growth factor, B polypeptide [Source:MGI Symbol;Acc:MGI:97528] |
| ENSMUSG00000001227 | Sema6b | 1.084326 | 162.3321 | 0.29151 | -3.71968 | 0.000199 | 0.001536 | 0.001861 | 233.7632 | 207.5294 | 105.1622 | 102.8737 | 20359 | sema domain, transmembrane domain (TM), and cytoplasmic domain, (semaphorin) 6B [Source:MGI Symbol;Acc:MGI:1202889] |
| ENSMUSG00000074802 | Gas2l3 | 1.086935 | 1626.331 | 0.106886 | -10.1682 | 2.75E-24 | 3.23E-22 | 3.92E-22 | 2257.243 | 2165.781 | 1027.846 | 1054.456 | 237436 | growth arrest-specific 2 like 3 [Source:MGI Symbol;Acc:MGI:1918780] |
| ENSMUSG00000032666 | 1700025G04Rik | 1.088412 | 343.0301 | 0.212678 | -5.11765 | 3.09E-07 | 4.53E-06 | 5.49E-06 | 503.4899 | 429.8121 | 217.64 | 221.1785 | 69399 | RIKEN cDNA 1700025G04 gene [Source:MGI Symbol;Acc:MGI:1916649] |
| ENSMUSG00000001542 | Ell2 | 1.088737 | 1103.595 | 0.154595 | -7.0425 | 1.89E-12 | 6.12E-11 | 7.41E-11 | 1672.306 | 1330.745 | 725.1619 | 686.1678 | 192657 | elongation factor for RNA polymerase II 2 [Source:MGI Symbol;Acc:MGI:2183438] |
| ENSMUSG00000033970 | Rfc3 | 1.089205 | 311.4718 | 0.226423 | -4.8105 | 1.51E-06 | 1.92E-05 | 2.32E-05 | 456.9488 | 390.47 | 186.5486 | 211.9199 | 69263 | replication factor C (activator 1) 3 [Source:MGI Symbol;Acc:MGI:1916513] |
| ENSMUSG00000057069 | Ero1b | 1.090099 | 344.9157 | 0.22652 | -4.81236 | 1.49E-06 | 1.9E-05 | 2.31E-05 | 502.4322 | 436.697 | 245.0736 | 195.4601 | 67475 | endoplasmic reticulum oxidoreductase 1 beta [Source:MGI Symbol;Acc:MGI:1914725] |
| ENSMUSG00000038005 | Hpf1 | 1.090421 | 469.3301 | 0.219943 | -4.95773 | 7.13E-07 | 9.68E-06 | 1.17E-05 | 740.4263 | 537.0192 | 287.1385 | 312.7362 | 72612 | histone PARylation factor 1 [Source:MGI Symbol;Acc:MGI:1919862] |
| ENSMUSG00000029366 | Dck | 1.09474 | 654.0377 | 0.15403 | -7.1073 | 1.18E-12 | 3.93E-11 | 4.77E-11 | 920.2442 | 861.5913 | 411.5042 | 422.8111 | 13178 | deoxycytidine kinase [Source:MGI Symbol;Acc:MGI:102726] |
| ENSMUSG00000031328 | Flna | 1.094758 | 10695.88 | 0.114641 | -9.54943 | 1.3E-21 | 1.13E-19 | 1.37E-19 | 16358.13 | 12782.24 | 6864.805 | 6778.35 | 192176 | filamin, alpha [Source:MGI Symbol;Acc:MGI:95556] |
| ENSMUSG00000091228 | Nme1nme2 | 1.094878 | 633.8535 | 0.166776 | -6.56497 | 5.2E-11 | 1.41E-09 | 1.7E-09 | 901.2046 | 826.1834 | 432.5367 | 375.4891 | | NME/NM23 nucleoside diphosphate kinase 1 and 2 readthrough [Source:MGI Symbol;Acc:MGI:5141855] |
| ENSMUSG00000003153 | Slc2a3 | 1.100479 | 261.7319 | 0.229549 | -4.79409 | 1.63E-06 | 2.06E-05 | 2.49E-05 | 349.0581 | 364.8977 | 167.345 | 165.6267 | 20527 | solute carrier family 2 (facilitated glucose transporter), member 3 [Source:MGI Symbol;Acc:MGI:95757] |
| ENSMUSG00000020492 | Ska2 | 1.100504 | 231.8258 | 0.259896 | -4.2344 | 2.29E-05 | 0.000225 | 0.000273 | 350.1159 | 282.2793 | 142.6548 | 152.2531 | 66140 | spindle and kinetochore associated complex subunit 2 [Source:MGI Symbol;Acc:MGI:1913390] |
| ENSMUSG00000032939 | Nup93 | 1.105252 | 630.3077 | 0.172964 | -6.39008 | 1.66E-10 | 4.17E-09 | 5.05E-09 | 939.2837 | 781.9236 | 389.5573 | 410.4662 | 71805 | nucleoporin 93 [Source:MGI Symbol;Acc:MGI:1919055] |
| ENSMUSG00000026708 | Cenpl | 1.107204 | 118.225 | 0.342025 | -3.2372 | 0.001207 | 0.007362 | 0.008919 | 174.5291 | 148.5163 | 76.81412 | 73.04035 | 70454 | centromere protein L [Source:MGI Symbol;Acc:MGI:1917704] |
| ENSMUSG00000074875 | Snord3a | 1.108677 | 204.8914 | 0.425435 | -2.60599 | 0.009161 | 0.039631 | 0.047872 | 276.0732 | 284.2464 | 189.2919 | 69.95414 | | small nucleolar RNA, C/D box 3A [Source:MGI Symbol;Acc:MGI:97977] |
| ENSMUSG00000022246 | Rai14 | 1.109929 | 594.2925 | 0.217679 | -5.09893 | 3.42E-07 | 4.95E-06 | 5.99E-06 | 946.688 | 677.6671 | 339.2624 | 413.5524 | 75646 | retinoic acid induced 14 [Source:MGI Symbol;Acc:MGI:1922896] |
| ENSMUSG00000050022 | Amz1 | 1.111617 | 258.5485 | 0.23126 | -4.80678 | 1.53E-06 | 1.95E-05 | 2.36E-05 | 362.8089 | 344.2431 | 164.6017 | 162.5405 | 231842 | archaelysin family metallopeptidase 1 [Source:MGI Symbol;Acc:MGI:2442258] |
| ENSMUSG00000026819 | Slc25a25 | 1.111765 | 447.4988 | 0.195744 | -5.6797 | 1.35E-08 | 2.58E-07 | 3.13E-07 | 660.0372 | 563.5751 | 267.0205 | 299.3626 | 227731 | solute carrier family 25 (mitochondrial carrier, phosphate carrier), member 25 [Source:MGI Symbol;Acc:MGI:1915913] |
| ENSMUSG00000013629 | Cad | 1.113753 | 558.8937 | 0.20761 | -5.36465 | 8.11E-08 | 1.36E-06 | 1.64E-06 | 870.5298 | 658.9796 | 384.0706 | 321.9948 | 69719 | carbamoyl-phosphate synthetase 2, aspartate transcarbamylase, and dihydroorotase [Source:MGI Symbol;Acc:MGI:1916969] |
| ENSMUSG00000096010 | H4c16 | 1.115337 | 513.7708 | 0.185417 | -6.01528 | 1.8E-09 | 3.88E-08 | 4.71E-08 | 743.5996 | 662.9138 | 350.2358 | 298.3338 | 320332 | H4 histone 16 [Source:MGI Symbol;Acc:MGI:2448443] |
| ENSMUSG00000022876 | Samsn1 | 1.115615 | 815.7392 | 0.142536 | -7.8269 | 5E-15 | 2.17E-13 | 2.63E-13 | 1164.585 | 1068.137 | 514.8375 | 515.3974 | 67742 | SAM domain, SH3 domain and nuclear localization signals, 1 [Source:MGI Symbol;Acc:MGI:1914992] |
| ENSMUSG00000025134 | Alyref | 1.118469 | 1062.915 | 0.127734 | -8.75622 | 2.02E-18 | 1.21E-16 | 1.47E-16 | 1509.412 | 1401.561 | 664.8079 | 675.8804 | 21681 | Aly/REF export factor [Source:MGI Symbol;Acc:MGI:1341044] |
| ENSMUSG00000064326 | Siva1 | 1.118618 | 172.7128 | 0.29881 | -3.74358 | 0.000181 | 0.001409 | 0.001706 | 266.5535 | 206.5459 | 109.7345 | 108.0174 | 30954 | SIVA1, apoptosis-inducing factor [Source:MGI Symbol;Acc:MGI:1353606] |
| ENSMUSG00000001918 | Slc1a5 | 1.12062 | 183.7607 | 0.270794 | -4.13827 | 3.5E-05 | 0.000332 | 0.000402 | 250.6872 | 252.7728 | 114.3067 | 117.2761 | 20514 | solute carrier family 1 (neutral amino acid transporter), member 5 [Source:MGI Symbol;Acc:MGI:105305] |
| ENSMUSG00000075254 | Heg1 | 1.121178 | 188.4821 | 0.358268 | -3.12944 | 0.001751 | 0.010014 | 0.01213 | 193.5686 | 322.605 | 97.84656 | 139.9083 | 77446 | heart development protein with EGF-like domains 1 [Source:MGI Symbol;Acc:MGI:1924696] |
| ENSMUSG00000032115 | Hyou1 | 1.121324 | 2571.436 | 0.099993 | -11.214 | 3.48E-29 | 6.36E-27 | 7.7E-27 | 3694.727 | 3351.944 | 1600.294 | 1638.779 | 12282 | hypoxia up-regulated 1 [Source:MGI Symbol;Acc:MGI:108030] |
| ENSMUSG00000015112 | Slc25a13 | 1.127093 | 303.0745 | 0.223999 | -5.03168 | 4.86E-07 | 6.85E-06 | 8.3E-06 | 444.2558 | 387.5194 | 198.4365 | 182.0865 | 50799 | solute carrier family 25 (mitochondrial carrier, adenine nucleotide translocator), member 13 [Source:MGI Symbol;Acc:MGI:1354721] |
| ENSMUSG00000020471 | Pold2 | 1.13076 | 166.2784 | 0.302654 | -3.73615 | 0.000187 | 0.001446 | 0.001752 | 255.976 | 200.6445 | 101.5044 | 106.9887 | 18972 | polymerase (DNA directed), delta 2, regulatory subunit [Source:MGI Symbol;Acc:MGI:1097163] |
| ENSMUSG00000027199 | Gatm | 1.131539 | 2478.858 | 0.142682 | -7.93048 | 2.18E-15 | 9.68E-14 | 1.17E-13 | 3863.968 | 2944.754 | 1622.241 | 1484.468 | 67092 | glycine amidinotransferase (L-arginine:glycine amidinotransferase) [Source:MGI Symbol;Acc:MGI:1914342] |
| ENSMUSG00000062075 | Lmnb2 | 1.131612 | 366.5166 | 0.202772 | -5.58072 | 2.4E-08 | 4.4E-07 | 5.33E-07 | 515.1252 | 491.7758 | 244.1592 | 215.0061 | 16907 | lamin B2 [Source:MGI Symbol;Acc:MGI:96796] |
| ENSMUSG00000027405 | Nop56 | 1.133127 | 1042.519 | 0.145135 | -7.80738 | 5.84E-15 | 2.51E-13 | 3.04E-13 | 1541.145 | 1322.877 | 620.9141 | 685.1391 | 67134 | NOP56 ribonucleoprotein [Source:MGI Symbol;Acc:MGI:1914384] |
| ENSMUSG00000028207 | Asph | 1.136673 | 2179.391 | 0.120458 | -9.43627 | 3.86E-21 | 3.23E-19 | 3.92E-19 | 3263.165 | 2729.356 | 1358.878 | 1366.163 | 65973 | aspartate-beta-hydroxylase [Source:MGI Symbol;Acc:MGI:1914186] |
| ENSMUSG00000046675 | Lyset | 1.141653 | 325.9489 | 0.216693 | -5.26853 | 1.38E-07 | 2.17E-06 | 2.63E-06 | 478.1039 | 418.993 | 194.7787 | 211.9199 | 320351 | lysosomal enzyme trafficking factor [Source:MGI Symbol;Acc:MGI:2443862] |
| ENSMUSG00000031530 | Dusp4 | 1.143625 | 203.3924 | 0.28089 | -4.07143 | 4.67E-05 | 0.000429 | 0.00052 | 314.1523 | 245.8879 | 119.7935 | 133.7359 | 319520 | dual specificity phosphatase 4 [Source:MGI Symbol;Acc:MGI:2442191] |
| ENSMUSG00000050107 | Haspin | 1.143768 | 163.3131 | 0.30183 | -3.78945 | 0.000151 | 0.001199 | 0.001453 | 244.3407 | 205.5623 | 109.7345 | 93.6151 | 14841 | histone H3 associated protein kinase [Source:MGI Symbol;Acc:MGI:1194498] |
| ENSMUSG00000060678 | H4c3 | 1.145444 | 2359.706 | 0.136703 | -8.37905 | 5.34E-17 | 2.82E-15 | 3.42E-15 | 3535.007 | 2966.392 | 1610.353 | 1327.071 | 319155 | H4 clustered histone 3 [Source:MGI Symbol;Acc:MGI:2448421] |
| ENSMUSG00000024525 | Impa2 | 1.145982 | 106.2335 | 0.370427 | -3.09368 | 0.001977 | 0.011053 | 0.013389 | 163.9515 | 128.8453 | 70.41294 | 61.72424 | 114663 | inositol monophosphatase 2 [Source:MGI Symbol;Acc:MGI:2149728] |
| ENSMUSG00000027635 | Dsn1 | 1.149158 | 221.2951 | 0.254349 | -4.51804 | 6.24E-06 | 6.99E-05 | 8.47E-05 | 315.2101 | 295.0655 | 146.3126 | 128.5922 | 66934 | DSN1 homolog, MIS12 kinetochore complex component [Source:MGI Symbol;Acc:MGI:1914184] |
| ENSMUSG00000030091 | Nup210 | 1.15569 | 229.8571 | 0.268606 | -4.30255 | 1.69E-05 | 0.000172 | 0.000209 | 356.4624 | 278.3451 | 150.8849 | 133.7359 | 54563 | nucleoporin 210 [Source:MGI Symbol;Acc:MGI:1859555] |
| ENSMUSG00000105987 | AI506816 | 1.163636 | 640.443 | 0.205349 | -5.66662 | 1.46E-08 | 2.76E-07 | 3.35E-07 | 1035.539 | 735.6967 | 383.1562 | 407.38 | | expressed sequence AI506816 [Source:MGI Symbol;Acc:MGI:2140929] |
| ENSMUSG00000057329 | Bcl2 | 1.164246 | 436.279 | 0.199016 | -5.85 | 4.92E-09 | 9.98E-08 | 1.21E-07 | 609.2651 | 597.0159 | 240.5014 | 298.3338 | 12043 | B cell leukemia/lymphoma 2 [Source:MGI Symbol;Acc:MGI:88138] |
| ENSMUSG00000112023 | Lilrb4b | 1.164475 | 6781.705 | 0.109019 | -10.6814 | 1.24E-26 | 1.94E-24 | 2.36E-24 | 10244.33 | 8513.623 | 3975.131 | 4393.737 | 14727 | leukocyte immunoglobulin-like receptor, subfamily B, member 4B [Source:MGI Symbol;Acc:MGI:102702] |
| ENSMUSG00000023272 | Creld2 | 1.164754 | 512.408 | 0.188707 | -6.1723 | 6.73E-10 | 1.55E-08 | 1.88E-08 | 771.1011 | 646.1935 | 299.0264 | 333.3109 | 76737 | cysteine-rich with EGF-like domains 2 [Source:MGI Symbol;Acc:MGI:1923987] |
| ENSMUSG00000031629 | Cenpu | 1.165216 | 117.212 | 0.347504 | -3.3531 | 0.000799 | 0.005187 | 0.006283 | 177.7023 | 146.5492 | 69.49849 | 75.09783 | 71876 | centromere protein U [Source:MGI Symbol;Acc:MGI:1919126] |
| ENSMUSG00000016239 | Lonrf3 | 1.167301 | 1377.676 | 0.152756 | -7.64161 | 2.15E-14 | 8.56E-13 | 1.04E-12 | 2140.89 | 1672.038 | 812.035 | 885.7429 | 74365 | LON peptidase N-terminal domain and ring finger 3 [Source:MGI Symbol;Acc:MGI:1921615] |
| ENSMUSG00000039231 | Suv39h1 | 1.168773 | 453.9118 | 0.203043 | -5.75629 | 8.6E-09 | 1.69E-07 | 2.05E-07 | 700.2318 | 556.6903 | 287.1385 | 271.5867 | 20937 | suppressor of variegation 3-9 1 [Source:MGI Symbol;Acc:MGI:1099440] |
| ENSMUSG00000115219 | Eef1akmt4 | 1.170761 | 138.1737 | 0.328743 | -3.56132 | 0.000369 | 0.002654 | 0.003215 | 213.6659 | 169.1709 | 89.61647 | 80.24151 | 110599566 | EEF1A lysine methyltransferase 4 [Source:MGI Symbol;Acc:MGI:5903914] |
| ENSMUSG00000024007 | Ppil1 | 1.171562 | 342.6673 | 0.20239 | -5.78865 | 7.1E-09 | 1.41E-07 | 1.71E-07 | 478.1039 | 471.1213 | 208.4955 | 212.9486 | 68816 | peptidylprolyl isomerase (cyclophilin)-like 1 [Source:MGI Symbol;Acc:MGI:1916066] |
| ENSMUSG00000030254 | Rad18 | 1.173175 | 153.8302 | 0.37894 | -3.09594 | 0.001962 | 0.010991 | 0.013314 | 271.8422 | 154.4176 | 84.12975 | 104.9312 | 58186 | RAD18 E3 ubiquitin protein ligase [Source:MGI Symbol;Acc:MGI:1890476] |
| ENSMUSG00000029687 | Ezh2 | 1.175644 | 362.5613 | 0.289164 | -4.06566 | 4.79E-05 | 0.000438 | 0.000531 | 610.3229 | 395.3878 | 262.4482 | 182.0865 | 14056 | enhancer of zeste 2 polycomb repressive complex 2 subunit [Source:MGI Symbol;Acc:MGI:107940] |
| ENSMUSG00000031403 | Dkc1 | 1.176067 | 362.0216 | 0.24288 | -4.84217 | 1.28E-06 | 1.66E-05 | 2.01E-05 | 582.8213 | 420.9601 | 206.6666 | 237.6383 | 245474 | dyskeratosis congenita 1, dyskerin [Source:MGI Symbol;Acc:MGI:1861727] |
| ENSMUSG00000026728 | Vim | 1.177206 | 3441.823 | 0.139205 | -8.45666 | 2.75E-17 | 1.51E-15 | 1.82E-15 | 4080.807 | 5464.613 | 2058.436 | 2163.435 | 22352 | vimentin [Source:MGI Symbol;Acc:MGI:98932] |
| ENSMUSG00000079685 | Ulbp1 | 1.18204 | 1040.707 | 0.16298 | -7.25269 | 4.09E-13 | 1.43E-11 | 1.73E-11 | 1577.108 | 1313.042 | 698.6427 | 574.0354 | 77777 | UL16 binding protein 1 [Source:MGI Symbol;Acc:MGI:1925027] |
| ENSMUSG00000020212 | Mdm1 | 1.18214 | 222.1757 | 0.254375 | -4.64723 | 3.36E-06 | 3.96E-05 | 4.8E-05 | 323.6721 | 293.0984 | 128.938 | 142.9945 | 17245 | MDM1 nuclear protein [Source:MGI Symbol;Acc:MGI:96951] |
| ENSMUSG00000022070 | Bora | 1.182772 | 123.4275 | 0.365627 | -3.23492 | 0.001217 | 0.007405 | 0.00897 | 204.1461 | 138.6808 | 76.81412 | 74.06909 | 77744 | bora, aurora kinase A activator [Source:MGI Symbol;Acc:MGI:1924994] |
| ENSMUSG00000017550 | Atad5 | 1.186305 | 367.2381 | 0.257317 | -4.61029 | 4.02E-06 | 4.67E-05 | 5.65E-05 | 614.5539 | 406.2068 | 233.1857 | 215.0061 | 237877 | ATPase family, AAA domain containing 5 [Source:MGI Symbol;Acc:MGI:2442925] |
| ENSMUSG00000097497 | Gm26652 | 1.187475 | 536.5292 | 0.239223 | -4.96388 | 6.91E-07 | 9.42E-06 | 1.14E-05 | 900.1469 | 591.1146 | 299.9409 | 354.9144 | | predicted gene, 26652 [Source:MGI Symbol;Acc:MGI:5477146] |
| ENSMUSG00000059323 | Tonsl | 1.191896 | 110.6009 | 0.359585 | -3.31465 | 0.000918 | 0.005826 | 0.007058 | 154.4318 | 153.4341 | 75.89967 | 58.63803 | 72749 | tonsoku-like, DNA repair protein [Source:MGI Symbol;Acc:MGI:1919899] |
| ENSMUSG00000037894 | H2az1 | 1.193734 | 5214.189 | 0.102457 | -11.6511 | 2.26E-31 | 5.72E-29 | 6.93E-29 | 7872.847 | 6639.957 | 3179.556 | 3164.396 | 51788 | H2A.Z variant histone 1 [Source:MGI Symbol;Acc:MGI:1888388] |
| ENSMUSG00000025586 | Cpeb1 | 1.193799 | 245.27 | 0.242952 | -4.91372 | 8.94E-07 | 1.19E-05 | 1.44E-05 | 324.7298 | 358.0128 | 156.3716 | 141.9658 | 12877 | cytoplasmic polyadenylation element binding protein 1 [Source:MGI Symbol;Acc:MGI:108442] |
| ENSMUSG00000020598 | Nrcam | 1.194536 | 470.1772 | 0.190182 | -6.28103 | 3.36E-10 | 8.06E-09 | 9.76E-09 | 608.2073 | 700.2888 | 269.7639 | 302.4488 | 319504 | neuronal cell adhesion molecule [Source:MGI Symbol;Acc:MGI:104750] |
| ENSMUSG00000007659 | Bcl2l1 | 1.196378 | 1313.541 | 0.115125 | -10.392 | 2.7E-25 | 3.61E-23 | 4.38E-23 | 1850.008 | 1807.768 | 783.6869 | 812.7025 | 12048 | BCL2-like 1 [Source:MGI Symbol;Acc:MGI:88139] |
| ENSMUSG00000045679 | Slc66a3 | 1.197554 | 180.0389 | 0.291995 | -4.10129 | 4.11E-05 | 0.000384 | 0.000466 | 280.3043 | 221.2991 | 111.5634 | 106.9887 | 217430 | solute carrier family 66 member 3 [Source:MGI Symbol;Acc:MGI:2444067] |
| ENSMUSG00000005087 | Cd44 | 1.198738 | 1030.546 | 0.144535 | -8.29375 | 1.1E-16 | 5.53E-15 | 6.7E-15 | 1527.394 | 1344.515 | 663.8935 | 586.3803 | 12505 | CD44 antigen [Source:MGI Symbol;Acc:MGI:88338] |
| ENSMUSG00000054150 | Syne3 | 1.200798 | 113.5721 | 0.345326 | -3.47728 | 0.000507 | 0.003507 | 0.004248 | 166.0671 | 150.4834 | 66.75513 | 70.98288 | 212073 | spectrin repeat containing, nuclear envelope family member 3 [Source:MGI Symbol;Acc:MGI:2442408] |
| ENSMUSG00000032076 | Cadm1 | 1.2013 | 1752.553 | 0.121429 | -9.89303 | 4.46E-23 | 4.64E-21 | 5.61E-21 | 2626.398 | 2259.218 | 1043.392 | 1081.203 | 54725 | cell adhesion molecule 1 [Source:MGI Symbol;Acc:MGI:1889272] |
| ENSMUSG00000027242 | Wdr76 | 1.20166 | 206.2419 | 0.277164 | -4.33556 | 1.45E-05 | 0.00015 | 0.000182 | 315.2101 | 259.6576 | 114.3067 | 135.7933 | 241627 | WD repeat domain 76 [Source:MGI Symbol;Acc:MGI:1926186] |
| ENSMUSG00000045751 | Mms22l | 1.202108 | 364.3325 | 0.200592 | -5.99279 | 2.06E-09 | 4.42E-08 | 5.36E-08 | 530.9915 | 484.891 | 221.2978 | 220.1498 | 212377 | MMS22-like, DNA repair protein [Source:MGI Symbol;Acc:MGI:2684980] |
| ENSMUSG00000024165 | Jpt2 | 1.202845 | 1583.865 | 0.133116 | -9.03609 | 1.62E-19 | 1.15E-17 | 1.39E-17 | 2404.27 | 2012.347 | 919.9405 | 998.904 | 52009 | Jupiter microtubule associated homolog 2 [Source:MGI Symbol;Acc:MGI:1196260] |
| ENSMUSG00000026134 | Prim2 | 1.205596 | 325.9279 | 0.212388 | -5.6764 | 1.38E-08 | 2.63E-07 | 3.18E-07 | 473.8729 | 435.7134 | 204.8377 | 189.2877 | 19076 | DNA primase, p58 subunit [Source:MGI Symbol;Acc:MGI:97758] |
| ENSMUSG00000074506 | Gm10705 | 1.206832 | 104.858 | 0.355679 | -3.39304 | 0.000691 | 0.004573 | 0.005539 | 148.0853 | 144.5821 | 64.01177 | 62.75298 | | predicted gene 10705 [Source:MGI Symbol;Acc:MGI:3708678] |
| ENSMUSG00000029430 | Ran | 1.21066 | 4863.246 | 0.107548 | -11.257 | 2.14E-29 | 4.16E-27 | 5.04E-27 | 7419.072 | 6164.902 | 2888.76 | 2980.252 | 19384 | RAN, member RAS oncogene family [Source:MGI Symbol;Acc:MGI:1333112] |
| ENSMUSG00000030528 | Blm | 1.21165 | 203.3815 | 0.301814 | -4.01455 | 5.96E-05 | 0.00053 | 0.000642 | 331.0763 | 237.036 | 110.6489 | 134.7646 | 12144 | Bloom syndrome, RecQ like helicase [Source:MGI Symbol;Acc:MGI:1328362] |
| ENSMUSG00000117613 | Gm2629 | 1.212358 | 148.9429 | 0.325575 | -3.72374 | 0.000196 | 0.001513 | 0.001833 | 215.7814 | 200.6445 | 104.2477 | 75.09783 | | predicted gene 2629 [Source:MGI Symbol;Acc:MGI:3780797] |
| ENSMUSG00000022346 | Myc | 1.213439 | 111.7566 | 0.362416 | -3.34819 | 0.000813 | 0.00526 | 0.006371 | 176.6446 | 135.7301 | 66.75513 | 67.89667 | 17869 | myelocytomatosis oncogene [Source:MGI Symbol;Acc:MGI:97250] |
| ENSMUSG00000027115 | Kif18a | 1.217239 | 246.8923 | 0.261827 | -4.64903 | 3.34E-06 | 3.93E-05 | 4.76E-05 | 383.9639 | 306.8681 | 160.9439 | 135.7933 | 228421 | kinesin family member 18A [Source:MGI Symbol;Acc:MGI:2446977] |
| ENSMUSG00000041406 | Firrm | 1.218607 | 166.0922 | 0.31038 | -3.92618 | 8.63E-05 | 0.000732 | 0.000887 | 257.0337 | 207.5294 | 88.70202 | 111.1036 | 381306 | FIGNL1 interacting regulator of recombination and mitosis [Source:MGI Symbol;Acc:MGI:3590554] |
| ENSMUSG00000054293 | P2ry10b | 1.220913 | 91.0879 | 0.411522 | -2.96682 | 0.003009 | 0.015673 | 0.01898 | 141.7388 | 113.1084 | 45.72269 | 63.78172 | 213438 | purinergic receptor P2Y, G-protein coupled 10B [Source:MGI Symbol;Acc:MGI:2441814] |
| ENSMUSG00000022351 | Sqle | 1.222905 | 275.6179 | 0.233816 | -5.23019 | 1.69E-07 | 2.63E-06 | 3.18E-06 | 385.0217 | 386.5358 | 150.8849 | 180.029 | 20775 | squalene epoxidase [Source:MGI Symbol;Acc:MGI:109296] |
| ENSMUSG00000058927 | Gm10053 | 1.223111 | 431.8474 | 0.195365 | -6.26065 | 3.83E-10 | 9.1E-09 | 1.1E-08 | 654.7484 | 554.7232 | 259.7049 | 258.2131 | | predicted gene 10053 [Source:MGI Symbol;Acc:MGI:3704493] |
| ENSMUSG00000139678 | Gm73848 | 1.22688 | 441.8086 | 0.213652 | -5.74242 | 9.33E-09 | 1.83E-07 | 2.21E-07 | 703.405 | 535.0521 | 271.5928 | 257.1843 | | predicted gene, 73848 [Source:MGI Symbol;Acc:MGI:7830440] |
| ENSMUSG00000062593 | Gm49339 | 1.226997 | 2079.609 | 0.133883 | -9.16471 | 4.97E-20 | 3.77E-18 | 4.56E-18 | 2999.784 | 2827.711 | 1100.088 | 1390.853 | | predicted gene, 49339 [Source:MGI Symbol;Acc:MGI:6121530] |
| ENSMUSG00000042759 | Apobr | 1.227754 | 426.4294 | 0.194398 | -6.31567 | 2.69E-10 | 6.55E-09 | 7.93E-09 | 638.8822 | 556.6903 | 264.2772 | 245.8682 | 171504 | apolipoprotein B receptor [Source:MGI Symbol;Acc:MGI:2176230] |
| ENSMUSG00000027654 | Fam83d | 1.228812 | 104.8338 | 0.397563 | -3.09086 | 0.001996 | 0.011108 | 0.013456 | 155.4895 | 138.6808 | 76.81412 | 48.35066 | 71878 | family with sequence similarity 83, member D [Source:MGI Symbol;Acc:MGI:1919128] |
| ENSMUSG00000040010 | Slc7a5 | 1.229183 | 109.1398 | 0.389973 | -3.15197 | 0.001622 | 0.0094 | 0.011387 | 173.4713 | 132.7795 | 76.81412 | 53.49434 | 20539 | solute carrier family 7 (cationic amino acid transporter, y+ system), member 5 [Source:MGI Symbol;Acc:MGI:1298205] |
| ENSMUSG00000026192 | Atic | 1.230856 | 949.4132 | 0.16498 | -7.46066 | 8.61E-14 | 3.26E-12 | 3.94E-12 | 1497.777 | 1165.509 | 570.6192 | 563.7481 | 108147 | 5-aminoimidazole-4-carboxamide ribonucleotide formyltransferase/IMP cyclohydrolase [Source:MGI Symbol;Acc:MGI:1351352] |
| ENSMUSG00000041147 | Brca2 | 1.231217 | 294.7006 | 0.276692 | -4.44978 | 8.6E-06 | 9.41E-05 | 0.000114 | 500.3167 | 326.5392 | 180.1474 | 171.7991 | 12190 | breast cancer 2, early onset [Source:MGI Symbol;Acc:MGI:109337] |
| ENSMUSG00000091625 | Lsm5 | 1.237291 | 146.2101 | 0.314859 | -3.92967 | 8.51E-05 | 0.000723 | 0.000876 | 215.7814 | 194.7432 | 78.64303 | 95.67257 | 66373 | LSM5 homolog, U6 small nuclear RNA and mRNA degradation associated [Source:MGI Symbol;Acc:MGI:1913623] |
| ENSMUSG00000120992 | D17H6S56E-5 | 1.239436 | 4222.171 | 0.127571 | -9.71568 | 2.59E-22 | 2.45E-20 | 2.97E-20 | 6647.971 | 5216.758 | 2637.285 | 2386.671 | 110956 | DNA segment, Chr 17, human D6S56E 5 [Source:MGI Symbol;Acc:MGI:1306799] |
| ENSMUSG00000054717 | Hmgb2 | 1.242844 | 546.6939 | 0.199769 | -6.22142 | 4.93E-10 | 1.16E-08 | 1.4E-08 | 793.3139 | 744.5486 | 372.1827 | 276.7303 | 97165 | high mobility group box 2 [Source:MGI Symbol;Acc:MGI:96157] |
| ENSMUSG00000027342 | Pcna | 1.246475 | 3475.378 | 0.127838 | -9.7504 | 1.84E-22 | 1.77E-20 | 2.15E-20 | 5424.152 | 4355.167 | 1934.07 | 2188.124 | 18538 | proliferating cell nuclear antigen [Source:MGI Symbol;Acc:MGI:97503] |
| ENSMUSG00000023805 | Synj2 | 1.247965 | 290.7247 | 0.243287 | -5.12961 | 2.9E-07 | 4.28E-06 | 5.19E-06 | 446.3713 | 371.7825 | 155.4571 | 189.2877 | 20975 | synaptojanin 2 [Source:MGI Symbol;Acc:MGI:1201671] |
| ENSMUSG00000034906 | Ncaph | 1.249353 | 742.6895 | 0.168562 | -7.41182 | 1.25E-13 | 4.61E-12 | 5.59E-12 | 1155.065 | 936.3412 | 446.2535 | 433.0984 | 215387 | non-SMC condensin I complex, subunit H [Source:MGI Symbol;Acc:MGI:2444777] |
| ENSMUSG00000028066 | Pmf1 | 1.251429 | 245.664 | 0.25135 | -4.97883 | 6.4E-07 | 8.76E-06 | 1.06E-05 | 375.5019 | 316.7036 | 153.6282 | 136.8221 | 67037 | polyamine-modulated factor 1 [Source:MGI Symbol;Acc:MGI:1914287] |
| ENSMUSG00000001707 | Eef1e1 | 1.257402 | 261.0978 | 0.239438 | -5.25148 | 1.51E-07 | 2.37E-06 | 2.87E-06 | 386.0794 | 350.1444 | 143.5692 | 164.598 | 66143 | eukaryotic translation elongation factor 1 epsilon 1 [Source:MGI Symbol;Acc:MGI:1913393] |
| ENSMUSG00000069305 | H4c18 | 1.260699 | 3362.763 | 0.125398 | -10.0536 | 8.86E-24 | 9.88E-22 | 1.2E-21 | 5295.106 | 4195.832 | 2029.173 | 1930.94 | 319161 | H4 clustered histone 18 [Source:MGI Symbol;Acc:MGI:4843992] |
| ENSMUSG00000020571 | Pdia6 | 1.261016 | 5459.911 | 0.093497 | -13.4872 | 1.86E-41 | 1.23E-38 | 1.48E-38 | 8193.346 | 7217.302 | 3289.29 | 3139.706 | 71853 | protein disulfide isomerase associated 6 [Source:MGI Symbol;Acc:MGI:1919103] |
| ENSMUSG00000062070 | Pgk1 | 1.261572 | 6027.536 | 0.104435 | -12.08 | 1.35E-33 | 4.35E-31 | 5.27E-31 | 9293.408 | 7720.881 | 3559.054 | 3536.799 | 18655 | phosphoglycerate kinase 1 [Source:MGI Symbol;Acc:MGI:97555] |
| ENSMUSG00000049588 | Ccdc69 | 1.264485 | 66.19644 | 0.464113 | -2.72452 | 0.006439 | 0.029606 | 0.035767 | 86.73566 | 100.3223 | 44.80824 | 32.9196 | 52570 | coiled-coil domain containing 69 [Source:MGI Symbol;Acc:MGI:1196234] |
| ENSMUSG00000059791 | Nrm | 1.268785 | 115.8653 | 0.347799 | -3.64805 | 0.000264 | 0.001977 | 0.002394 | 170.2981 | 157.3683 | 74.07076 | 61.72424 | 106582 | nurim (nuclear envelope membrane protein) [Source:MGI Symbol;Acc:MGI:2146855] |
| ENSMUSG00000034591 | Slc41a2 | 1.27173 | 194.7103 | 0.272326 | -4.66988 | 3.01E-06 | 3.59E-05 | 4.35E-05 | 292.9973 | 257.6905 | 108.82 | 119.3335 | 338365 | solute carrier family 41, member 2 [Source:MGI Symbol;Acc:MGI:2442940] |
| ENSMUSG00000021493 | Pdlim7 | 1.275735 | 97.00967 | 0.390963 | -3.26306 | 0.001102 | 0.006813 | 0.008253 | 157.605 | 117.0427 | 55.78168 | 57.60929 | 67399 | PDZ and LIM domain 7 [Source:MGI Symbol;Acc:MGI:1914649] |
| ENSMUSG00000032575 | Manf | 1.279948 | 1296.439 | 0.143919 | -8.89355 | 5.92E-19 | 3.87E-17 | 4.69E-17 | 2028.768 | 1644.498 | 747.1088 | 765.3806 | 74840 | mesencephalic astrocyte-derived neurotrophic factor [Source:MGI Symbol;Acc:MGI:1922090] |
| ENSMUSG00000079139 | Gm4204 | 1.288832 | 83.3216 | 0.445335 | -2.89407 | 0.003803 | 0.019163 | 0.023175 | 135.3922 | 101.3058 | 58.52504 | 38.06328 | | predicted gene 4204 [Source:MGI Symbol;Acc:MGI:3782381] |
| ENSMUSG00000028613 | Lrp8 | 1.290498 | 253.0898 | 0.243555 | -5.29859 | 1.17E-07 | 1.87E-06 | 2.27E-06 | 387.1372 | 331.4569 | 143.5692 | 150.1957 | 16975 | low density lipoprotein receptor-related protein 8, apolipoprotein e receptor [Source:MGI Symbol;Acc:MGI:1340044] |
| ENSMUSG00000033952 | Aspm | 1.291452 | 1152.813 | 0.180036 | -7.17331 | 7.32E-13 | 2.49E-11 | 3.02E-11 | 1855.297 | 1419.265 | 746.1943 | 590.4952 | 12316 | abnormal spindle microtubule assembly [Source:MGI Symbol;Acc:MGI:1334448] |
| ENSMUSG00000022769 | Sdf2l1 | 1.293065 | 1168.75 | 0.155459 | -8.31774 | 8.97E-17 | 4.62E-15 | 5.6E-15 | 1861.643 | 1458.607 | 667.5513 | 687.1966 | 64136 | stromal cell-derived factor 2-like 1 [Source:MGI Symbol;Acc:MGI:2149842] |
| ENSMUSG00000078762 | Haus5 | 1.300882 | 189.9807 | 0.29036 | -4.48024 | 7.46E-06 | 8.24E-05 | 9.98E-05 | 304.6326 | 236.0524 | 114.3067 | 104.9312 | 71909 | HAUS augmin-like complex, subunit 5 [Source:MGI Symbol;Acc:MGI:1919159] |
| ENSMUSG00000040658 | Dnph1 | 1.302573 | 68.45837 | 0.472868 | -2.75462 | 0.005876 | 0.027423 | 0.033128 | 115.295 | 79.66769 | 43.89378 | 34.97707 | 381101 | 2'-deoxynucleoside 5'-phosphate N-hydrolase 1 [Source:MGI Symbol;Acc:MGI:3039376] |
| ENSMUSG00000050335 | Lgals3 | 1.304285 | 2777.619 | 0.096017 | -13.584 | 4.98E-42 | 3.44E-39 | 4.16E-39 | 4036.381 | 3871.259 | 1536.282 | 1666.555 | 16854 | lectin, galactose binding, soluble 3 [Source:MGI Symbol;Acc:MGI:96778] |
| ENSMUSG00000024691 | Fam111a | 1.310981 | 2493.32 | 0.127767 | -10.2607 | 1.06E-24 | 1.3E-22 | 1.57E-22 | 3944.357 | 3164.086 | 1409.173 | 1455.663 | 107373 | family with sequence similarity 111, member A [Source:MGI Symbol;Acc:MGI:1915508] |
| ENSMUSG00000032294 | Pkm | 1.313048 | 17783.52 | 0.087717 | -14.9691 | 1.17E-50 | 2.96E-47 | 3.58E-47 | 27243.46 | 23477.38 | 10124.83 | 10288.4 | 18746 | pyruvate kinase, muscle [Source:MGI Symbol;Acc:MGI:97591] |
| ENSMUSG00000036067 | Slc2a6 | 1.314902 | 683.8103 | 0.163852 | -8.02495 | 1.02E-15 | 4.74E-14 | 5.74E-14 | 1038.712 | 912.736 | 412.4187 | 371.3742 | 227659 | solute carrier family 2 (facilitated glucose transporter), member 6 [Source:MGI Symbol;Acc:MGI:2443286] |
| ENSMUSG00000058290 | Espl1 | 1.318105 | 326.7199 | 0.225931 | -5.83411 | 5.41E-09 | 1.1E-07 | 1.33E-07 | 513.0097 | 419.9766 | 193.8642 | 180.029 | 105988 | extra spindle pole bodies 1, separase [Source:MGI Symbol;Acc:MGI:2146156] |
| ENSMUSG00000079445 | B3gnt7 | 1.319631 | 85.52878 | 0.399105 | -3.30648 | 0.000945 | 0.005976 | 0.00724 | 131.1612 | 113.1084 | 48.46605 | 49.37939 | 227327 | UDP-GlcNAc:betaGal beta-1,3-N-acetylglucosaminyltransferase 7 [Source:MGI Symbol;Acc:MGI:2384394] |
| ENSMUSG00000006398 | Cdc20 | 1.321597 | 544.8421 | 0.195982 | -6.74347 | 1.55E-11 | 4.47E-10 | 5.41E-10 | 869.4721 | 687.5026 | 329.2034 | 293.1901 | 107995 | cell division cycle 20 [Source:MGI Symbol;Acc:MGI:1859866] |
| ENSMUSG00000020534 | Shmt1 | 1.321756 | 145.344 | 0.339801 | -3.88979 | 0.0001 | 0.000834 | 0.00101 | 243.2829 | 172.1215 | 87.78757 | 78.18404 | 20425 | serine hydroxymethyltransferase 1 (soluble) [Source:MGI Symbol;Acc:MGI:98299] |
| ENSMUSG00000001525 | Tubb5 | 1.322839 | 11428.47 | 0.106709 | -12.3966 | 2.73E-35 | 1.06E-32 | 1.28E-32 | 18026.21 | 14633.28 | 6719.407 | 6334.965 | 22154 | tubulin, beta 5 class I [Source:MGI Symbol;Acc:MGI:107812] |
| ENSMUSG00000027018 | Hat1 | 1.323498 | 759.1196 | 0.155563 | -8.50782 | 1.77E-17 | 9.84E-16 | 1.19E-15 | 1163.527 | 1006.173 | 431.6222 | 435.1559 | 107435 | histone aminotransferase 1 [Source:MGI Symbol;Acc:MGI:96013] |
| ENSMUSG00000040322 | Slc25a24 | 1.323592 | 493.1784 | 0.207923 | -6.36577 | 1.94E-10 | 4.83E-09 | 5.85E-09 | 803.8915 | 605.8678 | 286.224 | 276.7303 | 229731 | solute carrier family 25 (mitochondrial carrier, phosphate carrier), member 24 [Source:MGI Symbol;Acc:MGI:1917160] |
| ENSMUSG00000029304 | Spp1 | 1.324129 | 31596.13 | 0.100134 | -13.2236 | 6.41E-40 | 3.47E-37 | 4.21E-37 | 45478.04 | 44834.22 | 16170.29 | 19901.95 | 20750 | secreted phosphoprotein 1 [Source:MGI Symbol;Acc:MGI:98389] |
| ENSMUSG00000032555 | Topbp1 | 1.324442 | 2075.096 | 0.147262 | -8.99379 | 2.39E-19 | 1.65E-17 | 1.99E-17 | 3384.806 | 2547.399 | 1201.592 | 1166.588 | 235559 | topoisomerase (DNA) II binding protein 1 [Source:MGI Symbol;Acc:MGI:1920018] |
| ENSMUSG00000005732 | Ranbp1 | 1.324618 | 1072.373 | 0.137554 | -9.62977 | 5.99E-22 | 5.57E-20 | 6.75E-20 | 1640.573 | 1425.166 | 612.6841 | 611.07 | 19385 | RAN binding protein 1 [Source:MGI Symbol;Acc:MGI:96269] |
| ENSMUSG00000055612 | Cdca7 | 1.325187 | 188.1395 | 0.351177 | -3.77355 | 0.000161 | 0.001268 | 0.001536 | 325.7876 | 212.4472 | 128.938 | 85.3852 | 66953 | cell division cycle associated 7 [Source:MGI Symbol;Acc:MGI:1914203] |
| ENSMUSG00000028560 | Usp1 | 1.327143 | 753.443 | 0.187617 | -7.07369 | 1.51E-12 | 4.94E-11 | 5.99E-11 | 1237.57 | 917.6537 | 439.8523 | 418.6961 | 230484 | ubiquitin specific peptidase 1 [Source:MGI Symbol;Acc:MGI:2385198] |
| ENSMUSG00000080875 | Gm7332 | 1.32779 | 81.49246 | 0.411005 | -3.23059 | 0.001235 | 0.007496 | 0.00908 | 126.9302 | 106.2236 | 47.5516 | 45.26444 | | predicted gene 7332 [Source:MGI Symbol;Acc:MGI:3643344] |
| ENSMUSG00000063142 | Kcnma1 | 1.328218 | 97.76184 | 0.388634 | -3.41765 | 0.000632 | 0.004238 | 0.005134 | 159.7205 | 119.9933 | 55.78168 | 55.55182 | 16531 | potassium large conductance calcium-activated channel, subfamily M, alpha member 1 [Source:MGI Symbol;Acc:MGI:99923] |
| ENSMUSG00000022385 | Gtse1 | 1.330456 | 189.6941 | 0.278555 | -4.77628 | 1.79E-06 | 2.22E-05 | 2.69E-05 | 295.1128 | 247.855 | 108.82 | 106.9887 | 29870 | G two S phase expressed protein 1 [Source:MGI Symbol;Acc:MGI:1352755] |
| ENSMUSG00000027353 | Mcm8 | 1.332877 | 93.84833 | 0.404657 | -3.29384 | 0.000988 | 0.006215 | 0.007529 | 157.605 | 111.1413 | 52.12387 | 54.52308 | 66634 | minichromosome maintenance 8 homologous recombination repair factor [Source:MGI Symbol;Acc:MGI:1913884] |
| ENSMUSG00000022221 | Ripk3 | 1.333034 | 137.3046 | 0.331015 | -4.02711 | 5.65E-05 | 0.000505 | 0.000612 | 221.0701 | 172.1215 | 76.81412 | 79.21278 | 56532 | receptor-interacting serine-threonine kinase 3 [Source:MGI Symbol;Acc:MGI:2154952] |
| ENSMUSG00000039470 | Zdhhc2 | 1.337003 | 207.2393 | 0.259187 | -5.15844 | 2.49E-07 | 3.73E-06 | 4.52E-06 | 306.7481 | 287.1971 | 119.7935 | 115.2186 | 70546 | zinc finger, DHHC domain containing 2 [Source:MGI Symbol;Acc:MGI:1923452] |
| ENSMUSG00000024084 | Qpct | 1.337373 | 338.9694 | 0.223042 | -5.99605 | 2.02E-09 | 4.35E-08 | 5.26E-08 | 532.0492 | 439.6476 | 202.0943 | 182.0865 | 70536 | glutaminyl-peptide cyclotransferase (glutaminyl cyclase) [Source:MGI Symbol;Acc:MGI:1917786] |
| ENSMUSG00000040675 | Mthfd1l | 1.341864 | 181.511 | 0.312379 | -4.29563 | 1.74E-05 | 0.000177 | 0.000214 | 302.517 | 218.3485 | 111.5634 | 93.6151 | 270685 | methylenetetrahydrofolate dehydrogenase (NADP+ dependent) 1-like [Source:MGI Symbol;Acc:MGI:1924836] |
| ENSMUSG00000034349 | Smc4 | 1.342873 | 1894.093 | 0.12993 | -10.3354 | 4.88E-25 | 6.27E-23 | 7.59E-23 | 2979.687 | 2454.945 | 1103.746 | 1037.996 | 70099 | structural maintenance of chromosomes 4 [Source:MGI Symbol;Acc:MGI:1917349] |
| ENSMUSG00000022881 | Rfc4 | 1.347736 | 162.9838 | 0.302046 | -4.46202 | 8.12E-06 | 8.93E-05 | 0.000108 | 248.5717 | 219.332 | 83.2153 | 100.8163 | 106344 | replication factor C (activator 1) 4 [Source:MGI Symbol;Acc:MGI:2146571] |
| ENSMUSG00000026547 | Tagln2 | 1.348351 | 770.1239 | 0.160919 | -8.37904 | 5.34E-17 | 2.82E-15 | 3.42E-15 | 1182.567 | 1029.779 | 461.7992 | 406.3513 | 21346 | transgelin 2 [Source:MGI Symbol;Acc:MGI:1312985] |
| ENSMUSG00000111394 | Gm49759 | 1.349094 | 337.2306 | 0.244038 | -5.52822 | 3.23E-08 | 5.84E-07 | 7.07E-07 | 561.6663 | 407.1904 | 193.8642 | 186.2015 | | predicted gene, 49759 [Source:MGI Symbol;Acc:MGI:6215263] |
| ENSMUSG00000036086 | Zranb3 | 1.350074 | 235.6346 | 0.254544 | -5.3039 | 1.13E-07 | 1.83E-06 | 2.21E-06 | 369.1554 | 307.8517 | 130.7669 | 134.7646 | 226409 | zinc finger, RAN-binding domain containing 3 [Source:MGI Symbol;Acc:MGI:1918362] |
| ENSMUSG00000052512 | Nav2 | 1.352876 | 199.196 | 0.403648 | -3.35162 | 0.000803 | 0.00521 | 0.006312 | 372.3287 | 200.6445 | 143.5692 | 80.24151 | 78286 | neuron navigator 2 [Source:MGI Symbol;Acc:MGI:2183691] |
| ENSMUSG00000052688 | Rab7b | 1.355717 | 2236.53 | 0.106724 | -12.703 | 5.69E-37 | 2.54E-34 | 3.07E-34 | 3399.615 | 3033.273 | 1259.203 | 1254.031 | 226421 | RAB7B, member RAS oncogene family [Source:MGI Symbol;Acc:MGI:2442295] |
| ENSMUSG00000031799 | Tpm4 | 1.36574 | 4989.085 | 0.1185 | -11.5253 | 9.84E-31 | 2.26E-28 | 2.74E-28 | 8023.048 | 6354.727 | 2792.742 | 2785.821 | 326618 | tropomyosin 4 [Source:MGI Symbol;Acc:MGI:2449202] |
| ENSMUSG00000023345 | Poc1a | 1.366697 | 140.7699 | 0.318539 | -4.29051 | 1.78E-05 | 0.00018 | 0.000218 | 216.8391 | 188.8419 | 74.07076 | 83.32773 | 70235 | POC1 centriolar protein A [Source:MGI Symbol;Acc:MGI:1917485] |
| ENSMUSG00000062248 | Cks2 | 1.370385 | 220.7695 | 0.29109 | -4.70778 | 2.5E-06 | 3.03E-05 | 3.67E-05 | 373.3864 | 263.5919 | 129.8524 | 116.2473 | 66197 | CDC28 protein kinase regulatory subunit 2 [Source:MGI Symbol;Acc:MGI:1913447] |
| ENSMUSG00000121491 | | 1.371794 | 61.96384 | 0.52413 | -2.61728 | 0.008863 | 0.038619 | 0.046648 | 111.064 | 67.86507 | 41.15042 | 27.77591 | 77022 | NDC80 homolog, kinetochore complex component pseudogene [Source:NCBI gene (formerly Entrezgene);Acc:77022] |
| ENSMUSG00000009687 | Fxyd5 | 1.379255 | 476.7539 | 0.200782 | -6.86941 | 6.45E-12 | 1.98E-10 | 2.4E-10 | 729.8488 | 648.1606 | 294.4541 | 234.5521 | 18301 | FXYD domain-containing ion transport regulator 5 [Source:MGI Symbol;Acc:MGI:1201785] |
| ENSMUSG00000028832 | Stmn1 | 1.382372 | 1888.772 | 0.125525 | -11.0127 | 3.32E-28 | 5.59E-26 | 6.77E-26 | 2885.547 | 2575.922 | 1126.607 | 967.0131 | 16765 | stathmin 1 [Source:MGI Symbol;Acc:MGI:96739] |
| ENSMUSG00000063954 | H2ac19 | 1.384011 | 3180.334 | 0.145458 | -9.51483 | 1.82E-21 | 1.55E-19 | 1.88E-19 | 5244.334 | 3953.878 | 1887.433 | 1635.692 | 319192 | H2A clustered histone 19 [Source:MGI Symbol;Acc:MGI:2448283] |
| ENSMUSG00000029363 | Rfc5 | 1.388189 | 465.9232 | 0.198278 | -7.00124 | 2.54E-12 | 8.1E-11 | 9.81E-11 | 745.7151 | 602.9172 | 257.876 | 257.1843 | 72151 | replication factor C (activator 1) 5 [Source:MGI Symbol;Acc:MGI:1919401] |
| ENSMUSG00000070348 | Ccnd1 | 1.394501 | 2856.867 | 0.151852 | -9.18329 | 4.18E-20 | 3.22E-18 | 3.9E-18 | 4809.598 | 3468.987 | 1523.48 | 1625.405 | 12443 | cyclin D1 [Source:MGI Symbol;Acc:MGI:88313] |
| ENSMUSG00000025044 | Msr1 | 1.405538 | 3353.996 | 0.116261 | -12.0895 | 1.2E-33 | 3.96E-31 | 4.79E-31 | 5301.453 | 4437.785 | 1753.008 | 1923.739 | 20288 | macrophage scavenger receptor 1 [Source:MGI Symbol;Acc:MGI:98257] |
| ENSMUSG00000078636 | Gm7336 | 1.406482 | 373.3379 | 0.271411 | -5.1821 | 2.19E-07 | 3.32E-06 | 4.03E-06 | 656.8639 | 427.845 | 228.6135 | 180.029 | | predicted gene 7336 [Source:MGI Symbol;Acc:MGI:3644851] |
| ENSMUSG00000039081 | Zfp503 | 1.413004 | 56.30764 | 0.505162 | -2.79713 | 0.005156 | 0.024662 | 0.0298 | 75.10039 | 88.51965 | 25.60471 | 36.00581 | 218820 | zinc finger protein 503 [Source:MGI Symbol;Acc:MGI:1353644] |
| ENSMUSG00000031697 | Orc6 | 1.413532 | 277.356 | 0.259552 | -5.44605 | 5.15E-08 | 8.96E-07 | 1.09E-06 | 464.3531 | 342.276 | 145.3982 | 157.3968 | 56452 | origin recognition complex, subunit 6 [Source:MGI Symbol;Acc:MGI:1929285] |
| ENSMUSG00000020089 | Ppa1 | 1.421425 | 276.344 | 0.230974 | -6.15404 | 7.55E-10 | 1.72E-08 | 2.09E-08 | 425.2163 | 379.651 | 147.2271 | 153.2819 | 67895 | pyrophosphatase (inorganic) 1 [Source:MGI Symbol;Acc:MGI:97831] |
| ENSMUSG00000019961 | Tmpo | 1.427608 | 2860.109 | 0.128647 | -11.0971 | 1.3E-28 | 2.21E-26 | 2.67E-26 | 4643.531 | 3697.171 | 1603.952 | 1495.784 | 21917 | thymopoietin [Source:MGI Symbol;Acc:MGI:106920] |
| ENSMUSG00000028551 | Cdkn2c | 1.43015 | 705.7816 | 0.169189 | -8.45299 | 2.84E-17 | 1.54E-15 | 1.87E-15 | 1120.159 | 939.2919 | 400.5308 | 363.1443 | 12580 | cyclin dependent kinase inhibitor 2C [Source:MGI Symbol;Acc:MGI:105388] |
| ENSMUSG00000025161 | Slc16a3 | 1.433672 | 701.0585 | 0.195255 | -7.34257 | 2.1E-13 | 7.64E-12 | 9.25E-12 | 1180.451 | 866.509 | 394.1296 | 363.1443 | 80879 | solute carrier family 16 (monocarboxylic acid transporters), member 3 [Source:MGI Symbol;Acc:MGI:1933438] |
| ENSMUSG00000032397 | Tipin | 1.436079 | 350.0264 | 0.241472 | -5.94719 | 2.73E-09 | 5.76E-08 | 6.98E-08 | 577.5325 | 444.5654 | 169.174 | 208.8337 | 66131 | timeless interacting protein [Source:MGI Symbol;Acc:MGI:1921571] |
| ENSMUSG00000031016 | Wee1 | 1.43719 | 220.2314 | 0.294151 | -4.88589 | 1.03E-06 | 1.36E-05 | 1.64E-05 | 380.7907 | 262.6083 | 116.1356 | 121.391 | 22390 | WEE 1 homolog 1 (S. pombe) [Source:MGI Symbol;Acc:MGI:103075] |
| ENSMUSG00000005481 | Ddx39a | 1.437779 | 880.8437 | 0.174465 | -8.2411 | 1.71E-16 | 8.44E-15 | 1.02E-14 | 1456.524 | 1117.315 | 493.8051 | 455.7306 | 68278 | DEAD box helicase 39a [Source:MGI Symbol;Acc:MGI:1915528] |
| ENSMUSG00000023004 | Tuba1b | 1.438344 | 10127.18 | 0.078345 | -18.3591 | 2.79E-75 | 4.23E-71 | 5.13E-71 | 15462.22 | 14128.72 | 5541.59 | 5376.181 | 22143 | tubulin, alpha 1B [Source:MGI Symbol;Acc:MGI:107804] |
| ENSMUSG00000048007 | Timm8a1 | 1.438727 | 213.331 | 0.297842 | -4.8305 | 1.36E-06 | 1.75E-05 | 2.12E-05 | 359.6357 | 263.5919 | 101.5044 | 128.5922 | 30058 | translocase of inner mitochondrial membrane 8A1 [Source:MGI Symbol;Acc:MGI:1353433] |
| ENSMUSG00000021418 | Rpp40 | 1.43886 | 79.50717 | 0.459515 | -3.13126 | 0.001741 | 0.009959 | 0.012065 | 142.7965 | 89.5032 | 38.40706 | 47.32192 | 208366 | ribonuclease P 40 subunit [Source:MGI Symbol;Acc:MGI:1346084] |
| ENSMUSG00000028884 | Rpa2 | 1.43908 | 333.1601 | 0.254315 | -5.65865 | 1.53E-08 | 2.88E-07 | 3.49E-07 | 573.3015 | 400.3055 | 172.8318 | 186.2015 | 19891 | replication protein A2 [Source:MGI Symbol;Acc:MGI:1339939] |
| ENSMUSG00000061482 | H4c4 | 1.454231 | 2022.091 | 0.150906 | -9.6367 | 5.6E-22 | 5.24E-20 | 6.34E-20 | 3292.782 | 2633.951 | 1191.533 | 970.0993 | 319156 | H4 clustered histone 4 [Source:MGI Symbol;Acc:MGI:2448423] |
| ENSMUSG00000021360 | Gcnt2 | 1.454602 | 159.8926 | 0.294737 | -4.93525 | 8E-07 | 1.08E-05 | 1.31E-05 | 231.6477 | 237.036 | 89.61647 | 81.27025 | 14538 | glucosaminyl (N-acetyl) transferase 2 (I blood group) [Source:MGI Symbol;Acc:MGI:1100870] |
| ENSMUSG00000018362 | Kpna2 | 1.459093 | 2432.55 | 0.118198 | -12.3444 | 5.22E-35 | 1.98E-32 | 2.4E-32 | 3880.892 | 3254.573 | 1314.985 | 1279.749 | 16647 | karyopherin subunit alpha 2 [Source:MGI Symbol;Acc:MGI:103561] |
| ENSMUSG00000112808 | Gm4739 | 1.459381 | 82.31207 | 0.470361 | -3.10268 | 0.001918 | 0.01079 | 0.013071 | 136.45 | 105.24 | 56.69614 | 30.86212 | | predicted gene 4739 [Source:MGI Symbol;Acc:MGI:3642984] |
| ENSMUSG00000027221 | Chst1 | 1.468995 | 70.75752 | 0.461267 | -3.1847 | 0.001449 | 0.008563 | 0.010373 | 119.526 | 88.51965 | 42.06488 | 32.9196 | 76969 | carbohydrate sulfotransferase 1 [Source:MGI Symbol;Acc:MGI:1924219] |
| ENSMUSG00000029413 | Naaa | 1.473535 | 105.6256 | 0.407664 | -3.61459 | 0.000301 | 0.002212 | 0.00268 | 121.6415 | 188.8419 | 50.29496 | 61.72424 | 67111 | N-acylethanolamine acid amidase [Source:MGI Symbol;Acc:MGI:1914361] |
| ENSMUSG00000021391 | Cenpp | 1.474302 | 55.39681 | 0.522576 | -2.82122 | 0.004784 | 0.023154 | 0.027995 | 95.19767 | 67.86507 | 33.83479 | 24.6897 | 66336 | centromere protein P [Source:MGI Symbol;Acc:MGI:1913586] |
| ENSMUSG00000047534 | Mis18bp1 | 1.475 | 441.4583 | 0.220091 | -6.70179 | 2.06E-11 | 5.87E-10 | 7.11E-10 | 739.3686 | 559.6409 | 248.7314 | 218.0923 | 217653 | MIS18 binding protein 1 [Source:MGI Symbol;Acc:MGI:2145099] |
| ENSMUSG00000055745 | Rtl6 | 1.475632 | 179.6655 | 0.300953 | -4.9032 | 9.43E-07 | 1.25E-05 | 1.51E-05 | 228.4744 | 299.9833 | 91.44538 | 98.75879 | 223732 | retrotransposon Gag like 6 [Source:MGI Symbol;Acc:MGI:2675858] |
| ENSMUSG00000052565 | H1f3 | 1.475888 | 2670.784 | 0.334078 | -4.41779 | 9.97E-06 | 0.000108 | 0.000131 | 4456.309 | 3402.105 | 1680.766 | 1143.956 | 14957 | H1.3 linker histone, cluster member [Source:MGI Symbol;Acc:MGI:107502] |
| ENSMUSG00000026605 | Cenpf | 1.476155 | 1177.341 | 0.177822 | -8.30131 | 1.03E-16 | 5.22E-15 | 6.33E-15 | 1934.628 | 1530.406 | 703.215 | 541.1158 | 108000 | centromere protein F [Source:MGI Symbol;Acc:MGI:1313302] |
| ENSMUSG00000025758 | Plk4 | 1.485477 | 412.4433 | 0.235281 | -6.31363 | 2.73E-10 | 6.62E-09 | 8.02E-09 | 705.5205 | 510.4633 | 233.1857 | 200.6038 | 20873 | polo like kinase 4 [Source:MGI Symbol;Acc:MGI:101783] |
| ENSMUSG00000031995 | St14 | 1.492154 | 72.78309 | 0.491663 | -3.03491 | 0.002406 | 0.013011 | 0.015761 | 120.5837 | 94.42096 | 49.38051 | 26.74717 | 19143 | suppression of tumorigenicity 14 (colon carcinoma) [Source:MGI Symbol;Acc:MG[illegible] |
| ENSMUSG00000069273 | H3c6 | 1.496952 | 1296.94 | 0.169888 | -8.81142 | 1.24E-18 | 7.62E-17 | 9.23E-17 | 2097.522 | 1734.002 | 770.8846 | 585.3516 | 319151 | H3 clustered histone 6 [Source:MGI Symbol;Acc:MGI:2448326] |
| ENSMUSG00000063234 | Gpr84 | 1.497805 | 814.1528 | 0.447336 | -3.34828 | 0.000813 | 0.00526 | 0.006371 | 1573.935 | 831.1012 | 405.103 | 446.472 | 80910 | G protein-coupled receptor 84 [Source:MGI Symbol;Acc:MGI:1934129] |
| ENSMUSG00000041957 | Pkp2 | 1.49814 | 133.5754 | 0.34549 | -4.33628 | 1.45E-05 | 0.00015 | 0.000182 | 227.4167 | 167.2038 | 67.66958 | 72.01161 | 67451 | plakophilin 2 [Source:MGI Symbol;Acc:MGI:1914701] |
| ENSMUSG00000021822 | Plau | 1.499685 | 2098.2 | 0.174485 | -8.5949 | 8.33E-18 | 4.77E-16 | 5.78E-16 | 3624.916 | 2574.938 | 982.1234 | 1210.824 | 18792 | plasminogen activator, urokinase [Source:MGI Symbol;Acc:MGI:97611] |
| ENSMUSG00000036752 | Tubb4b | 1.50004 | 2578.043 | 0.124299 | -12.068 | 1.56E-33 | 4.93E-31 | 5.97E-31 | 3811.08 | 3806.345 | 1197.935 | 1496.813 | 227613 | tubulin, beta 4B class IVB [Source:MGI Symbol;Acc:MGI:1915472] |

| | | | | | | | | | | | | | |
|---|---|---|---|---|---|---|---|---|---|---|---|---|---|
| ENSMUSG00000041488 | Stx3 | 1.649437 | 248.6356 | 0.282865 | -5.83117 | 5.5E-09 | 1.11E-07 | 1.35E-07 | 412.5232 | 341.2924 | 98.76101 | 141.9658 | 20908 | syntaxin 3 [Source:MGI Symbol;Acc:MGI:103077] |
| ENSMUSG00000038463 | Olfml2b | 1.654301 | 190.8002 | 0.321658 | -5.14305 | 2.7E-07 | 4.01E-06 | 4.86E-06 | 350.1159 | 229.1675 | 92.35984 | 91.55762 | 320078 | olfactomedin-like 2B [Source:MGI Symbol;Acc:MGI:2443310] |
| ENSMUSG00000024742 | Fen1 | 1.654565 | 949.6153 | 0.156072 | -10.6013 | 2.94E-26 | 4.45E-24 | 5.4E-24 | 1585.57 | 1297.305 | 452.6546 | 462.9318 | 14156 | flap structure specific endonuclease 1 [Source:MGI Symbol;Acc:MGI:102779] |
| ENSMUSG00000025324 | Atp10a | 1.654717 | 88.63919 | 0.425842 | -3.88575 | 0.000102 | 0.000847 | 0.001026 | 151.2585 | 118.0262 | 50.29496 | 34.97707 | 11982 | ATPase, class V, type 10A [Source:MGI Symbol;Acc:MGI:1330809] |
| ENSMUSG00000069270 | H2ac6 | 1.655273 | 400.9707 | 0.220037 | -7.52271 | 5.37E-14 | 2.06E-12 | 2.49E-12 | 682.25 | 535.0521 | 182.8908 | 203.69 | 319164 | H2A clustered histone 6 [Source:MGI Symbol;Acc:MGI:2448287] |
| ENSMUSG00000019942 | Cdk1 | 1.65892 | 1213.164 | 0.149025 | -11.1318 | 8.78E-29 | 1.53E-26 | 1.85E-26 | 2028.768 | 1657.285 | 609.0262 | 557.5756 | 12534 | cyclin dependent kinase 1 [Source:MGI Symbol;Acc:MGI:88351] |
| ENSMUSG00000028614 | Ndc1 | 1.659125 | 641.8459 | 0.197188 | -8.41391 | 3.97E-17 | 2.13E-15 | 2.58E-15 | 1114.871 | 835.0354 | 294.4541 | 323.0235 | 72787 | NDC1 transmembrane nucleoporin [Source:MGI Symbol;Acc:MGI:1920037] |
| ENSMUSG00000026873 | Phf19 | 1.662413 | 124.9158 | 0.334539 | -4.96927 | 6.72E-07 | 9.19E-06 | 1.11E-05 | 197.7996 | 181.9571 | 61.26841 | 58.63803 | 74016 | PHD finger protein 19 [Source:MGI Symbol;Acc:MGI:1921266] |
| ENSMUSG00000070407 | Hs3st3b1 | 1.669976 | 71.09273 | 0.460886 | -3.6234 | 0.000291 | 0.002147 | 0.002601 | 125.8725 | 90.48676 | 32.00588 | 36.00581 | 54710 | heparan sulfate (glucosamine) 3-O-sulfotransferase 3B1 [Source:MGI Symbol;Acc:MGI:1333853] |
| ENSMUSG00000041431 | Ccnb1 | 1.675174 | 1137.201 | 0.156305 | -10.7173 | 8.44E-27 | 1.33E-24 | 1.61E-24 | 1937.802 | 1526.472 | 541.3567 | 543.1733 | 268697 | cyclin B1 [Source:MGI Symbol;Acc:MGI:88302] |
| ENSMUSG00000025395 | Prim1 | 1.68314 | 324.3021 | 0.248423 | -6.7753 | 1.24E-11 | 3.63E-10 | 4.39E-10 | 572.2438 | 417.0259 | 153.6282 | 154.3106 | 19075 | DNA primase, p49 subunit [Source:MGI Symbol;Acc:MGI:97757] |
| ENSMUSG00000026646 | Suv39h2 | 1.68521 | 73.90431 | 0.453981 | -3.71207 | 0.000206 | 0.001577 | 0.001911 | 129.0457 | 96.38807 | 31.09143 | 39.09202 | 64707 | suppressor of variegation 3-9 2 [Source:MGI Symbol;Acc:MGI:1890396] |
| ENSMUSG00000026784 | Pdss1 | 1.685718 | 89.62973 | 0.411715 | -4.09438 | 4.23E-05 | 0.000394 | 0.000478 | 156.5473 | 117.0427 | 44.80824 | 40.12076 | 56075 | prenyl (solanesyl) diphosphate synthase, subunit 1 [Source:MGI Symbol;Acc:MGI:1889278] |
| ENSMUSG00000021714 | Cenpk | 1.693778 | 82.8281 | 0.432361 | -3.9175 | 8.95E-05 | 0.000755 | 0.000914 | 127.988 | 124.9111 | 31.09143 | 47.32192 | 60411 | centromere protein K [Source:MGI Symbol;Acc:MGI:1926210] |
| ENSMUSG00000041859 | Mcm3 | 1.697918 | 1548.024 | 0.144704 | -11.7338 | 8.56E-32 | 2.28E-29 | 2.76E-29 | 2644.38 | 2089.064 | 733.392 | 725.2598 | 17215 | minichromosome maintenance complex component 3 [Source:MGI Symbol;Acc:MGI:101845] |
| ENSMUSG00000053398 | Phgdh | 1.699941 | 490.9433 | 0.176145 | -9.65083 | 4.88E-22 | 4.59E-20 | 5.56E-20 | 755.2349 | 746.5157 | 237.758 | 224.2647 | 236539 | 3-phosphoglycerate dehydrogenase [Source:MGI Symbol;Acc:MGI:1355330] |
| ENSMUSG00000028678 | Kif2c | 1.702334 | 482.7296 | 0.195273 | -8.71772 | 2.84E-18 | 1.68E-16 | 2.04E-16 | 809.1802 | 667.8316 | 220.3834 | 233.5234 | 73804 | kinesin family member 2C [Source:MGI Symbol;Acc:MGI:1921054] |
| ENSMUSG00000073705 | Cenps | 1.705623 | 48.59537 | 0.550123 | -3.10044 | 0.001932 | 0.010856 | 0.01315 | 87.79341 | 60.9802 | 21.94689 | 23.66096 | 69928 | centromere protein S [Source:MGI Symbol;Acc:MGI:1917178] |
| ENSMUSG00000028555 | Ttc39a | 1.706396 | 44.86489 | 0.600435 | -2.84193 | 0.004484 | 0.021912 | 0.026492 | 81.4469 | 56.06245 | 26.51916 | 15.43106 | 230603 | tetratricopeptide repeat domain 39A [Source:MGI Symbol;Acc:MGI:2444350] |
| ENSMUSG00000005233 | Spc25 | 1.710934 | 275.0021 | 0.278532 | -6.14268 | 8.11E-10 | 1.85E-08 | 2.24E-08 | 500.3167 | 342.276 | 121.6224 | 135.7933 | 66442 | SPC25, NDC80 kinetochore complex component, homolog (S. cerevisiae) [Source:MGI Symbol;Acc:MGI:1913692] |
| ENSMUSG00000034023 | Fancd2 | 1.711654 | 263.7521 | 0.241023 | -7.10162 | 1.23E-12 | 4.09E-11 | 4.96E-11 | 432.6205 | 375.7167 | 125.2802 | 121.391 | 211651 | Fanconi anemia, complementation group D2 [Source:MGI Symbol;Acc:MGI:2448480] |
| ENSMUSG00000050410 | Tcf19 | 1.712767 | 285.9554 | 0.23022 | -7.43969 | 1.01E-13 | 3.8E-12 | 4.6E-12 | 464.3531 | 412.1082 | 132.5958 | 134.7646 | 106795 | transcription factor 19 [Source:MGI Symbol;Acc:MGI:103180] |
| ENSMUSG00000030978 | Rrm1 | 1.71462 | 2236.245 | 0.148844 | -11.5195 | 1.05E-30 | 2.38E-28 | 2.88E-28 | 3915.798 | 2940.82 | 1086.371 | 1001.99 | 20133 | ribonucleotide reductase M1 [Source:MGI Symbol;Acc:MGI:98180] |
| ENSMUSG00000023919 | Cenpq | 1.714913 | 143.8647 | 0.323356 | -5.30348 | 1.14E-07 | 1.83E-06 | 2.22E-06 | 228.4744 | 212.4472 | 59.4395 | 75.09783 | 83815 | centromere protein Q [Source:MGI Symbol;Acc:MGI:1933744] |
| ENSMUSG00000091405 | H4c14 | 1.720437 | 446.2277 | 0.209815 | -8.19979 | 2.41E-16 | 1.17E-14 | 1.42E-14 | 764.7546 | 604.8843 | 216.7256 | 198.5463 | 97122 | H4 clustered histone 14 [Source:MGI Symbol;Acc:MGI:2140113] |
| ENSMUSG00000035365 | Parpbp | 1.730487 | 192.7264 | 0.32468 | -5.32982 | 9.83E-08 | 1.61E-06 | 1.95E-06 | 357.5201 | 235.0689 | 96.01765 | 82.29899 | 75317 | PARP1 binding protein [Source:MGI Symbol;Acc:MGI:1922567] |
| ENSMUSG00000030103 | Bhlhe40 | 1.730628 | 195.0799 | 0.271921 | -6.36445 | 1.96E-10 | 4.86E-09 | 5.89E-09 | 290.8818 | 308.8352 | 94.18874 | 86.41394 | 20893 | basic helix-loop-helix family, member e40 [Source:MGI Symbol;Acc:MGI:1097714] |
| ENSMUSG00000028175 | Depdc1a | 1.730844 | 166.8288 | 0.325742 | -5.31355 | 1.08E-07 | 1.74E-06 | 2.11E-06 | 299.3438 | 213.4307 | 72.24185 | 82.29899 | 76131 | DEP domain containing 1a [Source:MGI Symbol;Acc:MGI:1923381] |
| ENSMUSG00000027397 | Slc20a1 | 1.731062 | 1109.568 | 0.167622 | -10.3272 | 5.31E-25 | 6.71E-23 | 8.13E-23 | 1938.859 | 1472.377 | 532.2121 | 494.8227 | 20515 | solute carrier family 20, member 1 [Source:MGI Symbol;Acc:MGI:108392] |
| ENSMUSG00000026039 | Sgo2a | 1.731154 | 461.5773 | 0.209164 | -8.27655 | 1.27E-16 | 6.37E-15 | 7.71E-15 | 797.5449 | 621.6047 | 220.3834 | 206.7762 | 68549 | shugoshin 2A [Source:MGI Symbol;Acc:MGI:1098767] |
| ENSMUSG00000019303 | Psmc3ip | 1.731733 | 53.84279 | 0.522252 | -3.3159 | 0.000913 | 0.005808 | 0.007035 | 86.73566 | 78.68414 | 20.11798 | 29.83338 | 19183 | proteasome (prosome, macropain) 26S subunit, ATPase 3, interacting protein [Source:MGI Symbol;Acc:MGI:1098610] |
| ENSMUSG00000023015 | Racgap1 | 1.731801 | 937.3797 | 0.162957 | -10.6273 | 2.22E-26 | 3.44E-24 | 4.17E-24 | 1603.552 | 1278.617 | 443.5101 | 423.8398 | 26934 | Rac GTPase-activating protein 1 [Source:MGI Symbol;Acc:MGI:1349423] |
| ENSMUSG00000027331 | Knstrn | 1.734293 | 477.3466 | 0.207288 | -8.36657 | 5.93E-17 | 3.12E-15 | 3.78E-15 | 801.776 | 666.848 | 243.2447 | 197.5176 | 51944 | kinetochore-localized astrin/SPAG5 binding [Source:MGI Symbol;Acc:MGI:1289298] |
| ENSMUSG00000046223 | Plaur | 1.736886 | 221.1467 | 0.270765 | -6.41473 | 1.41E-10 | 3.62E-09 | 4.38E-09 | 377.6174 | 302.9339 | 104.2477 | 99.78752 | 18793 | plasminogen activator, urokinase receptor [Source:MGI Symbol;Acc:MGI:97612] |
| ENSMUSG00000038943 | Prc1 | 1.736952 | 917.2197 | 0.191113 | -9.08861 | 1E-19 | 7.24E-18 | 8.77E-18 | 1651.151 | 1171.41 | 438.9378 | 407.38 | 233406 | protein regulator of cytokinesis 1 [Source:MGI Symbol;Acc:MGI:1858961] |
| ENSMUSG00000007646 | Rad51c | 1.738128 | 36.71203 | 0.633892 | -2.742 | 0.006107 | 0.028334 | 0.034229 | 68.75387 | 44.25983 | 17.37462 | 16.4598 | 114714 | RAD51 paralog C [Source:MGI Symbol;Acc:MGI:2150020] |
| ENSMUSG00000031762 | Mt2 | 1.743684 | 97.96881 | 0.533448 | -3.2687 | 0.00108 | 0.006703 | 0.00812 | 80.38915 | 221.2991 | 43.89378 | 46.29318 | 17750 | metallothionein 2 [Source:MGI Symbol;Acc:MGI:97172] |
| ENSMUSG00000027994 | Mcub | 1.747633 | 134.1256 | 0.341483 | -5.11777 | 3.09E-07 | 4.53E-06 | 5.49E-06 | 217.8969 | 195.7268 | 70.41294 | 52.4656 | 66815 | mitochondrial calcium uniporter dominant negative beta subunit [Source:MGI Symbol;Acc:MGI:1914065] |
| ENSMUSG00000026274 | Pask | 1.751077 | 135.8268 | 0.402515 | -4.35034 | 1.36E-05 | 0.000142 | 0.000172 | 254.9182 | 164.2531 | 76.81412 | 47.32192 | 269224 | PAS domain containing serine/threonine kinase [Source:MGI Symbol;Acc:MGI:2155936] |
| ENSMUSG00000033762 | Recql4 | 1.759211 | 72.09404 | 0.438978 | -4.00752 | 6.14E-05 | 0.000543 | 0.000658 | 117.4105 | 105.24 | 33.83479 | 31.89086 | 79456 | RecQ protein-like 4 [Source:MGI Symbol;Acc:MGI:1931028] |
| ENSMUSG00000029730 | Mcm7 | 1.761106 | 939.359 | 0.181023 | -9.72862 | 2.28E-22 | 2.18E-20 | 2.65E-20 | 1672.306 | 1229.44 | 438.0234 | 417.6674 | 17220 | minichromosome maintenance complex component 7 [Source:MGI Symbol;Acc:MGI:1298398] |
| ENSMUSG00000048327 | Ckap2l | 1.762984 | 314.5989 | 0.288221 | -6.11677 | 9.55E-10 | 2.15E-08 | 2.6E-08 | 596.5721 | 375.7167 | 155.4571 | 130.6496 | 70466 | cytoskeleton associated protein 2-like [Source:MGI Symbol;Acc:MGI:1917716] |
| ENSMUSG00000034206 | Polq | 1.7674 | 272.3328 | 0.288396 | -6.12837 | 8.88E-10 | 2.01E-08 | 2.43E-08 | 508.7787 | 333.424 | 129.8524 | 117.2761 | 77782 | polymerase (DNA directed), theta [Source:MGI Symbol;Acc:MGI:2155399] |
| ENSMUSG00000003779 | Kif20a | 1.769229 | 692.664 | 0.180464 | -9.80376 | 1.08E-22 | 1.09E-20 | 1.32E-20 | 1184.682 | 957.9793 | 333.7756 | 294.2189 | 19348 | kinesin family member 20A [Source:MGI Symbol;Acc:MGI:1201682] |
| ENSMUSG00000037466 | Tedc1 | 1.770198 | 92.19181 | 0.395577 | -4.47498 | 7.64E-06 | 8.43E-05 | 0.000102 | 152.3163 | 132.7795 | 38.40706 | 45.26444 | 104732 | tubulin epsilon and delta complex 1 [Source:MGI Symbol;Acc:MGI:2144738] |
| ENSMUSG00000031273 | Col4a6 | 1.773251 | 78.31811 | 0.631053 | -2.80999 | 0.004954 | 0.023833 | 0.028807 | 59.23411 | 182.9406 | 23.7758 | 47.32192 | 94216 | collagen, type IV, alpha 6 [Source:MGI Symbol;Acc:MGI:2152695] |
| ENSMUSG00000027469 | Tpx2 | 1.775664 | 916.5417 | 0.174579 | -10.1711 | 2.67E-24 | 3.16E-22 | 3.83E-22 | 1603.552 | 1234.357 | 435.28 | 392.9777 | 72119 | TPX2, microtubule-associated [Source:MGI Symbol;Acc:MGI:1919369] |
| ENSMUSG00000099583 | H3c4 | 1.777368 | 387.1595 | 0.228762 | -7.7695 | 7.88E-15 | 3.32E-13 | 4.02E-13 | 664.2682 | 535.0521 | 192.9498 | 156.3681 | 319149 | H3 clustered histone 4 [Source:MGI Symbol;Acc:MGI:2448322] |
| ENSMUSG00000027306 | Nusap1 | 1.777741 | 645.6853 | 0.182051 | -9.76509 | 1.59E-22 | 1.56E-20 | 1.88E-20 | 1100.062 | 899.9498 | 309.0854 | 273.6441 | 108907 | nucleolar and spindle associated protein 1 [Source:MGI Symbol;Acc:MGI:2675669] |
| ENSMUSG00000022673 | Mcm4 | 1.778734 | 1001.625 | 0.168985 | -10.526 | 6.56E-26 | 9.47E-24 | 1.15E-23 | 1759.041 | 1343.532 | 455.398 | 448.5295 | 17217 | minichromosome maintenance complex component 4 [Source:MGI Symbol;Acc:MGI:103199] |
| ENSMUSG00000016758 | Bik | 1.778929 | 31.1635 | 0.680853 | -2.6128 | 0.00898 | 0.038993 | 0.0471 | 58.17636 | 38.35852 | 13.71681 | 14.40232 | 12124 | BCL2-interacting killer [Source:MGI Symbol;Acc:MGI:1206591] |
| ENSMUSG00000025059 | Gk | 1.78494 | 526.2943 | 0.201254 | -8.88178 | 6.58E-19 | 4.28E-17 | 5.19E-17 | 921.3019 | 711.1079 | 234.1002 | 238.6671 | 14933 | glycerol kinase [Source:MGI Symbol;Acc:MGI:106594] |
| ENSMUSG00000064220 | H2ac18 | 1.796465 | 791.3933 | 0.152944 | -11.7459 | 7.41E-32 | 2.04E-29 | 2.48E-29 | 1307.381 | 1150.755 | 356.637 | 350.7994 | 15267 | H2A clustered histone 18 [Source:MGI Symbol;Acc:MGI:96097] |
| ENSMUSG00000027203 | Dut | 1.79735 | 190.7393 | 0.317325 | -5.66407 | 1.48E-08 | 2.8E-07 | 3.39E-07 | 345.8849 | 246.8715 | 95.1032 | 75.09783 | 110074 | deoxyuridine triphosphatase [Source:MGI Symbol;Acc:MGI:1346051] |
| ENSMUSG00000050370 | Ch25h | 1.798864 | 370.6392 | 0.3385 | -5.31423 | 1.07E-07 | 1.74E-06 | 2.1E-06 | 783.7942 | 367.8483 | 159.115 | 171.7991 | 12642 | cholesterol 25-hydroxylase [Source:MGI Symbol;Acc:MGI:1333869] |
| ENSMUSG00000062510 | Nsl1 | 1.799159 | 133.3394 | 0.336014 | -5.35442 | 8.58E-08 | 1.43E-06 | 1.73E-06 | 228.4744 | 185.8913 | 60.35395 | 58.63803 | 381318 | NSL1, MIS12 kinetochore complex component [Source:MGI Symbol;Acc:MGI:2685830] |
| ENSMUSG00000074476 | Spc24 | 1.799368 | 127.3206 | 0.340799 | -5.27986 | 1.29E-07 | 2.06E-06 | 2.49E-06 | 212.6081 | 182.9406 | 53.03832 | 60.6955 | 67629 | SPC24, NDC80 kinetochore complex component, homolog (S. cerevisiae) [Source:MGI Symbol;Acc:MGI:1914879] |
| ENSMUSG00000024590 | Lmnb1 | 1.805375 | 560.6246 | 0.223466 | -8.07898 | 6.53E-16 | 3.08E-14 | 3.74E-14 | 1026.019 | 717.9927 | 267.0205 | 231.4659 | 16906 | lamin B1 [Source:MGI Symbol;Acc:MGI:96795] |
| ENSMUSG00000141158 | Gm74286 | 1.806561 | 121.9992 | 0.371082 | -4.86836 | 1.13E-06 | 1.47E-05 | 1.78E-05 | 205.2039 | 174.0886 | 43.89378 | 64.81045 | | predicted gene, 74286 [Source:MGI Symbol;Acc:MGI:7831310] |
| ENSMUSG00000022021 | Diaph3 | 1.807254 | 397.6461 | 0.236268 | -7.64917 | 2.02E-14 | 8.14E-13 | 9.86E-13 | 716.098 | 521.2824 | 186.5486 | 166.6555 | 56419 | diaphanous related formin 3 [Source:MGI Symbol;Acc:MGI:1927222] |
| ENSMUSG00000032783 | Troap | 1.816181 | 57.50386 | 0.506041 | -3.589 | 0.000332 | 0.002419 | 0.00293 | 87.79341 | 91.47031 | 30.17698 | 20.57475 | 78733 | trophinin associated protein [Source:MGI Symbol;Acc:MGI:1925983] |
| ENSMUSG00000027074 | Slc43a3 | 1.821442 | 228.4729 | 0.278457 | -6.54119 | 6.1E-11 | 1.63E-09 | 1.98E-09 | 399.8302 | 312.7694 | 109.7345 | 91.55762 | 58207 | solute carrier family 43, member 3 [Source:MGI Symbol;Acc:MGI:1931054] |
| ENSMUSG00000006378 | Gcat | 1.821573 | 192.5312 | 0.290023 | -6.28078 | 3.37E-10 | 8.06E-09 | 9.76E-09 | 330.0186 | 270.4767 | 91.44538 | 78.18404 | 26912 | glycine C-acetyltransferase (2-amino-3-ketobutyrate-coenzyme A ligase) [Source:MGI Symbol;Acc:MGI:1349389] |
| ENSMUSG00000001403 | Ube2c | 1.831993 | 268.6946 | 0.254806 | -7.18976 | 6.49E-13 | 2.23E-11 | 2.7E-11 | 467.5263 | 371.7825 | 124.3657 | 111.1036 | 68612 | ubiquitin-conjugating enzyme E2C [Source:MGI Symbol;Acc:MGI:1915862] |
| ENSMUSG00000026437 | Cdk18 | 1.832916 | 198.581 | 0.30191 | -6.07106 | 1.27E-09 | 2.82E-08 | 3.41E-08 | 360.6934 | 259.6576 | 89.61647 | 84.35646 | 18557 | cyclin dependent kinase 18 [Source:MGI Symbol;Acc:MGI:97518] |
| ENSMUSG00000059060 | Rad51b | 1.835613 | 40.78338 | 0.580579 | -3.16169 | 0.001569 | 0.009169 | 0.011107 | 64.52287 | 62.94731 | 19.20353 | 16.4598 | 19363 | RAD51 paralog B [Source:MGI Symbol;Acc:MGI:1099436] |
| ENSMUSG00000020330 | Hmmr | 1.837258 | 655.2378 | 0.201863 | -9.1015 | 8.91E-20 | 6.46E-18 | 7.83E-18 | 1183.624 | 864.5419 | 298.1119 | 274.6729 | 15366 | hyaluronan mediated motility receptor (RHAMM) [Source:MGI Symbol;Acc:MGI:104667] |
| ENSMUSG00000047757 | Fancb | 1.838082 | 78.52859 | 0.458815 | -4.00615 | 6.17E-05 | 0.000546 | 0.000661 | 149.143 | 96.38807 | 35.6637 | 32.9196 | 237211 | Fanconi anemia, complementation group B [Source:MGI Symbol;Acc:MGI:2448558] |
| ENSMUSG00000035860 | Cdhr3 | 1.842638 | 37.63337 | 0.611569 | -3.01297 | 0.002587 | 0.013798 | 0.016709 | 51.82984 | 65.89796 | 17.37462 | 15.43106 | 68764 | cadherin-related family member 3 [Source:MGI Symbol;Acc:MGI:1916014] |
| ENSMUSG00000037544 | Dlgap5 | 1.847639 | 493.4236 | 0.197536 | -9.35342 | 8.49E-21 | 6.96E-19 | 8.43E-19 | 850.4325 | 694.3875 | 223.1267 | 205.7475 | 218977 | DLG associated protein 5 [Source:MGI Symbol;Acc:MGI:2183453] |
| ENSMUSG00000029910 | Mad2l1 | 1.848957 | 493.7209 | 0.217597 | -8.49715 | 1.94E-17 | 1.07E-15 | 1.3E-15 | 893.8004 | 652.0948 | 213.9822 | 215.0061 | 56150 | MAD2 mitotic arrest deficient-like 1 [Source:MGI Symbol;Acc:MGI:1860374] |
| ENSMUSG00000028312 | Smc2 | 1.851204 | 1654.085 | 0.150864 | -12.2707 | 1.3E-34 | 4.7E-32 | 5.69E-32 | 2913.049 | 2268.07 | 756.2533 | 678.9667 | 14211 | structural maintenance of chromosomes 2 [Source:MGI Symbol;Acc:MGI:106067] |
| ENSMUSG00000018819 | Lsp1 | 1.851612 | 346.6422 | 0.272822 | -6.78687 | 1.15E-11 | 3.36E-10 | 4.07E-10 | 609.2651 | 477.0226 | 181.9763 | 118.3048 | 16985 | lymphocyte specific 1 [Source:MGI Symbol;Acc:MGI:96832] |
| ENSMUSG00000022102 | Dok2 | 1.85556 | 498.3298 | 0.187816 | -9.87967 | 5.1E-23 | 5.23E-21 | 6.33E-21 | 841.9705 | 719.9598 | 219.4689 | 211.9199 | 13449 | docking protein 2 [Source:MGI Symbol;Acc:MGI:1332623] |
| ENSMUSG00000031146 | Plp2 | 1.855666 | 211.8071 | 0.343313 | -5.40517 | 6.47E-08 | 1.11E-06 | 1.34E-06 | 422.043 | 241.9537 | 97.84656 | 85.3852 | 18824 | proteolipid protein 2 [Source:MGI Symbol;Acc:MGI:1298382] |
| ENSMUSG00000027496 | Aurka | 1.856544 | 371.2619 | 0.230426 | -8.05701 | 7.82E-16 | 3.67E-14 | 4.45E-14 | 626.1891 | 538.0028 | 181.9763 | 138.8795 | 20878 | aurora kinase A [Source:MGI Symbol;Acc:MGI:894678] |
| ENSMUSG00000047246 | H2bc6 | 1.856799 | 1260.719 | 0.173031 | -10.731 | 7.28E-27 | 1.16E-24 | 1.41E-24 | 2251.954 | 1700.561 | 593.4805 | 496.8801 | 319179 | H2B clustered histone 6 [Source:MGI Symbol;Acc:MGI:2448380] |
| ENSMUSG00000005470 | Asf1b | 1.857135 | 338.5303 | 0.216743 | -8.56839 | 1.05E-17 | 5.98E-16 | 7.25E-16 | 562.724 | 498.6607 | 151.7993 | 140.937 | 66929 | anti-silencing function 1B histone chaperone [Source:MGI Symbol;Acc:MGI:1914179] |
| ENSMUSG00000033031 | Cip2a | 1.860171 | 598.2399 | 0.211468 | -8.79645 | 1.41E-18 | 8.63E-17 | 1.05E-16 | 1079.965 | 796.6769 | 280.7373 | 235.5809 | 224171 | cell proliferation regulating inhibitor of protein phosphatase 2A [Source:MGI Symbol;Acc:MGI:2146335] |
| ENSMUSG00000036875 | Dna2 | 1.871432 | 433.0562 | 0.209789 | -8.92053 | 4.64E-19 | 3.06E-17 | 3.71E-17 | 749.9461 | 610.7856 | 196.6076 | 174.8853 | 327762 | DNA replication helicase/nuclease 2 [Source:MGI Symbol;Acc:MGI:2443732] |
| ENSMUSG00000049932 | H2ax | 1.871693 | 597.0818 | 0.213574 | -8.76369 | 1.89E-18 | 1.14E-16 | 1.38E-16 | 1099.004 | 777.0058 | 263.3627 | 248.9544 | 15270 | H2A.X variant histone [Source:MGI Symbol;Acc:MGI:102688] |
| ENSMUSG00000056737 | Capg | 1.876726 | 2853.79 | 0.13249 | -14.1651 | 1.51E-45 | 1.9E-42 | 2.31E-42 | 4910.084 | 4063.052 | 1326.872 | 1115.151 | 12332 | capping actin protein, gelsolin like [Source:MGI Symbol;Acc:MGI:1098259] |
| ENSMUSG00000000567 | Sox9 | 1.878739 | 34.24772 | 0.718157 | -2.61606 | 0.008895 | 0.038691 | 0.046736 | 33.84806 | 73.76638 | 11.8879 | 17.48853 | 20682 | SRY (sex determining region Y)-box 9 [Source:MGI Symbol;Acc:MGI:98371] |
| ENSMUSG00000028873 | Cdca8 | 1.884929 | 270.1028 | 0.235476 | -8.00477 | 1.2E-15 | 5.53E-14 | 6.7E-14 | 442.1403 | 408.174 | 117.9645 | 112.1324 | 52276 | cell division cycle associated 8 [Source:MGI Symbol;Acc:MGI:1196274] |
| ENSMUSG00000019992 | Mtfr2 | 1.887371 | 29.13718 | 0.722289 | -2.61304 | 0.008974 | 0.038978 | 0.047082 | 47.59884 | 44.25983 | 16.46017 | 8.229899 | 71804 | mitochondrial fission regulator 2 [Source:MGI Symbol;Acc:MGI:1919054] |
| ENSMUSG00000040084 | Bub1b | 1.887763 | 958.9115 | 0.194223 | -9.71959 | 2.49E-22 | 2.37E-20 | 2.87E-20 | 1669.133 | 1351.4 | 474.6015 | 340.5121 | 12236 | BUB1B, mitotic checkpoint serine/threonine kinase [Source:MGI Symbol;Acc:MGI:1333889] |
| ENSMUSG00000047798 | Cd300lf | 1.89156 | 65.26311 | 0.504962 | -3.74594 | 0.00018 | 0.001399 | 0.001695 | 126.9302 | 78.68414 | 25.60471 | 29.83338 | 246746 | CD300 molecule like family member F [Source:MGI Symbol;Acc:MGI:2442359] |
| ENSMUSG00000026683 | Nuf2 | 1.89504 | 541.3166 | 0.207571 | -9.12959 | 6.88E-20 | 5.06E-18 | 6.13E-18 | 972.074 | 734.7131 | 241.4158 | 217.0636 | 66977 | NUF2, NDC80 kinetochore complex component [Source:MGI Symbol;Acc:MGI:1914227] |
| ENSMUSG00000012443 | Kif11 | 1.895693 | 1390.907 | 0.158316 | -11.9741 | 4.86E-33 | 1.5E-30 | 1.82E-30 | 2403.212 | 1982.84 | 652.92 | 524.656 | 16551 | kinesin family member 11 [Source:MGI Symbol;Acc:MGI:1098231] |
| ENSMUSG00000021176 | Efcab11 | 1.898706 | 60.35852 | 0.485339 | -3.91212 | 9.15E-05 | 0.00077 | 0.000933 | 93.08217 | 97.37162 | 28.34807 | 22.63222 | 78767 | EF-hand calcium binding domain 11 [Source:MGI Symbol;Acc:MGI:1926017] |
| ENSMUSG00000046591 | Ticrr | 1.903628 | 105.8104 | 0.43612 | -4.36492 | 1.27E-05 | 0.000134 | 0.000162 | 207.3194 | 126.8782 | 53.03832 | 36.00581 | 77011 | TOPBP1-interacting checkpoint and replication regulator [Source:MGI Symbol;Acc:MGI:1924261] |
| ENSMUSG00000023940 | Sgo1 | 1.904359 | 184.7025 | 0.282488 | -6.74137 | 1.57E-11 | 4.52E-10 | 5.48E-10 | 306.7481 | 276.378 | 79.55748 | 76.12656 | 72415 | shugoshin 1 [Source:MGI Symbol;Acc:MGI:1919665] |
| ENSMUSG00000020897 | Aurkb | 1.906458 | 478.1497 | 0.212542 | -8.96981 | 2.97E-19 | 2.02E-17 | 2.45E-17 | 795.4294 | 715.0421 | 231.3568 | 170.7704 | 20877 | aurora kinase B [Source:MGI Symbol;Acc:MGI:107168] |
| ENSMUSG00000058385 | H2bc8 | 1.906573 | 1417.718 | 0.1756 | -10.8575 | 1.84E-27 | 3E-25 | 3.63E-25 | 2597.839 | 1879.567 | 641.0321 | 552.432 | 319181 | H2B clustered histone 8 [Source:MGI Symbol;Acc:MGI:2448386] |
| ENSMUSG00000068855 | H2ac20 | 1.907677 | 835.8203 | 0.166093 | -11.4856 | 1.56E-30 | 3.48E-28 | 4.21E-28 | 1459.698 | 1180.262 | 356.637 | 346.6845 | 319176 | H2A clustered histone 20 [Source:MGI Symbol;Acc:MGI:2448316] |
| ENSMUSG00000060093 | H4c1 | 1.908305 | 1007.879 | 0.165569 | -11.5257 | 9.79E-31 | 2.26E-28 | 2.74E-28 | 1790.774 | 1392.709 | 416.9909 | 431.041 | 326619 | H4 clustered histone 1 [Source:MGI Symbol;Acc:MGI:2448419] |
| ENSMUSG00000069265 | H3c1 | 1.909255 | 720.4402 | 0.206243 | -9.25731 | 2.1E-20 | 1.67E-18 | 2.03E-18 | 1257.667 | 1018.96 | 352.0647 | 253.0694 | 360198 | H3 clustered histone 1 [Source:MGI Symbol;Acc:MGI:2668828] |
| ENSMUSG00000038379 | Ttk | 1.911093 | 210.181 | 0.299043 | -6.3907 | 1.65E-10 | 4.16E-09 | 5.04E-09 | 385.0217 | 279.3287 | 95.1032 | 81.27025 | 22137 | Ttk protein kinase [Source:MGI Symbol;Acc:MGI:1194921] |
| ENSMUSG00000067455 | H4c11 | 1.911733 | 676.7806 | 0.181409 | -10.5383 | 5.76E-26 | 8.39E-24 | 1.02E-23 | 1188.913 | 950.1109 | 294.4541 | 273.6441 | 319159 | H4 clustered histone 11 [Source:MGI Symbol;Acc:MGI:2448436] |
| ENSMUSG00000024791 | Cdca5 | 1.914268 | 159.1623 | 0.326537 | -5.86233 | 4.56E-09 | 9.31E-08 | 1.13E-07 | 289.824 | 213.4307 | 68.58404 | 64.81045 | 67849 | cell division cycle associated 5 [Source:MGI Symbol;Acc:MGI:1915099] |
| ENSMUSG00000028337 | Coro2a | 1.914279 | 209.6708 | 0.291932 | -6.55728 | 5.48E-11 | 1.47E-09 | 1.78E-09 | 379.7329 | 283.2629 | 92.35984 | 83.32773 | 107684 | coronin, actin binding protein 2A [Source:MGI Symbol;Acc:MGI:1345966] |
| ENSMUSG00000026355 | Mcm6 | 1.915251 | 2264.264 | 0.136898 | -13.9903 | 1.79E-44 | 1.5E-41 | 1.82E-41 | 4000.418 | 3159.168 | 983.9523 | 913.5188 | 17219 | minichromosome maintenance complex component 6 [Source:MGI Symbol;Acc:MGI:1298227] |
| ENSMUSG00000056394 | Lig1 | 1.917087 | 558.6195 | 0.202742 | -9.45581 | 3.21E-21 | 2.72E-19 | 3.29E-19 | 1002.749 | 764.2197 | 243.2447 | 224.2647 | 16881 | ligase I, DNA, ATP-dependent [Source:MGI Symbol;Acc:MGI:101789] |
| ENSMUSG00000071516 | H2ac13 | 1.918762 | 201.8875 | 0.279636 | -6.86163 | 6.81E-12 | 2.07E-10 | 2.51E-10 | 343.7694 | 295.0655 | 90.53093 | 78.18404 | 319191 | H2A clustered histone 13 [Source:MGI Symbol;Acc:MGI:2448457] |
| ENSMUSG00000031137 | Fgf13 | 1.918926 | 32.08714 | 0.690671 | -2.77835 | 0.005464 | 0.025873 | 0.031254 | 60.29186 | 41.30917 | 16.46017 | 10.28737 | 14168 | fibroblast growth factor 13 [Source:MGI Symbol;Acc:MGI:109178] |
| ENSMUSG00000078773 | Rad54b | 1.92598 | 126.8513 | 0.340845 | -5.6506 | 1.6E-08 | 3.01E-07 | 3.64E-07 | 215.7814 | 185.8913 | 51.20941 | 54.52308 | 623474 | RAD54 homolog B (S. cerevisiae) [Source:MGI Symbol;Acc:MGI:3605986] |
| ENSMUSG00000024795 | Kif20b | 1.925998 | 676.1249 | 0.196954 | -9.77894 | 1.39E-22 | 1.37E-20 | 1.66E-20 | 1198.433 | 943.2261 | 311.8288 | 251.0119 | 240641 | kinesin family member 20B [Source:MGI Symbol;Acc:MGI:2444576] |
| ENSMUSG00000020493 | Prr11 | 1.927199 | 157.9141 | 0.373776 | -5.15603 | 2.52E-07 | 3.78E-06 | 4.57E-06 | 305.6903 | 194.7432 | 77.72857 | 53.49434 | 270906 | proline rich 11 [Source:MGI Symbol;Acc:MGI:2444496] |
| ENSMUSG00000048922 | Cdca2 | 1.928958 | 447.6551 | 0.213918 | -9.01728 | 1.93E-19 | 1.35E-17 | 1.63E-17 | 789.0829 | 629.4731 | 200.2654 | 171.7991 | 108912 | cell division cycle associated 2 [Source:MGI Symbol;Acc:MGI:1919787] |
| ENSMUSG00000028718 | Stil | 1.931524 | 181.2012 | 0.348858 | -5.5367 | 3.08E-08 | 5.59E-07 | 6.77E-07 | 339.5384 | 235.0689 | 90.53093 | 59.66677 | 20460 | Scl/Tal1 interrupting locus [Source:MGI Symbol;Acc:MGI:107477] |
| ENSMUSG00000074403 | H3c13 | 1.931633 | 677.871 | 0.25114 | -7.69147 | 1.45E-14 | 5.96E-13 | 7.22E-13 | 1234.396 | 914.7031 | 348.4069 | 213.9774 | 319154 | H3 clustered histone 13 [Source:MGI Symbol;Acc:MGI:2448351] |
| ENSMUSG00000037313 | Tacc3 | 1.933322 | 856.9348 | 0.186171 | -10.3847 | 2.91E-25 | 3.84E-23 | 4.65E-23 | 1535.856 | 1181.246 | 388.6429 | 321.9948 | 21335 | transforming, acidic coiled-coil containing protein 3 [Source:MGI Symbol;Acc:MGI:1341163] |
| ENSMUSG00000046245 | Pilra | 1.937677 | 73.63096 | 0.452323 | -4.28384 | 1.84E-05 | 0.000185 | 0.000224 | 133.2767 | 100.3223 | 31.09143 | 29.83338 | 231805 | paired immunoglobin-like type 2 receptor alpha [Source:MGI Symbol;Acc:MGI:2450529] |
| ENSMUSG00000031756 | Cenpn | 1.938324 | 152.9004 | 0.340871 | -5.68639 | 1.3E-08 | 2.5E-07 | 3.02E-07 | 284.5353 | 200.6445 | 66.75513 | 59.66677 | 72155 | centromere protein N [Source:MGI Symbol;Acc:MGI:1919405] |
| ENSMUSG00000039994 | Timeless | 1.93862 | 271.2556 | 0.29527 | -6.56557 | 5.18E-11 | 1.4E-09 | 1.7E-09 | 496.0856 | 364.8977 | 133.5103 | 90.52889 | 21853 | timeless circadian clock 1 [Source:MGI Symbol;Acc:MGI:1321393] |
| ENSMUSG00000025747 | Tyms | 1.939146 | 365.3544 | 0.218718 | -8.86596 | 7.58E-19 | 4.85E-17 | 5.88E-17 | 628.3046 | 531.1179 | 160.0294 | 141.9658 | 22171 | thymidylate synthase [Source:MGI Symbol;Acc:MGI:98878] |
| ENSMUSG00000030867 | Plk1 | 1.93924 | 188.6046 | 0.351145 | -5.52262 | 3.34E-08 | 6.01E-07 | 7.28E-07 | 373.3864 | 225.2333 | 86.87311 | 68.9254 | 18817 | polo like kinase 1 [Source:MGI Symbol;Acc:MGI:97621] |
| ENSMUSG00000020808 | Pimreg | 1.941434 | 266.5009 | 0.270817 | -7.1688 | 7.57E-13 | 2.57E-11 | 3.11E-11 | 467.5263 | 378.6674 | 126.1946 | 93.6151 | 109212 | PICALM interacting mitotic regulator [Source:MGI Symbol;Acc:MGI:1924434] |
| ENSMUSG00000037572 | Wdhd1 | 1.941652 | 429.8277 | 0.226941 | -8.55576 | 1.17E-17 | 6.63E-16 | 8.03E-16 | 784.8519 | 579.3119 | 171.0029 | 184.144 | 218973 | WD repeat and HMG-box DNA binding protein 1 [Source:MGI Symbol;Acc:MGI:2443514] |
| ENSMUSG00000026429 | Ube2t | 1.951476 | 75.6294 | 0.439121 | -4.44406 | 8.83E-06 | 9.63E-05 | 0.000117 | 131.1612 | 109.1742 | 29.26252 | 32.9196 | 67196 | ubiquitin-conjugating enzyme E2T [Source:MGI Symbol;Acc:MGI:1914446] |
| ENSMUSG00000031821 | Gins2 | 1.95223 | 294.1426 | 0.252633 | -7.72753 | 1.1E-14 | 4.56E-13 | 5.52E-13 | 529.9337 | 405.2233 | 126.1946 | 115.2186 | 272551 | GINS complex subunit 2 [Source:MGI Symbol;Acc:MGI:1921019] |
| ENSMUSG00000032586 | Traip | 1.956704 | 105.0065 | 0.388655 | -5.03455 | 4.79E-07 | 6.76E-06 | 8.19E-06 | 191.4531 | 142.615 | 44.80824 | 41.14949 | 22036 | TRAF-interacting protein [Source:MGI Symbol;Acc:MGI:1096377] |
| ENSMUSG00000027326 | Knl1 | 1.967962 | 741.792 | 0.229893 | -8.56032 | 1.13E-17 | 6.39E-16 | 7.74E-16 | 1365.558 | 998.305 | 358.4659 | 244.8395 | 76464 | kinetochore scaffold 1 [Source:MGI Symbol;Acc:MGI:1923714] |
| ENSMUSG00000021569 | Trip13 | 1.972728 | 154.8801 | 0.370428 | -5.32553 | 1.01E-07 | 1.65E-06 | 1.99E-06 | 298.286 | 195.7268 | 74.07076 | 51.43687 | 69716 | thyroid hormone receptor interactor 13 [Source:MGI Symbol;Acc:MGI:1916966] |
| ENSMUSG00000041064 | Pif1 | 1.973753 | 135.484 | 0.329184 | -5.9959 | 2.02E-09 | 4.35E-08 | 5.26E-08 | 214.7236 | 217.3649 | 59.4395 | 50.40813 | 208084 | PIF1 5'-to-3' DNA helicase [Source:MGI Symbol;Acc:MGI:2143057] |
| ENSMUSG00000026779 | Mastl | 1.983425 | 353.9832 | 0.254089 | -7.80603 | 5.9E-15 | 2.53E-13 | 3.06E-13 | 663.2104 | 467.1871 | 151.7993 | 133.7359 | 67121 | microtubule associated serine/threonine kinase-like [Source:MGI Symbol;Acc:MGI:1914371] |
| ENSMUSG00000004891 | Nes | 1.990035 | 609.6436 | 0.167981 | -11.8468 | 2.24E-32 | 6.52E-30 | 7.9E-30 | 922.3597 | 1025.844 | 249.6459 | 240.7245 | 18008 | nestin [Source:MGI Symbol;Acc:MGI:101784] |
| ENSMUSG00000021508 | Cxcl14 | 2.002752 | 305.5894 | 0.257182 | -7.7873 | 6.85E-15 | 2.92E-13 | 3.53E-13 | 559.5508 | 418.993 | 131.6814 | 112.1324 | 57266 | C-X-C motif chemokine ligand 14 [Source:MGI Symbol;Acc:MGI:1888514] |
| ENSMUSG00000029516 | Cit | 2.005052 | 199.8201 | 0.37917 | -5.288 | 1.24E-07 | 1.97E-06 | 2.39E-06 | 392.426 | 247.855 | 102.4188 | 56.58055 | 12704 | citron [Source:MGI Symbol;Acc:MGI:105313] |
| ENSMUSG00000069268 | H2bc7 | 2.017055 | 266.5953 | 0.326452 | -6.17871 | 6.46E-10 | 1.5E-08 | 1.81E-08 | 503.4899 | 352.1115 | 132.5958 | 78.18404 | 319180 | H2B clustered histone 7 [Source:MGI Symbol;Acc:MGI:2448383] |
| ENSMUSG00000032218 | Ccnb2 | 2.019341 | 645.0878 | 0.215065 | -9.38942 | 6.03E-21 | 5E-19 | 6.06E-19 | 1138.141 | 932.407 | 299.9409 | 209.8624 | 12442 | cyclin B2 [Source:MGI Symbol;Acc:MGI:88311] |
| ENSMUSG00000035683 | Melk | 2.020098 | 296.97 | 0.252661 | -7.99529 | 1.29E-15 | 5.9E-14 | 7.15E-14 | 522.5294 | 430.7956 | 131.6814 | 102.8737 | 17279 | maternal embryonic leucine zipper kinase [Source:MGI Symbol;Acc:MGI:106924] |
| ENSMUSG00000037225 | Fgf2 | 2.023884 | 50.0262 | 0.536881 | -3.76971 | 0.000163 | 0.001286 | 0.001558 | 75.10039 | 85.569 | 21.94689 | 17.48853 | 14173 | fibroblast growth factor 2 [Source:MGI Symbol;Acc:MGI:95516] |
| ENSMUSG00000031506 | Ptpn7 | 2.024799 | 187.4474 | 0.322924 | -6.2702 | 3.61E-10 | 8.6E-09 | 1.04E-08 | 340.5961 | 261.6248 | 86.87311 | 60.6955 | 320139 | protein tyrosine phosphatase, non-receptor type 7 [Source:MGI Symbol;Acc:MGI:2156893] |
| ENSMUSG00000032400 | Zwilch | 2.033504 | 262.4676 | 0.274288 | -7.41375 | 1.23E-13 | 4.56E-12 | 5.53E-12 | 490.7969 | 353.0951 | 105.1622 | 100.8163 | 68014 | zwilch kinetochore protein [Source:MGI Symbol;Acc:MGI:1915264] |
| ENSMUSG00000036768 | Kif15 | 2.035177 | 600.5102 | 0.198681 | -10.2435 | 1.27E-24 | 1.52E-22 | 1.85E-22 | 1088.427 | 842.9038 | 250.5603 | 220.1498 | 209737 | kinesin family member 15 [Source:MGI Symbol;Acc:MGI:1098258] |
| ENSMUSG00000024056 | Ndc80 | 2.036485 | 384.3256 | 0.261762 | -7.77992 | 7.26E-15 | 3.07E-13 | 3.72E-13 | 704.4628 | 532.1015 | 178.3185 | 122.4197 | 67052 | NDC80 kinetochore complex component [Source:MGI Symbol;Acc:MGI:1914302] |
| ENSMUSG00000020914 | Top2a | 2.047125 | 4853.831 | 0.13665 | -14.9808 | 9.8E-51 | 2.96E-47 | 3.58E-47 | 8943.292 | 6690.119 | 1973.391 | 1808.52 | 21973 | topoisomerase (DNA) II alpha [Source:MGI Symbol;Acc:MGI:98790] |
| ENSMUSG00000069910 | Spdl1 | 2.047885 | 346.4754 | 0.245326 | -8.3476 | 6.97E-17 | 3.64E-15 | 4.41E-15 | 631.4779 | 484.891 | 148.1415 | 121.391 | 70385 | spindle apparatus coiled-coil protein 1 [Source:MGI Symbol;Acc:MGI:1917635] |
| ENSMUSG00000022945 | Chaf1b | 2.048396 | 251.1154 | 0.26879 | -7.6208 | 2.52E-14 | 1E-12 | 1.21E-12 | 445.3136 | 363.9141 | 108.82 | 86.41394 | 110749 | chromatin assembly factor 1, subunit B [Source:MGI Symbol;Acc:MGI:1314881] |
| ENSMUSG00000041498 | Kif14 | 2.051674 | 245.2557 | 0.319908 | -6.41334 | 1.42E-10 | 3.64E-09 | 4.41E-09 | 463.2953 | 327.5227 | 116.1356 | 74.06909 | 381293 | kinesin family member 14 [Source:MGI Symbol;Acc:MGI:1098226] |
| ENSMUSG00000045328 | Cenpe | 2.056747 | 1238.733 | 0.20058 | -10.254 | 1.14E-24 | 1.38E-22 | 1.67E-22 | 2278.398 | 1717.281 | 564.218 | 395.0351 | 229841 | centromere protein E [Source:MGI Symbol;Acc:MGI:1098230] |
| ENSMUSG00000001473 | Tubb6 | 2.05763 | 1047.324 | 0.149112 | -13.7993 | 2.58E-43 | 1.86E-40 | 2.25E-40 | 1821.449 | 1556.962 | 426.1355 | 384.7478 | 67951 | tubulin, beta 6 class V [Source:MGI Symbol;Acc:MGI:1915201] |
| ENSMUSG00000036777 | Anln | 2.058162 | 743.4704 | 0.19403 | -10.6074 | 2.75E-26 | 4.21E-24 | 5.11E-24 | 1343.345 | 1055.351 | 320.0588 | 255.1269 | 68743 | anillin, actin binding protein [Source:MGI Symbol;Acc:MGI:1920174] |
| ENSMUSG00000028044 | Cks1b | 2.06224 | 388.3055 | 0.224264 | -9.19558 | 3.73E-20 | 2.92E-18 | 3.53E-18 | 704.4628 | 548.8218 | 151.7993 | 148.1382 | 54124 | CDC28 protein kinase 1b [Source:MGI Symbol;Acc:MGI:1889208] |
| ENSMUSG00000042498 | Radx | 2.066855 | 92.90198 | 0.407191 | -5.07588 | 3.86E-07 | 5.53E-06 | 6.7E-06 | 169.2403 | 130.8124 | 36.57815 | 34.97707 | 102871 | RPA1 related single stranded DNA binding protein, X-linked [Source:MGI Symbol;Acc:MGI:2147848] |
| ENSMUSG00000003824 | Syce2 | 2.067139 | 159.0343 | 0.348778 | -5.9268 | 3.09E-09 | 6.47E-08 | 7.84E-08 | 273.9577 | 239.9866 | 75.89967 | 46.29318 | 71846 | synaptonemal complex central element protein 2 [Source:MGI Symbol;Acc:MGI:1919096] |
| ENSMUSG00000027454 | Gins1 | 2.073905 | 186.8998 | 0.294012 | -7.05381 | 1.74E-12 | 5.65E-11 | 6.85E-11 | 315.2101 | 289.1642 | 80.47194 | 62.75298 | 69270 | GINS complex subunit 1 [Source:MGI Symbol;Acc:MGI:1916520] |
| ENSMUSG00000029283 | Cdc7 | 2.081068 | 191.9866 | 0.314657 | -6.61376 | 3.75E-11 | 1.03E-09 | 1.25E-09 | 335.3074 | 286.2135 | 87.78757 | 58.63803 | 12545 | cell division cycle 7 [Source:MGI Symbol;Acc:MGI:1309511] |
| ENSMUSG00000069272 | H2ac8 | 2.082707 | 207.8866 | 0.275686 | -7.55464 | 4.2E-14 | 1.63E-12 | 1.98E-12 | 364.9244 | 307.8517 | 79.55748 | 79.21278 | 319166 | H2A clustered histone 8 [Source:MGI Symbol;Acc:MGI:2448290] |
| ENSMUSG00000048416 | Mlf1 | 2.083784 | 36.90788 | 0.642651 | -3.24248 | 0.001185 | 0.007239 | 0.008769 | 47.59884 | 71.79927 | 12.80235 | 15.43106 | 17349 | myeloid leukemia factor 1 [Source:MGI Symbol;Acc:MGI:1341819] |
| ENSMUSG00000069274 | H4c6 | 2.086156 | 1302.1 | 0.193388 | -10.7874 | 3.95E-27 | 6.37E-25 | 7.72E-25 | 2465.62 | 1750.722 | 556.9024 | 435.1559 | 319157 | H4 clustered histone 6 [Source:MGI Symbol;Acc:MGI:2448425] |
| ENSMUSG00000101972 | H3c11 | 2.086686 | 642.9746 | 0.196918 | -10.5967 | 3.09E-26 | 4.63E-24 | 5.61E-24 | 1099.004 | 983.5517 | 282.5662 | 206.7762 | 319153 | H3 clustered histone 11 [Source:MGI Symbol;Acc:MGI:2448350] |
| ENSMUSG00000101355 | H3c10 | 2.087599 | 609.0872 | 0.227844 | -9.16242 | 5.07E-20 | 3.83E-18 | 4.64E-18 | 1134.968 | 837.986 | 267.935 | 195.4601 | 319152 | H3 clustered histone 10 [Source:MGI Symbol;Acc:MGI:2448349] |
| ENSMUSG00000028702 | Rad54l | 2.092075 | 194.0562 | 0.294832 | -7.09583 | 1.29E-12 | 4.25E-11 | 5.15E-11 | 343.7694 | 285.23 | 81.38639 | 65.83919 | 19366 | RAD54 like (S. cerevisiae) [Source:MGI Symbol;Acc:MGI:894697] |
| ENSMUSG00000002870 | Mcm2 | 2.094349 | 1126.854 | 0.157429 | -13.3034 | 2.21E-40 | 1.24E-37 | 1.5E-37 | 2038.288 | 1614.008 | 426.1355 | 428.9835 | 17216 | minichromosome maintenance complex component 2 [Source:MGI Symbol;Acc:MGI:105380] |
| ENSMUSG00000063021 | H2ac15 | 2.096099 | 438.5547 | 0.22192 | -9.4453 | 3.54E-21 | 2.99E-19 | 3.62E-19 | 809.1802 | 612.7527 | 172.8318 | 159.4543 | 319169 | H2A clustered histone 15 [Source:MGI Symbol;Acc:MGI:2448297] |
| ENSMUSG00000024989 | Cep55 | 2.096539 | 364.6072 | 0.214917 | -9.75512 | 1.75E-22 | 1.7E-20 | 2.07E-20 | 631.4779 | 550.7889 | 144.4837 | 131.6784 | 74107 | centrosomal protein 55 [Source:MGI Symbol;Acc:MGI:1921357] |
| ENSMUSG00000039055 | Eme1 | 2.110486 | 106.7727 | 0.425602 | -4.95882 | 7.09E-07 | 9.64E-06 | 1.17E-05 | 194.6264 | 152.4505 | 51.20941 | 28.80465 | 268465 | essential meiotic structure-specific endonuclease 1 [Source:MGI Symbol;Acc:MGI:3576783] |
| ENSMUSG00000026622 | Nek2 | 2.115492 | 402.3526 | 0.232096 | -9.11475 | 7.89E-20 | 5.75E-18 | 6.96E-18 | 702.3473 | 605.8678 | 174.6607 | 126.5347 | 18005 | NIMA (never in mitosis gene a)-related expressed kinase 2 [Source:MGI Symbol;Acc:MGI:109359] |
| ENSMUSG00000069008 | Gm5537 | 2.117856 | 172.2237 | 0.449162 | -4.71513 | 2.42E-06 | 2.93E-05 | 3.54E-05 | 320.4988 | 239.0031 | 34.74925 | 94.64384 | | predicted gene 5537 [Source:MGI Symbol;Acc:MGI:3643179] |
| ENSMUSG00000015880 | Ncapg | 2.118831 | 685.6305 | 0.203595 | -10.4071 | 2.3E-25 | 3.12E-23 | 3.78E-23 | 1264.014 | 965.8478 | 285.3096 | 227.351 | 54392 | non-SMC condensin I complex, subunit G [Source:MGI Symbol;Acc:MGI:1930197] |
| ENSMUSG00000037628 | Cdkn3 | 2.120704 | 136.1019 | 0.422674 | -5.01735 | 5.24E-07 | 7.33E-06 | 8.88E-06 | 244.3407 | 198.6774 | 69.49849 | 31.89086 | 72391 | cyclin dependent kinase inhibitor 3 [Source:MGI Symbol;Acc:MGI:1919641] |
| ENSMUSG00000002055 | Spag5 | 2.132914 | 475.84 | 0.213578 | -9.98658 | 1.75E-23 | 1.85E-21 | 2.25E-21 | 847.2593 | 703.2395 | 197.522 | 155.3393 | 54141 | sperm associated antigen 5 [Source:MGI Symbol;Acc:MGI:1927470] |
| ENSMUSG00000124938 | Gm70475 | 2.133472 | 48.11344 | 0.55917 | -3.81542 | 0.000136 | 0.001091 | 0.001322 | 89.90891 | 66.88152 | 19.20353 | 16.4598 | | predicted gene, 70475 [Source:MGI Symbol;Acc:MGI:7823751] |
| ENSMUSG00000032113 | Chek1 | 2.138151 | 141.9298 | 0.37877 | -5.64499 | 1.65E-08 | 3.1E-07 | 3.76E-07 | 284.5353 | 178.0229 | 47.5516 | 57.60929 | 12649 | checkpoint kinase 1 [Source:MGI Symbol;Acc:MGI:1202065] |
| ENSMUSG00000026193 | Fn1 | 2.13848 | 6991.75 | 0.442385 | -4.83398 | 1.34E-06 | 1.72E-05 | 2.08E-05 | 14792.66 | 7998.242 | 2778.111 | 2397.987 | 14268 | fibronectin 1 [Source:MGI Symbol;Acc:MGI:95566] |
| ENSMUSG00000100210 | H3c7 | 2.141578 | 1325.629 | 0.149122 | -14.3612 | 9.06E-47 | 1.37E-43 | 1.66E-43 | 2402.155 | 1920.876 | 492.8906 | 486.5928 | 260423 | H3 clustered histone 7 [Source:MGI Symbol;Acc:MGI:2448329] |
| ENSMUSG00000105827 | H2bc18 | 2.141581 | 925.4508 | 0.222865 | -9.6093 | 7.3E-22 | 6.75E-20 | 8.18E-20 | 1727.309 | 1291.403 | 411.5042 | 271.5867 | 319189 | H2B clustered histone 18 [Source:MGI Symbol;Acc:MGI:2448413] |
| ENSMUSG00000094248 | H2ac23 | 2.145094 | 226.3072 | 0.330535 | -6.48977 | 8.6E-11 | 2.26E-09 | 2.74E-09 | 276.0732 | 462.2693 | 88.70202 | 78.18404 | 665433 | H2A clustered histone 23 [Source:MGI Symbol;Acc:MGI:2448302] |
| ENSMUSG00000027715 | Ccna2 | 2.147737 | 1519.791 | 0.152973 | -14.04 | 8.87E-45 | 8.97E-42 | 1.09E-41 | 2781.888 | 2178.567 | 588.9083 | 529.7997 | 12428 | cyclin A2 [Source:MGI Symbol;Acc:MGI:108069] |
| ENSMUSG00000027315 | Spint1 | 2.151195 | 24.56091 | 0.810378 | -2.65456 | 0.007941 | 0.03524 | 0.042564 | 37.02132 | 43.27627 | 12.80235 | 5.143687 | 20732 | serine protease inhibitor, Kunitz type 1 [Source:MGI Symbol;Acc:MGI:1338033] |
| ENSMUSG00000111713 | Gm20234 | 2.164963 | 120.4015 | 0.382402 | -5.66148 | 1.5E-08 | 2.84E-07 | 3.44E-07 | 231.6477 | 162.286 | 47.5516 | 40.12076 | | predicted gene, 20234 [Source:MGI Symbol;Acc:MGI:5012419] |
| ENSMUSG00000002835 | Chaf1a | 2.170726 | 310.8774 | 0.232311 | -9.34406 | 9.27E-21 | 7.56E-19 | 9.16E-19 | 543.6845 | 474.0719 | 119.7935 | 105.9599 | 27221 | chromatin assembly factor 1, subunit A [Source:MGI Symbol;Acc:MGI:1351331] |
| ENSMUSG00000039748 | Exo1 | 2.173475 | 170.6705 | 0.325873 | -6.6697 | 2.56E-11 | 7.22E-10 | 8.75E-10 | 325.7876 | 233.1018 | 63.09731 | 60.6955 | 26909 | exonuclease 1 [Source:MGI Symbol;Acc:MGI:1349427] |
| ENSMUSG00000072980 | Oip5 | 2.174182 | 107.6409 | 0.378504 | -5.74414 | 9.24E-09 | 1.81E-07 | 2.2E-07 | 189.3376 | 163.2696 | 42.97933 | 34.97707 | 70645 | Opa interacting protein 5 [Source:MGI Symbol;Acc:MGI:1917895] |
| ENSMUSG00000020185 | E2f7 | 2.179372 | 63.72646 | 0.504205 | -4.32239 | 1.54E-05 | 0.000159 | 0.000193 | 118.4682 | 90.48676 | 27.43361 | 18.51727 | 52679 | E2F transcription factor 7 [Source:MGI Symbol;Acc:MGI:1289147] |
| ENSMUSG00000001517 | Foxm1 | 2.181046 | 223.1963 | 0.32563 | -6.69794 | 2.11E-11 | 5.99E-10 | 7.26E-10 | 435.7938 | 296.0491 | 95.1032 | 65.83919 | 14235 | forkhead box M1 [Source:MGI Symbol;Acc:MGI:1347487] |
| ENSMUSG00000021707 | Dhfr | 2.181416 | 293.67 | 0.250855 | -8.69593 | 3.44E-18 | 2.03E-16 | 2.46E-16 | 541.569 | 420.9601 | 105.1622 | 106.9887 | 13361 | dihydrofolate reductase [Source:MGI Symbol;Acc:MGI:94890] |
| ENSMUSG00000019214 | Chtf18 | 2.183508 | 108.2207 | 0.400941 | -5.44596 | 5.15E-08 | 8.96E-07 | 1.09E-06 | 208.3771 | 146.5492 | 42.97933 | 34.97707 | 214901 | CTF18, chromosome transmission fidelity factor 18 [Source:MGI Symbol;Acc:MGI:2384887] |
| ENSMUSG00000032254 | Kif23 | 2.185236 | 507.7867 | 0.188805 | -11.574 | 5.58E-31 | 1.34E-28 | 1.63E-28 | 886.3961 | 778.9729 | 192.9498 | 172.8279 | 71819 | kinesin family member 23 [Source:MGI Symbol;Acc:MGI:1919069] |
| ENSMUSG00000034883 | Lrr1 | 2.197583 | 28.30567 | 0.710131 | -3.09462 | 0.001971 | 0.011022 | 0.013352 | 49.71434 | 43.27627 | 10.97345 | 9.258636 | 69706 | leucine rich repeat protein 1 [Source:MGI Symbol;Acc:MGI:1916956] |
| ENSMUSG00000017716 | Birc5 | 2.198633 | 417.0144 | 0.230198 | -9.55104 | 1.28E-21 | 1.12E-19 | 1.36E-19 | 758.4081 | 611.7692 | 168.2595 | 129.6209 | 11799 | baculoviral IAP repeat-containing 5 [Source:MGI Symbol;Acc:MGI:1203517] |
| ENSMUSG00000075266 | Cenpw | 2.202392 | 86.5706 | 0.422881 | -5.20807 | 1.91E-07 | 2.93E-06 | 3.55E-06 | 158.6628 | 125.8946 | 32.92034 | 28.80465 | 66311 | centromere protein W [Source:MGI Symbol;Acc:MGI:1913561] |
| ENSMUSG00000025001 | Hells | 2.205889 | 525.4718 | 0.210937 | -10.4576 | 1.35E-25 | 1.9E-23 | 2.3E-23 | 985.8248 | 741.598 | 181.0619 | 193.4026 | 15201 | helicase, lymphoid specific [Source:MGI Symbol;Acc:MGI:106209] |
| ENSMUSG00000026196 | Bard1 | 2.210032 | 144.3345 | 0.370913 | -5.95835 | 2.55E-09 | 5.4E-08 | 6.54E-08 | 290.8818 | 183.9242 | 52.12387 | 50.40813 | 12021 | BRCA1 associated RING domain 1 [Source:MGI Symbol;Acc:MGI:1328361] |
| ENSMUSG00000026069 | Il1rl1 | 2.215118 | 26.91505 | 0.760083 | -2.91431 | 0.003565 | 0.018078 | 0.021877 | 44.42558 | 44.25983 | 12.80235 | 6.172424 | 17082 | interleukin 1 receptor-like 1 [Source:MGI Symbol;Acc:MGI:98427] |
| ENSMUSG00000021965 | Ska3 | 2.221914 | 138.3229 | 0.326364 | -6.80808 | 9.89E-12 | 2.92E-10 | 3.53E-10 | 228.4744 | 227.2004 | 50.29496 | 47.32192 | 219114 | spindle and kinetochore associated complex subunit 3 [Source:MGI Symbol;Acc:MGI:3041235] |
| ENSMUSG00000031262 | Cenpi | 2.222618 | 278.6168 | 0.250501 | -8.87268 | 7.14E-19 | 4.59E-17 | 5.56E-17 | 507.7209 | 410.1411 | 97.84656 | 98.75879 | 102920 | centromere protein I [Source:MGI Symbol;Acc:MGI:2147897] |
| ENSMUSG00000037474 | Dtl | 2.223143 | 275.0684 | 0.27459 | -8.09623 | 5.67E-16 | 2.69E-14 | 3.26E-14 | 526.7605 | 379.651 | 103.3333 | 90.52889 | 76843 | denticleless E3 ubiquitin protein ligase [Source:MGI Symbol;Acc:MGI:1924093] |
| ENSMUSG00000064288 | H4c12 | 2.224416 | 511.3668 | 0.29211 | -7.61499 | 2.64E-14 | 1.04E-12 | 1.26E-12 | 1033.424 | 652.0948 | 223.1267 | 136.8221 | 319160 | H4 clustered histone 12 [Source:MGI Symbol;Acc:MGI:2448439] |
| ENSMUSG00000078452 | Raet1d;Raet1d; | 2.23136 | 94.09903 | 0.445348 | -5.01038 | 5.43E-07 | 7.58E-06 | 9.19E-06 | 178.7601 | 131.7959 | 41.15042 | 24.6897 | 19368;19369; | retinoic acid early transcript delta [Source:MGI Symbol;Acc:MGI:1861032];retinoic acid early transcript delta [Source:MGI Symbol;Acc:MGI:1861032];retinoic acid early transcript delta [Source:MGI Symbol;Acc:MGI:1861032];retinoic acid early transcript delta [Source:MGI Symbol;Acc:MGI:1861032] |
| ENSMUSG00000017724 | Etv4 | 2.232245 | 62.46965 | 0.543495 | -4.10721 | 4E-05 | 0.000377 | 0.000456 | 132.219 | 73.76638 | 19.20353 | 24.6897 | 18612 | ets variant 4 [Source:MGI Symbol;Acc:MGI:99423] |
| ENSMUSG00000022322 | Shcbp1 | 2.234221 | 334.4797 | 0.301232 | -7.41694 | 1.2E-13 | 4.48E-12 | 5.42E-12 | 650.5174 | 453.4173 | 144.4837 | 89.50015 | 20419 | Shc SH2-domain binding protein 1 [Source:MGI Symbol;Acc:MGI:1338802] |
| ENSMUSG00000049539 | H1f1 | 2.234984 | 1591.617 | 0.176875 | -12.6359 | 1.34E-36 | 5.8E-34 | 7.02E-34 | 2901.413 | 2350.689 | 648.3478 | 466.018 | 80838 | H1.1 linker histone, cluster member [Source:MGI Symbol;Acc:MGI:1931523] |
| ENSMUSG00000094777 | H2ac24 | 2.239458 | 967.7473 | 0.227648 | -9.83738 | 7.77E-23 | 7.91E-21 | 9.58E-21 | 1981.169 | 1213.703 | 360.2948 | 315.8224 | 319171 | H2A clustered histone 24 [Source:MGI Symbol;Acc:MGI:3710573] |
| ENSMUSG00000019773 | Fbxo5 | 2.240682 | 229.7551 | 0.345727 | -6.48107 | 9.11E-11 | 2.38E-09 | 2.88E-09 | 444.2558 | 314.7365 | 102.4188 | 57.60929 | 67141 | F-box protein 5 [Source:MGI Symbol;Acc:MGI:1914391] |
| ENSMUSG00000071478 | H2ac7 | 2.257384 | 435.6845 | 0.224322 | -10.0631 | 8.04E-24 | 9.16E-22 | 1.11E-21 | 797.5449 | 644.2264 | 168.2595 | 132.7071 | 319165 | H2A clustered histone 7 [Source:MGI Symbol;Acc:MGI:2448289] |
| ENSMUSG00000034311 | Kif4 | 2.259504 | 590.9905 | 0.24626 | -9.17529 | 4.5E-20 | 3.45E-18 | 4.18E-18 | 1134.968 | 821.2657 | 244.1592 | 163.5692 | 16571 | kinesin family member 4 [Source:MGI Symbol;Acc:MGI:108389] |
| ENSMUSG00000042489 | Clspn | 2.270511 | 314.6104 | 0.298197 | -7.61413 | 2.65E-14 | 1.05E-12 | 1.27E-12 | 580.7058 | 462.2693 | 136.2536 | 79.21278 | 269582 | claspin [Source:MGI Symbol;Acc:MGI:2445153] |
| ENSMUSG00000051378 | Kif18b | 2.270649 | 228.5491 | 0.333436 | -6.80985 | 9.77E-12 | 2.89E-10 | 3.5E-10 | 440.0248 | 317.6872 | 97.84656 | 58.63803 | 70218 | kinesin family member 18B [Source:MGI Symbol;Acc:MGI:2446979] |
| ENSMUSG00000095217 | H2bc15 | 2.271841 | 578.6892 | 0.226457 | -10.0321 | 1.1E-23 | 1.22E-21 | 1.48E-21 | 1124.39 | 793.7262 | 217.64 | 179.0003 | 319187 | H2B clustered histone 15 [Source:MGI Symbol;Acc:MGI:2448407] |
| ENSMUSG00000062727 | H2bc12 | 2.275062 | 1391.664 | 0.195422 | -11.6418 | 2.53E-31 | 6.28E-29 | 7.61E-29 | 2603.127 | 2011.363 | 563.3036 | 388.8627 | 319184 | H2B clustered histone 12 [Source:MGI Symbol;Acc:MGI:2448399] |
| ENSMUSG00000061615 | H2ac4 | 2.280667 | 1074.659 | 0.20142 | -11.3229 | 1.01E-29 | 2.04E-27 | 2.47E-27 | 2118.677 | 1446.805 | 397.7874 | 335.3684 | 319172 | H2A clustered histone 4 [Source:MGI Symbol;Acc:MGI:2448306] |
| ENSMUSG00000051220 | Ercc6l | 2.285013 | 332.6524 | 0.263215 | -8.68117 | 3.92E-18 | 2.28E-16 | 2.76E-16 | 643.1132 | 461.2857 | 124.3657 | 101.845 | 236930 | excision repair cross-complementing rodent repair deficiency complementation group 6 like [Source:MGI Symbol;Acc:MGI:2654144] |
| ENSMUSG00000078521 | Aunip | 2.287191 | 29.92087 | 0.699656 | -3.26902 | 0.001079 | 0.006698 | 0.008114 | 56.06085 | 43.27627 | 10.05899 | 10.28737 | 69885 | aurora kinase A and ninein interacting protein [Source:MGI Symbol;Acc:MGI:1917135] |
| ENSMUSG00000018983 | E2f2 | 2.287689 | 146.1549 | 0.359724 | -6.35957 | 2.02E-10 | 5E-09 | 6.06E-09 | 287.7085 | 197.6939 | 53.95278 | 45.26444 | 242705 | E2F transcription factor 2 [Source:MGI Symbol;Acc:MGI:1096341] |
| ENSMUSG00000017146 | Brca1 | 2.29335 | 493.2216 | 0.221647 | -10.3469 | 4.32E-25 | 5.6E-23 | 6.79E-23 | 943.5147 | 695.371 | 175.5751 | 158.4256 | 12189 | breast cancer 1, early onset [Source:MGI Symbol;Acc:MGI:104537] |
| ENSMUSG00000029591 | Ung | 2.299803 | 118.3013 | 0.378802 | -6.07125 | 1.27E-09 | 2.82E-08 | 3.41E-08 | 221.0701 | 172.1215 | 34.74925 | 45.26444 | 22256 | uracil DNA glycosylase [Source:MGI Symbol;Acc:MGI:109352] |
| ENSMUSG00000069308 | H2bc22 | 2.301323 | 60.65366 | 0.56992 | -4.03797 | 5.39E-05 | 0.000485 | 0.000587 | 116.3527 | 85.569 | 28.34807 | 12.34485 | 319188 | H2B clustered histone 22 [Source:MGI Symbol;Acc:MGI:2448409] |
| ENSMUSG00000002068 | Ccne1 | 2.308213 | 156.4262 | 0.361263 | -6.38928 | 1.67E-10 | 4.18E-09 | 5.07E-09 | 315.2101 | 205.5623 | 57.61059 | 47.32192 | 12447 | cyclin E1 [Source:MGI Symbol;Acc:MGI:88316] |
| ENSMUSG00000034853 | Acot11 | 2.318892 | 66.95296 | 0.527609 | -4.3951 | 1.11E-05 | 0.000118 | 0.000143 | 80.38915 | 142.615 | 20.11798 | 24.6897 | 329910 | acyl-CoA thioesterase 11 [Source:MGI Symbol;Acc:MGI:1913736] |
| ENSMUSG00000050936 | Gm42743 | 2.319387 | 263.5186 | 0.30345 | -7.64338 | 2.12E-14 | 8.47E-13 | 1.03E-12 | 521.4717 | 357.0293 | 101.5044 | 74.06909 | | predicted gene 42743 [Source:MGI Symbol;Acc:MGI:5662880] |
| ENSMUSG00000020014 | Cfap54 | 2.334159 | 69.74186 | 0.759297 | -3.0741 | 0.002111 | 0.011646 | 0.014107 | 35.96357 | 196.7103 | 16.46017 | 29.83338 | 380654 | cilia and flagella associated protein 54 [Source:MGI Symbol;Acc:MGI:1922208] |
| ENSMUSG00000075031 | H2bc3 | 2.338036 | 1506.751 | 0.175659 | -13.3101 | 2.02E-40 | 1.18E-37 | 1.43E-37 | 2898.24 | 2134.307 | 545.9289 | 448.5295 | 319178 | H2B clustered histone 3 [Source:MGI Symbol;Acc:MGI:2448377] |
| ENSMUSG00000020788 | Atp2a3 | 2.33883 | 131.9152 | 0.387315 | -6.03857 | 1.55E-09 | 3.4E-08 | 4.12E-08 | 242.2252 | 198.6774 | 54.86723 | 31.89086 | 53313 | ATPase, Ca++ transporting, ubiquitous [Source:MGI Symbol;Acc:MGI:1194503] |
| ENSMUSG00000094338 | H2bc13 | 2.346433 | 216.5135 | 0.305206 | -7.68804 | 1.49E-14 | 6.09E-13 | 7.38E-13 | 409.35 | 314.7365 | 82.30084 | 59.66677 | 319185 | H2B clustered histone 13 [Source:MGI Symbol;Acc:MGI:2448403] |
| ENSMUSG00000035455 | Fignl1 | 2.346562 | 484.1633 | 0.210548 | -11.145 | 7.57E-29 | 1.35E-26 | 1.64E-26 | 885.3384 | 732.746 | 142.6548 | 175.9141 | 60530 | fidgetin-like 1 [Source:MGI Symbol;Acc:MGI:1890648] |
| ENSMUSG00000023505 | Cdca3 | 2.347013 | 390.2097 | 0.246111 | -9.53641 | 1.48E-21 | 1.27E-19 | 1.53E-19 | 712.9248 | 592.0981 | 150.8849 | 104.9312 | 14793 | cell division cycle associated 3 [Source:MGI Symbol;Acc:MGI:1315198] |
| ENSMUSG00000120697 | Gm36496 | 2.357526 | 39.43773 | 0.659836 | -3.5729 | 0.000353 | 0.002556 | 0.003096 | 75.10039 | 57.046 | 17.37462 | 8.229899 | | predicted gene, 36496 [Source:MGI Symbol;Acc:MGI:5595655] |
| ENSMUSG00000005410 | Mcm5 | 2.360449 | 1266.794 | 0.167935 | -14.0557 | 7.11E-45 | 7.7E-42 | 9.32E-42 | 2430.714 | 1810.719 | 414.2476 | 411.4949 | 17218 | minichromosome maintenance complex component 5 [Source:MGI Symbol;Acc:MGI:103197] |
| ENSMUSG00000069267 | H3c2 | 2.362589 | 1295.944 | 0.16237 | -14.5507 | 5.78E-48 | 9.74E-45 | 1.18E-44 | 2348.209 | 1992.676 | 474.6015 | 368.288 | 319150 | H3 clustered histone 2 [Source:MGI Symbol;Acc:MGI:2448319] |
| ENSMUSG00000031004 | Mki67 | 2.364303 | 5884.867 | 0.35603 | -6.64074 | 3.12E-11 | 8.64E-10 | 1.05E-09 | 11738.93 | 7972.67 | 2214.807 | 1613.06 | 17345 | antigen identified by monoclonal antibody Ki 67 [Source:MGI Symbol;Acc:MGI:106035] |
| ENSMUSG00000034329 | Brip1 | 2.378078 | 389.7886 | 0.266461 | -8.92468 | 4.47E-19 | 2.96E-17 | 3.59E-17 | 757.3504 | 550.7889 | 148.1415 | 102.8737 | 237911 | BRCA1 interacting protein C-terminal helicase 1 [Source:MGI Symbol;Acc:MGI:2442836] |
| ENSMUSG00000047361 | Gm973 | 2.382542 | 41.52927 | 0.885973 | -2.68918 | 0.007163 | 0.03234 | 0.039072 | 22.21279 | 117.0427 | 7.315631 | 19.54601 | 381260 | predicted gene 973 [Source:MGI Symbol;Acc:MGI:2685819] |
| ENSMUSG00000032815 | Fanca | 2.39633 | 212.1731 | 0.309314 | -7.74723 | 9.39E-15 | 3.93E-13 | 4.77E-13 | 399.8302 | 313.753 | 79.55748 | 55.55182 | 14087 | Fanconi anemia, complementation group A [Source:MGI Symbol;Acc:MGI:1341823] |
| ENSMUSG00000007080 | Pole | 2.403691 | 507.3302 | 0.268878 | -8.9397 | 3.9E-19 | 2.61E-17 | 3.16E-17 | 995.3446 | 712.0914 | 198.4365 | 123.4485 | 18973 | polymerase (DNA directed), epsilon [Source:MGI Symbol;Acc:MGI:1196391] |
| ENSMUSG00000069300 | H2bc11 | 2.403778 | 971.6565 | 0.200919 | -11.9639 | 5.49E-33 | 1.67E-30 | 2.02E-30 | 1856.355 | 1413.364 | 356.637 | 260.2706 | 319183 | H2B clustered histone 11 [Source:MGI Symbol;Acc:MGI:2448388] |
| ENSMUSG00000022033 | Pbk | 2.405248 | 514.2788 | 0.257808 | -9.32962 | 1.06E-20 | 8.62E-19 | 1.04E-18 | 996.4023 | 734.7131 | 198.4365 | 127.5634 | 52033 | PDZ binding kinase [Source:MGI Symbol;Acc:MGI:1289156] |
| ENSMUSG00000056498 | Tmem154 | 2.407469 | 68.86927 | 0.54513 | -4.41632 | 1E-05 | 0.000109 | 0.000131 | 111.064 | 120.9769 | 31.09143 | 12.34485 | 320782 | transmembrane protein 154 [Source:MGI Symbol;Acc:MGI:2444725] |
| ENSMUSG00000058773 | H1f5 | 2.42154 | 1252.785 | 0.183451 | -13.1999 | 8.78E-40 | 4.44E-37 | 5.38E-37 | 2375.711 | 1848.094 | 449.9113 | 337.4259 | 56702 | H1.5 linker histone, cluster member [Source:MGI Symbol;Acc:MGI:1861461] |
| ENSMUSG00000026669 | Mcm10 | 2.447767 | 305.1002 | 0.24412 | -10.0269 | 1.16E-23 | 1.28E-21 | 1.55E-21 | 569.0705 | 462.2693 | 92.35984 | 96.70131 | 70024 | minichromosome maintenance 10 replication initiation factor [Source:MGI Symbol;Acc:MGI:1917274] |
| ENSMUSG00000026185 | Igfbp5 | 2.448257 | 271.5148 | 0.400089 | -6.11928 | 9.4E-10 | 2.11E-08 | 2.56E-08 | 305.6903 | 611.7692 | 58.52504 | 110.0749 | 16011 | insulin-like growth factor binding protein 5 [Source:MGI Symbol;Acc:MGI:96440] |
| ENSMUSG00000025473 | Adam8 | 2.450064 | 3092.847 | 0.290435 | -8.43585 | 3.29E-17 | 1.77E-15 | 2.15E-15 | 6289.393 | 4168.292 | 961.091 | 952.6108 | 11501 | a disintegrin and metallopeptidase domain 8 [Source:MGI Symbol;Acc:MGI:107825] |
| ENSMUSG00000058624 | Gda | 2.452151 | 77.06162 | 0.468257 | -5.23676 | 1.63E-07 | 2.55E-06 | 3.08E-06 | 153.374 | 107.2071 | 21.94689 | 25.71843 | 14544 | guanine deaminase [Source:MGI Symbol;Acc:MGI:95678] |
| ENSMUSG00000069309 | H2ac22 | 2.45253 | 144.3222 | 0.353984 | -6.92836 | 4.26E-12 | 1.34E-10 | 1.63E-10 | 284.5353 | 203.5952 | 43.89378 | 45.26444 | 319170 | H2A clustered histone 22 [Source:MGI Symbol;Acc:MGI:2448300] |
| ENSMUSG00000028587 | Orc1 | 2.453883 | 65.66263 | 0.490571 | -5.00209 | 5.67E-07 | 7.85E-06 | 9.51E-06 | 124.8147 | 97.37162 | 21.94689 | 18.51727 | 18392 | origin recognition complex, subunit 1 [Source:MGI Symbol;Acc:MGI:1328337] |
| ENSMUSG00000114456 | H2bc9 | 2.461204 | 621.4956 | 0.208549 | -11.8015 | 3.83E-32 | 1.1E-29 | 1.33E-29 | 1174.105 | 930.4399 | 215.8111 | 165.6267 | 319182 | H2B clustered histone 9 [Source:MGI Symbol;Acc:MGI:2448387] |
| ENSMUSG00000046295 | Ankle1 | 2.463876 | 65.91704 | 0.496754 | -4.95995 | 7.05E-07 | 9.6E-06 | 1.16E-05 | 126.9302 | 96.38807 | 22.86135 | 17.48853 | 234396 | ankyrin repeat and LEM domain containing 1 [Source:MGI Symbol;Acc:MGI:1918775] |
| ENSMUSG00000029414 | Kntc1 | 2.465308 | 597.2589 | 0.236 | -10.4462 | 1.52E-25 | 2.12E-23 | 2.57E-23 | 1140.257 | 883.2294 | 219.4689 | 146.0807 | 208628 | kinetochore associated 1 [Source:MGI Symbol;Acc:MGI:2673709] |
| ENSMUSG00000069310 | H3c3 | 2.472972 | 725.8336 | 0.164349 | -15.0471 | 3.61E-51 | 1.37E-47 | 1.66E-47 | 1311.612 | 1148.788 | 225.8701 | 217.0636 | 319148 | H3 clustered histone 3 [Source:MGI Symbol;Acc:MGI:2448320] |
| ENSMUSG00000025574 | Tk1 | 2.477829 | 457.7865 | 0.238082 | -10.4075 | 2.29E-25 | 3.12E-23 | 3.78E-23 | 869.4721 | 683.5684 | 161.8583 | 116.2473 | 21877 | thymidine kinase 1 [Source:MGI Symbol;Acc:MGI:98763] |
| ENSMUSG00000017861 | Mybl2 | 2.48566 | 106.8915 | 0.399824 | -6.21689 | 5.07E-10 | 1.19E-08 | 1.44E-08 | 209.4349 | 153.4341 | 33.83479 | 30.86212 | 17865 | myeloblastosis oncogene-like 2 [Source:MGI Symbol;Acc:MGI:101785] |
| ENSMUSG00000027379 | Bub1 | 2.487009 | 700.8102 | 0.198853 | -12.5068 | 6.85E-36 | 2.74E-33 | 3.31E-33 | 1348.634 | 1030.762 | 230.4424 | 193.4026 | 12235 | BUB1, mitotic checkpoint serine/threonine kinase [Source:MGI Symbol;Acc:MGI:1100510] |
| ENSMUSG00000082016 | Pgam1-ps2 | 2.493415 | 18.91362 | 0.924744 | -2.69633 | 0.007011 | 0.03179 | 0.038406 | 33.84806 | 30.4901 | 8.230084 | 3.086212 | | phosphoglycerate mutase 1, pseudogene 2 [Source:MGI Symbol;Acc:MGI:3645709] |
| ENSMUSG00000000028 | Cdc45 | 2.494637 | 209.5536 | 0.30494 | -8.18075 | 2.82E-16 | 1.37E-14 | 1.66E-14 | 407.2345 | 304.901 | 69.49849 | 56.58055 | 12544 | cell division cycle 45 [Source:MGI Symbol;Acc:MGI:1338073] |
| ENSMUSG00000020649 | Rrm2 | 2.495627 | 468.8302 | 0.266571 | -9.36195 | 7.83E-21 | 6.45E-19 | 7.81E-19 | 979.4783 | 613.7363 | 154.5427 | 127.5634 | 20135 | ribonucleotide reductase M2 [Source:MGI Symbol;Acc:MGI:98181] |
| ENSMUSG00000022034 | Esco2 | 2.49716 | 267.6064 | 0.303577 | -8.2258 | 1.94E-16 | 9.48E-15 | 1.15E-14 | 519.3562 | 390.47 | 97.84656 | 62.75298 | 71988 | establishment of sister chromatid cohesion N-acetyltransferase 2 [Source:MGI Symbol;Acc:MGI:1919238] |
| ENSMUSG00000040204 | Pclaf | 2.50688 | 881.3405 | 0.186887 | -13.4139 | 5.02E-41 | 3.17E-38 | 3.84E-38 | 1716.731 | 1281.568 | 277.0795 | 249.9832 | 68026 | PCNA clamp associated factor [Source:MGI Symbol;Acc:MGI:1915276] |
| ENSMUSG00000051457 | Spn | 2.535387 | 22.95464 | 0.803338 | -3.15607 | 0.001599 | 0.00931 | 0.011277 | 37.02132 | 41.30917 | 7.315631 | 6.172424 | 20737 | sialophorin [Source:MGI Symbol;Acc:MGI:98384] |
| ENSMUSG00000069301 | H2ac11 | 2.548695 | 470.6704 | 0.248997 | -10.2358 | 1.37E-24 | 1.64E-22 | 1.98E-22 | 900.1469 | 707.1737 | 109.7345 | 165.6267 | 319167 | H2A clustered histone 11 [Source:MGI Symbol;Acc:MGI:2448293] |
| ENSMUSG00000028212 | Ccne2 | 2.560148 | 260.0493 | 0.289865 | -8.83221 | 1.03E-18 | 6.43E-17 | 7.79E-17 | 524.6449 | 364.8977 | 78.64303 | 72.01161 | 12448 | cyclin E2 [Source:MGI Symbol;Acc:MGI:1329034] |
| ENSMUSG00000085295 | 4930430E12Rik | 2.56935 | 450.9124 | 0.26897 | -9.55254 | 1.27E-21 | 1.11E-19 | 1.34E-19 | 910.7244 | 632.4237 | 105.1622 | 155.3393 | | RIKEN cDNA 4930430E12 gene [Source:MGI Symbol;Acc:MGI:1918889] |
| ENSMUSG00000023908 | Pkmyt1 | 2.575169 | 68.61525 | 0.481344 | -5.34996 | 8.8E-08 | 1.46E-06 | 1.76E-06 | 113.1795 | 121.9604 | 22.86135 | 16.4598 | 268930 | protein kinase, membrane associated tyrosine/threonine 1 [Source:MGI Symbol;Acc:MGI:2137630] |
| ENSMUSG00000046179 | E2f8 | 2.598487 | 100.2749 | 0.445895 | -5.82757 | 5.62E-09 | 1.13E-07 | 1.37E-07 | 212.6081 | 131.7959 | 32.00588 | 24.6897 | 108961 | E2F transcription factor 8 [Source:MGI Symbol;Acc:MGI:1922038] |
| ENSMUSG00000061991 | H2ac10 | 2.602202 | 128.973 | 0.376068 | -6.91949 | 4.53E-12 | 1.43E-10 | 1.73E-10 | 255.976 | 186.8748 | 32.92034 | 40.12076 | 319173 | H2A clustered histone 10 [Source:MGI Symbol;Acc:MGI:2448309] |
| ENSMUSG00000036223 | Ska1 | 2.604402 | 71.62812 | 0.470788 | -5.532 | 3.17E-08 | 5.73E-07 | 6.94E-07 | 130.1035 | 116.0591 | 22.86135 | 17.48853 | 66468 | spindle and kinetochore associated complex subunit 1 [Source:MGI Symbol;Acc:MGI:1913718] |
| ENSMUSG00000022422 | Dscc1 | 2.605441 | 57.46269 | 0.57118 | -4.56151 | 5.08E-06 | 5.8E-05 | 7.03E-05 | 111.064 | 86.55255 | 21.94689 | 10.28737 | 72107 | DNA replication and sister chromatid cohesion 1 [Source:MGI Symbol;Acc:MGI:1919357] |
| ENSMUSG00000017499 | Cdc6 | 2.618061 | 163.1827 | 0.333925 | -7.84028 | 4.5E-15 | 1.97E-13 | 2.39E-13 | 316.2678 | 244.9044 | 41.15042 | 50.40813 | 23834 | cell division cycle 6 [Source:MGI Symbol;Acc:MGI:1345150] |
| ENSMUSG00000033715 | Akr1c14 | 2.625993 | 31.59737 | 0.774459 | -3.39074 | 0.000697 | 0.004605 | 0.005579 | 34.90581 | 73.76638 | 6.401177 | 11.31611 | 105387 | aldo-keto reductase family 1, member C14 [Source:MGI Symbol;Acc:MGI:2145458] |
| ENSMUSG00000114279 | H2bc14 | 2.635285 | 889.1293 | 0.17951 | -14.6804 | 8.61E-49 | 1.68E-45 | 2.04E-45 | 1713.558 | 1350.416 | 265.1916 | 227.351 | 319186 | H2B clustered histone 14 [Source:MGI Symbol;Acc:MGI:2448404] |
| ENSMUSG00000027323 | Rad51 | 2.65798 | 118.257 | 0.398709 | -6.66646 | 2.62E-11 | 7.33E-10 | 8.88E-10 | 242.2252 | 166.2202 | 34.74925 | 29.83338 | 19361 | RAD51 recombinase [Source:MGI Symbol;Acc:MGI:97890] |
| ENSMUSG00000027015 | Cybrd1 | 2.65904 | 25.14678 | 0.965157 | -2.75503 | 0.005869 | 0.027397 | 0.033096 | 16.92403 | 69.83217 | 4.572269 | 9.258636 | 73649 | cytochrome b reductase 1 [Source:MGI Symbol;Acc:MGI:2654575] |
| ENSMUSG00000099517 | H3c8 | 2.675932 | 516.5491 | 0.252705 | -10.5891 | 3.35E-26 | 4.97E-24 | 6.03E-24 | 1052.463 | 734.7131 | 162.7728 | 116.2473 | 97908 | H3 clustered histone 8 [Source:MGI Symbol;Acc:MGI:2145541] |
| ENSMUSG00000022371 | Col14a1 | 2.689264 | 878.6483 | 0.241581 | -11.1319 | 8.77E-29 | 1.53E-26 | 1.85E-26 | 1828.853 | 1214.686 | 280.7373 | 190.3164 | 12818 | collagen, type XIV, alpha 1 [Source:MGI Symbol;Acc:MGI:1341272] |
| ENSMUSG00000029648 | Flt1 | 2.718019 | 114.4425 | 0.419392 | -6.48085 | 9.12E-11 | 2.38E-09 | 2.88E-09 | 228.4744 | 169.1709 | 37.49261 | 22.63222 | 14254 | FMS-like tyrosine kinase 1 [Source:MGI Symbol;Acc:MGI:95558] |
| ENSMUSG00000030346 | Rad51ap1 | 2.72971 | 257.5102 | 0.285664 | -9.55568 | 1.23E-21 | 1.09E-19 | 1.32E-19 | 509.8364 | 385.5523 | 74.98521 | 59.66677 | 19362 | RAD51 associated protein 1 [Source:MGI Symbol;Acc:MGI:1098224] |
| ENSMUSG00000117390 | Gm50080 | 2.750827 | 20.43005 | 0.920102 | -2.9897 | 0.002793 | 0.014697 | 0.017798 | 47.59884 | 23.60524 | 6.401177 | 4.114949 | | predicted gene, 50080 [Source:MGI Symbol;Acc:MGI:6275410] |
| ENSMUSG00000039396 | Neil3 | 2.761911 | 138.237 | 0.366606 | -7.53374 | 4.93E-14 | 1.91E-12 | 2.32E-12 | 255.976 | 226.2169 | 42.97933 | 27.77591 | 234258 | nei like 3 (E. coli) [Source:MGI Symbol;Acc:MGI:2384588] |
| ENSMUSG00000001228 | Uhrf1 | 2.771228 | 924.6191 | 0.188798 | -14.6782 | 8.89E-49 | 1.68E-45 | 2.04E-45 | 1864.817 | 1361.236 | 236.8435 | 235.5809 | 18140 | ubiquitin-like, containing PHD and RING finger domains, 1 [Source:MGI Symbol;Acc:MGI:1338889] |
| ENSMUSG00000120514 | Gm56644 | 2.77682 | 23.03157 | 0.863504 | -3.21576 | 0.001301 | 0.007825 | 0.009479 | 51.82984 | 28.523 | 4.572269 | 7.201161 | | predicted gene, 56644 [Source:MGI Symbol;Acc:MGI:7337202] |
| ENSMUSG00000040751 | Lat2 | 2.861686 | 760.756 | 0.214495 | -13.3415 | 1.33E-40 | 8.05E-38 | 9.75E-38 | 1560.184 | 1115.348 | 203.9232 | 163.5692 | 56743 | linker for activation of T cells family, member 2 [Source:MGI Symbol;Acc:MGI:1926479] |
| ENSMUSG00000034773 | Hrob | 2.973833 | 25.29723 | 0.89453 | -3.32446 | 0.000886 | 0.005665 | 0.006863 | 61.34961 | 28.523 | 8.230084 | 3.086212 | 217216 | homologous recombination factor with OB-fold [Source:MGI Symbol;Acc:MGI:2387601] |
| ENSMUSG00000004709 | Cd244a | 2.979736 | 89.21736 | 0.491905 | -6.05755 | 1.38E-09 | 3.05E-08 | 3.7E-08 | 186.1643 | 130.8124 | 26.51916 | 13.37359 | 18106 | CD244 molecule A [Source:MGI Symbol;Acc:MGI:109294] |
| ENSMUSG00000016494 | Cd34 | 2.997787 | 259.2478 | 0.299577 | -10.0067 | 1.42E-23 | 1.54E-21 | 1.87E-21 | 524.6449 | 397.3549 | 67.66958 | 47.32192 | 12490 | CD34 antigen [Source:MGI Symbol;Acc:MGI:88329] |
| ENSMUSG00000081058 | H3c15 | 3.036603 | 589.1427 | 0.216554 | -14.0224 | 1.14E-44 | 1.01E-41 | 1.23E-41 | 1205.837 | 895.032 | 135.3392 | 120.3623 | 97114 | H3 clustered histone 15 [Source:MGI Symbol;Acc:MGI:2448357] |
| ENSMUSG00000069302 | H2ac12 | 3.052532 | 47.43511 | 0.590145 | -5.17251 | 2.31E-07 | 3.48E-06 | 4.22E-06 | 85.6779 | 83.60189 | 9.144538 | 11.31611 | 319168 | H2A clustered histone 12 [Source:MGI Symbol;Acc:MGI:2448295] |
| ENSMUSG00000026548 | Slamf9 | 3.076918 | 288.297 | 0.313397 | -9.81796 | 9.42E-23 | 9.53E-21 | 1.15E-20 | 625.1314 | 406.2068 | 70.41294 | 51.43687 | 98365 | SLAM family member 9 [Source:MGI Symbol;Acc:MGI:1923692] |
| ENSMUSG00000026068 | Il18rap | 3.134503 | 37.80376 | 0.66394 | -4.72106 | 2.35E-06 | 2.85E-05 | 3.46E-05 | 70.86938 | 64.91441 | 8.230084 | 7.201161 | 16174 | interleukin 18 receptor accessory protein [Source:MGI Symbol;Acc:MGI:1338888] |
| ENSMUSG00000045655 | Fam216b | 3.567588 | 25.47099 | 1.233331 | -2.89264 | 0.00382 | 0.019238 | 0.023265 | 6.346511 | 87.5361 | 1.828904 | 6.172424 | 219170 | family with sequence similarity 216, member B [Source:MGI Symbol;Acc:MGI:2145738] |
| ENSMUSG00000084989 | Crocc2 | 3.678189 | 17.10548 | 1.342071 | -2.74068 | 0.006131 | 0.028396 | 0.034303 | 6.346511 | 57.046 | 0.914454 | 4.114949 | 381284 | ciliary rootlet coiled-coil, rootletin family member 2 [Source:MGI Symbol;Acc:MGI:3045962] |
| ENSMUSG00000071335 | Mfsd4b3-ps | 5.023032 | 8.213611 | 1.89073 | -2.65666 | 0.007892 | | 0.042373 | 19.03953 | 12.78617 | 0 | 1.028737 | | major facilitator superfamily domain containing 4B3, pseudogene [Source:MGI Symbol;Acc:MGI:3773841] |
| ENSMUSG00000094708 | Gm10359 | 5.542008 | 11.61894 | 1.780179 | -3.11317 | 0.001851 | 0.010464 | 0.012676 | 16.92403 | 28.523 | 0 | 1.028737 | | predicted gene 10359 [Source:MGI Symbol;Acc:MGI:3708724] |
| ENSMUSG00000095937 | Gm12671 | 5.542008 | 11.61894 | 1.780179 | -3.11317 | 0.001851 | 0.010464 | 0.012676 | 16.92403 | 28.523 | 0 | 1.028737 | | predicted gene 12671 [Source:MGI Symbol;Acc:MGI:3651684] |
| ENSMUSG00000138098 | Gm71269 | 6.173554 | 6.443999 | 2.093373 | -2.94909 | 0.003187 | | 0.019939 | 16.92403 | 8.851965 | 0 | 0 | | predicted gene, 71269 [Source:MGI Symbol;Acc:MGI:7825319] |
| ENSMUSG00000089942 | Pira2 | 6.239641 | 6.731724 | 2.144203 | -2.91 | 0.003614 | | 0.022129 | 5.28876 | 21.63814 | 0 | 0 | 18725 | paired-Ig-like receptor A2 [Source:MGI Symbol;Acc:MGI:1195970] |
| ENSMUSG00000084071 | Gm13378 | 6.465773 | 7.896039 | 2.232318 | -2.89644 | 0.003774 | | 0.023016 | 29.61705 | 1.967103 | 0 | 0 | | predicted gene 13378 [Source:MGI Symbol;Acc:MGI:3650302] |

### Supplementary Table S2

### Down genes (after LPS administration)

#### I. Major biological programs

| **A. Microglia state genes** | **83** | | | | | | | | | | | |
|---|---|---|---|---|---|---|---|---|---|---|---|---|
| A1. microglia identity and DAM identity genes | 6 | | | | | | | | | | | |
| A1a Microglial identity | | 2 | Spic | Adgre1 | | | | | | | | |
| A1b DAM identity genes | | 4 | Apoe | Tyrobp | Trem2 | Axl | | | | | | |
| A2. Innate immune receptors | 19 | | | | | | | | | | | |
| A2a. Patterm-recognition receprors (PRPs) | | 4 | Cd4 | Nlrc4 | Ifih1 | Tlr8 | | | | | | |
| A2b. C-type lectin receptors | | 8 | Cd209f | Cd209g | Clec10a | Clec4a1 | Clec4a2 | Clec4a3 | Clec4n | Clec2d | | |
| A2c. Myeloid immunoregulatory receptors | | 5 | Adgre4 | Fcgr2b | Fcgr3 | Fcgrt | Siglec1 | | | | | |
| A2d. CD300 family receptors | | 2 | Cd300a | Cd300c2 | | | | | | | | |
| A2e. SLAM family receptors | | 0 | | | | | | | | | | |
| A2f. Other immune receptors | | 0 | | | | | | | | | | |
| A3. Cytokine / inflammatory signaling | 18 | | | | | | | | | | | |
| A3a. Cytokines | | 5 | Il1a | Il1b | Il6 | Il10 | Il16 | | | | | |
| A3b. Cytokine receptor genes | | 7 | Il10ra | Il2rg | Il6ra | Tnfrsf1b | Tnfrsf19 | Ifngr1 | Ifnar2 | | | |
| A3c. Chemokines | | 6 | Ccl3 | Ccl5 | Ccl6 | Ccl7 | Ccl8 | Ccl24 | | | | |
| A4. Antigen processing / phagocytosis | 38 | | | | | | | | | | | |
| A4a. Antigen processing | | 2 | Ifi30 | Tapbp | | | | | | | | |
| A4b. Antigen uptake / lysosomal | | 5 | Cd68 | Hexb | Lamp1 | Lyz1 | Lyz2 | | | | | |
| A4c. Lysosomal proteases | | 5 | Ctsb | Ctsc | Ctsh | Ctsf | Ctss | | | | | |
| A4d. Interferon-stimulated antiviral effectors | | 13 | Ifit1 | Ifit2 | Ifit3 | Ifit3b | Ifitm1 | Ifitm2 | Ifitm3 | Oas2 | Oas3 | Rsad2 |
| | | | Samhd1 | Bst2 | Rtp4 | | | | | | | |
| A4e. Cytosolic nucleic-acid sensing | | 1 | Zbp1 | | | | | | | | | |
| A4f. IFN-inducible GTPases | | 12 | Gbp2 | Gbp3 | Gbp5 | Gbp6 | Gbp7 | Gbp8 | Gbp9 | Iigp1 | Iigp1c | Irgm1 |
| | | | Irgm2 | Tgtp2 | | | | | | | | |
| A5. Innate immune transcriptional regulators | 2 | | | | | | | | | | | |
| A5a. Core transcription factors | | 1 | Spic | | | | | | | | | |
| A5b. Immune-response regulators | | 1 | Nfil3 | | | | | | | | | |

| **B. Cell-state regulators** | **46** | | | | | | | | | | | |
|---|---|---|---|---|---|---|---|---|---|---|---|---|
| B1. Transcriptional / Cell-state regulators | 28 | | | | | | | | | | | |
| | | | Irf1 | Irf4 | Irf7 | Irf8 | Stat1 | Stat2 | Rel | Fos | Fosb | Fosl2 |
| B1a. Core TFs | | 24 | Jun | Junb | Jund | Egr2 | Egr3 | Nr4a1 | Nr4a2 | Klf3 | Klf9 | Nfil3 |
| | | | Nfe2l3 | Prdm1 | Prdm6 | Zbtb46 | | | | | | |
| B1b. Transcriptional regulators | | 4 | Irf2bp2 | Zfp36 | Zfp36l1 | Zfp36l2 | | | | | | |
| B2. Signal-adaptation regulators | 18 | | | | | | | | | | | |
| B2a. MAPK / stress-response modulators | | 11 | Dusp1 | Dusp2 | Dusp7 | Dusp15 | Map3k8 | Mapk14 | Ier2 | Ier3 | Ier5 | Sik1 |
| | | | Plk2 | | | | | | | | | |
| B2b. PI3K–Akt / survival regulators | | 0 | | | | | | | | | | |
| B2c. Immune signaling adaptors | | 5 | Cish | Socs6 | Irf2bp2 | Nfkbid | Nfkbiz | | | | | |
| B2d. Cell-state stabilization regulators | | 2 | Btg1 | Btg2 | | | | | | | | |
| B3. Epigenetic / chromatin state regulators | 0 | | | | | | | | | | | |
| B3a. Histone methylation regulators | | 0 | | | | | | | | | | |
| B3b. Polycomb-associated regulators | | 0 | | | | | | | | | | |
| B3c. Chromatin remodeling factors | | 0 | | | | | | | | | | |
| B3d. DNA methylation / replication-coupled chromatin regulators | | 0 | | | | | | | | | | |
| B3e. Chromosome architecture regulators | | 0 | | | | | | | | | | |

| **C. Myelin / axon–glia interaction genes** | **30** | | | | | | | | | | | |
|---|---|---|---|---|---|---|---|---|---|---|---|---|
| C1. Myelin-associated genes | 12 | | Mbp | Mog | Mag | Mobp | Plp1 | Opalin | Ugt8a | Mal | Tspan2 | Ermn |
| | | | Mpzl2 | Cldn11 | | | | | | | | |
| C2. Axon guidance / neurite interaction | 2 | | Sema4a | Sema4d | | | | | | | | |
| C3. Neuron–glia adhesion molecules | 1 | | Ncam1 | | | | | | | | | |
| C4. Neurotrophic / developmental signaling | 7 | | **Bmp2** | Olig1 | Olig2 | Sox8 | Myrf | S1pr5 | Gpr37 | | | |
| C5. Synaptic / neuron–glia communication | 6 | | Adora2b | Ptgds | Gjb1 | Gjb2 | Gjb6 | Gjc2 | | | | |
| C6. Glial structural support / neural architecture | 1 | | Vcan | | | | | | | | | |
| C7. Neurovascular / neural niche interaction | 1 | | Angptl4 | | | | | | | | | |

| **D. Neuron–glia communication genes** | **30** | | | | | | | | | |
|---|---|---|---|---|---|---|---|---|---|---|
| D1. Growth factor / trophic signaling | 1 | | **Bmp2** | | | | | | | |
| D2. Guidance / axon interaction | 4 | | Sema4a | Sema4d | Dscam | Ptprd | | | | |
| D3. Neuroactive receptors | 5 | | **Adora2b** | Gria2 | Gria3 | Grik2 | Grik3 | | | |
| D4. Cell-cell communication | 4 | | Ncam1 | Nrxn2 | Nrxn3 | Lrrtm3 | | | | |
| D5. Cytoskeletal/synapse-contact | 0 | | | | | | | | | |
| D6. Vesicle trafficking / phagosome–synapse interface | 0 | | | | | | | | | |
| D7. Neuron-responsive TFs | 8 | | Fos | Fosb | Jun | Junb | Egr2 | Egr3 | Nr4a1 | Nr4a2 |
| D8. Neuroimmune communication | 8 | | Il1a | Il1b | Il6 | Cxcl10 | Kcna1 | Kcnq2 | Cacna1b | Cacna1c |

| **E. Astrocyte-like / glial-environment genes** | **26** | | | | | | | | | | | |
|---|---|---|---|---|---|---|---|---|---|---|---|---|
| E1. Astrocyte identity/glial differentiation | 4 | | **Gja1** | **Ptprz1** | Tnr | | | | | | | |
| E2. Metabolic support | 1 | | Angptl4 | | | | | | | | | |
| E3. Lipid/cholesterol support | 0 | | | | | | | | | | | |
| E4. Astrocyte ECM / neurovascular niche | 13 | | Vcan | Epyc | Pi15 | Col1a1 | Col11a1 | Col12a1 | Col8a1 | Thbs2 | Tnr | Ptprz1 |
| | | | Ddr1 | Gja1 | Ccn3 | | | | | | | |
| E5. Reactive astrocyte / glial-environment response | 8 | | Bmp2 | **Ccn3** | Thbs2 | Adamts5 | Adamts12 | Mmp16 | Ddr1 | Pi15 | | |

| **F. ECM-related genes** | **37** | | | | | | | | | | |
|---|---|---|---|---|---|---|---|---|---|---|---|
| F1. ECM structural components | 9 | | **Vcan** | Fbln5 | Col4a5 | **Col1a1** | **Col11a1** | **Col12a1** | **Col8a1** | **Tnr** | **Epyc** |
| F2. ECM remodeling enzymes | 8 | | **Adamts4** | **Adamts5** | **Adamts14** | **Adamts12** | **Mmp13** | **Mmp14** | **Mmp16** | **Prss35** | |
| F3. Matrisome-associated secreted proteins | 6 | | Spp2 | **Angptl4** | **Pi15** | Ccn3 | **Thbs2** | | | | |
| F4. ECM glycan / proteoglycan regulators | 2 | | Chst2 | Chst11 | | | | | | | |
| F5. ECM adhesion receptors | 2 | | Itga11 | **Ddr1** | | | | | | | |
| F6. Adhesion complex / focal adhesion | 1 | | Parvb | | | | | | | | |

| | | | | | | |
|---|---|---|---|---|---|---|
| F7. Cell–matrix interaction receptors | 2 | Cd44 | Ptprz1 | | | |
| F8. Migration / invasion regulators | 0 | | | | | |
| F9. Cytoskeletal ECM-response machinery | 0 | | | | | |
| F10. ECM-associated signaling molecules | 1 | Gja1 | | | | |
| F11. Tissue remodeling growth factors | 1 | Bmp2 | | | | |
| F12. Leukocyte trafficking / tissue recruitment | 5 | Ccl3 | Ccl5 | Ccl7 | Ccl8 | Cxcl10 |

| **G. Lipid / metabolic program genes** | **25** | | | | | | |
|---|---|---|---|---|---|---|---|
| G1. Lipid uptake / transport/ handling | 6 | Apoe | Apoc1 | Apoc2 | Abca1 | Abca6 | Abca9 |
| G2. Fatty-acid activation and turnover | 2 | Abhd4 | Abhd15 | | | | |
| G3. Lipid storage / lipid droplet program | 0 | | | | | | |
| G4. Cholesterol / sterol biosynthesis | 4 | Hmgcs2 | Cyp27a1 | Gramd1a | Gramd1b | | |
| G5. Membrane lipid remodeling / ether lipid metabolism | 1 | Hsd17b11 | | | | | |
| G6. Fatty-acid synthesis / elongation | 0 | | | | | | |
| G7. Eicosanoid / inflammatory lipid metabolism | 1 | Ptgs2 | | | | | |
| G8. Glycolysis / metabolic adaptation | 1 | Gpd1 | | | | | |
| G9. Pentose phosphate pathway | 0 | | | | | | |
| G10. Serine biosynthesis | 0 | | | | | | |
| G11. Purine / pyrimidine biosynthesis | 0 | | | | | | |
| G12. Amino-acid metabolism | 3 | Aldh6a1 | Slc7a7 | Slc43a2 |  |  |  |
| G13. Redox / antioxidant metabolism | 2 | Gstm2 | | | | | |
| G14. Lysosome–lipid metabolism axis | 1 | Slc29a3 | | | | | |
| G15. Mitochondrial metabolic adaptation | 3 | Aldh2 | Slc25a28 | Slc25a36 | | | |
| G16. Metabolic stress adaptation | 1 | Kmo | | | | | |

| **H. Cell cycle / DNA replication / mitosis** | **9** | | | | | |
|---|---|---|---|---|---|---|
| H1. Cell-cycle regulators | 4 | Ccnd2 | Cdkn1a | Cdkn1b | Ccng2 | |
| H2. Replication licensing | 0 | | | | | |
| H3. DNA synthesis | 0 | | | | | |
| H4. Replication checkpoint | 0 | | | | | |
| H5. Cohesion/condensin | 0 | | | | | |
| H6. Centromere/kinetochore | 0 | | | | | |
| H7. Spindle assembly checkpoint | 0 | | | | | |
| H8. Mitotic kinases | 0 | | | | | |
| H9. Mitotic motors/spindel dynamics | 0 | | | | | |
| H10. Cytokinesis | 0 | | | | | |
| H11. Proliferation markers | 0 | | | | | |
| H12. DNA repair/genome maintenance | 5 | Dclre1c | Eepd1 | Fbh1 | Pot1b | Trex1 |
| H13. Histone/chromatin module | 0 | | | | | |

| **I. Cell death / survival control** | **19** | | | | | | | |
|---|---|---|---|---|---|---|---|---|
| I1. Anti-apoptotic / survival | 3 | Birc2 | Birc3 | Cflar | | | | |
| I2. Pro-apoptotic | 4 | Bcl2l11 | Bmf | Trp53inp1 | Cdip1 | | | |
| I3. Caspase machinery | 0 | | | | | | | |
| I4. Cell-fate regulators | 5 | Foxo3 | Hbp1 | Tgif1 | Rbl2 | Peg3 | | |
| I5. Survival adaptation | 7 | Sesn1 | Deptor | Txnip | Tsc22d1 | Tsc22d2 | Tsc22d3 | Ppp1r15a |

#### II. Cellular machinery / homeostatic cellular systems

| **J. Ribosome biogenesis / translation program** | **5** | | | |
|---|---|---|---|---|
| J1. Ribosome biogenesis / nucleolar assembly | 1 | Nop53 | | |
| J2. RNA helicases involved in ribosome assembly | 0 | | | |
| J3. Translation initiation | 3 | Eif4a2 | Eif4b | Eif4ebp2 |
| J4. Translation elongation / termination | 1 | Eef2k | | |
| J5. Aminoacyl-tRNA synthetases | 0 | | | |
| J6. tRNA / RNA modification machinery | 0 | | | |
| J7. rRNA transcription support | 0 | | | |

| **K. RNA processing / splicing program** | **16** | | | | | |
|---|---|---|---|---|---|---|
| K1. Core spliceosome components | 2 | Rbm39 | Rbm5 | | | |
| K2. RNA-binding proteins (mRNA processing) | 5 | Tia1 | Ptbp2 | Stau1 | Ago3 | Ago4 |
| K3. mRNA export / nuclear transport | 2 | Ranbp10 | Ranbp3l | | | |
| K4. RNA helicases | 2 | Dhx57 | Dhx58 | | | |
| K5. RNA processing regulators | 3 | Tut4 | Tut7 | Cnot6l | | |
| K6. RNA modification / maturation | 2 | Ints12 | Ssu72 | | | |
| K7. RNA surveillance / degradation | 0 | | | | | |
| K8. Small RNA / snoRNA machinery | 0 | | | | | |

| **L. Protein homeostasis / proteostasis program** | **28** | | | | | | | | | | |
|---|---|---|---|---|---|---|---|---|---|---|---|
| L1. Proteasome complex | | | | | | | | | | | |
| L2. Ubiquitin / ubiquitin-like system | 11 | Itch | Wwp1 | Herc1 | Herc4 | Herc6 | Rnf145 | Rnf150 | Rnf169 | Rnf213 | Fbxo3 |
| | | Fbxl20 | | | | | | | | | |
| L3. Protein folding / chaperone system | 1 | Serpinh1 | | | | | | | | | |
| L4. ER quality control / ERAD | 2 | Herpud1 | Ccpg1 | | | | | | | | |
| L5. Vesicle trafficking / membrane transport | 5 | Cltc | Ap2a2 | Vps39 | Vps41 | Gapvd1 | | | | | |
| L6. Nuclear pore complex | 0 | | | | | | | | | | |
| L7. Lysosomal proteostasis | 9 | Lamp1 | Ctsb | Ctsh | Ctsc | Tpp1 | Man2b1 | Man2b2 | Galc | Glb1 | |
| L8. Protein maturation / assembly machinery | 0 | | | | | | | | | | |

| **M. Mitochondrial program** | **20** | | |
|---|---|---|---|
| M1. Mitochondrial respiratory chain / OXPHOS | 2 | Cox7a2l | Ndufa6 |
| M2. Mitochondrial ATP synthesis | 0 | | |

| | | | | | | | | | | | |
|---|---|---|---|---|---|---|---|---|---|---|---|
| M3. Mitochondrial transport / carrier system | 2 | Slc25a28 | Slc25a36 | | | | | | | | |
| M4. Mitochondrial protein import / membrane organization | 1 | Tafazzin | | | | | | | | | |
| M5. Mitochondrial translation / ribosome | 0 | | | | | | | | | | |
| M6. Metabolism | 9 | Aldh2 | Aldh6a1 | Bckdha | Coq8a | Hmgcs2 | Kmo | Mthfr | Pnpla8 | Scp2 | |
| M7. Mitochondrial redox / stress adaptation | 3 | Cat | Mgst1 | Pyroxd2 | | | | | | | |
| M8. Mitochondrial dynamics / quality control | 3 | Pink1 | Prkn | Atp13a2 | | | | | | | |
| **N. Vesicle trafficking / endolysosomal program** | **39** | | | | | | | | | | |
| N1. Rab-family vesicle trafficking | 2 | Rab5b | Rab30 | | | | | | | | |
| N2. Endosomal sorting / retromer machinery | 5 | Snx5 | Snx6 | Snx11 | Snx13 | Snx29 | | | | | |
| N3. Cargo adaptor complexes | 3 | Ap1b1 | Ap2a2 | Ap4m1 | | | | | | | |
| N4. ER–Golgi / secretory trafficking | 2 | Stx5a | Use1 | | | | | | | | |
| N5. Vesicle fusion and membrane dynamics | 3 | Cltc | Eps15 | Fcho2 | | | | | | | |
| N6. Endocytosis / phagosome trafficking | 8 | Mertk | Axl | Fcgr2b | Fcgr3 | Cd47 | Tyrobp | Trem2 | P2rx7 | | |
| N7. Lysosome / endolysosomal compartment | 14 | Scarb2 | Tmem175 | Atp13a2 | Lamp1 | Laptm5 | Tpp1 | Hexa | Hexd | Galc | Glb1 |
| | | Man2b1 | Man2b2 | Idua | Gns | | | | | | |
| N8. Lysosomal acidification machinery | 0 | | | | | | | | | | |
| N9. ESCRT / membrane remodeling | 2 | Vps39 | Vps41 | | | | | | | | |
| **O. Nuclear transport / chromosome organization program** | **9** | | | | | | | | | | |
| O1. Nuclear pore complex | 0 | | | | | | | | | | |
| O2. Nuclear import/export machinery | 2 | Ranbp10 | Ranbp3l | | | | | | | | |
| O3. Nuclear envelope/lamina organization | 2 | Syne1 | Syne3 | | | | | | | | |
| O4. Chromosome architecture / genome organization | 1 | Nipbl | | | | | | | | | |
| O5. Chromatin structural proteins | 2 | Mecp2 | H1f0 | | | | | | | | |
| O6. Centrosome / chromosome organization apparatus | 2 | Cep68 | Kiz | | | | | | | | |
| O7. Nuclear positioning / nucleo-cytoskeletal coupling | 0 | | | | | | | | | | |
| O8. Chromosome topology facor | 0 | | | | | | | | | | |
| **P. Biosynthetic / nucleotide–cofactor metabolism program** | **9** | | | | | | | | | | |
| P1. Purine biosynthesis | 0 | | | | | | | | | | |
| P2. Pyrimidine biosynthesis | 0 | | | | | | | | | | |
| P3. One-carbon / folate metabolism | 2 | Mthfr | Mthfs | | | | | | | | |
| P4. Nucleotide precursor synthesis | 1 | Tk2 | | | | | | | | | |
| P5. Purine salvage / nucleotide interconversion | 1 | Cmpk2 | | | | | | | | | |
| P6. Cofactor / methyl-donor metabolism | 1 | Pts | | | | | | | | | |
| P7. mino-acid biosynthesis and polyamine metabolism | 2 | Sat1 | Paox | | | | | | | | |
| P8. Hexosamine / nucleotide-sugar biosynthesis | 1 | Gnpda1 | | | | | | | | | |
| P9. Ether-lipid precursor biosynthesis | 1 | Agpat3 | | | | | | | | | |
| **Q. Ubiquitin–proteasome / protein degradation program** | **39** | | | | | | | | | | |
| Q1. Proteasome core complex | 0 | | | | | | | | | | |
| Q2. Proteasome regulatory complex | 0 | | | | | | | | | | |
| Q3. Ubiquitin conjugation machinery | 1 | Uba7 | | | | | | | | | |
| Q4. Deubiquitination machinery | 1 | Otud5 | | | | | | | | | |
| Q5. E3 ubiquitin ligases and ubiquitin-associated factors | 20 | Nedd4 | Wwp1 | Itch | Rnf145 | Rnf150 | Rnf167 | Rnf169 | Rnf19b | Rnf213 | Fbxo3 |
| | | Fbxl20 | Cul7 | Cul9 | Herc1 | Herc4 | Herc6 | Arel1 | Dcaf6 | Dcaf8 | Dcaf12 |
| Q6. Autophagy / degradation-associated factors | 17 | Ulk1 | Ulk2 | Atg2a | Atg14 | Map1lc3a | Map1lc3b | Wipi2 | Rb1cc1 | Nbr1 | Wdr45 |
| | | Ncoa4 | Calcoco1 | Ccpg1 | Retreg3 | Rubcn | Rubcnl | Sidt2 | | | |
| Q7. COP9 signalosome and protein quality-control degradation | 0 | | | | | | | | | | |
| **R. Cytoskeleton / intracellular architecture program** | **38** | | | | | | | | | | |
| R1. Actin cytoskeleton core | 2 | Marcks | Coro1b | | | | | | | | |
| R2. Actin nucleation and branching machinery | 1 | Fmnl2 | | | | | | | | | |
| R3. Septin cytoskeleton | 2 | Septin4 | Septin8 | | | | | | | | |
| R4. Microtubule architecture | 5 | Clasp2 | Map1b | Mtcl3 | Tubb3 | Tppp | | | | | |
| R5. Intermediate filament and scaffold proteins | 1 | Ina | | | | | | | | | |
| R6. Membrane–cytoskeleton linker proteins | 2 | Afap1l1 | Snta1 | | | | | | | | |
| R7. Rho-family regulators | 15 | Arhgap12 | Arhgap15 | Arhgap19 | Arhgap29 | Arhgap45 | Arhgef3 | Arhgef12 | Dock5 | Dock9 | Prex1 |
| | | Vav2 | Rasa1 | Rasa4 | Rasal2 | Rhou | | | | | |
| R8. Structural adaptors | 4 | Iqgap2 | Specc1l | Tln2 | Afdn | | | | | | |
| R9. Cell polarity and cortical organization | 2 | Nckap1 | Abi2 | | | | | | | | |
| R10. Intracellular architecture regulators | 4 | Gas7 | Cdc42ep2 | Plekhg1 | Plekhg5 | | | | | | |
| **S. General signaling / adaptor program** | **42** | | | | | | | | | | |
| S1. Heterotrimeric G-protein signaling | 6 | Gnai1 | Gna12 | Rgs2 | Rgs3 | Rgs8 | Rgs10 | | | | |
| S2. Small GTPase signaling module | 0 | | | | | | | | | | |
| S3. Kinase/phosphatase-associated signaling | 12 | Abl1 | Mapk14 | Map3k8 | Dyrk1b | Dyrk2 | Prkcz | Jak1 | Tyk2 | Ppp1r15a | Ppp1r16b |
| | | Ppp1r9a | Ppp2r2c | | | | | | | | |
| S4. Membrane-associated signaling organizers | 7 | Gab1 | Gab2 | Afdn | Akap6 | Akap13 | Plekhg1 | Plekhg5 | | | |
| S5. Signal relay and signal-integration proteins | 10 | Ralgds | Ralgps2 | Ralgapa2 | Cyth1 | Cyth3 | Tifa | Peli1 | Peli2 | Traf4 | Trafd1 |
| S6. Miscellaneous adaptors | 7 | Nckap1 | Abi2 | Src | Cask | Erbin | Reps2 | Ifrd1 | | | |
| **T. Core cellular maintenance / housekeeping program** | **12** | | | | | | | | | | |
| T1. Glycolysis / glucose utilization | 0 | | | | | | | | | | |
| T2. Core energy metabolism | 3 | Insr | Igf1r | Igf1 | | | | | | | |

| | | | | | | | |
|---|---|---|---|---|---|---|---|
| T3. Redox maintenance / antioxidant maintenance | 0 | | | | | | |
| T4. Energy bufferingand phosphate homeostasis | 3 | Inpp4b | Inpp5k | Ip6k1 | | | |
| T5. Amino-acid / nitrogen metabolism | 0 | | | | | | |
| T6. General cellular maintenance | 6 | Tpt1 | Tmbim6 | Flcn | Narf | Oxr1 | Creg1 |

| **U. ER stress / secretory adaptation program** | **13** | | | |
|---|---|---|---|---|
| U1. Canonical UPR regulators | 0 | | | |
| U2. ER chaperones | 3 | Serpinh1 | Fkbp10 | Herpud1 |
| U3. Folding machinery | 2 | Use1 | Rcn3 | |
| U4. ER quality control / ERAD | 3 | Ccpg1 | Tmbim6 | Os9 |
| U5. Secretory pathway adaptation | 3 | Tmcc1 | Tmcc3 | Sft2d2 |
| U6. ER stress-associated metabolic adaptation | 2 | Crebrf | Eif4ebp2 | |

| **V. Glycosylation / membrane biogenesis** | **8** | | | |
|---|---|---|---|---|
| V1. N-linked glycosylation | 3 | Man1c1 | Man2a2 | Mgat3 |
| V2. O-linked glycosylation | 0 | | | |
| V3. Glycan extension / sugar donor metabolism | 3 | Gxylt2 | B3gat1 | B3galnt1 |
| V4. GPI-anchor biosynthesis | 2 | Pgap2 | Pgap6 | |
| V5. Golgi membrane biogenesis / glycoprotein trafficking | 0 | | | |

| **W. Sterol / membrane lipid remodeling program** | **14** | | | | | |
|---|---|---|---|---|---|---|
| W1. Cholesterol biosynthesis | 0 | | | | | |
| W2. Sterol sensing / transcriptional control | 2 | Rxra | Rxrb | | | |
| W3. Intracellular cholesterol transport | 5 | Npc2 | Gramd1a | Gramd1b | Osbpl9 | Osbpl11 |
| W4. Cholesterol turnover | 4 | Abca1 | Abcg1 | Cyp27a1 | Hpgd | |
| W5. Membrane sterol–lipid remodeling | 3 | Adipor1 | Mylip | Scp2 | | |

| **X. Redox / oxidative stress adaptation** | **8** | | | |
|---|---|---|---|---|
| X1. Glutathione system | 2 | Mgst1 | Gstm2 | |
| X2. Thioredoxin system | 0 | | | |
| X3. Peroxiredoxin / peroxide detoxification | 0 | | | |
| X4. Heme / oxidative stress response | 1 | Blvrb | | |
| X5 Selenium-dependent redox regulation | 2 | Selenop | Selenbp1 | |
| X6 Oxidative stress adaptation / survival | 3 | Cat | Oxr1 | Pyroxd2 |

| **Y. Core amino-acid / methyl-donor adaptation** | **0** |
|---|---|
| Y1. Polyamine biosynthesis | 0 |
| Y2. Methionine salvage / methyl-donor recycling | 0 |
| Y3. One-carbon flux | 0 |

**Gene of interest: Rtl6 (its UTR transcripts were upregulated in Rtl6 KO)**

#### Supplementary Table S3

#### UP genes (after LPS administration)

##### I. Major biological programs

| | | | | | | | | | | | | |
|---|---|---|---|---|---|---|---|---|---|---|---|---|
| **A. Microglia state genes** | **49** | | | | | | | | | | | |
| A1. microglia identity and DAM identity genes | 8 | | | | | | | | | | | |
| A1a Microglial identity | | 1 | Tmem119* | | | | | | | | | |
| A1b DAM identity genes | | 7 | Spp1 | Gpnmb | Lgals3 | Lgals1 | Ch25h | Fabp5 | Msr1 | | | |
| A2. Innate immune receptors | 20 | | | | | | | | | | | |
| A2a. Pattern-recognition receptors (PRRs) | | 0 | | | | | | | | | | |
| A2b. C-type lectin receptors | | 5 | Clec4e | Clec5a | Clec7a | Clec12a | Colec12 | | | | | |
| A2c. Myeloid immunoregulatory receptors | | 7 | Pira2 | Fcgr4 | Pilra | Lilrb4a | Lilrb4b | Lair1 | Havcr2 | | | |
| A2d. CD300 family receptors | | 3 | Cd300lb | Cd300ld | Cd300lf | | | | | | | |
| A2e. SLAM family receptors | | 2 | Slamf9 | Cd244a | | | | | | | | |
| A2f. Other immune receptors | | 3 | Gpr84 | P2ry10b | Cd180 | | | | | | | |
| A3 Cytokine / inflammatory signaling | 8 | | | | | | | | | | | |
| A3a. Cytokines | | 0 | | | | | | | | | | |
| A3b. Cytokine receptor genes | | 6 | Il18rap | Il1rl1 | Osmr | Tnfrsf11a | Tnfrsf21 | Cd180 | | | | |
| A3c. Chemokines | | 2 | Ccl12 | Cxcl16 | | | | | | | | |
| A4 Antigen processing / phagocytosis | 6 | | | | | | | | | | | |
| A4a. Antigen processing | | 3 | Ifi30 | Ctss | Hexb | | | | | | | |
| A4b. Antigen uptake / lusosomal | | 3 | Cd68 | Cd63 | Lamp1 | | | | | | | |
| A5. Innate immune transcriptional regulators | 7 | | | | | | | | | | | |
| A5a. Core transcription factors | | 2 | Bhlhe40 | Irf5 | | | | | | | | |
| A5b. mmune-response regulators | | 5 | Dusp4 | Dusp3 | Dusp6 | Dusp16 | C9orf72 | | | | | |
| **B. Cell-state regulators** | **133** | | | | | | | | | | | |
| B1. Transcriptional / Cell-state regulators | 38 | | | | | | | | | | | |
| | | | Bhlhe40 | Bhlhe41 | Klf2 | Klf6 | Klf10 | Etv4 | Etv5 | E2f1 | E2f2 | E2f3 |
| B1a. Core TFs | | 21 | E2f6 | E2f7 | E2f8 | Foxm1 | Atf4 | Atf6 | Sox9 | Myc | Id2 | Mafg |
| | | | Rbpj | | | | | | | | | |
| B1b. Transcriptional regulators | | 17 | Nab1 | Nab2 | Tfdp1 | Lin9 | Lin54 | Phf19 | Suz12 | Eed | Ezh2 | Mbd2 |
| | | | Mbd3 | Smyd5 | Nsd2 | Suv39h1 | Suv39h2 | Baz1a | Baz1b | | | |
| B2 Signal-adaptation regulators | 52 | | | | | | | | | | | |
| B2a. MAPK / stress-response modulators | | 9 | Camkk1 | Dusp3 | Dusp4 | Dusp6 | Dusp16 | Mapkapk2 | Map2k1 | Mapk6 | Spred1 | |
| B2b. PI3K–Akt / survival regulators | | 12 | Pdk1 | Rasa2 | Rasgrp3 | Rapgef5 | Ptk2b | Vav1 | Vav3 | Dgkz | Plekho1 | Plekho2 |
| | | | Sh2b2 | Bcl2l1 | | | | | | | | |
| B2c. Immune signaling adaptors | | 17 | Dok2 | Tollip | Card9 | Pycard | Rhbdf2 | Tnfaip1 | Trim27 | Ubash3b | Lrrfip1 | Cgas |
| | | | Casp1 | Pros1 | Ltbr | Il17ra | Tgfbr1 | Cd180 | Tnfrsf21 | | | |
| B2d. Cell-state stabilization regulators | | 14 | C9orf72 | Vps13c | Sbno2 | Sgk1 | Macir | Nacc1 | Limd1 | Rhob | Rhoa | Arl11 |
| | | | Dock10 | Elmo2 | Iqgap1 | Iqgap3 | | | | | | |
| B3 Epigenetic / chromatin state regulators | 43 | | | | | | | | | | | |
| B3a. Histone methylation regulators | | 6 | Suv39h1 | Suv39h2 | Ezh2 | Nsd2 | Smyd5 | Kmt5a | | | | |
| B3b. Polycomb-associated regulators | | 5 | Eed | Suz12 | Phf19 | Mbd2 | Mbd3 | | | | | |
| B3c. Chromatin remodeling factors | | 12 | Baz1a | Baz1b | Smarca5 | Smarcb1 | Smarcc1 | Actl6a | Hmgb1 | Hmgb2 | Hmgb3 | Hmgn1 |
| | | | Hmgn2 | Dek | | | | | | | | |
| B3d. DNA methylation / replication-coupled chromatin regulators | | 10 | Dnmt1 | Uhrf1 | Asf1b | Chaf1a | Chaf1b | Hat1 | Hirip3 | Nasp | Slbp | Rbbp7 |
| B3e. Chromosome architecture regulators | | 10 | Cbx3 | Cbx5 | Rad21 | Smc1a | Smc2 | Smc4 | Smc5 | Smc6 | Ncapd2 | Ncapd3 |
| **C. Myelin / axon–glia interaction genes** | **33** | | | | | | | | | | | |
| C1. Myelin-associated genes | 3 | | Plp2 | Pmp22 | Cd9 | | | | | | | |
| C2. Axon guidance / neurite interaction | 5 | | Nrp2 | Sema4c | Sema6b | Fat3 | Nav2 | | | | | |
| C3. Neuron–glia adhesion molecules | 4 | | Nrcam | Cadm1 | Pcdhgc3 | Sdc3 | | | | | | |
| C4. Neurotrophic / developmental signaling | 9 | | Jag1 | Fgf2 | Fgf13 | Igfbp5 | Igf2r | Pros1 | Pdgfb | Tgfbr1 | Rbpj | |
| C5. Synaptic / neuron–glia communication | 3 | | Ptk2b | Lrp8 | Dab2 | | | | | | | |
| C6. Glial structural support / neural architecture | 8 | | Nes | Tmem119 | Heg1 | Ttyh2 | Syne1 | Syne3 | Sox9 | Vim | | |
| C7. Neurovascular / neural niche interaction | 1 | | Cxcl14 | | | | | | | | | |
| **D. Neuron–glia communication genes** | **75** | | | | | | | | | | | |
| D1. Growth factor / trophic signaling | 8 | | Fgf2 | Fgf13 | Pdgfb | Igfbp5 | Igf2r | Jag1 | Pros1 | Metrnl | | |
| D2. Guidance / axon interaction | 8 | | Nrcam | Cadm1 | Sema4c | Sema6b | Nrp2 | Ptprc | Ptpre | Ptk2b | | |
| D3. Neuroactive receptors | 12 | | Calcrl | Gpr180 | Gpr183 | Cnr2 | P2ry10b | Il1rl1 | Osmr | Il17ra | Tnfrsf11a | Tnfrsf21 |
| | | | Ltbr | Cd180 | | | | | | | | |
| D4. Cell-cell communication | 10 | | Cd44 | Cd34 | Cd69 | Cd52 | Cd9 | Spn | Fxyd5 | Emp1 | Emp3 | Pmp22 |
| D5. Cytoskeletal/synapse-contact | 11 | | Ezr | Msn | Actn1 | Actn4 | Flna | Iqgap1 | Pacsin2 | Myo1b | Myo1e | Myo10 |
| | | | Ptk2b | | | | | | | | | |
| D6. Vesicle trafficking / phagosome–synapse interface | 8 | | Rab27a | Rab32 | Rab11a | Rab5c | Rab7 | Eea1 | Stx7 | Tpcn2 | | |
| D7. Neuron-responsive TFs | 10 | | Bhlhe40 | Bhlhe41 | Klf2 | Klf6 | Klf10 | Etv4 | Etv5 | Nab1 | Nab2 | Id2 |
| D8. Neuroimmune communication | 8 | | Lgals3 | Lgals1 | Gpnmb | Spp1 | Ccl12 | Cxcl14 | Hexb | Abca7 | | |
| **E. Astrocyte-like / glial-environment genes** | **70** | | | | | | | | | | | |
| E1. Astrocyte identity/glial differentiation | 8 | | Sox9 | Nes | Vim | Jag1 | Rbpj | Fgf2 | Fgf13 | Epas1 | | |
| E2. Metabolic support | 10 | | Slc1a5 | Slc7a1 | Slc7a5 | Slc2a1 | Slc2a3 | Slc16a1 | Slc16a3 | Igf2r | Ldlr | Apobr |
| E3. Lipid/cholesterol support | 17 | | Hmgcr | Hmgcs1 | Sqle | Cyp51 | Sc5d | Insig1 | Idi1 | Fdps | Stard4 | Srebf2 |
| | | | Acsl3 | Acsl4 | Acsl5 | Abhd5 | Plin2 | Elovl5 | Hsd17b12 | | | |
| E4. Astrocyte ECM / neurovascular niche | 15 | | Fn1 | Col14a1 | Col4a6 | Lamc1 | Emilin2 | Olfml2b | Olfml3 | Tgfbi | Spp1 | Flt1 |
| | | | Calcrl | Nrp2 | Gpr180 | Pdgfb | Heg1 | | | | | |
| E5. Reactive astrocyte / glial-environment response | 20 | | Osmr | Il1rl1 | Il17ra | Ltbr | Cd44 | Pros1 | Atf4 | Atf6 | Xbp1 | Bhlhe40 |
| | | | Bhlhe41 | Hmox1 | Srxn1 | Creld2 | Manf | Hyou1 | Niban1 | Niban2 | Gpnmb | Lgals3 |
| **F. ECM-related genes** | **80** | | | | | | | | | | | |
| F1. ECM structural components | 8 | | Fn1 | Col14a1 | Col4a6 | Lamc1 | Emilin2 | Olfml2b | Olfml3 | Tgfbi | | |
| F2. ECM remodeling enzymes | 12 | | Adam8 | Adam9 | Adam10 | Adam17 | Adamts1 | Ctss | Lgmn | Plau | Plaur | Enpp1 |
| | | | Dnase2a | Plod1 | | | | | | | | |
| F3. Matrisome-associated secreted proteins | 6 | | Spp1 | Gpnmb | Lgals3 | Pros1 | Metrnl | Serpine2 | | | | |
| F4. ECM glycan / proteoglycan regulators | 3 | | Hs3st3b1 | B4galt7 | Csgalnact2 | | | | | | | |
| F5. ECM adhesion receptors | 7 | | Itga5 | Itga6 | Itgav | Itgb1 | Itgb2 | Itgb3 | Itgb5 | | | |
| F6. Adhesion complex / focal adhesion | 7 | | Fermt3 | Tln1 | Lims1 | Itgb1bp1 | Apbb1ip | Parvg | Lpxn | | | |
| F7. Cell–matrix interaction receptors | 8 | | Cd44 | Cd34 | Cadm1 | Nrcam | Heg1 | Nrp2 | Calcrl | Gpr180 | | |
| F8. Migration / invasion regulators | 10 | | Dock10 | Elmo2 | Arhgef7 | Arhgef39 | Arhgef40 | Arhgap22 | Arhgap24 | Arhgap25 | Arhgap30 | Rapgef5 |
| F9. Cytoskeletal ECM-response machinery | 11 | | Actn1 | Actn4 | Flna | Ezr | Msn | Pacsin2 | Plekha2 | Plekha8 | Fmn1 | Diaph1 |
| | | | Diaph3 | | | | | | | | | |
| F10. ECM-associated signaling molecules | 3 | | Ptk2b | Iqgap1 | Dab2 | | | | | | | |
| F11. Tissue remodeling growth factors | 2 | | Fgf2 | Pdgfb | | | | | | | | |
| F12. Leukocyte trafficking / tissue recruitment | 3 | | Selplg | Sdc3 | Pkp2 | | | | | | | |
| **G. Lipid / metabolic program genes** | **141** | | | | | | | | | | | |
| G1. Lipid uptake / transport/ handling | 8 | | Fabp4 | Fabp5 | Abca7 | Apobr | Lrp8 | Lrp12 | Igf2r | Nceh1 | | |
| G2. Fatty-acid activation and turnover | 8 | | Acsl3 | Acsl4 | Acsl5 | Acot9 | Acot11 | Acat1 | Acat2 | Agpat5 | | |
| G3. Lipid storage / lipid droplet program | 2 | | Plin2 | Abhd5 | | | | | | | | |
| G4. Cholesterol / sterol biosynthesis | 15 | | Acat2 | Hmgcr | Hmgcs1 | Sqle | Msmo1 | Cyp51 | Sc5d | Idi1 | Fdps | Insig1 |
| | | | Stard4 | Srebf2 | Tmem97 | Ldlr | Por | | | | | |
| G5. Membrane lipid remodeling / ether lipid metabolism | 7 | | Agps | Osbpl8 | Plaat3 | Far1 | Peds1 | Agps | Pon2 | | | |
| G6. Fatty-acid synthesis / elongation | 9 | | Fasn | Acaca | Elovl1 | Elovl5 | Hsd17b12 | Degs1 | Cers2 | Cers5 | Cers6 | |
| G7. Eicosanoid / inflammatory lipid metabolism | 4 | | Pla2g4a | Alox5ap | Ptges3 | Ch25h | | | | | | |
| G8. Glycolysis / metabolic adaptation | 18 | | Pgm1 | Pfkp | Pfkl | Pgk1 | Pgam1 | Gapdh | Eno1 | Tpi1 | Aldoa | Gpi1 |
| | | | Ldha | Pkm | Pdk1 | Tigar | Slc2a1 | Slc2a3 | Slc2a6 | Hk3 | | |
| G9. Pentose phosphate pathway | 3 | | Pgd | Tkt | Taldo1 | | | | | | | |
| G10. Serine biosynthesis | 2 | | Phgdh | Psat1 | | | | | | | | |
| G11. Purine / pyrimidine biosynthesis | 10 | | Cad | Pfas | Paics | Atic | Ctps1 | Umps | Aprt | Adsl | Gart | Ppat |
| G12. Amino-acid metabolism | 7 | | Asns | Aldh18a1 | Oat | Got1 | Got2 | Slc7a5 | Slc1a5 | | | |
| G13. Redox / antioxidant metabolism | 20 | | Gclc | Gsr | Txn1 | Txnrd1 | Txndc5 | Txndc9 | Txndc11 | Prdx1 | Prdx2 | Prdx4 |
| | | | Hmox1 | Srxn1 | Blvra | Park7 | Gpx1 | Selenof | Selenoh | Sephs1 | Sephs2 | Ptgr1 |
| G14. Lysosome–lipid metabolism axis | 4 | | Hexb | Gba1 | Manba | Naaa | | | | | | |
| G15. Mitochondrial metabolic adaptation | 13 | | Acat1 | Gatm | Gcat | Got1 | Got2 | Me2 | Oxct1 | Pgm1 | Hmbs | Pdss1 |
| | | | Cisd1 | Gpd2 | Flvcr1 | | | | | | | |
| G16. Metabolic stress adaptation | 11 | | Atf4 | Atf6 | Xbp1 | Niban1 | Niban2 | Manf | Creld2 | Hyou1 | Retreg1 | Mif |
| | | | Helz2 | | | | | | | | | |
| **H. Cell cycle / DNA replication / mitosis** | **292** | | | | | | | | | | | |
| | | | Cdk1 | Ccna2 | Ccnb1 | Ccnb2 | Ccne1 | Ccne2 | Cdc25b | Cdc25c | Cdc7 | Wee1 |
| H1. Cell-cycle regulators | 33 | | Pkmyt1 | Mastl | Foxm1 | Mybl2 | E2f1 | E2f2 | E2f3 | E2f6 | E2f7 | E2f8 |

| | | | | | | | | | | | |
|---|---|---|---|---|---|---|---|---|---|---|---|
| [illegible] | [illegible] | Tcf19 | Skp2 | Fbxo5 | Cks1b | Cks2 | Cdca2 | Cdca3 | Cdca4 | Cdca5 | Cdca7 |
| | | Cdca7l | Cdca8 | Cdkn3 | | | | | | | |
| H2. Replication licensing | 19 | Orc1 | Orc2 | Orc6 | Cdc6 | Cdt1 | Cdc45 | Mcm2 | Mcm3 | Mcm4 | Mcm5 |
| | | Mcm6 | Mcm7 | Mcm8 | Mcm10 | Gins1 | Gins2 | Ticrr | Dbf4 | Wdhd1 | |
| H3. DNA synthesis | 26 | Pold1 | Pold2 | Pole | Pole2 | Pole3 | Pole4 | Pola1 | Pola2 | Prim1 | Prim2 |
| | | Lig1 | Fen1 | Rrm1 | Rrm2 | Tk1 | Tyms | Dut | Dhfr | Ung | Pcna |
| | | Pclaf | Dscc1 | Rfc1 | Rfc2 | Rfc3 | Rfc4 | | | | |
| H4. Replication checkpoint | 11 | Chek1 | Clspn | Timeless | Tipin | Hus1 | Rad9a | Topbp1 | Rpa1 | Rpa2 | Atr |
| | | Traip | | | | | | | | | |
| H5. Cohesion/condensin | 22 | Esco2 | Sgo1 | Sgo2a | Smc1a | Smc2 | Smc4 | Smc5 | Smc6 | Ncapg | Ncapg2 |
| | | Ncapd2 | Ncapd3 | Ncaph | Ncaph2 | Rad21 | Pds5a | Incenp | Tmpo | Lmnb1 | Lmnb2 |
| | | Cenpc1 | Ndc1 | | | | | | | | |
| H6. Centromere/kinetochore | 24 | Cenpe | Cenpf | Cenph | Cenpi | Cenpk | Cenpl | Cenpn | Cenpp | Cenpq | Cenps |
| | | Cenpt | Cenpu | Cenpw | Ndc80 | Nuf2 | Knl1 | Zwilch | Zwint | Spc24 | Spc25 |
| | | Dsn1 | Mis18a | Mis18bp1 | Hjurp | | | | | | |
| H7. Spindle assenbly checkpoint | 8 | Bub1 | Bub1b | Bub3 | Mad1l1 | Mad2l1 | Ttk | Kntc1 | Nsl1 | | |
| H8. Mitotic kinases | 7 | Plk1 | Plk4 | Aurka | Aurkb | Nek2 | Haspin | Melk | | | |
| H9. Mitotic motors/spindel dynamics | 23 | Kif2c | Kif4 | Kif11 | Kif14 | Kif15 | Kif18a | Kif18b | Kif20a | Kif20b | Kif22 |
| | | Kif23 | Kif24 | Kifc1 | Spag5 | Tacc3 | Tpx2 | Stil | Aspm | Spdl1 | Ska1 |
| | | Ska2 | Ska3 | Knstrn | | | | | | | |
| H10. Cytokinesis | 10 | Anln | Racgap1 | Prc1 | Cit | Ect2 | Cep55 | Gas2l3 | Shcbp1 | Troap | Prr11 |
| H11. Proliferation markers | 18 | Mki67 | Pimreg | Dlgap5 | Nusap1 | Hmmr | Pbk | Birc5 | Oip5 | Cip2a | Fam83d |
| | | Depdc1a | Depdc1b | Gtse1 | Ccnf | Atad2 | Ckap2 | Ckap2l | Poc1a | | |
| H12. DNA repair/genome maintenance | 39 | Brca1 | Brca2 | Bard1 | Brip1 | Fanca | Fancb | Fancd2 | Fanci | Fancm | Rad51 |
| | | Rad51ap1 | Rad51b | Rad51c | Rad54b | Rad54l | Gen1 | Eme1 | Exo1 | Recql4 | Rtel1 |
| | | Zranb3 | Rbbp8 | Nbn | Chek2 | Polh | Polq | Rfwd3 | Swi5 | Firrm | Ddias |
| | | Ankle1 | Blm | Dna2 | Hrob | Mms22l | Tonsl | Atm | Ube2t | Fancl | |
| H13. Histone/chromatin module | 52 | H1f1 | H1f3 | H1f4 | H1f5 | H2ac4 | H2ac6 | H2ac7 | H2ac8 | H2ac10 | H2ac11 |
| | | H2ac13 | H2ac15 | H2ac18 | H2ac19 | H2ac20 | H2ac22 | H2ac23 | H2ac24 | H2bc3 | H2bc4 |
| | | H2bc6 | H2bc7 | H2bc8 | H2bc9 | H2bc11 | H2bc12 | H2bc13 | H2bc14 | H2bc15 | H2bc18 |
| | | H2bc22 | H3c1 | H3c2 | H3c3 | H3c4 | H3c6 | H3c7 | H3c10 | H3c11 | H3c13 |
| | | H3c15 | H4c1 | H4c2 | H4c3 | H4c4 | H4c6 | H4c8 | H4c11 | H4c12 | H4c14 |
| | | H4c16 | H4c18 | | | | | | | | |

| | | | | | | | | | |
|---|---|---|---|---|---|---|---|---|---|
| **I. Cell death / survival control** | **27** | | | | | | | | |
| I1. Anti-apoptotic / survival | 8 | **Bcl2** | **Bcl2l1** | **Bcl2a1d** | Api5 | Birc2 | Birc3 | Cflar | Mcl1 |
| I2. Pro-apoptotic | 8 | **Bak1** | Bid | Bik | Bax | Bad | Bbc3 | Pmaip1 | **Bcl2l11** |
| I3. Caspase machinery | 3 | Casp7 | Casp8 | Casp9 | | | | | |
| I4. Cell-fate regulators | 6 | Morrbid | Faf1 | Siva1 | Bcl2l11 | Dapk1 | Dapk2 | | |
| I5. Survival adaptation | 2 | Tnfaip8 | Tnfaip8l2 | | | | | | |

#### II. Cellular machinery / homeostatic cellular systems

| | | | | | | | | | | | |
|---|---|---|---|---|---|---|---|---|---|---|---|
| **J. Ribosome biogenesis / translation program** | **94** | | | | | | | | | | |
| J1. Ribosome biogenesis / nucleolar assembly | 34 | Nolc1 | Nol10 | Nol11 | Nop2 | Nop9 | Nop56 | Nop58 | Nifk | Utp4 | Utp6 |
| | | Utp14b | Utp15 | Utp18 | Utp20 | Rrp7a | Rrp12 | Rrp15 | Rpf1 | Rpf2 | Pwp1 |
| | | Pwp2 | Nip7 | Bop1 | Bud23 | Npm1 | Rrs1 | Rbm19 | Urb1 | Urb2 | Krr1 |
| | | Mak16 | Heatr1 | Heatr3 | Brix1 | | | | | | |
| J2. RNA helicases involved in ribosome assembly | 11 | Ddx1 | Ddx11 | Ddx21 | Ddx27 | Ddx31 | Ddx39a | Ddx39b | Ddx56 | Dhx9 | Dhx15 |
| | | Dhx29 | | | | | | | | | |
| J3. Translation initiation | 20 | Eif1a | Eif1ax | Eif1ad | Eif2s1 | Eif2s2 | Eif2b3 | Eif2b5 | Eif3a | Eif3b | Eif3c |
| | | Eif3g | Eif3j1 | Eif3l | Eif4a1 | Eif4a3 | Eif4e | Eif4g1 | Eif4g2 | Eif4h | Etf1 |
| J4. Translation elongation / termination | 4 | Eef1e1 | Eef1g | Gspt1 | Denr | | | | | | |
| J5. Aminoacyl-tRNA synthetases | 17 | Aars1 | Aarsd1 | Farsa | Farsb | Gars1 | Hars1 | Iars1 | Iars2 | Kars1 | Lars1 |
| | | Mars1 | Rars1 | Sars1 | Tars1 | Wars1 | Vars1 | Yars1 | | | |
| J6. tRNA / RNA modification machinery | 6 | Mettl1 | Nsun2 | Ftsj3 | Pus7 | Trmt6 | Trmt10c | | | | |
| J7. rRNA transcription support | 2 | Polr1a | Mybbp1a | | | | | | | | |

| | | | | | | | | | | | |
|---|---|---|---|---|---|---|---|---|---|---|---|
| **K. RNA processing / splicing program** | **113** | | | | | | | | | | |
| K1. Core spliceosome components | 14 | Snrpa | Snrpa1 | Snrpb | Snrpd1 | Snrpe | Snrpf | Snrnp40 | Snu13 | Sf3a3 | Sf3b3 |
| | | Sf3b5 | Sf3b6 | U2af1 | U2af2 | | | | | | |
| K2. RNA-binding proteins (mRNA processing) | 14 | Hnrnpa1 | Hnrnpa3 | Hnrnpab | Hnrnpf | Hnrnpk | Hnrnpl | Hnrnpm | Hnrnpu | Khdrbs1 | Ptbp1 |
| | | Caprin1 | Fmr1 | Tardbp | Ybx1 | | | | | | |
| K3. mRNA export / nuclear transport | 11 | Alyref | Ncbp1 | Xpo1 | Xpo4 | Kpna1 | Kpna2 | Kpnb1 | Nutf2 | Ipo5 | Ipo7 |
| | | Ipo9 | | | | | | | | | |
| K4. RNA helicases | 12 | Ddx1 | Ddx11 | Ddx21 | Ddx27 | Ddx31 | Ddx39a | Ddx39b | Ddx56 | Dhx9 | Dhx15 |
| | | Dhx29 | Dhx32 | | | | | | | | |
| K5. RNA processing regulators | 25 | Srsf1 | Srsf2 | Srsf3 | Srsf6 | Srsf7 | Srsf9 | Srsf10 | Tra2b | Rbms1 | Rbm3 |
| | | Rbm15 | Rbm19 | Rbm44 | Rbmxl1 | Cstf2 | Tcerg1 | Prpf19 | Prpf31 | Prpf40a | Dazap1 |
| | | Larp1 | Larp1b | Larp4 | Larp4b | Larp7 | | | | | |
| K6. RNA modification / maturation | 10 | Nsun2 | Mettl1 | Pus7 | Trmt6 | Trmt10c | Ftsj3 | Nop2 | Nhp2 | Dkc1 | Tep1 |
| K7. RNA surveillance / degradation | 7 | Exosc2 | Exosc8 | Exosc9 | Eri1 | Eri2 | Cnot6 | Cnot9 | | | |
| K8. Small RNA / snoRNA machinery | 20 | Nop56 | Nop58 | Pno1 | Pwp1 | Pwp2 | Nol10 | Nol11 | Nolc1 | Rrp12 | Rrp15 |
| | | Rrp7a | Utp4 | Utp6 | Utp18 | Utp20 | Utp15 | Utp14b | Imp4 | Bop1 | Bud23 |

| | | | | | | | | | | | |
|---|---|---|---|---|---|---|---|---|---|---|---|
| **L. Protein homeostasis / proteostasis program** | **165** | | | | | | | | | | |
| L1. Proteasome complex | 35 | Psma1 | Psma2 | Psma3 | Psma4 | Psma5 | Psma6 | Psma7 | Psmb1 | Psmb2 | Psmb3 |
| | | Psmb4 | Psmb5 | Psmb6 | Psmb7 | Psmb8 | Psmc1 | Psmc2 | Psmc3 | Psmc4 | Psmc5 |
| | | Psmc6 | Psmd1 | Psmd2 | Psmd3 | Psmd4 | Psmd5 | Psmd6 | Psmd8 | Psmd11 | Psmd12 |
| | | Psmd13 | Psmd14 | Psme1 | Psme2 | Psme3 | | | | | |
| L2. Ubiquitin / ubiquitin-like system | 24 | Uba2 | Uba3 | Uba6 | Ube2c | Ube2i | Ube2l3 | Ube2m | Ube2n | Ube2s | Ube2t |
| | | Ube2v2 | Rnf4 | Rnf128 | Rnf168 | Trim27 | Trim28 | Trim37 | Trim59 | Fbxl14 | Skp2 |
| | | Dcun1d5 | Senp5 | Sae1 | Nploc4 | | | | | | |
| L3. Protein folding / chaperone system | 33 | Hspa4 | Hspa4l | Hspa5 | Hspa9 | Hspa13 | Hspa14 | Hsp90aa1 | Hsp90b1 | Hspd1 | Hspe1 |
| | | Dnaja1 | Dnaja2 | Dnajb11 | Dnajc2 | Dnajc3 | Dnajc9 | Dnajc10 | Dnajc21 | Fkbp2 | Fkbp3 |
| | | Fkbp4 | Fkbp5 | Stip1 | Calr | Canx | Pdia3 | Pdia4 | Pdia6 | Erp29 | Erp44 |
| | | Ppia | Ppib | Ahsa1 | | | | | | | |
| L4. ER quality control / ERAD | 21 | Edem1 | Edem3 | Uggt1 | Selenos | Retreg1 | Creld2 | Manf | Hyou1 | Sdf2l1 | Tmx1 |
| | | Tmx4 | Ermp1 | Rpn1 | Rpn2 | Ddost | Mlec | Ostc | Stt3b | Alg6 | Alg8 |
| | | Alg10b | | | | | | | | | |
| L5. Vesicle trafficking / membrane transport | 28 | Rab1a | Rab5c | Rab7 | Rab8a | Rab11a | Rab27a | Rab32 | Arf1 | Arf6 | Ykt6 |
| | | Vps13c | Vps29 | Vps35 | Vps36 | Ap1s1 | Ap3m1 | Ap3s1 | Snx1 | Snx2 | Snx10 |
| | | Snx24 | Cops3 | Cops4 | Cops5 | Cops6 | Nsf | Cltb | Eea1 | | |
| L6. Nuclear pore complex | 15 | Nup35 | Nup43 | Nup50 | Nup62 | Nup85 | Nup88 | Nup93 | Nup107 | Nup153 | Nup155 |
| | | Nup160 | Nup188 | Nup205 | Nup210 | Pom121 | | | | | |
| L7. Lysosomal proteostasis | 5 | Ctss | Lgmn | Dnase2a | Atg3 | Ostm1 | | | | | |
| L8. Protein maturation / assembly machinery | 4 | Get3 | Get4 | Afg2a | Pomp | | | | | | |

| | | | | | | | | | | | |
|---|---|---|---|---|---|---|---|---|---|---|---|
| **M. Mitochondrial program** | **86** | | | | | | | | | | |
| M1. Mitochondrial respiratory chain / OXPHOS | 17 | Cyc1 | Cox5b | Cox6a1 | Cox7a2 | Uqcrb | Uqcrc1 | Uqcrc2 | Uqcrq | Ndufa9 | Ndufa12 |
| | | Ndufb6 | Ndufb7 | Ndufb8 | Ndufs1 | Ndufs4 | Ndufs6 | Ndufs8 | | | |
| M2. Mitochondrial ATP synthesis | 8 | Atp5f1a | Atp5f1b | Atp5if1 | Atp5mc1 | Atp5mc3 | Atp5mf | Atp5pb | Atp5po | | |
| M3. Mitochondrial transport / carrier system | 6 | Slc25a5 | Slc25a13 | Slc25a24 | Slc25a25 | Slc25a32 | Slc25a44 | | | | |
| M4. Mitochondrial protein import / membrane organization | 11 | Tomm5 | Tomm22 | Tomm40 | Tomm70a | Timm8a1 | Timm13 | Timm17a | Timm23 | Timm50 | Samm50 |
| | | Immt | | | | | | | | | |
| M5. Mitochondrial translation / ribosome | 17 | Mrpl13 | Mrpl15 | Mrpl16 | Mrpl18 | Mrpl19 | Mrpl20 | Mrpl21 | Mrpl33 | Mrpl41 | Mrpl42 |
| | | Mrpl51 | Mrpl52 | Mrpl54 | Mrps11 | Mrps14 | Mrps18b | Mrps28 | | | |
| M6. Metabolism | 12 | Dld | Mdh1 | Mdh2 | Gpd2 | Cs | Ak2 | Ak6 | Oat | Hmbs | Ppat |
| | | Ptgr1 | Oxct1 | | | | | | | | |
| M7. Mitochondrial redox / stress adaptation | 5 | Cisd1 | Fdx2 | Nmral1 | Pdk1 | Gatm | | | | | |
| M8. Mitochondrial dynamics / quality control | 10 | Dnm1l | Opa3 | Tspo | Vdac1 | Vdac2 | Vdac3 | Pgam5 | Mtfr2 | Mcub | Mcoln2 |

| | | | | | | | | | | | |
|---|---|---|---|---|---|---|---|---|---|---|---|
| **N. Vesicle trafficking / endolysosomal program** | **61** | | | | | | | | | | |
| N1. Rab-family vesicle trafficking | 8 | Rab1a | Rab5c | Rab7 | Rab7b | Rab8a | Rab11a | Rab27a | Rab32 | | |
| N2. Endosomal sorting / retromer machinery | 8 | Snx1 | Snx2 | Snx10 | Snx24 | Vps13c | Vps29 | Vps35 | Vps36 | | |
| N3. Cargo adaptor complexes | 3 | Ap1s1 | Ap3m1 | Ap3s1 | | | | | | | |
| N4. ER–Golgi / secretory trafficking | 14 | Tmed2 | Tmed5 | Tmed8 | Tmed9 | Tmed10 | Arcn1 | Sec13 | Sec23b | Sec24d | Golm1 |
| | | Get3 | Get4 | Surf4 | Rer1 | | | | | | |
| N5. Vesicle fusion and membrane dynamics | 4 | Stx3 | Stx7 | Ykt6 | Nsf | | | | | | |
| N6. Endocytosis / phagosome trafficking | 9 | Eea1 | Myof | Plekhf2 | Rabggtb | Vac14 | Rufy1 | Dennd4a | Dennd4c | Dennd10 | |
| N7. Lysosome / endolysosomal compartment | 8 | Lyset | Cln5 | Hexb | Cd68 | Cd63 | Lamp1 | Mcoln2 | Tpcn2 | | |

| | | | | | | | | | | | |
|---|---|---|---|---|---|---|---|---|---|---|---|
| N8. Lysosomal acidification machinery | 6 | Atp6v0a2 | Atp6v0c | Atp6v1a | Atp6v1c1 | Atp6v1d | Atp6v1h | | | | |
| N9. ESCRT / membrane remodeling | 1 | Chmp6 | | | | | | | | | |
| **O. Nuclear transport / chromosome organization program** | **64** | | | | | | | | | | |
| O1. Nuclear pore complex | 14 | Nup35 | Nup43 | Nup50 | Nup62 | Nup85 | Nup88 | Nup93 | Nup107 | Nup153 | Nup155 |
| | | Nup160 | Nup188 | Nup205 | Nup210 | | | | | | |
| O2. Nuclear import/export machinery | 13 | Kpna1 | Kpna2 | Kpnb1 | Ipo5 | Ipo7 | Ipo9 | Xpo1 | Xpo4 | Nutf2 | Ran |
| | | Ranbp1 | Ranbp2 | Rangap1 | | | | | | | |
| O3. Nuclear envelope/lamina organization | 8 | Lmna | Lmnb1 | Lmnb2 | Tmpo | Sun2 | Syne1 | Syne3 | Lbr | | |
| O4. Chromosome architecture / genome organization | 6 | Smchd1 | Rif1 | Paxip1 | Mcph1 | Banf1 | Nucks1 | | | | |
| O5. Chromatin structural proteins | 8 | Hmgb1* | Hmgb2* | Hmgb3* | Hmgn1* | Hmgn2* | Dek* | Rcc1 | Rcc1l | | |
| O6. Centrosome / chromosome organization apparatus | 11 | Nde1 | Nin | Odf2 | Cep83 | Cep128 | Cep152 | Cep192 | Cep250 | Cep295 | Tubg1 |
| | | Tubgcp2 | | | | | | | | | |
| O7. Nuclear positioning / nucleo-cytoskeletal coupling | 3 | Plec | Map4 | Gpsm2 | | | | | | | |
| O8. Chromosome topology facor | 1 | Top2a** | | | | | | | | | |
| **P. Biosynthetic / nucleotide–cofactor metabolism program** | **51** | | | | | | | | | | |
| P1. Purine biosynthesis | 6 | Ppat | Gart | Pfas | Paics | Atic | Adsl | | | | |
| P2 .Pyrimidine biosynthesis | 4 | Cad | Ctps1 | Umps | Dtymk | | | | | | |
| P3. One-carbon / folate metabolism | 8 | Phgdh | Psat1 | Shmt1 | Shmt2 | Mthfd1 | Mthfd1l | Mthfd2 | Gcsh | | |
| P4. Nucleotide precursor synthesis | 5 | Rrm1 | Rrm2 | Tk1 | Dck | Aprt | | | | | |
| P5. Purine salvage / nucleotide interconversion | 7 | Ampd2 | Ak2 | Ak6 | Impdh2 | Gmpr | Nt5dc2 | Adk | | | |
| P6. Cofactor / methyl-donor metabolism | 5 | Ahcy | Mtap | Pdss1 | Nans | Nmral1 | | | | | |
| P7. mino-acid biosynthesis and polyamine metabolism | 7 | Asns | Aldh18a1 | Oat | Azin1 | Srm | Sms | Amd1 | | | |
| P8. Hexosamine / nucleotide-sugar biosynthesis | 6 | Gnpnat1 | Uap1l1 | Gale | Ugdh | Man2a1 | Gmppb | | | | |
| P9. Ether-lipid precursor biosynthesis | 3 | Agps | Peds1 | Isyna1 | | | | | | | |
| **Q. Ubiquitin–proteasome / protein degradation program** | **81** | | | | | | | | | | |
| Q1. Proteasome core complex | 15 | Psma1 | Psma2 | Psma3 | Psma4 | Psma5 | Psma6 | Psma7 | Psmb1 | Psmb2 | Psmb3 |
| | | Psmb4 | Psmb5 | Psmb6 | Psmb7 | Psmb8 | | | | | |
| Q2. Proteasome regulatory complex | 21 | Psmc1 | Psmc2 | Psmc3 | Psmc4 | Psmc5 | Psmc6 | Psmd1 | Psmd2 | Psmd3 | Psmd4 |
| | | Psmd5 | Psmd6 | Psmd8 | Psmd11 | Psmd12 | Psmd13 | Psmd14 | Psme1 | Psme2 | Psme3 |
| | | Pomp | | | | | | | | | |
| Q3. Ubiquitin conjugation machinery | 10 | Uba2 | Uba3 | Uba6 | Ube2c | Ube2i | Ube2m | Ube2n | Ube2s | Ube2t | Ube2v2 |
| Q4. Deubiquitination machinery | 11 | Usp1 | Usp4 | Usp5 | Usp7 | Usp10 | Usp14 | Usp15 | Usp31 | Usp46 | Otud4 |
| | | Otud6b | | | | | | | | | |
| Q5. E3 ubiquitin ligases and ubiquitin-associated factors | 14 | Rnf4 | Rnf128 | Rnf168 | Trim27 | Trim28 | Trim37 | Trim59 | Rfwd3 | Fbxl14 | Skp2 |
| | | Wsb1 | Wsb2 | Dcun1d5 | Nploc4 | | | | | | |
| Q6. Autophagy / degradation-associated factors | 3 | Atg3 | Pgam5 | Dnase2a | | | | | | | |
| Q7. COP9 signalosome and protein quality-control degradation | 7 | Cops3 | Cops4 | Cops5 | Cops6 | Gps1 | Sem1 | Vcp | | | |
| **R. Cytoskeleton / intracellular architecture program** | **53** | | | | | | | | | | |
| R1. Actin cytoskeleton core | 5 | Actb | Actg1 | Cfl1 | Cap1 | Capza1 | | | | | |
| R2. Actin nucleation and branching machinery | 7 | Arpc1b | Arpc2 | Arpc3 | Arpc4 | Arpc5 | Actr2 | Actr3 | | | |
| R3. Septin cytoskeleton | 3 | Septin6 | Septin7 | Septin11 | | | | | | | |
| R4. Microtubule architecture | 10 | Tuba1b | Tuba1c | Tuba4a | Tubb2a | Tubb4b | Tubb5 | Tubb6 | Tbcb | Tbcd | Ttl |
| R5. Intermediate filament and scaffold proteins | 2 | Vim | Plec | | | | | | | | |
| R6. Membrane–cytoskeleton linker proteins | 6 | Ezr | Msn | Anxa2 | Anxa3 | Anxa4 | Anxa7 | | | | |
| R7. Rho-family regulators | 7 | Rhoa | Rhob | Rhoc | Arhgdia | Arhgdib | Cyrib | Abr | | | |
| R8. Structural adaptors | 6 | Cnn2 | Lasp1 | Crk | Pfn1 | Bcar3 | Sh3bp2 | | | | |
| R9. Cell polarity and cortical organization | 4 | Scrib | Pea15a | Ppp1r18 | Gpsm2 | | | | | | |
| R10. Intracellular architecture regulators | 3 | Map4 | Tacc1 | Iqgap1 | | | | | | | |
| **S. General signaling / adaptor program** | **43** | | | | | | | | | | |
| S1. Heterotrimeric G-protein signaling | 6 | Gnai2 | Gnai3 | Gnb1 | Gnb2 | Gng10 | Rgs7bp | | | | |
| S2. Small GTPase signaling module | 5 | Rap2a | Arf1 | Arf6 | Rabggtb | Ralb | | | | | |
| S3. Kinase/phosphatase-associated signaling | 15 | Camk1 | Camkk1 | Camk2g | Map2k1 | Mapkapk2 | Mapk6 | Pdk1 | Acp1 | Ptpa | Ppp1ca |
| | | Ppp1cb | Ppp1cc | Ppp2ca | Ppp1r8 | Ppp4r3a | | | | | |
| S4. Membrane-associated signaling organizers | 6 | Plekha2 | Plekha8 | Plekho1 | Plekho2 | Parvg | Itgb1bp1 | | | | |
| S5. Signal relay and signal-integration proteins | 6 | Ywhah | Ywhag | Ywhae | Ywhaz | Ywhaq | Pin1 | | | | |
| S6. Miscellaneous adaptors | 5 | Cacul1 | Ccny | Adprs | Gpr180 | Rasa2* | | | | | |
| **T. Core cellular maintenance / housekeeping program** | **44** | | | | | | | | | | |
| T1. Glycolysis / glucose utilization | 11 | Gapdh | Pgk1 | Pgam1 | Eno1 | Tpi1 | Pfkp | Pfkl | Pkm | Gpi1 | Aldoa |
| | | Ldha | | | | | | | | | |
| T2. Core energy metabolism | 5 | Cs | Idh3a | Dlst | Mdh1 | Me2 | | | | | |
| T3. Redox maintenance / antioxidant maintenance | 9 | Txn1 | Txnrd1 | Prdx1 | Prdx2 | Prdx4 | Gsr | Gpx1 | Srxn1 | Blvra | |
| T4. Energy bufferingand phosphate homeostasis | 8 | Ckb | Ak2 | Ak6 | Ppa1 | Itpa | Nme1 | Nme2 | Nme6 | | |
| T5. Amino-acid / nitrogen metabolism | 4 | Oat | Hmbs | Alad | Ampd2 | | | | | | |
| T6. General cellular maintenance | 7 | Comtd1 | Ciapin1 | Adk | Aprt | Higd1a | Nmral1 | Cisd1 | | | |
| **U. ER stress / secretory adaptation program** | **72** | | | | | | | | | | |
| U1. Canonical UPR regulators | 4 | Atf4 | Atf6 | Xbp1 | Manf | | | | | | |
| U2. ER chaperones | 13 | Hspa5 | Hsp90b1 | Calr | Canx | Dnajb11 | Dnajc3 | Hspa4 | Hspa4l | Hspa9 | Hspa13 |
| | | Hspa14 | Stip1 | Erp29 | | | | | | | |
| U3. Folding machinery | 14 | Pdia3 | Pdia4 | Pdia6 | P4hb | Ero1a | Ero1b | Fkbp2 | Fkbp3 | Fkbp4 | Fkbp5 |
| | | Txndc5 | Txndc11 | Tmx1 | Tmx4 | | | | | | |
| U4. ER quality control / ERAD | 7 | Edem1 | Edem3 | Uggt1 | Selenos | Ermp1 | Rer1 | Lyset | | | |
| U5. Secretory pathway adaptation | 23 | Golm1 | Surf4 | Lman1 | Mlec | Calu | Rpn1 | Rpn2 | Ddost | Sec11c | Ostc |
| | | Stt3b | Alg6 | Alg8 | Alg10b | Tmed2 | Tmed5 | Tmed8 | Tmed9 | Tmed10 | Emc1 |
| | | Emc2 | Sec61a1 | Sec61b | | | | | | | |
| U6. ER stress-associated metabolic adaptation | 11 | Hyou1 | Asns | Slc1a5 | Slc7a5 | Slc38a1 | Psat1 | Phgdh | Mthfd1 | Mthfd1l | Shmt1 |
| | | Shmt2 | | | | | | | | | |
| **V. Glycosylation / membrane biogenesis** | **28** | | | | | | | | | | |
| V1. N-linked glycosylation | 6 | Alg6 | Alg8 | Alg10b | Mgat2 | Mgat5 | Man2a1 | | | | |
| V2. O-linked glycosylation | 7 | Galnt6 | Galnt7 | Galnt12 | C1galt1 | Gcnt1 | Gcnt2 | B4galnt1 | | | |
| V3. Glycan extension / sugar donor metabolism | 6 | B4galt5 | B4galt7 | Gnpnat1 | Uap1l1 | Ugdh | Gale | | | | |
| V4. GPI-anchor biosynthesis | 1 | Piga | | | | | | | | | |
| V5. Golgi membrane biogenesis / glycoprotein trafficking | 8 | Golm1 | Tmed2 | Tmed5 | Tmed8 | Tmed9 | Tmed10 | Surf4 | Lman1 | | |
| **W. Sterol / membrane lipid remodeling program** | **29** | | | | | | | | | | |
| W1. Cholesterol biosynthesis | 8 | Hmgcr | Hmgcs1 | Fdps | Idi1 | Sqle | Cyp51 | Msmo1 | Sc5d | | |
| W2. Sterol sensing / transcriptional control | 2 | Srebf2 | Insig1 | | | | | | | | |
| W3. Intracellular cholesterol transport | 3 | Stard4 | Osbpl8 | Npc2 | | | | | | | |
| W4. Cholesterol turnover | 3 | Nceh1 | Abhd5 | Plin2 | | | | | | | |
| W5. Membrane sterol–lipid remodeling | 13 | Acsl3 | Acsl4 | Acsl5 | Elovl1 | Elovl5 | Degs1 | Cers2 | Cers5 | Cers6 | Far1 |
| | | Gpat4 | Hsd17b12 | Plaat3 | | | | | | | |
| **X. Redox / oxidative stress adaptation** | **32** | | | | | | | | | | |
| X1. Glutathione system | 9 | Gclc | Gclm | Gss | Gsr | Gstm1 | Gstm2 | Gstm3 | Gstt1 | Mgst1 | |
| X2. Thioredoxin system | 4 | Txn1 | Txn2 | Txnrd1 | Txnrd2 | | | | | | |
| X3. Peroxiredoxin / peroxide detoxification | 6 | Prdx1 | Prdx2 | Prdx4 | Prdx5 | Prdx6 | Srxn1 | | | | |
| X4. Heme / oxidative stress response | 3 | Hmox1 | Blvrb | Slc40a1 | | | | | | | |
| X5. Selenium-dependent redox regulation | 4 | Sephs1 | Sephs2 | Selenow | Selenoh | | | | | | |
| X6. Oxidative stress adaptation / survival | 6 | Park7 | Oxr1 | Sesn1 | Sesn3 | Tigar | Fdx2 | | | | |
| **Y. Core amino-acid / methyl-donor adaptation** | **8** | | | | | | | | | | |
| Y1. Polyamine biosynthesis | 3 | Amd1 | Srm | Sms | | | | | | | |
| Y2. Methionine salvage / methyl-donor recycling | 2 | Ahcy | Mtap | | | | | | | | |
| Y3. One-carbon flux | 3 | Shmt1 | Shmt2 | Mthfd2 | | | | | | | |

**Gene of interest: Rtl6 (its UTR transcripts were upregulated in Rtl6 KO)**

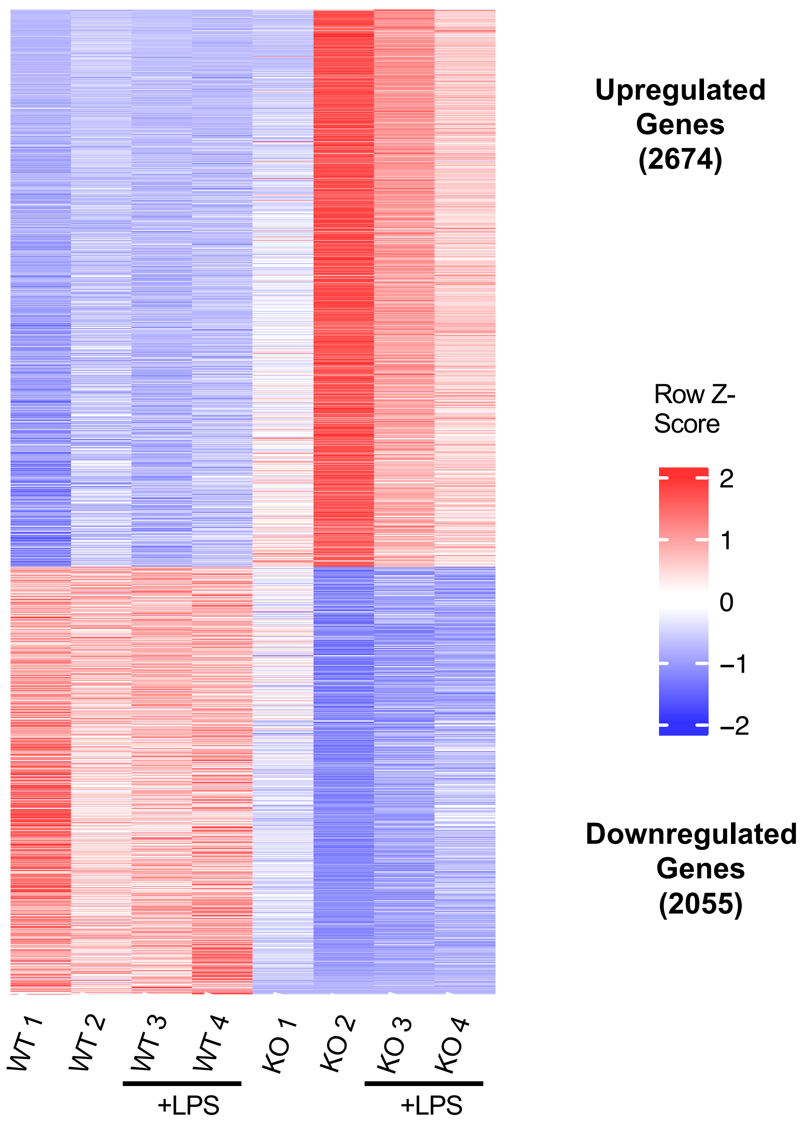

**Supplementary Fig S1**

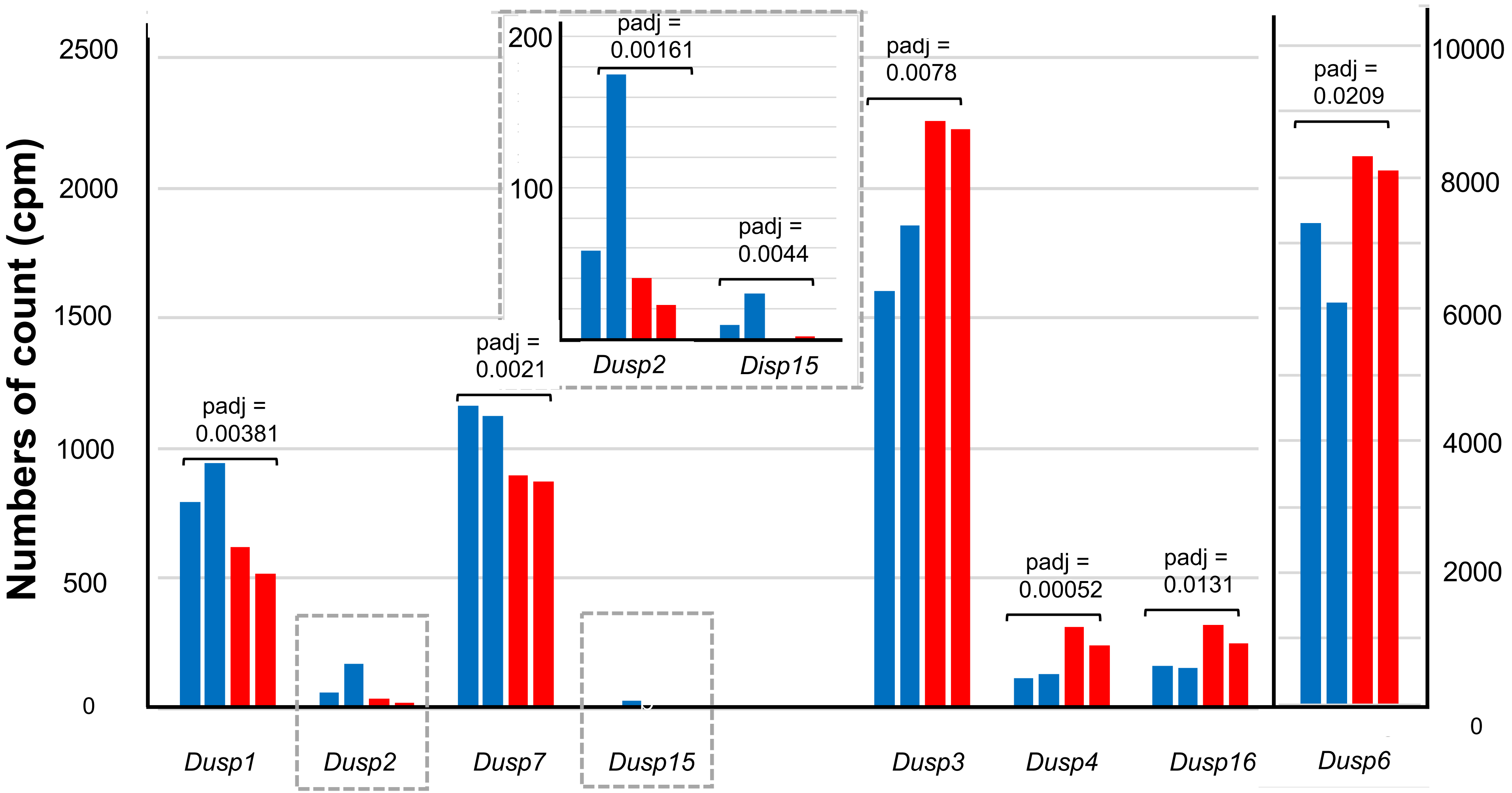

**Supplementary Fig S2**
